# Common Marmoset Gut Microbiome Profiles in Health and Intestinal Disease

**DOI:** 10.1101/2020.08.27.268524

**Authors:** Alexander Sheh, Stephen C. Artim, Monika A. Burns, Jose Arturo Molina-Mora, Mary Anne Lee, JoAnn Dzink-Fox, Sureshkumar Muthupalani, James G. Fox

**Affiliations:** Division of Comparative Medicine, Massachusetts Institute of Technology, Cambridge, Massachusetts; Centro de Investigación en Enfermedades Tropicales (CIET), Universidad de Costa Rica, San José, Costa Rica; Department of Biological Sciences, Wellesley College, Wellesley, Massachusetts

**Author notes:** These authors contributed equally to this work (Alexander Sheh, Stephen C. Artim).

**Keywords:** marmoset, *Prevotella copri*, *Clostridium perfringens*, inflammatory bowel disease, stricture, microbiome, enteritis

## Abstract

Chronic gastrointestinal (GI) diseases are the most common diseases in captive marmosets. The gut microbiome of healthy (n=91), inflammatory bowel disease (IBD) (n=59), and duodenal ulcer/stricture (n=23) captive marmosets was characterized. Healthy marmosets exhibited a “humanized,” *Bacteroidetes-dominant* microbiome. Despite standardized conditions, cohorts subdivided into *Prevotella*- and *Bacteroides-dominant* groups based on marmoset source. IBD was highest in a *Prevotella*-dominant cohort while strictures were highest in a *Bacteroides*-dominant cohort. Stricture-associated dysbiosis was characterized by *Anaerobiospirillum* loss and *Clostridium perfringens* increases. Stricture tissue presented upregulation of lipid metabolism genes and increased abundance of *C. perfringens*, a causative agent of GI diseases and intestinal strictures in humans. IBD was associated with a lower *Bacteroides*:*P. copri* ratio within each source. Consistent with *Prevotella*-linked diseases, pro-inflammatory genes were upregulated. This report highlights the humanization of the captive marmoset microbiome and its potential as a “humanized” animal model of *C. perfringens-induced* enteritis/strictures and *P. copri-associated* IBD.

## Background

Over 3.5 million people worldwide are affected by inflammatory bowel disease (IBD), a chronic gastrointestinal (GI) inflammatory disease triggered by interactions between host, microbes and the environment^1–5^. Two common forms of IBD are Crohn’s disease (CD), which affects the small and large intestines, and ulcerative colitis (UC), which localizes to the large intestine. Over 200 genomic loci may confer increased IBD risk, with many of these genes associated with regulating host-microbe interactions^1^. The human GI tract harbors trillions of microorganisms from at least 400 species that compose the intestinal microbiota^6,7^. In healthy individuals, the microbiome influences many physiological functions such as extracting nutrients, maintaining the gut mucosal barrier, training immune cells and protecting against pathogens^8^. Dysbiosis occurs due to loss of beneficial microbes, expansion of pathobionts (opportunistic microbes), or reduction of microbial diversity. Dysbiosis has been associated with human diseases, including irritable bowel syndrome, obesity, psoriasis, rheumatoid arthritis, autism spectrum disorders, *Clostridioles difficile* infection and IBD^8,9^. Changes in the intestinal microbiota observed in IBD patients have included reduction of short chain fatty acid (SCFA) producing bacteria, reduced alpha diversity, decreased *Firmicutes* abundance, and increased abundance of facultative anaerobes, *Proteobacteria* and *Bacleroideles*^2–4,10–12^.

In captive common marmosets, GI diseases are the most common and widespread clinical finding^13, 14^. IBD prevalence is reported to be as high as 28-60% in captive marmosets and presents with diarrhea, weight loss, enteritis, muscle atrophy, alopecia, hypoproteinemia, anemia, elevated liver enzymes, and a failure to thrive^13,15^. The IBD diagnosis can be refined to chronic lymphocytic enteritis (CLE) with histologic findings, such as small intestinal localization, shortened villi, crypt epithelial hyperplasia, and lymphocytic infiltration of the lamina propria^13,14^. Potential marmoset biomarkers include calprotectin and matrix metalloproteinase 9^16,17^, but clinical interventions involving glucocorticoids, gluten-free diets, Giardia treatment, etc. have yielded mixed results^18–20^. In addition to IBD, a novel chronic GI disease has been described in young adult marmosets characterized by duodenal dilation or stricture near the major duodenal papilla^21,22^. Clinical signs, such as diarrhea, weight loss, or poor weight gain, resemble IBD but increased vomiting is also observed. This syndrome was associated with hypoalbuminemia, hypoglobulinemia, hypoproteinemia, hypocalcemia (total), elevated alkaline phosphatase, anemia, and in some cases, leukocytosis^22^. Histologically, duodenal mucosal ulcerations with associated chronic-active granulocytic and lympho-histiocytic inflammation were observed.

As the microbiome has been associated to human GI diseases, factors affecting the microbiome in non-human primates (NHP) are being explored, such as species, social structure, environment and diet^23–26^. Captivity and captive diets have been associated with microbial diversity loss, shifts in the *Firmicutes*:*Bacteroidetes* ratio, and increased GI disease and mortality^23,26,27^. Dietary specialists, such as marmosets, are more susceptible to captivity-associated dietary changes^26^. Marmosets are exudivores that consume large amounts of indigestible oligosaccharides from tree gums^28^, and may harbor specific gut microbes dedicated to carbohydrate metabolism. Currently, few reports on the marmoset microbiome are available^29–34^. In this study, we evaluated microbiome, serum chemistry and complete blood count (CBC) samples from healthy marmosets (n=91) and marmosets with IBD (n=59) or duodenal ulcer/strictures (n=23), collected during physical examinations or necropsies over a two-year period. ‘Healthy’ controls were defined as individuals not clinically diagnosed with IBD or strictures and not receiving chronic drug treatments during the study period. Unique microbial profiles were associated with the four sources that populated the MIT colony. We identified changes in both microbial communities and blood parameters that may serve as marmoset biomarkers for IBD and strictures, and propose that marmosets may be useful animal models to study CD and *Clostridium*-driven GI disorders, such as duodenal strictures.

## Results

### Microbial Diversity in the Intestinal Microbiota of the Common Marmoset

303 samples from 91 healthy marmosets were analyzed to determine the normal microbiota (**Table 1**). 99% of the average microbial abundance in feces was captured by *Bacteroidetes, Firmicutes, Proteobacteria, Fusobacteria* and *Actinobacteria* (**Fig. 1a**). The microbiome profile observed in healthy, MIT marmosets resembles the microbiome observed in human stool with dominance of the phylum *Bacteroidetes* (average 63.2%), followed distantly by *Firmicutes* and *Proteobacteria*^7^. As observed in humans^7^, *Bacteroidetes* abundance varied significantly, ranging from 8-86%. *Bacteroidetes* were predominantly represented by *Bacteroides, Prevotella 9* and *Parabacteroides*. The most abundant *Firmicutes* were *Megamonas, Megasphaera* and *Phascolartcobacterium. Anaerobiospirillum, Sutterella* and *Escherichia-Shigella* were the most common *Proteobacteria*. Notably, *Bifidobacterium* were present in low abundance compared to other reported marmoset microbiomes^29,30^ (**Supp. Table 1**).

**Figure 1.**
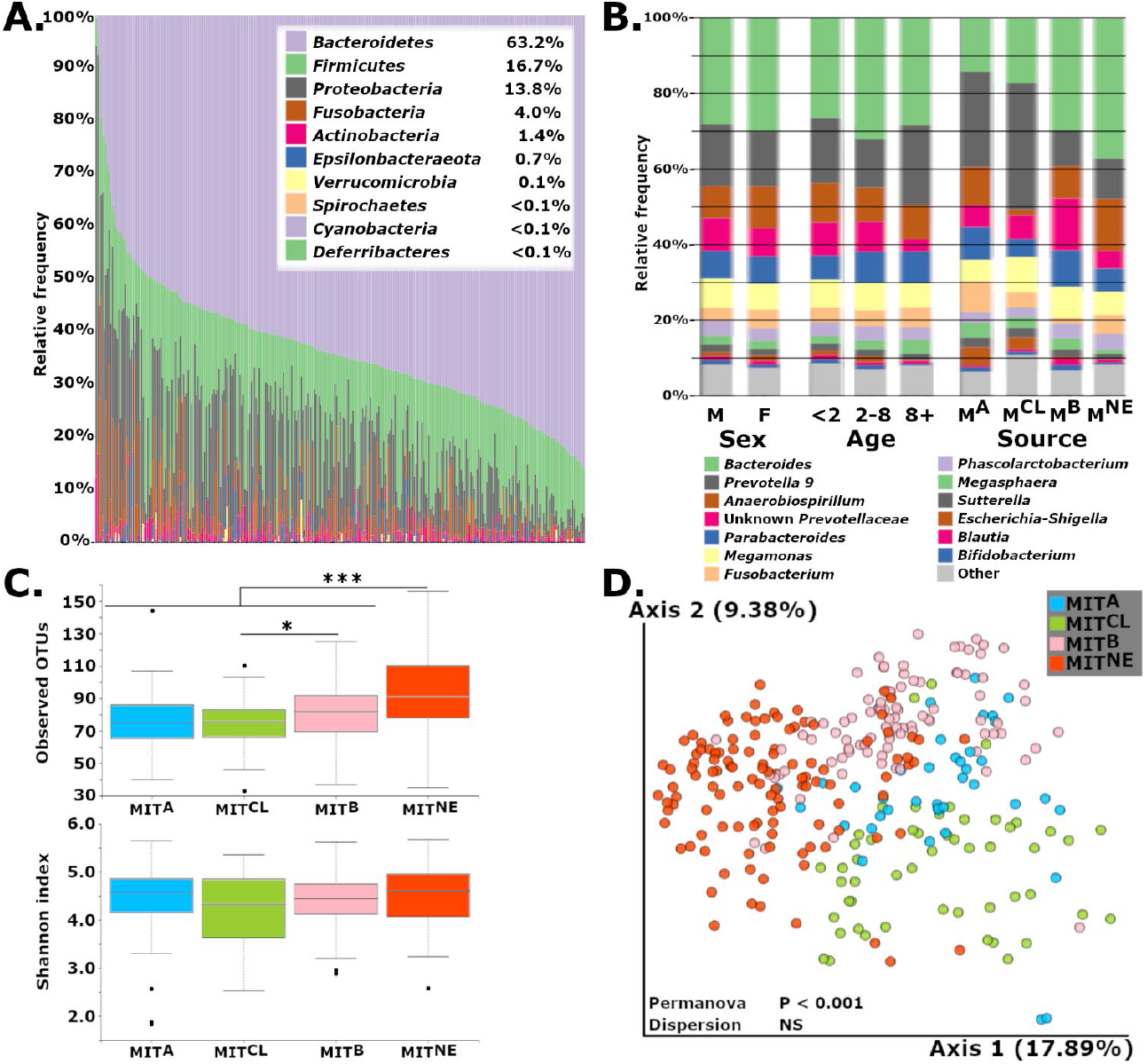
A) Gut microbiome profiles of healthy, common marmosets at the phylum level exhibit a *Bacteroidetes*-dominant and human-like microbiome. B) Averaged relative abundances at the genus level show differences associated with source but few differences based on sex or age. C) Observed OTUs were increased in MIT^NE^ vs. all sources and MIT^B^ compared to MITA and MIT^CL^, but metrics involving evenness, such as Shannon’s diversity index, showed no difference. D) PCoA plot using Unweighted UniFrac metric shows clustering of microbiome profiles based on marmoset source. *, *P<0.05;* **, *P<0.01* and ***, *P<0.001*.

**Table 1.** Demographics of samples used classified by Sex, Age, Sample Type and Source. Columns break down samples by disease status.

|  |  | Healthy | Stricture | IBD |
| --- | --- | --- | --- | --- |
| Sex | Male | 156 | 25 | 95 |
|  | Female | 147 | 37 | 105 |
| Age | 2 and under | 145 | 19 | 36 |
|  | 2 to 8 | 139 | 42 | 131 |
|  | Over 8 | 19 | 1 | 33 |
| Type | Rectal | 207 | 24 | 111 |
|  | Fecal | 96 | 38 | 89 |
| Source | MIT<sup>NE</sup> | 117 | 57 | 89 |
|  | MIT<sup>B</sup> | 94 | 3 | 30 |
|  | MIT<sup>CL</sup> | 53 | 2 | 61 |
|  | MIT<sup>A</sup> | 39 | 0 | 20 |

### Source population impacted microbiome diversity

Having established the baseline microbiome for healthy, MIT marmosets, we explored the effects of age, sex and original source, and found that source strongly influenced composition (**Fig 1b**). MIT’s colony originally received marmosets from four sources (A, B, CLEA and NEPRC), which we designated MIT^A^, MIT^B^, MIT^CL^ and MIT^NE^ following importation. Marmosets were housed in two buildings and provided standardized diet, husbandry and veterinary care. For this study, marmosets were co-housed with same-source animals. Using multiple estimators for alpha diversity, we noted that species richness estimators significantly differed between healthy marmosets by source, but not sex or age. MIT^NE^ marmosets had higher observed OTUs and Chao1 values compared to other sources (P<0.001 vs. each source, both metrics) (**Fig 1c, Supp. Fig 1**). MITB had significantly higher alpha diversity compared to MIT^CL^ (observed OTUs, P<0.05; Chao1, P<0.01) and MITA (Chao1, P<0.05). However, differences were not observed when accounting for evenness (Shannon diversity or Pielou’s evenness). Clustering of samples based on source (Unweighted UniFrac: PERMANOVA, P<0.001; beta-dispersion, NS) (**Fig 1d**), but not sex or age, was also observed (**Supp. Fig 2**).

### *Bacteroides* and *Prevotella* Define Microbial Communities of Sources

We next identified 63 differentially abundant genera between the 4 sources in the lower gut using ANCOM (Analysis of Composition of Microbiomes). 13 genera were present at relative abundances greater than 1% in at least one source (**Supp. Table 2**). High abundance of *Bacteroides* characterized MIT^NE^ and MIT^B^ samples, while MIT^CL^ and MIT^A^ were primarily colonized by genus *Prevotella 9* (**Fig. 1b**). The *Bacteroidaceae:Prevotellaceae* ratios for MITA, MIT^CL^, MITB, and MIT^NE^ (0.44, 0.39, 1.23 and 2.17, respectively) emphasize source-associated differences reflected in these two genera. *Anaerobiospirillum*, another highly abundant genus, represented 8.5-13.8% of bacterial in three sources but had low numbers in MIT^CL^ marmosets (1.5%).

Next, we explored the ability of classification models to identify marmoset source based on microbiome data (**Supp. Fig 3a**). After evaluating multiple models, we developed random forest (RF) classification models to identify healthy marmosets by source. After ranking ASVs based on their importance to the model, we iteratively created new models to determine the minimum number of ASVs required to achieve stability in accuracy, and selected an optimized model using 10 ASVs (**Supp. Fig 3b, 3c**). The optimized model achieved an accuracy of 93% with 100% sensitivity and 95% specificity. The RF model confirmed that despite importation and assimilation, unique source-specific signature microbiota were retained by cohousing same-source animals.

### Prevalence of GI disease in MIT-housed Marmosets

To study the effects of GI disease on the microbiome, marmosets were categorized as healthy (n=91), IBD (n=59) or duodenal stricture (n=23). Strictures were mainly observed in MIT^NE^ (21 of 23 cases) with a 26% prevalence in this cohort. IBD was observed throughout the colony with varied prevalence (MIT^CL^, 55%; MIT^NE^, 29%; MITA, 27%; and MIT^B^, 22%) (**Supp. Table 3**).

### Effects of Duodenal Strictures on the microbiome and blood analysis

As strictures predominantly affected MIT^NE^, we next investigated the effects of duodenal strictures on the marmoset microbiome in this cohort. “Progressors,” marmosets that had or developed strictures, had markedly different microbiomes compared to “non-progressors,” animals that remained healthy or developed other diseases (**Fig. 2a, Supp. Fig. 4a**). On average, a 32% decrease in *Bacteroides* was observed in stricture cases (35.8% abundance in non-progressors vs. 24.5% in progressors), which decreased the *Bacteroides*:*Prevotella 9* ratio from 3.1 in non-progressors to 1.4 in progressors. *Anaerobiospirillum*, the second most abundant genus in non-stricture marmosets (13%), decreased to 4.6% in stricture cases. Concurrently, a 50% increase in *Megamonas* was observed in progressors (**Fig. 2a**). ANCOM identified *Anaerobiospirillum* and *Clostridium sensu stricto 1* as differentially expressed, with *Clostridium sensu stricto 1* increases measured in progressors.

**Figure 2.**
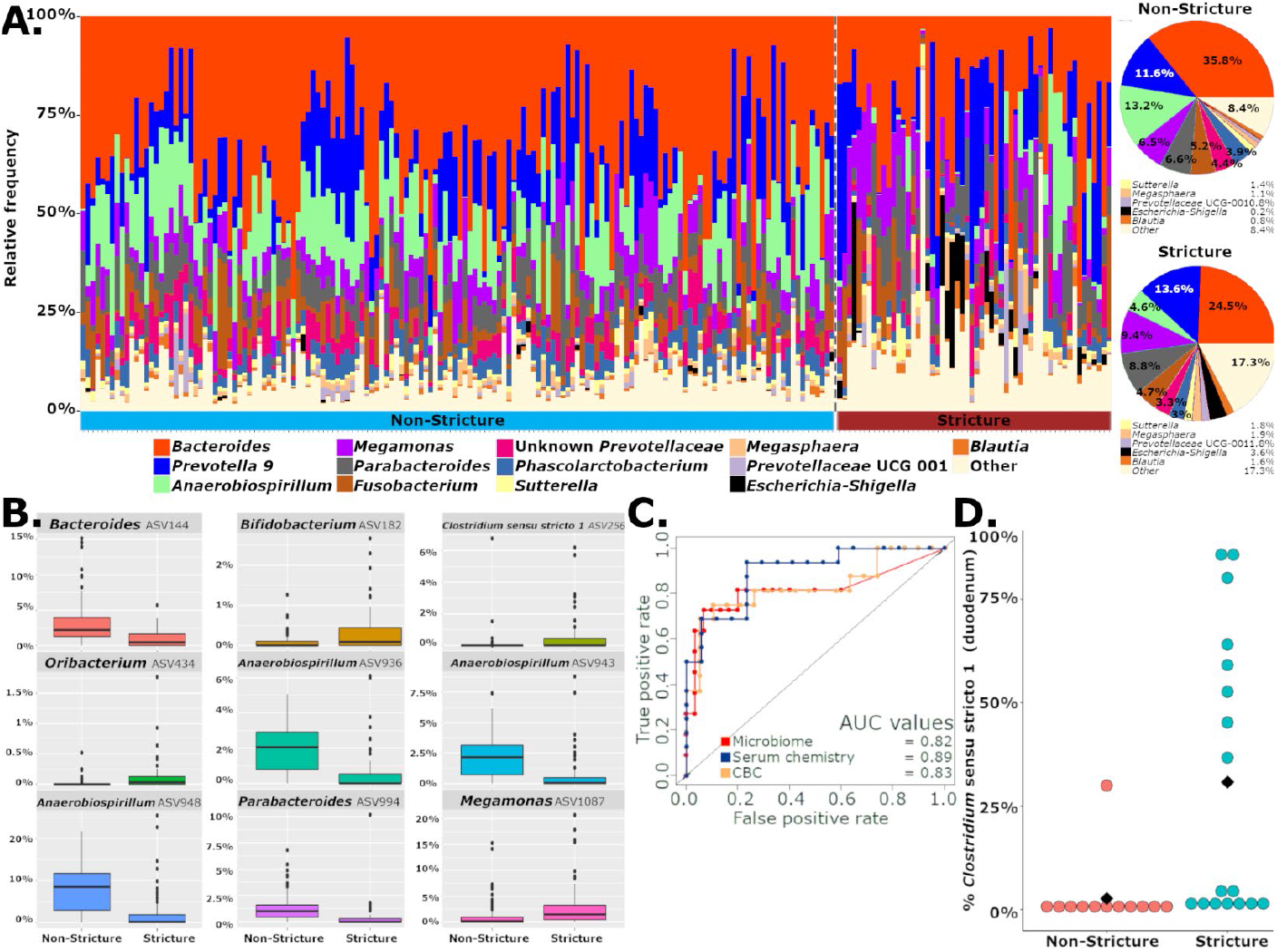
A) Microbiome composition of samples at the genus level and pie charts with average bacterial abundances of stricture progressors and non-progressors show dysbiosis associated with stricture characterized by decreased *Bacteroides* and *Anaerobiospirillum* and increased *Megamonas*. B) Nine ASVs identified by a random forest model that can correctly classify stricture and non-stricture samples with a 85% accuracy. C) Area under the curve (AUC) of receiver operating characteristic (ROC) curves for random forest models using microbiome, serum chemistry or complete blood count show strong performance of models in classifying strictures and non-strictures. D) Relative abundance of *Clostridium sensu stricto 1* reads in duodenal tissues is increased in stricture cases compared to non-stricture cases.

Despite changes in microbial composition, no changes in alpha diversity were observed using multiple metrics. We then optimized a 9 ASV RF model that minimized the number of ASVs while maximizing accuracy of stricture classification (**Fig. 2b, Supp. Fig. 4b, 4c**). Of these 9 ASVs, 3 *Anaerobiospirillum* ASVs, as well as *Bacteroides* and *Parabacteroides* ASVs, decreased in progressors. Increases were observed in ASVs from *Bifidobacterium, Clostridium sensu stricto 1, Oribacterium*, and *Megamonas*. The receiver operating characteristic (ROC) curve for this model had an area under the curve (AUC) of 0.82 (**Fig. 2c**) with an accuracy of 85%, a sensitivity of 100% and a specificity of 45%.

As ANCOM and our model highlighted the role of *Clostridium sensu stricto 1*, we investigated its presence in strictures. *Clostridium sensu stricto 1* encompasses *Clostridium* species *C. tetani, C. botulinum, C. kluyveri, C. acetobutylicum, C. novyi, C. perfringens* and *C. beijerinckii*, which are considered pathogenic and indicate less healthy and less diverse microbiota^35^. Using representative sequences assigned to *Clostridium sensu stricto 1* ASVs, we determined that 232,156 (69%) reads shared >99% identity over the 370 bp sequence with *C. perfringens*. Remaining reads matched with *C. baratii* (19%), *C. colicanis* (7%) and unknown *Clostridium* species (6%). Importantly, ASV256, which increased 6-fold in progressors, shared 100% identity with *C. perfringens*. We then sought to confirm the presence of this organism by culture and 16S rRNA Sanger sequencing of clinical isolates. *C. perfringens* was isolated in 4 of 9 duodenum samples from marmosets with histologically confirmed strictures. *C. baratii* or *C. sardiniense*, a rare causative agent of botulism^36^, was isolated from 1 of 9 samples. As the pathogen was isolated at the stricture site, we analyzed the microbiome in duodenal samples from stricture (n=17) and non-stricture cases (n=12) to determine if increased gut abundance reflected a duodenal infection. *Clostridium sensu stricto 1* was observed at greater than 1% abundance in 76% of strictures (13/17) but only in 16% of non-stricture cases (2/12). In 8 stricture cases, the bacterium was the most abundant genus with abundances ranging from 37-87%. Interestingly, one non-stricture sample with 30% abundance of *Clostridium sensu stricto 1* had duodenal pathology characterized by mild duodenal mucosal congestion (**Fig 2d**).

Next, we developed RF models using serum chemistry profiles or CBC data to categorize stricture progressors and non-progressors. 4 serum chemistry parameters (total protein, lipase, GGT and amylase) classified progressors and non-progressors with 84.8% accuracy, a sensitivity of 76.5%, a specificity of 93.8% and AUC of 0.89 (**Fig 2c, Supp. Fig. 4d-f**). Total protein and GGT decreased in stricture cases, while lipase and amylase increased. Using CBC data, the RF classifier used HCT, HGB, RBC, RDW, MCH and lymphocyte percentage to identify strictures with an accuracy of 82.8%, sensitivity of 89.4%, a specificity of 75% and AUC of 0.83 (**Fig. 2c, Supp. Fig. 4g-i**). All variables, except RDW, decreased in strictures. Of note, weight was excluded from these models as severe weight loss is observed, which masked the contribution of other parameters. Exclusion of weight data did not decrease the predictive power of serum chemistry or CBC models.

### Effects of IBD on the microbiome and blood analysis

While strictures were observed predominantly in MIT^NE^, IBD was diagnosed in the four sources. “Progressors” were diagnosed with or developed IBD during the study, while “non-progressors” remained healthy or developed non-IBD diseases. Across the colony, microbiome richness decreased in IBD progressors (Chao1, P<0.001; Observed OTUs, P<0.001) but these changes were not observed when accounting for evenness (Shannon’s index and Pielou’s evenness) (**Fig. 3a**). We used PCA to determine if progressors converged at a common dysbiotic state (**Supp. Fig. 5a**). Similar to human IBD studies^3,5,37^, overall differences in the gut microbiota were observed but no individual microbes were consistently associated with IBD across all sources. Even in disease, community structures were source-dependent. However, positive, IBD-associated shifts along the first principal component (PC) were observed within sources (**Fig. 3b, Supp. Fig. 5a**). Changes in PC1 position were significantly different between healthy and IBD cases in the entire dataset (All, P<0.001), and in 3 of 4 sources (MITB, P<0.01; MIT^CL^, P<0.001; MITA, P<0.05; MIT^NE^, P=0.6) (**Fig 3b**). While no single dysbiotic IBD state existed, source-specific, healthy states could become source-specific, IBD states through similar perturbations of the microbiome.

**Figure 3.**
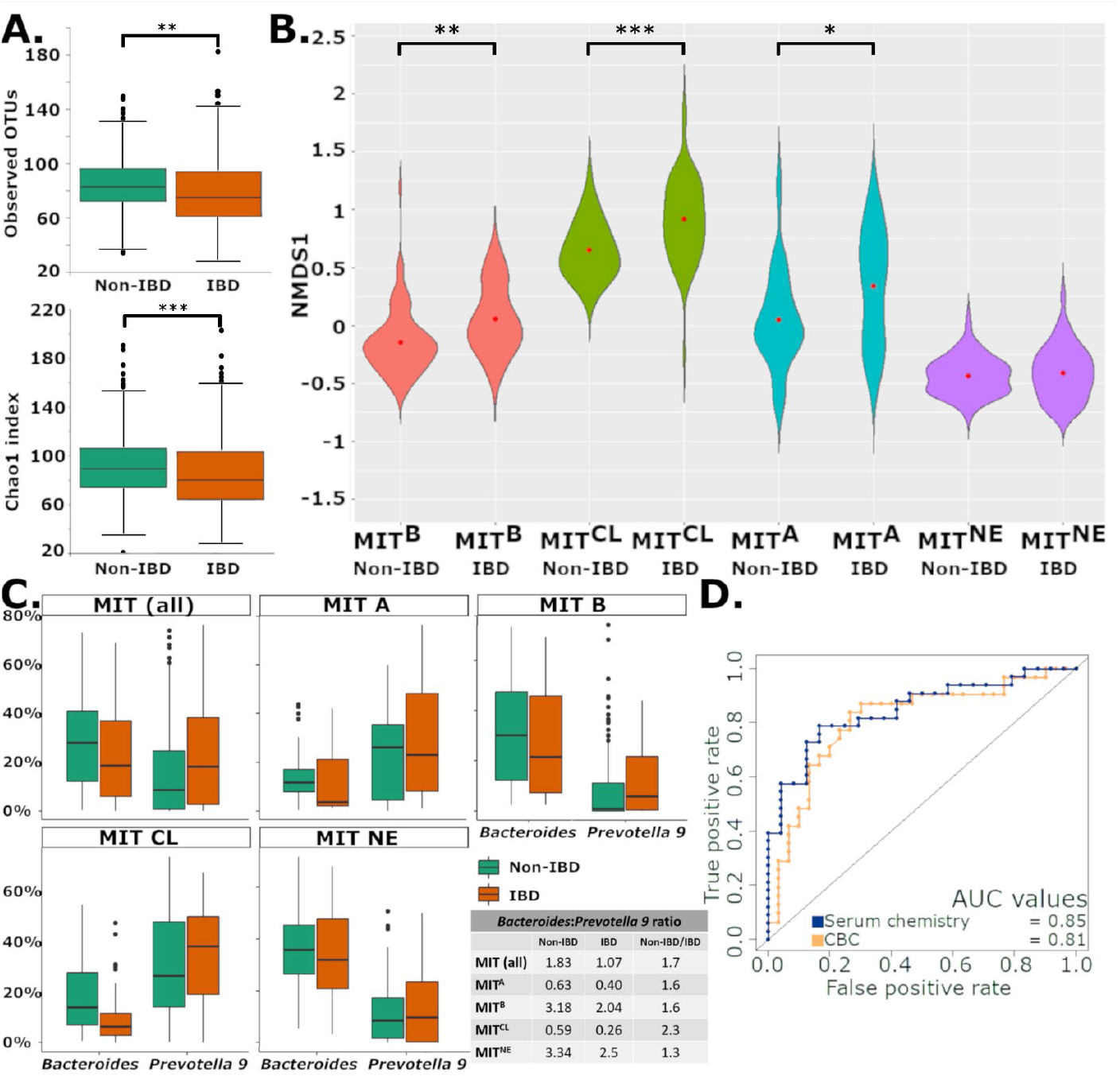
A) Decreased richness was observed in IBD marmosets (Observed OTUs and Chao1) compared to non-IBD marmosets similar to what is observed in humans. B) Increases in PC1 relative to source-specific, non-IBD controls were observed in 3 of 4 sources. C) *Bacteroides* and *Prevotella 9* levels are shown by source and IBD status. A lower overall and source-specific *Bacteroides:Prevotella 9* ratio is observed in IBD cases regardless of source-specific differences in abundances of these two genera. D) AUC of ROC for random forest models using serum chemistry and CBC show strong performance of models in classifying IBD progressors and non-progressors. *, *P<0.05;* **, *P<0.01* and ***, *P<0.001*.

To identify IBD-associated patterns observed in the dataset and within source-specific subsets, we examined ASVs correlated with PC1. Five *Prevotellaceae* ASVs (*Prevotella 9* and unclassified genera) and 3 *Megasphaera* ASVs were positively correlated with PC1, while 5 *Parabacteroides* ASVs and 3 *Bacteroides* ASVs were anti-correlated to PC1. Due to their importance in our analysis and the human gut microbiome^7,38^, we examined the relationship between *Bacteroides* and *Prevotella 9* in marmoset IBD. Using BLAST, 99.93% of *Prevotella 9* reads matched *P. copri* with a >99% identity. *Bacteroides* reads matched multiple species including *B. plebeius* (48.3%), *B. vulgatus* (16.8%), *B. uniformis* (6.4%), *B. dorei* (4.3%), *B. massiliensis* (3,2%), *B. thetaiotaomicron* (1.9%), *B. ovatus* (1.9%) and *B. coprocola* (1%). Decreases in *Bacteroides* were observed in IBD progressors, while *Prevotella 9* remained level or increased (**Fig. 3c**). Overall, the ratio of average *Bacteroides* abundance to average *Prevotella 9* abundance was 1.83 in non-progressors and 1.07 in IBD progressors, yielding a non-progressor/progressor ratio of 1.7. Similar non-progressor/progressor ratios were observed in all subsets, implying increased *P. copri* levels relative to *Bacteroides* spp. in marmoset IBD (**Fig. 3c**). We also developed 4 RF models to classify progressors and non-progressors using data from the entire colony, MIT^B^, MIT^CL^ and MIT^NE^ (MITA excluded due to insufficient n). The top 25 ASVs in each model were compared, and 8 ASVs shared by at least 3 models were identified (**Supp. Table 4**). ASVs belonged to *Sutterella* (3), *Megamonas* (2)*, Bacteroides, Asteroleplasma*, and *Prevotella 9*. Overlap in genera was determined by collapsing the 4 lists of ASVs by genus. Half of these ASVs belonged to 5 genera: *Bacteroides* (n=20), *Sutterella* (n=10), *Megamonas* (n=8), *Bifidobacterium* (n=7) and *Prevotella 9* (n=7). These results suggest that shifts in *Bacteroides*, *P. copri*, *Megamonas* and *Sutterella* are observed in IBD progressors relative to source-specific, healthy states.

Unlike the microbiome data, source-dependent clustering was not observed in serum chemistry or CBC PCA plots of IBD marmosets (**Supp. Fig. 5b-c**). Therefore, RF classifiers were trained solely on IBD status. The serum chemistry RF model used 7 variables (calcium, GGT, albumin, A:G ratio, amylase, cholesterol, and alkaline phosphatase), and had an accuracy of 77%, a sensitivity of 79%, a specificity of 76% and AUC of 0.85 (**Fig. 3d, Supp. Fig. 5d**). The optimized CBC RF model used HGB, RBC, RDW, MPV and neutrophil percentage, and had an accuracy of 77%, a sensitivity of 73%, a specificity of 83% and AUC of 0.81 (**Fig. 3d, Supp. Fig. 5e**). In these models, calcium, hemoglobin and RBC were the most important in classifying IBD.

### Effects of GI disease on gene expression of the small intestine

We tested whether strictures or IBD significantly altered marmoset transcriptomic profiles using RNA sequencing (RNAseq) on samples from IBD (n=3) or stricture (n=3) marmosets. Marmosets with strictures presented with gross thickening, duodenal stricture or ulceration (0.5-1cm aboral to the major duodenal papilla). Duodenal tissue evaluated was immediately distal to the lesion (“stricture”) or in an equivalent anatomic region in IBD animals (“non-stricture”). While IBD animals served as non-stricture controls, thickened intestines were observed grossly, and duodenitis was noted. As the most affected intestinal site in IBD^13^, we selected the jejunum to evaluate IBD effects. Unlike the IBD duodenum, the jejunum of stricture cases presented minimal pathology^22^, and were used as “non-IBD,” jejunum controls.

Comparing stricture and non-stricture duodenums, we identified 1,183 differentially expressed genes (DEG) (FDR <0.05) (**Fig 4a, Supp. Table 5**). To perform Gene ontology (GO) analysis, marmoset genes with official names were matched to *Homo sapiens* genes to retrieve Entrez IDs with associated GO categories. Analysis of this gene subset identified 903 DEGs with GO annotations. The top 15 biological processes (BP) with significant enrichment are listed in **Table 2** (complete list – **Supp. Table 6**). Stricture samples enriched BP sets involved with intestinal absorption, and lipid metabolism, localization and transport (**Fig 4b, Supp. Fig. 6a**). Stricture upregulated genes encompassed cholesterol-associated genes including apoliproteins (*APOB, APOA1* and *APOA4*), transport genes (*ABCG5, ABCG8, GRAMD1B*, and *STARD3*), metabolic genes (*DGAT1, CYP11A1*, and *CYP27A1*) and binding/absorption genes (*SOAT2, NPC1L1* and *SCARB1*) (**Supp. Table 5a**). Other lipid-associated genes upregulated by stricture included genes associated with fatty acid binding proteins (*FABP1* and *FABP2*), peroxisomes (*PPARA, ABCD1, ACAA1* and *EPHX2*), ketogenesis (*HMGCS2*) and lipid synthesis (*GPAM, SREBF1, SCAP*, and *ACACB*). Enriched cellular membrane GO sets shared these lipid-associated genes due to functional overlap (**Supp. Table 6, Supp. Fig. 6b**). Interestingly, immunity-associated genes were more highly expressed in non-stricture duodenums (**Fig 4b, Supp. Fig. 7**), possibly due to the enteritis observed in IBD marmosets. These genes included antimicrobial responses (*LCN2, LYZ, MUC20*), toll-like receptors (*TLR2* and *TLR4*) superoxide-generating NADPH oxidase activity (*NOX1* and *DUOX2*) killer cell lectin-like receptor genes (*KLRB1, KLRC1, KLRD1*, and *KLRF1*), and chemokine activity and receptor binding (*CXCL1, CXCL10, TFF2* and *PF4*) (**Supp. Table 5b**). The transcriptional profile implies the activity of natural killer (NK) cells, neutrophils and MHC class I protein complex binding.

**Figure 4.**
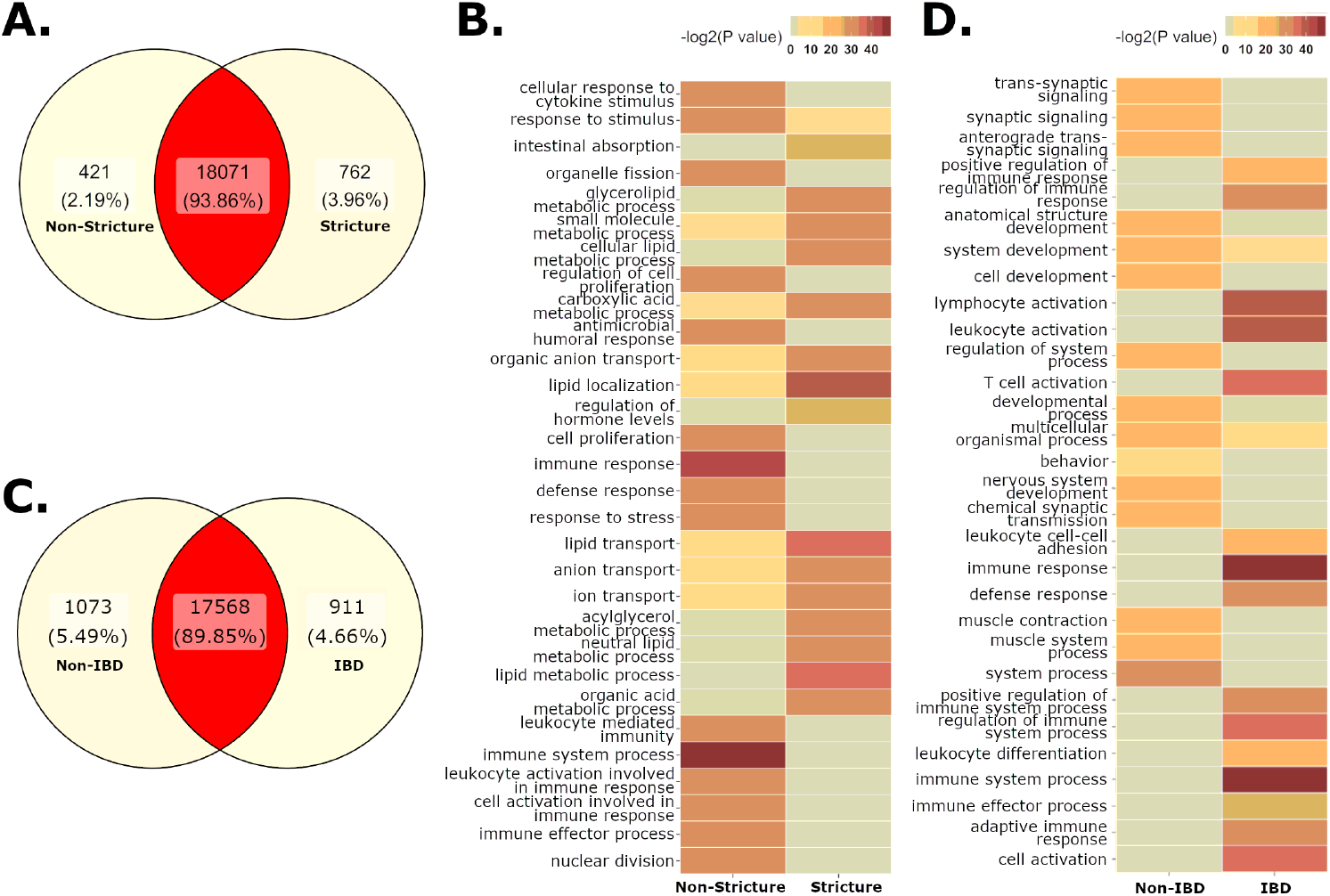
A) Differentially expressed genes (DEG)(FDR <0.05) in the duodenum of non-stricture and stricture cases. B) Gene ontology (GO) sets enriched in stricture cases show upregulation of lipid metabolism, transport and localization. Non-stricture cases have enrichment of immune processes, possibly due to underlying pathology caused by IBD. C) DEG (FDR <0.05) in the jejunum of non-IBD and IBD cases. D) IBD samples are enrich GO sets associated with immunity and immune cell activation.

**Table 2.**
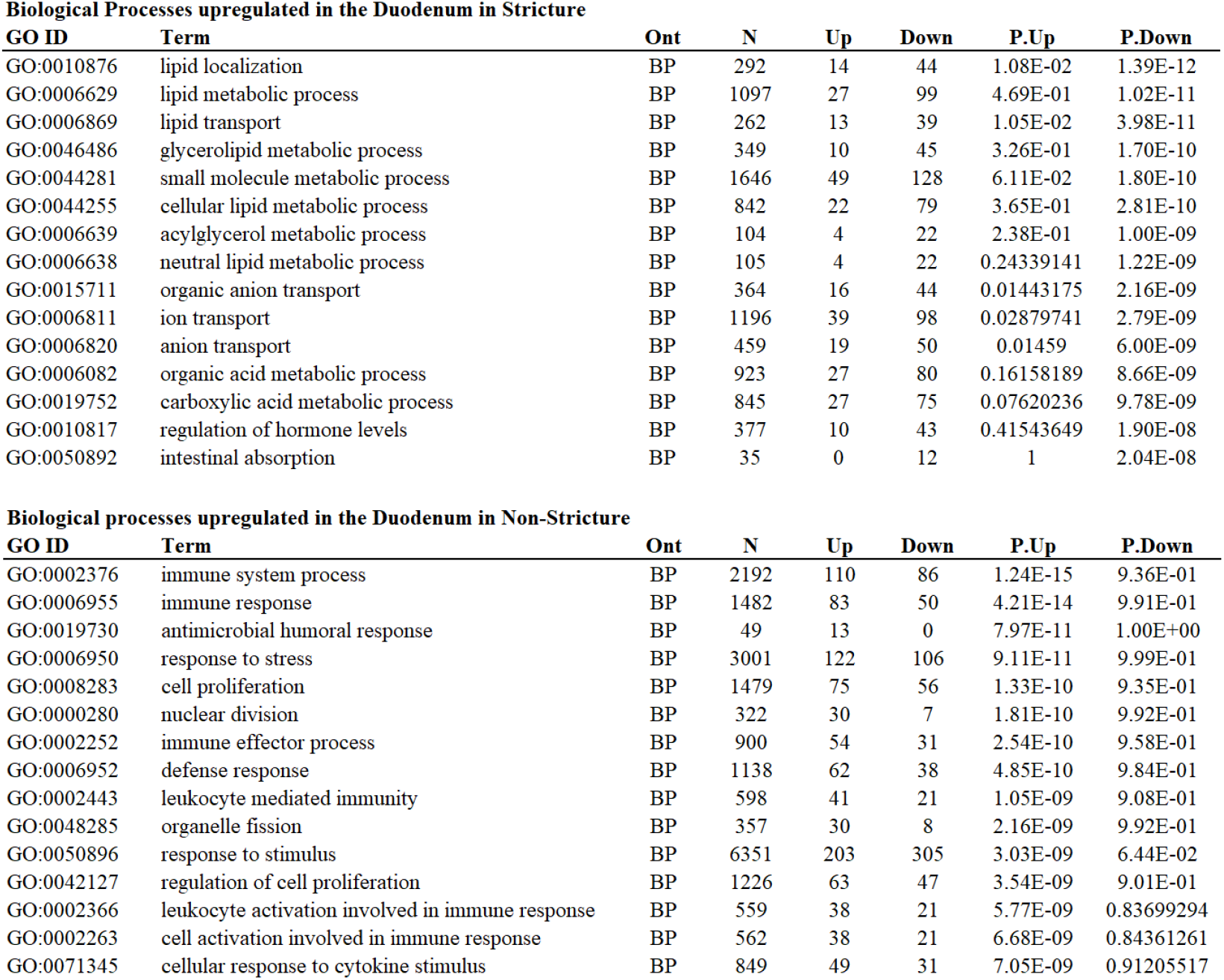
Top Gene Ontology sets observed in RNAseq analysis of stricture progressors and non-progressors

| GO ID | Term | Ont | N | Up | Down | P.Up | P.Down |
| --- | --- | --- | --- | --- | --- | --- | --- |
| GO:0010876 | lipid localization | BP | 292 | 14 | 44 | 1.08E-02 | 1.39E-12 |
| GO:0006629 | lipid metabolic process | BP | 1097 | 27 | 99 | 4.69E-01 | 1.02E-11 |
| GO:0006869 | lipid transport | BP | 262 | 13 | 39 | 1.05E-02 | 3.98E-11 |
| GO:0046486 | glycerolipid metabolic process | BP | 349 | 10 | 45 | 3.26E-01 | 1.70E-10 |
| GO:0044281 | small molecule metabolic process | BP | 1646 | 49 | 128 | 6.11E-02 | 1.80E-10 |
| GO:0044255 | cellular lipid metabolic process | BP | 842 | 22 | 79 | 3.65E-01 | 2.81E-10 |
| GO:0006639 | acylglycerol metabolic process | BP | 104 | 4 | 22 | 2.38E-01 | 1.00E-09 |
| GO:0006638 | neutral lipid metabolic process | BP | 105 | 4 | 22 | 0.24339141 | 1.22E-09 |
| GO:0015711 | organic anion transport | BP | 364 | 16 | 44 | 0.01443175 | 2.16E-09 |
| GO:0006811 | ion transport | BP | 1196 | 39 | 98 | 0.02879741 | 2.79E-09 |
| GO:0006820 | anion transport | BP | 459 | 19 | 50 | 0.01459 | 6.00E-09 |
| GO:0006082 | organic acid metabolic process | BP | 923 | 27 | 80 | 0.16158189 | 8.66E-09 |
| GO:0019752 | carboxylic acid metabolic process | BP | 845 | 27 | 75 | 0.07620236 | 9.78E-09 |
| GO:0010817 | regulation of hormone levels | BP | 377 | 10 | 43 | 0.41543649 | 1.90E-08 |
| GO:0050892 | intestinal absorption | BP | 35 | 0 | 12 | 1 | 2.04E-08 |

**Biological processes upregulated in the Duodenum in Non-Stricture**
| GO ID | Term | Ont | N | Up | Down | P.Up | P.Down |
| --- | --- | --- | --- | --- | --- | --- | --- |
| GO:0002376 | immune system process | BP | 2192 | 110 | 86 | 1.24E-15 | 9.36E-01 |
| GO:0006955 | immune response | BP | 1482 | 83 | 50 | 4.21E-14 | 9.91E-01 |
| GO:0019730 | antimicrobial humoral response | BP | 49 | 13 | 0 | 7.97E-11 | 1.00E+00 |
| GO:0006950 | response to stress | BP | 3001 | 122 | 106 | 9.11E-11 | 9.99E-01 |
| GO:0008283 | cell proliferation | BP | 1479 | 75 | 56 | 1.33E-10 | 9.35E-01 |
| GO:0000280 | nuclear division | BP | 322 | 30 | 7 | 1.81E-10 | 9.92E-01 |
| GO:0002252 | immune effector process | BP | 900 | 54 | 31 | 2.54E-10 | 9.58E-01 |
| GO:0006952 | defense response | BP | 1138 | 62 | 38 | 4.85E-10 | 9.84E-01 |
| GO:0002443 | leukocyte mediated immunity | BP | 598 | 41 | 21 | 1.05E-09 | 9.08E-01 |
| GO:0048285 | organelle fission | BP | 357 | 30 | 8 | 2.16E-09 | 9.92E-01 |
| GO:0050896 | response to stimulus | BP | 6351 | 203 | 305 | 3.03E-09 | 6.44E-02 |
| GO:0042127 | regulation of cell proliferation | BP | 1226 | 63 | 47 | 3.54E-09 | 9.01E-01 |
| GO:0002366 | leukocyte activation involved in immune response | BP | 559 | 38 | 21 | 5.77E-09 | 0.83699294 |
| GO:0002263 | cell activation involved in immune response | BP | 562 | 38 | 21 | 6.68E-09 | 0.84361261 |
| GO:0071345 | cellular response to cytokine stimulus | BP | 849 | 49 | 31 | 7.05E-09 | 0.91205517 |

1,984 DEGs were identified when comparing jejunums from IBD and non-IBD marmosets (**Fig 4c, Supp. Table 7**) following the exclusion of an IBD sample that did not cluster with other samples (**Supp. Fig. 8**). GO annotations were assigned to 1,586 DEGs, and the top 15 BP are summarized in **Table 3a** (complete list - **Supp. Table 8**). As observed in non-stricture duodenum (IBD) samples, the jejunum of IBD animals enriched GOs associated with host immunity, such as T cell activation, adaptive immune responses, and regulation of immune response (**Fig 4d, Supp. Fig. 9**). As observed in the non-stricture duodenum, genes associated with killer cell lectin-like receptors (*KLRB1, KLRC1, KLRC2, KLRF1*, and *KLRK1*) and antimicrobial responses (*LCN2, LYZ*, and *MUC20*) were upregulated in the jejunum of IBD marmosets. Genes involved in the adaptive immunity and T cell activation (*EOMES, PRF1, IFNG, FYN, CD160, CD244, CD3G, TBX21, CD27, PTPRC*, and *IL18R1*) had increased expression in IBD samples. (**Supp. Table 7**). In non-IBD animals, top GOs associated with homeostatic functions, such as synaptic signaling, development, and muscle contraction (**Table 3b, Supp. Fig. 10**).

**Table 3.**
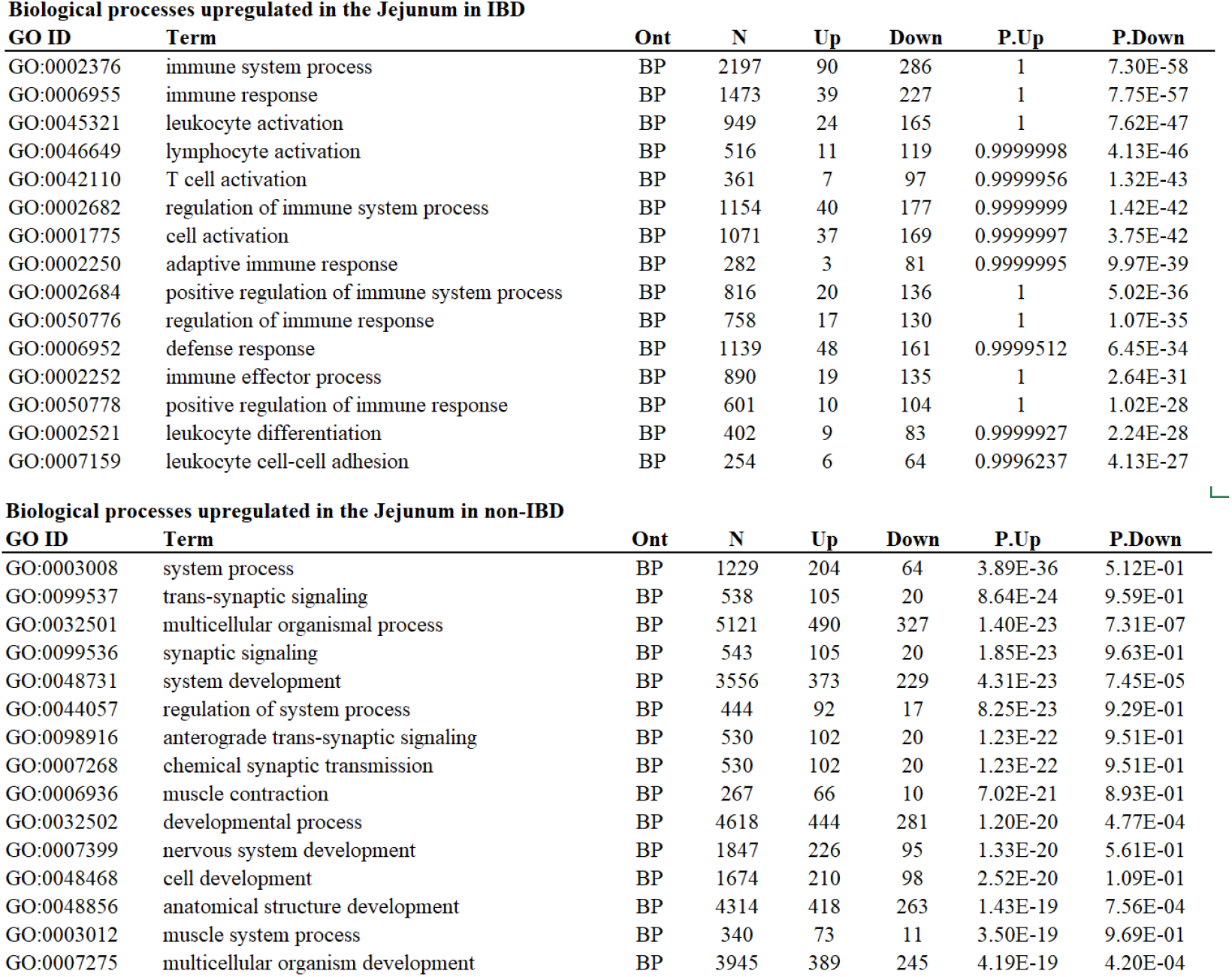
Top Gene Ontology sets observed in RNAseq analysis of IBD progressors and non-progressors

| GO ID | Term | Ont | N | Up | Down | P.Up | P.Down |
| --- | --- | --- | --- | --- | --- | --- | --- |
| GO:0002376 | immune system process | BP | 2197 | 90 | 286 | 1 | 7.30E-58 |
| GO:0006955 | immune response | BP | 1473 | 39 | 227 | 1 | 7.75E-57 |
| GO:0045321 | leukocyte activation | BP | 949 | 24 | 165 | 1 | 7.62E-47 |
| GO:0046649 | lymphocyte activation | BP | 516 | 11 | 119 | 0.9999998 | 4.13E-46 |
| GO:0042110 | T cell activation | BP | 361 | 7 | 97 | 0.9999956 | 1.32E-43 |
| GO:0002682 | regulation of immune system process | BP | 1154 | 40 | 177 | 0.9999999 | 1.42E-42 |
| GO:0001775 | cell activation | BP | 1071 | 37 | 169 | 0.9999997 | 3.75E-42 |
| GO:0002250 | adaptive immune response | BP | 282 | 3 | 81 | 0.9999995 | 9.97E-39 |
| GO:0002684 | positive regulation of immune system process | BP | 816 | 20 | 136 | 1 | 5.02E-36 |
| GO:0050776 | regulation of immune response | BP | 758 | 17 | 130 | 1 | 1.07E-35 |
| GO:0006952 | defense response | BP | 1139 | 48 | 161 | 0.9999512 | 6.45E-34 |
| GO:0002252 | immune effector process | BP | 890 | 19 | 135 | 1 | 2.64E-31 |
| GO:0050778 | positive regulation of immune response | BP | 601 | 10 | 104 | 1 | 1.02E-28 |
| GO:0002521 | leukocyte differentiation | BP | 402 | 9 | 83 | 0.9999927 | 2.24E-28 |
| GO:0007159 | leukocyte cell-cell adhesion | BP | 254 | 6 | 64 | 0.9996237 | 4.13E-27 |

**Biological processes upregulated in the Jejunum in non-IBD**
| GO ID | Term | Ont | N | Up | Down | P.Up | P.Down |
| --- | --- | --- | --- | --- | --- | --- | --- |
| GO:0003008 | system process | BP | 1229 | 204 | 64 | 3.89E-36 | 5.12E-01 |
| GO:0099537 | trans-synaptic signaling | BP | 538 | 105 | 20 | 8.64E-24 | 9.59E-01 |
| GO:0032501 | multicellular organismal process | BP | 5121 | 490 | 327 | 1.40E-23 | 7.31E-07 |
| GO:0099536 | synaptic signaling | BP | 543 | 105 | 20 | 1.85E-23 | 9.63E-01 |
| GO:0048731 | system development | BP | 3556 | 373 | 229 | 4.31E-23 | 7.45E-05 |
| GO:0044057 | regulation of system process | BP | 444 | 92 | 17 | 8.25E-23 | 9.29E-01 |
| GO:0098916 | anterograde trans-synaptic signaling | BP | 530 | 102 | 20 | 1.23E-22 | 9.51E-01 |
| GO:0007268 | chemical synaptic transmission | BP | 530 | 102 | 20 | 1.23E-22 | 9.51E-01 |
| GO:0006936 | muscle contraction | BP | 267 | 66 | 10 | 7.02E-21 | 8.93E-01 |
| GO:0032502 | developmental process | BP | 4618 | 444 | 281 | 1.20E-20 | 4.77E-04 |
| GO:0007399 | nervous system development | BP | 1847 | 226 | 95 | 1.33E-20 | 5.61E-01 |
| GO:0048468 | cell development | BP | 1674 | 210 | 98 | 2.52E-20 | 1.09E-01 |
| GO:0048856 | anatomical structure development | BP | 4314 | 418 | 263 | 1.43E-19 | 7.56E-04 |
| GO:0003012 | muscle system process | BP | 340 | 73 | 11 | 3.50E-19 | 9.69E-01 |
| GO:0007275 | multicellular organism development | BP | 3945 | 389 | 245 | 4.19E-19 | 4.20E-04 |

## Discussion

GI diseases are the most prevalent clinical disease in captive common marmosets^13,14,39^, but the role of the microbiome is largely unknown. Recent literature demonstrates that NHP captivity affects bacterial composition, reduces alpha diversity, and alters host responses to disease^23,26,40^. In captivity, NHP microbiomes lose distinctive, wild microbiota and become dominated by *Prevotella* and *Bacteroides*, the most abundant genera in the modern human gut microbiome^7,23,38^. In the largest marmoset microbiome study to date, our data supports the hypothesis that captivity humanizes the primate microbiome, as *Bacteroides* and *Prevotella 9* were the most abundant genera with levels similar to those observed in human feces^7,38^. In humans, *Prevotella* and *Bacteroides* abundances are anticorrelated, signifying that competitive advantages in metabolism determine the dominant bacteria^41,42^. *Prevotella* increases have been associated with high-fiber, plant-based diets and non-industrialized populations, while *Bacteroides* increases were linked to Westernized populations with diets rich in animal fat and protein^41,42^. Diets influence levels of fibers, fermentation products, SCFA and bile acids (BA), which determine bacterial communities^42^. As our marmosets were fed a standardized diet, dietary differences cannot account for the *Prevotella*- and *Bacteroides*-dominant profiles observed stably in our colony. Most bacteria observed were acetate- or propionate-producers, such as *Bacteroides, Prevotella, Anaerobiospirillum, Phascolarctobacterium, Megamonas*, and *Megasphaera*, with a low abundance of butyrate producers, such as *Lachnospiraceae*^43^. However, *Megasphaera* has been known to produce butyrate under specific conditions^33,44^. Inter-institutional differences greatly affect marmoset microbiomes, as previous studies report marmoset gut microbiota dominated by *Actinobacteria^29,30^, Firmicutes^33,34^, Proteobacteria*^24,45,46^ and *Bacteroidetes*^31,32,46^. At the Biomedical Primate Research Centre (BPRC) (Rijswijk, the Netherlands), *Actinobacteria*, represented by *Bifidobacterium* and *Collinsella*, was the most abundant phylum (66%), while *Bacteroides* and *Prevotella* represented <5% of the microbiome each^29^. BPRC marmosets have access to outdoor and indoor enclosures, as well as food enrichment, such as insects and gums, several times a week^29^. We hypothesize that increased environmental exposure and enrichment promotes a wild-like microbiome, rich in bifidobacteria that help metabolize oligosaccharide-rich tree gums, a common food source for wild marmosets^45,47^. High abundances of *Actinobacteria* are observed in wild callitrichids, but not in captive and semi-captive marmosets^24^. Unexpectedly, Ross et al. also reported high *Bifidobacterium* levels in marmosets housed within a specific-pathogen free (SPF) barrier facility and at the Southwest National Primate Resource Center (SNPRC)^30^. In contrast to the BPRC, SPF marmosets fed exclusively irradiated feed, nuts, seeds and dried fruits had median *Bifidobacterium* abundances of 17%^30^. This was much higher than the non-SPF parent colony at SNPRC, which had median *Bifidobacterium* frequencies of 4% and high levels of *Fusobacterium*^30^. However, a follow-up report from the barrier facility showed bacterial shifts with an increased *Bacteroidetes* abundance (35%) and a slight decrease in *Bifidobacteriaceae* (12%)^31^. Similar to our study, few age-related changes in the microbiome were observed^31^. In a colony with a microbiome similar to the MIT profile, microbiome synchronization occurred within a year in imported marmosets, characterized by expansion of *Bacteroidetes*^46^. Imported cohorts retained unique features following microbiome synchronization^46^, supporting our findings that source-specific microbiomes persist despite standardization of husbandry and diet. These studies demonstrate that inter-institutional differences can promote stable microbiomes in clinically healthy animals across a large range of bacterial compositions. In other NHP, wild-like microbiota may prevent captivity-associated illnesses^23^. The resilience to perturbations of different bacterial compositions in marmosets is unknown. Understanding and manipulating the marmoset microbiome may help prevent disease, and due to their importance as research models in neuroscience, aging, and toxicology, having marmosets with “humanized” microbiota may better represent the human condition.

In this study, we evaluated a marmoset colony with a “humanized” microbiota^48^ and compared the microbiota of clinically healthy individuals with marmosets with two GI diseases. While captivity increases susceptibility to GI disease, we observed source-specific differences in disease prevalence. MIT^NE^ marmosets had the highest *Bacteroidaceae* abundance (37%) and the lowest *Prevotellaceae* levels (17%), and were most susceptible to strictures, a novel GI disease in marmosets^21,22^. This duodenal syndrome was found in 21.9% of necropsy cases in an institution^21^, while MIT^NE^ marmosets had a 26% prevalence. Clinical signs include vomiting, bloating, weight loss and a palpable thickening of the duodenum that can be visualized through radiography and ultrasound^21,22^. Stricture-associated dysbiosis featured reductions in *Bacteroides* and *Anaerobiospirillum*, and *Megamonas* increases. Our analysis of strictures highlighted the importance of decreases in *Anaerobiospirillum* and increases in *Clostridium sensu stricto 1*. While *Anaerobiospirillum* has been previously reported in healthy marmosets, dogs and cats^45,49^, these bacteria may cause GI disease in humans^49^. However, *Anaerobiospirillum* was present in high abundances in our healthy marmosets, and reduced levels were seen in disease.

As *C. perfringens* was detected at higher levels in the duodenal lesions of diseased animals by culture and sequencing, we propose that *C. perfringens* is a potential causative agent of duodenal disease in marmosets. *C. perfringens* is a known GI pathogen that can encode multiple toxins (alpha, beta, epsilon, iota, perfringolysin O, and enterotoxin)^35^. In marmosets and other NHP, *C. perfringens* can cause gas gangrene and gastric dilatation syndrome^50–52^. Of note, *C. perfringens*- induced gas gangrene was reported in the institution that first reported duodenal strictures^50^. In the United Kingdom, *C. perfringens* is one of the top 5 causes of foodborne death^53^, and has been linked to diarrhea, *Clostridial* necrotizing enteritis (CNE), necrotizing enterocolitis (NEC), UC and enterotoxemia in humans and other mammals^35,54^. CNE is a necrotizing inflammation of the small intestine that can induce mild diarrhea or severe abdominal pain, vomiting and ulcers^35^. NEC predominantly affects infants due to intestinal immaturity or dysbiosis^35,54^. While these symptoms match the clinical presentation of duodenal strictures in marmosets, they are non-specific. However, intestinal strictures developed in 11-29.5% of NEC infants in both small and large intestines and could occur up to 20 months post-diagnosis^55,56^. Based on the site of *C. perfringens* infection at the junction of the duodenum and the common bile duct, we hypothesize that BA deregulation due to dysbiosis or antibiotic treatment may facilitate *C. perfringens* infection. Antibiotic usage in infants has been linked with increased NEC risk^57^, and antibiotics are commonly prescribed to treat NHP GI diseases. Furthermore, *C. perfringens* was overrepresented in dogs with chronic enteropathy, an IBD-like disease, and bacterial abundance was regulated by secondary BAs, deoxycholic acid and lithocholic acid, that are produced by gut bacteria^58,59^.

In addition to the role of *C. perfringens*, our serum chemistry and CBC-based RF models were highly sensitive in classifying strictures. Decreased total protein levels are often observed with GI disease and may indicate poor digestion/absorption. The importance of amylase and lipase in our stricture model is supported by clinical findings of cholecystitis and secondary pancreatitis^22^. Secondary pancreatitis, attributed to extension from the duodenal ulcer, was observed in 15 of 17 cases scored^22^. In the CBC-based model, HCT, HGB, RBC, RDW, and MCH relate to red blood cell function and suggested anemia. Anemia, a common finding in marmosets with strictures and IBD^15,22^, is also a risk factor for NEC in humans^60^. Interestingly, transcriptomic analysis of strictures showed enrichment of lipid metabolism and intestinal absorption genes, which may reflect enterocyte damage and is consistent with lipidomic alterations induced by *C. perfringens* alpha-toxin, a phospholipase C^61^. Increased expression of *FABP1* and *FABP2* was observed. These genes encode for liver and intestinal fatty-acid binding proteins (LFABP and IFABP), respectively, and are often used as biomarkers of GI diseases, including NEC^57^. To our knowledge, correlations of gut *FABP2* levels with serum IFABP levels have not been described, but we hypothesize that increased expression might be a compensatory mechanism triggered by enteritis. While increased inflammatory responses were not observed due to the lack of healthy control tissue, based on the *C. perfringens* infection, development of enteritis, anemia and strictures and deregulation of lipid metabolism, we believe marmosets could be developed as a model to investigate the mechanisms of bacterially-driven CNE/NEC.

In contrast, a unique microbial signature for IBD was not evident. Consistent with human studies, marmoset IBD decreased alpha diversity^3,11,37^. Human IBD is characterized by the loss of health-associated genera, such as *Roseburia, Faecalibacterium, Eubacterium, Ruminococcus* and *Subdoligranulum*^2,3,37,62^, but these bacteria have not been found in high abundance in marmosets^29–31^. Other potentially beneficial taxa in humans that have been observed in marmosets include *Bifidobacterium, Bacteroides, Collinsella*, and *Phascolarctobacterium*^3,62^. Increases in *Lactobacillus, Ruminococcus gnavus, Enterobacteriaceae*, *Pasteurellaceae*, *Veillonellaceae*, and *Fusobacteriaceae* have been associated with IBD^3,37,62^. While convergence to a single dysbiotic IBD state was not observed, multiple, source-specific states were associated with IBD. Within each source population, IBD progressors had higher average abundances of *P. copri* and *Megamonas*, as well as decreased abundance of *Bacteroides*, relative to controls. Our RF models also highlighted *Sutterella*, a bacteria associated with negative fecal microbiota transplantation outcomes, shorter remission periods in UC patients^63,64^, and its ability to dampen immune responses^65^. *Megamonas*, along with *B. plebeius*, deregulate BA metabolism in CD patients^66^, which could cause dysbiosis and opportunistic pathogen infections. However, while *Megamonas* increases were observed*, Bacteroides* decreased in marmoset IBD. Most *Bacteroides* reads matched *B. plebeius*, a non-*B. fragilis* group species^67^. *B. plebeius* ASVs were the most abundant in the two *Bacteroides*-dominated cohorts, and only 20% of *Bacteroides* reads matched members of the *B. fragilis* group, the most frequently isolated and virulent species in clinical specimens^68^. Furthermore, the role of the *B. fragilis* group in IBD is inconclusive, as they both modulate immunity and cause infections^3,68–70^.

While the effects of *Bacteroides* and *Prevotella* spp. in IBD patients have not been understood^3,71,72^, *Prevotella* have been considered inflammophilic pathobionts, commensal bacteria known to thrive in inflammatory environments and promote inflammatory diseases, such as periodontitis, bacterial vaginosis, rheumatoid arthritis (RA), and metabolic disorders^73–75^. *Prevotella*, including *P. copri*, activate TLR2, elicit specific IgA and IgG responses and promote the release of IL-1, IL-8, IL-6, IL-17, IL-23, and CCL20, which leads to neutrophil recruitment, reduced T helper 2 (Th2) cells and induction of Th17 cells^73–77^. In the gut, *Prevotella* has been linked to diarrhea, HIV-induced gut dysbiosis, irritable bowel syndrome and more severe colitis^78–80^. In a small study, higher levels of *Prevotella* were observed in marmosets with IBD compared to controls^46^. Furthermore, models of RA and colitis have shown that transfer of *Prevotella*- or *P. copri-rich* microbiota to mice transmitted disease phenotypes^74,77,78^. A possible mechanism could be linked to cycles of expansion and relaxation observed in *P. copri* abundance in healthy individuals, but absent in IBD patients^5^. Constant *P. copri* signals might promote chronic inflammation, but natural control of *P. copri* in the microbiome might prevent disease-causing chronic inflammatory states. In our study, IBD-associated enteritis upregulated pro-inflammatory immune responses in the duodenum and jejunum. Multiple genes associated with NK cell functions were upregulated by IBD, including genes associated with high cytolytic effector activity, cytotoxicity and IFN-γ production (*CD244, CD160, IL18R1, FYN*, and *IFNG*)^81,82^. In addition to *IFNG*, genes associated with Th1 cells (*TBX21, CCR2, CCR5*, and *IL2RB*) were also upregulated. In humans, killer immunoglobulin receptor (KIR) polymorphisms have linked NK cells with CD^83^. Further studies are needed to determine if *P. copri* causes enteritis and IBD in marmosets via NK cells.

This study is the largest evaluation of the captive marmoset microbiome, and is the first to systematically compare clinically healthy marmosets and marmosets with two GI disorders. The common marmoset may be a useful model to investigate *C. perfringens*-associated enteritis and intestinal strictures, as well as *P. copri*-mediated IBD. As observed in humans, a range of stable microbiome profiles may exist in clinically healthy marmosets. Better understanding of these profiles, the effects of diet and husbandry, and their inherent robustness to insults and disease will be helpful in promoting animal health, developing better models of human disease and understanding how to modulate microbial communities.

## Materials and Methods

### Animals

Common marmosets (*Callithrix jacchus*) were housed at the Massachusetts Institute of Technology in Cambridge, MA, from marmosets sourced from the New England Primate Research Center (NEPRC), an international primate center (CLEA Japan Inc.), and two biotech companies (A and B). Subsequently, the four sources will be referred to as MIT^NE^, MIT^CL^, MITA, and MIT^B^. All animals were housed in pairs or family groups within two vivaria at MIT, an AAALAC International accredited facility. Of the animals evaluated in this survey, 85 were male and 88 were female. All marmosets included in this study were on an animal use protocol approved by the MIT Institutional Care and Use Committee (IACUC).

The animal holding room temperature was maintained at 74.0 +/- 2°F with a relative humidity of 30 – 70%. The light cycle was maintained at a 12:12h light:dark cycle. Marmosets were housed in cages composed of stainless-steel bars and polycarbonate perches with the following dimensions: 30” W x 32” D x 67” H). Each cage had a nest box made of polycarbonate attached the outside of the cage. Other cage furniture present in the cages included hammocks, hanging toys, and manzanita wood branches. Foraging enrichment in the form of dried acacia gum-filled branches and forage board were provided weekly. Cages were removed for sanitization on a biweekly rotation.

All animals received a base chow diet of biscuits (Teklad New World Primate Diet 8794). Initially, biscuits were soaked in water for at least 20 minutes, but the practice was then changed to a pour-on/pour-off soak only. About halfway through the two-year period encompassing this study, biscuit prep protocol reverted to the original practice of a 20-minute soak to alleviate any concerns that soaking duration could be contributing to the development of duodenal ulcers. In addition to the base chow, a cafeteria-style supplemental offering of fruits, vegetables, and additional protein sources including hard-boiled eggs, mealworms, cottage cheese or ZuPreem (Premium Nutritional Products, Inc., Mission, KS).

On a semiannual basis, preventative health physical exams were performed on all colony animals. Rectal swabs and fecal samples were collected and screened for potentially pathogenic bacteria (including *Salmonella* spp., *Shigella* spp, beta-hemolytic *E.coli, Klebsiella* spp., and *Campylobacter* spp.) and parasites (including *Enterobius* spp., *Entamoeba* spp., *Giardia* spp., *Taenia* spp., and *Cryptosporidium* spp.). Intradermal testing for *Mycobacterium tuberculosis* was performed semiannually as well. All animals derived from progenitor stock were negative for squirrel monkey cytomegalovirus, *Saimiriine herpesvirus 1, Saimiriine herpesvirus 2*, and measles virus. Complete blood count and serum chemistry analysis were performed on an annual basis and during diagnostic workup of clinical cases. Hematology analysis was performed by the MIT DCM diagnostic laboratory using a HemaVet 950 veterinary hematology analyzer (Drew Scientific, Oxford, CT). Serum chemistry analysis was performed by Idexx Laboratories (Westbrook, ME). Serum chemistry and complete blood counts data were collected from the clinical records from the MIT colony. Fecal (n = 223) and rectal swab (n=342) were collected from common marmosets (*Callithrix jacchus*) (n = 565 samples, 173 individuals) between 2016-2018.

### Bacterial Culture Methods

Stricture samples containing duodenal tissue and duodenal contents were collected from animals during necropsies performed by clinical veterinarians and veterinary pathologists. Representative sections of major organs were collected, fixed in 10% neutral buffered formalin, embedded in paraffin, sectioned at 5 μm, and stained using hematoxylin and eosin (HE) for scoring by a boarded veterinary pathologist. Stricture samples were flash frozen in vials containing Brucella broth in 20% glycerol and frozen at −80°C. The tissues were thawed in an anaerobic atmosphere (10% CO2, 10% H2, 80% N2), and were homogenized with freeze medium with tissue grinders. The homogenate was divided into the following aliquots. For aerobic culture, the homogenates were plated onto chocolate agar, blood agar, MacConkey agar, and Brucella Broth medium containing 10% FCS. The plates were incubated at 37°C in 5% CO2 for 24-48 hours. For anaerobic culture, the homogenates were plated onto pre-reduced Brucella Blood Agar plates (BBL) and inoculated into thioglycollate broth. The cultures were incubated at 37° C in an anaerobic chamber (Coy Lab Products) with mixed gas (10% CO2, 10% H2, 80% N2) for 48 hours. For microaerobic culture to detect the growth of *Helicobacter* spp., the homogenates were plated onto selective antibiotic impregnated plates (50 μg/ml amphotericin B, 100 μg/ml vancomycin, 3.3 μg/ml polymyxin B, 200 μg/ml bacitracin, and 10.7 μg/ml nalidixic acid)^84^ and Brucella Blood Agar plates after passing through 0.65 μm syringe filter. The plates were placed into a vented jar filled with mixed gas (10% CO2, 10% H2, 80% N2) and incubated at 37°C for up to 3 weeks. The plates were checked every 2-3 days for growth. Aliquots of the homogenates were also used for DNA extraction. All bacterial strains isolated from the different culture conditions were identified by 16s rRNA sequencing.

### 16S microbiome profiling

Fecal DNA was extracted using the DNeasy PowerLyzer PowerSoil Kit, and DNA was amplified using universal primers of F515 (GTGYCAGCMGCCGCGGTAA) and R926 (CCGYCAATTYMTTTRAGTTT) to target the V4 and V5 regions of bacterial 16S rRNA fused to Illumina adaptors and barcode sequences as described previously.^85^ Individual samples were barcoded and pooled to construct the sequencing library, followed by sequencing with an Illumina MiSeq instrument to generate pair-ended 300 × 300 reads. Sequencing quality was inspected using FastQC^86^. Reads were processed using QIIME 2-2018.6 within the MicrobiomeHelper v. 2.3.0 virtual box^85,87^. Briefly, primer sequences were trimmed using the cutadapt plugin^88^. Forward and reverse reads were truncated at 243 and 195 bases, respectively, prior to stitching and denoising reads into amplicon sequence variants (ASV) using DADA2. Samples with fewer than 7,500 reads were excluded. ASVs present in fewer than 3 samples and with less than 24 counts were also excluded. A total of 1085 ASVs were retained after filtering. Taxonomic classification was assigned using the custom 16S V4/V5 region classifier based on the SILVA 132 database (SSU Ref NR 99)^89^. Phylogenetic trees, composition, alpha rarefaction, beta diversity metrics and ANCOM (Analysis of Composition of Microbiome)^90^ were evaluated using built-in QIIME2 functions^91^. Microsoft Excel and R (v 3.6.3 at http://www.R-project.org/) were used to perform statistical analyses and graphically represent data. Additionally R libraries ggplot2 (2.2.1)^92^, caret^93^, vegan^94^, pROC^95^, and gtools^96^ were used to model microbiome data. Classifiers were trained on 80% of the samples and the discovered signatures were used to predict the populations on the remaining 20% of samples (testing). We analyzed the *Bacteroides/Prevotella* abundance ratio by taking the ratio of the averaged *Bacteroides* abundance and the averaged *Prevotella* abundance.

### RNAseq

Tissues were collected from the duodenum and jejunum from marmosets with either stricture or IBD during necropsies performed by clinical veterinarians and veterinary pathologists. In stricture cases, duodenal samples were distal of the site of stricture (n=3), and in IBD cases, the same region of the duodenum presented with mild thickening based on gross observations (n=3). In IBD cases, the jejunum presented with increased thickening (n=3), while in stricture cases, the jejunum was grossly normal (n=3). Tissues were flash frozed in liquid nitrogen and stored at −80°C. RNA was extracted using TRIzol reagent according to manufacturer’s instructions (Thermo Fisher Scientific). Total RNA was shipped on dry ice to Arraystar, Inc. (Rockville, MD) for quality control, rRNA depletion and sequencing on an Illumina HiSeq4000. FASTA files and the NCBI RefSeq GTF files for *Callithrix jacchus* based on the March 2009 (WUBSC 3.2/calJac3) assembly were obtained from the UCSC Genome browser^97^. Raw sequencing reads were mapped to an index built from *C. jacchus* FASTA files using Rsubread^98^. Feature counts were obtained from the bam files using annotated exons in the *C. jacchus* GTF files. Analysis was then performed using edgeR^99,100^. Lowly expressed exons were removed using a cutoff of 10 counts per million (CPM). Normalization was performed using the Trimmed Mean of M-values (TMM) method. Multidimensional scaling (MDS) plots and heatmaps were used to evaluate grouping of biological samples. Data was fitted using the glmQLFit function that uses a generalized linear model (GLM) implementing a quasi-likelihood (QL) fitting method. Quasi-likelihood F-tests were performed to test for differential expression based on False Discovery Rate (FDR) adjusted P-values of 0.05. To retrieve Gene Ontology (GO) classifications, *C. jacchus* genes that matched *Homo sapiens* gene names were assigned both the *C. jacchus* and *Homo sapiens* Entrez IDs. GO analysis was performed using limma^101^, AnnotationDbi^102^, GO.db^103^, topGO^104^, mygene^105^ and org.Hs.eg.db. Data was visualized using ggplot2, gplots, Rgraphviz^106^, colorspace^107^ and ggVennDiagram^108^. Analysis of the IBD dataset demonstrated that the expression profile of one sample differed from the remaining samples and was excluded from the analysis presented.

## Data availability

RNAseq data is available under NCBI GEO accession number GSE156839. Microbiome data is available under NCBI BioProject PRJNA659472.

## Acknowledgements

This work was supported in part by a grant from the MIT McGovern Institute, NIH grant T32 OD010978 and by the National Institute of Environmental Health Sciences of the NIH under award P30-ES002109.

## Author contributions

Conception and design: AS, SCA, MAB, SM, JGF. Data acquisition, analysis and interpretation: AS, SCA, MAB, JAM, MAL, JD, SM. Manuscript: AS, SCA, JGF

