## Supplemental Figures for "Common Marmoset Gut Microbiome Profiles in Health and Intestinal Disease"

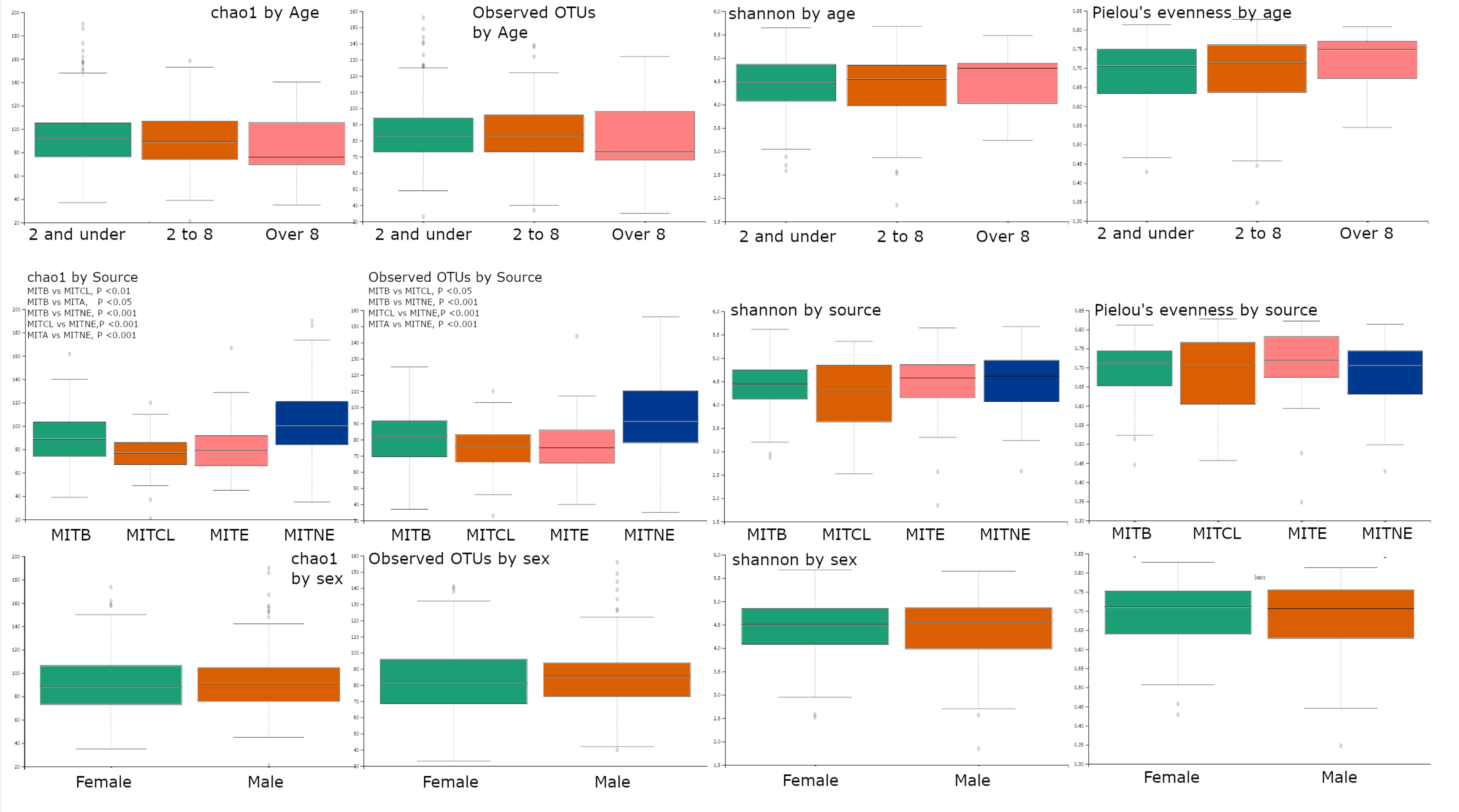


Supplemental Figure 1. Multiple alpha diversity metrics based on Age, Source and Sex. Significant differences were only observed when classifying by source with metrics accounting for richness but not evenness

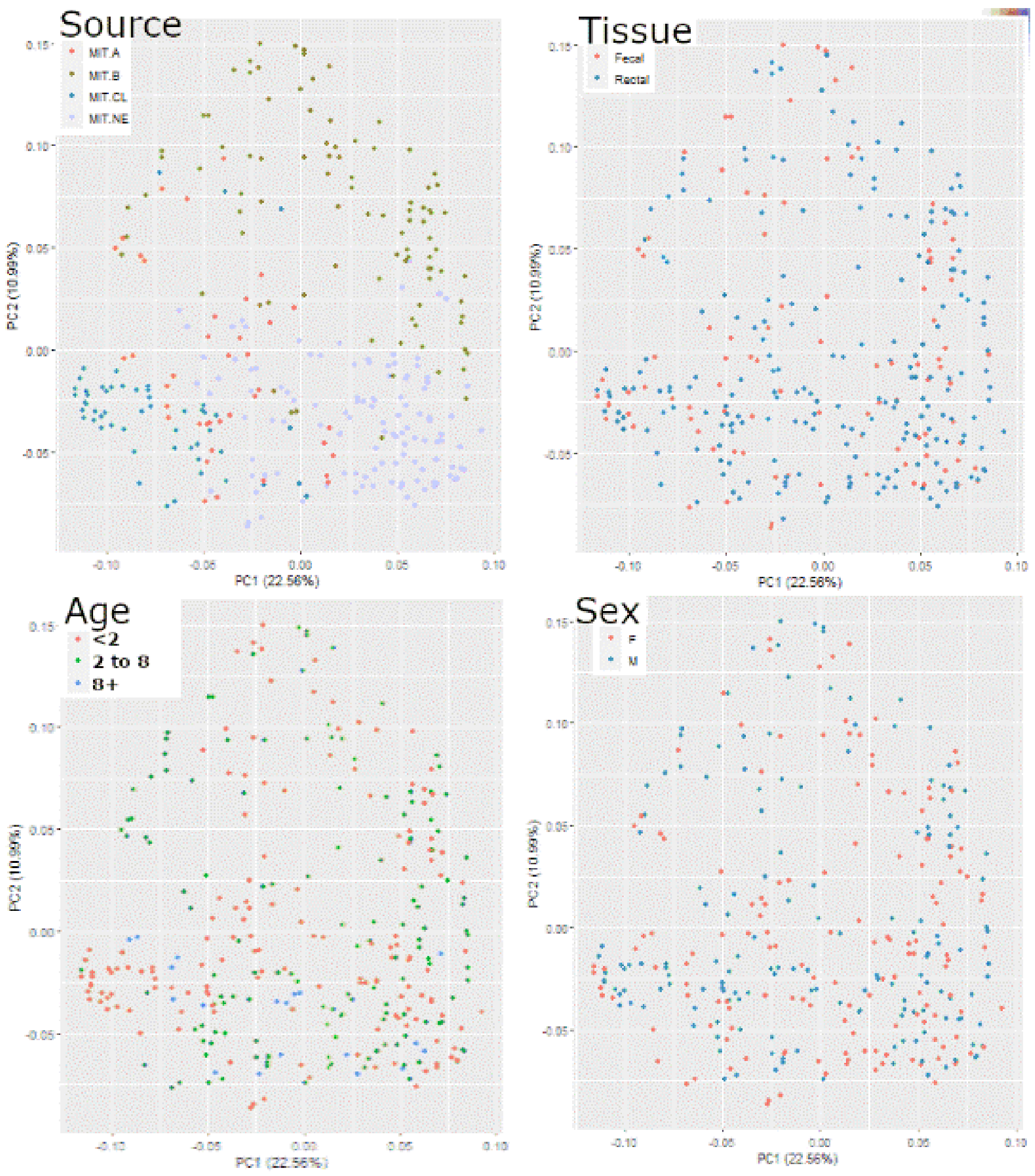


Supplemental Figure 2. PCA plots of healthy marmosets categorized by source, sample type (tissue), age and sex. Classification by source shows clustering, while other classifications show poor separation.

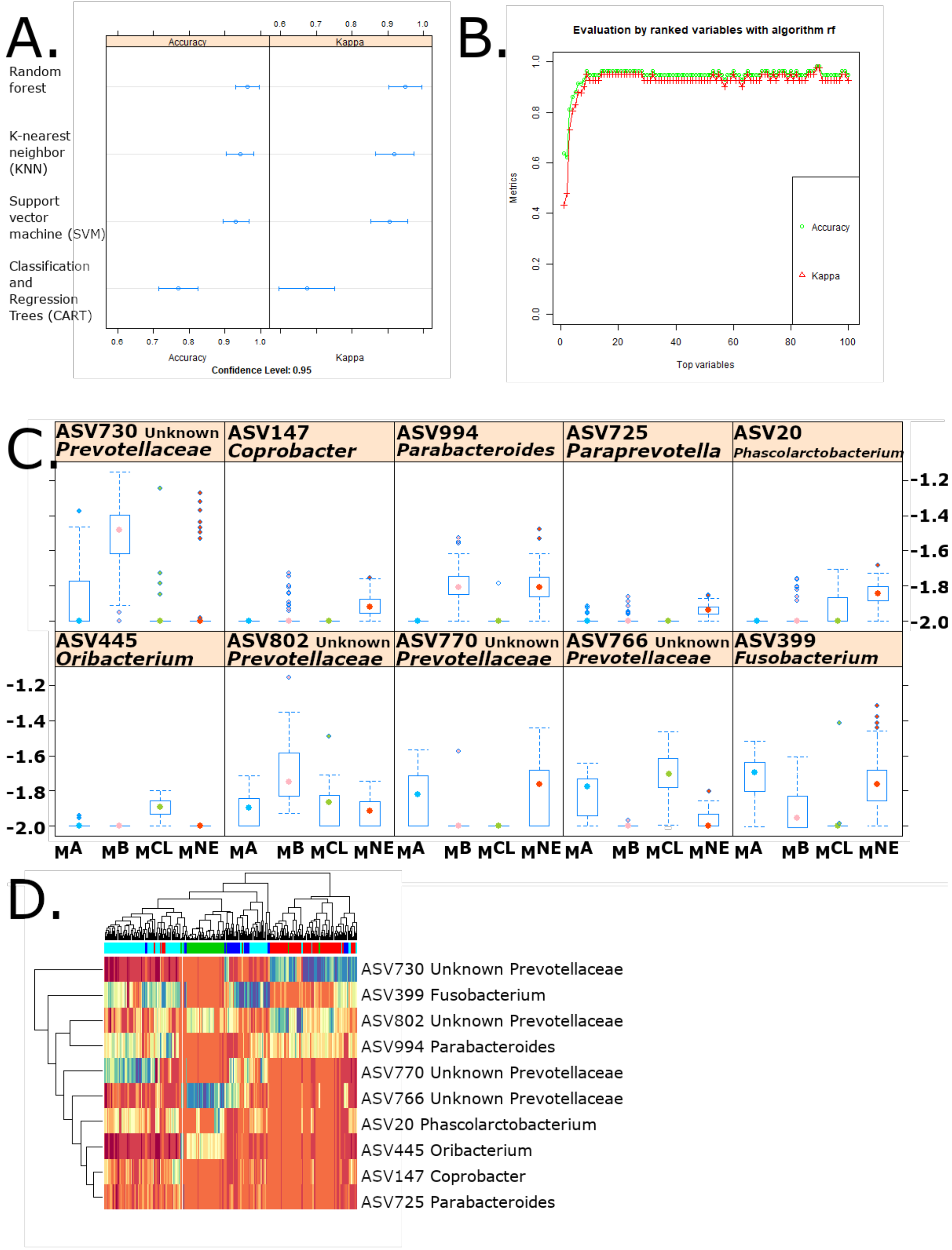


Supplemental Figure 3. A) Comparison of classifier models used to classify healthy microbiomes based on source included random forest (RF), K-nearest neighbor, support vector machines, and classification and regression trees (CART). RF consistently outperformed the other classifiers. B) Accuracy of RF model stabilizes with 10 variables. C) Boxplots of 10 ASVs selected by RF model show source-specific differences. D) Heatmap of ASV abundances showing classification of training data using 9 ASVs. Color bar on top indicates true identity.

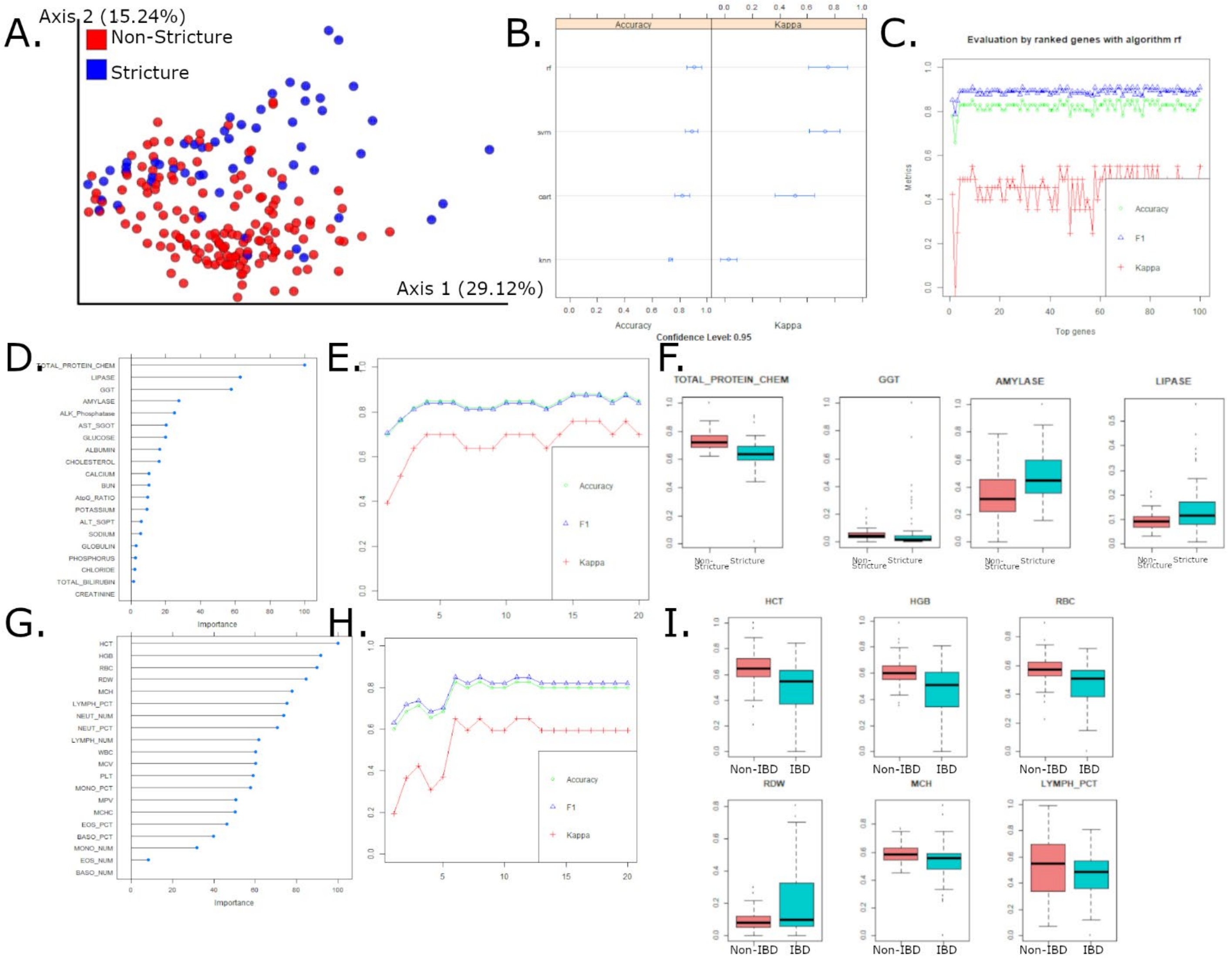


Supplemental Figure 4. A) PCA plot of stricture progressors and non-progressors shows marked differences in bacterial communities B) Accuracy comparison of RF, SVM, CART and KNN models show RF outperforms other models using microbiome data to classify strictures. C) Accuracy stabilizes for microbiome stricture RF model with 9 ASVs. D) Variables of importance for RF model using serum chemistry data to classify strictures. E) 4 serum chemistry parameters are required for optimal accuracy in serum chemistry RF model. F) 4 most important parameters include Total Protein, GGT, amylase and lipase. Amylase and lipase are elevated in stricture cases suggesting pancreatic disease. G) Variables of importance for RF model using complete blood counts (CBC) to classify strictures. H) 6 CBC parameters required for optimal accuracy. I) 6 CBC parameters include HCT, HGB, RBC, RDW, MCH and lymphocyte %. 5 of 6 parameters indicate anemia.

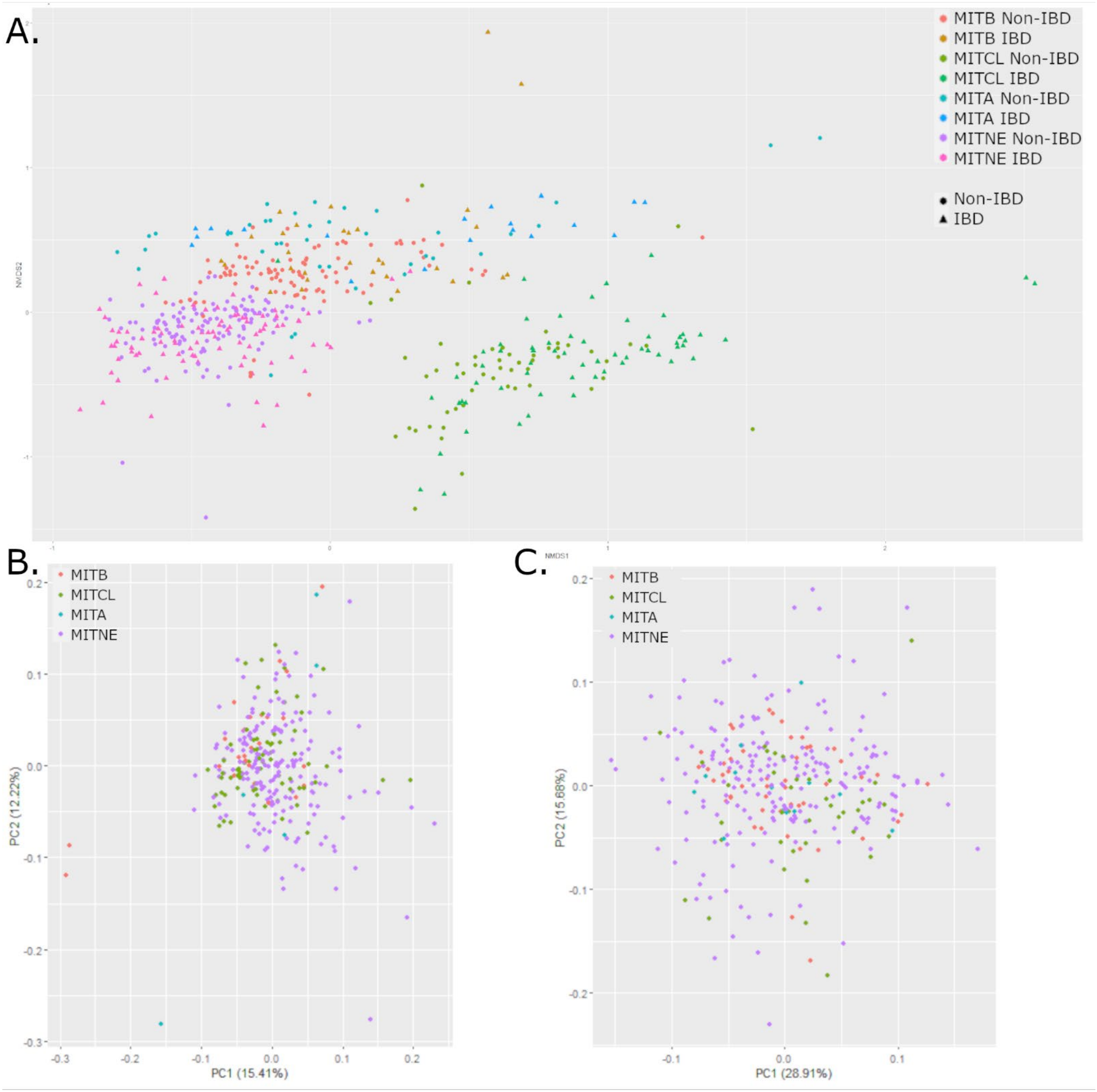


Supplemental Figure 5. A) PCA plots of microbiome data classified by source and IBD status shows slight PC1 shifts due to IBD within each source population. B) No separation observed in serum chemistry PCA plot by source. C) No separation in CBC PCA plot by source.

### Supp. Fig 6

#### A. Network of Biological Processes enriched in the duodenum of stricture cases

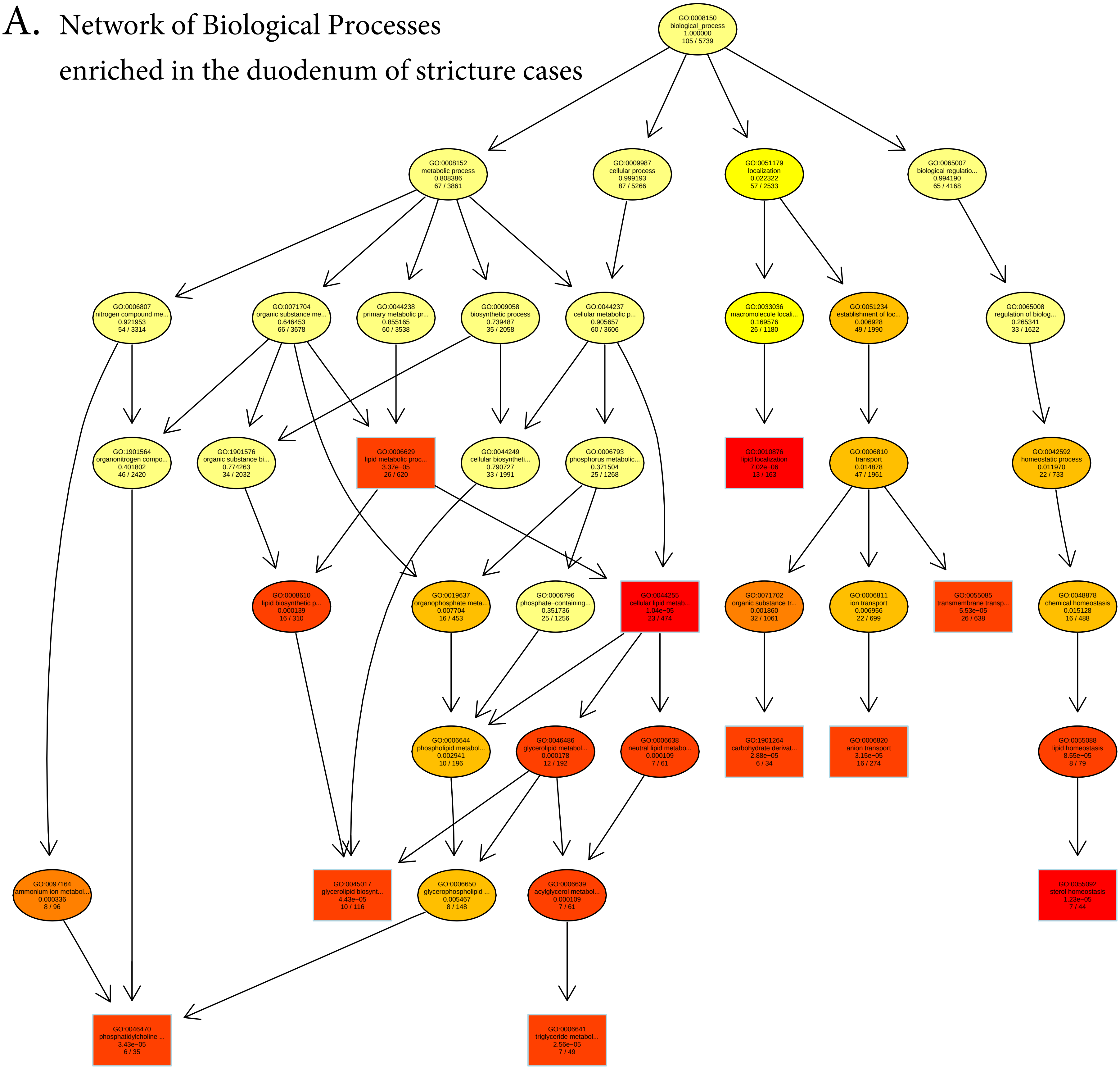

**B.** Network of Cellular Components enriched in the duodenum of stricture cases

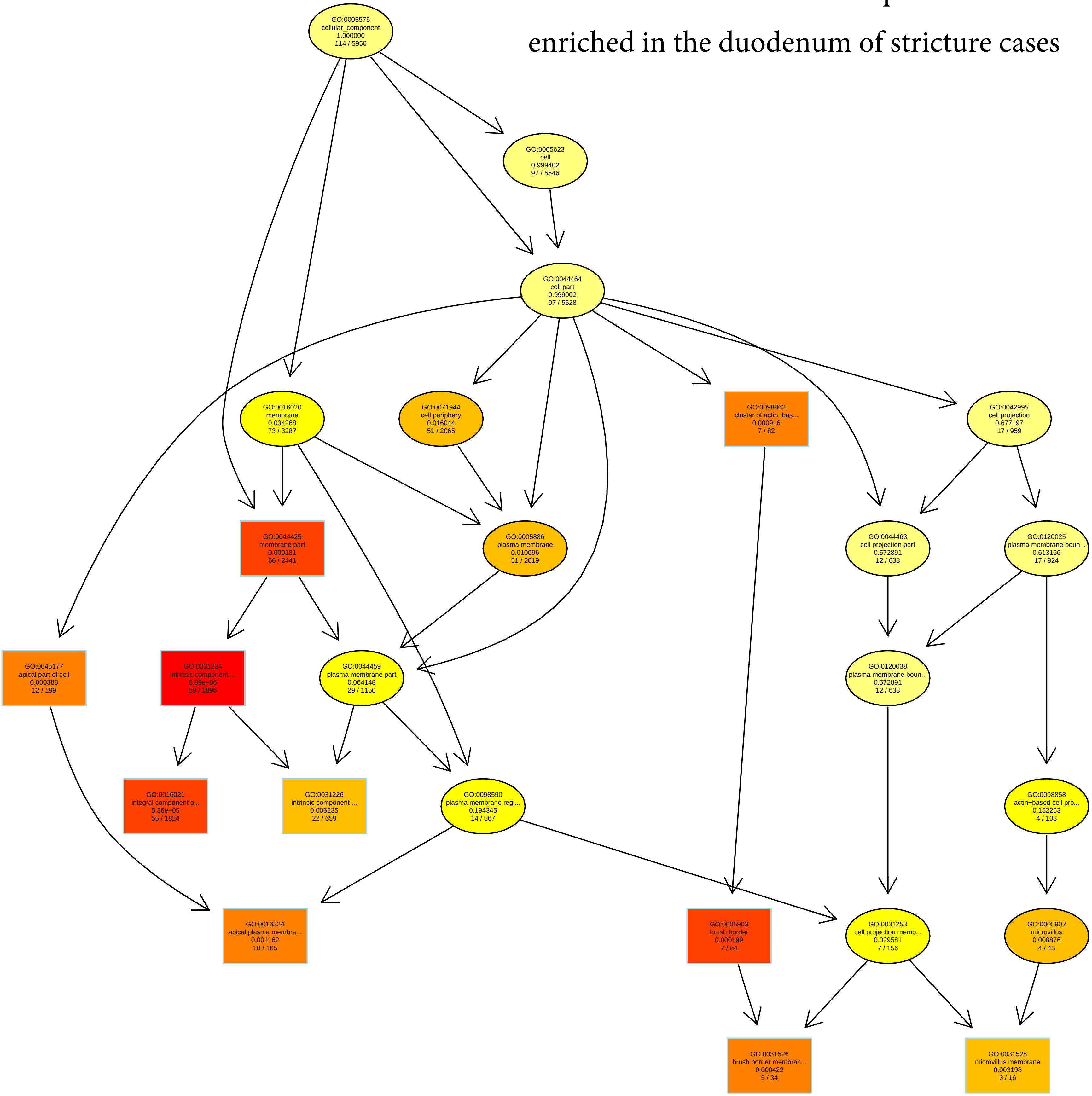

**Supp. Fig. 7.** Network of Biological Processes enriched in the duodenum of non-stricture cases

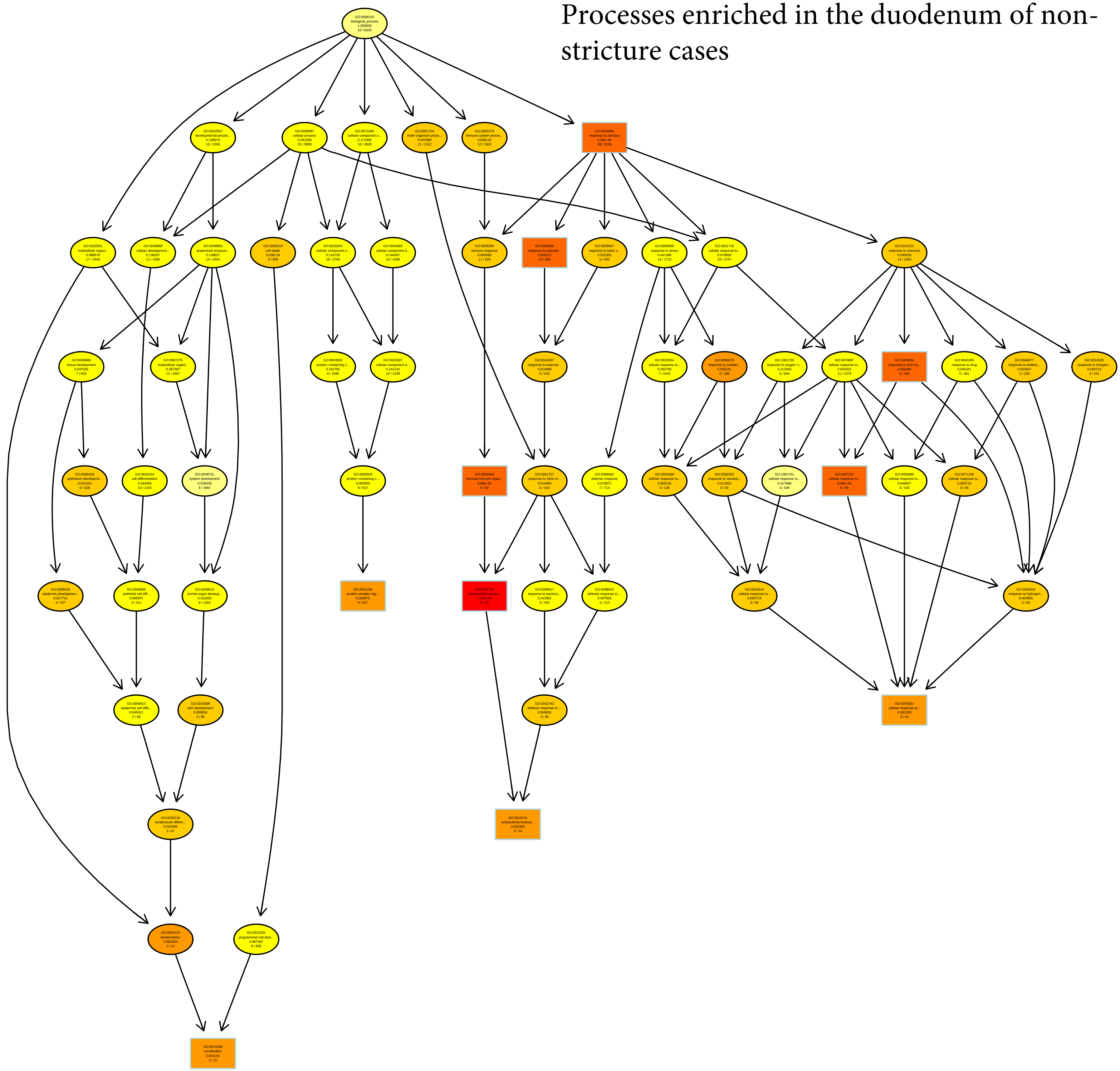

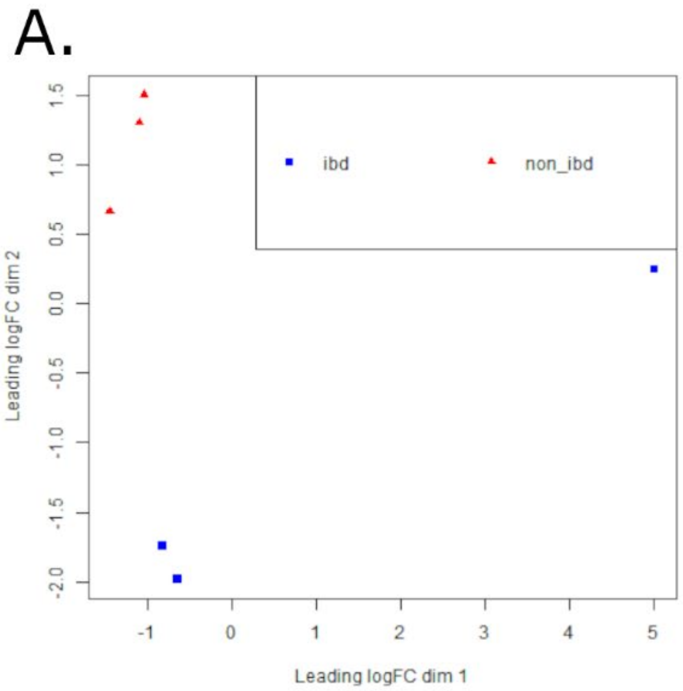


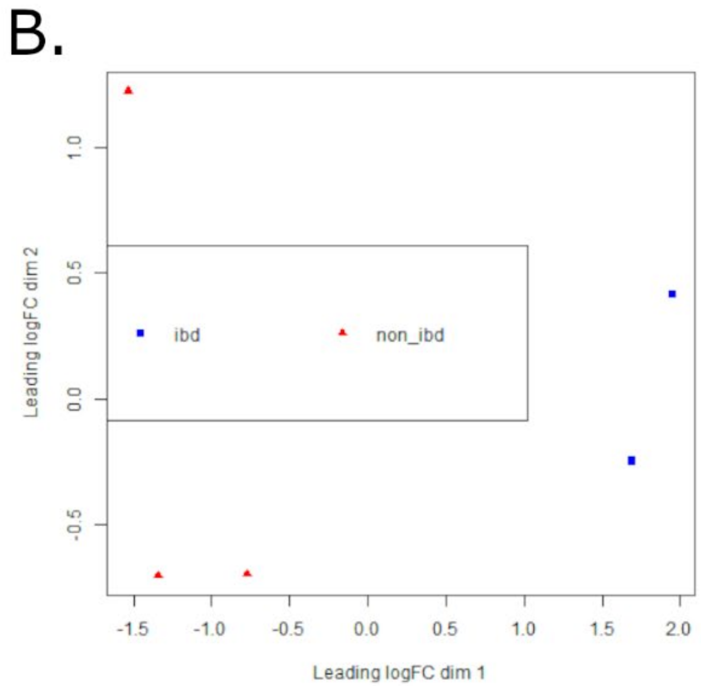


Supp Fig 8. A) PCA plot of RNAseq data for non-IBD and IBD samples from the jejunum show a single sample that differs from other 5 samples. B) PCA plot after removal of outlier.

Supp. Fig 9 Biological Processes enriched in the jejunum of IBD cases

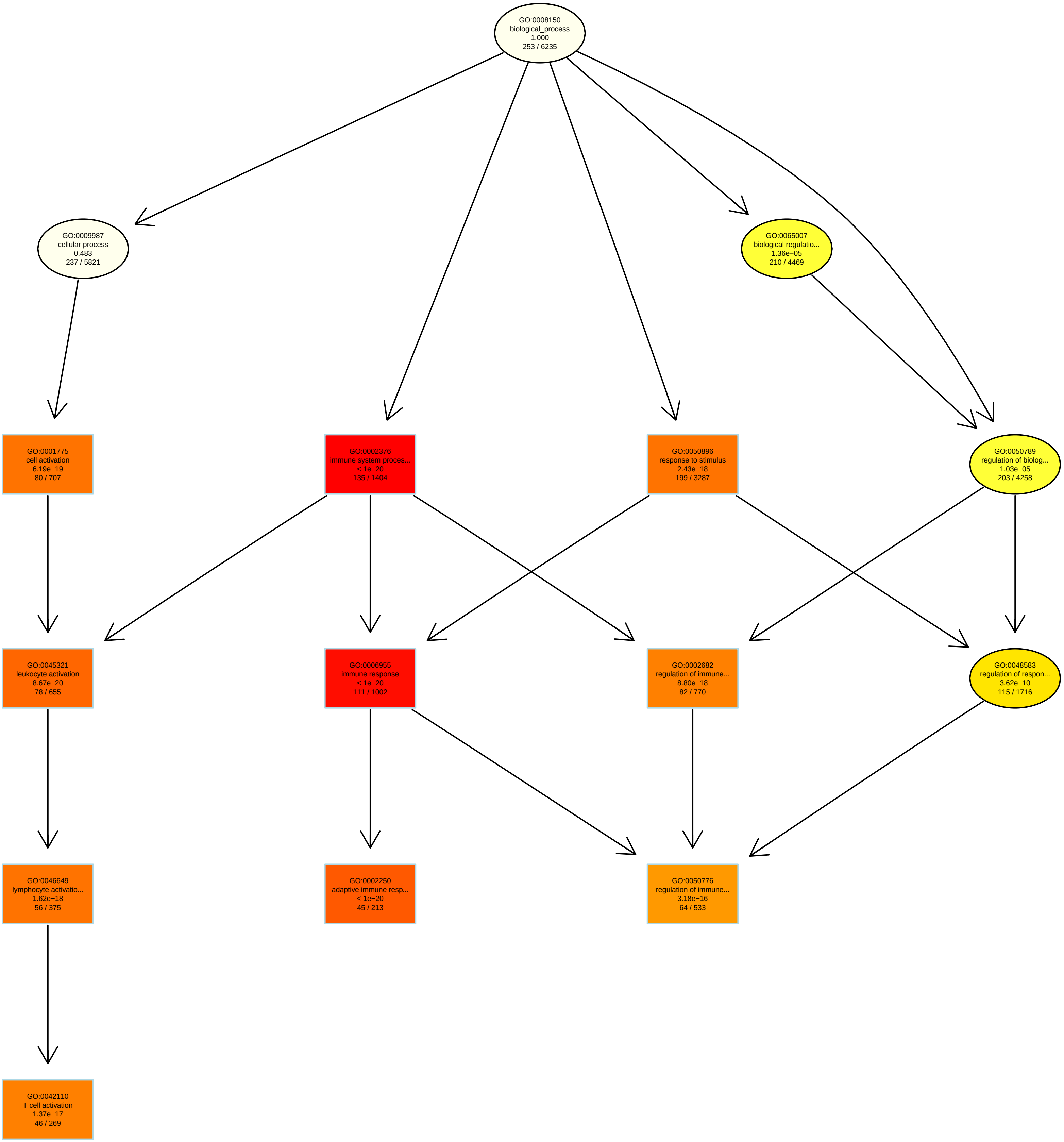

Supp. Fig. 10 Biological processes enriched in the jejunum of non-IBD cases

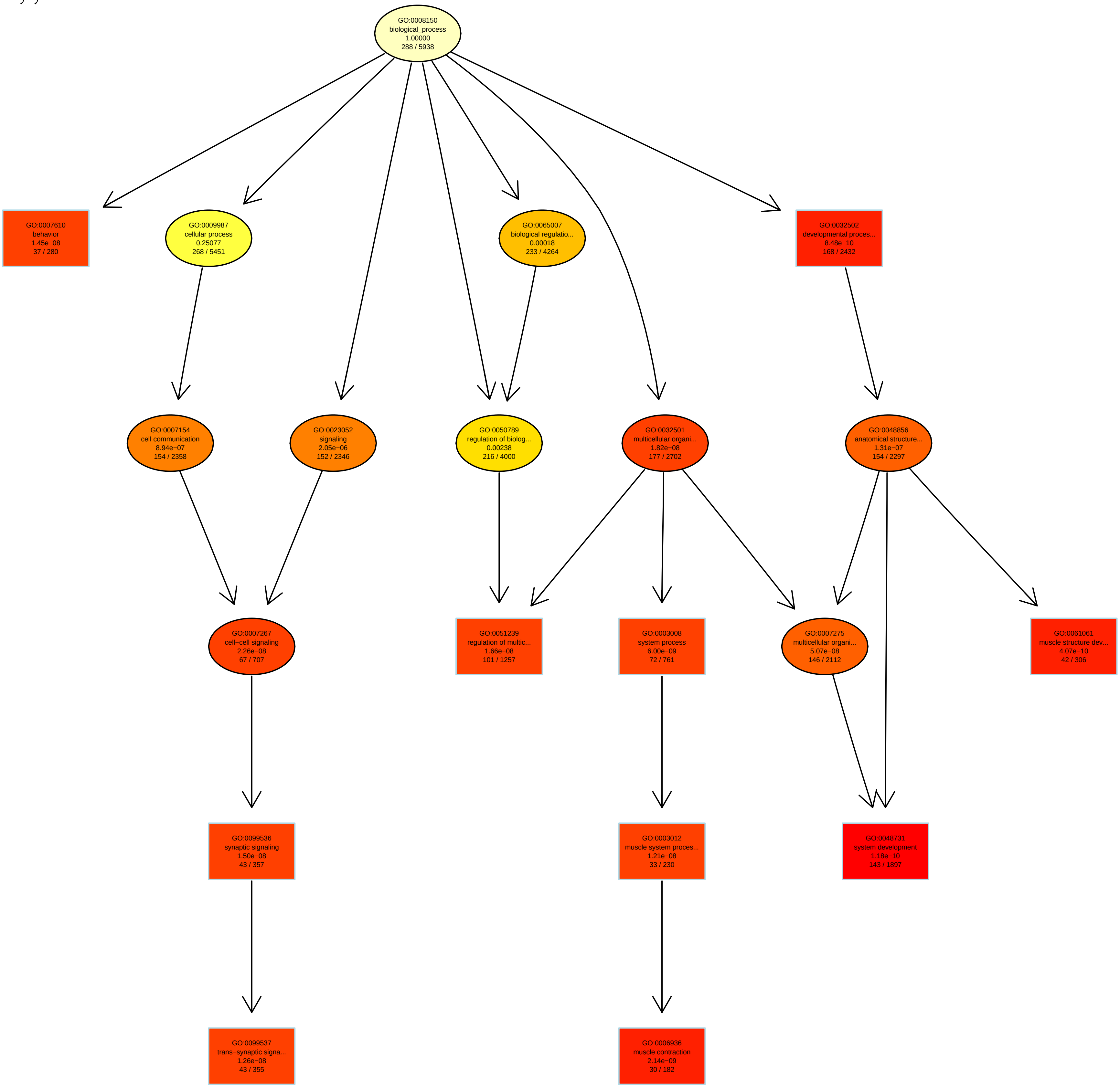
