## Supplemental Tables for "Common Marmoset Gut Microbiome Profiles in Health and Intestinal Disease"

**Supplemental Table 1**

| Phylum | % | Family | % | Genus | % |
|---|---|---|---|---|---|
| *Bacteroidetes* | 63.20% | *Bacteroidaceae* | 28.40% | *Bacteroides* | 28.40% |
| | | *Prevotellaceae* | 25.80% | *Prevotella 9* | 15.50% |
| | | | | *Unknown Prevotellaceae* | 8.10% |
| | | *Tannerellaceae* | 7.50% | *Parabacteroides* | 7.50% |
| *Firmicutes* | 16.70% | *Veillonellaceae* | 9.40% | *Megamonas* | 7.20% |
| | | | | *Megasphaera* | 2.20% |
| | | *Acidaminococcaceae* | 3.80% | *Phascolarctobacterium* | 3.80% |
| | | *Lachnospiraceae* | 1.90% | *Blautia* | 1.00% |
| *Proteobacteria* | 13.80% | *Succinivibrionaceae* | 9.70% | *Anaerobiospirillum* | 9.60% |
| | | *Burkholderiaceae* | 1.90% | *Sutterella* | 1.90% |
| | | *Enterobacteriaceae* | 1.80% | *Escherichia-Shigella* | 1.70% |
| *Fusobacteria* | 4.00% | *Fusobacteriaceae* | 4.00% | *Fusobacterium* | 4.00% |
| *Actinobacteria* | 1.40% | *Bifidobacteriaceae* | 1.00% | *Bifidobacterium* | 1.00% |
| *Epsilonbacteraeota* | 0.70% | | | | |
| *Verrucomicrobia* | 0.10% | | | | |
| *Spirochaetes* | 0.00% | | | | |
| *Cyanobacteria* | 0.00% | | | | |
| *Deferribacteres* | 0.00% | | | | |

Supplemental Table 2. ANCOM analysis by source in healthy marmosets

| Phylum | Class | Order | Family | Genus | W | $\text{MIT}^{NE}$ | $\text{MIT}^{CL}$ | $\text{MIT}^{B}$ | $\text{MIT}^{A}$ | MIT Abundance |
|---|---|---|---|---|---|---|---|---|---|---|
| Bacteroidetes | Bacteroidia | Bacteroidales | Prevotellaceae | Prevotella 9 | 136 | 10.64% | 33.40% | 9.44% | 25.17% | 15.40% |
| Bacteroidetes | Bacteroidia | Bacteroidales | Prevotellaceae | Alloprevotella | 136 | 0.00% | 2.51% | 0.00% | 0.00% | 0.38% |
| Fusobacteria | Fusobacteriia | Fusobacteriales | Fusobacteriaceae | Fusobacterium | 136 | 5.01% | 3.96% | 1.63% | 8.24% | 4.13% |
| Proteobacteria | Gammaproteobacteria | Aeromonadales | Succinivibrionaceae | Anaerobiospirillum | 136 | 13.77% | 1.47% | 8.50% | 10.17% | 9.77% |
| Proteobacteria | Gammaproteobacteria | Enterobacteriales | Enterobacteriaceae | Escherichia-Shigella | 136 | 0.13% | 3.16% | 0.61% | 4.92% | 1.30% |
| Firmicutes | Clostridia | Clostridiales | Lachnospiraceae | Blautia | 133 | 0.80% | 0.65% | 1.68% | 0.43% | 1.02% |
| Bacteroidetes | Bacteroidia | Bacteroidales | Prevotellaceae | Unknown | 131 | 4.78% | 6.37% | 13.83% | 5.84% | 8.07% |
| Actinobacteria | Actinobacteria | Bifidobacteriales | Bifidobacteriaceae | Bifidobacterium | 130 | 0.53% | 0.96% | 1.42% | 1.19% | 0.96% |
| Bacteroidetes | Bacteroidia | Bacteroidales | Tannerellaceae | Parabacteroides | 126 | 6.06% | 4.71% | 9.61% | 8.62% | 7.30% |
| Bacteroidetes | Bacteroidia | Bacteroidales | Bacteroidaceae | Bacteroides | 126 | 37.37% | 17.44% | 29.94% | 14.39% | 29.27% |
| Firmicutes | Negativicutes | Selenomonadales | Acidaminococcaceae | Phascolarctobacterium | 125 | 4.27% | 2.76% | 3.85% | 2.78% | 3.73% |
| Firmicutes | Negativicutes | Selenomonadales | Veillonellaceae | Megamonas | 125 | 6.21% | 9.37% | 8.22% | 5.62% | 7.28% |
| Firmicutes | Negativicutes | Selenomonadales | Veillonellaceae | Megasphaera | 116 | 1.04% | 2.69% | 2.96% | 3.96% | 2.25% |
| Bacteroidetes | Bacteroidia | Bacteroidales | Barnesiellaceae | Coprobacter | 136 | 0.27% | 0.00% | 0.08% | 0.00% | 0.14% |
| Bacteroidetes | Bacteroidia | Bacteroidales | Muribaculaceae | __ | 136 | 0.15% | 0.42% | 0.13% | 0.05% | 0.17% |
| Bacteroidetes | Bacteroidia | Bacteroidales | Muribaculaceae | uncultured bacterium | 136 | 0.38% | 0.08% | 0.82% | 0.36% | 0.48% |
| Bacteroidetes | Bacteroidia | Bacteroidales | Rikenellaceae | Alistipes | 136 | 0.18% | 0.08% | 0.08% | 0.02% | 0.11% |
| Firmicutes | Clostridia | Clostridiales | Lachnospiraceae | __ | 136 | 0.06% | 0.01% | 0.11% | 0.05% | 0.07% |
| Firmicutes | Clostridia | Clostridiales | Lachnospiraceae | Lachnoclostridium | 136 | 0.08% | 0.00% | 0.22% | 0.18% | 0.13% |
| Firmicutes | Clostridia | Clostridiales | Ruminococcaceae | Ruminiclostridium 5 | 136 | 0.04% | 0.02% | 0.01% | 0.00% | 0.02% |
| Firmicutes | Erysipelotrichia | Erysipelotrichales | Erysipelotrichaceae | Asteroleplasma | 136 | 0.20% | 0.11% | 0.70% | 0.40% | 0.37% |
| Proteobacteria | Deltaproteobacteria | Desulfovibrionales | Desulfovibrionaceae | Bilophila | 136 | 0.01% | 0.13% | 0.15% | 0.16% | 0.09% |
| Proteobacteria | Gammaproteobacteria | __ | __ | __ | 136 | 0.00% | 0.00% | 0.00% | 0.14% | 0.02% |
| Proteobacteria | Gammaproteobacteria | Aeromonadales | Succinivibrionaceae | Succinatimonas | 136 | 0.02% | 0.13% | 0.05% | 0.01% | 0.05% |
| Bacteroidetes | Bacteroidia | Bacteroidales | Barnesiellaceae | Barnesiella | 135 | 0.63% | 0.28% | 0.06% | 0.14% | 0.33% |
| Bacteroidetes | Bacteroidia | Bacteroidales | Marinifilaceae | Butyricimonas | 135 | 0.28% | 0.07% | 0.02% | 0.03% | 0.14% |
| Bacteroidetes | Bacteroidia | Bacteroidales | Marinifilaceae | Odoribacter | 134 | 0.12% | 0.00% | 0.01% | 0.00% | 0.05% |
| Firmicutes | Clostridia | Clostridiales | Lachnospiraceae | [Ruminococcus] gnavus group | 134 | 0.05% | 0.21% | 0.00% | 0.05% | 0.06% |
| Firmicutes | Clostridia | Clostridiales | Lachnospiraceae | Howardella | 134 | 0.00% | 0.00% | 0.03% | 0.01% | 0.01% |
| Firmicutes | Clostridia | Clostridiales | Lachnospiraceae | uncultured | 134 | 0.04% | 0.12% | 0.19% | 0.05% | 0.10% |
| Proteobacteria | Deltaproteobacteria | Desulfovibrionales | Desulfovibrionaceae | Desulfovibrio | 134 | 0.12% | 0.00% | 0.01% | 0.00% | 0.05% |
| Actinobacteria | Coriobacteriia | Coriobacteriales | Atopobiaceae | Olsenella | 133 | 0.11% | 0.05% | 0.15% | 0.03% | 0.10% |
| Firmicutes | Clostridia | Clostridiales | Lachnospiraceae | Lachnospiraceae UCG-004 | 133 | 0.00% | 0.05% | 0.00% | 0.00% | 0.01% |
| Proteobacteria | Gammaproteobacteria | Enterobacteriales | Enterobacteriaceae | Proteus | 133 | 0.01% | 0.00% | 0.00% | 0.08% | 0.01% |
| Firmicutes | Clostridia | Clostridiales | Peptococcaceae | Peptococcus | 132 | 0.08% | 0.01% | 0.01% | 0.01% | 0.04% |
| Firmicutes | Negativicutes | Selenomonadales | Veillonellaceae | Allisonella | 132 | 0.00% | 0.04% | 0.01% | 0.02% | 0.01% |
| Firmicutes | Negativicutes | Selenomonadales | Veillonellaceae | Dialister | 132 | 0.00% | 0.22% | 0.01% | 0.00% | 0.04% |
| Bacteroidetes | Bacteroidia | Bacteroidales | Prevotellaceae | Paraprevotella | 131 | 0.87% | 0.47% | 0.79% | 0.65% | 0.76% |
| Actinobacteria | Coriobacteriia | Coriobacteriales | Atopobiaceae | Libanicoccus | 130 | 0.06% | 0.02% | 0.10% | 0.03% | 0.06% |
| Firmicutes | Clostridia | Clostridiales | Lachnospiraceae | Oribacterium | 128 | 0.39% | 0.81% | 0.62% | 0.57% | 0.55% |
| Spirochaetes | Spirochaetia | Spirochaetales | Spirochaetaceae | Sphaerochaeta | 127 | 0.05% | 0.00% | 0.00% | 0.00% | 0.02% |
| Bacteroidetes | Bacteroidia | Bacteroidales | Porphyromonadaceae | Porphyromonas | 126 | 0.00% | 0.02% | 0.01% | 0.00% | 0.01% |
| Firmicutes | Clostridia | Clostridiales | Lachnospiraceae | GCA-900066575 | 126 | 0.04% | 0.00% | 0.03% | 0.00% | 0.02% |
| Firmicutes | Clostridia | Clostridiales | Ruminococcaceae | Ruminococcaceae UCG-014 | 125 | 0.09% | 0.00% | 0.00% | 0.00% | 0.04% |
| Firmicutes | Clostridia | Clostridiales | Lachnospiraceae | Lachnospiraceae NK4A136 group | 124 | 0.04% | 0.00% | 0.00% | 0.00% | 0.02% |
| Proteobacteria | Alphaproteobacteria | Rhodospirillales | uncultured | __ | 124 | 0.09% | 0.00% | 0.02% | 0.00% | 0.04% |
| Firmicutes | Clostridia | Clostridiales | Ruminococcaceae | Fournierella | 123 | 0.00% | 0.01% | 0.00% | 0.00% | 0.00% |
| Firmicutes | Clostridia | Clostridiales | Clostridiaceae 1 | Sarcina | 122 | 0.06% | 0.03% | 0.04% | 0.00% | 0.04% |
| Firmicutes | Clostridia | Clostridiales | Ruminococcaceae | __ | 122 | 0.01% | 0.00% | 0.00% | 0.00% | 0.01% |
| Bacteroidetes | Bacteroidia | Bacteroidales | Barnesiellaceae | uncultured | 121 | 0.03% | 0.00% | 0.02% | 0.00% | 0.02% |
| Firmicutes | Clostridia | Clostridiales | Ruminococcaceae | Ruminococcaceae UCG-004 | 121 | 0.02% | 0.00% | 0.00% | 0.00% | 0.01% |
| Firmicutes | Bacilli | Lactobacillales | Aerococcaceae | Aerococcus | 120 | 0.00% | 0.00% | 0.00% | 0.41% | 0.05% |
| Firmicutes | Clostridia | Clostridiales | Ruminococcaceae | Ruminiclostridium 9 | 120 | 0.05% | 0.01% | 0.02% | 0.00% | 0.03% |
| Proteobacteria | Gammaproteobacteria | Enterobacteriales | Enterobacteriaceae | __ | 120 | 0.00% | 0.00% | 0.00% | 0.08% | 0.01% |
| Actinobacteria | Coriobacteriia | Coriobacteriales | Coriobacteriaceae | Collinsella | 119 | 0.24% | 0.18% | 0.34% | 0.09% | 0.25% |
| Verrucomicrobia | Verrucomicrobiae | Verrucomicrobiales | Akkermansiaceae | Akkermansia | 112 | 0.25% | 0.05% | 0.00% | 0.00% | 0.11% |
| Firmicutes | Bacilli | Lactobacillales | Leuconostocaceae | Leuconostoc | 110 | 0.00% | 0.06% | 0.00% | 0.00% | 0.01% |
| Firmicutes | Clostridia | Clostridiales | Peptostreptococcaceae | Peptoclostridium | 109 | 0.00% | 0.05% | 0.00% | 0.00% | 0.01% |
| Firmicutes | Clostridia | Clostridiales | Ruminococcaceae | Oscillibacter | 107 | 0.01% | 0.01% | 0.00% | 0.00% | 0.01% |
| Epsilonbacteraeota | Campylobacteria | Campylobacterales | Helicobacteraceae | Helicobacter | 104 | 1.00% | 0.20% | 0.63% | 0.65% | 0.72% |
| Firmicutes | Clostridia | Clostridiales | Family XIII | Family XIII UCG-001 | 103 | 0.00% | 0.00% | 0.00% | 0.00% | 0.00% |
| Cyanobacteria | Melainabacteria | Gastranaerophilales | __ | __ | 102 | 0.01% | 0.00% | 0.00% | 0.00% | 0.00% |
| Firmicutes | Clostridia | Clostridiales | Ruminococcaceae | Subdoligranulum | 102 | 0.00% | 0.00% | 0.00% | 0.00% | 0.00% |

Supplemental Table 3. Number of marmosets and associated health status for animals on study

| | Healthy | Stricture | IBD |
|---|---|---|---|
| $MIT^{A}$ | 8 (73%) | 0 (0%) | 3 (27%) |
| $MIT^{B}$ | 27 (75%) | 1 (3%) | 8 (22%) |
| $MIT^{CL}$ | 19 (43%) | 1 (2%) | 24 (55%) |
| $MIT^{NE}$ | 37 (45%) | 21 (26%) | 24 (29%) |
| $MIT^{ALL}$ | 91 (53%) | 23 (13%) | 59 (34%) |

Supplemental Table 4. Overlap of top 25 ASVs in 4 IBD models

**ASVs of importance in 3+ models**

| | Family | Genus |
|---|---|---|
| ASV219 | *Burkholderiaceae* | *Sutterella* |
| ASV115 | *Bacteroidaceae* | *Bacteroides* |
| ASV207 | *Burkholderiaceae* | *Sutterella* |
| ASV209 | *Burkholderiaceae* | *Sutterella* |
| ASV346 | *Erysipelotrichaceae* | *Asteroleplasma* |
| ASV779 | *Prevotellaceae* | *Prevotella 9* |
| ASV1044 | *Veillonellaceae* | *Megamonas* |
| ASV1087 | *Veillonellaceae* | *Megamonas* |

**Genera of importance in 3+ models with number of ASVs per model**

| Genus | MIT (all) | MIT B | MIT CL | MIT NE | Total |
|---|---|---|---|---|---|
| *Bacteroides* | 8 | 4 | 4 | 4 | 20 |
| *Sutterella* | 3 | 3 | 1 | 3 | 10 |
| *Megamonas* | 2 | 3 | 3 | 0 | 8 |
| *Bifidobacterium* | 1 | 4 | 2 | 0 | 7 |
| *Prevotella 9* | 2 | 0 | 2 | 3 | 7 |
| *Unknown Prevotellaceae* | 0 | 2 | 2 | 1 | 5 |
| *Parabacteroides* | 1 | 1 | 0 | 3 | 5 |
| *Prevotellaceae UCG-001* | 2 | 0 | 1 | 1 | 4 |
| *Asteroleplasma* | 1 | 1 | 1 | 0 | 3 |

Supplemental Table 5. Differentially expressed genes in the duodenum between stricture and non-stricture

**Supp. Table 5a. Genes upregulated in Stricture**

| Gene name | logFC | logCPM | F | PValue | FDR |
|---|---|---|---|---|---|
| HYDIN | -5.57275 | 8.235763 | 255.7079 | 9.64E-09 | 0.000186 |
| FCN2 | -6.2382 | 5.877223 | 181.0315 | 9.24E-08 | 0.000677 |
| LOC100385035 | -4.39198 | 4.35918 | 159.023 | 1.06E-07 | 0.000677 |
| KIFC3 | -3.72529 | 6.16206 | 147.199 | 1.55E-07 | 0.000745 |
| FRMD1 | -4.56494 | 4.611455 | 129.1226 | 2.96E-07 | 0.001139 |
| GOLT1A | -3.66778 | 5.210277 | 124.2568 | 3.57E-07 | 0.001146 |
| LOC100415170 | -4.60077 | 4.395378 | 112.2745 | 6.08E-07 | 0.001456 |
| LOC100390425 | -4.07882 | 7.969966 | 112.2804 | 6.21E-07 | 0.001456 |
| SLC2A7 | -2.82598 | 6.173383 | 104.4619 | 8.31E-07 | 0.0016 |
| TUBAL3 | -3.41415 | 6.772312 | 100.0865 | 1.02E-06 | 0.001788 |
| LOC103792726 | -3.50238 | 4.766558 | 92.43918 | 1.50E-06 | 0.001872 |
| AQP10 | -3.17156 | 6.834658 | 90.41122 | 1.66E-06 | 0.001872 |
| ESPL1 | -2.35296 | 7.029238 | 90.60278 | 1.65E-06 | 0.001872 |
| CHAD | -2.32012 | 6.502384 | 94.50606 | 1.35E-06 | 0.001872 |
| LOC108590320 | -3.69941 | 3.094203 | 81.63781 | 2.70E-06 | 0.001911 |
| PRAP1 | -3.50931 | 9.558516 | 86.09876 | 2.10E-06 | 0.001911 |
| EPDR1 | -3.28843 | 5.553597 | 85.40134 | 2.18E-06 | 0.001911 |
| TREH | -3.22115 | 8.869108 | 81.9639 | 2.65E-06 | 0.001911 |
| GLIS1 | -3.21171 | 4.448253 | 82.11207 | 2.63E-06 | 0.001911 |
| LOC103793334 | -2.94838 | 3.755427 | 81.14824 | 2.78E-06 | 0.001911 |
| LOC100385629 | -2.83254 | 7.023845 | 87.61152 | 1.93E-06 | 0.001911 |
| PDZD3 | -2.65957 | 6.514583 | 84.4437 | 2.30E-06 | 0.001911 |
| FER1L5 | -2.62937 | 6.461021 | 82.45264 | 2.58E-06 | 0.001911 |
| ADAMTSL5 | -2.32562 | 5.818443 | 85.2145 | 2.21E-06 | 0.001911 |
| ZNF385C | -3.46513 | 4.47279 | 79.46398 | 3.07E-06 | 0.001969 |
| MROH7 | -2.81291 | 7.598492 | 79.63771 | 3.04E-06 | 0.001969 |
| TGM1 | -3.90035 | 4.350572 | 78.98298 | 3.53E-06 | 0.001992 |
| LOC100415600 | -3.79525 | 7.829823 | 83.86128 | 3.29E-06 | 0.001992 |
| LOC100397175 | -3.35994 | 4.550014 | 77.18741 | 3.52E-06 | 0.001992 |
| MTHFR | -2.51542 | 8.221851 | 76.71136 | 3.62E-06 | 0.001992 |
| LOC100390661 | -2.52158 | 6.35154 | 75.78073 | 3.84E-06 | 0.002051 |
| NPB | -3.5985 | 3.559672 | 74.93623 | 4.04E-06 | 0.002103 |
| RTN4R | -2.67811 | 3.950289 | 73.36671 | 4.46E-06 | 0.002138 |
| PMM1 | -2.12833 | 6.086473 | 73.04871 | 4.55E-06 | 0.002138 |
| OXCT2 | -2.78732 | 3.062813 | 71.85792 | 4.92E-06 | 0.002254 |
| LOC100411074 | -4.05337 | 4.176173 | 72.85729 | 5.72E-06 | 0.002446 |
| LOC103790125 | -2.7728 | 4.293793 | 69.68456 | 5.67E-06 | 0.002446 |
| MFSD2A | -2.62376 | 7.406906 | 70.24467 | 5.46E-06 | 0.002446 |
| LOC103793410 | -5.22733 | 2.764468 | 70.13029 | 5.88E-06 | 0.002462 |
| AMN | -2.92841 | 8.875209 | 68.79194 | 6.02E-06 | 0.002466 |
| LOC103791695 | -3.79597 | 3.480268 | 68.11951 | 6.30E-06 | 0.002476 |

| | | | | | |
|---|---|---|---|---|---|
| APOA4 | -3.59877 | 13.91962 | 68.30061 | 6.22E-06 | 0.002476 |
| DNAL1 | -3.16518 | 5.44604 | 69.49632 | 6.46E-06 | 0.002488 |
| FAM131C | -3.41453 | 3.765668 | 66.74177 | 6.92E-06 | 0.002614 |
| RNF157 | -2.61665 | 5.063351 | 65.82983 | 7.38E-06 | 0.002732 |
| LOC100388936 | -1.85957 | 7.044496 | 65.39543 | 7.61E-06 | 0.002764 |
| LOC100406930 | -2.9716 | 3.483737 | 63.3751 | 8.79E-06 | 0.00303 |
| LOC100389703 | -2.8047 | 6.746623 | 64.03561 | 8.81E-06 | 0.00303 |
| KCNK10 | -2.61107 | 3.951578 | 63.53738 | 8.68E-06 | 0.00303 |
| APOA1 | -3.14159 | 13.41736 | 62.25827 | 9.53E-06 | 0.00322 |
| LOC103793414 | -7.86258 | 1.574829 | 84.48291 | 1.01E-05 | 0.003345 |
| DYRK1B | -2.17365 | 6.64759 | 60.13611 | 1.12E-05 | 0.003584 |
| ACACB | -2.48726 | 7.814934 | 58.92525 | 1.23E-05 | 0.003796 |
| PROCA1 | -2.3523 | 4.032119 | 58.70854 | 1.25E-05 | 0.003796 |
| CCDC120 | -2.15183 | 6.633082 | 58.54436 | 1.26E-05 | 0.003796 |
| B4GALNT1 | -2.17693 | 7.93829 | 57.18402 | 1.40E-05 | 0.004094 |
| SLC7A8 | -2.04912 | 7.679964 | 57.26598 | 1.39E-05 | 0.004094 |
| AZGP1 | -5.22926 | 10.8214 | 66.51713 | 1.49E-05 | 0.004109 |
| SLC6A19 | -3.333 | 9.621099 | 57.4679 | 1.55E-05 | 0.004109 |
| BMP8A | -3.07527 | 5.365014 | 59.06857 | 1.51E-05 | 0.004109 |
| RASD2 | -2.30582 | 5.129872 | 56.532 | 1.48E-05 | 0.004109 |
| LOC108591342 | -4.20888 | 3.606382 | 59.17617 | 1.63E-05 | 0.004155 |
| XPNPEP2 | -2.4449 | 8.070324 | 55.24302 | 1.64E-05 | 0.004155 |
| CLCN6 | -1.83064 | 7.333102 | 55.28878 | 1.63E-05 | 0.004155 |
| ENPP7 | -4.71176 | 9.177042 | 65.86635 | 1.81E-05 | 0.004458 |
| TM4SF5 | -1.79026 | 6.861173 | 53.65483 | 1.87E-05 | 0.004557 |
| PLA2G5 | -3.12706 | 7.40029 | 57.60181 | 1.95E-05 | 0.004693 |
| SLC13A5 | -5.32345 | 2.770513 | 56.41981 | 2.07E-05 | 0.004752 |
| SLC34A3 | -3.31517 | 8.458871 | 57.03799 | 2.02E-05 | 0.004752 |
| SLC28A2 | -2.48467 | 8.168879 | 52.50639 | 2.06E-05 | 0.004752 |
| DCC | -2.48288 | 2.713025 | 52.61706 | 2.04E-05 | 0.004752 |
| KISS1 | -3.40466 | 2.775631 | 52.28549 | 2.10E-05 | 0.004754 |
| GBX1 | -4.03541 | 2.810412 | 50.62631 | 2.51E-05 | 0.005345 |
| GLP2R | -1.92139 | 4.524471 | 50.15254 | 2.53E-05 | 0.005345 |
| NOTCH4 | -1.75396 | 4.944269 | 50.2015 | 2.52E-05 | 0.005345 |
| TM6SF2 | -1.90542 | 7.31556 | 49.74777 | 2.62E-05 | 0.00548 |
| LOC103790447 | -1.75459 | 4.590728 | 49.58742 | 2.66E-05 | 0.005499 |
| HSPA12A | -1.80184 | 6.767724 | 49.07731 | 2.78E-05 | 0.005577 |
| LOC103791387 | -3.75245 | 2.810745 | 48.358 | 2.97E-05 | 0.005891 |
| SLC28A1 | -2.77781 | 6.330027 | 50.79082 | 3.13E-05 | 0.00615 |
| LOC100408489 | -1.93819 | 8.099631 | 47.5832 | 3.19E-05 | 0.006197 |
| LOC100406601 | -2.13458 | 6.664264 | 47.01015 | 3.36E-05 | 0.006407 |
| UNC93A | -3.39734 | 6.298714 | 53.95622 | 3.66E-05 | 0.006455 |
| PRLR | -2.40899 | 6.160636 | 46.68305 | 3.70E-05 | 0.006455 |
| SLC11A2 | -2.24993 | 6.958316 | 46.13208 | 3.65E-05 | 0.006455 |
| SLC23A1 | -2.04656 | 6.722801 | 45.92798 | 3.72E-05 | 0.006455 |
| TMEM184A | -2.00428 | 6.459311 | 46.68719 | 3.46E-05 | 0.006455 |
| MACROD1 | -1.68103 | 5.386201 | 46.42347 | 3.55E-05 | 0.006455 |

| | | | | | |
|---|---|---|---|---|---|
| ADH4 | -4.02996 | 7.639672 | 57.24212 | 3.78E-05 | 0.006506 |
| IDO2 | -2.75052 | 5.704174 | 48.12519 | 3.91E-05 | 0.00655 |
| C11H11orf24 | -1.64126 | 7.948333 | 45.57235 | 3.85E-05 | 0.00655 |
| LOC100396914 | -1.55334 | 6.970336 | 45.06767 | 4.04E-05 | 0.006651 |
| MOGAT3 | -2.35912 | 6.912318 | 45.35695 | 4.20E-05 | 0.006855 |
| LOC100388923 | -1.94891 | 4.477449 | 44.51104 | 4.27E-05 | 0.006903 |
| SCARB1 | -3.23464 | 9.092432 | 47.3952 | 4.68E-05 | 0.007101 |
| VSTM2A | -2.91947 | 2.440076 | 43.63991 | 4.65E-05 | 0.007101 |
| THNSL2 | -2.22759 | 6.526544 | 43.79414 | 4.72E-05 | 0.007101 |
| OSGIN1 | -1.92141 | 6.46443 | 43.85633 | 4.55E-05 | 0.007101 |
| REEP6 | -1.87297 | 7.207593 | 43.82531 | 4.56E-05 | 0.007101 |
| PHOSPHO1 | -1.54052 | 6.63689 | 43.6474 | 4.65E-05 | 0.007101 |
| GRAMD1C | -1.71965 | 6.152752 | 43.18 | 4.87E-05 | 0.007264 |
| THSD4 | -1.83909 | 6.424424 | 42.76596 | 5.07E-05 | 0.007514 |
| LOC103789453 | -5.54894 | 2.784761 | 48.23797 | 5.54E-05 | 0.007613 |
| ASRGL1 | -2.4035 | 2.967362 | 42.11218 | 5.42E-05 | 0.007613 |
| SDR16C5 | -2.16854 | 8.118209 | 41.961 | 5.51E-05 | 0.007613 |
| PROB1 | -1.94276 | 4.055483 | 42.16371 | 5.39E-05 | 0.007613 |
| GRAMD1B | -1.90378 | 7.08198 | 42.02974 | 5.47E-05 | 0.007613 |
| SREBF1 | -1.77473 | 8.514473 | 42.39738 | 5.27E-05 | 0.007613 |
| SOAT2 | -1.75901 | 7.719748 | 41.6358 | 5.69E-05 | 0.007684 |
| ABCD1 | -1.53703 | 6.281787 | 41.6135 | 5.71E-05 | 0.007684 |
| SEC22C | -1.88058 | 5.939413 | 41.2986 | 5.90E-05 | 0.007883 |
| TAAR1 | -3.50955 | 3.047237 | 42.04662 | 5.94E-05 | 0.007885 |
| LOC100403801 | -2.06464 | 4.131871 | 41.12049 | 6.01E-05 | 0.007921 |
| FLVCR1 | -1.58537 | 6.157675 | 40.68933 | 6.28E-05 | 0.00823 |
| PCK1 | -3.68336 | 8.454741 | 47.39454 | 7.12E-05 | 0.008406 |
| LOC100405255 | -3.30232 | 3.282232 | 40.48565 | 7.12E-05 | 0.008406 |
| SMLR1 | -2.23127 | 5.765636 | 40.15412 | 6.94E-05 | 0.008406 |
| LOC108590858 | -2.20569 | 4.585738 | 39.84714 | 6.87E-05 | 0.008406 |
| YPEL4 | -2.1301 | 4.025831 | 39.76861 | 6.93E-05 | 0.008406 |
| CES2 | -2.00642 | 7.799968 | 39.76638 | 6.93E-05 | 0.008406 |
| SEC16B | -1.68258 | 7.911082 | 40.13558 | 6.66E-05 | 0.008406 |
| WNK2 | -1.63952 | 7.596727 | 40.08795 | 6.70E-05 | 0.008406 |
| LOC103791091 | -1.51542 | 6.351465 | 39.84427 | 6.87E-05 | 0.008406 |
| UNC5B | -1.40791 | 7.530625 | 39.54126 | 7.10E-05 | 0.008406 |
| PCYT2 | -1.35891 | 7.242178 | 39.64438 | 7.02E-05 | 0.008406 |
| SLC14A2 | -3.33697 | 1.96657 | 39.2372 | 7.34E-05 | 0.008414 |
| LOC108592485 | -3.07261 | 5.49195 | 44.76901 | 7.25E-05 | 0.008414 |
| LOC108593218 | -2.96407 | 2.967843 | 39.21834 | 7.35E-05 | 0.008414 |
| ENTPD7 | -1.71297 | 7.385326 | 39.40678 | 7.20E-05 | 0.008414 |
| SULT1A3 | -1.68323 | 6.97348 | 39.17565 | 7.39E-05 | 0.008414 |
| LOC108594072 | -2.26971 | 3.93214 | 38.42475 | 8.02E-05 | 0.008821 |
| LEXM | -2.00546 | 5.678384 | 38.47664 | 7.97E-05 | 0.008821 |
| CDHR5 | -1.96046 | 9.043005 | 38.61415 | 7.85E-05 | 0.008821 |
| LOC100390305 | -1.95961 | 4.450831 | 38.4502 | 7.99E-05 | 0.008821 |
| CEACAM20 | -1.93258 | 6.395399 | 38.55775 | 7.90E-05 | 0.008821 |

| | | | | | |
|---|---|---|---|---|---|
| KIF26B | -1.91556 | 5.384368 | 38.48847 | 7.96E-05 | 0.008821 |
| SLC25A16 | -1.84382 | 4.572326 | 38.15985 | 8.25E-05 | 0.008929 |
| BCL2L11 | -1.82275 | 5.999842 | 38.10804 | 8.30E-05 | 0.008931 |
| SLC23A3 | -1.64041 | 7.742108 | 37.99086 | 8.41E-05 | 0.008953 |
| LOC100413669 | -1.61548 | 6.475252 | 37.90127 | 8.50E-05 | 0.008987 |
| HAPLN4 | -1.56238 | 5.776982 | 37.79741 | 8.59E-05 | 0.009042 |
| LOC100410434 | -2.00605 | 4.074107 | 37.63453 | 8.75E-05 | 0.009109 |
| PI4KB | -1.31719 | 7.242373 | 37.31322 | 9.07E-05 | 0.009393 |
| TRPV3 | -5.77563 | 2.026932 | 40.00035 | 9.34E-05 | 0.00942 |
| G6PC | -2.41783 | 7.286758 | 39.11334 | 9.27E-05 | 0.00942 |
| EFNA1 | -1.97679 | 6.631637 | 37.08317 | 9.31E-05 | 0.00942 |
| GSTA3 | -2.45075 | 5.007147 | 38.09463 | 9.53E-05 | 0.009554 |
| OBSCN | -2.90393 | 3.624237 | 37.29942 | 9.82E-05 | 0.009727 |
| LOC103793386 | -3.09391 | 1.898164 | 36.39627 | 0.000101 | 0.009768 |
| GGT1 | -2.09361 | 8.349029 | 36.37642 | 0.000101 | 0.009768 |
| SGF29 | -1.73432 | 4.85253 | 36.3928 | 0.000101 | 0.009768 |
| IGFN1 | -3.25237 | 1.776767 | 36.32174 | 0.000102 | 0.00978 |
| OPRK1 | -1.52197 | 4.318585 | 36.2497 | 0.000102 | 0.009813 |
| TTYH2 | -2.15573 | 4.19639 | 36.15367 | 0.000104 | 0.009873 |
| LOC103789058 | -2.8075 | 2.862718 | 35.91728 | 0.000106 | 0.009999 |
| LOC108590272 | -2.06117 | 4.666976 | 35.92352 | 0.000106 | 0.009999 |
| NBL1 | -1.78383 | 7.300295 | 35.92933 | 0.000106 | 0.009999 |
| C11H11orf86 | -1.67796 | 7.783021 | 35.84463 | 0.000107 | 0.010015 |
| GPAT3 | -2.32196 | 8.016111 | 36.77238 | 0.00011 | 0.010133 |
| CYSRT1 | -2.31084 | 5.366887 | 36.68891 | 0.00011 | 0.010133 |
| VEGFA | -1.58513 | 9.183401 | 35.27538 | 0.000115 | 0.010374 |
| LOC100394035 | -1.52386 | 4.173799 | 35.28259 | 0.000115 | 0.010374 |
| LOC108591871 | -3.1064 | 2.042316 | 35.09445 | 0.000117 | 0.01042 |
| RNF186 | -2.48153 | 5.821194 | 37.47039 | 0.000119 | 0.01042 |
| NHSL1 | -2.21031 | 8.053972 | 35.62735 | 0.000119 | 0.01042 |
| CRYM | -2.1424 | 5.716248 | 35.80653 | 0.000116 | 0.01042 |
| LOC100392670 | -1.52854 | 6.604986 | 35.07188 | 0.000118 | 0.01042 |
| RIMS3 | -2.81624 | 1.957671 | 34.79111 | 0.000122 | 0.010459 |
| DCST1 | -1.71212 | 6.658861 | 34.79435 | 0.000121 | 0.010459 |
| LOC108590022 | -1.51853 | 5.116555 | 34.83216 | 0.000121 | 0.010459 |
| LOC108592791 | -2.18506 | 5.407879 | 35.27369 | 0.000124 | 0.010491 |
| LOC103795564 | -2.06619 | 3.204847 | 34.66141 | 0.000123 | 0.010491 |
| FPGS | -1.51218 | 8.38683 | 34.60796 | 0.000124 | 0.010491 |
| NTS | -6.5334 | 4.367162 | 52.36068 | 0.00013 | 0.010807 |
| PABPN1L | -3.78883 | 2.839289 | 36.13614 | 0.000132 | 0.010807 |
| METTL7B | -3.25186 | 5.754308 | 40.90603 | 0.000134 | 0.010807 |
| PHYHIP | -2.5228 | 2.363014 | 33.9369 | 0.000135 | 0.010807 |
| LOC103794023 | -2.05239 | 3.400316 | 33.94723 | 0.000135 | 0.010807 |
| FABP2 | -2.04792 | 6.619509 | 34.55045 | 0.000134 | 0.010807 |
| PLA2G4F | -1.83983 | 5.457942 | 34.21378 | 0.00013 | 0.010807 |
| FBP1 | -1.76437 | 7.994518 | 34.05792 | 0.000133 | 0.010807 |
| MAPKBP1 | -1.49534 | 6.332373 | 34.2719 | 0.000129 | 0.010807 |

| | | | | | |
|---|---|---|---|---|---|
| ACSF2 | -1.9474 | 7.85595 | 33.83738 | 0.000136 | 0.01086 |
| PDXP | -1.48895 | 4.49063 | 33.82817 | 0.000137 | 0.01086 |
| CDHR2 | -1.89306 | 10.28248 | 33.65214 | 0.000139 | 0.011042 |
| METRNL | -1.45773 | 5.419505 | 33.62472 | 0.00014 | 0.011042 |
| PPM1H | -1.78306 | 5.695887 | 33.46235 | 0.000143 | 0.011199 |
| PEPD | -1.46125 | 7.399388 | 33.41012 | 0.000144 | 0.011199 |
| FAM13A | -2.02128 | 8.922188 | 33.37297 | 0.000144 | 0.011205 |
| CD209 | -4.78584 | 1.17117 | 33.01364 | 0.000151 | 0.011574 |
| PLPP2 | -1.43632 | 6.483867 | 33.03138 | 0.000151 | 0.011574 |
| CYP27A1 | -1.3169 | 6.455843 | 32.91934 | 0.000153 | 0.011664 |
| DLL4 | -1.64434 | 4.495619 | 32.77577 | 0.000155 | 0.011782 |
| LOC100405885 | -1.71192 | 3.814422 | 32.69138 | 0.000157 | 0.01186 |
| RORC | -1.58399 | 7.613123 | 32.54894 | 0.00016 | 0.012026 |
| JPH1 | -1.86299 | 4.412357 | 32.31132 | 0.000165 | 0.012201 |
| ABCG5 | -1.75357 | 6.949116 | 32.31969 | 0.000165 | 0.012201 |
| TBX3 | -1.58151 | 9.100632 | 32.1377 | 0.000168 | 0.012352 |
| ABCC6 | -2.03973 | 6.230307 | 32.61003 | 0.000171 | 0.012444 |
| APLNR | -1.85778 | 5.30962 | 32.00661 | 0.000171 | 0.012444 |
| LOC103793353 | -4.66248 | 0.984046 | 31.96939 | 0.000172 | 0.012454 |
| APOC3 | -7.22676 | 7.492426 | 57.08593 | 0.000177 | 0.012528 |
| PRKG2 | -3.00215 | 4.888798 | 36.25779 | 0.000176 | 0.012528 |
| ENOSF1 | -1.57038 | 6.43576 | 31.75406 | 0.000177 | 0.012528 |
| DOPEY2 | -1.40748 | 7.188702 | 31.77801 | 0.000176 | 0.012528 |
| PLEKHG6 | -1.33151 | 7.268008 | 31.85982 | 0.000174 | 0.012528 |
| CHP2 | -2.63544 | 6.714842 | 35.6788 | 0.000178 | 0.012587 |
| ENPP1 | -3.2168 | 5.899907 | 38.56545 | 0.000179 | 0.012596 |
| GARNL3 | -1.79797 | 4.392528 | 31.59373 | 0.000181 | 0.012639 |
| HDAC11 | -1.30755 | 4.969302 | 31.5632 | 0.000181 | 0.012639 |
| SHBG | -3.81637 | 3.50839 | 35.85228 | 0.000187 | 0.012767 |
| ENPP6 | -2.59526 | 6.052078 | 35.05787 | 0.000187 | 0.012767 |
| IGSF22 | -2.37934 | 2.425526 | 31.39896 | 0.000185 | 0.012767 |
| TPSG1 | -2.22851 | 3.720258 | 31.29351 | 0.000188 | 0.012767 |
| FBN3 | -2.2966 | 2.19501 | 31.19896 | 0.00019 | 0.012879 |
| AGMO | -4.20135 | 3.396258 | 36.47682 | 0.000192 | 0.012883 |
| CPNE6 | -2.86424 | 2.943561 | 31.35794 | 0.000191 | 0.012883 |
| SLC38A8 | -3.17264 | 2.596527 | 31.16393 | 0.000196 | 0.012954 |
| SH3D21 | -1.94498 | 7.960587 | 31.17867 | 0.000195 | 0.012954 |
| C19H1orf115 | -1.5683 | 5.01285 | 30.96711 | 0.000196 | 0.012954 |
| IGSF23 | -2.11588 | 9.900842 | 30.7257 | 0.000202 | 0.013124 |
| BMF | -1.75241 | 5.109946 | 30.67679 | 0.000203 | 0.013124 |
| LOC108590009 | -1.71267 | 5.801992 | 30.75049 | 0.000201 | 0.013124 |
| ACAA1 | -1.36177 | 5.536639 | 30.6654 | 0.000204 | 0.013124 |
| SCAP | -1.19074 | 7.166496 | 30.66087 | 0.000204 | 0.013124 |
| TMEM92 | -1.67057 | 7.338278 | 30.34907 | 0.000212 | 0.013392 |
| FLVCR2 | -1.60731 | 7.481589 | 30.38939 | 0.000211 | 0.013392 |
| LOC100390517 | -1.47142 | 7.466838 | 30.31936 | 0.000213 | 0.013392 |
| IGF2BP2 | -1.35976 | 6.414562 | 30.42931 | 0.00021 | 0.013392 |

| | | | | | |
|---|---|---|---|---|---|
| FMNL3 | -1.21756 | 6.232693 | 30.36373 | 0.000212 | 0.013392 |
| ATP8B2 | -1.75238 | 7.559482 | 30.12407 | 0.000219 | 0.013611 |
| SLC6A4 | -1.63153 | 7.985218 | 30.1527 | 0.000218 | 0.013611 |
| NAALADL1 | -5.76393 | 8.26052 | 47.4339 | 0.000234 | 0.01377 |
| LOC103793654 | -3.97721 | 1.207918 | 29.91195 | 0.000225 | 0.01377 |
| SUGCT | -2.95885 | 3.611263 | 31.48823 | 0.000235 | 0.01377 |
| LOC108593527 | -2.12309 | 3.27744 | 29.5896 | 0.000235 | 0.01377 |
| CREB3L3 | -1.88423 | 9.510881 | 29.71829 | 0.000231 | 0.01377 |
| PHLPP2 | -1.57279 | 7.119383 | 29.63922 | 0.000234 | 0.01377 |
| MAF | -1.50151 | 7.451814 | 29.84552 | 0.000227 | 0.01377 |
| ENGASE | -1.46159 | 6.146499 | 29.79245 | 0.000229 | 0.01377 |
| GPD1 | -1.35451 | 8.353481 | 29.60918 | 0.000234 | 0.01377 |
| PER2 | -1.35022 | 6.433036 | 29.71064 | 0.000231 | 0.01377 |
| AGPAT3 | -1.33548 | 8.328424 | 29.68114 | 0.000232 | 0.01377 |
| SPNS3 | -3.14755 | 1.151106 | 29.39451 | 0.000241 | 0.013832 |
| DFNA5 | -3.06853 | 6.424012 | 35.79083 | 0.000241 | 0.013832 |
| TMEM82 | -2.2465 | 5.880844 | 31.37458 | 0.000239 | 0.013832 |
| CNTNAP2 | -1.62141 | 3.204843 | 29.42467 | 0.00024 | 0.013832 |
| ACSS1 | -1.21629 | 8.114223 | 29.40824 | 0.000241 | 0.013832 |
| ADAP1 | -1.18776 | 6.396984 | 29.44061 | 0.00024 | 0.013832 |
| LOC103789105 | -3.36721 | 1.334176 | 29.19507 | 0.000248 | 0.014129 |
| LOC100403233 | -2.18896 | 5.976703 | 30.81708 | 0.00025 | 0.014137 |
| ACVR2B | -1.70783 | 3.380104 | 29.10367 | 0.000251 | 0.014137 |
| PTPRD | -1.68183 | 7.518536 | 29.08599 | 0.000252 | 0.014137 |
| TICAM1 | -1.49546 | 7.537283 | 29.12253 | 0.000251 | 0.014137 |
| EEPD1 | -1.56268 | 5.33855 | 29.05921 | 0.000253 | 0.014139 |
| RHOD | -1.38016 | 6.557369 | 28.97146 | 0.000256 | 0.01419 |
| LOC100896731 | -2.31848 | 4.065997 | 29.30953 | 0.00026 | 0.014213 |
| TSPO2 | -2.12022 | 4.174129 | 28.78297 | 0.000262 | 0.014213 |
| LOC108592792 | -2.10235 | 3.039637 | 28.76888 | 0.000263 | 0.014213 |
| CNNM1 | -1.76509 | 3.272658 | 28.90987 | 0.000258 | 0.014213 |
| PRR5 | -1.69137 | 5.027315 | 28.83557 | 0.000261 | 0.014213 |
| ABTB2 | -1.68459 | 7.424868 | 28.7561 | 0.000263 | 0.014213 |
| LOC103789057 | -1.30519 | 5.15136 | 28.75348 | 0.000264 | 0.014213 |
| HRC | -2.42444 | 2.718239 | 28.5395 | 0.000271 | 0.014456 |
| DNAJB3 | -1.76559 | 5.302836 | 28.54673 | 0.000271 | 0.014456 |
| LRP12 | -1.63026 | 3.660829 | 28.58101 | 0.00027 | 0.014456 |
| RHOU | -1.45459 | 5.584301 | 28.53103 | 0.000272 | 0.014456 |
| LOC108592879 | -1.96138 | 2.82908 | 28.50917 | 0.000273 | 0.01446 |
| MLPH | -1.55465 | 6.177086 | 28.48309 | 0.000274 | 0.014472 |
| DPEP1 | -3.09251 | 7.689324 | 34.43743 | 0.000276 | 0.01449 |
| LOC100386920 | -2.63942 | 3.746892 | 29.53697 | 0.000278 | 0.01449 |
| NECAB2 | -2.0435 | 2.591213 | 28.32421 | 0.00028 | 0.01449 |
| LOC103795688 | -1.91718 | 4.783033 | 28.39792 | 0.000277 | 0.01449 |
| AAK1 | -1.23954 | 7.510114 | 28.38628 | 0.000277 | 0.01449 |
| SLC5A9 | -1.59148 | 7.253407 | 28.21926 | 0.000284 | 0.014614 |
| ZNF599 | -1.96198 | 5.413676 | 28.44204 | 0.000293 | 0.014988 |

| | | | | | |
|---|---|---|---|---|---|
| TMEM253 | -1.54389 | 6.498081 | 27.95376 | 0.000295 | 0.015009 |
| VAV2 | -1.40226 | 7.087092 | 27.97218 | 0.000294 | 0.015009 |
| NELL2 | -3.09544 | 4.232295 | 31.64204 | 0.000297 | 0.015036 |
| NATD1 | -2.04917 | 2.284876 | 27.84787 | 0.000299 | 0.015036 |
| CACNA2D4 | -1.99776 | 4.069379 | 27.89081 | 0.000297 | 0.015036 |
| MMP8 | -1.48755 | 4.800667 | 27.88335 | 0.000298 | 0.015036 |
| HRH2 | -3.00785 | 3.910971 | 30.78148 | 0.0003 | 0.015042 |
| ANPEP | -2.55968 | 12.97132 | 27.78356 | 0.000302 | 0.015042 |
| PFKFB2 | -1.30934 | 6.572199 | 27.77279 | 0.000302 | 0.015042 |
| GRIA1 | -2.27788 | 2.057786 | 27.72142 | 0.000305 | 0.015111 |
| SLC26A10 | -1.80465 | 6.130702 | 27.78477 | 0.000312 | 0.015386 |
| GUCD1 | -1.26372 | 6.457547 | 27.51942 | 0.000313 | 0.015433 |
| PICK1 | -1.30896 | 5.910563 | 27.41941 | 0.000318 | 0.015615 |
| NPC1L1 | -1.88944 | 10.18639 | 27.3888 | 0.000319 | 0.015644 |
| ISG15 | -2.30466 | 3.293388 | 27.36128 | 0.000321 | 0.015666 |
| FABP1 | -1.84375 | 11.31608 | 27.24566 | 0.000326 | 0.015873 |
| LOC103789448 | -5.74491 | 1.915511 | 30.53428 | 0.000331 | 0.015875 |
| KCNJ13 | -2.07869 | 5.852718 | 28.45097 | 0.000332 | 0.015875 |
| ME1 | -1.92431 | 4.919818 | 27.28356 | 0.000332 | 0.015875 |
| LOC108589132 | -1.52797 | 4.750925 | 26.97043 | 0.000339 | 0.016066 |
| DNPEP | -1.17139 | 7.641181 | 26.96092 | 0.00034 | 0.016066 |
| PDLIM2 | -1.45344 | 7.024154 | 26.906 | 0.000342 | 0.016137 |
| EPS8L3 | -1.44051 | 9.189871 | 26.89696 | 0.000343 | 0.016137 |
| TMEM151A | -2.11401 | 2.517769 | 26.77734 | 0.000349 | 0.01634 |
| KIAA1549 | -1.77558 | 5.619026 | 26.90382 | 0.00035 | 0.016361 |
| LOC100896180 | -1.58133 | 5.898145 | 26.71992 | 0.000352 | 0.016398 |
| VMO1 | -1.31214 | 6.826812 | 26.64405 | 0.000356 | 0.01654 |
| ASPDH | -2.72946 | 2.922849 | 26.86989 | 0.000358 | 0.016578 |
| LOC103796600 | -2.38421 | 3.688019 | 26.85636 | 0.000366 | 0.016803 |
| CAMK1 | -1.69114 | 5.035017 | 26.43047 | 0.000367 | 0.016803 |
| TMEM150B | -1.27033 | 6.316678 | 26.42234 | 0.000367 | 0.016803 |
| LOC100405958 | -3.61078 | 7.370473 | 34.91307 | 0.000377 | 0.016947 |
| GATM | -3.23043 | 7.82166 | 32.71223 | 0.000376 | 0.016947 |
| LRTM2 | -2.93924 | 2.121284 | 26.29699 | 0.000374 | 0.016947 |
| LOC103793587 | -1.89836 | 3.872656 | 26.2776 | 0.000375 | 0.016947 |
| SPATA31A3 | -1.68612 | 4.652228 | 26.31185 | 0.000373 | 0.016947 |
| LOC100402010 | -2.2249 | 2.72242 | 26.16904 | 0.000381 | 0.017078 |
| TOM1 | -1.21824 | 6.583134 | 26.18438 | 0.000381 | 0.017078 |
| TEF | -2.13998 | 5.140264 | 27.33359 | 0.000386 | 0.017212 |
| LPAR3 | -2.19014 | 3.7715 | 26.11259 | 0.000393 | 0.017355 |
| JARID2 | -1.18691 | 7.587121 | 25.99713 | 0.000391 | 0.017355 |
| SYT12 | -1.76964 | 5.424355 | 26.0804 | 0.000395 | 0.017398 |
| FLRT1 | -1.81184 | 4.559241 | 25.83566 | 0.000401 | 0.017573 |
| CHKA | -1.51828 | 6.104321 | 25.79292 | 0.000403 | 0.017573 |
| TTLL11 | -1.50092 | 3.615796 | 25.8373 | 0.000401 | 0.017573 |
| VDR | -1.47489 | 7.733791 | 25.80185 | 0.000403 | 0.017573 |
| GOT1 | -1.13026 | 6.640136 | 25.81633 | 0.000402 | 0.017573 |

| | | | | | |
|---|---|---|---|---|---|
| IYD | -2.39876 | 4.641335 | 27.57431 | 0.00041 | 0.017603 |
| SLC5A4 | -2.08193 | 7.112745 | 27.32652 | 0.000409 | 0.017603 |
| ZNF484 | -1.53964 | 5.099345 | 25.71099 | 0.000408 | 0.017603 |
| SLC6A18 | -5.47411 | 1.006452 | 25.77369 | 0.000413 | 0.017633 |
| AGPAT1 | -1.27295 | 7.165368 | 25.60653 | 0.000415 | 0.017633 |
| F10 | -2.04605 | 6.477297 | 27.12639 | 0.000416 | 0.017653 |
| PON1 | -3.16429 | 3.038823 | 27.27339 | 0.000417 | 0.017658 |
| SLC27A4 | -1.13636 | 6.868826 | 25.52812 | 0.00042 | 0.017725 |
| OSBPL1A | -1.5984 | 6.939202 | 25.49578 | 0.000422 | 0.017734 |
| ABCB1 | -2.3449 | 8.566054 | 27.52206 | 0.000425 | 0.017743 |
| ENTPD5 | -2.08979 | 7.658697 | 26.93236 | 0.000425 | 0.017743 |
| NCKAP5 | -1.24247 | 6.634573 | 25.43476 | 0.000426 | 0.017743 |
| MATN2 | -1.16522 | 6.661133 | 25.45915 | 0.000424 | 0.017743 |
| TMCO4 | -1.42053 | 5.386092 | 25.40244 | 0.000428 | 0.017792 |
| IMMP2L | -1.74884 | 3.546979 | 25.34272 | 0.000432 | 0.017915 |
| LOC103789374 | -3.94679 | 1.356348 | 25.32625 | 0.000433 | 0.017921 |
| CYP4V2 | -1.39986 | 4.770036 | 25.29656 | 0.000435 | 0.017964 |
| LOC100399071 | -2.33226 | 7.754171 | 27.76301 | 0.000444 | 0.018083 |
| COMT | -1.30802 | 6.964484 | 25.16717 | 0.000443 | 0.018083 |
| PCSK5 | -1.25272 | 8.904884 | 25.17852 | 0.000443 | 0.018083 |
| CDK18 | -1.20719 | 6.629806 | 25.1637 | 0.000444 | 0.018083 |
| ATOH7 | -2.99818 | 4.710056 | 29.33338 | 0.000452 | 0.01826 |
| LOC108587886 | -2.42678 | 3.036274 | 25.00655 | 0.000454 | 0.018267 |
| PKP1 | -3.7775 | 0.981232 | 24.93243 | 0.00046 | 0.018387 |
| ABHD6 | -1.7054 | 8.072001 | 24.92303 | 0.00046 | 0.018387 |
| ALDH18A1 | -1.31445 | 8.377336 | 24.85714 | 0.000465 | 0.018536 |
| DGAT1 | -1.36858 | 9.712168 | 24.82647 | 0.000467 | 0.018586 |
| LOC103791965 | -2.01584 | 4.432089 | 25.13056 | 0.000471 | 0.018619 |
| RAB31 | -1.45439 | 5.449715 | 24.77801 | 0.000471 | 0.018619 |
| TTC7B | -1.32549 | 4.605135 | 24.79284 | 0.00047 | 0.018619 |
| CNTN2 | -1.69889 | 2.988079 | 24.73181 | 0.000474 | 0.018666 |
| ACBD4 | -1.23333 | 7.306738 | 24.61274 | 0.000483 | 0.018859 |
| CIDEB | -1.70047 | 8.190227 | 24.45051 | 0.000495 | 0.019234 |
| F13B | -2.75745 | 2.480754 | 24.35075 | 0.000509 | 0.019578 |
| SEMA6A | -1.76164 | 5.245208 | 24.35583 | 0.000516 | 0.019756 |
| LHX5 | -2.86368 | 2.387253 | 24.24001 | 0.000522 | 0.019832 |
| FAM83G | -1.70844 | 6.762592 | 24.46979 | 0.000521 | 0.019832 |
| ARSF | -1.90913 | 8.828617 | 24.37187 | 0.000529 | 0.019937 |
| RBP2 | -1.79978 | 9.654315 | 24.01228 | 0.00053 | 0.019948 |
| LCT | -3.50647 | 8.856894 | 30.31104 | 0.000532 | 0.019962 |
| AGRN | -1.34366 | 8.493948 | 23.92064 | 0.000538 | 0.020121 |
| SCTR | -1.25863 | 5.721217 | 23.84579 | 0.000545 | 0.020282 |
| TUBA8 | -2.91232 | 2.515586 | 24.10141 | 0.000552 | 0.020453 |
| SLC5A1 | -2.07618 | 9.431478 | 24.33777 | 0.00056 | 0.020453 |
| LIN7B | -1.89612 | 4.00008 | 23.75351 | 0.000553 | 0.020453 |
| SLC2A5 | -1.7561 | 8.842447 | 23.74859 | 0.000553 | 0.020453 |
| SDSL | -1.65875 | 5.00795 | 23.70322 | 0.000557 | 0.020453 |

| | | | | | |
|---|---|---|---|---|---|
| CBLN3 | -1.51301 | 5.533566 | 23.68459 | 0.000559 | 0.020453 |
| SQSTM1 | -1.30926 | 8.051929 | 23.6333 | 0.000563 | 0.020517 |
| LOC103789763 | -3.26373 | 1.991487 | 23.79213 | 0.000569 | 0.020634 |
| BST1 | -1.81237 | 4.266157 | 23.55266 | 0.000571 | 0.020634 |
| NPPB | -7.92347 | 5.134456 | 40.55225 | 0.000587 | 0.020982 |
| LOC108592439 | -4.4765 | 1.38757 | 23.70887 | 0.00059 | 0.020982 |
| F11 | -3.04767 | 1.218697 | 23.32311 | 0.000592 | 0.020982 |
| CES3 | -2.1438 | 6.67186 | 25.64724 | 0.000586 | 0.020982 |
| SPNS2 | -1.79866 | 3.500125 | 23.32587 | 0.000592 | 0.020982 |
| SLC2A9 | -1.56631 | 7.217292 | 23.33037 | 0.000591 | 0.020982 |
| LDHAL6B | -1.51902 | 4.221023 | 23.36234 | 0.000588 | 0.020982 |
| PRKAG2 | -1.31352 | 5.953742 | 23.36636 | 0.000588 | 0.020982 |
| TMEM178A | -2.3623 | 2.016206 | 23.2732 | 0.000597 | 0.021087 |
| PDILT | -2.95539 | 2.17135 | 23.30156 | 0.000604 | 0.021245 |
| LOC103792310 | -2.88323 | 2.971628 | 24.15055 | 0.000623 | 0.021772 |
| LOC103789718 | -2.72659 | 7.696283 | 27.26542 | 0.000624 | 0.021772 |
| LOC100894293 | -2.39795 | 4.809546 | 25.20693 | 0.000633 | 0.022009 |
| ERBB2 | -1.32699 | 8.544567 | 22.90879 | 0.000633 | 0.022009 |
| MB21D2 | -1.49898 | 5.093211 | 22.8097 | 0.000644 | 0.022328 |
| EPHX1 | -1.77039 | 7.445422 | 23.32177 | 0.000651 | 0.022536 |
| ECE1 | -1.27653 | 8.371656 | 22.71375 | 0.000654 | 0.022601 |
| MEP1A | -5.97714 | 9.317535 | 36.25905 | 0.000656 | 0.02263 |
| LOC100391891 | -2.80295 | 4.997502 | 26.65708 | 0.000661 | 0.022699 |
| BVES | -1.35093 | 4.308014 | 22.61836 | 0.000664 | 0.022745 |
| NEURL3 | -2.13218 | 3.865652 | 22.92951 | 0.000667 | 0.022783 |
| LOC103792464 | -2.31641 | 7.022288 | 25.3857 | 0.000669 | 0.022793 |
| LOC108588221 | -1.23126 | 6.829305 | 22.52721 | 0.000674 | 0.022936 |
| CACNB2 | -1.08012 | 5.509675 | 22.45258 | 0.000683 | 0.023098 |
| BRSK1 | -2.02528 | 4.798187 | 23.38842 | 0.000686 | 0.02317 |
| SLC13A2 | -2.13334 | 7.247554 | 24.56476 | 0.000687 | 0.023176 |
| RNF224 | -2.72216 | 6.738477 | 26.99096 | 0.000695 | 0.023306 |
| LOC103793682 | -1.69481 | 3.265272 | 22.33536 | 0.000696 | 0.023306 |
| AGAP1 | -1.06843 | 6.545349 | 22.34784 | 0.000695 | 0.023306 |
| LOC108588075 | -2.13906 | 3.481017 | 22.4178 | 0.0007 | 0.023385 |
| RAB40C | -1.2348 | 5.335194 | 22.26252 | 0.000705 | 0.023468 |
| NEB | -1.43492 | 5.965707 | 22.22303 | 0.000709 | 0.023501 |
| CDC14A | -1.3174 | 6.390765 | 22.19094 | 0.000713 | 0.023587 |
| PHYH | -1.46684 | 5.951531 | 22.15159 | 0.000718 | 0.023702 |
| ZCCHC14 | -1.13008 | 8.363751 | 22.05103 | 0.00073 | 0.024063 |
| IL1RAP | -1.62759 | 4.258444 | 22.02569 | 0.000733 | 0.024083 |
| LOC100399775 | -1.36781 | 7.024572 | 22.03055 | 0.000732 | 0.024083 |
| ERICH4 | -2.29243 | 4.615991 | 23.76025 | 0.000744 | 0.024275 |
| NDFIP2 | -1.27493 | 6.289698 | 21.92806 | 0.000745 | 0.024275 |
| NOS2 | -1.24021 | 7.583356 | 21.94751 | 0.000743 | 0.024275 |
| UCN3 | -2.86376 | 1.53905 | 21.89745 | 0.000749 | 0.024359 |
| LRRC75B | -1.84013 | 4.114594 | 21.87449 | 0.000752 | 0.024413 |
| TBC1D13 | -0.98633 | 6.541914 | 21.75175 | 0.000768 | 0.024841 |

| | | | | | |
|---|---|---|---|---|---|
| CACNA1E | -1.6612 | 4.200586 | 21.71637 | 0.000772 | 0.024865 |
| ALAD | -1.14714 | 6.265611 | 21.72208 | 0.000772 | 0.024865 |
| GDPD2 | -2.52838 | 5.788546 | 25.25657 | 0.000776 | 0.024939 |
| PTPRT | -1.97139 | 2.436553 | 21.59836 | 0.000788 | 0.025242 |
| LOC103790597 | -1.77021 | 3.368767 | 21.51718 | 0.000799 | 0.025509 |
| GABBR2 | -2.18599 | 5.263232 | 23.3936 | 0.000815 | 0.025896 |
| TMEM184B | -1.14251 | 7.267277 | 21.3586 | 0.000821 | 0.025995 |
| SLC39A4 | -1.41913 | 8.239201 | 21.32781 | 0.000825 | 0.02609 |
| LOC103788937 | -2.22083 | 2.684876 | 21.29482 | 0.00083 | 0.026196 |
| LOC103788766 | -3.09324 | 4.810838 | 25.88476 | 0.000838 | 0.026357 |
| ALPP | -2.91997 | 3.342104 | 23.2696 | 0.000837 | 0.026357 |
| GCKR | -3.39529 | 2.960902 | 23.61255 | 0.00085 | 0.02648 |
| RAB30 | -1.96628 | 5.564994 | 22.55559 | 0.000845 | 0.02648 |
| CIB2 | -1.22354 | 4.693141 | 21.17093 | 0.000848 | 0.02648 |
| FLCN | -1.11111 | 6.4393 | 21.15705 | 0.00085 | 0.02648 |
| ENPP3 | -3.24573 | 8.123794 | 27.08304 | 0.000853 | 0.026538 |
| CIDEC | -1.99809 | 7.375341 | 22.76685 | 0.000864 | 0.026731 |
| JPH2 | -1.56469 | 5.942305 | 21.20029 | 0.000862 | 0.026731 |
| ARHGEF26 | -1.11753 | 5.735188 | 21.05728 | 0.000865 | 0.026731 |
| LOC108587861 | -2.69761 | 1.863572 | 21.01306 | 0.000871 | 0.026842 |
| TACR1 | -2.12755 | 1.740912 | 20.99585 | 0.000874 | 0.02688 |
| TNFRSF11A | -1.09861 | 6.529172 | 20.97843 | 0.000877 | 0.026918 |
| STKLD1 | -1.61602 | 3.146413 | 20.95343 | 0.00088 | 0.02695 |
| CCDC153 | -2.08437 | 2.055009 | 20.8612 | 0.000895 | 0.027256 |
| PER3 | -1.40229 | 5.575446 | 20.81276 | 0.000902 | 0.027356 |
| CDYL2 | -2.0253 | 5.353821 | 22.25444 | 0.000911 | 0.027454 |
| MPP1 | -1.30817 | 7.424612 | 20.74303 | 0.000913 | 0.027477 |
| LMTK2 | -1.44233 | 8.937966 | 20.6827 | 0.000923 | 0.027659 |
| IDUA | -1.40058 | 6.640199 | 20.67876 | 0.000924 | 0.027659 |
| LOC108588148 | -4.02687 | 1.629753 | 21.31631 | 0.000957 | 0.028556 |
| RAB7B | -2.16117 | 3.689569 | 21.00842 | 0.000958 | 0.028556 |
| COBL | -1.14364 | 7.469552 | 20.4478 | 0.000962 | 0.028586 |
| LOC103790969 | -2.02278 | 5.978161 | 22.19043 | 0.000978 | 0.028834 |
| PCYT1A | -0.98685 | 7.125284 | 20.37747 | 0.000974 | 0.028834 |
| KCNK17 | -2.32307 | 3.209079 | 20.75469 | 0.000988 | 0.028993 |
| MLXIPL | -1.37071 | 7.602458 | 20.29035 | 0.000989 | 0.028993 |
| APOB | -2.69262 | 14.04363 | 20.36427 | 0.001 | 0.029171 |
| LOC103789231 | -1.69606 | 2.538353 | 20.23826 | 0.000999 | 0.029171 |
| ZNF274 | -1.10584 | 5.553993 | 20.23805 | 0.000999 | 0.029171 |
| TMEM37 | -1.67051 | 6.98002 | 20.78068 | 0.001012 | 0.029474 |
| LOC103787661 | -3.47566 | 1.356794 | 19.93924 | 0.001053 | 0.029915 |
| C2CD4D | -2.60922 | 1.745154 | 19.9411 | 0.001053 | 0.029915 |
| PYY | -2.19202 | 3.109139 | 20.04872 | 0.001057 | 0.029915 |
| LOC100399428 | -2.0829 | 5.508665 | 21.70203 | 0.001067 | 0.029915 |
| LOC100387361 | -2.05413 | 3.705889 | 20.24968 | 0.001051 | 0.029915 |
| RHOV | -2.00009 | 3.939373 | 20.2549 | 0.001068 | 0.029915 |
| LOC108590596 | -1.98131 | 2.234501 | 20.03979 | 0.001035 | 0.029915 |

| | | | | | |
|---|---|---|---|---|---|
| SUCO | -1.626 | 7.27253 | 20.46056 | 0.001031 | 0.029915 |
| ABCG8 | -1.57746 | 7.706537 | 20.08582 | 0.001067 | 0.029915 |
| LOC103789177 | -1.51394 | 2.997858 | 19.90364 | 0.00106 | 0.029915 |
| MMP28 | -1.44947 | 5.432919 | 19.90071 | 0.001061 | 0.029915 |
| SIDT2 | -1.31294 | 8.191115 | 19.86724 | 0.001067 | 0.029915 |
| CCS | -1.29409 | 6.66961 | 19.87798 | 0.001065 | 0.029915 |
| ACY1 | -1.26938 | 6.259623 | 19.91542 | 0.001058 | 0.029915 |
| DPYD | -1.19285 | 6.639966 | 20.0476 | 0.001033 | 0.029915 |
| MAGI1 | -1.13299 | 6.771463 | 19.88499 | 0.001064 | 0.029915 |
| ACP2 | -1.11051 | 5.721654 | 20.02812 | 0.001037 | 0.029915 |
| DENND1A | -1.05359 | 6.739952 | 19.98776 | 0.001044 | 0.029915 |
| TBC1D10A | -0.99756 | 6.100317 | 19.9118 | 0.001059 | 0.029915 |
| C18H1orf226 | -0.98884 | 5.721421 | 19.9942 | 0.001043 | 0.029915 |
| MRO | -1.82587 | 5.17699 | 20.58634 | 0.00108 | 0.030038 |
| LOC103795595 | -1.67795 | 4.085584 | 19.77169 | 0.001086 | 0.030038 |
| LOC103793368 | -1.34828 | 5.037167 | 19.7546 | 0.001089 | 0.030088 |
| SEC14L2 | -1.761 | 6.860844 | 20.6661 | 0.001101 | 0.030339 |
| COL26A1 | -2.50437 | 2.187199 | 19.67854 | 0.001104 | 0.030373 |
| ILVBL | -1.14868 | 6.644376 | 19.67117 | 0.001106 | 0.030373 |
| FAM188B | -1.55257 | 4.479048 | 19.62857 | 0.001114 | 0.030566 |
| CHAT | -1.73907 | 2.178807 | 19.57008 | 0.001126 | 0.030849 |
| LOC103787324 | -1.31872 | 6.022212 | 19.56006 | 0.001128 | 0.030862 |
| LOC108591681 | -1.59171 | 5.439669 | 19.79268 | 0.001131 | 0.030885 |
| PDK4 | -2.48548 | 6.967425 | 23.05695 | 0.001147 | 0.03095 |
| TMEM200C | -1.80824 | 2.38838 | 19.47945 | 0.001145 | 0.03095 |
| EGLN3 | -1.61324 | 7.296416 | 19.8761 | 0.001152 | 0.03095 |
| ADGRG4 | -1.38998 | 5.22765 | 19.4605 | 0.001149 | 0.03095 |
| HDAC10 | -1.36369 | 8.326211 | 19.44437 | 0.001153 | 0.03095 |
| SLC25A25 | -1.34541 | 7.083778 | 19.47042 | 0.001147 | 0.03095 |
| UBALD2 | -1.11209 | 5.924743 | 19.48923 | 0.001143 | 0.03095 |
| FOXO4 | -0.96671 | 5.990056 | 19.49818 | 0.001141 | 0.03095 |
| LOC100414419 | -1.09613 | 8.126462 | 19.42549 | 0.001157 | 0.031014 |
| TMEM150A | -1.38018 | 4.410672 | 19.2898 | 0.001186 | 0.031577 |
| LOC100403563 | -1.3094 | 3.443709 | 19.2951 | 0.001185 | 0.031577 |
| SMAD9 | -1.04362 | 5.184534 | 19.29318 | 0.001185 | 0.031577 |
| EGFLAM | -2.09132 | 2.784508 | 19.2182 | 0.001201 | 0.031777 |
| SUSD2 | -1.55601 | 8.256002 | 19.31255 | 0.0012 | 0.031777 |
| MAB21L1 | -1.34728 | 3.8076 | 19.2188 | 0.001201 | 0.031777 |
| RSAD1 | -1.36748 | 3.567463 | 19.16495 | 0.001213 | 0.032003 |
| LOC108591882 | -2.30276 | 1.951032 | 19.12744 | 0.001222 | 0.032035 |
| LOC100389624 | -1.20854 | 6.358608 | 19.10274 | 0.001227 | 0.03211 |
| GPAM | -1.56857 | 7.76753 | 19.32105 | 0.001231 | 0.032123 |
| SLC25A34 | -1.55266 | 5.697503 | 19.2795 | 0.001241 | 0.032292 |
| BCO1 | -3.18378 | 6.287916 | 25.22789 | 0.001253 | 0.032486 |
| ZFYVE1 | -0.98921 | 6.789105 | 18.99039 | 0.001253 | 0.032486 |
| CARMIL3 | -1.23619 | 3.967949 | 18.97337 | 0.001257 | 0.032494 |
| LOC103796376 | -1.24062 | 8.410428 | 18.93344 | 0.001267 | 0.032519 |

| | | | | | |
|---|---|---|---|---|---|
| PPARA | -1.08519 | 6.371403 | 18.89814 | 0.001275 | 0.032617 |
| LOC100407810 | -1.49008 | 6.392133 | 19.04287 | 0.001289 | 0.032804 |
| SLC22A4 | -1.28757 | 4.933561 | 18.81668 | 0.001295 | 0.032804 |
| LOC100389765 | -1.46478 | 2.988032 | 18.77466 | 0.001305 | 0.032972 |
| ENTPD2 | -1.53668 | 6.598328 | 19.04139 | 0.001316 | 0.033066 |
| LOC103794406 | -1.27412 | 4.192781 | 18.75078 | 0.001311 | 0.033066 |
| MTMR11 | -1.18905 | 7.041702 | 18.72509 | 0.001317 | 0.033066 |
| PCTP | -1.80391 | 3.170019 | 18.70039 | 0.001323 | 0.033088 |
| SERPINE1 | -1.69445 | 5.387241 | 19.30881 | 0.00132 | 0.033088 |
| BAIAP2L2 | -1.27993 | 7.981726 | 18.67083 | 0.001331 | 0.033218 |
| LOC103790002 | -1.59254 | 5.609481 | 18.98606 | 0.001338 | 0.033295 |
| COL17A1 | -1.33112 | 9.462126 | 18.63966 | 0.001338 | 0.033295 |
| ZDHHC19 | -2.56385 | 2.490846 | 18.8447 | 0.00135 | 0.033529 |
| INSR | -1.24948 | 7.217431 | 18.57785 | 0.001354 | 0.033556 |
| DGKD | -1.08982 | 9.121374 | 18.56325 | 0.001358 | 0.033562 |
| LOC103794071 | -1.818 | 2.860495 | 18.49467 | 0.001376 | 0.033784 |
| PLIN2 | -1.67348 | 8.419678 | 18.90333 | 0.001372 | 0.033784 |
| LOC103793814 | -1.18829 | 4.625293 | 18.50072 | 0.001374 | 0.033784 |
| ARHGEF16 | -1.16182 | 7.325482 | 18.51002 | 0.001372 | 0.033784 |
| HMOX1 | -1.7207 | 5.650952 | 19.23495 | 0.001384 | 0.033809 |
| LOC100393797 | -1.64005 | 4.095538 | 18.47284 | 0.001381 | 0.033809 |
| LOC108588948 | -1.8511 | 2.744263 | 18.44431 | 0.001389 | 0.033831 |
| CD109 | -1.142 | 4.758707 | 18.41169 | 0.001397 | 0.033971 |
| PGM1 | -1.03407 | 5.7664 | 18.382 | 0.001405 | 0.03406 |
| FOLH1B | -4.35296 | 5.214732 | 27.15711 | 0.001425 | 0.034364 |
| TRPV6 | -2.68912 | 6.846854 | 22.5834 | 0.001428 | 0.034364 |
| TRPM4 | -1.87293 | 8.90997 | 19.07777 | 0.001428 | 0.034364 |
| LOC108590682 | -1.73255 | 3.649747 | 18.26067 | 0.001438 | 0.034377 |
| IL22RA1 | -1.21482 | 5.549897 | 18.25133 | 0.001441 | 0.034377 |
| NR1D2 | -1.20319 | 5.221196 | 18.25248 | 0.001441 | 0.034377 |
| LOC100896839 | -4.28458 | 1.075703 | 18.47624 | 0.001445 | 0.034383 |
| LOC108593885 | -1.27682 | 4.729236 | 18.20907 | 0.001453 | 0.034443 |
| MYH15 | -1.36465 | 4.678169 | 18.19648 | 0.001456 | 0.034445 |
| PDE1C | -1.36283 | 5.181575 | 18.18954 | 0.001458 | 0.034445 |
| FCGRT | -1.07445 | 8.103247 | 18.17582 | 0.001462 | 0.034469 |
| TMC5 | -1.83655 | 7.962371 | 19.25441 | 0.001465 | 0.03447 |
| LOC108591857 | -1.62238 | 4.802544 | 18.37666 | 0.001466 | 0.03447 |
| SFXN3 | -1.1798 | 6.475674 | 18.12963 | 0.001475 | 0.034631 |
| LOC103794997 | -2.04087 | 5.149036 | 19.69154 | 0.001478 | 0.034666 |
| ACY3 | -3.59961 | 3.329922 | 21.5473 | 0.001495 | 0.034814 |
| COL13A1 | -1.77727 | 3.397784 | 18.0664 | 0.001493 | 0.034814 |
| STX3 | -1.63232 | 6.202882 | 18.72908 | 0.001496 | 0.034814 |
| LOC108593960 | -2.53528 | 1.016082 | 18.00511 | 0.001511 | 0.03485 |
| DPP4 | -2.33995 | 7.823262 | 20.78988 | 0.001509 | 0.03485 |
| LOC100413731 | -2.21821 | 1.320555 | 18.01733 | 0.001507 | 0.03485 |
| MMP27 | -1.33633 | 5.254087 | 17.98393 | 0.001517 | 0.03485 |
| LOC103789529 | -2.41442 | 2.201798 | 17.94524 | 0.001528 | 0.034894 |

| | | | | | |
|---|---|---|---|---|---|
| NEDD4L | -1.18025 | 7.495302 | 17.94904 | 0.001527 | 0.034894 |
| MAP1LC3A | -1.19586 | 4.925993 | 17.91623 | 0.001537 | 0.034923 |
| LOC100390841 | -1.17965 | 6.301341 | 17.91197 | 0.001538 | 0.034923 |
| SUSD6 | -0.99693 | 8.28054 | 17.91788 | 0.001536 | 0.034923 |
| PXDC1 | -1.35126 | 7.754146 | 17.87642 | 0.001549 | 0.035123 |
| GPSM1 | -2.37888 | 5.553477 | 20.63419 | 0.001578 | 0.035664 |
| LOC108593950 | -5.67115 | 2.018631 | 21.24013 | 0.001584 | 0.03576 |
| LOC103796759 | -3.51589 | 4.460115 | 23.15602 | 0.001592 | 0.035842 |
| CEACAM18 | -1.50896 | 8.573421 | 17.8136 | 0.001592 | 0.035842 |
| LOC108591798 | -1.36528 | 3.081498 | 17.69119 | 0.001606 | 0.03603 |
| GJB6 | -1.90378 | 2.402124 | 17.66965 | 0.001612 | 0.036063 |
| LOC108589304 | -1.73892 | 4.046723 | 17.79035 | 0.00161 | 0.036063 |
| USP2 | -1.36415 | 5.5067 | 17.66868 | 0.001613 | 0.036063 |
| XYLB | -1.5339 | 3.959768 | 17.65019 | 0.001618 | 0.036109 |
| PLA2G15 | -1.26169 | 5.720482 | 17.6264 | 0.001626 | 0.03616 |
| SLC43A2 | -1.17941 | 8.398207 | 17.62876 | 0.001625 | 0.03616 |
| RPS6KA5 | -1.11548 | 5.70711 | 17.61458 | 0.00163 | 0.036194 |
| SLC3A2 | -1.13104 | 7.228288 | 17.58976 | 0.001638 | 0.036253 |
| LOC103788363 | -1.43718 | 2.924744 | 17.55712 | 0.001648 | 0.036394 |
| AVPI1 | -1.16151 | 3.837498 | 17.54725 | 0.001651 | 0.036423 |
| SEPP1 | -2.42868 | 10.83114 | 18.91261 | 0.00168 | 0.036438 |
| MME | -2.38935 | 9.749853 | 19.32205 | 0.001668 | 0.036438 |
| MATK | -1.45081 | 6.448458 | 17.66108 | 0.001681 | 0.036438 |
| LOC103796764 | -1.35188 | 4.639274 | 17.49772 | 0.001668 | 0.036438 |
| TESK2 | -1.12112 | 6.409208 | 17.52006 | 0.00166 | 0.036438 |
| NUAK2 | -1.11363 | 5.61044 | 17.49006 | 0.00167 | 0.036438 |
| XPNPEP1 | -1.02805 | 7.174286 | 17.50897 | 0.001664 | 0.036438 |
| TBC1D24 | -0.9603 | 6.220629 | 17.49112 | 0.00167 | 0.036438 |
| ACSL6 | -1.23315 | 4.756507 | 17.42933 | 0.00169 | 0.036607 |
| C18H1orf105 | -3.18784 | 0.604108 | 17.42151 | 0.001693 | 0.036622 |
| LOC103791639 | -2.21312 | 4.761875 | 19.17188 | 0.00171 | 0.036801 |
| ABHD2 | -1.42921 | 10.19133 | 17.36771 | 0.001711 | 0.036801 |
| GNPDA1 | -1.05989 | 4.944897 | 17.37697 | 0.001708 | 0.036801 |
| RASSF4 | -1.05652 | 6.232269 | 17.35718 | 0.001714 | 0.036801 |
| LOC103794660 | -2.65597 | 2.686393 | 17.96938 | 0.001772 | 0.037495 |
| CCK | -2.36591 | 5.77127 | 20.14722 | 0.001772 | 0.037495 |
| SH2D7 | -1.38997 | 3.920812 | 17.19081 | 0.001772 | 0.037495 |
| ATP7B | -1.08908 | 6.894239 | 17.21278 | 0.001764 | 0.037495 |
| MTMR3 | -0.89711 | 7.572402 | 17.19458 | 0.001771 | 0.037495 |
| ADRA2B | -1.94963 | 1.694217 | 17.16854 | 0.00178 | 0.037618 |
| HES3 | -2.31455 | 1.953819 | 17.11439 | 0.001799 | 0.037736 |
| COL9A1 | -2.14761 | 2.833408 | 17.22655 | 0.001795 | 0.037736 |
| LOC103792971 | -1.49893 | 4.476931 | 17.10893 | 0.001801 | 0.037736 |
| LOC103796510 | -1.02869 | 4.85973 | 17.12655 | 0.001795 | 0.037736 |
| LOC100397266 | -1.018 | 6.636772 | 17.06992 | 0.001815 | 0.037949 |
| GALNT2 | -0.96312 | 7.454956 | 17.01132 | 0.001837 | 0.038272 |
| AGMAT | -1.23998 | 4.841208 | 16.97401 | 0.00185 | 0.038476 |

| | | | | | |
|---|---|---|---|---|---|
| GGT7 | -2.40086 | 6.32634 | 20.01622 | 0.001879 | 0.038902 |
| CAMKV | -1.52071 | 3.647763 | 16.89774 | 0.001879 | 0.038902 |
| SLC25A42 | -1.48066 | 4.064611 | 16.8822 | 0.001885 | 0.038917 |
| LOC108591233 | -1.39342 | 3.235537 | 16.8867 | 0.001883 | 0.038917 |
| IRS2 | -0.93541 | 5.175099 | 16.86526 | 0.001891 | 0.03899 |
| LOC100389414 | -2.17066 | 2.622025 | 16.881 | 0.001902 | 0.03905 |
| MMP13 | -1.71079 | 2.428495 | 16.83858 | 0.001902 | 0.03905 |
| RFX2 | -1.42139 | 4.4854 | 16.84678 | 0.001898 | 0.03905 |
| ENPEP | -2.06794 | 8.778869 | 18.31468 | 0.001918 | 0.03916 |
| TTC22 | -1.12388 | 6.950024 | 16.79695 | 0.001918 | 0.03916 |
| KLHDC3 | -0.95088 | 6.124429 | 16.79613 | 0.001918 | 0.03916 |
| DBP | -1.86542 | 5.785317 | 18.11767 | 0.001939 | 0.039262 |
| LOC108594082 | -1.75331 | 1.993885 | 16.74573 | 0.001938 | 0.039262 |
| SLC7A7 | -1.47227 | 7.374323 | 17.01146 | 0.001934 | 0.039262 |
| IGDCC4 | -1.12783 | 4.032437 | 16.7451 | 0.001938 | 0.039262 |
| TRAK1 | -1.01344 | 9.167509 | 16.75688 | 0.001933 | 0.039262 |
| HIPK2 | -0.99749 | 8.495954 | 16.75466 | 0.001934 | 0.039262 |
| FCN1 | -5.80352 | 5.027269 | 25.972 | 0.001945 | 0.039301 |
| STON2 | -1.40621 | 6.336364 | 16.89875 | 0.001948 | 0.039301 |
| CELF3 | -1.25814 | 3.031489 | 16.71442 | 0.00195 | 0.039301 |
| KIAA1161 | -1.23357 | 6.252198 | 16.71322 | 0.00195 | 0.039301 |
| PAG1 | -0.96621 | 6.660355 | 16.68646 | 0.001961 | 0.039418 |
| KCNK5 | -1.26455 | 6.620959 | 16.62475 | 0.001986 | 0.039827 |
| HSD17B2 | -1.87011 | 6.409418 | 18.08727 | 0.002021 | 0.040362 |
| ACE | -1.60641 | 11.8066 | 16.53887 | 0.002021 | 0.040362 |
| NOCT | -1.31357 | 4.391325 | 16.5158 | 0.00203 | 0.04043 |
| IDNK | -1.2207 | 5.375858 | 16.50526 | 0.002035 | 0.040472 |
| MCF2L2 | -1.7721 | 4.946477 | 17.25522 | 0.002045 | 0.040515 |
| TLCD2 | -1.40556 | 4.026698 | 16.47729 | 0.002046 | 0.040515 |
| PIP5K1C | -1.18374 | 7.037187 | 16.4716 | 0.002049 | 0.040515 |
| SORBS3 | -1.10436 | 6.956385 | 16.48833 | 0.002042 | 0.040515 |
| YPEL3 | -0.88691 | 5.758976 | 16.46485 | 0.002052 | 0.040515 |
| UGT2A3 | -2.79953 | 6.402424 | 21.05271 | 0.002066 | 0.040618 |
| BCAS1 | -1.87067 | 7.801801 | 17.77894 | 0.00206 | 0.040618 |
| ELMO2 | -1.11457 | 5.957602 | 16.4292 | 0.002067 | 0.040618 |
| HKDC1 | -1.42851 | 9.034254 | 16.42112 | 0.00207 | 0.04063 |
| ABI3 | -1.01639 | 5.599504 | 16.38469 | 0.002086 | 0.040851 |
| TMEM8B | -1.30856 | 3.655472 | 16.37036 | 0.002092 | 0.040878 |
| PTK2 | -1.2164 | 8.488864 | 16.36882 | 0.002092 | 0.040878 |
| GLYCTK | -1.06592 | 6.501534 | 16.36247 | 0.002095 | 0.040878 |
| HMGCS2 | -4.29744 | 7.037072 | 24.5984 | 0.002103 | 0.040883 |
| LOC103793036 | -2.66157 | 0.805044 | 16.34146 | 0.002104 | 0.040883 |
| LOC103796719 | -1.73005 | 1.993301 | 16.34876 | 0.002101 | 0.040883 |
| TIMD4 | -2.85059 | 2.639298 | 17.51411 | 0.002109 | 0.040938 |
| GUCY2C | -1.65782 | 7.548609 | 17.07669 | 0.002116 | 0.040963 |
| BRINP1 | -2.31335 | 1.382128 | 16.29737 | 0.002123 | 0.041024 |
| RGL1 | -1.02525 | 5.820016 | 16.28464 | 0.002129 | 0.041074 |

| | | | | | |
|---|---|---|---|---|---|
| CRY2 | -1.08021 | 5.49305 | 16.27589 | 0.002133 | 0.041107 |
| LOC108588589 | -2.09251 | 2.120129 | 16.26143 | 0.002139 | 0.041148 |
| MYOM2 | -1.41046 | 2.724301 | 16.25412 | 0.002142 | 0.041169 |
| LOC103794042 | -1.70935 | 2.571361 | 16.23476 | 0.002151 | 0.041293 |
| LOC108591704 | -1.30356 | 3.568213 | 16.21098 | 0.002162 | 0.041372 |
| LOC100414398 | -1.68753 | 2.059481 | 16.18883 | 0.002172 | 0.041439 |
| VPS11 | -0.97182 | 5.849948 | 16.19524 | 0.002169 | 0.041439 |
| OTOP2 | -2.16522 | 2.34303 | 16.16235 | 0.002184 | 0.041589 |
| LOC108592647 | -1.76248 | 2.901502 | 16.15794 | 0.002186 | 0.041589 |
| LOC100397252 | -1.28841 | 4.374194 | 16.15707 | 0.002186 | 0.041589 |
| LGI3 | -1.63831 | 2.326812 | 16.12239 | 0.002202 | 0.041806 |
| ADM | -1.49574 | 4.650273 | 16.17902 | 0.002206 | 0.041846 |
| SLC15A1 | -2.12281 | 8.939189 | 17.6676 | 0.002213 | 0.0419 |
| RUNDC3B | -1.55609 | 3.944377 | 16.08958 | 0.002217 | 0.041927 |
| KDM4B | -0.93624 | 6.519658 | 15.96873 | 0.002273 | 0.04266 |
| ZER1 | -0.92619 | 7.137067 | 15.9709 | 0.002272 | 0.04266 |
| LOC100391255 | -1.00519 | 6.5392 | 15.95837 | 0.002278 | 0.042711 |
| LOC100398471 | -3.27829 | 0.833766 | 15.91868 | 0.002297 | 0.042941 |
| BRPF3 | -0.93271 | 5.764295 | 15.92075 | 0.002296 | 0.042941 |
| DGKQ | -1.34797 | 8.04968 | 15.9125 | 0.0023 | 0.042955 |
| LOC100400861 | -2.1825 | 2.817909 | 16.11959 | 0.002308 | 0.043063 |
| LOC103789992 | -2.1255 | 2.454866 | 15.87219 | 0.00232 | 0.043223 |
| LOC103793440 | -1.67259 | 2.719512 | 15.85606 | 0.002328 | 0.043223 |
| LOC100413713 | -1.19981 | 3.819241 | 15.86648 | 0.002322 | 0.043223 |
| SLC45A4 | -1.06585 | 6.106221 | 15.86179 | 0.002325 | 0.043223 |
| MIGA2 | -0.9552 | 7.305072 | 15.85515 | 0.002328 | 0.043223 |
| OTOP3 | -1.13729 | 7.600392 | 15.84423 | 0.002333 | 0.04328 |
| LOC103794639 | -1.84797 | 2.91689 | 15.79385 | 0.002358 | 0.043573 |
| LOC103789328 | -1.64963 | 3.175721 | 15.78337 | 0.002363 | 0.043588 |
| CYP11A1 | -3.55149 | 3.330825 | 19.0711 | 0.00238 | 0.043691 |
| LOC108592102 | -2.18509 | 1.693579 | 15.75505 | 0.002377 | 0.043691 |
| NID1 | -1.10355 | 7.500067 | 15.76361 | 0.002373 | 0.043691 |
| LOC108590327 | -1.12763 | 4.752673 | 15.74175 | 0.002384 | 0.043718 |
| LOC108589387 | -2.3359 | 3.417297 | 16.84324 | 0.002389 | 0.043766 |
| BGN | -1.17229 | 3.510533 | 15.71645 | 0.002397 | 0.043868 |
| HFE | -1.39537 | 3.745364 | 15.68345 | 0.002414 | 0.044008 |
| EPHX2 | -1.31409 | 8.419711 | 15.65579 | 0.002428 | 0.04414 |
| PPP1R15A | -1.12451 | 5.583979 | 15.66381 | 0.002424 | 0.04414 |
| ZSWIM5 | -1.09224 | 3.770983 | 15.65914 | 0.002426 | 0.04414 |
| RALGPS1 | -0.90065 | 6.826798 | 15.64852 | 0.002432 | 0.044167 |
| UNC79 | -1.20932 | 3.45723 | 15.63312 | 0.002439 | 0.044269 |
| CAPN12 | -3.32963 | 6.040773 | 21.40244 | 0.002489 | 0.044491 |
| MGAM | -2.10685 | 8.758658 | 17.15905 | 0.002478 | 0.044491 |
| TM4SF4 | -1.95756 | 3.504397 | 15.95054 | 0.002493 | 0.044491 |
| SLC25A45 | -1.58965 | 3.869684 | 15.55728 | 0.002479 | 0.044491 |
| SPIRE2 | -1.448 | 5.448251 | 15.73783 | 0.002494 | 0.044491 |
| FOXH1 | -1.32414 | 5.546328 | 15.56977 | 0.002478 | 0.044491 |

| | | | | | |
|---|---|---|---|---|---|
| FAM84A | -1.15189 | 6.077738 | 15.55804 | 0.002479 | 0.044491 |
| ATG2A | -0.96844 | 7.612029 | 15.58492 | 0.002465 | 0.044491 |
| BSDC1 | -0.86188 | 6.535432 | 15.53747 | 0.002489 | 0.044491 |
| CDIPT | -0.8353 | 6.48414 | 15.54777 | 0.002484 | 0.044491 |
| LOC100384909 | -2.31132 | 1.995698 | 15.51926 | 0.002499 | 0.044513 |
| SPATA6L | -1.70707 | 3.472852 | 15.50478 | 0.002507 | 0.044555 |
| FAM212B | -1.40294 | 4.182739 | 15.47582 | 0.002522 | 0.044555 |
| SAT2 | -1.17378 | 5.768622 | 15.47818 | 0.002521 | 0.044555 |
| KIF21B | -1.16126 | 7.902879 | 15.4829 | 0.002519 | 0.044555 |
| C2CD2L | -1.09507 | 7.552836 | 15.47777 | 0.002521 | 0.044555 |
| ZBTB7B | -1.06314 | 8.830997 | 15.48858 | 0.002515 | 0.044555 |
| ARHGAP10 | -0.93181 | 6.118912 | 15.47989 | 0.00252 | 0.044555 |
| LOC100393377 | -2.35768 | 2.342847 | 15.563 | 0.002564 | 0.045127 |
| LOC100896466 | -1.43663 | 2.95989 | 15.37337 | 0.002578 | 0.045138 |
| DQX1 | -1.17783 | 6.646107 | 15.36798 | 0.002581 | 0.045138 |
| GABARAPL1 | -1.02206 | 6.090846 | 15.37774 | 0.002576 | 0.045138 |
| NBEA | -0.87154 | 6.73047 | 15.37161 | 0.002579 | 0.045138 |
| LOC103796559 | -2.07725 | 1.261904 | 15.35773 | 0.002587 | 0.045188 |
| MUC17 | -1.78646 | 8.268161 | 16.4203 | 0.002589 | 0.045188 |
| DUSP1 | -1.17756 | 7.209172 | 15.31544 | 0.00261 | 0.045401 |
| ODF2L | -1.17524 | 4.258283 | 15.2969 | 0.002621 | 0.045427 |
| CDH23 | -1.24505 | 5.607545 | 15.2828 | 0.002629 | 0.045515 |
| FUT2 | -1.13102 | 6.885778 | 15.26944 | 0.002636 | 0.045605 |
| NGFR | -1.37729 | 4.385876 | 15.25866 | 0.002642 | 0.045628 |
| KIF13A | -0.91123 | 7.871926 | 15.2306 | 0.002658 | 0.045822 |
| CISH | -1.38377 | 3.566533 | 15.20356 | 0.002674 | 0.046008 |
| PRSS8 | -1.01738 | 8.208437 | 15.15943 | 0.002699 | 0.046241 |
| LOC103790721 | -1.92087 | 5.851372 | 16.72735 | 0.002704 | 0.046283 |
| PLEKHM2 | -1.04414 | 7.102691 | 15.12773 | 0.002718 | 0.046454 |
| LOC108593703 | -3.92299 | 0.786807 | 15.13599 | 0.002734 | 0.04654 |
| LOC103791515 | -1.40396 | 4.082485 | 15.10636 | 0.002731 | 0.04654 |
| OGDHL | -1.74406 | 1.954416 | 15.09554 | 0.002737 | 0.046553 |
| LOC108594128 | -1.70755 | 4.730544 | 15.69229 | 0.00276 | 0.04678 |
| LOC103787972 | -0.98679 | 5.423258 | 15.02738 | 0.002778 | 0.046955 |
| TCN2 | -1.4112 | 7.976246 | 15.15522 | 0.002795 | 0.047171 |
| VPS37C | -0.99211 | 6.544147 | 15.0017 | 0.002793 | 0.047171 |
| FSTL5 | -2.55203 | 1.167581 | 14.98661 | 0.002802 | 0.047207 |
| LOC103788228 | -1.01481 | 5.355215 | 14.98877 | 0.002801 | 0.047207 |
| STARD3 | -0.93249 | 6.093179 | 14.91801 | 0.002845 | 0.047832 |
| OLFML2A | -1.22534 | 3.815339 | 14.89223 | 0.002861 | 0.047895 |
| CNKSR1 | -1.08415 | 6.374643 | 14.89711 | 0.002858 | 0.047895 |
| LOC103790448 | -1.35542 | 3.432619 | 14.85209 | 0.002886 | 0.048147 |
| DENND5B | -0.85618 | 6.344894 | 14.84837 | 0.002888 | 0.048147 |
| GRAMD4 | -1.1745 | 8.205548 | 14.77391 | 0.002936 | 0.048818 |
| SIK3 | -0.82494 | 6.906306 | 14.77332 | 0.002936 | 0.048818 |
| PDXK | -1.07048 | 7.661301 | 14.7623 | 0.002943 | 0.048894 |
| LOC108587995 | -1.8606 | 3.431603 | 14.99687 | 0.00295 | 0.048962 |

| | | | | | |
|---|---|---|---|---|---|
| KIF5A | -1.25185 | 5.312234 | 14.73034 | 0.002964 | 0.049154 |
| ANKDD1B | -1.43291 | 5.78631 | 15.0787 | 0.002972 | 0.049158 |
| BRINP3 | -3.67152 | 1.328016 | 15.1462 | 0.002984 | 0.049304 |
| MAST1 | -1.08861 | 3.352852 | 14.69307 | 0.002988 | 0.049304 |
| ACVR1B | -1.06747 | 6.346891 | 14.69573 | 0.002987 | 0.049304 |
| CNDP2 | -1.07031 | 8.79309 | 14.67801 | 0.002998 | 0.049425 |
| LOC108588949 | -1.6428 | 3.379274 | 14.67367 | 0.003001 | 0.049431 |
| TOLLIP | -0.99243 | 5.599217 | 14.66833 | 0.003005 | 0.049446 |
| CPT1A | -0.85107 | 7.437087 | 14.65398 | 0.003014 | 0.049561 |
| LOC103795080 | -1.95369 | 1.819216 | 14.6244 | 0.003034 | 0.049563 |
| LOC108591905 | -1.04995 | 3.76681 | 14.6291 | 0.003031 | 0.049563 |
| AMOTL2 | -0.98051 | 5.576477 | 14.64054 | 0.003023 | 0.049563 |
| ENG | -0.85673 | 6.315066 | 14.63476 | 0.003027 | 0.049563 |
| CYP2D19 | -2.34713 | 3.534854 | 15.82841 | 0.003063 | 0.049977 |
| SLC39A14 | -1.42102 | 8.307041 | 14.72396 | 0.003068 | 0.049977 |
| LIMCH1 | -1.07936 | 4.70542 | 14.56972 | 0.003071 | 0.04998 |

**Supp. Table 5b. Genes upregulated in Non-Stricture**

| Gene name | logFC | logCPM | F | PValue | FDR |
|---|---|---|---|---|---|
| OLFM4 | 3.962044 | 9.226766 | 108.8616 | 6.81E-07 | 0.001456 |
| GSDMC | 2.445208 | 7.207831 | 89.46273 | 1.75E-06 | 0.001872 |
| GPX2 | 2.699347 | 5.381887 | 90.79005 | 1.63E-06 | 0.001872 |
| DUOXA2 | 3.071026 | 4.29129 | 90.73075 | 1.64E-06 | 0.001872 |
| DUOX2 | 2.364712 | 6.272863 | 77.02953 | 3.55E-06 | 0.001992 |
| EHF | 2.582943 | 5.993913 | 74.24658 | 4.22E-06 | 0.002131 |
| TFAP2A | 3.702115 | 2.81782 | 73.88935 | 4.32E-06 | 0.002131 |
| KCNA3 | 2.023745 | 4.512767 | 60.75597 | 1.07E-05 | 0.003478 |
| PLA2G3 | 2.916604 | 3.954147 | 58.82957 | 1.23E-05 | 0.003796 |
| LOC100395944 | 2.492541 | 6.833861 | 55.87768 | 1.56E-05 | 0.004109 |
| LOC100897014 | 2.63958 | 2.915863 | 56.1545 | 1.52E-05 | 0.004109 |
| LOC108591301 | 5.130923 | 3.243093 | 62.37386 | 1.47E-05 | 0.004109 |
| PPP1R1B | 2.475696 | 5.31787 | 54.53408 | 1.74E-05 | 0.004346 |
| KIAA1324 | 2.005451 | 4.732674 | 51.79839 | 2.19E-05 | 0.004899 |
| KRT18 | 1.795348 | 8.041112 | 51.50027 | 2.25E-05 | 0.004969 |
| LOC100399118 | 2.270286 | 3.752208 | 51.02851 | 2.34E-05 | 0.005118 |
| LOC103796010 | 2.634405 | 3.399831 | 49.14112 | 2.76E-05 | 0.005577 |
| LOC100896501 | 2.975348 | 9.929896 | 49.59439 | 2.75E-05 | 0.005577 |
| JCHAIN | 3.080872 | 9.162999 | 49.66322 | 3.32E-05 | 0.006388 |
| LRP4 | 1.697053 | 6.47698 | 46.55236 | 3.51E-05 | 0.006455 |
| ATP8B4 | 2.789379 | 4.292576 | 46.54699 | 3.63E-05 | 0.006455 |
| PYCR1 | 2.97523 | 3.931594 | 46.80927 | 3.54E-05 | 0.006455 |
| KLRG1 | 5.938983 | 3.593731 | 57.96486 | 3.70E-05 | 0.006455 |
| NPNT | 1.947806 | 6.250068 | 45.47412 | 3.89E-05 | 0.00655 |
| LOC100398523 | 1.790694 | 6.4347 | 45.1761 | 4.00E-05 | 0.006639 |
| ITLN1 | 2.076331 | 5.537288 | 44.37982 | 4.32E-05 | 0.006934 |
| LOC103788970 | 2.621533 | 3.484916 | 44.01368 | 4.48E-05 | 0.007101 |
| LOC100386050 | 2.853356 | 6.036297 | 47.2868 | 4.72E-05 | 0.007101 |

| | | | | | |
|---|---|---|---|---|---|
| CDC42EP5 | 2.056285 | 4.144682 | 42.53012 | 5.20E-05 | 0.007579 |
| CBS | 2.357826 | 2.709264 | 42.60377 | 5.16E-05 | 0.007579 |
| KRT19 | 2.173298 | 8.187196 | 42.31948 | 5.31E-05 | 0.007613 |
| WFDC2 | 2.669839 | 3.856664 | 42.10512 | 5.43E-05 | 0.007613 |
| SLC12A2 | 1.677458 | 7.294872 | 41.79088 | 5.60E-05 | 0.007652 |
| RNASE6 | 1.757274 | 4.463135 | 40.59913 | 6.34E-05 | 0.008253 |
| HK2 | 2.539275 | 4.389793 | 40.75906 | 6.43E-05 | 0.008314 |
| MB21D1 | 2.457943 | 3.493471 | 40.37836 | 6.49E-05 | 0.008334 |
| CTSC | 1.811502 | 5.645378 | 39.82204 | 6.89E-05 | 0.008406 |
| MYB | 2.543018 | 4.088127 | 40.09511 | 6.69E-05 | 0.008406 |
| LOC103796425 | 2.312639 | 4.449813 | 39.32659 | 7.27E-05 | 0.008414 |
| EPN3 | 1.880179 | 4.77258 | 38.24635 | 8.18E-05 | 0.008929 |
| BPIFB2 | 7.503519 | 1.296987 | 48.77528 | 8.22E-05 | 0.008929 |
| STX11 | 3.296873 | 2.216236 | 37.98531 | 8.42E-05 | 0.008953 |
| LCN2 | 4.166038 | 5.815036 | 49.86364 | 8.71E-05 | 0.009109 |
| LOC100395418 | 1.619053 | 6.626063 | 37.14162 | 9.25E-05 | 0.00942 |
| ACSL1 | 2.034153 | 4.745712 | 37.16719 | 9.22E-05 | 0.00942 |
| LOC103796419 | 1.885803 | 6.411231 | 36.75946 | 9.66E-05 | 0.009639 |
| MUC20 | 3.191131 | 1.903679 | 36.58984 | 9.85E-05 | 0.009727 |
| LOC108587745 | 1.579337 | 4.873034 | 36.45202 | 0.0001 | 0.009768 |
| ITGB8 | 1.827005 | 3.609076 | 35.82016 | 0.000108 | 0.010015 |
| HSP90B1 | 1.579922 | 9.245958 | 35.58773 | 0.000111 | 0.010143 |
| VSIG4 | 2.427386 | 2.861624 | 35.52375 | 0.000111 | 0.010171 |
| NUCB2 | 1.817569 | 4.257497 | 35.02631 | 0.000118 | 0.01042 |
| C21H21orf62 | 2.674306 | 2.532796 | 35.01878 | 0.000118 | 0.01042 |
| WARS | 1.779394 | 6.752129 | 34.78214 | 0.000122 | 0.010459 |
| LOC100413268 | 1.637642 | 7.617772 | 34.60998 | 0.000124 | 0.010491 |
| SOX9 | 1.438265 | 5.019244 | 34.31662 | 0.000129 | 0.010807 |
| ITM2A | 2.844543 | 2.904798 | 34.13901 | 0.000131 | 0.010807 |
| KLRB1 | 4.705906 | 2.692451 | 38.11415 | 0.000134 | 0.010807 |
| ASNS | 2.025318 | 5.071846 | 33.44198 | 0.000143 | 0.011199 |
| LOC108593906 | 3.378508 | 1.526441 | 33.21258 | 0.000147 | 0.011383 |
| DMBT1 | 2.734435 | 10.31888 | 34.21401 | 0.000154 | 0.011705 |
| BTLA | 1.96615 | 3.460636 | 32.49212 | 0.000161 | 0.012058 |
| PF4 | 3.688202 | 2.637275 | 33.75874 | 0.000162 | 0.012058 |
| PLA2G7 | 1.642063 | 4.63396 | 32.16366 | 0.000168 | 0.012352 |
| BTK | 1.670961 | 3.413705 | 32.12322 | 0.000169 | 0.012352 |
| IKZF2 | 2.5318 | 5.663541 | 34.92499 | 0.000174 | 0.012528 |
| LOC100406154 | 2.972265 | 8.350544 | 36.14314 | 0.000182 | 0.012639 |
| DMXL2 | 1.369716 | 7.260451 | 31.49181 | 0.000183 | 0.012668 |
| GSAP | 1.563377 | 5.150296 | 31.3306 | 0.000187 | 0.012767 |
| KIF20B | 1.878192 | 4.01424 | 31.11607 | 0.000192 | 0.012883 |
| SH2D1A | 4.408838 | 2.188382 | 32.71609 | 0.000193 | 0.012915 |
| EVI2A | 1.890409 | 3.238176 | 30.93327 | 0.000197 | 0.012967 |
| RAB27A | 2.053283 | 5.394442 | 31.33921 | 0.000199 | 0.013108 |
| LTV1 | 1.630539 | 4.301767 | 30.7045 | 0.000203 | 0.013124 |
| CYBB | 1.678891 | 4.721198 | 30.60433 | 0.000205 | 0.013178 |

| | | | | | |
|---|---|---|---|---|---|
| SERPINB1 | 1.542636 | 5.400874 | 30.36407 | 0.000212 | 0.013392 |
| PHLDA2 | 1.646706 | 3.678468 | 30.30818 | 0.000214 | 0.013392 |
| LOC100387926 | 4.240066 | 2.219473 | 31.67773 | 0.000219 | 0.013611 |
| SLC9B2 | 2.066209 | 3.188229 | 30.05736 | 0.000221 | 0.013668 |
| LRG1 | 3.221159 | 1.877805 | 30.02193 | 0.000222 | 0.013689 |
| TXNDC5 | 1.453488 | 6.823233 | 29.83243 | 0.000228 | 0.01377 |
| KIF11 | 2.274832 | 3.426722 | 29.59826 | 0.000235 | 0.01377 |
| SLC7A5 | 2.291466 | 4.288866 | 30.18492 | 0.000232 | 0.01377 |
| SLC1A3 | 2.297171 | 2.554417 | 29.9326 | 0.000224 | 0.01377 |
| GPR171 | 2.435082 | 2.749519 | 29.85668 | 0.000227 | 0.01377 |
| AGR2 | 2.468175 | 7.580354 | 32.84095 | 0.000228 | 0.01377 |
| SYN3 | 3.11333 | 4.901027 | 34.45665 | 0.000236 | 0.013781 |
| IQGAP3 | 2.225813 | 3.453123 | 29.30564 | 0.000244 | 0.013959 |
| LYZ | 3.276051 | 6.628272 | 36.57538 | 0.000252 | 0.014137 |
| ASPHD2 | 2.273795 | 2.599933 | 29.04028 | 0.000253 | 0.014139 |
| APOC1 | 2.829656 | 2.570012 | 28.98768 | 0.000255 | 0.01419 |
| CTSS | 1.593181 | 5.921056 | 28.75394 | 0.000264 | 0.014213 |
| ELOVL6 | 2.130804 | 5.001819 | 29.67329 | 0.000258 | 0.014213 |
| LOC100390453 | 2.973232 | 2.557663 | 28.78859 | 0.000262 | 0.014213 |
| LOC100386951 | 2.073651 | 3.935462 | 28.56975 | 0.00027 | 0.014456 |
| PDIA4 | 1.285278 | 7.778836 | 28.34876 | 0.000279 | 0.01449 |
| PLA2G2A | 1.525584 | 7.234126 | 28.29916 | 0.000281 | 0.01449 |
| TACSTD2 | 2.648335 | 2.463471 | 28.3379 | 0.000279 | 0.01449 |
| LOC103790605 | 3.046276 | 1.289483 | 28.30859 | 0.00028 | 0.01449 |
| ERAP2 | 1.974406 | 4.679599 | 28.26333 | 0.000286 | 0.014697 |
| C1H9orf152 | 4.500001 | 1.661164 | 28.28539 | 0.000299 | 0.015036 |
| ADGRE5 | 1.493354 | 5.6257 | 27.77211 | 0.000302 | 0.015042 |
| LOC100405150 | 2.675372 | 6.368434 | 31.90111 | 0.000307 | 0.015195 |
| ATP2C2 | 1.359516 | 6.214102 | 27.23059 | 0.000327 | 0.015873 |
| FAM46C | 1.453963 | 6.23223 | 27.21704 | 0.000327 | 0.015873 |
| LOC100396537 | 1.644237 | 5.104393 | 27.13946 | 0.000331 | 0.015875 |
| STC1 | 1.699325 | 3.165922 | 27.15062 | 0.00033 | 0.015875 |
| ERVMER34-1 | 1.763335 | 4.440378 | 27.14501 | 0.000331 | 0.015875 |
| GNL3 | 1.583539 | 4.739078 | 27.0656 | 0.000335 | 0.015942 |
| LOC100404432 | 2.069559 | 5.826944 | 28.36087 | 0.000335 | 0.015945 |
| BIRC5 | 2.292177 | 2.698946 | 26.83076 | 0.000346 | 0.016253 |
| LOC108589742 | 1.716485 | 5.139984 | 26.6067 | 0.000358 | 0.016578 |
| IL5RA | 2.644438 | 2.480751 | 26.50256 | 0.000363 | 0.016765 |
| CENPE | 2.187958 | 4.389327 | 27.08447 | 0.000366 | 0.016803 |
| PDIA6 | 1.268272 | 6.615737 | 26.36571 | 0.00037 | 0.016904 |
| FCRLA | 3.276244 | 1.018496 | 26.32737 | 0.000373 | 0.016947 |
| SLC39A10 | 1.829411 | 4.755317 | 26.0852 | 0.000386 | 0.017212 |
| CDC20 | 1.908052 | 3.53544 | 25.97174 | 0.000393 | 0.017355 |
| TXK | 2.827534 | 4.936877 | 29.79998 | 0.000392 | 0.017355 |
| PTMA | 1.410108 | 7.581205 | 25.77728 | 0.000404 | 0.017574 |
| LOC100405375 | 1.226075 | 6.246643 | 25.71769 | 0.000408 | 0.017603 |
| SAMD9L | 2.300417 | 5.1718 | 27.76453 | 0.000408 | 0.017603 |

| | | | | | |
|---|---|---|---|---|---|
| KNTC1 | 2.2701 | 2.842799 | 25.6102 | 0.000415 | 0.017633 |
| LOC103796238 | 2.916399 | 2.056822 | 25.60729 | 0.000415 | 0.017633 |
| LOC103795879 | 3.503258 | 1.4266 | 25.65456 | 0.000412 | 0.017633 |
| KLRC1 | 5.598641 | 3.849965 | 36.23567 | 0.000421 | 0.017734 |
| TFRC | 1.497568 | 5.777222 | 25.2635 | 0.000437 | 0.017981 |
| DCDC2 | 3.154254 | 1.185524 | 25.262 | 0.000437 | 0.017981 |
| SELPLG | 2.117974 | 4.333735 | 25.61368 | 0.000445 | 0.018083 |
| C4H6orf141 | 2.639864 | 3.060257 | 25.57113 | 0.000441 | 0.018083 |
| LOC103788928 | 1.658788 | 3.763883 | 25.02275 | 0.000453 | 0.01826 |
| SLC12A8 | 2.177124 | 3.424289 | 25.03182 | 0.000453 | 0.01826 |
| LOC103795187 | 4.247395 | 3.975976 | 32.18587 | 0.000451 | 0.01826 |
| CXCL1 | 2.628558 | 2.763026 | 25.06829 | 0.000456 | 0.018295 |
| TFF3 | 2.399044 | 5.940485 | 27.82107 | 0.000472 | 0.018619 |
| SAA1 | 2.246401 | 3.27342 | 24.71632 | 0.000475 | 0.018673 |
| LOC100897022 | 1.429493 | 8.474231 | 24.69151 | 0.000477 | 0.018706 |
| ANXA1 | 1.292348 | 4.978177 | 24.66929 | 0.000479 | 0.018733 |
| LOC108587665 | 1.185546 | 5.131457 | 24.51536 | 0.00049 | 0.019108 |
| FOXM1 | 2.481161 | 3.172033 | 24.77116 | 0.000495 | 0.019234 |
| NEK2 | 2.789172 | 1.655793 | 24.42342 | 0.000497 | 0.019266 |
| RARRES1 | 1.607155 | 5.786665 | 24.39375 | 0.0005 | 0.019317 |
| LOC100391553 | 1.543588 | 3.854755 | 24.3563 | 0.000503 | 0.019391 |
| CD47 | 1.209198 | 6.292447 | 24.27537 | 0.000509 | 0.019578 |
| LOC100894491 | 4.721867 | 0.739385 | 24.18633 | 0.000516 | 0.019756 |
| PLEK | 1.631599 | 2.94958 | 24.15637 | 0.000519 | 0.01981 |
| NDC80 | 2.503134 | 2.021327 | 24.12772 | 0.000521 | 0.019832 |
| CDC42EP1 | 1.280882 | 4.796458 | 24.09539 | 0.000524 | 0.019843 |
| LOC103790458 | 2.022817 | 3.990988 | 24.19225 | 0.000526 | 0.019902 |
| LOC100396946 | 1.645862 | 5.026691 | 24.05169 | 0.000527 | 0.019902 |
| TOP2A | 2.029955 | 5.010389 | 24.8964 | 0.000538 | 0.020121 |
| PAH | 3.154322 | 1.182442 | 23.88233 | 0.000541 | 0.020204 |
| FRMD4B | 1.224768 | 4.780587 | 23.67252 | 0.00056 | 0.020453 |
| DDX21 | 1.450035 | 5.512408 | 23.7411 | 0.000554 | 0.020453 |
| FOXA3 | 1.901619 | 4.285982 | 23.79371 | 0.000556 | 0.020453 |
| CXCL9 | 2.682374 | 1.70069 | 23.72181 | 0.000555 | 0.020453 |
| PIK3AP1 | 1.327925 | 5.936944 | 23.62939 | 0.000564 | 0.020517 |
| DNPH1 | 4.031472 | 1.796086 | 24.39427 | 0.000567 | 0.020592 |
| SSR1 | 1.142649 | 6.378694 | 23.54673 | 0.000571 | 0.020634 |
| LOC100397835 | 3.023333 | 1.71915 | 23.49014 | 0.000576 | 0.020783 |
| ENC1 | 1.548819 | 5.907908 | 23.36295 | 0.000588 | 0.020982 |
| GALNT3 | 1.571241 | 5.351755 | 23.3153 | 0.000593 | 0.020982 |
| UAP1 | 1.136713 | 5.798146 | 23.23446 | 0.000601 | 0.02118 |
| LOC108593911 | 2.372885 | 3.806075 | 24.13605 | 0.000605 | 0.021245 |
| LOC100395837 | 1.219576 | 5.120498 | 23.16524 | 0.000607 | 0.021302 |
| GPNMB | 2.144064 | 3.778696 | 23.42637 | 0.000613 | 0.021469 |
| RPLP2 | 1.139751 | 6.860548 | 22.64381 | 0.000661 | 0.022699 |
| ESCO2 | 2.618233 | 1.711383 | 22.65354 | 0.00066 | 0.022699 |
| LOC100399303 | 1.504627 | 3.205386 | 22.61006 | 0.000665 | 0.022745 |

| | | | | | |
|---|---|---|---|---|---|
| LOC108590090 | 2.055308 | 3.278972 | 22.51142 | 0.000676 | 0.022949 |
| LOC103788629 | 3.793723 | 1.046243 | 22.50239 | 0.000677 | 0.022949 |
| FGFBP1 | 5.936828 | 1.685591 | 24.95231 | 0.000692 | 0.02329 |
| LOC103795324 | 3.6083 | 0.942041 | 22.26618 | 0.000704 | 0.023468 |
| STK39 | 1.87751 | 4.567011 | 22.54042 | 0.000707 | 0.023501 |
| XCL1 | 4.668828 | 0.699268 | 22.22896 | 0.000708 | 0.023501 |
| LOC100399888 | 1.661558 | 6.061788 | 22.28666 | 0.000737 | 0.024175 |
| CLDN2 | 4.332263 | 4.668447 | 30.75448 | 0.000738 | 0.024175 |
| POLR3G | 2.417725 | 1.658617 | 21.83807 | 0.000757 | 0.024522 |
| LOC100397874 | 1.447451 | 4.040956 | 21.72756 | 0.000771 | 0.024865 |
| SFN | 1.733245 | 6.809038 | 22.37455 | 0.000777 | 0.024939 |
| LOC100393754 | 1.526687 | 6.023471 | 21.52982 | 0.000798 | 0.025509 |
| ERCC6L | 2.244741 | 2.759264 | 21.45228 | 0.000808 | 0.025751 |
| LOC100386390 | 1.91778 | 2.470039 | 21.40242 | 0.000815 | 0.025896 |
| STARD10 | 1.30395 | 5.298548 | 21.38662 | 0.000817 | 0.025913 |
| CAPN2 | 1.158854 | 6.795211 | 21.17345 | 0.000847 | 0.02648 |
| NOP56 | 1.37741 | 6.608775 | 21.18951 | 0.000845 | 0.02648 |
| LOC108590372 | 2.559203 | 2.131239 | 21.05559 | 0.000865 | 0.026731 |
| CENPU | 2.116667 | 2.991071 | 21.04294 | 0.000867 | 0.026747 |
| LOC100391701 | 2.043431 | 5.796823 | 22.80399 | 0.00088 | 0.02695 |
| PPIF | 1.419525 | 5.941772 | 20.9108 | 0.000887 | 0.027107 |
| MYO5C | 1.545871 | 5.476093 | 20.8792 | 0.000892 | 0.027214 |
| LOC100405488 | 2.250259 | 9.131196 | 22.59652 | 0.000897 | 0.027294 |
| CHIT1 | 4.536876 | 0.552298 | 20.82673 | 0.0009 | 0.027334 |
| ETV4 | 1.554973 | 3.680077 | 20.80422 | 0.000904 | 0.027356 |
| PRKDC | 1.405178 | 7.806665 | 20.79426 | 0.000905 | 0.02736 |
| LOC108589051 | 2.662874 | 3.028487 | 21.66951 | 0.000909 | 0.027426 |
| LRP8 | 1.842327 | 3.62792 | 20.72242 | 0.000917 | 0.027534 |
| LOC100413711 | 3.517581 | 1.604719 | 20.62496 | 0.000955 | 0.028556 |
| CPVL | 2.199704 | 3.262316 | 20.69151 | 0.000961 | 0.028586 |
| GALNT12 | 1.406917 | 5.504553 | 20.35661 | 0.000978 | 0.028834 |
| TNFSF15 | 2.103854 | 2.502422 | 20.34958 | 0.000979 | 0.028834 |
| KIF14 | 2.246763 | 2.968217 | 20.44957 | 0.000978 | 0.028834 |
| BUB1 | 2.267356 | 2.154295 | 20.34701 | 0.000979 | 0.028834 |
| NCR1 | 3.520071 | 1.409638 | 20.33376 | 0.000982 | 0.028858 |
| CXCL10 | 2.122779 | 1.757314 | 20.07533 | 0.001028 | 0.0299 |
| EML4 | 1.145521 | 6.306883 | 19.85837 | 0.001069 | 0.029915 |
| DLGAP5 | 1.38145 | 3.382933 | 19.99339 | 0.001043 | 0.029915 |
| LOC100388032 | 1.447474 | 3.17729 | 19.92083 | 0.001057 | 0.029915 |
| LOC100402677 | 2.437237 | 3.547075 | 21.06522 | 0.001043 | 0.029915 |
| LOC100393780 | 2.56549 | 2.040716 | 19.9773 | 0.001046 | 0.029915 |
| SLAMF6 | 2.765272 | 3.653755 | 21.9826 | 0.001067 | 0.029915 |
| CKAP4 | 1.306427 | 4.736781 | 19.82929 | 0.001075 | 0.030029 |
| ADGRG1 | 1.301238 | 4.382813 | 19.81081 | 0.001078 | 0.030038 |
| SQLE | 1.392075 | 5.76372 | 19.79109 | 0.001082 | 0.030038 |
| CFB | 1.403535 | 5.543312 | 19.77715 | 0.001085 | 0.030038 |
| FANCD2 | 1.769095 | 3.310708 | 19.78254 | 0.001084 | 0.030038 |

| | | | | | |
|---|---|---|---|---|---|
| LOC108589199 | 1.826808 | 5.084022 | 20.53404 | 0.001081 | 0.030038 |
| LOC100388085 | 1.892179 | 2.971153 | 19.70708 | 0.001099 | 0.030305 |
| CYTIP | 2.634305 | 4.269632 | 21.91003 | 0.001149 | 0.03095 |
| LOC100895417 | 3.01098 | 0.789104 | 19.52662 | 0.001135 | 0.03095 |
| LOC108589745 | 3.129574 | 1.168168 | 19.50424 | 0.00114 | 0.03095 |
| LOC108593080 | 4.38098 | 4.589876 | 27.68612 | 0.00115 | 0.03095 |
| LOC103796360 | 1.91973 | 2.924127 | 19.40809 | 0.00116 | 0.03107 |
| CCDC34 | 1.825628 | 2.120862 | 19.31558 | 0.00118 | 0.031558 |
| MYC | 1.576575 | 5.37901 | 19.45648 | 0.001192 | 0.0317 |
| SERF1A | 3.071475 | 0.867039 | 19.22316 | 0.0012 | 0.031777 |
| CXCL2 | 2.696351 | 3.058183 | 20.37435 | 0.001209 | 0.031934 |
| NME2 | 1.066846 | 5.743998 | 19.14455 | 0.001218 | 0.03203 |
| EPRS | 1.087951 | 6.433843 | 19.13828 | 0.001219 | 0.03203 |
| ACPP | 4.625426 | 0.620708 | 19.14906 | 0.001217 | 0.03203 |
| LOC100395429 | 1.313032 | 5.034799 | 19.12281 | 0.001223 | 0.032035 |
| SMC2 | 1.428678 | 4.054713 | 19.08723 | 0.001231 | 0.032123 |
| LOC100407326 | 1.187536 | 7.30963 | 19.04312 | 0.001241 | 0.032292 |
| CD24 | 2.013107 | 3.916882 | 19.51884 | 0.001254 | 0.032486 |
| SHTN1 | 1.367495 | 3.971042 | 18.97375 | 0.001257 | 0.032494 |
| VARS | 1.166459 | 6.425386 | 18.96599 | 0.001259 | 0.032495 |
| LOC100397481 | 1.498998 | 4.720924 | 18.94731 | 0.001263 | 0.032496 |
| LOC103791604 | 1.550899 | 2.775697 | 18.94438 | 0.001264 | 0.032496 |
| KLRC2 | 6.996013 | 5.039238 | 30.79741 | 0.001262 | 0.032496 |
| TNFAIP8 | 1.516981 | 4.793353 | 18.92535 | 0.001269 | 0.032525 |
| ARHGAP15 | 1.920975 | 4.326283 | 19.50926 | 0.001276 | 0.032617 |
| STIP1 | 1.065201 | 6.709516 | 18.88377 | 0.001279 | 0.032648 |
| LOC100386315 | 1.159588 | 6.800498 | 18.8088 | 0.001297 | 0.032804 |
| LOC100403703 | 1.492405 | 2.765146 | 18.84305 | 0.001288 | 0.032804 |
| ASPM | 1.583245 | 4.809872 | 18.90968 | 0.001293 | 0.032804 |
| TFCP2L1 | 2.124858 | 2.017568 | 18.81581 | 0.001295 | 0.032804 |
| LOC103790854 | 2.626156 | 2.469526 | 19.12371 | 0.001292 | 0.032804 |
| LOC108590058 | 1.280533 | 4.756243 | 18.73675 | 0.001314 | 0.033066 |
| CCNB1 | 2.37225 | 1.985729 | 18.72455 | 0.001317 | 0.033066 |
| FOXC1 | 2.886339 | 1.20113 | 18.70268 | 0.001323 | 0.033088 |
| PCSK9 | 2.36609 | 4.099131 | 20.29529 | 0.001332 | 0.033218 |
| TLR2 | 1.493581 | 2.594226 | 18.5854 | 0.001352 | 0.033551 |
| MRC1 | 1.651361 | 4.648271 | 18.71953 | 0.001357 | 0.033562 |
| NOLC1 | 1.361701 | 5.740632 | 18.53705 | 0.001365 | 0.033686 |
| FCRL5 | 1.644624 | 3.939038 | 18.48627 | 0.001378 | 0.033795 |
| LOC100411120 | 1.375596 | 6.495657 | 18.46813 | 0.001383 | 0.033809 |
| ICA1 | 1.464265 | 4.28832 | 18.44054 | 0.00139 | 0.033831 |
| PLXNC1 | 1.630382 | 4.561149 | 18.53317 | 0.001387 | 0.033831 |
| GALNT5 | 2.152535 | 2.131986 | 18.40542 | 0.001399 | 0.033971 |
| LOC100396448 | 1.120166 | 6.275311 | 18.37843 | 0.001406 | 0.03406 |
| XBP1 | 1.031272 | 6.227034 | 18.34937 | 0.001414 | 0.034206 |
| LOC100413674 | 1.084534 | 5.775585 | 18.29237 | 0.00143 | 0.034364 |
| LOC100409403 | 1.201767 | 7.787503 | 18.30912 | 0.001425 | 0.034364 |

| | | | | | |
|---|---|---|---|---|---|
| RDH11 | 1.242226 | 4.729279 | 18.28269 | 0.001432 | 0.034377 |
| LOC100390149 | 1.720938 | 2.698753 | 18.27478 | 0.001434 | 0.034377 |
| USP44 | 2.85178 | 2.347952 | 18.88358 | 0.001437 | 0.034377 |
| MS4A6A | 1.770903 | 3.091149 | 18.23877 | 0.001444 | 0.034383 |
| LOC100399654 | 1.110884 | 6.784278 | 18.2111 | 0.001452 | 0.034443 |
| LOC108590528 | 2.627671 | 3.692248 | 20.05842 | 0.00145 | 0.034443 |
| SHMT2 | 1.480861 | 4.629504 | 18.19461 | 0.001457 | 0.034445 |
| FBL | 1.119566 | 5.13127 | 18.17313 | 0.001463 | 0.034469 |
| LOC108589971 | 1.652024 | 2.952254 | 18.11037 | 0.00148 | 0.034674 |
| ALDH1L2 | 1.672523 | 4.83261 | 18.41857 | 0.001486 | 0.034772 |
| MANF | 1.124886 | 5.000069 | 18.05175 | 0.001497 | 0.034814 |
| E2F2 | 2.033464 | 4.175093 | 18.90392 | 0.001495 | 0.034814 |
| PIGR | 1.790004 | 10.94658 | 18.04541 | 0.001499 | 0.034814 |
| SNRPA1 | 1.059546 | 5.121509 | 18.02702 | 0.001504 | 0.03485 |
| WDR3 | 1.132447 | 5.345905 | 17.97226 | 0.00152 | 0.03485 |
| LOC100409427 | 1.477427 | 4.720219 | 17.97173 | 0.00152 | 0.03485 |
| NME1 | 1.540037 | 2.841279 | 18.01974 | 0.001506 | 0.03485 |
| NKRF | 1.608315 | 3.169737 | 17.98109 | 0.001518 | 0.03485 |
| LOC108593753 | 1.673721 | 3.426887 | 17.98563 | 0.001516 | 0.03485 |
| FLG2 | 2.861807 | 2.898113 | 19.43981 | 0.001512 | 0.03485 |
| THOP1 | 1.24501 | 4.754664 | 17.93449 | 0.001531 | 0.034894 |
| CFTR | 1.449889 | 6.86274 | 18.11891 | 0.00153 | 0.034894 |
| LOC100414401 | 1.474931 | 6.517827 | 18.1479 | 0.001531 | 0.034894 |
| CD84 | 1.964033 | 5.663841 | 19.43505 | 0.001563 | 0.035405 |
| LOC100412268 | 2.516391 | 3.98806 | 19.69603 | 0.001568 | 0.035465 |
| LMNB1 | 1.226695 | 5.770912 | 17.69313 | 0.001605 | 0.03603 |
| PSME1 | 1.365768 | 3.742439 | 17.70047 | 0.001603 | 0.03603 |
| LOC100401712 | 1.591029 | 3.355596 | 17.65881 | 0.001616 | 0.03609 |
| CARD16 | 1.835553 | 3.295741 | 17.62533 | 0.001626 | 0.03616 |
| DCTPP1 | 2.189986 | 2.179537 | 17.60219 | 0.001634 | 0.03624 |
| KCNN4 | 1.386351 | 4.037021 | 17.58863 | 0.001638 | 0.036253 |
| LOC108591201 | 1.228205 | 5.099273 | 17.56544 | 0.001646 | 0.036376 |
| CALR | 1.113682 | 9.422265 | 17.46641 | 0.001678 | 0.036438 |
| HMGCS1 | 1.184844 | 7.047126 | 17.52629 | 0.001658 | 0.036438 |
| LAT2 | 1.39051 | 3.661931 | 17.47442 | 0.001675 | 0.036438 |
| TNS4 | 1.471057 | 6.665396 | 17.75387 | 0.001658 | 0.036438 |
| EIF4EBP1 | 1.539304 | 3.913801 | 17.47776 | 0.001674 | 0.036438 |
| LOC100413337 | 1.983899 | 1.93723 | 17.46072 | 0.00168 | 0.036438 |
| SCARA5 | 2.177305 | 3.480202 | 18.15572 | 0.001662 | 0.036438 |
| ABRACL | 1.33673 | 3.886855 | 17.41537 | 0.001695 | 0.036625 |
| HMGA1 | 1.147309 | 6.690269 | 17.38776 | 0.001704 | 0.036784 |
| ACAT2 | 1.452184 | 3.790443 | 17.36063 | 0.001713 | 0.036801 |
| PLP2 | 1.093591 | 5.447458 | 17.33204 | 0.001723 | 0.03684 |
| LOC100414675 | 1.192997 | 5.742812 | 17.32934 | 0.001724 | 0.03684 |
| CKS2 | 2.307596 | 2.01932 | 17.3379 | 0.001721 | 0.03684 |
| KNSTRN | 2.334276 | 2.005209 | 17.34156 | 0.00172 | 0.03684 |
| CKAP2L | 1.779679 | 2.052867 | 17.29077 | 0.001737 | 0.037006 |

| | | | | | |
|---|---|---|---|---|---|
| PSAT1 | 2.101496 | 2.435198 | 17.29156 | 0.001737 | 0.037006 |
| LOC108588608 | 2.6307 | 2.038203 | 17.36399 | 0.001738 | 0.037006 |
| KRT7 | 2.234057 | 4.204789 | 18.64 | 0.001769 | 0.037495 |
| UGT8 | 2.330671 | 2.255916 | 17.13803 | 0.001791 | 0.037723 |
| LOC100895947 | 2.8063 | 1.32955 | 17.14339 | 0.001789 | 0.037723 |
| HES5 | 3.312041 | 0.725867 | 17.14152 | 0.001789 | 0.037723 |
| HYAL1 | 1.436249 | 2.998329 | 17.11002 | 0.001801 | 0.037736 |
| MLEC | 1.068235 | 8.480678 | 17.08833 | 0.001809 | 0.037851 |
| LOC100409880 | 1.070698 | 5.987672 | 17.06374 | 0.001818 | 0.037955 |
| LOC108589888 | 1.426068 | 6.223953 | 17.22992 | 0.001828 | 0.038129 |
| KCNE3 | 1.75241 | 3.045151 | 16.9932 | 0.001843 | 0.03837 |
| ANP32E | 1.354259 | 4.855269 | 16.94431 | 0.001862 | 0.038665 |
| DAPP1 | 1.491027 | 3.80386 | 16.92229 | 0.00187 | 0.038794 |
| FABP3 | 1.491209 | 3.940794 | 16.87986 | 0.001886 | 0.038917 |
| LOC100409607 | 1.482282 | 3.437615 | 16.83887 | 0.001901 | 0.03905 |
| KIF15 | 1.969701 | 3.370786 | 17.02143 | 0.001905 | 0.039052 |
| B4GALNT3 | 4.812193 | 2.947803 | 21.78586 | 0.00191 | 0.039123 |
| PCNA | 1.647789 | 3.728099 | 16.74955 | 0.001936 | 0.039262 |
| TDRD6 | 1.778586 | 2.496518 | 16.76398 | 0.00193 | 0.039262 |
| LOC100409955 | 1.325995 | 3.114207 | 16.71084 | 0.001951 | 0.039301 |
| SLC7A11 | 1.792649 | 2.21002 | 16.68579 | 0.001961 | 0.039418 |
| ADD3 | 1.093888 | 5.968031 | 16.67056 | 0.001967 | 0.039499 |
| LOC100414020 | 1.178355 | 5.514962 | 16.60997 | 0.001992 | 0.039905 |
| ACER2 | 1.509103 | 2.701429 | 16.56183 | 0.002011 | 0.040257 |
| POLR3GL | 1.101513 | 3.763957 | 16.52447 | 0.002027 | 0.04043 |
| PROKR2 | 1.851497 | 3.574578 | 16.70008 | 0.002031 | 0.04043 |
| KLF2 | 2.005341 | 4.281745 | 17.48935 | 0.002042 | 0.040515 |
| TNFRSF9 | 4.511883 | 0.530983 | 16.46743 | 0.002051 | 0.040515 |
| CLPB | 1.029735 | 4.544768 | 16.43928 | 0.002062 | 0.040618 |
| LOC103787291 | 1.066171 | 4.72058 | 16.42755 | 0.002067 | 0.040618 |
| LOC100409130 | 0.943297 | 6.873103 | 16.40628 | 0.002076 | 0.040712 |
| LOC100387472 | 0.980821 | 5.414426 | 16.35685 | 0.002098 | 0.040878 |
| NOP58 | 1.148328 | 5.367798 | 16.36162 | 0.002096 | 0.040878 |
| LOC100894398 | 1.441071 | 2.90822 | 16.31571 | 0.002115 | 0.040963 |
| TYROBP | 1.943225 | 3.684623 | 16.75584 | 0.002117 | 0.040963 |
| PA2G4 | 1.011919 | 5.539565 | 16.29547 | 0.002124 | 0.041024 |
| GADD45GIP1 | 1.133167 | 3.967807 | 16.26293 | 0.002139 | 0.041148 |
| OIP5 | 2.808096 | 1.060109 | 16.22773 | 0.002154 | 0.041311 |
| BRCA2 | 1.883492 | 3.767705 | 16.58823 | 0.002159 | 0.04137 |
| CD44 | 1.566395 | 7.138637 | 16.80289 | 0.002171 | 0.041439 |
| CCR8 | 2.629887 | 1.145381 | 16.12795 | 0.002199 | 0.041799 |
| IDO1 | 1.719489 | 4.002884 | 16.28125 | 0.00221 | 0.041881 |
| ECT2 | 1.557378 | 2.940277 | 16.07649 | 0.002223 | 0.041999 |
| LOC103789502 | 1.046859 | 8.441214 | 16.04863 | 0.002236 | 0.042202 |
| SNX5 | 1.01107 | 5.319219 | 16.03288 | 0.002243 | 0.042279 |
| LOC100389406 | 2.753241 | 0.64139 | 16.03046 | 0.002244 | 0.042279 |
| LOC100387896 | 1.228007 | 5.284298 | 15.99896 | 0.002259 | 0.042516 |

| | | | | | |
|---|---|---|---|---|---|
| LOC100404789 | 1.590797 | 6.09747 | 16.66737 | 0.002269 | 0.042659 |
| LOC108593890 | 1.848424 | 2.150207 | 15.92269 | 0.002295 | 0.042941 |
| LOC100414476 | 0.942737 | 5.330667 | 15.83155 | 0.00234 | 0.043319 |
| DEPDC5 | 1.344239 | 4.979497 | 15.83081 | 0.00234 | 0.043319 |
| DDAH1 | 0.929951 | 5.04614 | 15.81496 | 0.002348 | 0.043422 |
| MET | 1.226106 | 5.891851 | 15.78314 | 0.002363 | 0.043588 |
| FBLN1 | 1.446553 | 5.932659 | 16.05903 | 0.002376 | 0.043691 |
| KIF4A | 1.869559 | 2.803192 | 15.75116 | 0.002379 | 0.043691 |
| RNF24 | 1.100028 | 4.070667 | 15.70974 | 0.0024 | 0.043889 |
| CCDC112 | 1.286717 | 3.50409 | 15.70009 | 0.002405 | 0.043937 |
| LOC100406435 | 1.383076 | 6.104665 | 15.85419 | 0.002413 | 0.044008 |
| LOC100394108 | 2.001357 | 5.181215 | 17.14037 | 0.002443 | 0.044284 |
| DBF4 | 1.510936 | 2.772847 | 15.61709 | 0.002448 | 0.044322 |
| ORC1 | 1.887449 | 3.313517 | 15.78216 | 0.002449 | 0.044322 |
| LOC100896443 | 0.904236 | 5.596524 | 15.53916 | 0.002489 | 0.044491 |
| PDIA3 | 0.97595 | 8.202938 | 15.55208 | 0.002482 | 0.044491 |
| LOC100401056 | 1.169673 | 5.651031 | 15.5865 | 0.002464 | 0.044491 |
| PEX5L | 1.848255 | 4.034248 | 16.04755 | 0.002496 | 0.044491 |
| NOX1 | 2.571612 | 1.219195 | 15.5564 | 0.002479 | 0.044491 |
| SAMD3 | 4.055956 | 2.057644 | 17.47391 | 0.002466 | 0.044491 |
| SMC4 | 1.07883 | 5.781002 | 15.51467 | 0.002502 | 0.044515 |
| EPHB2 | 2.317331 | 5.255696 | 17.96528 | 0.002515 | 0.044555 |
| CEP55 | 2.353133 | 2.084733 | 15.44705 | 0.002538 | 0.044788 |
| HOPX | 1.840614 | 2.448911 | 15.44144 | 0.002541 | 0.044801 |
| MKI67 | 1.784718 | 7.479972 | 16.64167 | 0.002545 | 0.044841 |
| LOC100404972 | 1.742797 | 3.547518 | 15.46371 | 0.002568 | 0.045138 |
| LOC108592653 | 2.679016 | 2.119774 | 15.75814 | 0.002571 | 0.045138 |
| LOC103794276 | 3.303172 | 0.71678 | 15.36908 | 0.00258 | 0.045138 |
| KLRF1 | 2.639756 | 2.231304 | 15.7583 | 0.0026 | 0.045346 |
| SYNGR2 | 0.898451 | 5.584826 | 15.32472 | 0.002605 | 0.045393 |
| LOC100405988 | 4.036453 | 1.837691 | 16.75873 | 0.00261 | 0.045401 |
| LOC100385683 | 1.11048 | 5.668566 | 15.30568 | 0.002616 | 0.045414 |
| LOC100411388 | 1.728188 | 3.333113 | 15.30677 | 0.002615 | 0.045414 |
| UHRF1 | 2.000292 | 4.427166 | 16.45492 | 0.002621 | 0.045427 |
| LOC103791620 | 2.066147 | 1.403451 | 15.26288 | 0.00264 | 0.045628 |
| TRIM46 | 3.381021 | 0.741894 | 15.23085 | 0.002658 | 0.045822 |
| C4H6orf58 | 6.531583 | 4.92668 | 24.33325 | 0.00267 | 0.045979 |
| AMD1 | 1.032767 | 4.690635 | 15.16195 | 0.002698 | 0.046241 |
| NUP205 | 1.091447 | 5.685766 | 15.1595 | 0.002699 | 0.046241 |
| C12H16orf54 | 2.298458 | 3.686813 | 16.41606 | 0.002697 | 0.046241 |
| KLRD1 | 4.598889 | 4.670712 | 22.25233 | 0.002698 | 0.046241 |
| LOC100411071 | 0.96556 | 5.853255 | 15.12407 | 0.00272 | 0.046454 |
| GLIPR1 | 1.426132 | 3.643117 | 15.12169 | 0.002722 | 0.046454 |
| SORCS2 | 2.523254 | 4.358781 | 17.42894 | 0.002732 | 0.04654 |
| CD180 | 1.397655 | 3.793638 | 15.07977 | 0.002746 | 0.046662 |
| FTSJ1 | 1.568431 | 2.311491 | 15.07449 | 0.002749 | 0.046662 |
| KLHL32 | 1.572002 | 2.053406 | 15.07249 | 0.002751 | 0.046662 |

| | | | | | |
|---|---|---|---|---|---|
| CCDC18 | 1.235262 | 3.862858 | 15.03699 | 0.002772 | 0.046899 |
| WNT1 | 2.658361 | 1.308298 | 15.04027 | 0.00277 | 0.046899 |
| LOC100397769 | 0.896832 | 5.35837 | 14.97536 | 0.002809 | 0.047282 |
| BRIP1 | 1.992566 | 1.82577 | 14.91426 | 0.002847 | 0.047832 |
| NUSAP1 | 1.880262 | 2.571069 | 14.90661 | 0.002852 | 0.04787 |
| PARP8 | 1.099117 | 5.542143 | 14.89316 | 0.00286 | 0.047895 |
| TRAFD1 | 0.998991 | 6.654068 | 14.87926 | 0.002869 | 0.047948 |
| CRLF1 | 2.996076 | 0.505019 | 14.88015 | 0.002868 | 0.047948 |
| MCM6 | 2.3308 | 4.460813 | 16.83804 | 0.002881 | 0.048103 |
| PRIM1 | 1.603365 | 3.339379 | 14.81368 | 0.00291 | 0.048472 |
| SLC16A6 | 1.874971 | 2.727054 | 14.72293 | 0.002969 | 0.049158 |
| LOC100396470 | 2.987914 | 3.362363 | 16.98448 | 0.002971 | 0.049158 |
| ZBTB38 | 1.028801 | 6.696967 | 14.643 | 0.003022 | 0.049563 |
| SLFN5 | 1.431124 | 4.466525 | 14.65231 | 0.003035 | 0.049563 |
| FIGNL1 | 1.565183 | 3.549367 | 14.64086 | 0.003023 | 0.049563 |
| FCGR2B | 2.113251 | 2.357262 | 14.63275 | 0.003028 | 0.049563 |
| LOC108588225 | 3.031068 | 0.526374 | 14.57384 | 0.003068 | 0.049977 |

**Supplemental Table 6. GO sets differentially expressed genes in the duodenum between stricture and non-stricture**

**Supp. Table 6a. GO sets upregulated in stricture**

| GO ID | Term | Ont | N | Up | Down | P-value |
|---|---|---|---|---|---|---|
| GO:0031224 | intrinsic component of membrane | CC | 3410 | 97 | 236 | 2.83E-14 |
| GO:0071944 | cell periphery | CC | 3686 | 122 | 250 | 2.91E-14 |
| GO:0005886 | plasma membrane | CC | 3600 | 118 | 245 | 4.50E-14 |
| GO:0016021 | integral component of membrane | CC | 3303 | 95 | 226 | 5.68E-13 |
| GO:0010876 | lipid localization | BP | 292 | 14 | 44 | 1.39E-12 |
| GO:0005215 | transporter activity | MF | 782 | 20 | 79 | 7.34E-12 |
| GO:0006629 | lipid metabolic process | BP | 1097 | 27 | 99 | 1.02E-11 |
| GO:0044425 | membrane part | CC | 4478 | 124 | 278 | 2.63E-11 |
| GO:0006869 | lipid transport | BP | 262 | 13 | 39 | 3.98E-11 |
| GO:0016324 | apical plasma membrane | CC | 253 | 12 | 38 | 5.43E-11 |
| GO:0045177 | apical part of cell | CC | 309 | 13 | 42 | 1.34E-10 |
| GO:0046486 | glycerolipid metabolic process | BP | 349 | 10 | 45 | 1.70E-10 |
| GO:0044281 | small molecule metabolic process | BP | 1646 | 49 | 128 | 1.80E-10 |
| GO:0044255 | cellular lipid metabolic process | BP | 842 | 22 | 79 | 2.81E-10 |
| GO:0006639 | acylglycerol metabolic process | BP | 104 | 4 | 22 | 1.00E-09 |
| GO:0006638 | neutral lipid metabolic process | BP | 105 | 4 | 22 | 1.22E-09 |
| GO:0015711 | organic anion transport | BP | 364 | 16 | 44 | 2.16E-09 |
| GO:0006811 | ion transport | BP | 1196 | 39 | 98 | 2.79E-09 |
| GO:0006820 | anion transport | BP | 459 | 19 | 50 | 6.00E-09 |
| GO:0006082 | organic acid metabolic process | BP | 923 | 27 | 80 | 8.66E-09 |
| GO:0019752 | carboxylic acid metabolic process | BP | 845 | 27 | 75 | 9.78E-09 |
| GO:0010817 | regulation of hormone levels | BP | 377 | 10 | 43 | 1.90E-08 |
| GO:0050892 | intestinal absorption | BP | 35 | 0 | 12 | 2.04E-08 |
| GO:0043436 | oxoacid metabolic process | BP | 910 | 27 | 78 | 2.22E-08 |
| GO:0006641 | triglyceride metabolic process | BP | 84 | 4 | 18 | 2.72E-08 |
| GO:0022857 | transmembrane transporter activity | MF | 707 | 18 | 65 | 2.78E-08 |
| GO:0055088 | lipid homeostasis | BP | 115 | 4 | 21 | 3.80E-08 |
| GO:0015294 | solute:cation symporter activity | MF | 71 | 4 | 16 | 7.62E-08 |
| GO:0032787 | monocarboxylic acid metabolic process | BP | 499 | 15 | 50 | 9.03E-08 |
| GO:0044459 | plasma membrane part | CC | 2005 | 77 | 138 | 9.38E-08 |
| GO:0015908 | fatty acid transport | BP | 72 | 6 | 16 | 9.39E-08 |
| GO:0008610 | lipid biosynthetic process | BP | 580 | 18 | 55 | 1.28E-07 |
| GO:0007586 | digestion | BP | 103 | 7 | 19 | 1.43E-07 |
| GO:0044282 | small molecule catabolic process | BP | 362 | 10 | 40 | 1.43E-07 |
| GO:0017144 | drug metabolic process | BP | 613 | 28 | 57 | 1.45E-07 |
| GO:0008238 | exopeptidase activity | MF | 84 | 2 | 17 | 1.60E-07 |
| GO:0015293 | symporter activity | MF | 104 | 5 | 19 | 1.68E-07 |
| GO:0042445 | hormone metabolic process | BP | 159 | 3 | 24 | 1.82E-07 |
| GO:0046503 | glycerolipid catabolic process | BP | 50 | 2 | 13 | 2.13E-07 |
| GO:0008289 | lipid binding | MF | 567 | 19 | 53 | 3.47E-07 |
| GO:0043168 | anion binding | MF | 2228 | 70 | 147 | 4.36E-07 |
| GO:0016125 | sterol metabolic process | BP | 132 | 7 | 21 | 4.36E-07 |
| GO:0031226 | intrinsic component of plasma membrane | CC | 1128 | 44 | 86 | 7.38E-07 |
| GO:0008202 | steroid metabolic process | BP | 248 | 8 | 30 | 8.10E-07 |
| GO:0046942 | carboxylic acid transport | BP | 250 | 12 | 30 | 9.61E-07 |
| GO:0015849 | organic acid transport | BP | 250 | 12 | 30 | 9.61E-07 |
| GO:0015370 | solute:sodium symporter activity | MF | 48 | 3 | 12 | 9.95E-07 |
| GO:0019637 | organophosphate metabolic process | BP | 933 | 28 | 74 | 1.13E-06 |
| GO:0006810 | transport | BP | 3949 | 115 | 231 | 1.27E-06 |
| GO:0015909 | long-chain fatty acid transport | BP | 50 | 5 | 12 | 1.60E-06 |
| GO:0004177 | aminopeptidase activity | MF | 42 | 1 | 11 | 1.71E-06 |
| GO:0098856 | intestinal lipid absorption | BP | 15 | 0 | 7 | 1.72E-06 |
| GO:1901615 | organic hydroxy compound metabolic process | BP | 413 | 15 | 41 | 1.72E-06 |
| GO:0005319 | lipid transporter activity | MF | 99 | 2 | 17 | 1.84E-06 |
| GO:0008203 | cholesterol metabolic process | BP | 121 | 7 | 19 | 1.88E-06 |
| GO:0043171 | peptide catabolic process | BP | 21 | 1 | 8 | 1.97E-06 |
| GO:0044242 | cellular lipid catabolic process | BP | 181 | 4 | 24 | 2.03E-06 |
| GO:0051186 | cofactor metabolic process | BP | 492 | 17 | 46 | 2.12E-06 |
| GO:0005887 | integral component of plasma membrane | CC | 1072 | 43 | 81 | 2.31E-06 |
| GO:0097164 | ammonium ion metabolic process | BP | 159 | 9 | 22 | 2.65E-06 |

| | | | | | | |
|---|---|---|---|---|---|---|
| GO:0046394 | carboxylic acid biosynthetic process | BP | 363 | 17 | 37 | 3.01E-06 |
| GO:0098862 | cluster of actin-based cell projections | CC | 125 | 6 | 19 | 3.11E-06 |
| GO:1902652 | secondary alcohol metabolic process | BP | 125 | 7 | 19 | 3.11E-06 |
| GO:0016053 | organic acid biosynthetic process | BP | 364 | 17 | 37 | 3.21E-06 |
| GO:0016020 | membrane | CC | 6503 | 180 | 347 | 3.56E-06 |
| GO:0032000 | positive regulation of fatty acid beta-oxidation | BP | 7 | 0 | 5 | 3.58E-06 |
| GO:0044283 | small molecule biosynthetic process | BP | 661 | 28 | 56 | 3.64E-06 |
| GO:0015291 | secondary active transmembrane transporter activity | MF | 175 | 7 | 23 | 3.86E-06 |
| GO:0019216 | regulation of lipid metabolic process | BP | 338 | 11 | 35 | 3.91E-06 |
| GO:0051234 | establishment of localization | BP | 4049 | 122 | 233 | 4.16E-06 |
| GO:0015075 | ion transmembrane transporter activity | MF | 586 | 18 | 51 | 4.76E-06 |
| GO:0017127 | cholesterol transporter activity | MF | 17 | 0 | 7 | 4.80E-06 |
| GO:0120020 | intermembrane cholesterol transfer activity | MF | 17 | 0 | 7 | 4.80E-06 |
| GO:0050996 | positive regulation of lipid catabolic process | BP | 17 | 0 | 7 | 4.80E-06 |
| GO:0030258 | lipid modification | BP | 203 | 4 | 25 | 4.85E-06 |
| GO:0045017 | glycerolipid biosynthetic process | BP | 219 | 5 | 26 | 6.19E-06 |
| GO:0008324 | cation transmembrane transporter activity | MF | 422 | 12 | 40 | 7.16E-06 |
| GO:0006650 | glycerophospholipid metabolic process | BP | 276 | 8 | 30 | 7.40E-06 |
| GO:0120015 | intermembrane sterol transfer activity | MF | 18 | 0 | 7 | 7.55E-06 |
| GO:0006694 | steroid biosynthetic process | BP | 157 | 6 | 21 | 7.66E-06 |
| GO:0098590 | plasma membrane region | CC | 915 | 30 | 70 | 8.00E-06 |
| GO:0015748 | organophosphate ester transport | BP | 88 | 5 | 15 | 8.14E-06 |
| GO:0006631 | fatty acid metabolic process | BP | 292 | 9 | 31 | 8.45E-06 |
| GO:0022804 | active transmembrane transporter activity | MF | 264 | 9 | 29 | 8.55E-06 |
| GO:0055085 | transmembrane transport | BP | 1146 | 32 | 83 | 8.86E-06 |
| GO:0046470 | phosphatidylcholine metabolic process | BP | 68 | 6 | 13 | 8.93E-06 |
| GO:0046320 | regulation of fatty acid oxidation | BP | 25 | 1 | 8 | 8.94E-06 |
| GO:0006644 | phospholipid metabolic process | BP | 366 | 10 | 36 | 9.15E-06 |
| GO:0031667 | response to nutrient levels | BP | 396 | 18 | 38 | 9.17E-06 |
| GO:0005903 | brush border | CC | 89 | 4 | 15 | 9.39E-06 |
| GO:0008374 | O-acyltransferase activity | MF | 41 | 0 | 10 | 1.02E-05 |
| GO:0015318 | inorganic molecular entity transmembrane transporter activity | MF | 538 | 15 | 47 | 1.03E-05 |
| GO:0046464 | acylglycerol catabolic process | BP | 33 | 1 | 9 | 1.06E-05 |
| GO:0046461 | neutral lipid catabolic process | BP | 33 | 1 | 9 | 1.06E-05 |
| GO:0030299 | intestinal cholesterol absorption | BP | 13 | 0 | 6 | 1.07E-05 |
| GO:0032934 | sterol binding | MF | 50 | 0 | 11 | 1.07E-05 |
| GO:0008237 | metallopeptidase activity | MF | 136 | 2 | 19 | 1.10E-05 |
| GO:0006066 | alcohol metabolic process | BP | 296 | 10 | 31 | 1.11E-05 |
| GO:0005576 | extracellular region | CC | 2794 | 111 | 169 | 1.18E-05 |
| GO:0005543 | phospholipid binding | MF | 356 | 10 | 35 | 1.23E-05 |
| GO:0046395 | carboxylic acid catabolic process | BP | 230 | 9 | 26 | 1.50E-05 |
| GO:0016054 | organic acid catabolic process | BP | 230 | 9 | 26 | 1.50E-05 |
| GO:0071702 | organic substance transport | BP | 2233 | 55 | 140 | 1.51E-05 |
| GO:0022600 | digestive system process | BP | 82 | 6 | 14 | 1.59E-05 |
| GO:0050994 | regulation of lipid catabolic process | BP | 43 | 1 | 10 | 1.61E-05 |
| GO:0043178 | alcohol binding | MF | 72 | 0 | 13 | 1.71E-05 |
| GO:0044241 | lipid digestion | BP | 14 | 0 | 6 | 1.81E-05 |
| GO:0009991 | response to extracellular stimulus | BP | 424 | 20 | 39 | 1.87E-05 |
| GO:0044214 | spanning component of plasma membrane | CC | 5 | 0 | 4 | 1.98E-05 |
| GO:0019217 | regulation of fatty acid metabolic process | BP | 73 | 4 | 13 | 2.00E-05 |
| GO:0120013 | intermembrane lipid transfer activity | MF | 21 | 0 | 7 | 2.45E-05 |
| GO:0015918 | sterol transport | BP | 86 | 3 | 14 | 2.79E-05 |
| GO:0031998 | regulation of fatty acid beta-oxidation | BP | 15 | 0 | 6 | 2.90E-05 |
| GO:0015248 | sterol transporter activity | MF | 29 | 0 | 8 | 3.02E-05 |
| GO:0031526 | brush border membrane | CC | 46 | 2 | 10 | 3.02E-05 |
| GO:0046434 | organophosphate catabolic process | BP | 214 | 6 | 24 | 3.61E-05 |
| GO:0051179 | localization | BP | 5081 | 153 | 276 | 3.63E-05 |
| GO:0072329 | monocarboxylic acid catabolic process | BP | 111 | 1 | 16 | 3.63E-05 |
| GO:0033344 | cholesterol efflux | BP | 38 | 1 | 9 | 3.66E-05 |
| GO:0042180 | cellular ketone metabolic process | BP | 201 | 7 | 23 | 3.81E-05 |
| GO:0015144 | carbohydrate transmembrane transporter activity | MF | 30 | 0 | 8 | 3.96E-05 |
| GO:0016042 | lipid catabolic process | BP | 258 | 7 | 27 | 4.13E-05 |
| GO:0062012 | regulation of small molecule metabolic process | BP | 378 | 17 | 35 | 4.37E-05 |
| GO:0006140 | regulation of nucleotide metabolic process | BP | 113 | 5 | 16 | 4.54E-05 |

| | | | | | | |
|---|---|---|---|---|---|---|
| GO:0005496 | steroid binding | MF | 79 | 0 | 13 | 4.77E-05 |
| GO:0044421 | extracellular region part | CC | 2383 | 104 | 145 | 4.79E-05 |
| GO:0065008 | regulation of biological quality | BP | 3088 | 104 | 180 | 5.09E-05 |
| GO:0005996 | monosaccharide metabolic process | BP | 234 | 3 | 25 | 5.63E-05 |
| GO:0015718 | monocarboxylic acid transport | BP | 115 | 8 | 16 | 5.65E-05 |
| GO:0030300 | regulation of intestinal cholesterol absorption | BP | 6 | 0 | 4 | 5.72E-05 |
| GO:1904729 | regulation of intestinal lipid absorption | BP | 6 | 0 | 4 | 5.72E-05 |
| GO:0036094 | small molecule binding | MF | 2001 | 60 | 125 | 5.92E-05 |
| GO:0046321 | positive regulation of fatty acid oxidation | BP | 11 | 0 | 5 | 6.76E-05 |
| GO:1903587 | regulation of blood vessel endothelial cell proliferation involved in | BP | 11 | 1 | 5 | 6.76E-05 |
| GO:1905039 | carboxylic acid transmembrane transport | BP | 105 | 4 | 15 | 7.03E-05 |
| GO:1903825 | organic acid transmembrane transport | BP | 105 | 4 | 15 | 7.03E-05 |
| GO:0019915 | lipid storage | BP | 61 | 1 | 11 | 7.67E-05 |
| GO:0072330 | monocarboxylic acid biosynthetic process | BP | 268 | 11 | 27 | 7.97E-05 |
| GO:0022890 | inorganic cation transmembrane transporter activity | MF | 390 | 11 | 35 | 8.24E-05 |
| GO:0030808 | regulation of nucleotide biosynthetic process | BP | 95 | 2 | 14 | 8.64E-05 |
| GO:0019433 | triglyceride catabolic process | BP | 25 | 1 | 7 | 8.65E-05 |
| GO:0008410 | CoA-transferase activity | MF | 3 | 0 | 3 | 9.13E-05 |
| GO:0055092 | sterol homeostasis | BP | 73 | 4 | 12 | 9.37E-05 |
| GO:0006805 | xenobiotic metabolic process | BP | 73 | 1 | 12 | 9.37E-05 |
| GO:0005402 | carbohydrate:cation symporter activity | MF | 18 | 0 | 6 | 9.57E-05 |
| GO:0015145 | monosaccharide transmembrane transporter activity | MF | 18 | 0 | 6 | 9.57E-05 |
| GO:0044262 | cellular carbohydrate metabolic process | BP | 242 | 4 | 25 | 9.73E-05 |
| GO:0015485 | cholesterol binding | MF | 43 | 0 | 9 | 0.0001 |
| GO:0015914 | phospholipid transport | BP | 63 | 5 | 11 | 0.0001 |
| GO:0019432 | triglyceride biosynthetic process | BP | 34 | 1 | 8 | 0.0001 |
| GO:0090407 | organophosphate biosynthetic process | BP | 575 | 15 | 46 | 0.00011 |
| GO:0045923 | positive regulation of fatty acid metabolic process | BP | 26 | 2 | 7 | 0.00011 |
| GO:1904478 | regulation of intestinal absorption | BP | 7 | 0 | 4 | 0.00013 |
| GO:0062013 | positive regulation of small molecule metabolic process | BP | 112 | 4 | 15 | 0.00015 |
| GO:0008235 | metalloexopeptidase activity | MF | 45 | 1 | 9 | 0.00015 |
| GO:0046463 | acylglycerol biosynthetic process | BP | 36 | 1 | 8 | 0.00016 |
| GO:0046460 | neutral lipid biosynthetic process | BP | 36 | 1 | 8 | 0.00016 |
| GO:0010565 | regulation of cellular ketone metabolic process | BP | 152 | 6 | 18 | 0.00017 |
| GO:0046341 | CDP-diacylglycerol metabolic process | BP | 13 | 0 | 5 | 0.00017 |
| GO:0008514 | organic anion transmembrane transporter activity | MF | 153 | 5 | 18 | 0.00018 |
| GO:0016126 | sterol biosynthetic process | BP | 67 | 5 | 11 | 0.00018 |
| GO:0051119 | sugar transmembrane transporter activity | MF | 20 | 0 | 6 | 0.00018 |
| GO:0045834 | positive regulation of lipid metabolic process | BP | 115 | 5 | 15 | 0.0002 |
| GO:0005975 | carbohydrate metabolic process | BP | 507 | 10 | 41 | 0.00021 |
| GO:0005615 | extracellular space | CC | 2237 | 99 | 134 | 0.00021 |
| GO:0046873 | metal ion transmembrane transporter activity | MF | 315 | 10 | 29 | 0.00022 |
| GO:0016052 | carbohydrate catabolic process | BP | 169 | 4 | 19 | 0.00022 |
| GO:0048878 | chemical homeostasis | BP | 890 | 35 | 63 | 0.00022 |
| GO:0019318 | hexose metabolic process | BP | 212 | 3 | 22 | 0.00024 |
| GO:0042737 | drug catabolic process | BP | 92 | 3 | 13 | 0.00024 |
| GO:0034371 | chylomicron remodeling | BP | 8 | 0 | 4 | 0.00025 |
| GO:1905952 | regulation of lipid localization | BP | 105 | 5 | 14 | 0.00026 |
| GO:0009165 | nucleotide biosynthetic process | BP | 303 | 9 | 28 | 0.00026 |
| GO:0016805 | dipeptidase activity | MF | 14 | 0 | 5 | 0.00026 |
| GO:0051188 | cofactor biosynthetic process | BP | 288 | 12 | 27 | 0.00026 |
| GO:0007588 | excretion | BP | 39 | 2 | 8 | 0.00029 |
| GO:1900371 | regulation of purine nucleotide biosynthetic process | BP | 94 | 2 | 13 | 0.00029 |
| GO:0042446 | hormone biosynthetic process | BP | 60 | 1 | 10 | 0.00032 |
| GO:0007623 | circadian rhythm | BP | 160 | 5 | 18 | 0.00032 |
| GO:1901293 | nucleoside phosphate biosynthetic process | BP | 307 | 9 | 28 | 0.00032 |
| GO:0034754 | cellular hormone metabolic process | BP | 83 | 1 | 12 | 0.00033 |
| GO:0051196 | regulation of coenzyme metabolic process | BP | 83 | 2 | 12 | 0.00033 |
| GO:0034377 | plasma lipoprotein particle assembly | BP | 22 | 1 | 6 | 0.00033 |
| GO:0015171 | amino acid transmembrane transporter activity | MF | 50 | 3 | 9 | 0.00035 |
| GO:0034363 | intermediate-density lipoprotein particle | CC | 4 | 0 | 3 | 0.00035 |
| GO:0042632 | cholesterol homeostasis | BP | 72 | 4 | 11 | 0.00035 |
| GO:0015077 | monovalent inorganic cation transmembrane transporter activity | MF | 233 | 9 | 23 | 0.00036 |
| GO:0043167 | ion binding | MF | 4756 | 112 | 254 | 0.00036 |

| | | | | | | |
|---|---|---|---|---|---|---|
| GO:0046943 | carboxylic acid transmembrane transporter activity | MF | 109 | 4 | 14 | 0.00038 |
| GO:0005342 | organic acid transmembrane transporter activity | MF | 109 | 4 | 14 | 0.00038 |
| GO:1900542 | regulation of purine nucleotide metabolic process | BP | 109 | 5 | 14 | 0.00038 |
| GO:0046474 | glycerophospholipid biosynthetic process | BP | 191 | 4 | 20 | 0.0004 |
| GO:0001523 | retinoid metabolic process | BP | 73 | 2 | 11 | 0.0004 |
| GO:0046890 | regulation of lipid biosynthetic process | BP | 163 | 8 | 18 | 0.0004 |
| GO:0006721 | terpenoid metabolic process | BP | 85 | 3 | 12 | 0.00041 |
| GO:0006695 | cholesterol biosynthetic process | BP | 62 | 5 | 10 | 0.00042 |
| GO:1902653 | secondary alcohol biosynthetic process | BP | 62 | 5 | 10 | 0.00042 |
| GO:0008509 | anion transmembrane transporter activity | MF | 221 | 7 | 22 | 0.00042 |
| GO:0009649 | entrainment of circadian clock | BP | 23 | 0 | 6 | 0.00043 |
| GO:0015175 | neutral amino acid transmembrane transporter activity | MF | 23 | 2 | 6 | 0.00043 |
| GO:0034378 | chylomicron assembly | BP | 9 | 0 | 4 | 0.00043 |
| GO:0005984 | disaccharide metabolic process | BP | 9 | 0 | 4 | 0.00043 |
| GO:0089717 | spanning component of membrane | CC | 9 | 0 | 4 | 0.00043 |
| GO:0031406 | carboxylic acid binding | MF | 137 | 7 | 16 | 0.00045 |
| GO:0006732 | coenzyme metabolic process | BP | 345 | 12 | 30 | 0.00045 |
| GO:0019842 | vitamin binding | MF | 99 | 2 | 13 | 0.00049 |
| GO:0030301 | cholesterol transport | BP | 75 | 3 | 11 | 0.00051 |
| GO:0005355 | glucose transmembrane transporter activity | MF | 16 | 0 | 5 | 0.00053 |
| GO:0015149 | hexose transmembrane transporter activity | MF | 16 | 0 | 5 | 0.00053 |
| GO:0034308 | primary alcohol metabolic process | BP | 64 | 1 | 10 | 0.00054 |
| GO:0006654 | phosphatidic acid biosynthetic process | BP | 33 | 1 | 7 | 0.00056 |
| GO:0043177 | organic acid binding | MF | 140 | 8 | 16 | 0.00057 |
| GO:0019218 | regulation of steroid metabolic process | BP | 101 | 4 | 13 | 0.0006 |
| GO:1901605 | alpha-amino acid metabolic process | BP | 183 | 10 | 19 | 0.00061 |
| GO:0072524 | pyridine-containing compound metabolic process | BP | 169 | 6 | 18 | 0.00062 |
| GO:0032922 | circadian regulation of gene expression | BP | 54 | 2 | 9 | 0.00063 |
| GO:0016101 | diterpenoid metabolic process | BP | 77 | 2 | 11 | 0.00064 |
| GO:0006090 | pyruvate metabolic process | BP | 128 | 3 | 15 | 0.00064 |
| GO:0006720 | isoprenoid metabolic process | BP | 102 | 3 | 13 | 0.00065 |
| GO:0008643 | carbohydrate transport | BP | 115 | 2 | 14 | 0.00066 |
| GO:0046473 | phosphatidic acid metabolic process | BP | 34 | 1 | 7 | 0.00068 |
| GO:0055081 | anion homeostasis | BP | 44 | 5 | 8 | 0.00069 |
| GO:0045161 | neuronal ion channel clustering | BP | 10 | 0 | 4 | 0.00069 |
| GO:0034379 | very-low-density lipoprotein particle assembly | BP | 10 | 1 | 4 | 0.00069 |
| GO:0065005 | protein-lipid complex assembly | BP | 25 | 1 | 6 | 0.0007 |
| GO:0015850 | organic hydroxy compound transport | BP | 185 | 4 | 19 | 0.0007 |
| GO:0072525 | pyridine-containing compound biosynthetic process | BP | 129 | 5 | 15 | 0.0007 |
| GO:0031253 | cell projection membrane | CC | 261 | 7 | 24 | 0.00075 |
| GO:0006701 | progesterone biosynthetic process | BP | 5 | 0 | 3 | 0.00085 |
| GO:0007168 | receptor guanylyl cyclase signaling pathway | BP | 5 | 0 | 3 | 0.00085 |
| GO:0032309 | icosanoid secretion | BP | 26 | 4 | 6 | 0.00087 |
| GO:0034220 | ion transmembrane transport | BP | 830 | 26 | 57 | 0.00094 |
| GO:0015081 | sodium ion transmembrane transporter activity | MF | 106 | 4 | 13 | 0.00095 |
| GO:0002043 | blood vessel endothelial cell proliferation involved in sprouting ang | BP | 18 | 1 | 5 | 0.00096 |
| GO:0034364 | high-density lipoprotein particle | CC | 18 | 2 | 5 | 0.00096 |
| GO:0070006 | metalloaminopeptidase activity | MF | 18 | 1 | 5 | 0.00096 |
| GO:0071827 | plasma lipoprotein particle organization | BP | 36 | 3 | 7 | 0.00098 |
| GO:0055086 | nucleobase-containing small molecule metabolic process | BP | 618 | 22 | 45 | 0.00101 |
| GO:0042627 | chylomicron | CC | 11 | 1 | 4 | 0.00105 |
| GO:0031982 | vesicle | CC | 2976 | 105 | 166 | 0.00105 |
| GO:0097367 | carbohydrate derivative binding | MF | 1767 | 58 | 106 | 0.00107 |
| GO:0042562 | hormone binding | MF | 70 | 1 | 10 | 0.00111 |
| GO:0009062 | fatty acid catabolic process | BP | 95 | 1 | 12 | 0.00113 |
| GO:0009914 | hormone transport | BP | 238 | 7 | 22 | 0.00114 |
| GO:0006006 | glucose metabolic process | BP | 178 | 3 | 18 | 0.00114 |
| GO:0006656 | phosphatidylcholine biosynthetic process | BP | 37 | 3 | 7 | 0.00116 |
| GO:0098656 | anion transmembrane transport | BP | 208 | 7 | 20 | 0.00117 |
| GO:0043153 | entrainment of circadian clock by photoperiod | BP | 19 | 0 | 5 | 0.00126 |
| GO:0045821 | positive regulation of glycolytic process | BP | 19 | 1 | 5 | 0.00126 |
| GO:0033218 | amide binding | MF | 240 | 13 | 22 | 0.00127 |
| GO:0006766 | vitamin metabolic process | BP | 110 | 3 | 13 | 0.00134 |
| GO:0042304 | regulation of fatty acid biosynthetic process | BP | 38 | 3 | 7 | 0.00137 |

| | | | | | | |
|---|---|---|---|---|---|---|
| GO:0006520 | cellular amino acid metabolic process | BP | 289 | 11 | 25 | 0.0014 |
| GO:0034381 | plasma lipoprotein particle clearance | BP | 49 | 2 | 8 | 0.00143 |
| GO:0009056 | catabolic process | BP | 2163 | 49 | 125 | 0.00149 |
| GO:0032488 | Cdc42 protein signal transduction | BP | 12 | 1 | 4 | 0.00152 |
| GO:0016024 | CDP-diacylglycerol biosynthetic process | BP | 12 | 0 | 4 | 0.00152 |
| GO:0055089 | fatty acid homeostasis | BP | 12 | 1 | 4 | 0.00152 |
| GO:0034384 | high-density lipoprotein particle clearance | BP | 12 | 0 | 4 | 0.00152 |
| GO:0034370 | triglyceride-rich lipoprotein particle remodeling | BP | 12 | 0 | 4 | 0.00152 |
| GO:0097006 | regulation of plasma lipoprotein particle levels | BP | 73 | 4 | 10 | 0.00155 |
| GO:0071825 | protein-lipid complex subunit organization | BP | 39 | 3 | 7 | 0.0016 |
| GO:0023061 | signal release | BP | 357 | 10 | 29 | 0.00161 |
| GO:0016411 | acylglycerol O-acyltransferase activity | MF | 20 | 0 | 5 | 0.00162 |
| GO:0051197 | positive regulation of coenzyme metabolic process | BP | 20 | 1 | 5 | 0.00162 |
| GO:0030813 | positive regulation of nucleotide catabolic process | BP | 20 | 1 | 5 | 0.00162 |
| GO:0048511 | rhythmic process | BP | 229 | 6 | 21 | 0.00162 |
| GO:0019359 | nicotinamide nucleotide biosynthetic process | BP | 126 | 4 | 14 | 0.00162 |
| GO:0019363 | pyridine nucleotide biosynthetic process | BP | 126 | 4 | 14 | 0.00162 |
| GO:0043467 | regulation of generation of precursor metabolites and energy | BP | 126 | 6 | 14 | 0.00162 |
| GO:0015749 | monosaccharide transmembrane transport | BP | 86 | 2 | 11 | 0.00162 |
| GO:0034366 | spherical high-density lipoprotein particle | CC | 6 | 0 | 3 | 0.00165 |
| GO:0006753 | nucleoside phosphate metabolic process | BP | 545 | 19 | 40 | 0.00165 |
| GO:0031669 | cellular response to nutrient levels | BP | 199 | 10 | 19 | 0.00167 |
| GO:0008654 | phospholipid biosynthetic process | BP | 230 | 5 | 21 | 0.00171 |
| GO:0003333 | amino acid transmembrane transport | BP | 62 | 4 | 9 | 0.00174 |
| GO:0005768 | endosome | CC | 743 | 17 | 51 | 0.00175 |
| GO:0016787 | hydrolase activity | MF | 1952 | 45 | 114 | 0.00175 |
| GO:0019395 | fatty acid oxidation | BP | 87 | 2 | 11 | 0.00178 |
| GO:0090181 | regulation of cholesterol metabolic process | BP | 51 | 3 | 8 | 0.00187 |
| GO:0009108 | coenzyme biosynthetic process | BP | 232 | 7 | 21 | 0.0019 |
| GO:0046879 | hormone secretion | BP | 232 | 7 | 21 | 0.0019 |
| GO:0055090 | acylglycerol homeostasis | BP | 30 | 1 | 6 | 0.00193 |
| GO:1901571 | fatty acid derivative transport | BP | 30 | 4 | 6 | 0.00193 |
| GO:0071715 | icosanoid transport | BP | 30 | 4 | 6 | 0.00193 |
| GO:0070328 | triglyceride homeostasis | BP | 30 | 1 | 6 | 0.00193 |
| GO:0034219 | carbohydrate transmembrane transport | BP | 88 | 2 | 11 | 0.00196 |
| GO:0034440 | lipid oxidation | BP | 88 | 3 | 11 | 0.00196 |
| GO:1901568 | fatty acid derivative metabolic process | BP | 115 | 6 | 13 | 0.00201 |
| GO:0046361 | 2-oxobutyrate metabolic process | BP | 2 | 0 | 2 | 0.00203 |
| GO:0030881 | beta-2-microglobulin binding | MF | 2 | 0 | 2 | 0.00203 |
| GO:0001300 | chronological cell aging | BP | 2 | 0 | 2 | 0.00203 |
| GO:0097037 | heme export | BP | 2 | 0 | 2 | 0.00203 |
| GO:0090675 | intermicrovillar adhesion | BP | 2 | 0 | 2 | 0.00203 |
| GO:0060752 | intestinal phytosterol absorption | BP | 2 | 0 | 2 | 0.00203 |
| GO:0080030 | methyl indole-3-acetate esterase activity | MF | 2 | 0 | 2 | 0.00203 |
| GO:1904479 | negative regulation of intestinal absorption | BP | 2 | 0 | 2 | 0.00203 |
| GO:0045796 | negative regulation of intestinal cholesterol absorption | BP | 2 | 0 | 2 | 0.00203 |
| GO:1904730 | negative regulation of intestinal lipid absorption | BP | 2 | 0 | 2 | 0.00203 |
| GO:0010949 | negative regulation of intestinal phytosterol absorption | BP | 2 | 0 | 2 | 0.00203 |
| GO:0060168 | positive regulation of adenosine receptor signaling pathway | BP | 2 | 0 | 2 | 0.00203 |
| GO:0015680 | protein maturation by copper ion transfer | BP | 2 | 0 | 2 | 0.00203 |
| GO:0060931 | sinoatrial node cell development | BP | 2 | 0 | 2 | 0.00203 |
| GO:0060921 | sinoatrial node cell differentiation | BP | 2 | 0 | 2 | 0.00203 |
| GO:0003163 | sinoatrial node development | BP | 2 | 0 | 2 | 0.00203 |
| GO:0009648 | photoperiodism | BP | 21 | 0 | 5 | 0.00205 |
| GO:0051193 | regulation of cofactor metabolic process | BP | 102 | 4 | 12 | 0.0021 |
| GO:0046459 | short-chain fatty acid metabolic process | BP | 13 | 0 | 4 | 0.00211 |
| GO:0046165 | alcohol biosynthetic process | BP | 144 | 7 | 15 | 0.00214 |
| GO:0051181 | cofactor transport | BP | 41 | 3 | 7 | 0.00216 |
| GO:0009117 | nucleotide metabolic process | BP | 538 | 19 | 39 | 0.00234 |
| GO:0009135 | purine nucleoside diphosphate metabolic process | BP | 117 | 3 | 13 | 0.00234 |
| GO:0009179 | purine ribonucleoside diphosphate metabolic process | BP | 117 | 3 | 13 | 0.00234 |
| GO:1901292 | nucleoside phosphate catabolic process | BP | 160 | 5 | 16 | 0.00236 |
| GO:0070011 | peptidase activity, acting on L-amino acid peptides | MF | 417 | 11 | 32 | 0.00236 |
| GO:0051187 | cofactor catabolic process | BP | 42 | 2 | 7 | 0.0025 |

| | | | | | | |
|---|---|---|---|---|---|---|
| GO:0006633 | fatty acid biosynthetic process | BP | 118 | 5 | 13 | 0.00253 |
| GO:0031528 | microvillus membrane | CC | 22 | 1 | 5 | 0.00255 |
| GO:0052689 | carboxylic ester hydrolase activity | MF | 91 | 3 | 11 | 0.00256 |
| GO:0008233 | peptidase activity | MF | 437 | 11 | 33 | 0.00263 |
| GO:0071466 | cellular response to xenobiotic stimulus | BP | 119 | 3 | 13 | 0.00273 |
| GO:0009185 | ribonucleoside diphosphate metabolic process | BP | 119 | 3 | 13 | 0.00273 |
| GO:0034383 | low-density lipoprotein particle clearance | BP | 32 | 1 | 6 | 0.00273 |
| GO:0017046 | peptide hormone binding | MF | 32 | 0 | 6 | 0.00273 |
| GO:0030810 | positive regulation of nucleotide biosynthetic process | BP | 32 | 1 | 6 | 0.00273 |
| GO:1900373 | positive regulation of purine nucleotide biosynthetic process | BP | 32 | 1 | 6 | 0.00273 |
| GO:0031012 | extracellular matrix | CC | 404 | 13 | 31 | 0.00275 |
| GO:0019203 | carbohydrate phosphatase activity | MF | 7 | 0 | 3 | 0.00279 |
| GO:0005412 | glucose:sodium symporter activity | MF | 7 | 0 | 3 | 0.00279 |
| GO:0032275 | luteinizing hormone secretion | BP | 7 | 0 | 3 | 0.00279 |
| GO:1903589 | positive regulation of blood vessel endothelial cell proliferation inv | BP | 7 | 1 | 3 | 0.00279 |
| GO:0072530 | purine-containing compound transmembrane transport | BP | 7 | 1 | 3 | 0.00279 |
| GO:0043654 | recognition of apoptotic cell | BP | 7 | 0 | 3 | 0.00279 |
| GO:0050308 | sugar-phosphatase activity | MF | 7 | 0 | 3 | 0.00279 |
| GO:0098657 | import into cell | BP | 687 | 30 | 47 | 0.0028 |
| GO:0046496 | nicotinamide nucleotide metabolic process | BP | 163 | 5 | 16 | 0.00285 |
| GO:0019362 | pyridine nucleotide metabolic process | BP | 163 | 5 | 16 | 0.00285 |
| GO:0001991 | regulation of systemic arterial blood pressure by circulatory renin-a | BP | 14 | 0 | 4 | 0.00286 |
| GO:0030246 | carbohydrate binding | MF | 179 | 15 | 17 | 0.00302 |
| GO:0050886 | endocrine process | BP | 55 | 1 | 8 | 0.00306 |
| GO:2001169 | regulation of ATP biosynthetic process | BP | 80 | 2 | 10 | 0.00309 |
| GO:1900543 | negative regulation of purine nucleotide metabolic process | BP | 23 | 1 | 5 | 0.00314 |
| GO:0034369 | plasma lipoprotein particle remodeling | BP | 23 | 3 | 5 | 0.00314 |
| GO:0034368 | protein-lipid complex remodeling | BP | 23 | 3 | 5 | 0.00314 |
| GO:0006110 | regulation of glycolytic process | BP | 68 | 2 | 9 | 0.00333 |
| GO:0097746 | regulation of blood vessel diameter | BP | 108 | 1 | 12 | 0.0034 |
| GO:0050880 | regulation of blood vessel size | BP | 108 | 1 | 12 | 0.0034 |
| GO:0035296 | regulation of tube diameter | BP | 108 | 1 | 12 | 0.0034 |
| GO:0035150 | regulation of tube size | BP | 108 | 1 | 12 | 0.0034 |
| GO:0006164 | purine nucleotide biosynthetic process | BP | 229 | 8 | 20 | 0.00364 |
| GO:0030811 | regulation of nucleotide catabolic process | BP | 69 | 2 | 9 | 0.00368 |
| GO:0030809 | negative regulation of nucleotide biosynthetic process | BP | 15 | 0 | 4 | 0.00376 |
| GO:1900372 | negative regulation of purine nucleotide biosynthetic process | BP | 15 | 0 | 4 | 0.00376 |
| GO:0046337 | phosphatidylethanolamine metabolic process | BP | 15 | 0 | 4 | 0.00376 |
| GO:0090208 | positive regulation of triglyceride metabolic process | BP | 15 | 0 | 4 | 0.00376 |
| GO:0043691 | reverse cholesterol transport | BP | 15 | 0 | 4 | 0.00376 |
| GO:0034385 | triglyceride-rich plasma lipoprotein particle | CC | 15 | 1 | 4 | 0.00376 |
| GO:0034361 | very-low-density lipoprotein particle | CC | 15 | 1 | 4 | 0.00376 |
| GO:0045980 | negative regulation of nucleotide metabolic process | BP | 24 | 1 | 5 | 0.00382 |
| GO:0016486 | peptide hormone processing | BP | 24 | 0 | 5 | 0.00382 |
| GO:0034367 | protein-containing complex remodeling | BP | 24 | 3 | 5 | 0.00382 |
| GO:0007205 | protein kinase C-activating G protein-coupled receptor signaling pa | BP | 24 | 0 | 5 | 0.00382 |
| GO:0006575 | cellular modified amino acid metabolic process | BP | 153 | 6 | 15 | 0.00385 |
| GO:1903578 | regulation of ATP metabolic process | BP | 96 | 5 | 11 | 0.0039 |
| GO:0042578 | phosphoric ester hydrolase activity | MF | 312 | 2 | 25 | 0.00392 |
| GO:0044248 | cellular catabolic process | BP | 1940 | 45 | 111 | 0.00399 |
| GO:1901617 | organic hydroxy compound biosynthetic process | BP | 216 | 10 | 19 | 0.00419 |
| GO:0009064 | glutamine family amino acid metabolic process | BP | 58 | 5 | 8 | 0.00428 |
| GO:0052646 | alditol phosphate metabolic process | BP | 8 | 0 | 3 | 0.00431 |
| GO:0032274 | gonadotropin secretion | BP | 8 | 0 | 3 | 0.00431 |
| GO:0010896 | regulation of triglyceride catabolic process | BP | 8 | 0 | 3 | 0.00431 |
| GO:0009750 | response to fructose | BP | 8 | 1 | 3 | 0.00431 |
| GO:0008216 | spermidine metabolic process | BP | 8 | 1 | 3 | 0.00431 |
| GO:0046415 | urate metabolic process | BP | 8 | 0 | 3 | 0.00431 |
| GO:0097084 | vascular smooth muscle cell development | BP | 8 | 0 | 3 | 0.00431 |
| GO:1905953 | negative regulation of lipid localization | BP | 35 | 2 | 6 | 0.00436 |
| GO:0009395 | phospholipid catabolic process | BP | 35 | 1 | 6 | 0.00436 |
| GO:0008645 | hexose transmembrane transport | BP | 84 | 2 | 10 | 0.00442 |
| GO:1901681 | sulfur compound binding | MF | 186 | 8 | 17 | 0.00448 |
| GO:0042221 | response to chemical | BP | 3265 | 110 | 175 | 0.00453 |

| | | | | | | |
|---|---|---|---|---|---|---|
| GO:0006757 | ATP generation from ADP | BP | 98 | 3 | 11 | 0.00457 |
| GO:0006096 | glycolytic process | BP | 98 | 3 | 11 | 0.00457 |
| GO:0010906 | regulation of glucose metabolic process | BP | 98 | 2 | 11 | 0.00457 |
| GO:0051194 | positive regulation of cofactor metabolic process | BP | 25 | 2 | 5 | 0.0046 |
| GO:0032094 | response to food | BP | 25 | 1 | 5 | 0.0046 |
| GO:0072521 | purine-containing compound metabolic process | BP | 454 | 16 | 33 | 0.00473 |
| GO:1902001 | fatty acid transmembrane transport | BP | 16 | 0 | 4 | 0.00483 |
| GO:0015101 | organic cation transmembrane transporter activity | MF | 16 | 0 | 4 | 0.00483 |
| GO:0001871 | pattern binding | MF | 16 | 0 | 4 | 0.00483 |
| GO:0004623 | phospholipase A2 activity | MF | 16 | 3 | 4 | 0.00483 |
| GO:0030247 | polysaccharide binding | MF | 16 | 0 | 4 | 0.00483 |
| GO:0042866 | pyruvate biosynthetic process | BP | 99 | 3 | 11 | 0.00494 |
| GO:0006898 | receptor-mediated endocytosis | BP | 204 | 10 | 18 | 0.00507 |
| GO:0048037 | cofactor binding | MF | 387 | 11 | 29 | 0.00517 |
| GO:0005539 | glycosaminoglycan binding | MF | 158 | 10 | 15 | 0.00519 |
| GO:0042594 | response to starvation | BP | 158 | 6 | 15 | 0.00519 |
| GO:0006754 | ATP biosynthetic process | BP | 114 | 3 | 12 | 0.00528 |
| GO:0006094 | gluconeogenesis | BP | 73 | 0 | 9 | 0.00537 |
| GO:0005902 | microvillus | CC | 73 | 3 | 9 | 0.00537 |
| GO:0010039 | response to iron ion | BP | 26 | 2 | 5 | 0.00549 |
| GO:0043230 | extracellular organelle | CC | 1679 | 70 | 97 | 0.00551 |
| GO:0042752 | regulation of circadian rhythm | BP | 87 | 3 | 10 | 0.00568 |
| GO:0030574 | collagen catabolic process | BP | 37 | 1 | 6 | 0.00579 |
| GO:0006073 | cellular glucan metabolic process | BP | 61 | 1 | 8 | 0.00584 |
| GO:0044042 | glucan metabolic process | BP | 61 | 1 | 8 | 0.00584 |
| GO:0003824 | catalytic activity | MF | 4476 | 101 | 231 | 0.00589 |
| GO:0060926 | cardiac pacemaker cell development | BP | 3 | 0 | 2 | 0.00591 |
| GO:0060920 | cardiac pacemaker cell differentiation | BP | 3 | 0 | 2 | 0.00591 |
| GO:0072338 | cellular lactam metabolic process | BP | 3 | 0 | 2 | 0.00591 |
| GO:0097167 | circadian regulation of translation | BP | 3 | 0 | 2 | 0.00591 |
| GO:0045163 | clustering of voltage-gated potassium channels | BP | 3 | 0 | 2 | 0.00591 |
| GO:0006824 | cobalt ion transport | BP | 3 | 0 | 2 | 0.00591 |
| GO:0002074 | extraocular skeletal muscle development | BP | 3 | 0 | 2 | 0.00591 |
| GO:0097629 | extrinsic component of omegasome membrane | CC | 3 | 0 | 2 | 0.00591 |
| GO:0036374 | glutathione hydrolase activity | MF | 3 | 0 | 2 | 0.00591 |
| GO:0004366 | glycerol-3-phosphate O-acyltransferase activity | MF | 3 | 0 | 2 | 0.00591 |
| GO:0070653 | high-density lipoprotein particle receptor binding | MF | 3 | 0 | 2 | 0.00591 |
| GO:0102953 | hypoglycin A gamma-glutamyl transpeptidase activity | MF | 3 | 0 | 2 | 0.00591 |
| GO:0103068 | leukotriene C4 gamma-glutamyl transferase activity | MF | 3 | 0 | 2 | 0.00591 |
| GO:1901750 | leukotriene D4 biosynthetic process | BP | 3 | 0 | 2 | 0.00591 |
| GO:1901748 | leukotriene D4 metabolic process | BP | 3 | 0 | 2 | 0.00591 |
| GO:0070573 | metallodipeptidase activity | MF | 3 | 0 | 2 | 0.00591 |
| GO:0034164 | negative regulation of toll-like receptor 9 signaling pathway | BP | 3 | 0 | 2 | 0.00591 |
| GO:0010903 | negative regulation of very-low-density lipoprotein particle remode | BP | 3 | 0 | 2 | 0.00591 |
| GO:0005415 | nucleoside:sodium symporter activity | MF | 3 | 1 | 2 | 0.00591 |
| GO:0000048 | peptidyltransferase activity | MF | 3 | 0 | 2 | 0.00591 |
| GO:0042822 | pyridoxal phosphate metabolic process | BP | 3 | 0 | 2 | 0.00591 |
| GO:0008321 | Ral guanyl-nucleotide exchange factor activity | MF | 3 | 0 | 2 | 0.00591 |
| GO:0060167 | regulation of adenosine receptor signaling pathway | BP | 3 | 0 | 2 | 0.00591 |
| GO:0010901 | regulation of very-low-density lipoprotein particle remodeling | BP | 3 | 0 | 2 | 0.00591 |
| GO:0102420 | sn-1-glycerol-3-phosphate C16:0-DCA-CoA acyl transferase activi | MF | 3 | 0 | 2 | 0.00591 |
| GO:0019229 | regulation of vasoconstriction | BP | 49 | 0 | 7 | 0.00606 |
| GO:1903963 | arachidonate transport | BP | 17 | 3 | 4 | 0.0061 |
| GO:0050482 | arachidonic acid secretion | BP | 17 | 3 | 4 | 0.0061 |
| GO:0034375 | high-density lipoprotein particle remodeling | BP | 17 | 1 | 4 | 0.0061 |
| GO:0006691 | leukotriene metabolic process | BP | 17 | 1 | 4 | 0.0061 |
| GO:0031065 | positive regulation of histone deacetylation | BP | 17 | 0 | 4 | 0.0061 |
| GO:0016746 | transferase activity, transferring acyl groups | MF | 208 | 5 | 18 | 0.00619 |
| GO:0003018 | vascular process in circulatory system | BP | 131 | 2 | 13 | 0.00621 |
| GO:0002003 | angiotensin maturation | BP | 9 | 0 | 3 | 0.00625 |
| GO:0005184 | neuropeptide hormone activity | MF | 9 | 0 | 3 | 0.00625 |
| GO:0008242 | omega peptidase activity | MF | 9 | 0 | 3 | 0.00625 |
| GO:0033700 | phospholipid efflux | BP | 9 | 1 | 3 | 0.00625 |
| GO:0022841 | potassium ion leak channel activity | MF | 9 | 0 | 3 | 0.00625 |

| | | | | | | |
|---|---|---|---|---|---|---|
| GO:0042448 | progesterone metabolic process | BP | 9 | 0 | 3 | 0.00625 |
| GO:0002002 | regulation of angiotensin levels in blood | BP | 9 | 0 | 3 | 0.00625 |
| GO:0046624 | sphingolipid transporter activity | MF | 9 | 0 | 3 | 0.00625 |
| GO:0005789 | endoplasmic reticulum membrane | CC | 849 | 20 | 54 | 0.00639 |
| GO:0072522 | purine-containing compound biosynthetic process | BP | 241 | 10 | 20 | 0.00641 |
| GO:0042310 | vasoconstriction | BP | 62 | 0 | 8 | 0.00645 |
| GO:0006812 | cation transport | BP | 812 | 25 | 52 | 0.00649 |
| GO:1990777 | lipoprotein particle | CC | 27 | 3 | 5 | 0.00649 |
| GO:0034358 | plasma lipoprotein particle | CC | 27 | 3 | 5 | 0.00649 |
| GO:0090276 | regulation of peptide hormone secretion | BP | 162 | 4 | 15 | 0.00653 |
| GO:1902475 | L-alpha-amino acid transmembrane transport | BP | 38 | 3 | 6 | 0.00661 |
| GO:0003044 | regulation of systemic arterial blood pressure mediated by a chemic | BP | 38 | 1 | 6 | 0.00661 |
| GO:0031668 | cellular response to extracellular stimulus | BP | 226 | 12 | 19 | 0.00678 |
| GO:0042277 | peptide binding | MF | 194 | 13 | 17 | 0.0068 |
| GO:0030072 | peptide hormone secretion | BP | 194 | 5 | 17 | 0.0068 |
| GO:0098827 | endoplasmic reticulum subcompartment | CC | 853 | 20 | 54 | 0.007 |
| GO:0019319 | hexose biosynthetic process | BP | 76 | 0 | 9 | 0.007 |
| GO:0043470 | regulation of carbohydrate catabolic process | BP | 76 | 2 | 9 | 0.007 |
| GO:0009132 | nucleoside diphosphate metabolic process | BP | 133 | 5 | 13 | 0.00704 |
| GO:0008217 | regulation of blood pressure | BP | 133 | 5 | 13 | 0.00704 |
| GO:0009605 | response to external stimulus | BP | 1735 | 74 | 99 | 0.00717 |
| GO:1903561 | extracellular vesicle | CC | 1677 | 69 | 96 | 0.00743 |
| GO:0015932 | nucleobase-containing compound transmembrane transporter activi | MF | 39 | 1 | 6 | 0.00753 |
| GO:0045981 | positive regulation of nucleotide metabolic process | BP | 39 | 2 | 6 | 0.00753 |
| GO:1900544 | positive regulation of purine nucleotide metabolic process | BP | 39 | 2 | 6 | 0.00753 |
| GO:0010884 | positive regulation of lipid storage | BP | 18 | 0 | 4 | 0.00756 |
| GO:0015295 | solute:proton symporter activity | MF | 18 | 0 | 4 | 0.00756 |
| GO:2001171 | positive regulation of ATP biosynthetic process | BP | 28 | 1 | 5 | 0.00762 |
| GO:0051180 | vitamin transport | BP | 28 | 0 | 5 | 0.00762 |
| GO:0030073 | insulin secretion | BP | 165 | 5 | 15 | 0.00769 |
| GO:0006733 | oxidoreduction coenzyme metabolic process | BP | 181 | 5 | 16 | 0.00784 |
| GO:0070062 | extracellular exosome | CC | 1661 | 69 | 95 | 0.00794 |
| GO:0006796 | phosphate-containing compound metabolic process | BP | 2620 | 76 | 142 | 0.00795 |
| GO:0007186 | G protein-coupled receptor signaling pathway | BP | 561 | 20 | 38 | 0.00804 |
| GO:0009986 | cell surface | CC | 598 | 35 | 40 | 0.00813 |
| GO:0016773 | phosphotransferase activity, alcohol group as acceptor | MF | 598 | 11 | 40 | 0.00813 |
| GO:0006793 | phosphorus metabolic process | BP | 2643 | 78 | 143 | 0.0082 |
| GO:0046883 | regulation of hormone secretion | BP | 198 | 6 | 17 | 0.00828 |
| GO:0098739 | import across plasma membrane | BP | 78 | 5 | 9 | 0.00828 |
| GO:0045540 | regulation of cholesterol biosynthetic process | BP | 40 | 3 | 6 | 0.00852 |
| GO:0010883 | regulation of lipid storage | BP | 40 | 0 | 6 | 0.00852 |
| GO:0106118 | regulation of sterol biosynthetic process | BP | 40 | 3 | 6 | 0.00852 |
| GO:1901264 | carbohydrate derivative transport | BP | 65 | 2 | 8 | 0.00856 |
| GO:0097756 | negative regulation of blood vessel diameter | BP | 65 | 0 | 8 | 0.00856 |
| GO:0046889 | positive regulation of lipid biosynthetic process | BP | 65 | 4 | 8 | 0.00856 |
| GO:0071501 | cellular response to sterol depletion | BP | 10 | 0 | 3 | 0.00864 |
| GO:0070508 | cholesterol import | BP | 10 | 0 | 3 | 0.00864 |
| GO:0019370 | leukotriene biosynthetic process | BP | 10 | 0 | 3 | 0.00864 |
| GO:0005324 | long-chain fatty acid transporter activity | MF | 10 | 1 | 3 | 0.00864 |
| GO:0032536 | regulation of cell projection size | BP | 10 | 0 | 3 | 0.00864 |
| GO:0030322 | stabilization of membrane potential | BP | 10 | 1 | 3 | 0.00864 |
| GO:0035376 | sterol import | BP | 10 | 0 | 3 | 0.00864 |
| GO:0016755 | transferase activity, transferring amino-acyl groups | MF | 10 | 0 | 3 | 0.00864 |
| GO:0034372 | very-low-density lipoprotein particle remodeling | BP | 10 | 0 | 3 | 0.00864 |
| GO:0050662 | coenzyme binding | MF | 248 | 6 | 20 | 0.00869 |
| GO:0046031 | ADP metabolic process | BP | 107 | 3 | 11 | 0.0088 |
| GO:0097009 | energy homeostasis | BP | 29 | 0 | 5 | 0.00887 |
| GO:0015804 | neutral amino acid transport | BP | 29 | 1 | 5 | 0.00887 |
| GO:0032994 | protein-lipid complex | CC | 29 | 3 | 5 | 0.00887 |
| GO:0050810 | regulation of steroid biosynthetic process | BP | 79 | 3 | 9 | 0.00899 |
| GO:0046906 | tetrapyrrole binding | MF | 79 | 4 | 9 | 0.00899 |
| GO:0007266 | Rho protein signal transduction | BP | 184 | 6 | 16 | 0.00912 |
| GO:0051049 | regulation of transport | BP | 1430 | 45 | 83 | 0.00912 |
| GO:0016405 | CoA-ligase activity | MF | 19 | 1 | 4 | 0.00924 |

| | | | | | | |
|---|---|---|---|---|---|---|
| GO:0051371 | muscle alpha-actinin binding | MF | 19 | 0 | 4 | 0.00924 |
| GO:1902930 | regulation of alcohol biosynthetic process | BP | 66 | 4 | 8 | 0.00937 |
| GO:0003073 | regulation of systemic arterial blood pressure | BP | 66 | 2 | 8 | 0.00937 |
| GO:0009152 | purine ribonucleotide biosynthetic process | BP | 217 | 8 | 18 | 0.00945 |
| GO:0009206 | purine ribonucleoside triphosphate biosynthetic process | BP | 123 | 5 | 12 | 0.00955 |
| GO:0042175 | nuclear outer membrane-endoplasmic reticulum membrane network | CC | 867 | 21 | 54 | 0.00956 |
| GO:0022838 | substrate-specific channel activity | MF | 267 | 5 | 21 | 0.00956 |
| GO:0032879 | regulation of localization | BP | 2160 | 67 | 119 | 0.0096 |
| GO:0032374 | regulation of cholesterol transport | BP | 41 | 2 | 6 | 0.00961 |
| GO:0032371 | regulation of sterol transport | BP | 41 | 2 | 6 | 0.00961 |
| GO:0016298 | lipase activity | MF | 94 | 4 | 10 | 0.0097 |
| GO:0032368 | regulation of lipid transport | BP | 80 | 5 | 9 | 0.00973 |
| GO:0008144 | drug binding | MF | 1376 | 46 | 80 | 0.01003 |
| GO:0031225 | anchored component of membrane | CC | 109 | 2 | 11 | 0.01006 |
| GO:0042592 | homeostatic process | BP | 1476 | 60 | 85 | 0.01013 |
| GO:0009145 | purine nucleoside triphosphate biosynthetic process | BP | 124 | 5 | 12 | 0.01016 |
| GO:0044432 | endoplasmic reticulum part | CC | 1082 | 28 | 65 | 0.01016 |
| GO:0008652 | cellular amino acid biosynthetic process | BP | 67 | 7 | 8 | 0.01023 |
| GO:0072341 | modified amino acid binding | MF | 67 | 2 | 8 | 0.01023 |
| GO:0051393 | alpha-actinin binding | MF | 30 | 0 | 5 | 0.01026 |
| GO:0004180 | carboxypeptidase activity | MF | 30 | 1 | 5 | 0.01026 |
| GO:0044058 | regulation of digestive system process | BP | 30 | 2 | 5 | 0.01026 |
| GO:0001990 | regulation of systemic arterial blood pressure by hormone | BP | 30 | 1 | 5 | 0.01026 |
| GO:0090207 | regulation of triglyceride metabolic process | BP | 30 | 0 | 5 | 0.01026 |
| GO:0003008 | system process | BP | 1201 | 28 | 71 | 0.0105 |
| GO:1904659 | glucose transmembrane transport | BP | 81 | 2 | 9 | 0.01053 |
| GO:0062014 | negative regulation of small molecule metabolic process | BP | 81 | 3 | 9 | 0.01053 |
| GO:0000287 | magnesium ion binding | MF | 187 | 5 | 16 | 0.01056 |
| GO:0006576 | cellular biogenic amine metabolic process | BP | 42 | 2 | 6 | 0.0108 |
| GO:0044275 | cellular carbohydrate catabolic process | BP | 42 | 0 | 6 | 0.0108 |
| GO:0031907 | microbody lumen | CC | 42 | 0 | 6 | 0.0108 |
| GO:0005782 | peroxisomal matrix | CC | 42 | 0 | 6 | 0.0108 |
| GO:0008015 | blood circulation | BP | 392 | 9 | 28 | 0.01105 |
| GO:0034637 | cellular carbohydrate biosynthetic process | BP | 68 | 0 | 8 | 0.01115 |
| GO:0009068 | aspartate family amino acid catabolic process | BP | 20 | 0 | 4 | 0.01115 |
| GO:0051184 | cofactor transmembrane transporter activity | MF | 20 | 0 | 4 | 0.01115 |
| GO:0046475 | glycerophospholipid catabolic process | BP | 20 | 1 | 4 | 0.01115 |
| GO:0003081 | regulation of systemic arterial blood pressure by renin-angiotensin | BP | 20 | 1 | 4 | 0.01115 |
| GO:0043648 | dicarboxylic acid metabolic process | BP | 82 | 4 | 9 | 0.01137 |
| GO:0046364 | monosaccharide biosynthetic process | BP | 82 | 0 | 9 | 0.01137 |
| GO:0006091 | generation of precursor metabolites and energy | BP | 393 | 11 | 28 | 0.01141 |
| GO:0046683 | response to organophosphorus | BP | 111 | 6 | 11 | 0.01145 |
| GO:0071077 | adenosine 3',5'-bisphosphate transmembrane transporter activity | MF | 4 | 0 | 2 | 0.01147 |
| GO:0004046 | aminoacylase activity | MF | 4 | 0 | 2 | 0.01147 |
| GO:0010815 | bradykinin catabolic process | BP | 4 | 0 | 2 | 0.01147 |
| GO:0004144 | diacylglycerol O-acyltransferase activity | MF | 4 | 0 | 2 | 0.01147 |
| GO:0097176 | epoxide metabolic process | BP | 4 | 0 | 2 | 0.01147 |
| GO:0086045 | membrane depolarization during AV node cell action potential | BP | 4 | 0 | 2 | 0.01147 |
| GO:1903588 | negative regulation of blood vessel endothelial cell proliferation inv | BP | 4 | 0 | 2 | 0.01147 |
| GO:2001186 | negative regulation of CD8-positive, alpha-beta T cell activation | BP | 4 | 0 | 2 | 0.01147 |
| GO:1903349 | omegasome membrane | CC | 4 | 0 | 2 | 0.01147 |
| GO:0044245 | polysaccharide digestion | BP | 4 | 1 | 2 | 0.01147 |
| GO:1900086 | positive regulation of peptidyl-tyrosine autophosphorylation | BP | 4 | 0 | 2 | 0.01147 |
| GO:0010898 | positive regulation of triglyceride catabolic process | BP | 4 | 0 | 2 | 0.01147 |
| GO:0019541 | propionate metabolic process | BP | 4 | 0 | 2 | 0.01147 |
| GO:0015252 | proton channel activity | MF | 4 | 0 | 2 | 0.01147 |
| GO:2000182 | regulation of progesterone biosynthetic process | BP | 4 | 0 | 2 | 0.01147 |
| GO:0051790 | short-chain fatty acid biosynthetic process | BP | 4 | 0 | 2 | 0.01147 |
| GO:0071918 | urea transmembrane transport | BP | 4 | 0 | 2 | 0.01147 |
| GO:0015204 | urea transmembrane transporter activity | MF | 4 | 0 | 2 | 0.01147 |
| GO:0015840 | urea transport | BP | 4 | 0 | 2 | 0.01147 |
| GO:0042760 | very long-chain fatty acid catabolic process | BP | 4 | 0 | 2 | 0.01147 |
| GO:0000295 | adenine nucleotide transmembrane transporter activity | MF | 11 | 0 | 3 | 0.01148 |
| GO:0006853 | carnitine shuttle | BP | 11 | 0 | 3 | 0.01148 |

| | | | | | | |
|---|---|---|---|---|---|---|
| GO:0008239 | dipeptidyl-peptidase activity | MF | 11 | 0 | 3 | 0.01148 |
| GO:0022840 | leak channel activity | MF | 11 | 0 | 3 | 0.01148 |
| GO:0022842 | narrow pore channel activity | MF | 11 | 0 | 3 | 0.01148 |
| GO:0032375 | negative regulation of cholesterol transport | BP | 11 | 2 | 3 | 0.01148 |
| GO:0032372 | negative regulation of sterol transport | BP | 11 | 2 | 3 | 0.01148 |
| GO:0038007 | netrin-activated signaling pathway | BP | 11 | 1 | 3 | 0.01148 |
| GO:0005346 | purine ribonucleotide transmembrane transporter activity | MF | 11 | 0 | 3 | 0.01148 |
| GO:0006991 | response to sterol depletion | BP | 11 | 0 | 3 | 0.01148 |
| GO:0006814 | sodium ion transport | BP | 173 | 6 | 15 | 0.01164 |
| GO:0090066 | regulation of anatomical structure size | BP | 412 | 12 | 29 | 0.01185 |
| GO:0070405 | ammonium ion binding | MF | 43 | 3 | 6 | 0.01209 |
| GO:0033293 | monocarboxylic acid binding | MF | 43 | 3 | 6 | 0.01209 |
| GO:0005548 | phospholipid transporter activity | MF | 43 | 1 | 6 | 0.01209 |
| GO:0009267 | cellular response to starvation | BP | 127 | 5 | 12 | 0.01215 |
| GO:0016788 | hydrolase activity, acting on ester bonds | MF | 595 | 9 | 39 | 0.01216 |
| GO:0006165 | nucleoside diphosphate phosphorylation | BP | 112 | 5 | 11 | 0.0122 |
| GO:0030001 | metal ion transport | BP | 651 | 20 | 42 | 0.01221 |
| GO:0050796 | regulation of insulin secretion | BP | 143 | 4 | 13 | 0.01258 |
| GO:0016791 | phosphatase activity | MF | 224 | 1 | 18 | 0.01283 |
| GO:0016575 | histone deacetylation | BP | 70 | 0 | 8 | 0.01317 |
| GO:0006700 | C21-steroid hormone biosynthetic process | BP | 21 | 0 | 4 | 0.01329 |
| GO:0090312 | positive regulation of protein deacetylation | BP | 21 | 0 | 4 | 0.01329 |
| GO:0006109 | regulation of carbohydrate metabolic process | BP | 176 | 5 | 15 | 0.01347 |
| GO:0009201 | ribonucleoside triphosphate biosynthetic process | BP | 129 | 5 | 12 | 0.01364 |
| GO:0046939 | nucleotide phosphorylation | BP | 114 | 5 | 11 | 0.01381 |
| GO:0009743 | response to carbohydrate | BP | 193 | 5 | 16 | 0.01397 |
| GO:0007589 | body fluid secretion | BP | 71 | 6 | 8 | 0.01428 |
| GO:0044070 | regulation of anion transport | BP | 71 | 5 | 8 | 0.01428 |
| GO:0042133 | neurotransmitter metabolic process | BP | 115 | 8 | 11 | 0.01468 |
| GO:0051503 | adenine nucleotide transport | BP | 12 | 0 | 3 | 0.01481 |
| GO:0016208 | AMP binding | MF | 12 | 0 | 3 | 0.01481 |
| GO:0050667 | homocysteine metabolic process | BP | 12 | 1 | 3 | 0.01481 |
| GO:0017110 | nucleoside-diphosphatase activity | MF | 12 | 0 | 3 | 0.01481 |
| GO:0006646 | phosphatidylethanolamine biosynthetic process | BP | 12 | 0 | 3 | 0.01481 |
| GO:0015216 | purine nucleotide transmembrane transporter activity | MF | 12 | 0 | 3 | 0.01481 |
| GO:0030730 | sequestering of triglyceride | BP | 12 | 0 | 3 | 0.01481 |
| GO:0046834 | lipid phosphorylation | BP | 58 | 0 | 7 | 0.01501 |
| GO:0003013 | circulatory system process | BP | 402 | 9 | 28 | 0.01508 |
| GO:1901575 | organic substance catabolic process | BP | 1804 | 44 | 100 | 0.01509 |
| GO:0034404 | nucleobase-containing small molecule biosynthetic process | BP | 195 | 8 | 16 | 0.01529 |
| GO:0020037 | heme binding | MF | 72 | 4 | 8 | 0.01545 |
| GO:1903579 | negative regulation of ATP metabolic process | BP | 22 | 1 | 4 | 0.01568 |
| GO:0032369 | negative regulation of lipid transport | BP | 22 | 2 | 4 | 0.01568 |
| GO:0009260 | ribonucleotide biosynthetic process | BP | 229 | 8 | 18 | 0.01578 |
| GO:0015672 | monovalent inorganic cation transport | BP | 351 | 13 | 25 | 0.01629 |
| GO:0005179 | hormone activity | MF | 46 | 1 | 6 | 0.01659 |
| GO:0030018 | Z disc | CC | 102 | 1 | 10 | 0.01663 |
| GO:0006690 | icosanoid metabolic process | BP | 73 | 3 | 8 | 0.01669 |
| GO:0015267 | channel activity | MF | 282 | 7 | 21 | 0.01689 |
| GO:0006163 | purine nucleotide metabolic process | BP | 424 | 15 | 29 | 0.01695 |
| GO:0046903 | secretion | BP | 1288 | 59 | 74 | 0.01697 |
| GO:0006775 | fat-soluble vitamin metabolic process | BP | 34 | 0 | 5 | 0.01729 |
| GO:0022803 | passive transmembrane transporter activity | MF | 283 | 7 | 21 | 0.01751 |
| GO:0010675 | regulation of cellular carbohydrate metabolic process | BP | 118 | 3 | 11 | 0.01751 |
| GO:0016301 | kinase activity | MF | 648 | 12 | 41 | 0.01772 |
| GO:0005977 | glycogen metabolic process | BP | 60 | 1 | 7 | 0.01785 |
| GO:0009127 | purine nucleoside monophosphate biosynthetic process | BP | 134 | 3 | 12 | 0.01797 |
| GO:0009168 | purine ribonucleoside monophosphate biosynthetic process | BP | 134 | 3 | 12 | 0.01797 |
| GO:0006767 | water-soluble vitamin metabolic process | BP | 74 | 3 | 8 | 0.018 |
| GO:0009066 | aspartate family amino acid metabolic process | BP | 47 | 1 | 6 | 0.01831 |
| GO:0015807 | L-amino acid transport | BP | 47 | 3 | 6 | 0.01831 |
| GO:0050661 | NADP binding | MF | 47 | 2 | 6 | 0.01831 |
| GO:0001786 | phosphatidylserine binding | MF | 47 | 1 | 6 | 0.01831 |
| GO:0033013 | tetrapyrrole metabolic process | BP | 47 | 0 | 6 | 0.01831 |

| | | | | | | |
|---|---|---|---|---|---|---|
| GO:0016878 | acid-thiol ligase activity | MF | 23 | 1 | 4 | 0.01832 |
| GO:0045940 | positive regulation of steroid metabolic process | BP | 23 | 1 | 4 | 0.01832 |
| GO:0003846 | 2-acylglycerol O-acyltransferase activity | MF | 5 | 0 | 2 | 0.01855 |
| GO:0036155 | acylglycerol acyl-chain remodeling | BP | 5 | 0 | 2 | 0.01855 |
| GO:0033564 | anterior/posterior axon guidance | BP | 5 | 0 | 2 | 0.01855 |
| GO:1904970 | brush border assembly | BP | 5 | 0 | 2 | 0.01855 |
| GO:0072501 | cellular divalent inorganic anion homeostasis | BP | 5 | 0 | 2 | 0.01855 |
| GO:0030643 | cellular phosphate ion homeostasis | BP | 5 | 0 | 2 | 0.01855 |
| GO:0071332 | cellular response to fructose stimulus | BP | 5 | 1 | 2 | 0.01855 |
| GO:0072502 | cellular trivalent inorganic anion homeostasis | BP | 5 | 0 | 2 | 0.01855 |
| GO:0006182 | cGMP biosynthetic process | BP | 5 | 0 | 2 | 0.01855 |
| GO:0005375 | copper ion transmembrane transporter activity | MF | 5 | 0 | 2 | 0.01855 |
| GO:0048788 | cytoskeleton of presynaptic active zone | CC | 5 | 0 | 2 | 0.01855 |
| GO:0042420 | dopamine catabolic process | BP | 5 | 0 | 2 | 0.01855 |
| GO:0006751 | glutathione catabolic process | BP | 5 | 0 | 2 | 0.01855 |
| GO:0006114 | glycerol biosynthetic process | BP | 5 | 0 | 2 | 0.01855 |
| GO:0043559 | insulin binding | MF | 5 | 0 | 2 | 0.01855 |
| GO:0030314 | junctional membrane complex | CC | 5 | 0 | 2 | 0.01855 |
| GO:0072674 | multinuclear osteoclast differentiation | BP | 5 | 0 | 2 | 0.01855 |
| GO:0090027 | negative regulation of monocyte chemotaxis | BP | 5 | 0 | 2 | 0.01855 |
| GO:0010891 | negative regulation of sequestering of triglyceride | BP | 5 | 0 | 2 | 0.01855 |
| GO:1990462 | omegasome | CC | 5 | 0 | 2 | 0.01855 |
| GO:0019800 | peptide cross-linking via chondroitin 4-sulfate glycosaminoglycan | BP | 5 | 0 | 2 | 0.01855 |
| GO:0060228 | phosphatidylcholine-sterol O-acyltransferase activator activity | MF | 5 | 1 | 2 | 0.01855 |
| GO:0071205 | protein localization to juxtaparanode region of axon | BP | 5 | 0 | 2 | 0.01855 |
| GO:0031282 | regulation of guanylate cyclase activity | BP | 5 | 0 | 2 | 0.01855 |
| GO:0033684 | regulation of luteinizing hormone secretion | BP | 5 | 0 | 2 | 0.01855 |
| GO:1900084 | regulation of peptidyl-tyrosine autophosphorylation | BP | 5 | 0 | 2 | 0.01855 |
| GO:0006566 | threonine metabolic process | BP | 5 | 0 | 2 | 0.01855 |
| GO:0042816 | vitamin B6 metabolic process | BP | 5 | 1 | 2 | 0.01855 |
| GO:0009109 | coenzyme catabolic process | BP | 13 | 1 | 3 | 0.01861 |
| GO:0006000 | fructose metabolic process | BP | 13 | 0 | 3 | 0.01861 |
| GO:0042447 | hormone catabolic process | BP | 13 | 0 | 3 | 0.01861 |
| GO:0006910 | phagocytosis, recognition | BP | 13 | 0 | 3 | 0.01861 |
| GO:0015868 | purine ribonucleotide transport | BP | 13 | 0 | 3 | 0.01861 |
| GO:0060841 | venous blood vessel development | BP | 13 | 0 | 3 | 0.01861 |
| GO:0032963 | collagen metabolic process | BP | 89 | 3 | 9 | 0.01872 |
| GO:0050770 | regulation of axonogenesis | BP | 151 | 5 | 13 | 0.01905 |
| GO:1903580 | positive regulation of ATP metabolic process | BP | 35 | 2 | 5 | 0.01944 |
| GO:0046390 | ribose phosphate biosynthetic process | BP | 235 | 8 | 18 | 0.01999 |
| GO:0006897 | endocytosis | BP | 598 | 25 | 38 | 0.02062 |
| GO:0050801 | ion homeostasis | BP | 598 | 26 | 38 | 0.02062 |
| GO:0048029 | monosaccharide binding | MF | 62 | 2 | 7 | 0.02106 |
| GO:1905954 | positive regulation of lipid localization | BP | 62 | 3 | 7 | 0.02106 |
| GO:0009110 | vitamin biosynthetic process | BP | 24 | 1 | 4 | 0.02122 |
| GO:0042995 | cell projection | CC | 1666 | 37 | 92 | 0.02147 |
| GO:0071496 | cellular response to external stimulus | BP | 289 | 14 | 21 | 0.02156 |
| GO:0007595 | lactation | BP | 36 | 3 | 5 | 0.02177 |
| GO:0008307 | structural constituent of muscle | MF | 36 | 1 | 5 | 0.02177 |
| GO:0051056 | regulation of small GTPase mediated signal transduction | BP | 307 | 7 | 22 | 0.02185 |
| GO:0009166 | nucleotide catabolic process | BP | 154 | 5 | 13 | 0.02204 |
| GO:0009142 | nucleoside triphosphate biosynthetic process | BP | 138 | 5 | 12 | 0.02212 |
| GO:1901616 | organic hydroxy compound catabolic process | BP | 49 | 0 | 6 | 0.02212 |
| GO:0009897 | external side of plasma membrane | CC | 221 | 17 | 17 | 0.02247 |
| GO:0015085 | calcium ion transmembrane transporter activity | MF | 107 | 1 | 10 | 0.02251 |
| GO:0005262 | calcium channel activity | MF | 92 | 0 | 9 | 0.02275 |
| GO:0005509 | calcium ion binding | MF | 490 | 15 | 32 | 0.02282 |
| GO:0048185 | activin binding | MF | 14 | 0 | 3 | 0.02291 |
| GO:0034435 | cholesterol esterification | BP | 14 | 1 | 3 | 0.02291 |
| GO:0006067 | ethanol metabolic process | BP | 14 | 0 | 3 | 0.02291 |
| GO:2001170 | negative regulation of ATP biosynthetic process | BP | 14 | 0 | 3 | 0.02291 |
| GO:0002689 | negative regulation of leukocyte chemotaxis | BP | 14 | 0 | 3 | 0.02291 |
| GO:0015865 | purine nucleotide transport | BP | 14 | 0 | 3 | 0.02291 |
| GO:0034433 | steroid esterification | BP | 14 | 1 | 3 | 0.02291 |

| | | | | | | |
|---|---|---|---|---|---|---|
| GO:0034434 | sterol esterification | BP | 14 | 1 | 3 | 0.02291 |
| GO:0001222 | transcription corepressor binding | MF | 14 | 0 | 3 | 0.02291 |
| GO:0014074 | response to purine-containing compound | BP | 123 | 7 | 11 | 0.0231 |
| GO:0031674 | I band | CC | 108 | 1 | 10 | 0.02385 |
| GO:0017001 | antibiotic catabolic process | BP | 37 | 1 | 5 | 0.02426 |
| GO:1901505 | carbohydrate derivative transmembrane transporter activity | MF | 37 | 1 | 5 | 0.02426 |
| GO:0009250 | glucan biosynthetic process | BP | 37 | 0 | 5 | 0.02426 |
| GO:0005978 | glycogen biosynthetic process | BP | 37 | 0 | 5 | 0.02426 |
| GO:0008207 | C21-steroid hormone metabolic process | BP | 25 | 0 | 4 | 0.02439 |
| GO:0005504 | fatty acid binding | MF | 25 | 3 | 4 | 0.02439 |
| GO:0015695 | organic cation transport | BP | 25 | 0 | 4 | 0.02439 |
| GO:0031063 | regulation of histone deacetylation | BP | 25 | 0 | 4 | 0.02439 |
| GO:0009150 | purine ribonucleotide metabolic process | BP | 401 | 14 | 27 | 0.02474 |
| GO:0055013 | cardiac muscle cell development | BP | 65 | 1 | 7 | 0.02659 |
| GO:0006865 | amino acid transport | BP | 110 | 5 | 10 | 0.02669 |
| GO:0030879 | mammary gland development | BP | 110 | 6 | 10 | 0.02669 |
| GO:0042805 | actinin binding | MF | 38 | 0 | 5 | 0.02694 |
| GO:0046658 | anchored component of plasma membrane | CC | 38 | 1 | 5 | 0.02694 |
| GO:0003254 | regulation of membrane depolarization | BP | 38 | 1 | 5 | 0.02694 |
| GO:0001221 | transcription cofactor binding | MF | 38 | 1 | 5 | 0.02694 |
| GO:0019401 | alditol biosynthetic process | BP | 6 | 0 | 2 | 0.027 |
| GO:0043190 | ATP-binding cassette (ABC) transporter complex | CC | 6 | 0 | 2 | 0.027 |
| GO:0032052 | bile acid binding | MF | 6 | 0 | 2 | 0.027 |
| GO:0047498 | calcium-dependent phospholipase A2 activity | MF | 6 | 2 | 2 | 0.027 |
| GO:0019614 | catechol-containing compound catabolic process | BP | 6 | 0 | 2 | 0.027 |
| GO:0042424 | catecholamine catabolic process | BP | 6 | 0 | 2 | 0.027 |
| GO:0015889 | cobalamin transport | BP | 6 | 0 | 2 | 0.027 |
| GO:0051185 | coenzyme transmembrane transporter activity | MF | 6 | 0 | 2 | 0.027 |
| GO:0035434 | copper ion transmembrane transport | BP | 6 | 0 | 2 | 0.027 |
| GO:0004301 | epoxide hydrolase activity | MF | 6 | 0 | 2 | 0.027 |
| GO:1901569 | fatty acid derivative catabolic process | BP | 6 | 0 | 2 | 0.027 |
| GO:0006072 | glycerol-3-phosphate metabolic process | BP | 6 | 0 | 2 | 0.027 |
| GO:0002227 | innate immune response in mucosa | BP | 6 | 0 | 2 | 0.027 |
| GO:0106001 | intestinal hexose absorption | BP | 6 | 0 | 2 | 0.027 |
| GO:2000323 | negative regulation of glucocorticoid receptor signaling pathway | BP | 6 | 1 | 2 | 0.027 |
| GO:0005042 | netrin receptor activity | MF | 6 | 0 | 2 | 0.027 |
| GO:0015205 | nucleobase transmembrane transporter activity | MF | 6 | 1 | 2 | 0.027 |
| GO:0004528 | phosphodiesterase I activity | MF | 6 | 0 | 2 | 0.027 |
| GO:0032927 | positive regulation of activin receptor signaling pathway | BP | 6 | 0 | 2 | 0.027 |
| GO:0099569 | presynaptic cytoskeleton | CC | 6 | 0 | 2 | 0.027 |
| GO:0072526 | pyridine-containing compound catabolic process | BP | 6 | 0 | 2 | 0.027 |
| GO:0015651 | quaternary ammonium group transmembrane transporter activity | MF | 6 | 0 | 2 | 0.027 |
| GO:0032489 | regulation of Cdc42 protein signal transduction | BP | 6 | 0 | 2 | 0.027 |
| GO:1903998 | regulation of eating behavior | BP | 6 | 0 | 2 | 0.027 |
| GO:0032276 | regulation of gonadotropin secretion | BP | 6 | 0 | 2 | 0.027 |
| GO:0032532 | regulation of microvillus length | BP | 6 | 0 | 2 | 0.027 |
| GO:0070472 | regulation of uterine smooth muscle contraction | BP | 6 | 0 | 2 | 0.027 |
| GO:0010269 | response to selenium ion | BP | 6 | 0 | 2 | 0.027 |
| GO:0044341 | sodium-dependent phosphate transport | BP | 6 | 0 | 2 | 0.027 |
| GO:0046790 | virion binding | MF | 6 | 0 | 2 | 0.027 |
| GO:0033500 | carbohydrate homeostasis | BP | 192 | 7 | 15 | 0.0273 |
| GO:0042593 | glucose homeostasis | BP | 192 | 7 | 15 | 0.0273 |
| GO:0005261 | cation channel activity | MF | 209 | 4 | 16 | 0.02733 |
| GO:0005776 | autophagosome | CC | 80 | 0 | 8 | 0.02745 |
| GO:0090100 | positive regulation of transmembrane receptor protein serine/threon | BP | 80 | 4 | 8 | 0.02745 |
| GO:0003841 | 1-acylglycerol-3-phosphate O-acyltransferase activity | MF | 15 | 0 | 3 | 0.0277 |
| GO:0002029 | desensitization of G protein-coupled receptor signaling pathway | BP | 15 | 0 | 3 | 0.0277 |
| GO:0018904 | ether metabolic process | BP | 15 | 0 | 3 | 0.0277 |
| GO:0042362 | fat-soluble vitamin biosynthetic process | BP | 15 | 0 | 3 | 0.0277 |
| GO:0006750 | glutathione biosynthetic process | BP | 15 | 1 | 3 | 0.0277 |
| GO:0042171 | lysophosphatidic acid acyltransferase activity | MF | 15 | 0 | 3 | 0.0277 |
| GO:0022401 | negative adaptation of signaling pathway | BP | 15 | 0 | 3 | 0.0277 |
| GO:0005328 | neurotransmitter:sodium symporter activity | MF | 15 | 1 | 3 | 0.0277 |
| GO:0006595 | polyamine metabolic process | BP | 15 | 1 | 3 | 0.0277 |

| | | | | | | |
|---|---|---|---|---|---|---|
| GO:0045723 | positive regulation of fatty acid biosynthetic process | BP | 15 | 2 | 3 | 0.0277 |
| GO:0032098 | regulation of appetite | BP | 15 | 1 | 3 | 0.0277 |
| GO:0010885 | regulation of cholesterol storage | BP | 15 | 0 | 3 | 0.0277 |
| GO:0003222 | ventricular trabecula myocardium morphogenesis | BP | 15 | 0 | 3 | 0.0277 |
| GO:0032553 | ribonucleotide binding | MF | 1542 | 43 | 85 | 0.0278 |
| GO:0001569 | branching involved in blood vessel morphogenesis | BP | 26 | 0 | 4 | 0.02782 |
| GO:0006536 | glutamate metabolic process | BP | 26 | 2 | 4 | 0.02782 |
| GO:0033198 | response to ATP | BP | 26 | 2 | 4 | 0.02782 |
| GO:0031410 | cytoplasmic vesicle | CC | 1849 | 61 | 100 | 0.02786 |
| GO:0005216 | ion channel activity | MF | 262 | 5 | 19 | 0.02837 |
| GO:0035023 | regulation of Rho protein signal transduction | BP | 127 | 3 | 11 | 0.02844 |
| GO:0120025 | plasma membrane bounded cell projection | CC | 1605 | 36 | 88 | 0.02854 |
| GO:0097708 | intracellular vesicle | CC | 1851 | 61 | 100 | 0.02858 |
| GO:0001505 | regulation of neurotransmitter levels | BP | 280 | 12 | 20 | 0.02873 |
| GO:0048167 | regulation of synaptic plasticity | BP | 144 | 4 | 12 | 0.02961 |
| GO:0033865 | nucleoside bisphosphate metabolic process | BP | 112 | 4 | 10 | 0.02977 |
| GO:0034032 | purine nucleoside bisphosphate metabolic process | BP | 112 | 4 | 10 | 0.02977 |
| GO:0033875 | ribonucleoside bisphosphate metabolic process | BP | 112 | 4 | 10 | 0.02977 |
| GO:0042887 | amide transmembrane transporter activity | MF | 39 | 0 | 5 | 0.02979 |
| GO:0043062 | extracellular structure organization | BP | 353 | 16 | 24 | 0.02984 |
| GO:1990845 | adaptive thermogenesis | BP | 128 | 5 | 11 | 0.0299 |
| GO:0007411 | axon guidance | BP | 229 | 4 | 17 | 0.03033 |
| GO:0097485 | neuron projection guidance | BP | 229 | 4 | 17 | 0.03033 |
| GO:0046034 | ATP metabolic process | BP | 212 | 7 | 16 | 0.03064 |
| GO:0006635 | fatty acid beta-oxidation | BP | 67 | 1 | 7 | 0.03078 |
| GO:0045913 | positive regulation of carbohydrate metabolic process | BP | 67 | 1 | 7 | 0.03078 |
| GO:0007611 | learning or memory | BP | 195 | 4 | 15 | 0.03078 |
| GO:0044264 | cellular polysaccharide metabolic process | BP | 82 | 1 | 8 | 0.03124 |
| GO:0032559 | adenyl ribonucleotide binding | MF | 1267 | 41 | 71 | 0.03138 |
| GO:0046621 | negative regulation of organ growth | BP | 27 | 0 | 4 | 0.03153 |
| GO:0043169 | cation binding | MF | 3223 | 63 | 165 | 0.03202 |
| GO:0009156 | ribonucleoside monophosphate biosynthetic process | BP | 146 | 3 | 12 | 0.03247 |
| GO:0004879 | nuclear receptor activity | MF | 40 | 0 | 5 | 0.03283 |
| GO:0098531 | transcription factor activity, direct ligand regulated sequence-speci | MF | 40 | 0 | 5 | 0.03283 |
| GO:0023058 | adaptation of signaling pathway | BP | 16 | 0 | 3 | 0.03298 |
| GO:0006525 | arginine metabolic process | BP | 16 | 1 | 3 | 0.03298 |
| GO:0048739 | cardiac muscle fiber development | BP | 16 | 0 | 3 | 0.03298 |
| GO:0071318 | cellular response to ATP | BP | 16 | 2 | 3 | 0.03298 |
| GO:0071617 | lysophospholipid acyltransferase activity | MF | 16 | 0 | 3 | 0.03298 |
| GO:0015215 | nucleotide transmembrane transporter activity | MF | 16 | 0 | 3 | 0.03298 |
| GO:0005088 | Ras guanyl-nucleotide exchange factor activity | MF | 130 | 1 | 11 | 0.03299 |
| GO:0006112 | energy reserve metabolic process | BP | 68 | 2 | 7 | 0.03303 |
| GO:0007584 | response to nutrient | BP | 180 | 11 | 14 | 0.03331 |
| GO:0046578 | regulation of Ras protein signal transduction | BP | 215 | 6 | 16 | 0.03424 |
| GO:0060089 | molecular transducer activity | MF | 719 | 31 | 43 | 0.0353 |
| GO:1901700 | response to oxygen-containing compound | BP | 1275 | 52 | 71 | 0.03533 |
| GO:0046717 | acid secretion | BP | 84 | 7 | 8 | 0.03538 |
| GO:1901606 | alpha-amino acid catabolic process | BP | 84 | 5 | 8 | 0.03538 |
| GO:0051899 | membrane depolarization | BP | 69 | 1 | 7 | 0.03539 |
| GO:0009072 | aromatic amino acid family metabolic process | BP | 28 | 2 | 4 | 0.03552 |
| GO:0000421 | autophagosome membrane | CC | 28 | 0 | 4 | 0.03552 |
| GO:0016831 | carboxy-lyase activity | MF | 28 | 1 | 4 | 0.03552 |
| GO:0005326 | neurotransmitter transporter activity | MF | 28 | 2 | 4 | 0.03552 |
| GO:0000407 | phagophore assembly site | CC | 28 | 0 | 4 | 0.03552 |
| GO:0009259 | ribonucleotide metabolic process | BP | 415 | 14 | 27 | 0.03593 |
| GO:0055114 | oxidation-reduction process | BP | 759 | 24 | 45 | 0.03601 |
| GO:0006749 | glutathione metabolic process | BP | 41 | 1 | 5 | 0.03605 |
| GO:0031638 | zymogen activation | BP | 41 | 0 | 5 | 0.03605 |
| GO:0000166 | nucleotide binding | MF | 1746 | 50 | 94 | 0.03616 |
| GO:1901265 | nucleoside phosphate binding | MF | 1747 | 50 | 94 | 0.03662 |
| GO:0006220 | pyrimidine nucleotide metabolic process | BP | 55 | 3 | 6 | 0.03665 |
| GO:1903670 | regulation of sprouting angiogenesis | BP | 55 | 4 | 6 | 0.03665 |
| GO:0034627 | 'de novo' NAD biosynthetic process | BP | 7 | 1 | 2 | 0.03668 |
| GO:0034354 | 'de novo' NAD biosynthetic process from tryptophan | BP | 7 | 1 | 2 | 0.03668 |

| | | | | | | |
|---|---|---|---|---|---|---|
| GO:0017002 | activin-activated receptor activity | MF | 7 | 0 | 2 | 0.03668 |
| GO:0048179 | activin receptor complex | CC | 7 | 0 | 2 | 0.03668 |
| GO:0035612 | AP-2 adaptor complex binding | MF | 7 | 0 | 2 | 0.03668 |
| GO:0005347 | ATP transmembrane transporter activity | MF | 7 | 0 | 2 | 0.03668 |
| GO:0036122 | BMP binding | MF | 7 | 0 | 2 | 0.03668 |
| GO:0006657 | CDP-choline pathway | BP | 7 | 0 | 2 | 0.03668 |
| GO:0030002 | cellular anion homeostasis | BP | 7 | 0 | 2 | 0.03668 |
| GO:0030320 | cellular monovalent inorganic anion homeostasis | BP | 7 | 0 | 2 | 0.03668 |
| GO:0050962 | detection of light stimulus involved in sensory perception | BP | 7 | 0 | 2 | 0.03668 |
| GO:0050908 | detection of light stimulus involved in visual perception | BP | 7 | 0 | 2 | 0.03668 |
| GO:0072505 | divalent inorganic anion homeostasis | BP | 7 | 0 | 2 | 0.03668 |
| GO:0019585 | glucuronate metabolic process | BP | 7 | 0 | 2 | 0.03668 |
| GO:0015232 | heme transporter activity | MF | 7 | 0 | 2 | 0.03668 |
| GO:0008035 | high-density lipoprotein particle binding | MF | 7 | 1 | 2 | 0.03668 |
| GO:0014701 | junctional sarcoplasmic reticulum membrane | CC | 7 | 0 | 2 | 0.03668 |
| GO:0071676 | negative regulation of mononuclear cell migration | BP | 7 | 0 | 2 | 0.03668 |
| GO:1901642 | nucleoside transmembrane transport | BP | 7 | 1 | 2 | 0.03668 |
| GO:0005337 | nucleoside transmembrane transporter activity | MF | 7 | 1 | 2 | 0.03668 |
| GO:0015858 | nucleoside transport | BP | 7 | 1 | 2 | 0.03668 |
| GO:0006107 | oxaloacetate metabolic process | BP | 7 | 0 | 2 | 0.03668 |
| GO:0016634 | oxidoreductase activity, acting on the CH-CH group of donors, oxy | MF | 7 | 0 | 2 | 0.03668 |
| GO:0019336 | phenol-containing compound catabolic process | BP | 7 | 0 | 2 | 0.03668 |
| GO:0055062 | phosphate ion homeostasis | BP | 7 | 0 | 2 | 0.03668 |
| GO:0090336 | positive regulation of brown fat cell differentiation | BP | 7 | 0 | 2 | 0.03668 |
| GO:0010886 | positive regulation of cholesterol storage | BP | 7 | 0 | 2 | 0.03668 |
| GO:0032464 | positive regulation of protein homooligomerization | BP | 7 | 0 | 2 | 0.03668 |
| GO:0009445 | putrescine metabolic process | BP | 7 | 0 | 2 | 0.03668 |
| GO:2000322 | regulation of glucocorticoid receptor signaling pathway | BP | 7 | 1 | 2 | 0.03668 |
| GO:0099151 | regulation of postsynaptic density assembly | BP | 7 | 0 | 2 | 0.03668 |
| GO:0034163 | regulation of toll-like receptor 9 signaling pathway | BP | 7 | 0 | 2 | 0.03668 |
| GO:0017153 | sodium:dicarboxylate symporter activity | MF | 7 | 1 | 2 | 0.03668 |
| GO:0015643 | toxic substance binding | MF | 7 | 0 | 2 | 0.03668 |
| GO:0072506 | trivalent inorganic anion homeostasis | BP | 7 | 0 | 2 | 0.03668 |
| GO:0006063 | uronic acid metabolic process | BP | 7 | 0 | 2 | 0.03668 |
| GO:0070471 | uterine smooth muscle contraction | BP | 7 | 0 | 2 | 0.03668 |
| GO:0034447 | very-low-density lipoprotein particle clearance | BP | 7 | 1 | 2 | 0.03668 |
| GO:0006776 | vitamin A metabolic process | BP | 7 | 0 | 2 | 0.03668 |
| GO:0030554 | adenyl nucleotide binding | MF | 1278 | 41 | 71 | 0.03691 |
| GO:0014070 | response to organic cyclic compound | BP | 722 | 25 | 43 | 0.03739 |
| GO:0006476 | protein deacetylation | BP | 85 | 0 | 8 | 0.03758 |
| GO:0055006 | cardiac cell development | BP | 70 | 1 | 7 | 0.03785 |
| GO:0005811 | lipid droplet | CC | 70 | 2 | 7 | 0.03785 |
| GO:0005938 | cell cortex | CC | 253 | 5 | 18 | 0.03787 |
| GO:0019932 | second-messenger-mediated signaling | BP | 307 | 14 | 21 | 0.03811 |
| GO:0009063 | cellular amino acid catabolic process | BP | 101 | 5 | 9 | 0.03849 |
| GO:0010878 | cholesterol storage | BP | 17 | 0 | 3 | 0.03874 |
| GO:0042755 | eating behavior | BP | 17 | 0 | 3 | 0.03874 |
| GO:0008211 | glucocorticoid metabolic process | BP | 17 | 0 | 3 | 0.03874 |
| GO:0019184 | nonribosomal peptide biosynthetic process | BP | 17 | 1 | 3 | 0.03874 |
| GO:1903861 | positive regulation of dendrite extension | BP | 17 | 0 | 3 | 0.03874 |
| GO:0042398 | cellular modified amino acid biosynthetic process | BP | 42 | 1 | 5 | 0.03947 |
| GO:0044273 | sulfur compound catabolic process | BP | 42 | 2 | 5 | 0.03947 |
| GO:0001917 | photoreceptor inner segment | CC | 29 | 0 | 4 | 0.03978 |
| GO:0006637 | acyl-CoA metabolic process | BP | 86 | 4 | 8 | 0.03987 |
| GO:0035383 | thioester metabolic process | BP | 86 | 4 | 8 | 0.03987 |
| GO:0009410 | response to xenobiotic stimulus | BP | 202 | 4 | 15 | 0.04012 |
| GO:0043269 | regulation of ion transport | BP | 495 | 18 | 31 | 0.04043 |
| GO:0016747 | transferase activity, transferring acyl groups other than amino-acyl | MF | 185 | 4 | 14 | 0.04054 |
| GO:0044306 | neuron projection terminus | CC | 102 | 1 | 9 | 0.0406 |
| GO:0009746 | response to hexose | BP | 168 | 5 | 13 | 0.04069 |
| GO:0007409 | axonogenesis | BP | 383 | 7 | 25 | 0.041 |
| GO:0032555 | purine ribonucleotide binding | MF | 1530 | 43 | 83 | 0.04105 |
| GO:0055001 | muscle cell development | BP | 135 | 2 | 11 | 0.04167 |
| GO:0016772 | transferase activity, transferring phosphorus-containing groups | MF | 768 | 16 | 45 | 0.04249 |

| | | | | | | |
|---|---|---|---|---|---|---|
| GO:0008201 | heparin binding | MF | 119 | 7 | 10 | 0.04251 |
| GO:1901607 | alpha-amino acid biosynthetic process | BP | 57 | 7 | 6 | 0.04259 |
| GO:0030145 | manganese ion binding | MF | 57 | 1 | 6 | 0.04259 |
| GO:0006790 | sulfur compound metabolic process | BP | 293 | 8 | 20 | 0.04304 |
| GO:0090277 | positive regulation of peptide hormone secretion | BP | 72 | 1 | 7 | 0.04312 |
| GO:0046620 | regulation of organ growth | BP | 72 | 2 | 7 | 0.04312 |
| GO:0044437 | vacuolar part | CC | 461 | 13 | 29 | 0.04418 |
| GO:1900015 | regulation of cytokine production involved in inflammatory respons | BP | 30 | 1 | 4 | 0.04433 |
| GO:0042572 | retinol metabolic process | BP | 30 | 1 | 4 | 0.04433 |
| GO:0044438 | microbody part | CC | 88 | 2 | 8 | 0.04473 |
| GO:0044439 | peroxisomal part | CC | 88 | 2 | 8 | 0.04473 |
| GO:0008519 | ammonium transmembrane transporter activity | MF | 18 | 1 | 3 | 0.04497 |
| GO:0036315 | cellular response to sterol | BP | 18 | 2 | 3 | 0.04497 |
| GO:0005112 | Notch binding | MF | 18 | 0 | 3 | 0.04497 |
| GO:0005343 | organic acid:sodium symporter activity | MF | 18 | 1 | 3 | 0.04497 |
| GO:0018149 | peptide cross-linking | BP | 18 | 1 | 3 | 0.04497 |
| GO:1903859 | regulation of dendrite extension | BP | 18 | 0 | 3 | 0.04497 |
| GO:0003947 | (N-acetylneuraminyl)-galactosylglucosylceramide N-acetylgalactos | MF | 1 | 0 | 1 | 0.04511 |
| GO:0033885 | 10-hydroxy-9-(phosphonooxy)octadecanoate phosphatase activity | MF | 1 | 0 | 1 | 0.04511 |
| GO:0046360 | 2-oxobutyrate biosynthetic process | BP | 1 | 0 | 1 | 0.04511 |
| GO:0019606 | 2-oxobutyrate catabolic process | BP | 1 | 0 | 1 | 0.04511 |
| GO:0008260 | 3-oxoacid CoA-transferase activity | MF | 1 | 0 | 1 | 0.04511 |
| GO:0031687 | A2A adenosine receptor binding | MF | 1 | 0 | 1 | 0.04511 |
| GO:0008775 | acetate CoA-transferase activity | MF | 1 | 0 | 1 | 0.04511 |
| GO:0047609 | acetylputrescine deacetylase activity | MF | 1 | 0 | 1 | 0.04511 |
| GO:0047611 | acetylspermidine deacetylase activity | MF | 1 | 0 | 1 | 0.04511 |
| GO:0061811 | ADP-ribosyl cyclase activity | MF | 1 | 0 | 1 | 0.04511 |
| GO:0031700 | adrenomedullin receptor binding | MF | 1 | 0 | 1 | 0.04511 |
| GO:0008783 | agmatinase activity | MF | 1 | 0 | 1 | 0.04511 |
| GO:0103046 | alanylglutamate dipeptidase activity | MF | 1 | 0 | 1 | 0.04511 |
| GO:0005503 | all-trans retinal binding | MF | 1 | 0 | 1 | 0.04511 |
| GO:0004555 | alpha,alpha-trehalase activity | MF | 1 | 0 | 1 | 0.04511 |
| GO:0015067 | amidinotransferase activity | MF | 1 | 0 | 1 | 0.04511 |
| GO:0047685 | amine sulfotransferase activity | MF | 1 | 0 | 1 | 0.04511 |
| GO:0019202 | amino acid kinase activity | MF | 1 | 0 | 1 | 0.04511 |
| GO:0001760 | aminocarboxymuconate-semialdehyde decarboxylase activity | MF | 1 | 0 | 1 | 0.04511 |
| GO:0004971 | AMPA glutamate receptor activity | MF | 1 | 0 | 1 | 0.04511 |
| GO:0016160 | amylase activity | MF | 1 | 0 | 1 | 0.04511 |
| GO:0120077 | angiogenic sprout fusion | BP | 1 | 0 | 1 | 0.04511 |
| GO:0060182 | apelin receptor activity | MF | 1 | 0 | 1 | 0.04511 |
| GO:0060183 | apelin receptor signaling pathway | BP | 1 | 0 | 1 | 0.04511 |
| GO:0004067 | asparaginase activity | MF | 1 | 0 | 1 | 0.04511 |
| GO:0006530 | asparagine catabolic process | BP | 1 | 0 | 1 | 0.04511 |
| GO:0033345 | asparagine catabolic process via L-aspartate | BP | 1 | 0 | 1 | 0.04511 |
| GO:0033735 | aspartate dehydrogenase activity | MF | 1 | 0 | 1 | 0.04511 |
| GO:0003167 | atrioventricular bundle cell differentiation | BP | 1 | 0 | 1 | 0.04511 |
| GO:1905222 | atrioventricular canal morphogenesis | BP | 1 | 0 | 1 | 0.04511 |
| GO:1990716 | axonemal central apparatus | CC | 1 | 0 | 1 | 0.04511 |
| GO:1990718 | axonemal central pair projection | CC | 1 | 0 | 1 | 0.04511 |
| GO:0097323 | B cell adhesion | BP | 1 | 0 | 1 | 0.04511 |
| GO:0045575 | basophil activation | BP | 1 | 0 | 1 | 0.04511 |
| GO:0002276 | basophil activation involved in immune response | BP | 1 | 0 | 1 | 0.04511 |
| GO:0019115 | benzaldehyde dehydrogenase activity | MF | 1 | 0 | 1 | 0.04511 |
| GO:0019483 | beta-alanine biosynthetic process | BP | 1 | 0 | 1 | 0.04511 |
| GO:1901810 | beta-carotene metabolic process | BP | 1 | 0 | 1 | 0.04511 |
| GO:0010816 | calcitonin catabolic process | BP | 1 | 0 | 1 | 0.04511 |
| GO:1902098 | calcitriol binding | MF | 1 | 0 | 1 | 0.04511 |
| GO:0098704 | carbohydrate import across plasma membrane | BP | 1 | 0 | 1 | 0.04511 |
| GO:0035538 | carbohydrate response element binding | MF | 1 | 0 | 1 | 0.04511 |
| GO:0003881 | CDP-diacylglycerol-inositol 3-phosphatidyltransferase activity | MF | 1 | 0 | 1 | 0.04511 |
| GO:0120078 | cell adhesion involved in sprouting angiogenesis | BP | 1 | 0 | 1 | 0.04511 |
| GO:0002752 | cell surface pattern recognition receptor signaling pathway | BP | 1 | 0 | 1 | 0.04511 |
| GO:0043605 | cellular amide catabolic process | BP | 1 | 0 | 1 | 0.04511 |
| GO:0072340 | cellular lactam catabolic process | BP | 1 | 0 | 1 | 0.04511 |

| | | | | | | |
|---|---|---|---|---|---|---|
| GO:0010106 | cellular response to iron ion starvation | BP | 1 | 0 | 1 | 0.04511 |
| GO:0099038 | ceramide-translocating ATPase activity | MF | 1 | 0 | 1 | 0.04511 |
| GO:0099040 | ceramide translocation | BP | 1 | 0 | 1 | 0.04511 |
| GO:0047749 | cholestanetriol 26-monooxygenase activity | MF | 1 | 0 | 1 | 0.04511 |
| GO:0031073 | cholesterol 26-hydroxylase activity | MF | 1 | 0 | 1 | 0.04511 |
| GO:0008386 | cholesterol monooxygenase (side-chain-cleaving) activity | MF | 1 | 0 | 1 | 0.04511 |
| GO:0004102 | choline O-acetyltransferase activity | MF | 1 | 0 | 1 | 0.04511 |
| GO:0034360 | chylomicron remnant | CC | 1 | 0 | 1 | 0.04511 |
| GO:0033961 | cis-stilbene-oxide hydrolase activity | MF | 1 | 0 | 1 | 0.04511 |
| GO:0015087 | cobalt ion transmembrane transporter activity | MF | 1 | 0 | 1 | 0.04511 |
| GO:0019811 | cocaine binding | MF | 1 | 0 | 1 | 0.04511 |
| GO:0005600 | collagen type XIII trimer | CC | 1 | 0 | 1 | 0.04511 |
| GO:1990708 | conditioned place preference | BP | 1 | 0 | 1 | 0.04511 |
| GO:0051431 | corticotropin-releasing hormone receptor 2 binding | MF | 1 | 0 | 1 | 0.04511 |
| GO:0046449 | creatinine metabolic process | BP | 1 | 0 | 1 | 0.04511 |
| GO:0010852 | cyclase inhibitor activity | MF | 1 | 0 | 1 | 0.04511 |
| GO:0061812 | cyclic ADP-ribose hydrolase | MF | 1 | 0 | 1 | 0.04511 |
| GO:0102008 | cytosolic dipeptidase activity | MF | 1 | 0 | 1 | 0.04511 |
| GO:0030209 | dermatan sulfate catabolic process | BP | 1 | 0 | 1 | 0.04511 |
| GO:0017113 | dihydropyrimidine dehydrogenase (NADP+) activity | MF | 1 | 0 | 1 | 0.04511 |
| GO:0046352 | disaccharide catabolic process | BP | 1 | 0 | 1 | 0.04511 |
| GO:0008613 | diuretic hormone activity | MF | 1 | 0 | 1 | 0.04511 |
| GO:0038048 | dynorphin receptor activity | MF | 1 | 0 | 1 | 0.04511 |
| GO:0036497 | eIF2alpha dephosphorylation in response to endoplasmic reticulum | BP | 1 | 0 | 1 | 0.04511 |
| GO:0034959 | endothelin maturation | BP | 1 | 0 | 1 | 0.04511 |
| GO:0071206 | establishment of protein localization to juxtaparanode region of axo | BP | 1 | 0 | 1 | 0.04511 |
| GO:0004306 | ethanolamine-phosphate cytidylyltransferase activity | MF | 1 | 0 | 1 | 0.04511 |
| GO:0006580 | ethanolamine metabolic process | BP | 1 | 0 | 1 | 0.04511 |
| GO:0015607 | fatty-acyl-CoA transmembrane transporter activity | MF | 1 | 0 | 1 | 0.04511 |
| GO:0010430 | fatty acid omega-oxidation | BP | 1 | 0 | 1 | 0.04511 |
| GO:0042132 | fructose 1,6-bisphosphate 1-phosphatase activity | MF | 1 | 0 | 1 | 0.04511 |
| GO:0032445 | fructose import | BP | 1 | 0 | 1 | 0.04511 |
| GO:1990539 | fructose import across plasma membrane | BP | 1 | 0 | 1 | 0.04511 |
| GO:0060112 | generation of ovulation cycle rhythm | BP | 1 | 0 | 1 | 0.04511 |
| GO:0004339 | glucan 1,4-alpha-glucosidase activity | MF | 1 | 0 | 1 | 0.04511 |
| GO:0046316 | gluconokinase activity | MF | 1 | 0 | 1 | 0.04511 |
| GO:0004350 | glutamate-5-semialdehyde dehydrogenase activity | MF | 1 | 0 | 1 | 0.04511 |
| GO:0004349 | glutamate 5-kinase activity | MF | 1 | 0 | 1 | 0.04511 |
| GO:0008887 | glycerate kinase activity | MF | 1 | 0 | 1 | 0.04511 |
| GO:0047390 | glycerophosphocholine cholinephosphodiesterase activity | MF | 1 | 0 | 1 | 0.04511 |
| GO:0047394 | glycerophosphoinositol inositolphosphodiesterase activity | MF | 1 | 0 | 1 | 0.04511 |
| GO:0050479 | glyceryl-ether monooxygenase activity | MF | 1 | 0 | 1 | 0.04511 |
| GO:0015068 | glycine amidinotransferase activity | MF | 1 | 0 | 1 | 0.04511 |
| GO:0030251 | guanylate cyclase inhibitor activity | MF | 1 | 0 | 1 | 0.04511 |
| GO:0030211 | heparin catabolic process | BP | 1 | 0 | 1 | 0.04511 |
| GO:0070829 | heterochromatin maintenance | BP | 1 | 0 | 1 | 0.04511 |
| GO:0060932 | His-Purkinje system cell differentiation | BP | 1 | 0 | 1 | 0.04511 |
| GO:0001697 | histamine-induced gastric acid secretion | BP | 1 | 0 | 1 | 0.04511 |
| GO:0043990 | histone H2A-S1 phosphorylation | BP | 1 | 0 | 1 | 0.04511 |
| GO:0002416 | IgG immunoglobulin transcytosis in epithelial cells mediated by Fc | BP | 1 | 0 | 1 | 0.04511 |
| GO:0002030 | inhibitory G protein-coupled receptor phosphorylation | BP | 1 | 0 | 1 | 0.04511 |
| GO:0004447 | iodide peroxidase activity | MF | 1 | 0 | 1 | 0.04511 |
| GO:0031773 | kisspeptin receptor binding | MF | 1 | 0 | 1 | 0.04511 |
| GO:0047801 | L-cysteine:2-oxoglutarate aminotransferase activity | MF | 1 | 0 | 1 | 0.04511 |
| GO:0050023 | L-fuconate dehydratase activity | MF | 1 | 0 | 1 | 0.04511 |
| GO:0003940 | L-iduronidase activity | MF | 1 | 0 | 1 | 0.04511 |
| GO:1903801 | L-leucine import across plasma membrane | BP | 1 | 0 | 1 | 0.04511 |
| GO:0004794 | L-threonine ammonia-lyase activity | MF | 1 | 0 | 1 | 0.04511 |
| GO:0000016 | lactase activity | MF | 1 | 0 | 1 | 0.04511 |
| GO:0015094 | lead ion transmembrane transporter activity | MF | 1 | 0 | 1 | 0.04511 |
| GO:0015692 | lead ion transport | BP | 1 | 0 | 1 | 0.04511 |
| GO:0098713 | leucine import across plasma membrane | BP | 1 | 0 | 1 | 0.04511 |
| GO:0015920 | lipopolysaccharide transport | BP | 1 | 0 | 1 | 0.04511 |
| GO:1902121 | lithocholic acid binding | MF | 1 | 0 | 1 | 0.04511 |

| | | | | | | |
|---|---|---|---|---|---|---|
| GO:0038186 | lithocholic acid receptor activity | MF | 1 | 0 | 1 | 0.04511 |
| GO:0015361 | low-affinity sodium:dicarboxylate symporter activity | MF | 1 | 0 | 1 | 0.04511 |
| GO:0051978 | lysophospholipid transporter activity | MF | 1 | 0 | 1 | 0.04511 |
| GO:0033925 | mannosyl-glycoprotein endo-beta-N-acetylglucosaminidase activity | MF | 1 | 0 | 1 | 0.04511 |
| GO:0034359 | mature chylomicron | CC | 1 | 0 | 1 | 0.04511 |
| GO:0004489 | methylenetetrahydrofolate reductase (NAD(P)H) activity | MF | 1 | 0 | 1 | 0.04511 |
| GO:0032034 | myosin II head/neck binding | MF | 1 | 0 | 1 | 0.04511 |
| GO:0032038 | myosin II heavy chain binding | MF | 1 | 0 | 1 | 0.04511 |
| GO:1904283 | negative regulation of antigen processing and presentation of endog | BP | 1 | 0 | 1 | 0.04511 |
| GO:2000986 | negative regulation of behavioral fear response | BP | 1 | 0 | 1 | 0.04511 |
| GO:0106089 | negative regulation of cell adhesion involved in sprouting angiogen | BP | 1 | 0 | 1 | 0.04511 |
| GO:0060354 | negative regulation of cell adhesion molecule production | BP | 1 | 0 | 1 | 0.04511 |
| GO:0060243 | negative regulation of cell growth involved in contact inhibition | BP | 1 | 0 | 1 | 0.04511 |
| GO:1903366 | negative regulation of fear response | BP | 1 | 0 | 1 | 0.04511 |
| GO:1903597 | negative regulation of gap junction assembly | BP | 1 | 0 | 1 | 0.04511 |
| GO:0031283 | negative regulation of guanylate cyclase activity | BP | 1 | 0 | 1 | 0.04511 |
| GO:1990787 | negative regulation of hh target transcription factor activity | BP | 1 | 0 | 1 | 0.04511 |
| GO:0010987 | negative regulation of high-density lipoprotein particle clearance | BP | 1 | 0 | 1 | 0.04511 |
| GO:0051141 | negative regulation of NK T cell proliferation | BP | 1 | 0 | 1 | 0.04511 |
| GO:1904985 | negative regulation of quinolinate biosynthetic process | BP | 1 | 0 | 1 | 0.04511 |
| GO:2000098 | negative regulation of smooth muscle cell-matrix adhesion | BP | 1 | 0 | 1 | 0.04511 |
| GO:0002626 | negative regulation of T cell antigen processing and presentation | BP | 1 | 0 | 1 | 0.04511 |
| GO:0061044 | negative regulation of vascular wound healing | BP | 1 | 0 | 1 | 0.04511 |
| GO:0061837 | neuropeptide processing | BP | 1 | 0 | 1 | 0.04511 |
| GO:0099583 | neurotransmitter receptor activity involved in regulation of postsyn | MF | 1 | 0 | 1 | 0.04511 |
| GO:0035444 | nickel cation transmembrane transport | BP | 1 | 0 | 1 | 0.04511 |
| GO:0015099 | nickel cation transmembrane transporter activity | MF | 1 | 0 | 1 | 0.04511 |
| GO:0015675 | nickel cation transport | BP | 1 | 0 | 1 | 0.04511 |
| GO:0046204 | nor-spermidine metabolic process | BP | 1 | 0 | 1 | 0.04511 |
| GO:0006592 | ornithine biosynthetic process | BP | 1 | 0 | 1 | 0.04511 |
| GO:0016713 | oxidoreductase activity, acting on paired donors, with incorporation | MF | 1 | 0 | 1 | 0.04511 |
| GO:0070826 | paraferritin complex | CC | 1 | 0 | 1 | 0.04511 |
| GO:0008241 | peptidyl-dipeptidase activity | MF | 1 | 0 | 1 | 0.04511 |
| GO:0090555 | phosphatidylethanolamine-translocating ATPase activity | MF | 1 | 0 | 1 | 0.04511 |
| GO:0052731 | phosphocholine phosphatase activity | MF | 1 | 0 | 1 | 0.04511 |
| GO:0052732 | phosphoethanolamine phosphatase activity | MF | 1 | 0 | 1 | 0.04511 |
| GO:0048244 | phytanoyl-CoA dioxygenase activity | MF | 1 | 0 | 1 | 0.04511 |
| GO:1905004 | picolinic acid biosynthetic process | BP | 1 | 0 | 1 | 0.04511 |
| GO:1905003 | picolinic acid metabolic process | BP | 1 | 0 | 1 | 0.04511 |
| GO:0106047 | polyamine deacetylation | BP | 1 | 0 | 1 | 0.04511 |
| GO:0004655 | porphobilinogen synthase activity | MF | 1 | 0 | 1 | 0.04511 |
| GO:0034263 | positive regulation of autophagy in response to ER overload | BP | 1 | 0 | 1 | 0.04511 |
| GO:1903917 | positive regulation of endoplasmic reticulum stress-induced eIF2 al | BP | 1 | 0 | 1 | 0.04511 |
| GO:0071848 | positive regulation of ERK1 and ERK2 cascade via TNFSF11-med | BP | 1 | 0 | 1 | 0.04511 |
| GO:0071812 | positive regulation of fever generation by positive regulation of pro | BP | 1 | 0 | 1 | 0.04511 |
| GO:0050668 | positive regulation of homocysteine metabolic process | BP | 1 | 0 | 1 | 0.04511 |
| GO:1904325 | positive regulation of inhibitory G protein-coupled receptor phosph | BP | 1 | 0 | 1 | 0.04511 |
| GO:0035491 | positive regulation of leukotriene production involved in inflammat | BP | 1 | 0 | 1 | 0.04511 |
| GO:1901331 | positive regulation of odontoblast differentiation | BP | 1 | 0 | 1 | 0.04511 |
| GO:2000170 | positive regulation of peptidyl-cysteine S-nitrosylation | BP | 1 | 0 | 1 | 0.04511 |
| GO:1903572 | positive regulation of protein kinase D signaling | BP | 1 | 0 | 1 | 0.04511 |
| GO:1904199 | positive regulation of regulation of vascular smooth muscle cell me | BP | 1 | 0 | 1 | 0.04511 |
| GO:0014064 | positive regulation of serotonin secretion | BP | 1 | 0 | 1 | 0.04511 |
| GO:2000640 | positive regulation of SREBP signaling pathway | BP | 1 | 0 | 1 | 0.04511 |
| GO:0031337 | positive regulation of sulfur amino acid metabolic process | BP | 1 | 0 | 1 | 0.04511 |
| GO:0045887 | positive regulation of synaptic growth at neuromuscular junction | BP | 1 | 0 | 1 | 0.04511 |
| GO:0061402 | positive regulation of transcription from RNA polymerase II promo | BP | 1 | 0 | 1 | 0.04511 |
| GO:0004925 | prolactin receptor activity | MF | 1 | 0 | 1 | 0.04511 |
| GO:0102009 | proline dipeptidase activity | MF | 1 | 0 | 1 | 0.04511 |
| GO:1904854 | proteasome core complex binding | MF | 1 | 0 | 1 | 0.04511 |
| GO:0015211 | purine nucleoside transmembrane transporter activity | MF | 1 | 0 | 1 | 0.04511 |
| GO:0032920 | putrescine acetylation | BP | 1 | 0 | 1 | 0.04511 |
| GO:0033388 | putrescine biosynthetic process from arginine | BP | 1 | 0 | 1 | 0.04511 |
| GO:0033389 | putrescine biosynthetic process from arginine, using agmatinase | BP | 1 | 0 | 1 | 0.04511 |

| | | | | | | |
|---|---|---|---|---|---|---|
| GO:0009443 | pyridoxal 5'-phosphate salvage | BP | 1 | 0 | 1 | 0.04511 |
| GO:0008478 | pyridoxal kinase activity | MF | 1 | 0 | 1 | 0.04511 |
| GO:0032361 | pyridoxal phosphate catabolic process | BP | 1 | 0 | 1 | 0.04511 |
| GO:0046127 | pyrimidine deoxyribonucleoside catabolic process | BP | 1 | 0 | 1 | 0.04511 |
| GO:0002061 | pyrimidine nucleobase binding | MF | 1 | 0 | 1 | 0.04511 |
| GO:1905012 | regulation of 'de novo' NAD biosynthetic process from tryptophan | BP | 1 | 0 | 1 | 0.04511 |
| GO:1904282 | regulation of antigen processing and presentation of endogenous pe | BP | 1 | 0 | 1 | 0.04511 |
| GO:0106088 | regulation of cell adhesion involved in sprouting angiogenesis | BP | 1 | 0 | 1 | 0.04511 |
| GO:0072366 | regulation of cellular ketone metabolic process by positive regulati | BP | 1 | 0 | 1 | 0.04511 |
| GO:1903916 | regulation of endoplasmic reticulum stress-induced eIF2 alpha depl | BP | 1 | 0 | 1 | 0.04511 |
| GO:1904323 | regulation of inhibitory G protein-coupled receptor phosphorylation | BP | 1 | 0 | 1 | 0.04511 |
| GO:0035490 | regulation of leukotriene production involved in inflammatory resp | BP | 1 | 0 | 1 | 0.04511 |
| GO:0061074 | regulation of neural retina development | BP | 1 | 0 | 1 | 0.04511 |
| GO:1903570 | regulation of protein kinase D signaling | BP | 1 | 0 | 1 | 0.04511 |
| GO:1904984 | regulation of quinolinate biosynthetic process | BP | 1 | 0 | 1 | 0.04511 |
| GO:0090067 | regulation of thalamus size | BP | 1 | 0 | 1 | 0.04511 |
| GO:0090357 | regulation of tryptophan metabolic process | BP | 1 | 0 | 1 | 0.04511 |
| GO:1990736 | regulation of vascular smooth muscle cell membrane depolarization | BP | 1 | 0 | 1 | 0.04511 |
| GO:1904681 | response to 3-methylcholanthrene | BP | 1 | 0 | 1 | 0.04511 |
| GO:1903165 | response to polycyclic arene | BP | 1 | 0 | 1 | 0.04511 |
| GO:1990267 | response to transition metal nanoparticle | BP | 1 | 0 | 1 | 0.04511 |
| GO:0033274 | response to vitamin B2 | BP | 1 | 0 | 1 | 0.04511 |
| GO:0001135 | RNA polymerase II transcription regulator recruiting activity | MF | 1 | 0 | 1 | 0.04511 |
| GO:0044317 | rod spherule | CC | 1 | 0 | 1 | 0.04511 |
| GO:1901702 | salt transmembrane transporter activity | MF | 1 | 0 | 1 | 0.04511 |
| GO:0008513 | secondary active organic cation transmembrane transporter activity | MF | 1 | 0 | 1 | 0.04511 |
| GO:0015055 | secretin receptor activity | MF | 1 | 0 | 1 | 0.04511 |
| GO:0070009 | serine-type aminopeptidase activity | MF | 1 | 0 | 1 | 0.04511 |
| GO:0106048 | spermidine deacetylation | BP | 1 | 0 | 1 | 0.04511 |
| GO:0032919 | spermine acetylation | BP | 1 | 0 | 1 | 0.04511 |
| GO:0048792 | spontaneous exocytosis of neurotransmitter | BP | 1 | 0 | 1 | 0.04511 |
| GO:0005983 | starch catabolic process | BP | 1 | 0 | 1 | 0.04511 |
| GO:0005982 | starch metabolic process | BP | 1 | 0 | 1 | 0.04511 |
| GO:0046272 | stilbene catabolic process | BP | 1 | 0 | 1 | 0.04511 |
| GO:0009810 | stilbene metabolic process | BP | 1 | 0 | 1 | 0.04511 |
| GO:0035003 | subapical complex | CC | 1 | 0 | 1 | 0.04511 |
| GO:0010814 | substance P catabolic process | BP | 1 | 0 | 1 | 0.04511 |
| GO:0016496 | substance P receptor activity | MF | 1 | 0 | 1 | 0.04511 |
| GO:0047369 | succinate-hydroxymethylglutarate CoA-transferase activity | MF | 1 | 0 | 1 | 0.04511 |
| GO:0005986 | sucrose biosynthetic process | BP | 1 | 0 | 1 | 0.04511 |
| GO:0046901 | tetrahydrofolylpolyglutamate biosynthetic process | BP | 1 | 0 | 1 | 0.04511 |
| GO:0004326 | tetrahydrofolylpolyglutamate synthase activity | MF | 1 | 0 | 1 | 0.04511 |
| GO:0047127 | thiomorpholine-carboxylate dehydrogenase activity | MF | 1 | 0 | 1 | 0.04511 |
| GO:0009088 | threonine biosynthetic process | BP | 1 | 0 | 1 | 0.04511 |
| GO:0004795 | threonine synthase activity | MF | 1 | 0 | 1 | 0.04511 |
| GO:0006214 | thymidine catabolic process | BP | 1 | 0 | 1 | 0.04511 |
| GO:0001594 | trace-amine receptor activity | MF | 1 | 0 | 1 | 0.04511 |
| GO:0030936 | transmembrane collagen trimer | CC | 1 | 0 | 1 | 0.04511 |
| GO:0015927 | trehalase activity | MF | 1 | 0 | 1 | 0.04511 |
| GO:0005993 | trehalose catabolic process | BP | 1 | 0 | 1 | 0.04511 |
| GO:0002058 | uracil binding | MF | 1 | 0 | 1 | 0.04511 |
| GO:0006212 | uracil catabolic process | BP | 1 | 0 | 1 | 0.04511 |
| GO:0015143 | urate transmembrane transporter activity | MF | 1 | 0 | 1 | 0.04511 |
| GO:0015747 | urate transport | BP | 1 | 0 | 1 | 0.04511 |
| GO:0015265 | urea channel activity | MF | 1 | 0 | 1 | 0.04511 |
| GO:0036113 | very long-chain fatty-acyl-CoA catabolic process | BP | 1 | 0 | 1 | 0.04511 |
| GO:0035238 | vitamin A biosynthetic process | BP | 1 | 0 | 1 | 0.04511 |
| GO:0042820 | vitamin B6 catabolic process | BP | 1 | 0 | 1 | 0.04511 |
| GO:0008431 | vitamin E binding | MF | 1 | 0 | 1 | 0.04511 |
| GO:0004856 | xylulokinase activity | MF | 1 | 0 | 1 | 0.04511 |
| GO:0005998 | xylulose catabolic process | BP | 1 | 0 | 1 | 0.04511 |
| GO:0009167 | purine ribonucleoside monophosphate metabolic process | BP | 241 | 7 | 17 | 0.04571 |
| GO:0098659 | inorganic cation import across plasma membrane | BP | 58 | 3 | 6 | 0.04577 |
| GO:0099587 | inorganic ion import across plasma membrane | BP | 58 | 3 | 6 | 0.04577 |

| | | | | | | |
|---|---|---|---|---|---|---|
| GO:0046914 | transition metal ion binding | MF | 812 | 12 | 47 | 0.0461 |
| GO:0006935 | chemotaxis | BP | 482 | 20 | 30 | 0.04627 |
| GO:0038023 | signaling receptor activity | MF | 675 | 31 | 40 | 0.04648 |
| GO:0005783 | endoplasmic reticulum | CC | 1499 | 34 | 81 | 0.04672 |
| GO:0009126 | purine nucleoside monophosphate metabolic process | BP | 242 | 7 | 17 | 0.0472 |
| GO:0016705 | oxidoreductase activity, acting on paired donors, with incorporation | MF | 105 | 2 | 9 | 0.04741 |
| GO:0046185 | aldehyde catabolic process | BP | 8 | 0 | 2 | 0.04747 |
| GO:0072488 | ammonium transmembrane transport | BP | 8 | 1 | 2 | 0.04747 |
| GO:0015867 | ATP transport | BP | 8 | 0 | 2 | 0.04747 |
| GO:0035620 | ceramide transporter activity | MF | 8 | 0 | 2 | 0.04747 |
| GO:0019695 | choline metabolic process | BP | 8 | 0 | 2 | 0.04747 |
| GO:0034382 | chylomicron remnant clearance | BP | 8 | 1 | 2 | 0.04747 |
| GO:0051182 | coenzyme transport | BP | 8 | 1 | 2 | 0.04747 |
| GO:0034650 | cortisol metabolic process | BP | 8 | 0 | 2 | 0.04747 |
| GO:0070567 | cytidylyltransferase activity | MF | 8 | 0 | 2 | 0.04747 |
| GO:0035907 | dorsal aorta development | BP | 8 | 0 | 2 | 0.04747 |
| GO:0035912 | dorsal aorta morphogenesis | BP | 8 | 0 | 2 | 0.04747 |
| GO:0060856 | establishment of blood-brain barrier | BP | 8 | 0 | 2 | 0.04747 |
| GO:0015886 | heme transport | BP | 8 | 0 | 2 | 0.04747 |
| GO:0042577 | lipid phosphatase activity | MF | 8 | 0 | 2 | 0.04747 |
| GO:0042711 | maternal behavior | BP | 8 | 0 | 2 | 0.04747 |
| GO:0060586 | multicellular organismal iron ion homeostasis | BP | 8 | 0 | 2 | 0.04747 |
| GO:0035529 | NADH pyrophosphatase activity | MF | 8 | 0 | 2 | 0.04747 |
| GO:0045602 | negative regulation of endothelial cell differentiation | BP | 8 | 0 | 2 | 0.04747 |
| GO:0033007 | negative regulation of mast cell activation involved in immune resp | BP | 8 | 1 | 2 | 0.04747 |
| GO:1990535 | neuron projection maintenance | BP | 8 | 0 | 2 | 0.04747 |
| GO:0038092 | nodal signaling pathway | BP | 8 | 0 | 2 | 0.04747 |
| GO:0015851 | nucleobase transport | BP | 8 | 1 | 2 | 0.04747 |
| GO:0019755 | one-carbon compound transport | BP | 8 | 0 | 2 | 0.04747 |
| GO:0060746 | parental behavior | BP | 8 | 0 | 2 | 0.04747 |
| GO:0042605 | peptide antigen binding | MF | 8 | 1 | 2 | 0.04747 |
| GO:0034638 | phosphatidylcholine catabolic process | BP | 8 | 1 | 2 | 0.04747 |
| GO:0036149 | phosphatidylinositol acyl-chain remodeling | BP | 8 | 1 | 2 | 0.04747 |
| GO:0010873 | positive regulation of cholesterol esterification | BP | 8 | 1 | 2 | 0.04747 |
| GO:0019870 | potassium channel inhibitor activity | MF | 8 | 0 | 2 | 0.04747 |
| GO:0015697 | quaternary ammonium group transport | BP | 8 | 0 | 2 | 0.04747 |
| GO:0021936 | regulation of cerebellar granule cell precursor proliferation | BP | 8 | 0 | 2 | 0.04747 |
| GO:1904889 | regulation of excitatory synapse assembly | BP | 8 | 0 | 2 | 0.04747 |
| GO:0033131 | regulation of glucokinase activity | BP | 8 | 0 | 2 | 0.04747 |
| GO:0150052 | regulation of postsynapse assembly | BP | 8 | 0 | 2 | 0.04747 |
| GO:0099150 | regulation of postsynaptic specialization assembly | BP | 8 | 0 | 2 | 0.04747 |
| GO:0019626 | short-chain fatty acid catabolic process | BP | 8 | 0 | 2 | 0.04747 |
| GO:0038203 | TORC2 signaling | BP | 8 | 0 | 2 | 0.04747 |
| GO:0099545 | trans-synaptic signaling by trans-synaptic complex | BP | 8 | 1 | 2 | 0.04747 |
| GO:0071830 | triglyceride-rich lipoprotein particle clearance | BP | 8 | 1 | 2 | 0.04747 |
| GO:0048845 | venous blood vessel morphogenesis | BP | 8 | 0 | 2 | 0.04747 |
| GO:0019935 | cyclic-nucleotide-mediated signaling | BP | 138 | 8 | 11 | 0.04758 |
| GO:0098793 | presynapse | CC | 389 | 9 | 25 | 0.04774 |
| GO:0022836 | gated channel activity | MF | 207 | 7 | 15 | 0.04794 |
| GO:0042330 | taxis | BP | 484 | 20 | 30 | 0.04838 |
| GO:0017076 | purine nucleotide binding | MF | 1543 | 43 | 83 | 0.04849 |
| GO:0072562 | blood microparticle | CC | 74 | 4 | 7 | 0.04885 |
| GO:0010008 | endosome membrane | CC | 390 | 13 | 25 | 0.04894 |
| GO:0048812 | neuron projection morphogenesis | BP | 542 | 11 | 33 | 0.04897 |
| GO:0006778 | porphyrin-containing compound metabolic process | BP | 31 | 0 | 4 | 0.04915 |
| GO:0005773 | vacuole | CC | 639 | 21 | 38 | 0.04919 |
| GO:0034284 | response to monosaccharide | BP | 173 | 5 | 13 | 0.04946 |
| GO:0009124 | nucleoside monophosphate biosynthetic process | BP | 156 | 3 | 12 | 0.04979 |

**Supp. Table 6b. GO sets upregulated in non-stricture**

| GO ID | Term | Ont | N | Up | Down | P-value |
|---|---|---|---|---|---|---|
| GO:0002376 | immune system process | BP | 2192 | 110 | 86 | 1.24E-15 |
| GO:0006955 | immune response | BP | 1482 | 83 | 50 | 4.21E-14 |

| | | | | | | |
|---|---|---|---|---|---|---|
| GO:0019730 | antimicrobial humoral response | BP | 49 | 13 | 0 | 7.97E-11 |
| GO:0044421 | extracellular region part | CC | 2383 | 104 | 145 | 8.70E-11 |
| GO:0006950 | response to stress | BP | 3001 | 122 | 106 | 9.11E-11 |
| GO:0008283 | cell proliferation | BP | 1479 | 75 | 56 | 1.33E-10 |
| GO:0005615 | extracellular space | CC | 2237 | 99 | 134 | 1.43E-10 |
| GO:0000280 | nuclear division | BP | 322 | 30 | 7 | 1.81E-10 |
| GO:0002252 | immune effector process | BP | 900 | 54 | 31 | 2.54E-10 |
| GO:0006952 | defense response | BP | 1138 | 62 | 38 | 4.85E-10 |
| GO:0002443 | leukocyte mediated immunity | BP | 598 | 41 | 21 | 1.05E-09 |
| GO:0048285 | organelle fission | BP | 357 | 30 | 8 | 2.16E-09 |
| GO:0050896 | response to stimulus | BP | 6351 | 203 | 305 | 3.03E-09 |
| GO:0042127 | regulation of cell proliferation | BP | 1226 | 63 | 47 | 3.54E-09 |
| GO:0005576 | extracellular region | CC | 2794 | 111 | 169 | 4.59E-09 |
| GO:0002366 | leukocyte activation involved in immune response | BP | 559 | 38 | 21 | 5.77E-09 |
| GO:0002263 | cell activation involved in immune response | BP | 562 | 38 | 21 | 6.68E-09 |
| GO:0071345 | cellular response to cytokine stimulus | BP | 849 | 49 | 31 | 7.05E-09 |
| GO:0001775 | cell activation | BP | 1079 | 57 | 35 | 8.68E-09 |
| GO:0006959 | humoral immune response | BP | 129 | 17 | 4 | 1.13E-08 |
| GO:0140014 | mitotic nuclear division | BP | 237 | 23 | 5 | 1.22E-08 |
| GO:0000278 | mitotic cell cycle | BP | 866 | 49 | 16 | 1.34E-08 |
| GO:0045321 | leukocyte activation | BP | 954 | 52 | 30 | 1.57E-08 |
| GO:0002682 | regulation of immune system process | BP | 1159 | 59 | 39 | 1.75E-08 |
| GO:0045087 | innate immune response | BP | 649 | 40 | 15 | 3.46E-08 |
| GO:0034097 | response to cytokine | BP | 923 | 50 | 34 | 3.79E-08 |
| GO:0007059 | chromosome segregation | BP | 261 | 23 | 2 | 7.40E-08 |
| GO:0002444 | myeloid leukocyte mediated immunity | BP | 449 | 31 | 19 | 1.15E-07 |
| GO:1990869 | cellular response to chemokine | BP | 59 | 11 | 1 | 1.24E-07 |
| GO:1990868 | response to chemokine | BP | 59 | 11 | 1 | 1.24E-07 |
| GO:0043299 | leukocyte degranulation | BP | 433 | 30 | 17 | 1.72E-07 |
| GO:0098813 | nuclear chromosome segregation | BP | 212 | 20 | 2 | 1.78E-07 |
| GO:0006954 | inflammatory response | BP | 534 | 34 | 22 | 1.88E-07 |
| GO:1903047 | mitotic cell cycle process | BP | 750 | 42 | 10 | 2.21E-07 |
| GO:0009605 | response to external stimulus | BP | 1735 | 74 | 99 | 3.20E-07 |
| GO:0070098 | chemokine-mediated signaling pathway | BP | 52 | 10 | 0 | 3.46E-07 |
| GO:0050776 | regulation of immune response | BP | 766 | 42 | 22 | 3.90E-07 |
| GO:0000819 | sister chromatid segregation | BP | 164 | 17 | 2 | 4.00E-07 |
| GO:0048518 | positive regulation of biological process | BP | 4680 | 154 | 193 | 5.83E-07 |
| GO:0051094 | positive regulation of developmental process | BP | 1043 | 51 | 44 | 6.70E-07 |
| GO:0046903 | secretion | BP | 1288 | 59 | 74 | 6.90E-07 |
| GO:0051301 | cell division | BP | 488 | 31 | 8 | 7.12E-07 |
| GO:0002275 | myeloid cell activation involved in immune response | BP | 440 | 29 | 19 | 7.86E-07 |
| GO:0009986 | cell surface | CC | 598 | 35 | 40 | 9.05E-07 |
| GO:0070887 | cellular response to chemical stimulus | BP | 2475 | 94 | 120 | 1.10E-06 |
| GO:0000070 | mitotic sister chromatid segregation | BP | 139 | 15 | 2 | 1.16E-06 |
| GO:0002274 | myeloid leukocyte activation | BP | 528 | 32 | 19 | 1.30E-06 |
| GO:0001817 | regulation of cytokine production | BP | 533 | 32 | 19 | 1.60E-06 |
| GO:0043230 | extracellular organelle | CC | 1679 | 70 | 97 | 1.71E-06 |
| GO:0008284 | positive regulation of cell proliferation | BP | 671 | 37 | 26 | 1.80E-06 |
| GO:0051716 | cellular response to stimulus | BP | 5330 | 168 | 238 | 2.04E-06 |
| GO:0034599 | cellular response to oxidative stress | BP | 248 | 20 | 7 | 2.16E-06 |
| GO:0002684 | positive regulation of immune system process | BP | 820 | 42 | 24 | 2.32E-06 |
| GO:0070062 | extracellular exosome | CC | 1661 | 69 | 95 | 2.36E-06 |
| GO:0045055 | regulated exocytosis | BP | 657 | 36 | 29 | 2.91E-06 |
| GO:0048584 | positive regulation of response to stimulus | BP | 1809 | 73 | 69 | 3.22E-06 |
| GO:1903561 | extracellular vesicle | CC | 1677 | 69 | 96 | 3.31E-06 |
| GO:0000793 | condensed chromosome | CC | 171 | 16 | 0 | 3.44E-06 |
| GO:0072686 | mitotic spindle | CC | 98 | 12 | 3 | 3.68E-06 |
| GO:0051240 | positive regulation of multicellular organismal process | BP | 1328 | 58 | 54 | 4.01E-06 |
| GO:0031428 | box C/D snoRNP complex | CC | 6 | 4 | 0 | 4.65E-06 |
| GO:0001816 | cytokine production | BP | 589 | 33 | 22 | 4.86E-06 |
| GO:0048522 | positive regulation of cellular process | BP | 4167 | 137 | 170 | 5.19E-06 |
| GO:0071310 | cellular response to organic substance | BP | 2072 | 80 | 90 | 5.23E-06 |
| GO:0042119 | neutrophil activation | BP | 408 | 26 | 16 | 5.53E-06 |
| GO:0051239 | regulation of multicellular organismal process | BP | 2390 | 89 | 111 | 5.62E-06 |

| | | | | | | |
|---|---|---|---|---|---|---|
| GO:0080134 | regulation of response to stress | BP | 1222 | 54 | 39 | 6.56E-06 |
| GO:0036230 | granulocyte activation | BP | 412 | 26 | 17 | 6.59E-06 |
| GO:0031982 | vesicle | CC | 2976 | 105 | 166 | 6.64E-06 |
| GO:0042379 | chemokine receptor binding | MF | 31 | 7 | 0 | 6.74E-06 |
| GO:0051783 | regulation of nuclear division | BP | 160 | 15 | 5 | 6.79E-06 |
| GO:1901701 | cellular response to oxygen-containing compound | BP | 890 | 43 | 47 | 7.68E-06 |
| GO:0008009 | chemokine activity | MF | 22 | 6 | 0 | 9.66E-06 |
| GO:0043312 | neutrophil degranulation | BP | 396 | 25 | 16 | 9.90E-06 |
| GO:0044459 | plasma membrane part | CC | 2005 | 77 | 138 | 1.02E-05 |
| GO:0007088 | regulation of mitotic nuclear division | BP | 146 | 14 | 5 | 1.06E-05 |
| GO:0001906 | cell killing | BP | 91 | 11 | 2 | 1.06E-05 |
| GO:0002283 | neutrophil activation involved in immune response | BP | 398 | 25 | 16 | 1.08E-05 |
| GO:0006887 | exocytosis | BP | 755 | 38 | 32 | 1.10E-05 |
| GO:0007049 | cell cycle | BP | 1505 | 62 | 35 | 1.14E-05 |
| GO:0005819 | spindle | CC | 301 | 21 | 4 | 1.17E-05 |
| GO:0019221 | cytokine-mediated signaling pathway | BP | 588 | 32 | 19 | 1.23E-05 |
| GO:0022402 | cell cycle process | BP | 1123 | 50 | 20 | 1.25E-05 |
| GO:0010033 | response to organic substance | BP | 2512 | 91 | 114 | 1.32E-05 |
| GO:0048248 | CXCR3 chemokine receptor binding | MF | 3 | 3 | 0 | 1.36E-05 |
| GO:0065007 | biological regulation | BP | 8644 | 241 | 357 | 1.40E-05 |
| GO:0007154 | cell communication | BP | 4596 | 146 | 208 | 1.45E-05 |
| GO:0006968 | cellular defense response | BP | 35 | 7 | 0 | 1.59E-05 |
| GO:0008219 | cell death | BP | 1688 | 67 | 50 | 1.60E-05 |
| GO:0051983 | regulation of chromosome segregation | BP | 95 | 11 | 2 | 1.61E-05 |
| GO:0033554 | cellular response to stress | BP | 1723 | 68 | 55 | 1.64E-05 |
| GO:0002446 | neutrophil mediated immunity | BP | 408 | 25 | 17 | 1.65E-05 |
| GO:0071944 | cell periphery | CC | 3686 | 122 | 250 | 1.95E-05 |
| GO:0051707 | response to other organism | BP | 659 | 34 | 19 | 1.99E-05 |
| GO:0043207 | response to external biotic stimulus | BP | 660 | 34 | 19 | 2.06E-05 |
| GO:0045236 | CXCR chemokine receptor binding | MF | 8 | 4 | 0 | 2.09E-05 |
| GO:0071219 | cellular response to molecule of bacterial origin | BP | 155 | 14 | 5 | 2.10E-05 |
| GO:0061844 | antimicrobial humoral immune response mediated by antimicrobial | BP | 25 | 6 | 0 | 2.16E-05 |
| GO:0007346 | regulation of mitotic cell cycle | BP | 551 | 30 | 11 | 2.32E-05 |
| GO:0009897 | external side of plasma membrane | CC | 221 | 17 | 17 | 2.34E-05 |
| GO:0006979 | response to oxidative stress | BP | 366 | 23 | 10 | 2.43E-05 |
| GO:0042592 | homeostatic process | BP | 1476 | 60 | 85 | 2.45E-05 |
| GO:0071216 | cellular response to biotic stimulus | BP | 178 | 15 | 6 | 2.45E-05 |
| GO:0044772 | mitotic cell cycle phase transition | BP | 498 | 28 | 8 | 2.47E-05 |
| GO:0030246 | carbohydrate binding | MF | 179 | 15 | 17 | 2.62E-05 |
| GO:0031347 | regulation of defense response | BP | 612 | 32 | 24 | 2.73E-05 |
| GO:0050793 | regulation of developmental process | BP | 1991 | 75 | 82 | 2.74E-05 |
| GO:0009617 | response to bacterium | BP | 421 | 25 | 18 | 2.79E-05 |
| GO:0001909 | leukocyte mediated cytotoxicity | BP | 67 | 9 | 1 | 2.90E-05 |
| GO:0048583 | regulation of response to stimulus | BP | 3224 | 109 | 132 | 2.92E-05 |
| GO:0071396 | cellular response to lipid | BP | 476 | 27 | 25 | 2.98E-05 |
| GO:0042221 | response to chemical | BP | 3265 | 110 | 175 | 3.07E-05 |
| GO:0032653 | regulation of interleukin-10 production | BP | 39 | 7 | 1 | 3.35E-05 |
| GO:0050789 | regulation of biological process | BP | 8170 | 229 | 324 | 3.38E-05 |
| GO:0030155 | regulation of cell adhesion | BP | 563 | 30 | 24 | 3.47E-05 |
| GO:0007166 | cell surface receptor signaling pathway | BP | 2252 | 82 | 83 | 3.54E-05 |
| GO:0019725 | cellular homeostasis | BP | 740 | 36 | 41 | 3.92E-05 |
| GO:0046649 | lymphocyte activation | BP | 512 | 28 | 13 | 4.06E-05 |
| GO:0051241 | negative regulation of multicellular organismal process | BP | 925 | 42 | 44 | 4.32E-05 |
| GO:0000777 | condensed chromosome kinetochore | CC | 88 | 10 | 0 | 4.62E-05 |
| GO:0005886 | plasma membrane | CC | 3600 | 118 | 245 | 4.76E-05 |
| GO:0009607 | response to biotic stimulus | BP | 688 | 34 | 19 | 4.77E-05 |
| GO:0010941 | regulation of cell death | BP | 1314 | 54 | 37 | 5.05E-05 |
| GO:0002250 | adaptive immune response | BP | 283 | 19 | 7 | 5.17E-05 |
| GO:0023052 | signaling | BP | 4581 | 143 | 203 | 5.27E-05 |
| GO:0000942 | condensed nuclear chromosome outer kinetochore | CC | 4 | 3 | 0 | 5.33E-05 |
| GO:0008608 | attachment of spindle microtubules to kinetochore | BP | 29 | 6 | 0 | 5.34E-05 |
| GO:0032940 | secretion by cell | BP | 1187 | 50 | 62 | 5.37E-05 |
| GO:0032613 | interleukin-10 production | BP | 42 | 7 | 2 | 5.52E-05 |
| GO:0009987 | cellular process | BP | 11175 | 290 | 473 | 5.53E-05 |

| | | | | | | |
|---|---|---|---|---|---|---|
| GO:0065008 | regulation of biological quality | BP | 3088 | 104 | 180 | 5.92E-05 |
| GO:0071222 | cellular response to lipopolysaccharide | BP | 149 | 13 | 5 | 5.95E-05 |
| GO:0072593 | reactive oxygen species metabolic process | BP | 215 | 16 | 11 | 6.05E-05 |
| GO:0022603 | regulation of anatomical structure morphogenesis | BP | 877 | 40 | 36 | 6.05E-05 |
| GO:0042581 | specific granule | CC | 129 | 12 | 9 | 6.13E-05 |
| GO:0012501 | programmed cell death | BP | 1592 | 62 | 46 | 6.31E-05 |
| GO:0140013 | meiotic nuclear division | BP | 110 | 11 | 3 | 6.41E-05 |
| GO:1904724 | tertiary granule lumen | CC | 43 | 7 | 2 | 6.46E-05 |
| GO:0044770 | cell cycle phase transition | BP | 527 | 28 | 9 | 6.72E-05 |
| GO:0099503 | secretory vesicle | CC | 761 | 36 | 33 | 6.99E-05 |
| GO:0000082 | G1/S transition of mitotic cell cycle | BP | 219 | 16 | 2 | 7.53E-05 |
| GO:0002697 | regulation of immune effector process | BP | 293 | 19 | 7 | 8.24E-05 |
| GO:1901700 | response to oxygen-containing compound | BP | 1275 | 52 | 71 | 8.67E-05 |
| GO:0002699 | positive regulation of immune effector process | BP | 156 | 13 | 3 | 9.54E-05 |
| GO:0070301 | cellular response to hydrogen peroxide | BP | 78 | 9 | 2 | 9.81E-05 |
| GO:0035690 | cellular response to drug | BP | 297 | 19 | 12 | 9.86E-05 |
| GO:0001910 | regulation of leukocyte mediated cytotoxicity | BP | 46 | 7 | 1 | 0.0001 |
| GO:0033993 | response to lipid | BP | 715 | 34 | 37 | 0.0001 |
| GO:0050727 | regulation of inflammatory response | BP | 273 | 18 | 14 | 0.0001 |
| GO:0005694 | chromosome | CC | 837 | 38 | 4 | 0.0001 |
| GO:0009628 | response to abiotic stimulus | BP | 993 | 43 | 41 | 0.0001 |
| GO:0005125 | cytokine activity | MF | 97 | 10 | 3 | 0.00011 |
| GO:0001913 | T cell mediated cytotoxicity | BP | 21 | 5 | 0 | 0.00011 |
| GO:0007052 | mitotic spindle organization | BP | 98 | 10 | 0 | 0.00012 |
| GO:0035580 | specific granule lumen | CC | 47 | 7 | 4 | 0.00012 |
| GO:0043235 | receptor complex | CC | 301 | 19 | 17 | 0.00012 |
| GO:0071236 | cellular response to antibiotic | BP | 119 | 11 | 3 | 0.00013 |
| GO:0070663 | regulation of leukocyte proliferation | BP | 161 | 13 | 7 | 0.00013 |
| GO:0007165 | signal transduction | BP | 4225 | 132 | 179 | 0.00013 |
| GO:0005871 | kinesin complex | CC | 48 | 7 | 5 | 0.00013 |
| GO:0030141 | secretory granule | CC | 636 | 31 | 26 | 0.00013 |
| GO:0051988 | regulation of attachment of spindle microtubules to kinetochore | BP | 12 | 4 | 0 | 0.00014 |
| GO:0000779 | condensed chromosome, centromeric region | CC | 100 | 10 | 0 | 0.00014 |
| GO:0051726 | regulation of cell cycle | BP | 1006 | 43 | 24 | 0.00014 |
| GO:0060548 | negative regulation of cell death | BP | 758 | 35 | 21 | 0.00014 |
| GO:0050778 | positive regulation of immune response | BP | 609 | 30 | 13 | 0.00014 |
| GO:0050794 | regulation of cellular process | BP | 7678 | 215 | 286 | 0.00014 |
| GO:0097237 | cellular response to toxic substance | BP | 185 | 14 | 8 | 0.00015 |
| GO:0044843 | cell cycle G1/S phase transition | BP | 232 | 16 | 3 | 0.00015 |
| GO:0060429 | epithelium development | BP | 883 | 39 | 32 | 0.00015 |
| GO:1902850 | microtubule cytoskeleton organization involved in mitosis | BP | 122 | 11 | 1 | 0.00016 |
| GO:0051179 | localization | BP | 5081 | 153 | 276 | 0.00017 |
| GO:0002703 | regulation of leukocyte mediated immunity | BP | 143 | 12 | 4 | 0.00017 |
| GO:0030071 | regulation of mitotic metaphase/anaphase transition | BP | 50 | 7 | 2 | 0.00017 |
| GO:0016192 | vesicle-mediated transport | BP | 1684 | 63 | 82 | 0.00017 |
| GO:0010564 | regulation of cell cycle process | BP | 647 | 31 | 11 | 0.00018 |
| GO:1903046 | meiotic cell cycle process | BP | 124 | 11 | 3 | 0.00019 |
| GO:1902099 | regulation of metaphase/anaphase transition of cell cycle | BP | 51 | 7 | 2 | 0.0002 |
| GO:0009636 | response to toxic substance | BP | 393 | 22 | 24 | 0.0002 |
| GO:0070661 | leukocyte proliferation | BP | 214 | 15 | 9 | 0.0002 |
| GO:0002237 | response to molecule of bacterial origin | BP | 263 | 17 | 13 | 0.0002 |
| GO:0090307 | mitotic spindle assembly | BP | 52 | 7 | 0 | 0.00022 |
| GO:0050672 | negative regulation of lymphocyte proliferation | BP | 52 | 7 | 2 | 0.00022 |
| GO:0032945 | negative regulation of mononuclear cell proliferation | BP | 52 | 7 | 2 | 0.00022 |
| GO:0002449 | lymphocyte mediated immunity | BP | 171 | 13 | 2 | 0.00024 |
| GO:0030595 | leukocyte chemotaxis | BP | 149 | 12 | 10 | 0.00024 |
| GO:0007051 | spindle organization | BP | 149 | 12 | 1 | 0.00024 |
| GO:0050900 | leukocyte migration | BP | 319 | 19 | 16 | 0.00025 |
| GO:0007091 | metaphase/anaphase transition of mitotic cell cycle | BP | 53 | 7 | 2 | 0.00025 |
| GO:0046651 | lymphocyte proliferation | BP | 195 | 14 | 7 | 0.00025 |
| GO:0023024 | MHC class I protein complex binding | MF | 6 | 3 | 0 | 0.00026 |
| GO:0051383 | kinetochore organization | BP | 14 | 4 | 0 | 0.00027 |
| GO:0002683 | negative regulation of immune system process | BP | 348 | 20 | 15 | 0.00027 |
| GO:0006915 | apoptotic process | BP | 1539 | 58 | 43 | 0.00028 |

| | | | | | | |
|---|---|---|---|---|---|---|
| GO:0072073 | kidney epithelium development | BP | 109 | 10 | 2 | 0.00028 |
| GO:0032943 | mononuclear cell proliferation | BP | 197 | 14 | 8 | 0.00028 |
| GO:0044784 | metaphase/anaphase transition of cell cycle | BP | 54 | 7 | 2 | 0.00028 |
| GO:2000026 | regulation of multicellular organismal development | BP | 1576 | 59 | 68 | 0.00029 |
| GO:0050670 | regulation of lymphocyte proliferation | BP | 152 | 12 | 6 | 0.00029 |
| GO:0030855 | epithelial cell differentiation | BP | 461 | 24 | 21 | 0.0003 |
| GO:0032944 | regulation of mononuclear cell proliferation | BP | 153 | 12 | 6 | 0.00031 |
| GO:0010965 | regulation of mitotic sister chromatid separation | BP | 55 | 7 | 2 | 0.00032 |
| GO:0002228 | natural killer cell mediated immunity | BP | 40 | 6 | 0 | 0.00035 |
| GO:0099513 | polymeric cytoskeletal fiber | CC | 494 | 25 | 18 | 0.00035 |
| GO:0051704 | multi-organism process | BP | 1833 | 66 | 50 | 0.00036 |
| GO:0070664 | negative regulation of leukocyte proliferation | BP | 56 | 7 | 3 | 0.00036 |
| GO:0038023 | signaling receptor activity | MF | 675 | 31 | 40 | 0.00038 |
| GO:0048002 | antigen processing and presentation of peptide antigen | BP | 157 | 12 | 5 | 0.00039 |
| GO:0034614 | cellular response to reactive oxygen species | BP | 135 | 11 | 5 | 0.0004 |
| GO:0030261 | chromosome condensation | BP | 27 | 5 | 0 | 0.0004 |
| GO:1901607 | alpha-amino acid biosynthetic process | BP | 57 | 7 | 6 | 0.0004 |
| GO:0072028 | nephron morphogenesis | BP | 57 | 7 | 2 | 0.0004 |
| GO:0031341 | regulation of cell killing | BP | 57 | 7 | 2 | 0.0004 |
| GO:0048870 | cell motility | BP | 1223 | 48 | 58 | 0.00041 |
| GO:0051674 | localization of cell | BP | 1223 | 48 | 58 | 0.00041 |
| GO:0071621 | granulocyte chemotaxis | BP | 75 | 8 | 3 | 0.00041 |
| GO:0001657 | ureteric bud development | BP | 75 | 8 | 2 | 0.00041 |
| GO:0042110 | T cell activation | BP | 360 | 20 | 10 | 0.00042 |
| GO:0043067 | regulation of programmed cell death | BP | 1226 | 48 | 34 | 0.00043 |
| GO:0060326 | cell chemotaxis | BP | 206 | 14 | 12 | 0.00044 |
| GO:0033018 | sarcoplasmic reticulum lumen | CC | 7 | 3 | 1 | 0.00044 |
| GO:0051306 | mitotic sister chromatid separation | BP | 58 | 7 | 2 | 0.00044 |
| GO:0007155 | cell adhesion | BP | 1095 | 44 | 51 | 0.00045 |
| GO:0007159 | leukocyte cell-cell adhesion | BP | 256 | 16 | 8 | 0.00045 |
| GO:0072163 | mesonephric epithelium development | BP | 76 | 8 | 2 | 0.00045 |
| GO:0072164 | mesonephric tubule development | BP | 76 | 8 | 2 | 0.00045 |
| GO:0050830 | defense response to Gram-positive bacterium | BP | 42 | 6 | 1 | 0.00045 |
| GO:0035272 | exocrine system development | BP | 42 | 6 | 1 | 0.00045 |
| GO:0048732 | gland development | BP | 335 | 19 | 20 | 0.00046 |
| GO:0001818 | negative regulation of cytokine production | BP | 207 | 14 | 8 | 0.00046 |
| GO:0022610 | biological adhesion | BP | 1099 | 44 | 52 | 0.00048 |
| GO:0042493 | response to drug | BP | 778 | 34 | 42 | 0.00049 |
| GO:1905818 | regulation of chromosome separation | BP | 59 | 7 | 2 | 0.00049 |
| GO:0045765 | regulation of angiogenesis | BP | 233 | 15 | 14 | 0.00049 |
| GO:0000776 | kinetochore | CC | 118 | 10 | 0 | 0.00053 |
| GO:1901342 | regulation of vasculature development | BP | 260 | 16 | 14 | 0.00053 |
| GO:1902105 | regulation of leukocyte differentiation | BP | 210 | 14 | 3 | 0.00054 |
| GO:0001823 | mesonephros development | BP | 78 | 8 | 2 | 0.00054 |
| GO:0019882 | antigen processing and presentation | BP | 186 | 13 | 5 | 0.00054 |
| GO:0005887 | integral component of plasma membrane | CC | 1072 | 43 | 81 | 0.00054 |
| GO:0044433 | cytoplasmic vesicle part | CC | 1205 | 47 | 65 | 0.00054 |
| GO:0030593 | neutrophil chemotaxis | BP | 60 | 7 | 3 | 0.00055 |
| GO:0051321 | meiotic cell cycle | BP | 163 | 12 | 5 | 0.00055 |
| GO:0002706 | regulation of lymphocyte mediated immunity | BP | 98 | 9 | 2 | 0.00056 |
| GO:0001912 | positive regulation of leukocyte mediated cytotoxicity | BP | 29 | 5 | 1 | 0.00056 |
| GO:0042542 | response to hydrogen peroxide | BP | 119 | 10 | 4 | 0.00057 |
| GO:0043316 | cytotoxic T cell degranulation | BP | 2 | 2 | 0 | 0.00057 |
| GO:0009893 | positive regulation of metabolic process | BP | 2817 | 92 | 116 | 0.00059 |
| GO:0042981 | regulation of apoptotic process | BP | 1210 | 47 | 34 | 0.00059 |
| GO:2000738 | positive regulation of stem cell differentiation | BP | 17 | 4 | 0 | 0.0006 |
| GO:0000302 | response to reactive oxygen species | BP | 189 | 13 | 7 | 0.00063 |
| GO:0048869 | cellular developmental process | BP | 3128 | 100 | 131 | 0.00066 |
| GO:0032652 | regulation of interleukin-1 production | BP | 62 | 7 | 1 | 0.00067 |
| GO:0000922 | spindle pole | CC | 144 | 11 | 3 | 0.00068 |
| GO:0051304 | chromosome separation | BP | 81 | 8 | 2 | 0.00069 |
| GO:0072009 | nephron epithelium development | BP | 81 | 8 | 2 | 0.00069 |
| GO:0016175 | superoxide-generating NADPH oxidase activity | MF | 8 | 3 | 0 | 0.0007 |
| GO:0001819 | positive regulation of cytokine production | BP | 347 | 19 | 9 | 0.0007 |

| | | | | | | |
|---|---|---|---|---|---|---|
| GO:0051249 | regulation of lymphocyte activation | BP | 320 | 18 | 9 | 0.00071 |
| GO:0000775 | chromosome, centromeric region | CC | 168 | 12 | 0 | 0.00072 |
| GO:0070820 | tertiary granule | CC | 123 | 10 | 4 | 0.00074 |
| GO:0001655 | urogenital system development | BP | 268 | 16 | 10 | 0.00074 |
| GO:0098641 | cadherin binding involved in cell-cell adhesion | MF | 18 | 4 | 0 | 0.00075 |
| GO:0030277 | maintenance of gastrointestinal epithelium | BP | 18 | 4 | 0 | 0.00075 |
| GO:0042629 | mast cell granule | CC | 18 | 4 | 0 | 0.00075 |
| GO:0010818 | T cell chemotaxis | BP | 18 | 4 | 0 | 0.00075 |
| GO:0002478 | antigen processing and presentation of exogenous peptide antigen | BP | 146 | 11 | 2 | 0.00077 |
| GO:0071634 | regulation of transforming growth factor beta production | BP | 31 | 5 | 0 | 0.00077 |
| GO:0017144 | drug metabolic process | BP | 613 | 28 | 57 | 0.0008 |
| GO:0042277 | peptide binding | MF | 194 | 13 | 17 | 0.0008 |
| GO:1900407 | regulation of cellular response to oxidative stress | BP | 64 | 7 | 1 | 0.00081 |
| GO:0051276 | chromosome organization | BP | 962 | 39 | 19 | 0.00082 |
| GO:0000228 | nuclear chromosome | CC | 465 | 23 | 2 | 0.00082 |
| GO:0031226 | intrinsic component of plasma membrane | CC | 1128 | 44 | 86 | 0.00083 |
| GO:0042802 | identical protein binding | MF | 1365 | 51 | 59 | 0.00083 |
| GO:0019884 | antigen processing and presentation of exogenous antigen | BP | 148 | 11 | 2 | 0.00086 |
| GO:0033047 | regulation of mitotic sister chromatid segregation | BP | 65 | 7 | 2 | 0.00089 |
| GO:0050704 | regulation of interleukin-1 secretion | BP | 32 | 5 | 1 | 0.0009 |
| GO:0060142 | regulation of syncytium formation by plasma membrane fusion | BP | 19 | 4 | 1 | 0.00094 |
| GO:0050867 | positive regulation of cell activation | BP | 248 | 15 | 7 | 0.00094 |
| GO:0045839 | negative regulation of mitotic nuclear division | BP | 48 | 6 | 1 | 0.00094 |
| GO:0005875 | microtubule associated complex | CC | 127 | 10 | 6 | 0.00094 |
| GO:0002694 | regulation of leukocyte activation | BP | 384 | 20 | 12 | 0.00095 |
| GO:0007093 | mitotic cell cycle checkpoint | BP | 150 | 11 | 3 | 0.00096 |
| GO:0042100 | B cell proliferation | BP | 66 | 7 | 3 | 0.00098 |
| GO:0001101 | response to acid chemical | BP | 276 | 16 | 14 | 0.00101 |
| GO:2000377 | regulation of reactive oxygen species metabolic process | BP | 151 | 11 | 5 | 0.00102 |
| GO:0000940 | condensed chromosome outer kinetochore | CC | 9 | 3 | 0 | 0.00102 |
| GO:0007094 | mitotic spindle assembly checkpoint | BP | 33 | 5 | 1 | 0.00104 |
| GO:0071174 | mitotic spindle checkpoint | BP | 33 | 5 | 1 | 0.00104 |
| GO:0071173 | spindle assembly checkpoint | BP | 33 | 5 | 1 | 0.00104 |
| GO:0031577 | spindle checkpoint | BP | 33 | 5 | 1 | 0.00104 |
| GO:0071604 | transforming growth factor beta production | BP | 33 | 5 | 0 | 0.00104 |
| GO:0006323 | DNA packaging | BP | 107 | 9 | 0 | 0.00105 |
| GO:0051093 | negative regulation of developmental process | BP | 718 | 31 | 27 | 0.00106 |
| GO:0030545 | receptor regulator activity | MF | 251 | 15 | 14 | 0.00106 |
| GO:0008652 | cellular amino acid biosynthetic process | BP | 67 | 7 | 8 | 0.00107 |
| GO:0003777 | microtubule motor activity | MF | 67 | 7 | 5 | 0.00107 |
| GO:0002708 | positive regulation of lymphocyte mediated immunity | BP | 67 | 7 | 0 | 0.00107 |
| GO:0098657 | import into cell | BP | 687 | 30 | 47 | 0.00107 |
| GO:0060089 | molecular transducer activity | MF | 719 | 31 | 43 | 0.00108 |
| GO:0032496 | response to lipopolysaccharide | BP | 252 | 15 | 13 | 0.00111 |
| GO:1903707 | negative regulation of hemopoiesis | BP | 108 | 9 | 3 | 0.00113 |
| GO:0050865 | regulation of cell activation | BP | 418 | 21 | 12 | 0.00113 |
| GO:0045597 | positive regulation of cell differentiation | BP | 753 | 32 | 33 | 0.00114 |
| GO:0032103 | positive regulation of response to external stimulus | BP | 227 | 14 | 7 | 0.00115 |
| GO:0045785 | positive regulation of cell adhesion | BP | 334 | 18 | 13 | 0.00116 |
| GO:0072080 | nephron tubule development | BP | 68 | 7 | 2 | 0.00117 |
| GO:0030888 | regulation of B cell proliferation | BP | 50 | 6 | 3 | 0.00117 |
| GO:0005876 | spindle microtubule | CC | 50 | 6 | 0 | 0.00117 |
| GO:1901987 | regulation of cell cycle phase transition | BP | 391 | 20 | 6 | 0.00119 |
| GO:0030667 | secretory granule membrane | CC | 228 | 14 | 12 | 0.0012 |
| GO:0045088 | regulation of innate immune response | BP | 363 | 19 | 9 | 0.0012 |
| GO:0030496 | midbody | CC | 155 | 11 | 3 | 0.00126 |
| GO:0031349 | positive regulation of defense response | BP | 393 | 20 | 13 | 0.00126 |
| GO:0043069 | negative regulation of programmed cell death | BP | 696 | 30 | 19 | 0.00131 |
| GO:0045930 | negative regulation of mitotic cell cycle | BP | 283 | 16 | 6 | 0.00131 |
| GO:0045937 | positive regulation of phosphate metabolic process | BP | 856 | 35 | 41 | 0.00132 |
| GO:0010562 | positive regulation of phosphorus metabolic process | BP | 856 | 35 | 41 | 0.00132 |
| GO:0002285 | lymphocyte activation involved in immune response | BP | 133 | 10 | 2 | 0.00134 |
| GO:0045787 | positive regulation of cell cycle | BP | 311 | 17 | 10 | 0.00135 |
| GO:0048018 | receptor ligand activity | MF | 231 | 14 | 13 | 0.00135 |

| | | | | | | |
|---|---|---|---|---|---|---|
| GO:1903037 | regulation of leukocyte cell-cell adhesion | BP | 231 | 14 | 7 | 0.00135 |
| GO:1902100 | negative regulation of metaphase/anaphase transition of cell cycle | BP | 35 | 5 | 1 | 0.00137 |
| GO:0045841 | negative regulation of mitotic metaphase/anaphase transition | BP | 35 | 5 | 1 | 0.00137 |
| GO:0051234 | establishment of localization | BP | 4049 | 122 | 233 | 0.00138 |
| GO:0061326 | renal tubule development | BP | 70 | 7 | 2 | 0.00139 |
| GO:0097530 | granulocyte migration | BP | 90 | 8 | 3 | 0.00139 |
| GO:0002705 | positive regulation of leukocyte mediated immunity | BP | 90 | 8 | 1 | 0.00139 |
| GO:1903409 | reactive oxygen species biosynthetic process | BP | 90 | 8 | 5 | 0.00139 |
| GO:0005732 | small nucleolar ribonucleoprotein complex | CC | 21 | 4 | 0 | 0.0014 |
| GO:0048871 | multicellular organismal homeostasis | BP | 368 | 19 | 20 | 0.0014 |
| GO:0005874 | microtubule | CC | 340 | 18 | 13 | 0.00141 |
| GO:0060205 | cytoplasmic vesicle lumen | CC | 285 | 16 | 10 | 0.00141 |
| GO:0036500 | ATF6-mediated unfolded protein response | BP | 10 | 3 | 0 | 0.00144 |
| GO:0006563 | L-serine metabolic process | BP | 10 | 3 | 1 | 0.00144 |
| GO:0051784 | negative regulation of nuclear division | BP | 52 | 6 | 1 | 0.00144 |
| GO:0016477 | cell migration | BP | 1126 | 43 | 55 | 0.00145 |
| GO:1901990 | regulation of mitotic cell cycle phase transition | BP | 369 | 19 | 5 | 0.00145 |
| GO:0072006 | nephron development | BP | 112 | 9 | 4 | 0.00146 |
| GO:0031325 | positive regulation of cellular metabolic process | BP | 2594 | 84 | 103 | 0.00146 |
| GO:0031983 | vesicle lumen | CC | 286 | 16 | 10 | 0.00146 |
| GO:0032501 | multicellular organismal process | BP | 5056 | 147 | 233 | 0.00149 |
| GO:0060993 | kidney morphogenesis | BP | 71 | 7 | 2 | 0.00151 |
| GO:1902882 | regulation of response to oxidative stress | BP | 71 | 7 | 2 | 0.00151 |
| GO:0046677 | response to antibiotic | BP | 260 | 15 | 8 | 0.00151 |
| GO:0004888 | transmembrane signaling receptor activity | MF | 517 | 24 | 29 | 0.00151 |
| GO:0002521 | leukocyte differentiation | BP | 399 | 20 | 8 | 0.00151 |
| GO:0048247 | lymphocyte chemotaxis | BP | 36 | 5 | 0 | 0.00156 |
| GO:0098687 | chromosomal region | CC | 288 | 16 | 0 | 0.00157 |
| GO:0097529 | myeloid leukocyte migration | BP | 136 | 10 | 9 | 0.00159 |
| GO:0042113 | B cell activation | BP | 184 | 12 | 3 | 0.00159 |
| GO:0072078 | nephron tubule morphogenesis | BP | 53 | 6 | 2 | 0.0016 |
| GO:1903201 | regulation of oxidative stress-induced cell death | BP | 53 | 6 | 0 | 0.0016 |
| GO:1990266 | neutrophil migration | BP | 72 | 7 | 3 | 0.00163 |
| GO:0051251 | positive regulation of lymphocyte activation | BP | 210 | 13 | 6 | 0.00165 |
| GO:0035295 | tube development | BP | 835 | 34 | 39 | 0.00165 |
| GO:0030515 | snoRNA binding | MF | 22 | 4 | 0 | 0.00167 |
| GO:0097149 | centralspindlin complex | CC | 3 | 2 | 0 | 0.00169 |
| GO:0036118 | hyaluranon cable assembly | BP | 3 | 2 | 0 | 0.00169 |
| GO:0036438 | maintenance of lens transparency | BP | 3 | 2 | 0 | 0.00169 |
| GO:1904746 | negative regulation of apoptotic process involved in development | BP | 3 | 2 | 0 | 0.00169 |
| GO:1902338 | negative regulation of apoptotic process involved in morphogenesis | BP | 3 | 2 | 0 | 0.00169 |
| GO:1903974 | positive regulation of cellular response to macrophage colony-stimu | BP | 3 | 2 | 0 | 0.00169 |
| GO:1900106 | positive regulation of hyaluranon cable assembly | BP | 3 | 2 | 0 | 0.00169 |
| GO:2001206 | positive regulation of osteoclast development | BP | 3 | 2 | 0 | 0.00169 |
| GO:1903971 | positive regulation of response to macrophage colony-stimulating f | BP | 3 | 2 | 0 | 0.00169 |
| GO:0032954 | regulation of cytokinetic process | BP | 3 | 2 | 0 | 0.00169 |
| GO:1900104 | regulation of hyaluranon cable assembly | BP | 3 | 2 | 0 | 0.00169 |
| GO:0001810 | regulation of type I hypersensitivity | BP | 3 | 2 | 0 | 0.00169 |
| GO:0016068 | type I hypersensitivity | BP | 3 | 2 | 0 | 0.00169 |
| GO:0051225 | spindle assembly | BP | 93 | 8 | 0 | 0.00172 |
| GO:1905819 | negative regulation of chromosome separation | BP | 37 | 5 | 1 | 0.00177 |
| GO:2000816 | negative regulation of mitotic sister chromatid separation | BP | 37 | 5 | 1 | 0.00177 |
| GO:0031343 | positive regulation of cell killing | BP | 37 | 5 | 2 | 0.00177 |
| GO:0001889 | liver development | BP | 116 | 9 | 7 | 0.00186 |
| GO:0002696 | positive regulation of leukocyte activation | BP | 239 | 14 | 7 | 0.00187 |
| GO:0071103 | DNA conformation change | BP | 213 | 13 | 1 | 0.00187 |
| GO:0001894 | tissue homeostasis | BP | 163 | 11 | 6 | 0.00188 |
| GO:0032612 | interleukin-1 production | BP | 74 | 7 | 1 | 0.00192 |
| GO:0072088 | nephron epithelium morphogenesis | BP | 55 | 6 | 2 | 0.00194 |
| GO:0006928 | movement of cell or subcellular component | BP | 1595 | 56 | 73 | 0.00194 |
| GO:0043020 | NADPH oxidase complex | CC | 11 | 3 | 0 | 0.00194 |
| GO:0045346 | regulation of MHC class II biosynthetic process | BP | 11 | 3 | 0 | 0.00194 |
| GO:0043066 | negative regulation of apoptotic process | BP | 683 | 29 | 19 | 0.00197 |
| GO:0098805 | whole membrane | CC | 1351 | 49 | 71 | 0.00199 |

| | | | | | | |
|---|---|---|---|---|---|---|
| GO:0006884 | cell volume homeostasis | BP | 23 | 4 | 2 | 0.00199 |
| GO:0050863 | regulation of T cell activation | BP | 241 | 14 | 6 | 0.00202 |
| GO:0010647 | positive regulation of cell communication | BP | 1422 | 51 | 59 | 0.00202 |
| GO:0031401 | positive regulation of protein modification process | BP | 945 | 37 | 42 | 0.00206 |
| GO:0050878 | regulation of body fluid levels | BP | 381 | 19 | 19 | 0.00209 |
| GO:0022414 | reproductive process | BP | 946 | 37 | 36 | 0.00209 |
| GO:0050729 | positive regulation of inflammatory response | BP | 96 | 8 | 5 | 0.0021 |
| GO:0032675 | regulation of interleukin-6 production | BP | 96 | 8 | 4 | 0.0021 |
| GO:0045596 | negative regulation of cell differentiation | BP | 531 | 24 | 21 | 0.00215 |
| GO:0072001 | renal system development | BP | 243 | 14 | 10 | 0.00218 |
| GO:0000003 | reproduction | BP | 949 | 37 | 36 | 0.00221 |
| GO:0023056 | positive regulation of signaling | BP | 1428 | 51 | 61 | 0.00221 |
| GO:0030154 | cell differentiation | BP | 2971 | 93 | 125 | 0.00221 |
| GO:0098552 | side of membrane | CC | 383 | 19 | 19 | 0.00221 |
| GO:0061008 | hepaticobiliary system development | BP | 119 | 9 | 7 | 0.00222 |
| GO:0034767 | positive regulation of ion transmembrane transport | BP | 119 | 9 | 6 | 0.00222 |
| GO:0042136 | neurotransmitter biosynthetic process | BP | 76 | 7 | 7 | 0.00224 |
| GO:0033045 | regulation of sister chromatid segregation | BP | 76 | 7 | 2 | 0.00224 |
| GO:0005882 | intermediate filament | CC | 57 | 6 | 1 | 0.00233 |
| GO:0061333 | renal tubule morphogenesis | BP | 57 | 6 | 2 | 0.00233 |
| GO:0099512 | supramolecular fiber | CC | 659 | 28 | 30 | 0.00233 |
| GO:0010669 | epithelial structure maintenance | BP | 24 | 4 | 0 | 0.00234 |
| GO:0042742 | defense response to bacterium | BP | 120 | 9 | 6 | 0.00235 |
| GO:0050801 | ion homeostasis | BP | 598 | 26 | 38 | 0.00243 |
| GO:0044283 | small molecule biosynthetic process | BP | 661 | 28 | 56 | 0.00243 |
| GO:0051130 | positive regulation of cellular component organization | BP | 1056 | 40 | 43 | 0.00247 |
| GO:0001934 | positive regulation of protein phosphorylation | BP | 758 | 31 | 33 | 0.00248 |
| GO:0005126 | cytokine receptor binding | MF | 169 | 11 | 6 | 0.0025 |
| GO:0048523 | negative regulation of cellular process | BP | 3678 | 111 | 130 | 0.0025 |
| GO:0098632 | cell-cell adhesion mediator activity | MF | 40 | 5 | 1 | 0.00252 |
| GO:0140253 | cell-cell fusion | BP | 40 | 5 | 4 | 0.00252 |
| GO:0050701 | interleukin-1 secretion | BP | 40 | 5 | 1 | 0.00252 |
| GO:0033048 | negative regulation of mitotic sister chromatid segregation | BP | 40 | 5 | 1 | 0.00252 |
| GO:0000768 | syncytium formation by plasma membrane fusion | BP | 40 | 5 | 4 | 0.00252 |
| GO:0040011 | locomotion | BP | 1402 | 50 | 67 | 0.00253 |
| GO:0048878 | chemical homeostasis | BP | 890 | 35 | 63 | 0.00253 |
| GO:0045342 | MHC class II biosynthetic process | BP | 12 | 3 | 0 | 0.00254 |
| GO:0010819 | regulation of T cell chemotaxis | BP | 12 | 3 | 0 | 0.00254 |
| GO:0051775 | response to redox state | BP | 12 | 3 | 0 | 0.00254 |
| GO:0005515 | protein binding | MF | 9002 | 238 | 354 | 0.00269 |
| GO:0000075 | cell cycle checkpoint | BP | 196 | 12 | 3 | 0.0027 |
| GO:0099081 | supramolecular polymer | CC | 666 | 28 | 30 | 0.00271 |
| GO:0003823 | antigen binding | MF | 25 | 4 | 3 | 0.00273 |
| GO:0051247 | positive regulation of protein metabolic process | BP | 1338 | 48 | 53 | 0.00276 |
| GO:0099080 | supramolecular complex | CC | 667 | 28 | 30 | 0.00276 |
| GO:0045132 | meiotic chromosome segregation | BP | 59 | 6 | 1 | 0.00278 |
| GO:0008285 | negative regulation of cell proliferation | BP | 542 | 24 | 20 | 0.0028 |
| GO:0001658 | branching involved in ureteric bud morphogenesis | BP | 41 | 5 | 1 | 0.00281 |
| GO:0070192 | chromosome organization involved in meiotic cell cycle | BP | 41 | 5 | 0 | 0.00281 |
| GO:0030858 | positive regulation of epithelial cell differentiation | BP | 41 | 5 | 2 | 0.00281 |
| GO:0090342 | regulation of cell aging | BP | 41 | 5 | 1 | 0.00281 |
| GO:0001664 | G protein-coupled receptor binding | MF | 172 | 11 | 9 | 0.00287 |
| GO:1903706 | regulation of hemopoiesis | BP | 363 | 18 | 8 | 0.00288 |
| GO:0005102 | signaling receptor binding | MF | 1066 | 40 | 53 | 0.00292 |
| GO:0009967 | positive regulation of signal transduction | BP | 1307 | 47 | 50 | 0.00292 |
| GO:0060249 | anatomical structure homeostasis | BP | 335 | 17 | 8 | 0.00295 |
| GO:0001666 | response to hypoxia | BP | 307 | 16 | 16 | 0.00299 |
| GO:0019886 | antigen processing and presentation of exogenous peptide antigen v | BP | 80 | 7 | 2 | 0.003 |
| GO:0002312 | B cell activation involved in immune response | BP | 60 | 6 | 0 | 0.00303 |
| GO:0048646 | anatomical structure formation involved in morphogenesis | BP | 867 | 34 | 44 | 0.00304 |
| GO:0009991 | response to extracellular stimulus | BP | 424 | 20 | 39 | 0.00306 |
| GO:0016020 | membrane | CC | 6503 | 180 | 347 | 0.0031 |
| GO:0051985 | negative regulation of chromosome segregation | BP | 42 | 5 | 2 | 0.00313 |
| GO:0033046 | negative regulation of sister chromatid segregation | BP | 42 | 5 | 2 | 0.00313 |

| | | | | | | |
|---|---|---|---|---|---|---|
| GO:0006949 | syncytium formation | BP | 42 | 5 | 4 | 0.00313 |
| GO:0070482 | response to oxygen levels | BP | 337 | 17 | 17 | 0.00314 |
| GO:0032309 | icosanoid secretion | BP | 26 | 4 | 6 | 0.00317 |
| GO:0050673 | epithelial cell proliferation | BP | 309 | 16 | 13 | 0.00319 |
| GO:0046983 | protein dimerization activity | MF | 970 | 37 | 47 | 0.0032 |
| GO:0002495 | antigen processing and presentation of peptide antigen via MHC cl | BP | 81 | 7 | 2 | 0.00322 |
| GO:0000778 | condensed nuclear chromosome kinetochore | CC | 13 | 3 | 0 | 0.00325 |
| GO:0050664 | oxidoreductase activity, acting on NAD(P)H, oxygen as acceptor | MF | 13 | 3 | 0 | 0.00325 |
| GO:0019763 | immunoglobulin receptor activity | MF | 4 | 2 | 0 | 0.00332 |
| GO:0072131 | kidney mesenchyme morphogenesis | BP | 4 | 2 | 0 | 0.00332 |
| GO:0006565 | L-serine catabolic process | BP | 4 | 2 | 1 | 0.00332 |
| GO:0001875 | lipopolysaccharide receptor activity | MF | 4 | 2 | 1 | 0.00332 |
| GO:0072133 | metanephric mesenchyme morphogenesis | BP | 4 | 2 | 0 | 0.00332 |
| GO:0007079 | mitotic chromosome movement towards spindle pole | BP | 4 | 2 | 0 | 0.00332 |
| GO:0002317 | plasma cell differentiation | BP | 4 | 2 | 0 | 0.00332 |
| GO:0032661 | regulation of interleukin-18 production | BP | 4 | 2 | 0 | 0.00332 |
| GO:0030485 | smooth muscle contractile fiber | CC | 4 | 2 | 0 | 0.00332 |
| GO:0061134 | peptidase regulator activity | MF | 127 | 9 | 3 | 0.00344 |
| GO:0002504 | antigen processing and presentation of peptide or polysaccharide ar | BP | 82 | 7 | 2 | 0.00345 |
| GO:0002377 | immunoglobulin production | BP | 82 | 7 | 1 | 0.00345 |
| GO:1903428 | positive regulation of reactive oxygen species biosynthetic process | BP | 43 | 5 | 3 | 0.00347 |
| GO:0090183 | regulation of kidney development | BP | 43 | 5 | 2 | 0.00347 |
| GO:0001822 | kidney development | BP | 230 | 13 | 10 | 0.00364 |
| GO:0032467 | positive regulation of cytokinesis | BP | 27 | 4 | 1 | 0.00365 |
| GO:0032635 | interleukin-6 production | BP | 105 | 8 | 5 | 0.00368 |
| GO:1901216 | positive regulation of neuron death | BP | 83 | 7 | 1 | 0.00369 |
| GO:0035239 | tube morphogenesis | BP | 682 | 28 | 31 | 0.00377 |
| GO:0045595 | regulation of cell differentiation | BP | 1396 | 49 | 57 | 0.00384 |
| GO:0055081 | anion homeostasis | BP | 44 | 5 | 8 | 0.00384 |
| GO:0006984 | ER-nucleus signaling pathway | BP | 44 | 5 | 4 | 0.00384 |
| GO:0002456 | T cell mediated immunity | BP | 63 | 6 | 2 | 0.00387 |
| GO:0051250 | negative regulation of lymphocyte activation | BP | 106 | 8 | 4 | 0.0039 |
| GO:0051246 | regulation of protein metabolic process | BP | 2196 | 71 | 78 | 0.00394 |
| GO:0046717 | acid secretion | BP | 84 | 7 | 8 | 0.00394 |
| GO:0036293 | response to decreased oxygen levels | BP | 316 | 16 | 17 | 0.00396 |
| GO:0048469 | cell maturation | BP | 130 | 9 | 7 | 0.00402 |
| GO:0002695 | negative regulation of leukocyte activation | BP | 130 | 9 | 6 | 0.00402 |
| GO:0045931 | positive regulation of mitotic cell cycle | BP | 130 | 9 | 3 | 0.00402 |
| GO:0043270 | positive regulation of ion transport | BP | 206 | 12 | 13 | 0.00403 |
| GO:0015630 | microtubule cytoskeleton | CC | 984 | 37 | 23 | 0.00405 |
| GO:0007584 | response to nutrient | BP | 180 | 11 | 14 | 0.00405 |
| GO:0034185 | apolipoprotein binding | MF | 14 | 3 | 1 | 0.00406 |
| GO:0071682 | endocytic vesicle lumen | CC | 14 | 3 | 2 | 0.00406 |
| GO:0050665 | hydrogen peroxide biosynthetic process | BP | 14 | 3 | 0 | 0.00406 |
| GO:0030889 | negative regulation of B cell proliferation | BP | 14 | 3 | 0 | 0.00406 |
| GO:0032693 | negative regulation of interleukin-10 production | BP | 14 | 3 | 1 | 0.00406 |
| GO:0038187 | pattern recognition receptor activity | MF | 14 | 3 | 2 | 0.00406 |
| GO:0090266 | regulation of mitotic cell cycle spindle assembly checkpoint | BP | 14 | 3 | 1 | 0.00406 |
| GO:1903504 | regulation of mitotic spindle checkpoint | BP | 14 | 3 | 1 | 0.00406 |
| GO:0090231 | regulation of spindle checkpoint | BP | 14 | 3 | 1 | 0.00406 |
| GO:0009070 | serine family amino acid biosynthetic process | BP | 14 | 3 | 1 | 0.00406 |
| GO:0008329 | signaling pattern recognition receptor activity | MF | 14 | 3 | 2 | 0.00406 |
| GO:0022407 | regulation of cell-cell adhesion | BP | 317 | 16 | 11 | 0.00409 |
| GO:0045730 | respiratory burst | BP | 28 | 4 | 1 | 0.00418 |
| GO:0001540 | amyloid-beta binding | MF | 64 | 6 | 5 | 0.00419 |
| GO:0045638 | negative regulation of myeloid cell differentiation | BP | 64 | 6 | 1 | 0.00419 |
| GO:0048146 | positive regulation of fibroblast proliferation | BP | 45 | 5 | 0 | 0.00424 |
| GO:0034764 | positive regulation of transmembrane transport | BP | 156 | 10 | 8 | 0.00431 |
| GO:0002440 | production of molecular mediator of immune response | BP | 156 | 10 | 6 | 0.00431 |
| GO:0090068 | positive regulation of cell cycle process | BP | 236 | 13 | 4 | 0.00453 |
| GO:0002700 | regulation of production of molecular mediator of immune respons | BP | 109 | 8 | 5 | 0.00462 |
| GO:0030097 | hemopoiesis | BP | 693 | 28 | 19 | 0.00469 |
| GO:0005539 | glycosaminoglycan binding | MF | 158 | 10 | 15 | 0.00472 |
| GO:0098609 | cell-cell adhesion | BP | 629 | 26 | 25 | 0.00475 |

| | | | | | | |
|---|---|---|---|---|---|---|
| GO:0050706 | regulation of interleukin-1 beta secretion | BP | 29 | 4 | 1 | 0.00476 |
| GO:0002761 | regulation of myeloid leukocyte differentiation | BP | 87 | 7 | 1 | 0.00479 |
| GO:0006897 | endocytosis | BP | 598 | 25 | 38 | 0.00485 |
| GO:0002460 | adaptive immune response based on somatic recombination of imm | BP | 185 | 11 | 4 | 0.00498 |
| GO:0019731 | antibacterial humoral response | BP | 15 | 3 | 0 | 0.00499 |
| GO:0001848 | complement binding | MF | 15 | 3 | 0 | 0.00499 |
| GO:0015037 | peptide disulfide oxidoreductase activity | MF | 15 | 3 | 0 | 0.00499 |
| GO:0004653 | polypeptide N-acetylgalactosaminyltransferase activity | MF | 15 | 3 | 1 | 0.00499 |
| GO:1901739 | regulation of myoblast fusion | BP | 15 | 3 | 0 | 0.00499 |
| GO:0001914 | regulation of T cell mediated cytotoxicity | BP | 15 | 3 | 0 | 0.00499 |
| GO:0009314 | response to radiation | BP | 353 | 17 | 17 | 0.005 |
| GO:0051128 | regulation of cellular component organization | BP | 2069 | 67 | 85 | 0.00502 |
| GO:0051129 | negative regulation of cellular component organization | BP | 600 | 25 | 21 | 0.00506 |
| GO:0050678 | regulation of epithelial cell proliferation | BP | 267 | 14 | 9 | 0.00506 |
| GO:0031410 | cytoplasmic vesicle | CC | 1849 | 61 | 100 | 0.00506 |
| GO:0090303 | positive regulation of wound healing | BP | 47 | 5 | 1 | 0.00512 |
| GO:0002637 | regulation of immunoglobulin production | BP | 47 | 5 | 0 | 0.00512 |
| GO:0060675 | ureteric bud morphogenesis | BP | 47 | 5 | 2 | 0.00512 |
| GO:0003774 | motor activity | MF | 111 | 8 | 7 | 0.00515 |
| GO:0097708 | intracellular vesicle | CC | 1851 | 61 | 100 | 0.00518 |
| GO:0033218 | amide binding | MF | 240 | 13 | 22 | 0.0052 |
| GO:0000281 | mitotic cytokinesis | BP | 67 | 6 | 1 | 0.00525 |
| GO:2000106 | regulation of leukocyte apoptotic process | BP | 67 | 6 | 2 | 0.00525 |
| GO:1901571 | fatty acid derivative transport | BP | 30 | 4 | 6 | 0.0054 |
| GO:0071715 | icosanoid transport | BP | 30 | 4 | 6 | 0.0054 |
| GO:0042554 | superoxide anion generation | BP | 30 | 4 | 0 | 0.0054 |
| GO:0045766 | positive regulation of angiogenesis | BP | 136 | 9 | 7 | 0.0054 |
| GO:0102336 | 3-oxo-arachidoyl-CoA synthase activity | MF | 5 | 2 | 0 | 0.00544 |
| GO:0102337 | 3-oxo-cerotoyl-CoA synthase activity | MF | 5 | 2 | 0 | 0.00544 |
| GO:0102338 | 3-oxo-lignoceronyl-CoA synthase activity | MF | 5 | 2 | 0 | 0.00544 |
| GO:0002322 | B cell proliferation involved in immune response | BP | 5 | 2 | 0 | 0.00544 |
| GO:0042262 | DNA protection | BP | 5 | 2 | 0 | 0.00544 |
| GO:0009922 | fatty acid elongase activity | MF | 5 | 2 | 0 | 0.00544 |
| GO:0034625 | fatty acid elongation, monounsaturated fatty acid | BP | 5 | 2 | 0 | 0.00544 |
| GO:0034626 | fatty acid elongation, polyunsaturated fatty acid | BP | 5 | 2 | 0 | 0.00544 |
| GO:0019367 | fatty acid elongation, saturated fatty acid | BP | 5 | 2 | 0 | 0.00544 |
| GO:0019368 | fatty acid elongation, unsaturated fatty acid | BP | 5 | 2 | 0 | 0.00544 |
| GO:0060729 | intestinal epithelial structure maintenance | BP | 5 | 2 | 0 | 0.00544 |
| GO:0072282 | metanephric nephron tubule morphogenesis | BP | 5 | 2 | 0 | 0.00544 |
| GO:0070487 | monocyte aggregation | BP | 5 | 2 | 0 | 0.00544 |
| GO:0001781 | neutrophil apoptotic process | BP | 5 | 2 | 0 | 0.00544 |
| GO:1901534 | positive regulation of hematopoietic progenitor cell differentiation | BP | 5 | 2 | 0 | 0.00544 |
| GO:0010572 | positive regulation of platelet activation | BP | 5 | 2 | 0 | 0.00544 |
| GO:1990405 | protein antigen binding | MF | 5 | 2 | 0 | 0.00544 |
| GO:1903972 | regulation of cellular response to macrophage colony-stimulating fa | BP | 5 | 2 | 0 | 0.00544 |
| GO:0002883 | regulation of hypersensitivity | BP | 5 | 2 | 0 | 0.00544 |
| GO:0033029 | regulation of neutrophil apoptotic process | BP | 5 | 2 | 0 | 0.00544 |
| GO:2001245 | regulation of phosphatidylcholine biosynthetic process | BP | 5 | 2 | 0 | 0.00544 |
| GO:1903969 | regulation of response to macrophage colony-stimulating factor | BP | 5 | 2 | 0 | 0.00544 |
| GO:0048254 | snoRNA localization | BP | 5 | 2 | 0 | 0.00544 |
| GO:0102756 | very-long-chain 3-ketoacyl-CoA synthase activity | MF | 5 | 2 | 0 | 0.00544 |
| GO:0042327 | positive regulation of phosphorylation | BP | 801 | 31 | 38 | 0.00558 |
| GO:0010604 | positive regulation of macromolecule metabolic process | BP | 2604 | 81 | 94 | 0.00559 |
| GO:0072171 | mesonephric tubule morphogenesis | BP | 48 | 5 | 2 | 0.00561 |
| GO:0055082 | cellular chemical homeostasis | BP | 605 | 25 | 36 | 0.00561 |
| GO:0032101 | regulation of response to external stimulus | BP | 605 | 25 | 27 | 0.00561 |
| GO:0032270 | positive regulation of cellular protein metabolic process | BP | 1247 | 44 | 50 | 0.00562 |
| GO:0046470 | phosphatidylcholine metabolic process | BP | 68 | 6 | 13 | 0.00564 |
| GO:0051050 | positive regulation of transport | BP | 769 | 30 | 38 | 0.0057 |
| GO:0030659 | cytoplasmic vesicle membrane | CC | 606 | 25 | 34 | 0.00573 |
| GO:0051928 | positive regulation of calcium ion transport | BP | 90 | 7 | 5 | 0.00576 |
| GO:2000045 | regulation of G1/S transition of mitotic cell cycle | BP | 138 | 9 | 0 | 0.00594 |
| GO:0044427 | chromosomal part | CC | 739 | 29 | 4 | 0.00603 |
| GO:0016864 | intramolecular oxidoreductase activity, transposing S-S bonds | MF | 16 | 3 | 1 | 0.00603 |

| | | | | | | |
|---|---|---|---|---|---|---|
| GO:0023023 | MHC protein complex binding | MF | 16 | 3 | 0 | 0.00603 |
| GO:0007076 | mitotic chromosome condensation | BP | 16 | 3 | 0 | 0.00603 |
| GO:0050765 | negative regulation of phagocytosis | BP | 16 | 3 | 0 | 0.00603 |
| GO:0004623 | phospholipase A2 activity | MF | 16 | 3 | 4 | 0.00603 |
| GO:0003756 | protein disulfide isomerase activity | MF | 16 | 3 | 1 | 0.00603 |
| GO:0034975 | protein folding in endoplasmic reticulum | BP | 16 | 3 | 1 | 0.00603 |
| GO:0048519 | negative regulation of biological process | BP | 4130 | 120 | 147 | 0.00604 |
| GO:1902107 | positive regulation of leukocyte differentiation | BP | 114 | 8 | 1 | 0.00604 |
| GO:0002698 | negative regulation of immune effector process | BP | 91 | 7 | 5 | 0.00612 |
| GO:0098631 | cell adhesion mediator activity | MF | 49 | 5 | 1 | 0.00612 |
| GO:0043588 | skin development | BP | 191 | 11 | 7 | 0.0063 |
| GO:0015718 | monocarboxylic acid transport | BP | 115 | 8 | 16 | 0.00636 |
| GO:0042133 | neurotransmitter metabolic process | BP | 115 | 8 | 11 | 0.00636 |
| GO:0019220 | regulation of phosphate metabolic process | BP | 1399 | 48 | 66 | 0.00648 |
| GO:0050864 | regulation of B cell activation | BP | 92 | 7 | 3 | 0.00649 |
| GO:0070542 | response to fatty acid | BP | 70 | 6 | 6 | 0.0065 |
| GO:0046394 | carboxylic acid biosynthetic process | BP | 363 | 17 | 37 | 0.00656 |
| GO:0051174 | regulation of phosphorus metabolic process | BP | 1401 | 48 | 66 | 0.00666 |
| GO:0015909 | long-chain fatty acid transport | BP | 50 | 5 | 12 | 0.00667 |
| GO:0016053 | organic acid biosynthetic process | BP | 364 | 17 | 37 | 0.00674 |
| GO:0007520 | myoblast fusion | BP | 32 | 4 | 2 | 0.00682 |
| GO:0007589 | body fluid secretion | BP | 71 | 6 | 8 | 0.00696 |
| GO:0036473 | cell death in response to oxidative stress | BP | 71 | 6 | 0 | 0.00696 |
| GO:0002824 | positive regulation of adaptive immune response based on somatic r | BP | 71 | 6 | 0 | 0.00696 |
| GO:1903426 | regulation of reactive oxygen species biosynthetic process | BP | 71 | 6 | 4 | 0.00696 |
| GO:0006810 | transport | BP | 3949 | 115 | 231 | 0.00701 |
| GO:0097367 | carbohydrate derivative binding | MF | 1767 | 58 | 106 | 0.00703 |
| GO:0031667 | response to nutrient levels | BP | 396 | 18 | 38 | 0.00706 |
| GO:0006338 | chromatin remodeling | BP | 142 | 9 | 2 | 0.00713 |
| GO:1903963 | arachidonate transport | BP | 17 | 3 | 4 | 0.00719 |
| GO:0050482 | arachidonic acid secretion | BP | 17 | 3 | 4 | 0.00719 |
| GO:0035338 | long-chain fatty-acyl-CoA biosynthetic process | BP | 17 | 3 | 1 | 0.00719 |
| GO:0045606 | positive regulation of epidermal cell differentiation | BP | 17 | 3 | 1 | 0.00719 |
| GO:0060143 | positive regulation of syncytium formation by plasma membrane fu | BP | 17 | 3 | 1 | 0.00719 |
| GO:0045806 | negative regulation of endocytosis | BP | 51 | 5 | 2 | 0.00726 |
| GO:0019933 | cAMP-mediated signaling | BP | 118 | 8 | 9 | 0.0074 |
| GO:0042770 | signal transduction in response to DNA damage | BP | 118 | 8 | 1 | 0.0074 |
| GO:0015908 | fatty acid transport | BP | 72 | 6 | 16 | 0.00744 |
| GO:0050707 | regulation of cytokine secretion | BP | 143 | 9 | 5 | 0.00745 |
| GO:0032502 | developmental process | BP | 4557 | 130 | 191 | 0.00753 |
| GO:0004869 | cysteine-type endopeptidase inhibitor activity | MF | 33 | 4 | 1 | 0.00762 |
| GO:0002244 | hematopoietic progenitor cell differentiation | BP | 144 | 9 | 4 | 0.00778 |
| GO:1903039 | positive regulation of leukocyte cell-cell adhesion | BP | 170 | 10 | 3 | 0.00781 |
| GO:0045944 | positive regulation of transcription by RNA polymerase II | BP | 924 | 34 | 34 | 0.00802 |
| GO:0000915 | actomyosin contractile ring assembly | BP | 6 | 2 | 0 | 0.00803 |
| GO:0000912 | assembly of actomyosin apparatus involved in cytokinesis | BP | 6 | 2 | 0 | 0.00803 |
| GO:0035381 | ATP-gated ion channel activity | MF | 6 | 2 | 0 | 0.00803 |
| GO:0047498 | calcium-dependent phospholipase A2 activity | MF | 6 | 2 | 2 | 0.00803 |
| GO:0031748 | D1 dopamine receptor binding | MF | 6 | 2 | 0 | 0.00803 |
| GO:0031232 | extrinsic component of external side of plasma membrane | CC | 6 | 2 | 1 | 0.00803 |
| GO:0002524 | hypersensitivity | BP | 6 | 2 | 0 | 0.00803 |
| GO:0032621 | interleukin-18 production | BP | 6 | 2 | 0 | 0.00803 |
| GO:0051454 | intracellular pH elevation | BP | 6 | 2 | 0 | 0.00803 |
| GO:0010032 | meiotic chromosome condensation | BP | 6 | 2 | 0 | 0.00803 |
| GO:0072173 | metanephric tubule morphogenesis | BP | 6 | 2 | 0 | 0.00803 |
| GO:0042824 | MHC class I peptide loading complex | CC | 6 | 2 | 0 | 0.00803 |
| GO:0051256 | mitotic spindle midzone assembly | BP | 6 | 2 | 0 | 0.00803 |
| GO:2001054 | negative regulation of mesenchymal cell apoptotic process | BP | 6 | 2 | 0 | 0.00803 |
| GO:0045852 | pH elevation | BP | 6 | 2 | 0 | 0.00803 |
| GO:0015379 | potassium:chloride symporter activity | MF | 6 | 2 | 0 | 0.00803 |
| GO:0042268 | regulation of cytolysis | BP | 6 | 2 | 0 | 0.00803 |
| GO:0046602 | regulation of mitotic centrosome separation | BP | 6 | 2 | 0 | 0.00803 |
| GO:0071229 | cellular response to acid chemical | BP | 171 | 10 | 9 | 0.00812 |
| GO:0042470 | melanosome | CC | 96 | 7 | 2 | 0.00815 |

| | | | | | | |
|---|---|---|---|---|---|---|
| GO:0048770 | pigment granule | CC | 96 | 7 | 2 | 0.00815 |
| GO:0031668 | cellular response to extracellular stimulus | BP | 226 | 12 | 19 | 0.0083 |
| GO:0043168 | anion binding | MF | 2228 | 70 | 147 | 0.00836 |
| GO:0032268 | regulation of cellular protein metabolic process | BP | 2041 | 65 | 74 | 0.00836 |
| GO:0007431 | salivary gland development | BP | 34 | 4 | 0 | 0.00847 |
| GO:0032040 | small-subunit processome | CC | 34 | 4 | 0 | 0.00847 |
| GO:0071453 | cellular response to oxygen levels | BP | 199 | 11 | 8 | 0.00847 |
| GO:0022409 | positive regulation of cell-cell adhesion | BP | 199 | 11 | 4 | 0.00847 |
| GO:0017000 | antibiotic biosynthetic process | BP | 18 | 3 | 1 | 0.00848 |
| GO:0061436 | establishment of skin barrier | BP | 18 | 3 | 0 | 0.00848 |
| GO:0060487 | lung epithelial cell differentiation | BP | 18 | 3 | 0 | 0.00848 |
| GO:0002313 | mature B cell differentiation involved in immune response | BP | 18 | 3 | 0 | 0.00848 |
| GO:0042535 | positive regulation of tumor necrosis factor biosynthetic process | BP | 18 | 3 | 0 | 0.00848 |
| GO:1901623 | regulation of lymphocyte chemotaxis | BP | 18 | 3 | 0 | 0.00848 |
| GO:0012506 | vesicle membrane | CC | 626 | 25 | 36 | 0.00851 |
| GO:0044877 | protein-containing complex binding | MF | 928 | 34 | 33 | 0.00854 |
| GO:0061135 | endopeptidase regulator activity | MF | 97 | 7 | 3 | 0.00861 |
| GO:0005788 | endoplasmic reticulum lumen | CC | 256 | 13 | 13 | 0.00878 |
| GO:0048534 | hematopoietic or lymphoid organ development | BP | 727 | 28 | 22 | 0.00882 |
| GO:1903708 | positive regulation of hemopoiesis | BP | 147 | 9 | 4 | 0.00886 |
| GO:0055080 | cation homeostasis | BP | 531 | 22 | 29 | 0.00886 |
| GO:0042116 | macrophage activation | BP | 75 | 6 | 2 | 0.00904 |
| GO:0030216 | keratinocyte differentiation | BP | 98 | 7 | 5 | 0.00909 |
| GO:0003697 | single-stranded DNA binding | MF | 98 | 7 | 1 | 0.00909 |
| GO:0071456 | cellular response to hypoxia | BP | 174 | 10 | 7 | 0.00913 |
| GO:0002548 | monocyte chemotaxis | BP | 35 | 4 | 4 | 0.00939 |
| GO:2000736 | regulation of stem cell differentiation | BP | 99 | 7 | 0 | 0.00958 |
| GO:0062012 | regulation of small molecule metabolic process | BP | 378 | 17 | 35 | 0.00963 |
| GO:1902106 | negative regulation of leukocyte differentiation | BP | 76 | 6 | 3 | 0.00963 |
| GO:0002821 | positive regulation of adaptive immune response | BP | 76 | 6 | 0 | 0.00963 |
| GO:0045786 | negative regulation of cell cycle | BP | 536 | 22 | 11 | 0.00982 |
| GO:0002520 | immune system development | BP | 767 | 29 | 22 | 0.00986 |
| GO:0071496 | cellular response to external stimulus | BP | 289 | 14 | 21 | 0.00988 |
| GO:0008574 | ATP-dependent microtubule motor activity, plus-end-directed | MF | 19 | 3 | 1 | 0.0099 |
| GO:0001530 | lipopolysaccharide binding | MF | 19 | 3 | 1 | 0.0099 |
| GO:0060479 | lung cell differentiation | BP | 19 | 3 | 0 | 0.0099 |
| GO:0002320 | lymphoid progenitor cell differentiation | BP | 19 | 3 | 1 | 0.0099 |
| GO:0033561 | regulation of water loss via skin | BP | 19 | 3 | 0 | 0.0099 |
| GO:0071674 | mononuclear cell migration | BP | 55 | 5 | 4 | 0.00995 |
| GO:0032651 | regulation of interleukin-1 beta production | BP | 55 | 5 | 1 | 0.00995 |
| GO:0009653 | anatomical structure morphogenesis | BP | 2095 | 66 | 94 | 0.00996 |
| GO:0050866 | negative regulation of cell activation | BP | 150 | 9 | 6 | 0.01004 |
| GO:0050679 | positive regulation of epithelial cell proliferation | BP | 150 | 9 | 5 | 0.01004 |
| GO:1902806 | regulation of cell cycle G1/S phase transition | BP | 150 | 9 | 1 | 0.01004 |
| GO:0001525 | angiogenesis | BP | 411 | 18 | 22 | 0.01014 |
| GO:0001932 | regulation of protein phosphorylation | BP | 1114 | 39 | 44 | 0.01023 |
| GO:0061640 | cytoskeleton-dependent cytokinesis | BP | 77 | 6 | 2 | 0.01024 |
| GO:0050702 | interleukin-1 beta secretion | BP | 36 | 4 | 1 | 0.01037 |
| GO:0042267 | natural killer cell mediated cytotoxicity | BP | 36 | 4 | 0 | 0.01037 |
| GO:1900408 | negative regulation of cellular response to oxidative stress | BP | 36 | 4 | 0 | 0.01037 |
| GO:0002762 | negative regulation of myeloid leukocyte differentiation | BP | 36 | 4 | 1 | 0.01037 |
| GO:1903202 | negative regulation of oxidative stress-induced cell death | BP | 36 | 4 | 0 | 0.01037 |
| GO:0005856 | cytoskeleton | CC | 1691 | 55 | 54 | 0.01039 |
| GO:0098771 | inorganic ion homeostasis | BP | 539 | 22 | 31 | 0.01043 |
| GO:0006869 | lipid transport | BP | 262 | 13 | 39 | 0.01053 |
| GO:0070665 | positive regulation of leukocyte proliferation | BP | 101 | 7 | 4 | 0.01064 |
| GO:1903036 | positive regulation of response to wounding | BP | 56 | 5 | 1 | 0.01072 |
| GO:0010876 | lipid localization | BP | 292 | 14 | 44 | 0.01076 |
| GO:1904018 | positive regulation of vasculature development | BP | 152 | 9 | 7 | 0.01089 |
| GO:0009888 | tissue development | BP | 1441 | 48 | 62 | 0.01103 |
| GO:0080135 | regulation of cellular response to stress | BP | 607 | 24 | 15 | 0.01104 |
| GO:0043062 | extracellular structure organization | BP | 353 | 16 | 24 | 0.01104 |
| GO:0044837 | actomyosin contractile ring organization | BP | 7 | 2 | 0 | 0.01107 |
| GO:0010997 | anaphase-promoting complex binding | MF | 7 | 2 | 0 | 0.01107 |

| | | | | | | |
|---|---|---|---|---|---|---|
| GO:0051305 | chromosome movement towards spindle pole | BP | 7 | 2 | 0 | 0.01107 |
| GO:0032490 | detection of molecule of bacterial origin | BP | 7 | 2 | 1 | 0.01107 |
| GO:0060272 | embryonic skeletal joint morphogenesis | BP | 7 | 2 | 0 | 0.01107 |
| GO:0004312 | fatty acid synthase activity | MF | 7 | 2 | 0 | 0.01107 |
| GO:0016742 | hydroxymethyl-, formyl- and related transferase activity | MF | 7 | 2 | 0 | 0.01107 |
| GO:0005828 | kinetochore microtubule | CC | 7 | 2 | 0 | 0.01107 |
| GO:0006564 | L-serine biosynthetic process | BP | 7 | 2 | 0 | 0.01107 |
| GO:0032808 | lacrimal gland development | BP | 7 | 2 | 0 | 0.01107 |
| GO:0010985 | negative regulation of lipoprotein particle clearance | BP | 7 | 2 | 1 | 0.01107 |
| GO:0071635 | negative regulation of transforming growth factor beta production | BP | 7 | 2 | 0 | 0.01107 |
| GO:0004859 | phospholipase inhibitor activity | MF | 7 | 2 | 0 | 0.01107 |
| GO:1903012 | positive regulation of bone development | BP | 7 | 2 | 0 | 0.01107 |
| GO:0045348 | positive regulation of MHC class II biosynthetic process | BP | 7 | 2 | 0 | 0.01107 |
| GO:0070761 | pre-snoRNP complex | CC | 7 | 2 | 0 | 0.01107 |
| GO:0002864 | regulation of acute inflammatory response to antigenic stimulus | BP | 7 | 2 | 0 | 0.01107 |
| GO:2001053 | regulation of mesenchymal cell apoptotic process | BP | 7 | 2 | 0 | 0.01107 |
| GO:0043232 | intracellular non-membrane-bounded organelle | CC | 3334 | 98 | 78 | 0.01112 |
| GO:1901991 | negative regulation of mitotic cell cycle phase transition | BP | 207 | 11 | 3 | 0.01119 |
| GO:0043486 | histone exchange | BP | 37 | 4 | 0 | 0.01141 |
| GO:0097150 | neuronal stem cell population maintenance | BP | 20 | 3 | 0 | 0.01144 |
| GO:0042451 | purine nucleoside biosynthetic process | BP | 20 | 3 | 0 | 0.01144 |
| GO:0046129 | purine ribonucleoside biosynthetic process | BP | 20 | 3 | 0 | 0.01144 |
| GO:0009147 | pyrimidine nucleoside triphosphate metabolic process | BP | 20 | 3 | 0 | 0.01144 |
| GO:0002082 | regulation of oxidative phosphorylation | BP | 20 | 3 | 2 | 0.01144 |
| GO:1904666 | regulation of ubiquitin protein ligase activity | BP | 20 | 3 | 0 | 0.01144 |
| GO:0033044 | regulation of chromosome organization | BP | 295 | 14 | 11 | 0.01169 |
| GO:0007586 | digestion | BP | 103 | 7 | 19 | 0.01177 |
| GO:0005488 | binding | MF | 10871 | 275 | 453 | 0.01178 |
| GO:0051173 | positive regulation of nitrogen compound metabolic process | BP | 2490 | 76 | 87 | 0.01178 |
| GO:0043228 | non-membrane-bounded organelle | CC | 3341 | 98 | 78 | 0.0118 |
| GO:0044430 | cytoskeletal part | CC | 1267 | 43 | 38 | 0.01187 |
| GO:0008144 | drug binding | MF | 1376 | 46 | 80 | 0.01205 |
| GO:0006935 | chemotaxis | BP | 482 | 20 | 30 | 0.0121 |
| GO:0045121 | membrane raft | CC | 267 | 13 | 10 | 0.01219 |
| GO:0007189 | adenylate cyclase-activating G protein-coupled receptor signaling p | BP | 80 | 6 | 4 | 0.01223 |
| GO:0009064 | glutamine family amino acid metabolic process | BP | 58 | 5 | 8 | 0.01237 |
| GO:0032410 | negative regulation of transporter activity | BP | 58 | 5 | 2 | 0.01237 |
| GO:0050871 | positive regulation of B cell activation | BP | 58 | 5 | 3 | 0.01237 |
| GO:0071840 | cellular component organization or biogenesis | BP | 5231 | 145 | 202 | 0.01241 |
| GO:0031424 | keratinization | BP | 38 | 4 | 3 | 0.01252 |
| GO:0043303 | mast cell degranulation | BP | 38 | 4 | 1 | 0.01252 |
| GO:1902883 | negative regulation of response to oxidative stress | BP | 38 | 4 | 1 | 0.01252 |
| GO:0032732 | positive regulation of interleukin-1 production | BP | 38 | 4 | 0 | 0.01252 |
| GO:0002712 | regulation of B cell mediated immunity | BP | 38 | 4 | 0 | 0.01252 |
| GO:0002889 | regulation of immunoglobulin mediated immune response | BP | 38 | 4 | 0 | 0.01252 |
| GO:0098857 | membrane microdomain | CC | 268 | 13 | 10 | 0.01254 |
| GO:0042330 | taxis | BP | 484 | 20 | 30 | 0.01261 |
| GO:1901605 | alpha-amino acid metabolic process | BP | 183 | 10 | 19 | 0.01271 |
| GO:0007219 | Notch signaling pathway | BP | 156 | 9 | 4 | 0.01276 |
| GO:0002822 | regulation of adaptive immune response based on somatic recombii | BP | 105 | 7 | 3 | 0.01299 |
| GO:0030856 | regulation of epithelial cell differentiation | BP | 105 | 7 | 6 | 0.01299 |
| GO:0031399 | regulation of protein modification process | BP | 1455 | 48 | 59 | 0.01304 |
| GO:0042325 | regulation of phosphorylation | BP | 1239 | 42 | 54 | 0.0131 |
| GO:0019835 | cytolysis | BP | 21 | 3 | 0 | 0.01311 |
| GO:0035336 | long-chain fatty-acyl-CoA metabolic process | BP | 21 | 3 | 2 | 0.01311 |
| GO:0090344 | negative regulation of cell aging | BP | 21 | 3 | 0 | 0.01311 |
| GO:2001024 | negative regulation of response to drug | BP | 21 | 3 | 0 | 0.01311 |
| GO:0050716 | positive regulation of interleukin-1 secretion | BP | 21 | 3 | 0 | 0.01311 |
| GO:0032879 | regulation of localization | BP | 2160 | 67 | 119 | 0.01314 |
| GO:0036294 | cellular response to decreased oxygen levels | BP | 184 | 10 | 8 | 0.01317 |
| GO:0031670 | cellular response to nutrient | BP | 59 | 5 | 5 | 0.01326 |
| GO:0050870 | positive regulation of T cell activation | BP | 157 | 9 | 3 | 0.01326 |
| GO:0010948 | negative regulation of cell cycle process | BP | 300 | 14 | 6 | 0.01338 |
| GO:0010942 | positive regulation of cell death | BP | 585 | 23 | 18 | 0.01351 |

| | | | | | | |
|---|---|---|---|---|---|---|
| GO:0032506 | cytokinetic process | BP | 39 | 4 | 1 | 0.01369 |
| GO:0050829 | defense response to Gram-negative bacterium | BP | 39 | 4 | 4 | 0.01369 |
| GO:0002279 | mast cell activation involved in immune response | BP | 39 | 4 | 2 | 0.01369 |
| GO:0043030 | regulation of macrophage activation | BP | 39 | 4 | 1 | 0.01369 |
| GO:0022600 | digestive system process | BP | 82 | 6 | 14 | 0.0137 |
| GO:0002573 | myeloid leukocyte differentiation | BP | 158 | 9 | 4 | 0.01378 |
| GO:0034774 | secretory granule lumen | CC | 272 | 13 | 9 | 0.01404 |
| GO:0098542 | defense response to other organism | BP | 302 | 14 | 8 | 0.0141 |
| GO:0110110 | positive regulation of animal organ morphogenesis | BP | 60 | 5 | 3 | 0.01419 |
| GO:0006801 | superoxide metabolic process | BP | 60 | 5 | 5 | 0.01419 |
| GO:0097164 | ammonium ion metabolic process | BP | 159 | 9 | 22 | 0.01431 |
| GO:0015711 | organic anion transport | BP | 364 | 16 | 44 | 0.01443 |
| GO:0050868 | negative regulation of T cell activation | BP | 83 | 6 | 4 | 0.01448 |
| GO:2000379 | positive regulation of reactive oxygen species metabolic process | BP | 83 | 6 | 3 | 0.01448 |
| GO:0097449 | astrocyte projection | CC | 8 | 2 | 0 | 0.01453 |
| GO:0071223 | cellular response to lipoteichoic acid | BP | 8 | 2 | 0 | 0.01453 |
| GO:0045009 | chitosome | CC | 8 | 2 | 0 | 0.01453 |
| GO:0033391 | chromatoid body | CC | 8 | 2 | 0 | 0.01453 |
| GO:0000796 | condensin complex | CC | 8 | 2 | 0 | 0.01453 |
| GO:0061575 | cyclin-dependent protein serine/threonine kinase activator activity | MF | 8 | 2 | 0 | 0.01453 |
| GO:0044539 | long-chain fatty acid import | BP | 8 | 2 | 1 | 0.01453 |
| GO:0033162 | melanosome membrane | CC | 8 | 2 | 0 | 0.01453 |
| GO:0097152 | mesenchymal cell apoptotic process | BP | 8 | 2 | 0 | 0.01453 |
| GO:0000022 | mitotic spindle elongation | BP | 8 | 2 | 0 | 0.01453 |
| GO:0042834 | peptidoglycan binding | MF | 8 | 2 | 0 | 0.01453 |
| GO:0090741 | pigment granule membrane | CC | 8 | 2 | 0 | 0.01453 |
| GO:0031915 | positive regulation of synaptic plasticity | BP | 8 | 2 | 0 | 0.01453 |
| GO:1905456 | regulation of lymphoid progenitor cell differentiation | BP | 8 | 2 | 1 | 0.01453 |
| GO:2001204 | regulation of osteoclast development | BP | 8 | 2 | 0 | 0.01453 |
| GO:1903909 | regulation of receptor clustering | BP | 8 | 2 | 0 | 0.01453 |
| GO:0070391 | response to lipoteichoic acid | BP | 8 | 2 | 0 | 0.01453 |
| GO:0006590 | thyroid hormone generation | BP | 8 | 2 | 1 | 0.01453 |
| GO:0006228 | UTP biosynthetic process | BP | 8 | 2 | 0 | 0.01453 |
| GO:0008150 | biological_process | BP | 12072 | 299 | 533 | 0.01456 |
| GO:0006820 | anion transport | BP | 459 | 19 | 50 | 0.01459 |
| GO:0000780 | condensed nuclear chromosome, centromeric region | CC | 22 | 3 | 0 | 0.01492 |
| GO:0008376 | acetylgalactosaminyltransferase activity | MF | 40 | 4 | 2 | 0.01493 |
| GO:0042743 | hydrogen peroxide metabolic process | BP | 40 | 4 | 1 | 0.01493 |
| GO:0002448 | mast cell mediated immunity | BP | 40 | 4 | 1 | 0.01493 |
| GO:1904063 | negative regulation of cation transmembrane transport | BP | 61 | 5 | 1 | 0.01516 |
| GO:0016853 | isomerase activity | MF | 134 | 8 | 6 | 0.01523 |
| GO:0042098 | T cell proliferation | BP | 134 | 8 | 4 | 0.01523 |
| GO:0098589 | membrane region | CC | 275 | 13 | 10 | 0.01526 |
| GO:0016798 | hydrolase activity, acting on glycosyl bonds | MF | 84 | 6 | 7 | 0.01529 |
| GO:1901988 | negative regulation of cell cycle phase transition | BP | 217 | 11 | 4 | 0.01547 |
| GO:0016887 | ATPase activity | MF | 306 | 14 | 10 | 0.01565 |
| GO:0042129 | regulation of T cell proliferation | BP | 109 | 7 | 3 | 0.0157 |
| GO:0045089 | positive regulation of innate immune response | BP | 307 | 14 | 8 | 0.01606 |
| GO:0019932 | second-messenger-mediated signaling | BP | 307 | 14 | 21 | 0.01606 |
| GO:0006695 | cholesterol biosynthetic process | BP | 62 | 5 | 10 | 0.01618 |
| GO:0046209 | nitric oxide metabolic process | BP | 62 | 5 | 4 | 0.01618 |
| GO:0032755 | positive regulation of interleukin-6 production | BP | 62 | 5 | 4 | 0.01618 |
| GO:0016528 | sarcoplasm | CC | 62 | 5 | 3 | 0.01618 |
| GO:1902653 | secondary alcohol biosynthetic process | BP | 62 | 5 | 10 | 0.01618 |
| GO:0010830 | regulation of myotube differentiation | BP | 41 | 4 | 0 | 0.01625 |
| GO:0002931 | response to ischemia | BP | 41 | 4 | 3 | 0.01625 |
| GO:0009069 | serine family amino acid metabolic process | BP | 41 | 4 | 3 | 0.01625 |
| GO:0005524 | ATP binding | MF | 1221 | 41 | 66 | 0.01639 |
| GO:1901214 | regulation of neuron death | BP | 248 | 12 | 8 | 0.0164 |
| GO:0035556 | intracellular signal transduction | BP | 2336 | 71 | 102 | 0.01657 |
| GO:0098761 | cellular response to interleukin-7 | BP | 23 | 3 | 2 | 0.01686 |
| GO:0034369 | plasma lipoprotein particle remodeling | BP | 23 | 3 | 5 | 0.01686 |
| GO:2000108 | positive regulation of leukocyte apoptotic process | BP | 23 | 3 | 0 | 0.01686 |
| GO:0034368 | protein-lipid complex remodeling | BP | 23 | 3 | 5 | 0.01686 |

| | | | | | | |
|---|---|---|---|---|---|---|
| GO:1903203 | regulation of oxidative stress-induced neuron death | BP | 23 | 3 | 0 | 0.01686 |
| GO:0042534 | regulation of tumor necrosis factor biosynthetic process | BP | 23 | 3 | 0 | 0.01686 |
| GO:0098760 | response to interleukin-7 | BP | 23 | 3 | 2 | 0.01686 |
| GO:0042533 | tumor necrosis factor biosynthetic process | BP | 23 | 3 | 0 | 0.01686 |
| GO:0071887 | leukocyte apoptotic process | BP | 86 | 6 | 3 | 0.017 |
| GO:0015914 | phospholipid transport | BP | 63 | 5 | 11 | 0.01724 |
| GO:2001057 | reactive nitrogen species metabolic process | BP | 63 | 5 | 4 | 0.01724 |
| GO:0046942 | carboxylic acid transport | BP | 250 | 12 | 30 | 0.01736 |
| GO:0015849 | organic acid transport | BP | 250 | 12 | 30 | 0.01736 |
| GO:0030866 | cortical actin cytoskeleton organization | BP | 42 | 4 | 1 | 0.01763 |
| GO:0007080 | mitotic metaphase plate congression | BP | 42 | 4 | 0 | 0.01763 |
| GO:0042130 | negative regulation of T cell proliferation | BP | 42 | 4 | 2 | 0.01763 |
| GO:0010524 | positive regulation of calcium ion transport into cytosol | BP | 42 | 4 | 2 | 0.01763 |
| GO:0060688 | regulation of morphogenesis of a branching structure | BP | 42 | 4 | 1 | 0.01763 |
| GO:0051347 | positive regulation of transferase activity | BP | 534 | 21 | 18 | 0.0178 |
| GO:0019935 | cyclic-nucleotide-mediated signaling | BP | 138 | 8 | 11 | 0.01789 |
| GO:0030968 | endoplasmic reticulum unfolded protein response | BP | 112 | 7 | 5 | 0.01798 |
| GO:0032680 | regulation of tumor necrosis factor production | BP | 112 | 7 | 2 | 0.01798 |
| GO:0021700 | developmental maturation | BP | 222 | 11 | 8 | 0.01802 |
| GO:0070371 | ERK1 and ERK2 cascade | BP | 222 | 11 | 9 | 0.01802 |
| GO:0006468 | protein phosphorylation | BP | 1557 | 50 | 64 | 0.01808 |
| GO:0055065 | metal ion homeostasis | BP | 470 | 19 | 28 | 0.01825 |
| GO:0031016 | pancreas development | BP | 64 | 5 | 3 | 0.01835 |
| GO:0060433 | bronchus development | BP | 9 | 2 | 0 | 0.01839 |
| GO:0071281 | cellular response to iron ion | BP | 9 | 2 | 2 | 0.01839 |
| GO:0006957 | complement activation, alternative pathway | BP | 9 | 2 | 0 | 0.01839 |
| GO:0098543 | detection of other organism | BP | 9 | 2 | 0 | 0.01839 |
| GO:0034663 | endoplasmic reticulum chaperone complex | CC | 9 | 2 | 0 | 0.01839 |
| GO:0031313 | extrinsic component of endosome membrane | CC | 9 | 2 | 0 | 0.01839 |
| GO:0021781 | glial cell fate commitment | BP | 9 | 2 | 0 | 0.01839 |
| GO:0006183 | GTP biosynthetic process | BP | 9 | 2 | 0 | 0.01839 |
| GO:0070486 | leukocyte aggregation | BP | 9 | 2 | 0 | 0.01839 |
| GO:0002921 | negative regulation of humoral immune response | BP | 9 | 2 | 0 | 0.01839 |
| GO:0050713 | negative regulation of interleukin-1 beta secretion | BP | 9 | 2 | 1 | 0.01839 |
| GO:0072079 | nephron tubule formation | BP | 9 | 2 | 0 | 0.01839 |
| GO:0071073 | positive regulation of phospholipid biosynthetic process | BP | 9 | 2 | 0 | 0.01839 |
| GO:0099612 | protein localization to axon | BP | 9 | 2 | 2 | 0.01839 |
| GO:1904748 | regulation of apoptotic process involved in development | BP | 9 | 2 | 1 | 0.01839 |
| GO:1902337 | regulation of apoptotic process involved in morphogenesis | BP | 9 | 2 | 1 | 0.01839 |
| GO:0051231 | spindle elongation | BP | 9 | 2 | 0 | 0.01839 |
| GO:0051255 | spindle midzone assembly | BP | 9 | 2 | 0 | 0.01839 |
| GO:0046051 | UTP metabolic process | BP | 9 | 2 | 0 | 0.01839 |
| GO:0050663 | cytokine secretion | BP | 166 | 9 | 8 | 0.01845 |
| GO:0003674 | molecular_function | MF | 12092 | 299 | 533 | 0.0185 |
| GO:0007569 | cell aging | BP | 88 | 6 | 5 | 0.01885 |
| GO:0034728 | nucleosome organization | BP | 88 | 6 | 0 | 0.01885 |
| GO:0016324 | apical plasma membrane | CC | 253 | 12 | 38 | 0.01888 |
| GO:0042056 | chemoattractant activity | MF | 24 | 3 | 1 | 0.01894 |
| GO:0002335 | mature B cell differentiation | BP | 24 | 3 | 0 | 0.01894 |
| GO:0036475 | neuron death in response to oxidative stress | BP | 24 | 3 | 0 | 0.01894 |
| GO:2000406 | positive regulation of T cell migration | BP | 24 | 3 | 0 | 0.01894 |
| GO:0034367 | protein-containing complex remodeling | BP | 24 | 3 | 5 | 0.01894 |
| GO:0042269 | regulation of natural killer cell mediated cytotoxicity | BP | 24 | 3 | 0 | 0.01894 |
| GO:0002828 | regulation of type 2 immune response | BP | 24 | 3 | 0 | 0.01894 |
| GO:0070228 | regulation of lymphocyte apoptotic process | BP | 43 | 4 | 2 | 0.01908 |
| GO:0002709 | regulation of T cell mediated immunity | BP | 43 | 4 | 2 | 0.01908 |
| GO:0007162 | negative regulation of cell adhesion | BP | 224 | 11 | 12 | 0.01912 |
| GO:0006909 | phagocytosis | BP | 224 | 11 | 10 | 0.01912 |
| GO:0043177 | organic acid binding | MF | 140 | 8 | 16 | 0.01933 |
| GO:0002690 | positive regulation of leukocyte chemotaxis | BP | 65 | 5 | 2 | 0.0195 |
| GO:0030003 | cellular cation homeostasis | BP | 475 | 19 | 25 | 0.02013 |
| GO:0008544 | epidermis development | BP | 226 | 11 | 10 | 0.02028 |
| GO:1903555 | regulation of tumor necrosis factor superfamily cytokine productio | BP | 115 | 7 | 2 | 0.02049 |
| GO:0032640 | tumor necrosis factor production | BP | 115 | 7 | 2 | 0.02049 |

| | | | | | | |
|---|---|---|---|---|---|---|
| GO:0051303 | establishment of chromosome localization | BP | 66 | 5 | 0 | 0.0207 |
| GO:0032611 | interleukin-1 beta production | BP | 66 | 5 | 1 | 0.0207 |
| GO:0051272 | positive regulation of cellular component movement | BP | 444 | 18 | 15 | 0.02072 |
| GO:0045111 | intermediate filament cytoskeleton | CC | 90 | 6 | 1 | 0.02082 |
| GO:0009913 | epidermal cell differentiation | BP | 142 | 8 | 5 | 0.02086 |
| GO:0005504 | fatty acid binding | MF | 25 | 3 | 4 | 0.02114 |
| GO:0050869 | negative regulation of B cell activation | BP | 25 | 3 | 0 | 0.02114 |
| GO:0001891 | phagocytic cup | CC | 25 | 3 | 2 | 0.02114 |
| GO:0032733 | positive regulation of interleukin-10 production | BP | 25 | 3 | 0 | 0.02114 |
| GO:0031669 | cellular response to nutrient levels | BP | 199 | 10 | 19 | 0.02158 |
| GO:0043065 | positive regulation of apoptotic process | BP | 545 | 21 | 18 | 0.02175 |
| GO:0016043 | cellular component organization | BP | 5066 | 139 | 202 | 0.02183 |
| GO:0004866 | endopeptidase inhibitor activity | MF | 91 | 6 | 3 | 0.02186 |
| GO:0045454 | cell redox homeostasis | BP | 67 | 5 | 2 | 0.02194 |
| GO:0050000 | chromosome localization | BP | 67 | 5 | 0 | 0.02194 |
| GO:0043154 | negative regulation of cysteine-type endopeptidase activity involved | BP | 67 | 5 | 4 | 0.02194 |
| GO:0016126 | sterol biosynthetic process | BP | 67 | 5 | 11 | 0.02194 |
| GO:0007010 | cytoskeleton organization | BP | 1101 | 37 | 46 | 0.02196 |
| GO:0044093 | positive regulation of molecular function | BP | 1428 | 46 | 61 | 0.02204 |
| GO:0072132 | mesenchyme morphogenesis | BP | 45 | 4 | 1 | 0.0222 |
| GO:0032655 | regulation of interleukin-12 production | BP | 45 | 4 | 0 | 0.0222 |
| GO:0016021 | integral component of membrane | CC | 3303 | 95 | 226 | 0.02239 |
| GO:0015377 | cation:chloride symporter activity | MF | 10 | 2 | 1 | 0.02263 |
| GO:0072203 | cell proliferation involved in metanephros development | BP | 10 | 2 | 0 | 0.02263 |
| GO:0055064 | chloride ion homeostasis | BP | 10 | 2 | 0 | 0.02263 |
| GO:0098581 | detection of external biotic stimulus | BP | 10 | 2 | 1 | 0.02263 |
| GO:0072498 | embryonic skeletal joint development | BP | 10 | 2 | 0 | 0.02263 |
| GO:0051938 | L-glutamate import | BP | 10 | 2 | 1 | 0.02263 |
| GO:0098712 | L-glutamate import across plasma membrane | BP | 10 | 2 | 1 | 0.02263 |
| GO:0034374 | low-density lipoprotein particle remodeling | BP | 10 | 2 | 1 | 0.02263 |
| GO:0002638 | negative regulation of immunoglobulin production | BP | 10 | 2 | 0 | 0.02263 |
| GO:0043301 | negative regulation of leukocyte degranulation | BP | 10 | 2 | 1 | 0.02263 |
| GO:0002887 | negative regulation of myeloid leukocyte mediated immunity | BP | 10 | 2 | 1 | 0.02263 |
| GO:0014029 | neural crest formation | BP | 10 | 2 | 0 | 0.02263 |
| GO:0005652 | nuclear lamina | CC | 10 | 2 | 0 | 0.02263 |
| GO:0102567 | phospholipase A2 activity (consuming 1,2-dipalmitoylphosphatidyl | MF | 10 | 2 | 2 | 0.02263 |
| GO:0102568 | phospholipase A2 activity consuming 1,2-dioleoylphosphatidyletha | MF | 10 | 2 | 2 | 0.02263 |
| GO:0045618 | positive regulation of keratinocyte differentiation | BP | 10 | 2 | 1 | 0.02263 |
| GO:0031274 | positive regulation of pseudopodium assembly | BP | 10 | 2 | 0 | 0.02263 |
| GO:0001916 | positive regulation of T cell mediated cytotoxicity | BP | 10 | 2 | 0 | 0.02263 |
| GO:1904668 | positive regulation of ubiquitin protein ligase activity | BP | 10 | 2 | 0 | 0.02263 |
| GO:0002328 | pro-B cell differentiation | BP | 10 | 2 | 1 | 0.02263 |
| GO:1901626 | regulation of postsynaptic membrane organization | BP | 10 | 2 | 0 | 0.02263 |
| GO:0031272 | regulation of pseudopodium assembly | BP | 10 | 2 | 0 | 0.02263 |
| GO:0042761 | very long-chain fatty acid biosynthetic process | BP | 10 | 2 | 0 | 0.02263 |
| GO:0033043 | regulation of organelle organization | BP | 1068 | 36 | 36 | 0.02274 |
| GO:0034976 | response to endoplasmic reticulum stress | BP | 260 | 12 | 6 | 0.0228 |
| GO:0006793 | phosphorus metabolic process | BP | 2643 | 78 | 143 | 0.02311 |
| GO:2001251 | negative regulation of chromosome organization | BP | 118 | 7 | 4 | 0.02322 |
| GO:0000794 | condensed nuclear chromosome | CC | 68 | 5 | 0 | 0.02324 |
| GO:0032760 | positive regulation of tumor necrosis factor production | BP | 68 | 5 | 2 | 0.02324 |
| GO:0043068 | positive regulation of programmed cell death | BP | 549 | 21 | 18 | 0.02334 |
| GO:0006873 | cellular ion homeostasis | BP | 483 | 19 | 27 | 0.02345 |
| GO:0090543 | Flemming body | CC | 26 | 3 | 0 | 0.02349 |
| GO:0015949 | nucleobase-containing small molecule interconversion | BP | 26 | 3 | 0 | 0.02349 |
| GO:0070207 | protein homotrimerization | BP | 26 | 3 | 0 | 0.02349 |
| GO:0002715 | regulation of natural killer cell mediated immunity | BP | 26 | 3 | 0 | 0.02349 |
| GO:0045604 | regulation of epidermal cell differentiation | BP | 46 | 4 | 2 | 0.02387 |
| GO:0002886 | regulation of myeloid leukocyte mediated immunity | BP | 46 | 4 | 2 | 0.02387 |
| GO:0042351 | 'de novo' GDP-L-fucose biosynthetic process | BP | 1 | 1 | 0 | 0.02393 |
| GO:0070650 | actin filament bundle distribution | BP | 1 | 1 | 0 | 0.02393 |
| GO:0061573 | actin filament bundle retrograde transport | BP | 1 | 1 | 0 | 0.02393 |
| GO:0004014 | adenosylmethionine decarboxylase activity | MF | 1 | 1 | 0 | 0.02393 |
| GO:0021541 | ammon gyrus development | BP | 1 | 1 | 0 | 0.02393 |

| | | | | | | |
|---|---|---|---|---|---|---|
| GO:0001788 | antibody-dependent cellular cytotoxicity | BP | 1 | 1 | 0 | 0.02393 |
| GO:0009095 | aromatic amino acid family biosynthetic process, prephenate pathw | BP | 1 | 1 | 0 | 0.02393 |
| GO:0036520 | astrocyte-dopaminergic neuron signaling | BP | 1 | 1 | 0 | 0.02393 |
| GO:0002344 | B cell affinity maturation | BP | 1 | 1 | 0 | 0.02393 |
| GO:0002450 | B cell antigen processing and presentation | BP | 1 | 1 | 0 | 0.02393 |
| GO:0032597 | B cell receptor transport into membrane raft | BP | 1 | 1 | 0 | 0.02393 |
| GO:0032595 | B cell receptor transport within lipid bilayer | BP | 1 | 1 | 0 | 0.02393 |
| GO:0000494 | box C/D snoRNA 3'-end processing | BP | 1 | 1 | 0 | 0.02393 |
| GO:0033967 | box C/D snoRNA metabolic process | BP | 1 | 1 | 0 | 0.02393 |
| GO:0034963 | box C/D snoRNA processing | BP | 1 | 1 | 0 | 0.02393 |
| GO:0062064 | box C/D snoRNP complex binding | MF | 1 | 1 | 0 | 0.02393 |
| GO:0062065 | box H/ACA snoRNP complex binding | MF | 1 | 1 | 0 | 0.02393 |
| GO:0110017 | cap-independent translational initiation of linear mRNA | BP | 1 | 1 | 0 | 0.02393 |
| GO:0070025 | carbon monoxide binding | MF | 1 | 1 | 0 | 0.02393 |
| GO:1990451 | cellular stress response to acidic pH | BP | 1 | 1 | 0 | 0.02393 |
| GO:0021685 | cerebellar granular layer structural organization | BP | 1 | 1 | 0 | 0.02393 |
| GO:0021588 | cerebellum formation | BP | 1 | 1 | 0 | 0.02393 |
| GO:0032600 | chemokine receptor transport out of membrane raft | BP | 1 | 1 | 0 | 0.02393 |
| GO:0033606 | chemokine receptor transport within lipid bilayer | BP | 1 | 1 | 0 | 0.02393 |
| GO:0031723 | CXCR4 chemokine receptor binding | MF | 1 | 1 | 0 | 0.02393 |
| GO:0097125 | cyclin B1-CDK1 complex | CC | 1 | 1 | 0 | 0.02393 |
| GO:0004122 | cystathionine beta-synthase activity | MF | 1 | 1 | 0 | 0.02393 |
| GO:0006535 | cysteine biosynthetic process from serine | BP | 1 | 1 | 0 | 0.02393 |
| GO:0004124 | cysteine synthase activity | MF | 1 | 1 | 0 | 0.02393 |
| GO:0015328 | cystine secondary active transmembrane transporter activity | MF | 1 | 1 | 0 | 0.02393 |
| GO:0015327 | cystine:glutamate antiporter activity | MF | 1 | 1 | 0 | 0.02393 |
| GO:0031431 | Dbf4-dependent protein kinase complex | CC | 1 | 1 | 0 | 0.02393 |
| GO:0006253 | dCTP catabolic process | BP | 1 | 1 | 0 | 0.02393 |
| GO:0047840 | dCTP diphosphatase activity | MF | 1 | 1 | 0 | 0.02393 |
| GO:0046065 | dCTP metabolic process | BP | 1 | 1 | 0 | 0.02393 |
| GO:0070694 | deoxyribonucleoside 5'-monophosphate N-glycosidase activity | MF | 1 | 1 | 0 | 0.02393 |
| GO:0071750 | dimeric IgA immunoglobulin complex | CC | 1 | 1 | 0 | 0.02393 |
| GO:0140206 | dipeptide import across plasma membrane | BP | 1 | 1 | 0 | 0.02393 |
| GO:0044796 | DNA polymerase processivity factor complex | CC | 1 | 1 | 0 | 0.02393 |
| GO:1990814 | DNA/DNA annealing activity | MF | 1 | 1 | 0 | 0.02393 |
| GO:1990175 | EH domain binding | MF | 1 | 1 | 0 | 0.02393 |
| GO:0008843 | endochitinase activity | MF | 1 | 1 | 0 | 0.02393 |
| GO:0061163 | endoplasmic reticulum polarization | BP | 1 | 1 | 0 | 0.02393 |
| GO:0060690 | epithelial cell differentiation involved in salivary gland developmer | BP | 1 | 1 | 0 | 0.02393 |
| GO:0060691 | epithelial cell maturation involved in salivary gland development | BP | 1 | 1 | 0 | 0.02393 |
| GO:0060517 | epithelial cell proliferation involved in prostatic bud elongation | BP | 1 | 1 | 0 | 0.02393 |
| GO:0070287 | ferritin receptor activity | MF | 1 | 1 | 0 | 0.02393 |
| GO:0021797 | forebrain anterior/posterior pattern specification | BP | 1 | 1 | 0 | 0.02393 |
| GO:0042350 | GDP-L-fucose biosynthetic process | BP | 1 | 1 | 0 | 0.02393 |
| GO:0008446 | GDP-mannose 4,6-dehydratase activity | MF | 1 | 1 | 0 | 0.02393 |
| GO:0035746 | granzyme A production | BP | 1 | 1 | 0 | 0.02393 |
| GO:0003421 | growth plate cartilage axis specification | BP | 1 | 1 | 0 | 0.02393 |
| GO:1990922 | hepatic stellate cell proliferation | BP | 1 | 1 | 0 | 0.02393 |
| GO:0005008 | hepatocyte growth factor-activated receptor activity | MF | 1 | 1 | 0 | 0.02393 |
| GO:0021576 | hindbrain formation | BP | 1 | 1 | 0 | 0.02393 |
| GO:1990259 | histone-glutamine methyltransferase activity | MF | 1 | 1 | 0 | 0.02393 |
| GO:1990258 | histone glutamine methylation | BP | 1 | 1 | 0 | 0.02393 |
| GO:2000775 | histone H3-S10 phosphorylation involved in chromosome condensa | BP | 1 | 1 | 0 | 0.02393 |
| GO:0019862 | IgA binding | MF | 1 | 1 | 0 | 0.02393 |
| GO:0071745 | IgA immunoglobulin complex | CC | 1 | 1 | 0 | 0.02393 |
| GO:0071746 | IgA immunoglobulin complex, circulating | CC | 1 | 1 | 0 | 0.02393 |
| GO:0019770 | IgG receptor activity | MF | 1 | 1 | 0 | 0.02393 |
| GO:0071753 | IgM immunoglobulin complex | CC | 1 | 1 | 0 | 0.02393 |
| GO:0071754 | IgM immunoglobulin complex, circulating | CC | 1 | 1 | 0 | 0.02393 |
| GO:0002386 | immune response in mucosal-associated lymphoid tissue | BP | 1 | 1 | 0 | 0.02393 |
| GO:0042571 | immunoglobulin complex, circulating | CC | 1 | 1 | 0 | 0.02393 |
| GO:0002415 | immunoglobulin transcytosis in epithelial cells mediated by polyme | BP | 1 | 1 | 0 | 0.02393 |
| GO:0022894 | Intermediate conductance calcium-activated potassium channel acti | MF | 1 | 1 | 0 | 0.02393 |
| GO:0005260 | intracellularly ATP-gated chloride channel activity | MF | 1 | 1 | 0 | 0.02393 |

| | | | | | | |
|---|---|---|---|---|---|---|
| GO:0099142 | intracellularly ATP-gated ion channel activity | MF | 1 | 1 | 0 | 0.02393 |
| GO:0035622 | intrahepatic bile duct development | BP | 1 | 1 | 0 | 0.02393 |
| GO:0002192 | IRES-dependent translational initiation of linear mRNA | BP | 1 | 1 | 0 | 0.02393 |
| GO:0070981 | L-asparagine biosynthetic process | BP | 1 | 1 | 0 | 0.02393 |
| GO:0070982 | L-asparagine metabolic process | BP | 1 | 1 | 0 | 0.02393 |
| GO:0099607 | lateral attachment of mitotic spindle microtubules to kinetochore | BP | 1 | 1 | 0 | 0.02393 |
| GO:0010348 | lithium:proton antiporter activity | MF | 1 | 1 | 0 | 0.02393 |
| GO:0019772 | low-affinity IgG receptor activity | MF | 1 | 1 | 0 | 0.02393 |
| GO:0003796 | lysozyme activity | MF | 1 | 1 | 0 | 0.02393 |
| GO:0070685 | macropinocytic cup | CC | 1 | 1 | 0 | 0.02393 |
| GO:0035782 | mature natural killer cell chemotaxis | BP | 1 | 1 | 0 | 0.02393 |
| GO:0010705 | meiotic DNA double-strand break processing involved in reciproca | BP | 1 | 1 | 0 | 0.02393 |
| GO:0072289 | metanephric nephron tubule formation | BP | 1 | 1 | 0 | 0.02393 |
| GO:0023030 | MHC class Ib protein binding, via antigen binding groove | MF | 1 | 1 | 0 | 0.02393 |
| GO:0023025 | MHC class Ib protein complex binding | MF | 1 | 1 | 0 | 0.02393 |
| GO:0099606 | microtubule plus-end directed mitotic chromosome migration | BP | 1 | 1 | 0 | 0.02393 |
| GO:0021732 | midbrain-hindbrain boundary maturation | BP | 1 | 1 | 0 | 0.02393 |
| GO:0022004 | midbrain-hindbrain boundary maturation during brain development | BP | 1 | 1 | 0 | 0.02393 |
| GO:0005753 | mitochondrial proton-transporting ATP synthase complex | CC | 1 | 1 | 0 | 0.02393 |
| GO:0071748 | monomeric IgA immunoglobulin complex | CC | 1 | 1 | 0 | 0.02393 |
| GO:0070856 | myosin VI light chain binding | MF | 1 | 1 | 0 | 0.02393 |
| GO:0047263 | N-acylsphingosine galactosyltransferase activity | MF | 1 | 1 | 0 | 0.02393 |
| GO:0001814 | negative regulation of antibody-dependent cellular cytotoxicity | BP | 1 | 1 | 0 | 0.02393 |
| GO:1902257 | negative regulation of apoptotic process involved in outflow tract m | BP | 1 | 1 | 0 | 0.02393 |
| GO:0045769 | negative regulation of asymmetric cell division | BP | 1 | 1 | 0 | 0.02393 |
| GO:1903126 | negative regulation of centriole-centriole cohesion | BP | 1 | 1 | 0 | 0.02393 |
| GO:0043318 | negative regulation of cytotoxic T cell degranulation | BP | 1 | 1 | 0 | 0.02393 |
| GO:0032685 | negative regulation of granulocyte macrophage colony-stimulating | BP | 1 | 1 | 0 | 0.02393 |
| GO:0010920 | negative regulation of inositol phosphate biosynthetic process | BP | 1 | 1 | 0 | 0.02393 |
| GO:0032701 | negative regulation of interleukin-18 production | BP | 1 | 1 | 0 | 0.02393 |
| GO:0032384 | negative regulation of intracellular cholesterol transport | BP | 1 | 1 | 0 | 0.02393 |
| GO:0032378 | negative regulation of intracellular lipid transport | BP | 1 | 1 | 0 | 0.02393 |
| GO:0032381 | negative regulation of intracellular sterol transport | BP | 1 | 1 | 0 | 0.02393 |
| GO:1905596 | negative regulation of low-density lipoprotein particle receptor bin | BP | 1 | 1 | 0 | 0.02393 |
| GO:1905598 | negative regulation of low-density lipoprotein receptor activity | BP | 1 | 1 | 0 | 0.02393 |
| GO:1905032 | negative regulation of membrane repolarization during cardiac mus | BP | 1 | 1 | 0 | 0.02393 |
| GO:1905025 | negative regulation of membrane repolarization during ventricular | BP | 1 | 1 | 0 | 0.02393 |
| GO:1904782 | negative regulation of NMDA glutamate receptor activity | BP | 1 | 1 | 0 | 0.02393 |
| GO:2000276 | negative regulation of oxidative phosphorylation uncoupler activity | BP | 1 | 1 | 0 | 0.02393 |
| GO:0010900 | negative regulation of phosphatidylcholine catabolic process | BP | 1 | 1 | 0 | 0.02393 |
| GO:1905601 | negative regulation of receptor-mediated endocytosis involved in ch | BP | 1 | 1 | 0 | 0.02393 |
| GO:1903403 | negative regulation of renal phosphate excretion | BP | 1 | 1 | 0 | 0.02393 |
| GO:2001229 | negative regulation of response to gamma radiation | BP | 1 | 1 | 0 | 0.02393 |
| GO:2000518 | negative regulation of T-helper 1 cell activation | BP | 1 | 1 | 0 | 0.02393 |
| GO:0032913 | negative regulation of transforming growth factor beta3 production | BP | 1 | 1 | 0 | 0.02393 |
| GO:0001811 | negative regulation of type I hypersensitivity | BP | 1 | 1 | 0 | 0.02393 |
| GO:0002893 | negative regulation of type II hypersensitivity | BP | 1 | 1 | 0 | 0.02393 |
| GO:0001797 | negative regulation of type IIa hypersensitivity | BP | 1 | 1 | 0 | 0.02393 |
| GO:0014036 | neural crest cell fate specification | BP | 1 | 1 | 0 | 0.02393 |
| GO:0106028 | neuron projection retraction | BP | 1 | 1 | 0 | 0.02393 |
| GO:0005294 | neutral L-amino acid secondary active transmembrane transporter a | MF | 1 | 1 | 0 | 0.02393 |
| GO:0050421 | nitrite reductase (NO-forming) activity | MF | 1 | 1 | 0 | 0.02393 |
| GO:0098809 | nitrite reductase activity | MF | 1 | 1 | 0 | 0.02393 |
| GO:0004648 | O-phospho-L-serine:2-oxoglutarate aminotransferase activity | MF | 1 | 1 | 0 | 0.02393 |
| GO:0021623 | oculomotor nerve formation | BP | 1 | 1 | 0 | 0.02393 |
| GO:0021622 | oculomotor nerve morphogenesis | BP | 1 | 1 | 0 | 0.02393 |
| GO:0140205 | oligopeptide import across plasma membrane | BP | 1 | 1 | 0 | 0.02393 |
| GO:0090402 | oncogene-induced cell senescence | BP | 1 | 1 | 0 | 0.02393 |
| GO:0003409 | optic cup structural organization | BP | 1 | 1 | 0 | 0.02393 |
| GO:0003404 | optic vesicle morphogenesis | BP | 1 | 1 | 0 | 0.02393 |
| GO:0016662 | oxidoreductase activity, acting on other nitrogenous compounds as | MF | 1 | 1 | 0 | 0.02393 |
| GO:0043626 | PCNA complex | CC | 1 | 1 | 0 | 0.02393 |
| GO:0071756 | pentameric IgM immunoglobulin complex | CC | 1 | 1 | 0 | 0.02393 |
| GO:0061783 | peptidoglycan muralytic activity | MF | 1 | 1 | 0 | 0.02393 |

| | | | | | | |
|---|---|---|---|---|---|---|
| GO:0018106 | peptidyl-histidine phosphorylation | BP | 1 | 1 | 0 | 0.02393 |
| GO:0002343 | peripheral B cell selection | BP | 1 | 1 | 0 | 0.02393 |
| GO:0030845 | phospholipase C-inhibiting G protein-coupled receptor signaling pa | BP | 1 | 1 | 0 | 0.02393 |
| GO:0006649 | phospholipid transfer to membrane | BP | 1 | 1 | 0 | 0.02393 |
| GO:0071749 | polymeric IgA immunoglobulin complex | CC | 1 | 1 | 0 | 0.02393 |
| GO:0001792 | polymeric immunoglobulin receptor activity | MF | 1 | 1 | 0 | 0.02393 |
| GO:2000566 | positive regulation of CD8-positive, alpha-beta T cell proliferation | BP | 1 | 1 | 0 | 0.02393 |
| GO:0045795 | positive regulation of cell volume | BP | 1 | 1 | 0 | 0.02393 |
| GO:1903435 | positive regulation of constitutive secretory pathway | BP | 1 | 1 | 0 | 0.02393 |
| GO:1902161 | positive regulation of cyclic nucleotide-gated ion channel activity | BP | 1 | 1 | 0 | 0.02393 |
| GO:0014054 | positive regulation of gamma-aminobutyric acid secretion | BP | 1 | 1 | 0 | 0.02393 |
| GO:2000513 | positive regulation of granzyme A production | BP | 1 | 1 | 0 | 0.02393 |
| GO:1902038 | positive regulation of hematopoietic stem cell differentiation | BP | 1 | 1 | 0 | 0.02393 |
| GO:1904899 | positive regulation of hepatic stellate cell proliferation | BP | 1 | 1 | 0 | 0.02393 |
| GO:2000558 | positive regulation of immunoglobulin production in mucosal tissue | BP | 1 | 1 | 0 | 0.02393 |
| GO:0010925 | positive regulation of inositol-polyphosphate 5-phosphatase activity | BP | 1 | 1 | 0 | 0.02393 |
| GO:1903489 | positive regulation of lactation | BP | 1 | 1 | 0 | 0.02393 |
| GO:0140214 | positive regulation of long-chain fatty acid import into cell | BP | 1 | 1 | 0 | 0.02393 |
| GO:0090096 | positive regulation of metanephric cap mesenchymal cell proliferati | BP | 1 | 1 | 0 | 0.02393 |
| GO:0070430 | positive regulation of nucleotide-binding oligomerization domain c | BP | 1 | 1 | 0 | 0.02393 |
| GO:1900100 | positive regulation of plasma cell differentiation | BP | 1 | 1 | 0 | 0.02393 |
| GO:1901631 | positive regulation of presynaptic membrane organization | BP | 1 | 1 | 0 | 0.02393 |
| GO:2000176 | positive regulation of pro-T cell differentiation | BP | 1 | 1 | 0 | 0.02393 |
| GO:0009674 | potassium:sodium symporter activity | MF | 1 | 1 | 0 | 0.02393 |
| GO:0004673 | protein histidine kinase activity | MF | 1 | 1 | 0 | 0.02393 |
| GO:0032599 | protein transport out of membrane raft | BP | 1 | 1 | 0 | 0.02393 |
| GO:0045259 | proton-transporting ATP synthase complex | CC | 1 | 1 | 0 | 0.02393 |
| GO:0015390 | purine-specific nucleoside:sodium symporter activity | MF | 1 | 1 | 0 | 0.02393 |
| GO:0019810 | putrescine binding | MF | 1 | 1 | 0 | 0.02393 |
| GO:0032556 | pyrimidine deoxyribonucleotide binding | MF | 1 | 1 | 0 | 0.02393 |
| GO:0001813 | regulation of antibody-dependent cellular cytotoxicity | BP | 1 | 1 | 0 | 0.02393 |
| GO:0002622 | regulation of B cell antigen processing and presentation | BP | 1 | 1 | 0 | 0.02393 |
| GO:0002721 | regulation of B cell cytokine production | BP | 1 | 1 | 0 | 0.02393 |
| GO:0060668 | regulation of branching involved in salivary gland morphogenesis b | BP | 1 | 1 | 0 | 0.02393 |
| GO:0060305 | regulation of cell diameter | BP | 1 | 1 | 0 | 0.02393 |
| GO:0060784 | regulation of cell proliferation involved in tissue homeostasis | BP | 1 | 1 | 0 | 0.02393 |
| GO:1902159 | regulation of cyclic nucleotide-gated ion channel activity | BP | 1 | 1 | 0 | 0.02393 |
| GO:1901494 | regulation of cysteine metabolic process | BP | 1 | 1 | 0 | 0.02393 |
| GO:2000431 | regulation of cytokinesis, actomyosin contractile ring assembly | BP | 1 | 1 | 0 | 0.02393 |
| GO:0043317 | regulation of cytotoxic T cell degranulation | BP | 1 | 1 | 0 | 0.02393 |
| GO:2000511 | regulation of granzyme A production | BP | 1 | 1 | 0 | 0.02393 |
| GO:1904897 | regulation of hepatic stellate cell proliferation | BP | 1 | 1 | 0 | 0.02393 |
| GO:2000557 | regulation of immunoglobulin production in mucosal tissue | BP | 1 | 1 | 0 | 0.02393 |
| GO:0010924 | regulation of inositol-polyphosphate 5-phosphatase activity | BP | 1 | 1 | 0 | 0.02393 |
| GO:0140212 | regulation of long-chain fatty acid import into cell | BP | 1 | 1 | 0 | 0.02393 |
| GO:0090095 | regulation of metanephric cap mesenchymal cell proliferation | BP | 1 | 1 | 0 | 0.02393 |
| GO:1900098 | regulation of plasma cell differentiation | BP | 1 | 1 | 0 | 0.02393 |
| GO:2000174 | regulation of pro-T cell differentiation | BP | 1 | 1 | 0 | 0.02393 |
| GO:0010849 | regulation of proton-transporting ATPase activity, rotational mecha | BP | 1 | 1 | 0 | 0.02393 |
| GO:1903402 | regulation of renal phosphate excretion | BP | 1 | 1 | 0 | 0.02393 |
| GO:2000517 | regulation of T-helper 1 cell activation | BP | 1 | 1 | 0 | 0.02393 |
| GO:0044722 | renal phosphate excretion | BP | 1 | 1 | 0 | 0.02393 |
| GO:0046680 | response to DDT | BP | 1 | 1 | 0 | 0.02393 |
| GO:0009645 | response to low light intensity stimulus | BP | 1 | 1 | 0 | 0.02393 |
| GO:0046690 | response to tellurium ion | BP | 1 | 1 | 0 | 0.02393 |
| GO:0006557 | S-adenosylmethioninamine biosynthetic process | BP | 1 | 1 | 0 | 0.02393 |
| GO:0046499 | S-adenosylmethioninamine metabolic process | BP | 1 | 1 | 0 | 0.02393 |
| GO:0106138 | Sec61 translocon complex binding | MF | 1 | 1 | 0 | 0.02393 |
| GO:0071752 | secretory dimeric IgA immunoglobulin complex | CC | 1 | 1 | 0 | 0.02393 |
| GO:0071751 | secretory IgA immunoglobulin complex | CC | 1 | 1 | 0 | 0.02393 |
| GO:0030519 | snoRNP binding | MF | 1 | 1 | 0 | 0.02393 |
| GO:0008511 | sodium:potassium:chloride symporter activity | MF | 1 | 1 | 0 | 0.02393 |
| GO:0072086 | specification of loop of Henle identity | BP | 1 | 1 | 0 | 0.02393 |
| GO:0004506 | squalene monooxygenase activity | MF | 1 | 1 | 0 | 0.02393 |

| | | | | | | |
|---|---|---|---|---|---|---|
| GO:0030527 | structural constituent of chromatin | MF | 1 | 1 | 0 | 0.02393 |
| GO:0120146 | sulfatide binding | MF | 1 | 1 | 0 | 0.02393 |
| GO:1901680 | sulfur-containing amino acid secondary active transmembrane trans | MF | 1 | 1 | 0 | 0.02393 |
| GO:0070634 | transepithelial ammonium transport | BP | 1 | 1 | 0 | 0.02393 |
| GO:0004998 | transferrin receptor activity | MF | 1 | 1 | 0 | 0.02393 |
| GO:0042497 | triacyl lipopeptide binding | MF | 1 | 1 | 0 | 0.02393 |
| GO:0052666 | tRNA (cytosine-2'-O-)-methyltransferase activity | MF | 1 | 1 | 0 | 0.02393 |
| GO:0009020 | tRNA (guanosine-2'-O-)-methyltransferase activity | MF | 1 | 1 | 0 | 0.02393 |
| GO:0002128 | tRNA nucleoside ribose methylation | BP | 1 | 1 | 0 | 0.02393 |
| GO:0006571 | tyrosine biosynthetic process | BP | 1 | 1 | 0 | 0.02393 |
| GO:0035375 | zymogen binding | MF | 1 | 1 | 0 | 0.02393 |
| GO:0048754 | branching morphogenesis of an epithelial tube | BP | 119 | 7 | 6 | 0.02419 |
| GO:0008201 | heparin binding | MF | 119 | 7 | 10 | 0.02419 |
| GO:0034399 | nuclear periphery | CC | 119 | 7 | 1 | 0.02419 |
| GO:0002819 | regulation of adaptive immune response | BP | 119 | 7 | 3 | 0.02419 |
| GO:0071706 | tumor necrosis factor superfamily cytokine production | BP | 119 | 7 | 2 | 0.02419 |
| GO:0061138 | morphogenesis of a branching epithelium | BP | 147 | 8 | 7 | 0.02505 |
| GO:0010951 | negative regulation of endopeptidase activity | BP | 147 | 8 | 6 | 0.02505 |
| GO:0006898 | receptor-mediated endocytosis | BP | 204 | 10 | 18 | 0.02509 |
| GO:1903038 | negative regulation of leukocyte cell-cell adhesion | BP | 94 | 6 | 5 | 0.02518 |
| GO:0006282 | regulation of DNA repair | BP | 94 | 6 | 2 | 0.02518 |
| GO:2000241 | regulation of reproductive process | BP | 94 | 6 | 3 | 0.02518 |
| GO:0005815 | microtubule organizing center | CC | 621 | 23 | 11 | 0.02525 |
| GO:0044425 | membrane part | CC | 4478 | 124 | 278 | 0.02529 |
| GO:1902476 | chloride transmembrane transport | BP | 47 | 4 | 3 | 0.02561 |
| GO:0030865 | cortical cytoskeleton organization | BP | 47 | 4 | 1 | 0.02561 |
| GO:0032615 | interleukin-12 production | BP | 47 | 4 | 0 | 0.02561 |
| GO:1990777 | lipoprotein particle | CC | 27 | 3 | 5 | 0.02597 |
| GO:0045920 | negative regulation of exocytosis | BP | 27 | 3 | 1 | 0.02597 |
| GO:0002701 | negative regulation of production of molecular mediator of immune | BP | 27 | 3 | 3 | 0.02597 |
| GO:0034358 | plasma lipoprotein particle | CC | 27 | 3 | 5 | 0.02597 |
| GO:0002714 | positive regulation of B cell mediated immunity | BP | 27 | 3 | 0 | 0.02597 |
| GO:0002891 | positive regulation of immunoglobulin mediated immune response | BP | 27 | 3 | 0 | 0.02597 |
| GO:2000403 | positive regulation of lymphocyte migration | BP | 27 | 3 | 0 | 0.02597 |
| GO:0033280 | response to vitamin D | BP | 27 | 3 | 1 | 0.02597 |
| GO:0042092 | type 2 immune response | BP | 27 | 3 | 0 | 0.02597 |
| GO:0097306 | cellular response to alcohol | BP | 70 | 5 | 2 | 0.02597 |
| GO:1903557 | positive regulation of tumor necrosis factor superfamily cytokine pı | BP | 70 | 5 | 2 | 0.02597 |
| GO:0030684 | preribosome | CC | 70 | 5 | 0 | 0.02597 |
| GO:0032649 | regulation of interferon-gamma production | BP | 70 | 5 | 1 | 0.02597 |
| GO:0033273 | response to vitamin | BP | 70 | 5 | 3 | 0.02597 |
| GO:0009416 | response to light stimulus | BP | 235 | 11 | 13 | 0.02611 |
| GO:0000226 | microtubule cytoskeleton organization | BP | 456 | 18 | 15 | 0.02619 |
| GO:0008203 | cholesterol metabolic process | BP | 121 | 7 | 19 | 0.0262 |
| GO:0044774 | mitotic DNA integrity checkpoint | BP | 95 | 6 | 2 | 0.02636 |
| GO:0030414 | peptidase inhibitor activity | MF | 95 | 6 | 3 | 0.02636 |
| GO:0034446 | substrate adhesion-dependent cell spreading | BP | 95 | 6 | 5 | 0.02636 |
| GO:0008017 | microtubule binding | MF | 206 | 10 | 8 | 0.02661 |
| GO:2001020 | regulation of response to DNA damage stimulus | BP | 177 | 9 | 2 | 0.02664 |
| GO:0048471 | perinuclear region of cytoplasm | CC | 591 | 22 | 31 | 0.02692 |
| GO:1901564 | organonitrogen compound metabolic process | BP | 5303 | 144 | 243 | 0.02703 |
| GO:0015172 | acidic amino acid transmembrane transporter activity | MF | 11 | 2 | 1 | 0.02722 |
| GO:0015296 | anion:cation symporter activity | MF | 11 | 2 | 1 | 0.02722 |
| GO:0051315 | attachment of mitotic spindle microtubules to kinetochore | BP | 11 | 2 | 0 | 0.02722 |
| GO:0036006 | cellular response to macrophage colony-stimulating factor stimulus | BP | 11 | 2 | 0 | 0.02722 |
| GO:0001886 | endothelial cell morphogenesis | BP | 11 | 2 | 2 | 0.02722 |
| GO:0030497 | fatty acid elongation | BP | 11 | 2 | 1 | 0.02722 |
| GO:0019215 | intermediate filament binding | MF | 11 | 2 | 1 | 0.02722 |
| GO:0045095 | keratin filament | CC | 11 | 2 | 0 | 0.02722 |
| GO:0005313 | L-glutamate transmembrane transporter activity | MF | 11 | 2 | 1 | 0.02722 |
| GO:0055102 | lipase inhibitor activity | MF | 11 | 2 | 2 | 0.02722 |
| GO:0007100 | mitotic centrosome separation | BP | 11 | 2 | 0 | 0.02722 |
| GO:1990023 | mitotic spindle midzone | CC | 11 | 2 | 0 | 0.02722 |
| GO:0032375 | negative regulation of cholesterol transport | BP | 11 | 2 | 3 | 0.02722 |

| | | | | | | |
|---|---|---|---|---|---|---|
| GO:0050711 | negative regulation of interleukin-1 secretion | BP | 11 | 2 | 1 | 0.02722 |
| GO:2000650 | negative regulation of sodium ion transmembrane transporter activi | BP | 11 | 2 | 1 | 0.02722 |
| GO:0032372 | negative regulation of sterol transport | BP | 11 | 2 | 3 | 0.02722 |
| GO:0002710 | negative regulation of T cell mediated immunity | BP | 11 | 2 | 1 | 0.02722 |
| GO:0001846 | opsonin binding | MF | 11 | 2 | 0 | 0.02722 |
| GO:0036148 | phosphatidylglycerol acyl-chain remodeling | BP | 11 | 2 | 2 | 0.02722 |
| GO:0045579 | positive regulation of B cell differentiation | BP | 11 | 2 | 0 | 0.02722 |
| GO:0010820 | positive regulation of T cell chemotaxis | BP | 11 | 2 | 0 | 0.02722 |
| GO:0097104 | postsynaptic membrane assembly | BP | 11 | 2 | 0 | 0.02722 |
| GO:0071435 | potassium ion export | BP | 11 | 2 | 0 | 0.02722 |
| GO:0097623 | potassium ion export across plasma membrane | BP | 11 | 2 | 0 | 0.02722 |
| GO:0031269 | pseudopodium assembly | BP | 11 | 2 | 0 | 0.02722 |
| GO:1901722 | regulation of cell proliferation involved in kidney development | BP | 11 | 2 | 1 | 0.02722 |
| GO:0002730 | regulation of dendritic cell cytokine production | BP | 11 | 2 | 1 | 0.02722 |
| GO:0033623 | regulation of integrin activation | BP | 11 | 2 | 0 | 0.02722 |
| GO:0036005 | response to macrophage colony-stimulating factor | BP | 11 | 2 | 0 | 0.02722 |
| GO:0017154 | semaphorin receptor activity | MF | 11 | 2 | 1 | 0.02722 |
| GO:0072520 | seminiferous tubule development | BP | 11 | 2 | 0 | 0.02722 |
| GO:0009071 | serine family amino acid catabolic process | BP | 11 | 2 | 2 | 0.02722 |
| GO:0001094 | TFIID-class transcription factor complex binding | MF | 11 | 2 | 0 | 0.02722 |
| GO:0042403 | thyroid hormone metabolic process | BP | 11 | 2 | 2 | 0.02722 |
| GO:0034766 | negative regulation of ion transmembrane transport | BP | 71 | 5 | 2 | 0.02741 |
| GO:0044070 | regulation of anion transport | BP | 71 | 5 | 8 | 0.02741 |
| GO:0045576 | mast cell activation | BP | 48 | 4 | 2 | 0.02743 |
| GO:0050891 | multicellular organismal water homeostasis | BP | 48 | 4 | 0 | 0.02743 |
| GO:0030101 | natural killer cell activation | BP | 48 | 4 | 1 | 0.02743 |
| GO:0045428 | regulation of nitric oxide biosynthetic process | BP | 48 | 4 | 3 | 0.02743 |
| GO:0072678 | T cell migration | BP | 48 | 4 | 0 | 0.02743 |
| GO:0030330 | DNA damage response, signal transduction by p53 class mediator | BP | 96 | 6 | 1 | 0.02757 |
| GO:1901343 | negative regulation of vasculature development | BP | 96 | 6 | 6 | 0.02757 |
| GO:0050671 | positive regulation of lymphocyte proliferation | BP | 96 | 6 | 4 | 0.02757 |
| GO:0031224 | intrinsic component of membrane | CC | 3410 | 97 | 236 | 0.02762 |
| GO:0008361 | regulation of cell size | BP | 150 | 8 | 8 | 0.02783 |
| GO:0032559 | adenyl ribonucleotide binding | MF | 1267 | 41 | 71 | 0.02817 |
| GO:2000027 | regulation of animal organ morphogenesis | BP | 208 | 10 | 4 | 0.02819 |
| GO:0042582 | azurophil granule | CC | 123 | 7 | 3 | 0.02832 |
| GO:0005766 | primary lysosome | CC | 123 | 7 | 3 | 0.02832 |
| GO:0014074 | response to purine-containing compound | BP | 123 | 7 | 11 | 0.02832 |
| GO:0071353 | cellular response to interleukin-4 | BP | 28 | 3 | 1 | 0.02858 |
| GO:0044364 | disruption of cells of other organism | BP | 28 | 3 | 2 | 0.02858 |
| GO:0046949 | fatty-acyl-CoA biosynthetic process | BP | 28 | 3 | 1 | 0.02858 |
| GO:0031640 | killing of cells of other organism | BP | 28 | 3 | 2 | 0.02858 |
| GO:0110111 | negative regulation of animal organ morphogenesis | BP | 28 | 3 | 1 | 0.02858 |
| GO:0045684 | positive regulation of epidermis development | BP | 28 | 3 | 1 | 0.02858 |
| GO:0002764 | immune response-regulating signaling pathway | BP | 461 | 18 | 12 | 0.02876 |
| GO:0006811 | ion transport | BP | 1196 | 39 | 98 | 0.0288 |
| GO:0032946 | positive regulation of mononuclear cell proliferation | BP | 97 | 6 | 4 | 0.02882 |
| GO:0038024 | cargo receptor activity | MF | 72 | 5 | 6 | 0.0289 |
| GO:0032481 | positive regulation of type I interferon production | BP | 72 | 5 | 1 | 0.0289 |
| GO:0045171 | intercellular bridge | CC | 49 | 4 | 0 | 0.02932 |
| GO:0031663 | lipopolysaccharide-mediated signaling pathway | BP | 49 | 4 | 2 | 0.02932 |
| GO:0050918 | positive chemotaxis | BP | 49 | 4 | 1 | 0.02932 |
| GO:0043525 | positive regulation of neuron apoptotic process | BP | 49 | 4 | 1 | 0.02932 |
| GO:1900024 | regulation of substrate adhesion-dependent cell spreading | BP | 49 | 4 | 2 | 0.02932 |
| GO:0060284 | regulation of cell development | BP | 734 | 26 | 30 | 0.02937 |
| GO:0010243 | response to organonitrogen compound | BP | 804 | 28 | 36 | 0.02943 |
| GO:0071346 | cellular response to interferon-gamma | BP | 124 | 7 | 2 | 0.02943 |
| GO:0005575 | cellular_component | CC | 12500 | 306 | 557 | 0.02955 |
| GO:0046982 | protein heterodimerization activity | MF | 333 | 14 | 12 | 0.0297 |
| GO:0052547 | regulation of peptidase activity | BP | 302 | 13 | 13 | 0.03006 |
| GO:0048144 | fibroblast proliferation | BP | 73 | 5 | 0 | 0.03044 |
| GO:2000117 | negative regulation of cysteine-type endopeptidase activity | BP | 73 | 5 | 4 | 0.03044 |
| GO:0002702 | positive regulation of production of molecular mediator of immune | BP | 73 | 5 | 1 | 0.03044 |
| GO:0048145 | regulation of fibroblast proliferation | BP | 73 | 5 | 0 | 0.03044 |

| | | | | | | |
|---|---|---|---|---|---|---|
| GO:0001678 | cellular glucose homeostasis | BP | 125 | 7 | 9 | 0.03056 |
| GO:0034620 | cellular response to unfolded protein | BP | 125 | 7 | 5 | 0.03056 |
| GO:1902652 | secondary alcohol metabolic process | BP | 125 | 7 | 19 | 0.03056 |
| GO:0010466 | negative regulation of peptidase activity | BP | 153 | 8 | 6 | 0.03081 |
| GO:2000147 | positive regulation of cell motility | BP | 432 | 17 | 14 | 0.03096 |
| GO:0045860 | positive regulation of protein kinase activity | BP | 432 | 17 | 16 | 0.03096 |
| GO:0016266 | O-glycan processing | BP | 50 | 4 | 2 | 0.03129 |
| GO:0010332 | response to gamma radiation | BP | 50 | 4 | 0 | 0.03129 |
| GO:0002639 | positive regulation of immunoglobulin production | BP | 29 | 3 | 0 | 0.03133 |
| GO:0032735 | positive regulation of interleukin-12 production | BP | 29 | 3 | 0 | 0.03133 |
| GO:0002711 | positive regulation of T cell mediated immunity | BP | 29 | 3 | 0 | 0.03133 |
| GO:0032994 | protein-lipid complex | CC | 29 | 3 | 5 | 0.03133 |
| GO:2000826 | regulation of heart morphogenesis | BP | 29 | 3 | 1 | 0.03133 |
| GO:1901031 | regulation of response to reactive oxygen species | BP | 29 | 3 | 0 | 0.03133 |
| GO:0051961 | negative regulation of nervous system development | BP | 242 | 11 | 9 | 0.03142 |
| GO:0030183 | B cell differentiation | BP | 99 | 6 | 0 | 0.03143 |
| GO:0030554 | adenyl nucleotide binding | MF | 1278 | 41 | 71 | 0.03179 |
| GO:0034644 | cellular response to UV | BP | 74 | 5 | 2 | 0.03203 |
| GO:0002438 | acute inflammatory response to antigenic stimulus | BP | 12 | 2 | 0 | 0.03216 |
| GO:0036109 | alpha-linolenic acid metabolic process | BP | 12 | 2 | 2 | 0.03216 |
| GO:0051299 | centrosome separation | BP | 12 | 2 | 0 | 0.03216 |
| GO:0006241 | CTP biosynthetic process | BP | 12 | 2 | 0 | 0.03216 |
| GO:0046036 | CTP metabolic process | BP | 12 | 2 | 0 | 0.03216 |
| GO:0002371 | dendritic cell cytokine production | BP | 12 | 2 | 1 | 0.03216 |
| GO:0006268 | DNA unwinding involved in DNA replication | BP | 12 | 2 | 0 | 0.03216 |
| GO:0021871 | forebrain regionalization | BP | 12 | 2 | 0 | 0.03216 |
| GO:1901070 | guanosine-containing compound biosynthetic process | BP | 12 | 2 | 0 | 0.03216 |
| GO:0043651 | linoleic acid metabolic process | BP | 12 | 2 | 1 | 0.03216 |
| GO:2001039 | negative regulation of cellular response to drug | BP | 12 | 2 | 0 | 0.03216 |
| GO:0010561 | negative regulation of glycoprotein biosynthetic process | BP | 12 | 2 | 0 | 0.03216 |
| GO:1903206 | negative regulation of hydrogen peroxide-induced cell death | BP | 12 | 2 | 0 | 0.03216 |
| GO:0043031 | negative regulation of macrophage activation | BP | 12 | 2 | 0 | 0.03216 |
| GO:0010832 | negative regulation of myotube differentiation | BP | 12 | 2 | 0 | 0.03216 |
| GO:1901032 | negative regulation of response to reactive oxygen species | BP | 12 | 2 | 0 | 0.03216 |
| GO:1902306 | negative regulation of sodium ion transmembrane transport | BP | 12 | 2 | 1 | 0.03216 |
| GO:1900025 | negative regulation of substrate adhesion-dependent cell spreading | BP | 12 | 2 | 0 | 0.03216 |
| GO:0001780 | neutrophil homeostasis | BP | 12 | 2 | 0 | 0.03216 |
| GO:2000193 | positive regulation of fatty acid transport | BP | 12 | 2 | 1 | 0.03216 |
| GO:0031268 | pseudopodium organization | BP | 12 | 2 | 0 | 0.03216 |
| GO:0032354 | response to follicle-stimulating hormone | BP | 12 | 2 | 0 | 0.03216 |
| GO:0070493 | thrombin-activated receptor signaling pathway | BP | 12 | 2 | 1 | 0.03216 |
| GO:1901654 | response to ketone | BP | 155 | 8 | 5 | 0.03292 |
| GO:0006986 | response to unfolded protein | BP | 155 | 8 | 5 | 0.03292 |
| GO:0071824 | protein-DNA complex subunit organization | BP | 184 | 9 | 1 | 0.03302 |
| GO:0051049 | regulation of transport | BP | 1430 | 45 | 83 | 0.03328 |
| GO:0032413 | negative regulation of ion transmembrane transporter activity | BP | 51 | 4 | 1 | 0.03334 |
| GO:0072676 | lymphocyte migration | BP | 75 | 5 | 1 | 0.03367 |
| GO:0035579 | specific granule membrane | CC | 75 | 5 | 4 | 0.03367 |
| GO:0002286 | T cell activation involved in immune response | BP | 75 | 5 | 2 | 0.03367 |
| GO:0001763 | morphogenesis of a branching structure | BP | 156 | 8 | 8 | 0.03401 |
| GO:0010212 | response to ionizing radiation | BP | 128 | 7 | 2 | 0.03414 |
| GO:0032508 | DNA duplex unwinding | BP | 101 | 6 | 1 | 0.0342 |
| GO:0009925 | basal plasma membrane | CC | 30 | 3 | 2 | 0.03421 |
| GO:0060428 | lung epithelium development | BP | 30 | 3 | 0 | 0.03421 |
| GO:0005720 | nuclear heterochromatin | CC | 30 | 3 | 0 | 0.03421 |
| GO:0032728 | positive regulation of interferon-beta production | BP | 30 | 3 | 1 | 0.03421 |
| GO:0090184 | positive regulation of kidney development | BP | 30 | 3 | 1 | 0.03421 |
| GO:0070670 | response to interleukin-4 | BP | 30 | 3 | 1 | 0.03421 |
| GO:0032941 | secretion by tissue | BP | 30 | 3 | 1 | 0.03421 |
| GO:0044454 | nuclear chromosome part | CC | 438 | 17 | 2 | 0.03462 |
| GO:1901617 | organic hydroxy compound biosynthetic process | BP | 216 | 10 | 19 | 0.03519 |
| GO:0045177 | apical part of cell | CC | 309 | 13 | 42 | 0.0352 |
| GO:0036094 | small molecule binding | MF | 2001 | 60 | 125 | 0.03523 |
| GO:0007275 | multicellular organism development | BP | 3883 | 108 | 160 | 0.03523 |

| | | | | | | |
|---|---|---|---|---|---|---|
| GO:0010035 | response to inorganic substance | BP | 439 | 17 | 20 | 0.03526 |
| GO:0045621 | positive regulation of lymphocyte differentiation | BP | 76 | 5 | 1 | 0.03537 |
| GO:1901532 | regulation of hematopoietic progenitor cell differentiation | BP | 76 | 5 | 1 | 0.03537 |
| GO:0050764 | regulation of phagocytosis | BP | 76 | 5 | 3 | 0.03537 |
| GO:0051302 | regulation of cell division | BP | 129 | 7 | 6 | 0.03539 |
| GO:0050853 | B cell receptor signaling pathway | BP | 52 | 4 | 0 | 0.03546 |
| GO:0015030 | Cajal body | CC | 52 | 4 | 0 | 0.03546 |
| GO:0006879 | cellular iron ion homeostasis | BP | 52 | 4 | 4 | 0.03546 |
| GO:0010812 | negative regulation of cell-substrate adhesion | BP | 52 | 4 | 1 | 0.03546 |
| GO:0050710 | negative regulation of cytokine secretion | BP | 52 | 4 | 2 | 0.03546 |
| GO:0045739 | positive regulation of DNA repair | BP | 52 | 4 | 2 | 0.03546 |
| GO:0030104 | water homeostasis | BP | 52 | 4 | 2 | 0.03546 |
| GO:0009891 | positive regulation of biosynthetic process | BP | 1586 | 49 | 56 | 0.0356 |
| GO:0070997 | neuron death | BP | 278 | 12 | 9 | 0.03562 |
| GO:0009615 | response to virus | BP | 247 | 11 | 5 | 0.03566 |
| GO:0006796 | phosphate-containing compound metabolic process | BP | 2620 | 76 | 142 | 0.0358 |
| GO:0050921 | positive regulation of chemotaxis | BP | 103 | 6 | 3 | 0.03711 |
| GO:0051281 | positive regulation of release of sequestered calcium ion into cytos( | BP | 31 | 3 | 2 | 0.03723 |
| GO:1901976 | regulation of cell cycle checkpoint | BP | 31 | 3 | 1 | 0.03723 |
| GO:0042455 | ribonucleoside biosynthetic process | BP | 31 | 3 | 0 | 0.03723 |
| GO:0001505 | regulation of neurotransmitter levels | BP | 280 | 12 | 20 | 0.03731 |
| GO:0008301 | DNA binding, bending | MF | 13 | 2 | 0 | 0.03741 |
| GO:0050780 | dopamine receptor binding | MF | 13 | 2 | 0 | 0.03741 |
| GO:0002070 | epithelial cell maturation | BP | 13 | 2 | 0 | 0.03741 |
| GO:0072075 | metanephric mesenchyme development | BP | 13 | 2 | 0 | 0.03741 |
| GO:0002385 | mucosal immune response | BP | 13 | 2 | 2 | 0.03741 |
| GO:0060457 | negative regulation of digestive system process | BP | 13 | 2 | 2 | 0.03741 |
| GO:1903019 | negative regulation of glycoprotein metabolic process | BP | 13 | 2 | 0 | 0.03741 |
| GO:0060192 | negative regulation of lipase activity | BP | 13 | 2 | 2 | 0.03741 |
| GO:0016502 | nucleotide receptor activity | MF | 13 | 2 | 0 | 0.03741 |
| GO:1900103 | positive regulation of endoplasmic reticulum unfolded protein respo | BP | 13 | 2 | 1 | 0.03741 |
| GO:0034116 | positive regulation of heterotypic cell-cell adhesion | BP | 13 | 2 | 0 | 0.03741 |
| GO:0140131 | positive regulation of lymphocyte chemotaxis | BP | 13 | 2 | 0 | 0.03741 |
| GO:0090026 | positive regulation of monocyte chemotaxis | BP | 13 | 2 | 1 | 0.03741 |
| GO:1901741 | positive regulation of myoblast fusion | BP | 13 | 2 | 0 | 0.03741 |
| GO:0001614 | purinergic nucleotide receptor activity | MF | 13 | 2 | 0 | 0.03741 |
| GO:0009209 | pyrimidine ribonucleoside triphosphate biosynthetic process | BP | 13 | 2 | 0 | 0.03741 |
| GO:0004857 | enzyme inhibitor activity | MF | 249 | 11 | 8 | 0.03747 |
| GO:0051310 | metaphase plate congression | BP | 53 | 4 | 0 | 0.03766 |
| GO:0048856 | anatomical structure development | BP | 4256 | 117 | 177 | 0.03785 |
| GO:0052548 | regulation of endopeptidase activity | BP | 281 | 12 | 13 | 0.03817 |
| GO:1901698 | response to nitrogen compound | BP | 859 | 29 | 39 | 0.03818 |
| GO:0098772 | molecular function regulator | MF | 1259 | 40 | 51 | 0.03846 |
| GO:0098739 | import across plasma membrane | BP | 78 | 5 | 9 | 0.03891 |
| GO:2001022 | positive regulation of response to DNA damage stimulus | BP | 78 | 5 | 2 | 0.03891 |
| GO:0010646 | regulation of cell communication | BP | 2749 | 79 | 119 | 0.03917 |
| GO:0016125 | sterol metabolic process | BP | 132 | 7 | 21 | 0.03932 |
| GO:1902400 | intracellular signal transduction involved in G1 DNA damage checl | BP | 54 | 4 | 0 | 0.03994 |
| GO:0033260 | nuclear DNA replication | BP | 54 | 4 | 0 | 0.03994 |
| GO:1904427 | positive regulation of calcium ion transmembrane transport | BP | 54 | 4 | 3 | 0.03994 |
| GO:0050766 | positive regulation of phagocytosis | BP | 54 | 4 | 2 | 0.03994 |
| GO:0016529 | sarcoplasmic reticulum | CC | 54 | 4 | 3 | 0.03994 |
| GO:0072431 | signal transduction involved in mitotic G1 DNA damage checkpoin | BP | 54 | 4 | 0 | 0.03994 |
| GO:0051208 | sequestering of calcium ion | BP | 105 | 6 | 5 | 0.04018 |
| GO:0034762 | regulation of transmembrane transport | BP | 413 | 16 | 22 | 0.04019 |
| GO:0003899 | DNA-directed 5'-3' RNA polymerase activity | MF | 32 | 3 | 0 | 0.04037 |
| GO:0002437 | inflammatory response to antigenic stimulus | BP | 32 | 3 | 0 | 0.04037 |
| GO:2000516 | positive regulation of CD4-positive, alpha-beta T cell activation | BP | 32 | 3 | 1 | 0.04037 |
| GO:0010922 | positive regulation of phosphatase activity | BP | 32 | 3 | 2 | 0.04037 |
| GO:0051233 | spindle midzone | CC | 32 | 3 | 0 | 0.04037 |
| GO:0014070 | response to organic cyclic compound | BP | 722 | 25 | 43 | 0.04062 |
| GO:0034765 | regulation of ion transmembrane transport | BP | 348 | 14 | 19 | 0.04069 |
| GO:0000910 | cytokinesis | BP | 133 | 7 | 6 | 0.0407 |
| GO:0046916 | cellular transition metal ion homeostasis | BP | 79 | 5 | 6 | 0.04076 |

| | | | | | | |
|---|---|---|---|---|---|---|
| GO:0032609 | interferon-gamma production | BP | 79 | 5 | 1 | 0.04076 |
| GO:0032414 | positive regulation of ion transmembrane transporter activity | BP | 79 | 5 | 4 | 0.04076 |
| GO:0006890 | retrograde vesicle-mediated transport, Golgi to ER | BP | 79 | 5 | 2 | 0.04076 |
| GO:0051051 | negative regulation of transport | BP | 381 | 15 | 18 | 0.04095 |
| GO:0055086 | nucleobase-containing small molecule metabolic process | BP | 618 | 22 | 45 | 0.04105 |
| GO:0070555 | response to interleukin-1 | BP | 162 | 8 | 7 | 0.04106 |
| GO:0051054 | positive regulation of DNA metabolic process | BP | 192 | 9 | 2 | 0.04153 |
| GO:0071356 | cellular response to tumor necrosis factor | BP | 223 | 10 | 5 | 0.04224 |
| GO:1903670 | regulation of sprouting angiogenesis | BP | 55 | 4 | 6 | 0.04229 |
| GO:0072413 | signal transduction involved in mitotic cell cycle checkpoint | BP | 55 | 4 | 0 | 0.04229 |
| GO:1902402 | signal transduction involved in mitotic DNA damage checkpoint | BP | 55 | 4 | 0 | 0.04229 |
| GO:1902403 | signal transduction involved in mitotic DNA integrity checkpoint | BP | 55 | 4 | 0 | 0.04229 |
| GO:0046890 | regulation of lipid biosynthetic process | BP | 163 | 8 | 18 | 0.04233 |
| GO:0012505 | endomembrane system | CC | 3471 | 97 | 166 | 0.04238 |
| GO:0004197 | cysteine-type endopeptidase activity | MF | 80 | 5 | 3 | 0.04266 |
| GO:0032368 | regulation of lipid transport | BP | 80 | 5 | 9 | 0.04266 |
| GO:0007017 | microtubule-based process | BP | 621 | 22 | 21 | 0.04291 |
| GO:0010369 | chromocenter | CC | 14 | 2 | 0 | 0.04297 |
| GO:0009595 | detection of biotic stimulus | BP | 14 | 2 | 1 | 0.04297 |
| GO:0042921 | glucocorticoid receptor signaling pathway | BP | 14 | 2 | 2 | 0.04297 |
| GO:0070365 | hepatocyte differentiation | BP | 14 | 2 | 1 | 0.04297 |
| GO:0048012 | hepatocyte growth factor receptor signaling pathway | BP | 14 | 2 | 0 | 0.04297 |
| GO:0030214 | hyaluronan catabolic process | BP | 14 | 2 | 0 | 0.04297 |
| GO:0019865 | immunoglobulin binding | MF | 14 | 2 | 1 | 0.04297 |
| GO:0072234 | metanephric nephron tubule development | BP | 14 | 2 | 0 | 0.04297 |
| GO:0090185 | negative regulation of kidney development | BP | 14 | 2 | 1 | 0.04297 |
| GO:1901017 | negative regulation of potassium ion transmembrane transporter act | BP | 14 | 2 | 1 | 0.04297 |
| GO:0004550 | nucleoside diphosphate kinase activity | MF | 14 | 2 | 0 | 0.04297 |
| GO:0036035 | osteoclast development | BP | 14 | 2 | 0 | 0.04297 |
| GO:0036152 | phosphatidylethanolamine acyl-chain remodeling | BP | 14 | 2 | 2 | 0.04297 |
| GO:0090280 | positive regulation of calcium ion import | BP | 14 | 2 | 1 | 0.04297 |
| GO:0070230 | positive regulation of lymphocyte apoptotic process | BP | 14 | 2 | 0 | 0.04297 |
| GO:0045672 | positive regulation of osteoclast differentiation | BP | 14 | 2 | 0 | 0.04297 |
| GO:1903209 | positive regulation of oxidative stress-induced cell death | BP | 14 | 2 | 0 | 0.04297 |
| GO:0002830 | positive regulation of type 2 immune response | BP | 14 | 2 | 0 | 0.04297 |
| GO:0016540 | protein autoprocessing | BP | 14 | 2 | 0 | 0.04297 |
| GO:0035590 | purinergic nucleotide receptor signaling pathway | BP | 14 | 2 | 0 | 0.04297 |
| GO:0009208 | pyrimidine ribonucleoside triphosphate metabolic process | BP | 14 | 2 | 0 | 0.04297 |
| GO:0090189 | regulation of branching involved in ureteric bud morphogenesis | BP | 14 | 2 | 1 | 0.04297 |
| GO:0071071 | regulation of phospholipid biosynthetic process | BP | 14 | 2 | 0 | 0.04297 |
| GO:0034134 | toll-like receptor 2 signaling pathway | BP | 14 | 2 | 0 | 0.04297 |
| GO:0070633 | transepithelial transport | BP | 14 | 2 | 1 | 0.04297 |
| GO:0048514 | blood vessel morphogenesis | BP | 484 | 18 | 24 | 0.04309 |
| GO:0006875 | cellular metal ion homeostasis | BP | 417 | 16 | 25 | 0.04322 |
| GO:0043271 | negative regulation of ion transport | BP | 107 | 6 | 4 | 0.0434 |
| GO:0051656 | establishment of organelle localization | BP | 451 | 17 | 17 | 0.04363 |
| GO:0034062 | 5'-3' RNA polymerase activity | MF | 33 | 3 | 0 | 0.04365 |
| GO:1990939 | ATP-dependent microtubule motor activity | MF | 33 | 3 | 2 | 0.04365 |
| GO:0070268 | cornification | BP | 33 | 3 | 3 | 0.04365 |
| GO:0002704 | negative regulation of leukocyte mediated immunity | BP | 33 | 3 | 2 | 0.04365 |
| GO:0016504 | peptidase activator activity | MF | 33 | 3 | 0 | 0.04365 |
| GO:0030890 | positive regulation of B cell proliferation | BP | 33 | 3 | 3 | 0.04365 |
| GO:0030501 | positive regulation of bone mineralization | BP | 33 | 3 | 2 | 0.04365 |
| GO:2001222 | regulation of neuron migration | BP | 33 | 3 | 1 | 0.04365 |
| GO:2000404 | regulation of T cell migration | BP | 33 | 3 | 0 | 0.04365 |
| GO:0097747 | RNA polymerase activity | MF | 33 | 3 | 0 | 0.04365 |
| GO:0014902 | myotube differentiation | BP | 81 | 5 | 2 | 0.04461 |
| GO:0051188 | cofactor biosynthetic process | BP | 288 | 12 | 27 | 0.04462 |
| GO:0090398 | cellular senescence | BP | 56 | 4 | 2 | 0.04472 |
| GO:0008194 | UDP-glycosyltransferase activity | MF | 108 | 6 | 3 | 0.04507 |
| GO:0002253 | activation of immune response | BP | 487 | 18 | 13 | 0.04528 |
| GO:1901565 | organonitrogen compound catabolic process | BP | 1092 | 35 | 48 | 0.04605 |
| GO:0010721 | negative regulation of cell development | BP | 258 | 11 | 11 | 0.04638 |
| GO:0031406 | carboxylic acid binding | MF | 137 | 7 | 16 | 0.0465 |

| | | | | | | |
|---|---|---|---|---|---|---|
| GO:0034341 | response to interferon-gamma | BP | 137 | 7 | 2 | 0.0465 |
| GO:0002367 | cytokine production involved in immune response | BP | 82 | 5 | 5 | 0.04662 |
| GO:0050708 | regulation of protein secretion | BP | 355 | 14 | 18 | 0.0467 |
| GO:0032392 | DNA geometric change | BP | 109 | 6 | 1 | 0.04678 |
| GO:1904064 | positive regulation of cation transmembrane transport | BP | 109 | 6 | 5 | 0.04678 |
| GO:0034508 | centromere complex assembly | BP | 34 | 3 | 0 | 0.04705 |
| GO:0035337 | fatty-acyl-CoA metabolic process | BP | 34 | 3 | 3 | 0.04705 |
| GO:0045429 | positive regulation of nitric oxide biosynthetic process | BP | 34 | 3 | 3 | 0.04705 |
| GO:0001941 | postsynaptic membrane organization | BP | 34 | 3 | 0 | 0.04705 |
| GO:0051445 | regulation of meiotic cell cycle | BP | 34 | 3 | 2 | 0.04705 |
| GO:0009268 | response to pH | BP | 34 | 3 | 2 | 0.04705 |
| GO:0005764 | lysosome | CC | 558 | 20 | 33 | 0.04711 |
| GO:0009258 | 10-formyltetrahydrofolate catabolic process | BP | 2 | 1 | 0 | 0.04729 |
| GO:0031247 | actin rod assembly | BP | 2 | 1 | 1 | 0.04729 |
| GO:1905069 | allantois development | BP | 2 | 1 | 0 | 0.04729 |
| GO:0080144 | amino acid homeostasis | BP | 2 | 1 | 0 | 0.04729 |
| GO:1990184 | amino acid transport complex | CC | 2 | 1 | 1 | 0.04729 |
| GO:0003275 | apoptotic process involved in outflow tract morphogenesis | BP | 2 | 1 | 0 | 0.04729 |
| GO:0006529 | asparagine biosynthetic process | BP | 2 | 1 | 0 | 0.04729 |
| GO:0004066 | asparagine synthase (glutamine-hydrolyzing) activity | MF | 2 | 1 | 0 | 0.04729 |
| GO:0060018 | astrocyte fate commitment | BP | 2 | 1 | 0 | 0.04729 |
| GO:0002368 | B cell cytokine production | BP | 2 | 1 | 0 | 0.04729 |
| GO:0034769 | basement membrane disassembly | BP | 2 | 1 | 0 | 0.04729 |
| GO:0034512 | box C/D snoRNA binding | MF | 2 | 1 | 0 | 0.04729 |
| GO:0060532 | bronchus cartilage development | BP | 2 | 1 | 0 | 0.04729 |
| GO:1904954 | canonical Wnt signaling pathway involved in midbrain dopaminerg | BP | 2 | 1 | 0 | 0.04729 |
| GO:0060722 | cell proliferation involved in embryonic placenta development | BP | 2 | 1 | 0 | 0.04729 |
| GO:0071283 | cellular response to iron(III) ion | BP | 2 | 1 | 0 | 0.04729 |
| GO:1904482 | cellular response to tetrahydrofolate | BP | 2 | 1 | 0 | 0.04729 |
| GO:0097533 | cellular stress response to acid chemical | BP | 2 | 1 | 1 | 0.04729 |
| GO:0021551 | central nervous system morphogenesis | BP | 2 | 1 | 0 | 0.04729 |
| GO:0021698 | cerebellar cortex structural organization | BP | 2 | 1 | 0 | 0.04729 |
| GO:0021693 | cerebellar Purkinje cell layer structural organization | BP | 2 | 1 | 0 | 0.04729 |
| GO:0006032 | chitin catabolic process | BP | 2 | 1 | 0 | 0.04729 |
| GO:0006030 | chitin metabolic process | BP | 2 | 1 | 0 | 0.04729 |
| GO:0004568 | chitinase activity | MF | 2 | 1 | 0 | 0.04729 |
| GO:0050897 | cobalt ion binding | MF | 2 | 1 | 0 | 0.04729 |
| GO:0097058 | CRLF-CLCF1 complex | CC | 2 | 1 | 0 | 0.04729 |
| GO:0019343 | cysteine biosynthetic process via cystathionine | BP | 2 | 1 | 0 | 0.04729 |
| GO:0060327 | cytoplasmic actin-based contraction involved in cell motility | BP | 2 | 1 | 0 | 0.04729 |
| GO:0031751 | D4 dopamine receptor binding | MF | 2 | 1 | 0 | 0.04729 |
| GO:0001651 | dense fibrillar component | CC | 2 | 1 | 0 | 0.04729 |
| GO:0009159 | deoxyribonucleoside monophosphate catabolic process | BP | 2 | 1 | 0 | 0.04729 |
| GO:0042496 | detection of diacyl bacterial lipopeptide | BP | 2 | 1 | 0 | 0.04729 |
| GO:0016046 | detection of fungus | BP | 2 | 1 | 0 | 0.04729 |
| GO:0032497 | detection of lipopolysaccharide | BP | 2 | 1 | 1 | 0.04729 |
| GO:0042495 | detection of triacyl bacterial lipopeptide | BP | 2 | 1 | 0 | 0.04729 |
| GO:0071633 | dihydroceramidase activity | MF | 2 | 1 | 0 | 0.04729 |
| GO:0016403 | dimethylargininase activity | MF | 2 | 1 | 0 | 0.04729 |
| GO:0030337 | DNA polymerase processivity factor activity | MF | 2 | 1 | 0 | 0.04729 |
| GO:0009330 | DNA topoisomerase complex (ATP-hydrolyzing) | CC | 2 | 1 | 0 | 0.04729 |
| GO:0061505 | DNA topoisomerase II activity | MF | 2 | 1 | 0 | 0.04729 |
| GO:0003918 | DNA topoisomerase type II (ATP-hydrolyzing) activity | MF | 2 | 1 | 0 | 0.04729 |
| GO:1990918 | double-strand break repair involved in meiotic recombination | BP | 2 | 1 | 0 | 0.04729 |
| GO:0097681 | double-strand break repair via alternative nonhomologous end joini | BP | 2 | 1 | 0 | 0.04729 |
| GO:1903981 | enterobactin binding | MF | 2 | 1 | 0 | 0.04729 |
| GO:0007174 | epidermal growth factor catabolic process | BP | 2 | 1 | 0 | 0.04729 |
| GO:0010142 | farnesyl diphosphate biosynthetic process, mevalonate pathway | BP | 2 | 1 | 1 | 0.04729 |
| GO:0002316 | follicular B cell differentiation | BP | 2 | 1 | 0 | 0.04729 |
| GO:0016155 | formyltetrahydrofolate dehydrogenase activity | MF | 2 | 1 | 0 | 0.04729 |
| GO:0009449 | gamma-aminobutyric acid biosynthetic process | BP | 2 | 1 | 0 | 0.04729 |
| GO:0046368 | GDP-L-fucose metabolic process | BP | 2 | 1 | 0 | 0.04729 |
| GO:0018992 | germ-line sex determination | BP | 2 | 1 | 0 | 0.04729 |
| GO:0072139 | glomerular parietal epithelial cell differentiation | BP | 2 | 1 | 0 | 0.04729 |

| | | | | | | |
|---|---|---|---|---|---|---|
| GO:0090461 | glutamate homeostasis | BP | 2 | 1 | 0 | 0.04729 |
| GO:0034775 | glutathione transmembrane transport | BP | 2 | 1 | 0 | 0.04729 |
| GO:0006545 | glycine biosynthetic process | BP | 2 | 1 | 0 | 0.04729 |
| GO:0019264 | glycine biosynthetic process from serine | BP | 2 | 1 | 0 | 0.04729 |
| GO:0004372 | glycine hydroxymethyltransferase activity | MF | 2 | 1 | 0 | 0.04729 |
| GO:0003920 | GMP reductase activity | MF | 2 | 1 | 0 | 0.04729 |
| GO:1902560 | GMP reductase complex | CC | 2 | 1 | 0 | 0.04729 |
| GO:0071613 | granzyme B production | BP | 2 | 1 | 0 | 0.04729 |
| GO:0036117 | hyaluranon cable | CC | 2 | 1 | 0 | 0.04729 |
| GO:0004421 | hydroxymethylglutaryl-CoA synthase activity | MF | 2 | 1 | 1 | 0.04729 |
| GO:0002436 | immune complex clearance by monocytes and macrophages | BP | 2 | 1 | 0 | 0.04729 |
| GO:0002426 | immunoglobulin production in mucosal tissue | BP | 2 | 1 | 0 | 0.04729 |
| GO:0002414 | immunoglobulin transcytosis in epithelial cells | BP | 2 | 1 | 1 | 0.04729 |
| GO:0051389 | inactivation of MAPKK activity | BP | 2 | 1 | 0 | 0.04729 |
| GO:0033754 | indoleamine 2,3-dioxygenase activity | MF | 2 | 1 | 1 | 0.04729 |
| GO:0034686 | integrin alphav-beta8 complex | CC | 2 | 1 | 0 | 0.04729 |
| GO:0038043 | interleukin-5-mediated signaling pathway | BP | 2 | 1 | 0 | 0.04729 |
| GO:0004914 | interleukin-5 receptor activity | MF | 2 | 1 | 0 | 0.04729 |
| GO:1902767 | isoprenoid biosynthetic process via mevalonate | BP | 2 | 1 | 1 | 0.04729 |
| GO:0034276 | kynurenic acid biosynthetic process | BP | 2 | 1 | 0 | 0.04729 |
| GO:0034275 | kynurenic acid metabolic process | BP | 2 | 1 | 0 | 0.04729 |
| GO:0008732 | L-allo-threonine aldolase activity | MF | 2 | 1 | 0 | 0.04729 |
| GO:0140009 | L-aspartate import across plasma membrane | BP | 2 | 1 | 0 | 0.04729 |
| GO:1904556 | L-tryptophan transmembrane transport | BP | 2 | 1 | 0 | 0.04729 |
| GO:0015196 | L-tryptophan transmembrane transporter activity | MF | 2 | 1 | 0 | 0.04729 |
| GO:0050902 | leukocyte adhesive activation | BP | 2 | 1 | 0 | 0.04729 |
| GO:0140042 | lipid droplet formation | BP | 2 | 1 | 0 | 0.04729 |
| GO:0061145 | lung smooth muscle development | BP | 2 | 1 | 0 | 0.04729 |
| GO:0035692 | macrophage migration inhibitory factor receptor complex | CC | 2 | 1 | 0 | 0.04729 |
| GO:0019100 | male germ-line sex determination | BP | 2 | 1 | 0 | 0.04729 |
| GO:0000706 | meiotic DNA double-strand break processing | BP | 2 | 1 | 0 | 0.04729 |
| GO:1901145 | mesenchymal cell apoptotic process involved in nephron morphoge | BP | 2 | 1 | 0 | 0.04729 |
| GO:0072038 | mesenchymal stem cell maintenance involved in nephron morphoge | BP | 2 | 1 | 0 | 0.04729 |
| GO:0072185 | metanephric cap development | BP | 2 | 1 | 0 | 0.04729 |
| GO:0090094 | metanephric cap mesenchymal cell proliferation involved in metane | BP | 2 | 1 | 0 | 0.04729 |
| GO:0072186 | metanephric cap morphogenesis | BP | 2 | 1 | 0 | 0.04729 |
| GO:0072174 | metanephric tubule formation | BP | 2 | 1 | 0 | 0.04729 |
| GO:0002397 | MHC class I protein complex assembly | BP | 2 | 1 | 0 | 0.04729 |
| GO:0070853 | myosin VI binding | MF | 2 | 1 | 1 | 0.04729 |
| GO:0033842 | N-acetyl-beta-glucosaminyl-glycoprotein 4-beta-N-acetylgalactosa | MF | 2 | 1 | 0 | 0.04729 |
| GO:0043132 | NAD transport | BP | 2 | 1 | 0 | 0.04729 |
| GO:0070401 | NADP+ binding | MF | 2 | 1 | 0 | 0.04729 |
| GO:0031262 | Ndc80 complex | CC | 2 | 1 | 0 | 0.04729 |
| GO:0002865 | negative regulation of acute inflammatory response to antigenic stir | BP | 2 | 1 | 0 | 0.04729 |
| GO:0090191 | negative regulation of branching involved in ureteric bud morphoge | BP | 2 | 1 | 0 | 0.04729 |
| GO:0106135 | negative regulation of cardiac muscle cell contraction | BP | 2 | 1 | 0 | 0.04729 |
| GO:0045957 | negative regulation of complement activation, alternative pathway | BP | 2 | 1 | 0 | 0.04729 |
| GO:1905408 | negative regulation of creatine transmembrane transporter activity | BP | 2 | 1 | 0 | 0.04729 |
| GO:1905450 | negative regulation of Fc-gamma receptor signaling pathway involv | BP | 2 | 1 | 0 | 0.04729 |
| GO:1900450 | negative regulation of glutamate receptor signaling pathway | BP | 2 | 1 | 0 | 0.04729 |
| GO:0002884 | negative regulation of hypersensitivity | BP | 2 | 1 | 0 | 0.04729 |
| GO:0033624 | negative regulation of integrin activation | BP | 2 | 1 | 0 | 0.04729 |
| GO:0045081 | negative regulation of interleukin-10 biosynthetic process | BP | 2 | 1 | 0 | 0.04729 |
| GO:0032707 | negative regulation of interleukin-23 production | BP | 2 | 1 | 0 | 0.04729 |
| GO:0072040 | negative regulation of mesenchymal cell apoptotic process involvec | BP | 2 | 1 | 0 | 0.04729 |
| GO:1901740 | negative regulation of myoblast fusion | BP | 2 | 1 | 0 | 0.04729 |
| GO:1900138 | negative regulation of phospholipase A2 activity | BP | 2 | 1 | 0 | 0.04729 |
| GO:1902303 | negative regulation of potassium ion export | BP | 2 | 1 | 0 | 0.04729 |
| GO:1903765 | negative regulation of potassium ion export across plasma membrai | BP | 2 | 1 | 0 | 0.04729 |
| GO:0060051 | negative regulation of protein glycosylation | BP | 2 | 1 | 0 | 0.04729 |
| GO:0090285 | negative regulation of protein glycosylation in Golgi | BP | 2 | 1 | 0 | 0.04729 |
| GO:1904673 | negative regulation of somatic stem cell population maintenance | BP | 2 | 1 | 0 | 0.04729 |
| GO:1903946 | negative regulation of ventricular cardiac muscle cell action potenti | BP | 2 | 1 | 0 | 0.04729 |
| GO:0072076 | nephrogenic mesenchyme development | BP | 2 | 1 | 0 | 0.04729 |

| | | | | | | |
|---|---|---|---|---|---|---|
| GO:0072134 | nephrogenic mesenchyme morphogenesis | BP | 2 | 1 | 0 | 0.04729 |
| GO:0014034 | neural crest cell fate commitment | BP | 2 | 1 | 0 | 0.04729 |
| GO:0004983 | neuropeptide Y receptor activity | MF | 2 | 1 | 0 | 0.04729 |
| GO:0070026 | nitric oxide binding | MF | 2 | 1 | 0 | 0.04729 |
| GO:0002537 | nitric oxide production involved in inflammatory response | BP | 2 | 1 | 0 | 0.04729 |
| GO:0015391 | nucleobase:cation symporter activity | MF | 2 | 1 | 1 | 0.04729 |
| GO:0070538 | oleic acid binding | MF | 2 | 1 | 0 | 0.04729 |
| GO:0016657 | oxidoreductase activity, acting on NAD(P)H, nitrogenous group as | MF | 2 | 1 | 0 | 0.04729 |
| GO:0070557 | PCNA-p21 complex | CC | 2 | 1 | 0 | 0.04729 |
| GO:1990667 | PCSK9-AnxA2 complex | CC | 2 | 1 | 0 | 0.04729 |
| GO:1990666 | PCSK9-LDLR complex | CC | 2 | 1 | 0 | 0.04729 |
| GO:1904090 | peptidase inhibitor complex | CC | 2 | 1 | 0 | 0.04729 |
| GO:0002502 | peptide antigen assembly with MHC class I protein complex | BP | 2 | 1 | 0 | 0.04729 |
| GO:0002501 | peptide antigen assembly with MHC protein complex | BP | 2 | 1 | 0 | 0.04729 |
| GO:0005052 | peroxisome matrix targeting signal-1 binding | MF | 2 | 1 | 0 | 0.04729 |
| GO:0097195 | pilomotor reflex | BP | 2 | 1 | 0 | 0.04729 |
| GO:1901953 | positive regulation of anterograde dense core granule transport | BP | 2 | 1 | 0 | 0.04729 |
| GO:2000538 | positive regulation of B cell chemotaxis | BP | 2 | 1 | 0 | 0.04729 |
| GO:1904172 | positive regulation of bleb assembly | BP | 2 | 1 | 0 | 0.04729 |
| GO:0002678 | positive regulation of chronic inflammatory response | BP | 2 | 1 | 0 | 0.04729 |
| GO:1904811 | positive regulation of dense core granule transport | BP | 2 | 1 | 0 | 0.04729 |
| GO:0061184 | positive regulation of dermatome development | BP | 2 | 1 | 0 | 0.04729 |
| GO:0071663 | positive regulation of granzyme B production | BP | 2 | 1 | 0 | 0.04729 |
| GO:0110090 | positive regulation of hippocampal neuron apoptotic process | BP | 2 | 1 | 0 | 0.04729 |
| GO:0010729 | positive regulation of hydrogen peroxide biosynthetic process | BP | 2 | 1 | 0 | 0.04729 |
| GO:1903797 | positive regulation of inorganic anion transmembrane transport | BP | 2 | 1 | 0 | 0.04729 |
| GO:0050717 | positive regulation of interleukin-1 alpha secretion | BP | 2 | 1 | 0 | 0.04729 |
| GO:0032805 | positive regulation of low-density lipoprotein particle receptor cata | BP | 2 | 1 | 0 | 0.04729 |
| GO:1905458 | positive regulation of lymphoid progenitor cell differentiation | BP | 2 | 1 | 0 | 0.04729 |
| GO:0034241 | positive regulation of macrophage fusion | BP | 2 | 1 | 0 | 0.04729 |
| GO:1903490 | positive regulation of mitotic cytokinesis | BP | 2 | 1 | 0 | 0.04729 |
| GO:1903438 | positive regulation of mitotic cytokinetic process | BP | 2 | 1 | 0 | 0.04729 |
| GO:2000768 | positive regulation of nephron tubule epithelial cell differentiation | BP | 2 | 1 | 0 | 0.04729 |
| GO:0070434 | positive regulation of nucleotide-binding oligomerization domain c | BP | 2 | 1 | 0 | 0.04729 |
| GO:0070426 | positive regulation of nucleotide-binding oligomerization domain c | BP | 2 | 1 | 0 | 0.04729 |
| GO:1903615 | positive regulation of protein tyrosine phosphatase activity | BP | 2 | 1 | 0 | 0.04729 |
| GO:0045870 | positive regulation of single stranded viral RNA replication via dou | BP | 2 | 1 | 0 | 0.04729 |
| GO:1904395 | positive regulation of skeletal muscle acetylcholine-gated channel c | BP | 2 | 1 | 0 | 0.04729 |
| GO:2000845 | positive regulation of testosterone secretion | BP | 2 | 1 | 0 | 0.04729 |
| GO:0070172 | positive regulation of tooth mineralization | BP | 2 | 1 | 0 | 0.04729 |
| GO:0001812 | positive regulation of type I hypersensitivity | BP | 2 | 1 | 0 | 0.04729 |
| GO:0001805 | positive regulation of type III hypersensitivity | BP | 2 | 1 | 0 | 0.04729 |
| GO:1902943 | positive regulation of voltage-gated chloride channel activity | BP | 2 | 1 | 0 | 0.04729 |
| GO:0060516 | primary prostatic bud elongation | BP | 2 | 1 | 0 | 0.04729 |
| GO:0007538 | primary sex determination | BP | 2 | 1 | 0 | 0.04729 |
| GO:0007542 | primary sex determination, germ-line | BP | 2 | 1 | 0 | 0.04729 |
| GO:0002572 | pro-T cell differentiation | BP | 2 | 1 | 0 | 0.04729 |
| GO:0097179 | protease inhibitor complex | CC | 2 | 1 | 0 | 0.04729 |
| GO:0036009 | protein-glutamine N-methyltransferase activity | MF | 2 | 1 | 0 | 0.04729 |
| GO:0044314 | protein K27-linked ubiquitination | BP | 2 | 1 | 0 | 0.04729 |
| GO:0070560 | protein secretion by platelet | BP | 2 | 1 | 0 | 0.04729 |
| GO:0072047 | proximal/distal pattern formation involved in nephron development | BP | 2 | 1 | 0 | 0.04729 |
| GO:0000701 | purine-specific mismatch base pair DNA N-glycosylase activity | MF | 2 | 1 | 0 | 0.04729 |
| GO:1904823 | purine nucleobase transmembrane transport | BP | 2 | 1 | 1 | 0.04729 |
| GO:0015860 | purine nucleoside transmembrane transport | BP | 2 | 1 | 1 | 0.04729 |
| GO:0015950 | purine nucleotide interconversion | BP | 2 | 1 | 0 | 0.04729 |
| GO:0015951 | purine ribonucleotide interconversion | BP | 2 | 1 | 0 | 0.04729 |
| GO:0008615 | pyridoxine biosynthetic process | BP | 2 | 1 | 0 | 0.04729 |
| GO:0008614 | pyridoxine metabolic process | BP | 2 | 1 | 0 | 0.04729 |
| GO:0015389 | pyrimidine- and adenine-specific:sodium symporter activity | MF | 2 | 1 | 1 | 0.04729 |
| GO:0009213 | pyrimidine deoxyribonucleoside triphosphate catabolic process | BP | 2 | 1 | 0 | 0.04729 |
| GO:0005350 | pyrimidine nucleobase transmembrane transporter activity | MF | 2 | 1 | 1 | 0.04729 |
| GO:0009149 | pyrimidine nucleoside triphosphate catabolic process | BP | 2 | 1 | 0 | 0.04729 |
| GO:0004735 | pyrroline-5-carboxylate reductase activity | MF | 2 | 1 | 0 | 0.04729 |

| | | | | | | |
|---|---|---|---|---|---|---|
| GO:0038025 | reelin receptor activity | MF | 2 | 1 | 0 | 0.04729 |
| GO:1904717 | regulation of AMPA glutamate receptor clustering | BP | 2 | 1 | 0 | 0.04729 |
| GO:1901951 | regulation of anterograde dense core granule transport | BP | 2 | 1 | 0 | 0.04729 |
| GO:1902256 | regulation of apoptotic process involved in outflow tract morphoge | BP | 2 | 1 | 0 | 0.04729 |
| GO:0009786 | regulation of asymmetric cell division | BP | 2 | 1 | 0 | 0.04729 |
| GO:2000537 | regulation of B cell chemotaxis | BP | 2 | 1 | 0 | 0.04729 |
| GO:1904170 | regulation of bleb assembly | BP | 2 | 1 | 0 | 0.04729 |
| GO:0060723 | regulation of cell proliferation involved in embryonic placenta deve | BP | 2 | 1 | 0 | 0.04729 |
| GO:0030451 | regulation of complement activation, alternative pathway | BP | 2 | 1 | 0 | 0.04729 |
| GO:1903433 | regulation of constitutive secretory pathway | BP | 2 | 1 | 0 | 0.04729 |
| GO:1905407 | regulation of creatine transmembrane transporter activity | BP | 2 | 1 | 0 | 0.04729 |
| GO:1904809 | regulation of dense core granule transport | BP | 2 | 1 | 0 | 0.04729 |
| GO:0106104 | regulation of glutamate receptor clustering | BP | 2 | 1 | 0 | 0.04729 |
| GO:1903786 | regulation of glutathione biosynthetic process | BP | 2 | 1 | 0 | 0.04729 |
| GO:0071661 | regulation of granzyme B production | BP | 2 | 1 | 0 | 0.04729 |
| GO:1903547 | regulation of growth hormone activity | BP | 2 | 1 | 0 | 0.04729 |
| GO:0090264 | regulation of immune complex clearance by monocytes and macrop | BP | 2 | 1 | 0 | 0.04729 |
| GO:1903487 | regulation of lactation | BP | 2 | 1 | 0 | 0.04729 |
| GO:1905595 | regulation of low-density lipoprotein particle receptor binding | BP | 2 | 1 | 0 | 0.04729 |
| GO:0034239 | regulation of macrophage fusion | BP | 2 | 1 | 0 | 0.04729 |
| GO:0072039 | regulation of mesenchymal cell apoptotic process involved in neph | BP | 2 | 1 | 0 | 0.04729 |
| GO:1903436 | regulation of mitotic cytokinetic process | BP | 2 | 1 | 0 | 0.04729 |
| GO:0070428 | regulation of nucleotide-binding oligomerization domain containin | BP | 2 | 1 | 0 | 0.04729 |
| GO:2000275 | regulation of oxidative phosphorylation uncoupler activity | BP | 2 | 1 | 0 | 0.04729 |
| GO:1901629 | regulation of presynaptic membrane organization | BP | 2 | 1 | 0 | 0.04729 |
| GO:2001228 | regulation of response to gamma radiation | BP | 2 | 1 | 0 | 0.04729 |
| GO:0060721 | regulation of spongiotrophoblast cell proliferation | BP | 2 | 1 | 0 | 0.04729 |
| GO:2000843 | regulation of testosterone secretion | BP | 2 | 1 | 0 | 0.04729 |
| GO:2000609 | regulation of thyroid hormone generation | BP | 2 | 1 | 0 | 0.04729 |
| GO:0001803 | regulation of type III hypersensitivity | BP | 2 | 1 | 0 | 0.04729 |
| GO:1902941 | regulation of voltage-gated chloride channel activity | BP | 2 | 1 | 0 | 0.04729 |
| GO:0072034 | renal vesicle induction | BP | 2 | 1 | 0 | 0.04729 |
| GO:1990418 | response to insulin-like growth factor stimulus | BP | 2 | 1 | 0 | 0.04729 |
| GO:0009624 | response to nematode | BP | 2 | 1 | 0 | 0.04729 |
| GO:1904481 | response to tetrahydrofolate | BP | 2 | 1 | 0 | 0.04729 |
| GO:0060221 | retinal rod cell differentiation | BP | 2 | 1 | 0 | 0.04729 |
| GO:0004522 | ribonuclease A activity | MF | 2 | 1 | 0 | 0.04729 |
| GO:0035985 | senescence-associated heterochromatin focus | CC | 2 | 1 | 0 | 0.04729 |
| GO:0035986 | senescence-associated heterochromatin focus assembly | BP | 2 | 1 | 0 | 0.04729 |
| GO:0072081 | specification of nephron tubule identity | BP | 2 | 1 | 0 | 0.04729 |
| GO:0060061 | Spemann organizer formation | BP | 2 | 1 | 0 | 0.04729 |
| GO:0007284 | spermatogonial cell division | BP | 2 | 1 | 0 | 0.04729 |
| GO:0006597 | spermine biosynthetic process | BP | 2 | 1 | 0 | 0.04729 |
| GO:0060720 | spongiotrophoblast cell proliferation | BP | 2 | 1 | 0 | 0.04729 |
| GO:0090214 | spongiotrophoblast layer developmental growth | BP | 2 | 1 | 0 | 0.04729 |
| GO:0097532 | stress response to acid chemical | BP | 2 | 1 | 1 | 0.04729 |
| GO:0036269 | swimming behavior | BP | 2 | 1 | 0 | 0.04729 |
| GO:0020005 | symbiont-containing vacuole membrane | CC | 2 | 1 | 0 | 0.04729 |
| GO:0035711 | T-helper 1 cell activation | BP | 2 | 1 | 0 | 0.04729 |
| GO:0035936 | testosterone secretion | BP | 2 | 1 | 0 | 0.04729 |
| GO:0004793 | threonine aldolase activity | MF | 2 | 1 | 0 | 0.04729 |
| GO:0035354 | Toll-like receptor 1-Toll-like receptor 2 protein complex | CC | 2 | 1 | 0 | 0.04729 |
| GO:0035377 | transepithelial water transport | BP | 2 | 1 | 0 | 0.04729 |
| GO:0035443 | tripeptide transmembrane transport | BP | 2 | 1 | 0 | 0.04729 |
| GO:0106050 | tRNA 2'-O-methyltransferase activity | MF | 2 | 1 | 0 | 0.04729 |
| GO:0004833 | tryptophan 2,3-dioxygenase activity | MF | 2 | 1 | 1 | 0.04729 |
| GO:0001802 | type III hypersensitivity | BP | 2 | 1 | 0 | 0.04729 |
| GO:0003977 | UDP-N-acetylglucosamine diphosphorylase activity | MF | 2 | 1 | 0 | 0.04729 |
| GO:0034189 | very-low-density lipoprotein particle binding | MF | 2 | 1 | 0 | 0.04729 |
| GO:0019035 | viral integration complex | CC | 2 | 1 | 0 | 0.04729 |
| GO:0098655 | cation transmembrane transport | BP | 593 | 21 | 33 | 0.04737 |
| GO:0000323 | lytic vacuole | CC | 559 | 20 | 33 | 0.04783 |
| GO:0010639 | negative regulation of organelle organization | BP | 324 | 13 | 7 | 0.04828 |
| GO:0030879 | mammary gland development | BP | 110 | 6 | 10 | 0.04853 |

| | | | | | | |
|---|---|---|---|---|---|---|
| GO:0030148 | sphingolipid biosynthetic process | BP | 83 | 5 | 2 | 0.04868 |
| GO:0051338 | regulation of transferase activity | BP | 807 | 27 | 23 | 0.0488 |
| GO:0098590 | plasma membrane region | CC | 915 | 30 | 70 | 0.0488 |
| GO:0048532 | anatomical structure arrangement | BP | 15 | 2 | 0 | 0.04882 |
| GO:0071397 | cellular response to cholesterol | BP | 15 | 2 | 2 | 0.04882 |
| GO:0031958 | corticosteroid receptor signaling pathway | BP | 15 | 2 | 2 | 0.04882 |
| GO:0009084 | glutamine family amino acid biosynthetic process | BP | 15 | 2 | 1 | 0.04882 |
| GO:0072074 | kidney mesenchyme development | BP | 15 | 2 | 0 | 0.04882 |
| GO:0072170 | metanephric tubule development | BP | 15 | 2 | 0 | 0.04882 |
| GO:0051782 | negative regulation of cell division | BP | 15 | 2 | 1 | 0.04882 |
| GO:0002251 | organ or tissue specific immune response | BP | 15 | 2 | 2 | 0.04882 |
| GO:0045723 | positive regulation of fatty acid biosynthetic process | BP | 15 | 2 | 3 | 0.04882 |
| GO:0043032 | positive regulation of macrophage activation | BP | 15 | 2 | 1 | 0.04882 |
| GO:0010744 | positive regulation of macrophage derived foam cell differentiation | BP | 15 | 2 | 1 | 0.04882 |
| GO:0032816 | positive regulation of natural killer cell activation | BP | 15 | 2 | 1 | 0.04882 |
| GO:2001224 | positive regulation of neuron migration | BP | 15 | 2 | 1 | 0.04882 |
| GO:0051770 | positive regulation of nitric-oxide synthase biosynthetic process | BP | 15 | 2 | 0 | 0.04882 |
| GO:0002902 | regulation of B cell apoptotic process | BP | 15 | 2 | 1 | 0.04882 |
| GO:0002861 | regulation of inflammatory response to antigenic stimulus | BP | 15 | 2 | 0 | 0.04882 |
| GO:0010984 | regulation of lipoprotein particle clearance | BP | 15 | 2 | 1 | 0.04882 |
| GO:0016074 | snoRNA metabolic process | BP | 15 | 2 | 0 | 0.04882 |
| GO:0044703 | multi-organism reproductive process | BP | 630 | 22 | 24 | 0.04885 |
| GO:0040017 | positive regulation of locomotion | BP | 458 | 17 | 17 | 0.04913 |
| GO:0048232 | male gamete generation | BP | 325 | 13 | 12 | 0.04926 |
| GO:0007186 | G protein-coupled receptor signaling pathway | BP | 561 | 20 | 38 | 0.04929 |
| GO:0051640 | organelle localization | BP | 631 | 22 | 22 | 0.04954 |
| GO:0061572 | actin filament bundle organization | BP | 139 | 7 | 7 | 0.04959 |
| GO:0032989 | cellular component morphogenesis | BP | 953 | 31 | 52 | 0.04979 |
| GO:0070227 | lymphocyte apoptotic process | BP | 58 | 4 | 3 | 0.0498 |
| GO:0061515 | myeloid cell development | BP | 58 | 4 | 2 | 0.0498 |
| GO:0030500 | regulation of bone mineralization | BP | 58 | 4 | 5 | 0.0498 |

Supplemental Table 7. Differentially expressed genes in the jejunum between IBD and non-IBD

**Supp. Table 7a. Genes upregulated in IBD**

| Gene name | logFC | logCPM | F | PValue | FDR |
|---|---|---|---|---|---|
| EOMES | -6.26773 | 4.795712 | 238.9221 | 1.98E-09 | 2.28E-05 |
| SYN3 | -4.17353 | 5.720445 | 228.1557 | 2.33E-09 | 2.28E-05 |
| LOC103795187 | -4.55675 | 3.79983 | 180.8202 | 9.12E-09 | 3.57E-05 |
| GZMK | -4.89805 | 4.464442 | 172.9042 | 1.18E-08 | 3.86E-05 |
| KLRG1 | -5.88748 | 3.721534 | 153.6252 | 2.97E-08 | 7.26E-05 |
| KLRC1 | -4.9475 | 3.527199 | 133.8629 | 5.18E-08 | 0.000113 |
| DUOX2 | -3.40933 | 5.928407 | 122.5008 | 8.59E-08 | 0.000146 |
| KLRB1 | -5.28025 | 2.629178 | 118.2924 | 1.05E-07 | 0.000146 |
| TXK | -3.36023 | 4.941728 | 119.5514 | 9.86E-08 | 0.000146 |
| LOC100895140 | -4.02086 | 5.912802 | 115.6012 | 2.15E-07 | 0.000248 |
| CD3D | -3.78618 | 3.869036 | 104.699 | 2.08E-07 | 0.000248 |
| PDE7A | -2.73119 | 5.34521 | 100.6988 | 2.59E-07 | 0.000281 |
| CD244 | -3.68561 | 3.37237 | 97.5835 | 3.08E-07 | 0.000301 |
| ABCC3 | -2.47864 | 6.861872 | 98.05979 | 3.00E-07 | 0.000301 |
| IKZF2 | -2.67602 | 5.370504 | 92.4573 | 4.15E-07 | 0.000386 |
| CD53 | -2.83256 | 5.323494 | 90.63373 | 4.63E-07 | 0.000412 |
| LOC108588707 | -5.0431 | 5.364641 | 127.3316 | 5.01E-07 | 0.000412 |
| AVPR2 | -2.86618 | 3.943468 | 88.6834 | 5.22E-07 | 0.000412 |
| CD247 | -2.56034 | 4.571049 | 88.52382 | 5.27E-07 | 0.000412 |
| NUP93 | -2.50603 | 5.857745 | 87.11616 | 5.76E-07 | 0.000433 |
| TAGAP | -2.85887 | 4.188696 | 86.22245 | 6.09E-07 | 0.000441 |
| LOC100894383 | -4.31135 | 2.55545 | 82.77349 | 7.62E-07 | 0.000456 |
| AIM2 | -3.91148 | 3.006566 | 85.05951 | 6.56E-07 | 0.000456 |
| CD27 | -3.48817 | 4.607047 | 85.56855 | 7.69E-07 | 0.000456 |
| THEMIS | -3.27922 | 5.134477 | 86.49603 | 6.78E-07 | 0.000456 |
| LOC100391701 | -2.75544 | 5.518541 | 83.43982 | 7.29E-07 | 0.000456 |
| GPX2 | -2.99275 | 4.830252 | 80.43918 | 8.90E-07 | 0.000497 |
| LOC100396924 | -2.99046 | 4.482877 | 80.59521 | 8.81E-07 | 0.000497 |
| SHC3 | -4.06627 | 3.776964 | 84.80491 | 9.17E-07 | 0.000498 |
| KLRK1 | -4.86872 | 5.062932 | 112.2974 | 1.04E-06 | 0.00052 |
| PTPRC | -3.36332 | 7.131769 | 85.46964 | 1.02E-06 | 0.00052 |
| LPXN | -2.63593 | 3.989824 | 78.39021 | 1.02E-06 | 0.00052 |
| HEATR9 | -3.81794 | 2.251433 | 76.4331 | 1.17E-06 | 0.000534 |
| TFAP2A | -3.46682 | 2.656949 | 76.72571 | 1.15E-06 | 0.000534 |
| LOC108591227 | -2.57277 | 4.080455 | 76.59817 | 1.16E-06 | 0.000534 |
| IL18RAP | -4.06138 | 2.323753 | 76.10247 | 1.20E-06 | 0.000534 |
| FYN | -2.19883 | 5.957969 | 74.93555 | 1.31E-06 | 0.000561 |
| IKZF1 | -2.42936 | 6.260121 | 73.91857 | 1.41E-06 | 0.000585 |
| SKAP1 | -3.38222 | 3.241197 | 71.57797 | 1.67E-06 | 0.000653 |
| RASGRP1 | -2.15074 | 5.633474 | 71.5818 | 1.67E-06 | 0.000653 |
| IKZF3 | -3.60237 | 5.220942 | 82.5873 | 1.91E-06 | 0.000725 |
| ATP8B4 | -2.95838 | 4.389976 | 69.4484 | 1.96E-06 | 0.000725 |
| RDM1 | -2.63064 | 3.866767 | 69.43866 | 1.96E-06 | 0.000725 |

| | | | | | |
|---|---|---|---|---|---|
| KLRC2 | -6.48222 | 5.111541 | 126.6293 | 2.20E-06 | 0.000767 |
| CD84 | -2.44017 | 5.598777 | 67.11256 | 2.35E-06 | 0.00078 |
| LOC108592781 | -2.24374 | 5.13526 | 67.28218 | 2.32E-06 | 0.00078 |
| JAKMIP1 | -2.70694 | 4.146603 | 64.74679 | 2.85E-06 | 0.000898 |
| DOCK2 | -2.69334 | 6.523762 | 65.55449 | 2.85E-06 | 0.000898 |
| PLA2G7 | -2.45859 | 4.718895 | 64.80808 | 2.83E-06 | 0.000898 |
| ZNF831 | -3.06328 | 4.46691 | 66.01493 | 3.02E-06 | 0.000923 |
| C12H16orf54 | -2.79807 | 3.775013 | 64.10314 | 3.00E-06 | 0.000923 |
| CRTAM | -6.34073 | 1.513288 | 63.51297 | 3.15E-06 | 0.000936 |
| ITGA4 | -2.50932 | 6.434677 | 63.38693 | 3.18E-06 | 0.000936 |
| RNASE1 | -2.31678 | 5.939168 | 63.29406 | 3.21E-06 | 0.000936 |
| LOC100896501 | -3.18453 | 8.770704 | 67.6651 | 3.45E-06 | 0.000973 |
| LOC108588705 | -2.27719 | 4.352408 | 62.56445 | 3.41E-06 | 0.000973 |
| FAM78A | -2.28399 | 4.681806 | 61.72797 | 3.66E-06 | 0.000993 |
| DOK2 | -3.01513 | 4.032622 | 61.98682 | 3.76E-06 | 0.001008 |
| ST8SIA4 | -2.55768 | 4.118578 | 60.92925 | 3.92E-06 | 0.001035 |
| LOC100414230 | -2.00331 | 4.782956 | 60.71014 | 3.99E-06 | 0.00104 |
| RAB37 | -2.34752 | 5.44925 | 60.48781 | 4.07E-06 | 0.001046 |
| LOC103794002 | -4.48477 | 1.710079 | 59.98993 | 4.25E-06 | 0.001064 |
| TMEM71 | -2.75578 | 3.18316 | 59.31439 | 4.50E-06 | 0.001087 |
| FYB | -2.40787 | 6.397899 | 59.34485 | 4.49E-06 | 0.001087 |
| LOC100407327 | -2.33718 | 3.821222 | 58.67762 | 4.76E-06 | 0.001122 |
| SELPLG | -2.65981 | 4.415366 | 58.06128 | 5.03E-06 | 0.001145 |
| LOC108589051 | -3.70631 | 3.067609 | 58.79461 | 5.35E-06 | 0.001189 |
| IPCEF1 | -2.96521 | 2.90878 | 56.24065 | 5.93E-06 | 0.001275 |
| LOC103795350 | -2.63442 | 4.750967 | 56.56338 | 6.05E-06 | 0.001286 |
| LCP1 | -2.86957 | 7.8152 | 60.36103 | 6.39E-06 | 0.001343 |
| SLAIN1 | -2.28825 | 3.471724 | 54.33638 | 7.09E-06 | 0.001443 |
| CST7 | -1.93954 | 4.557635 | 54.12948 | 7.23E-06 | 0.001456 |
| E2F2 | -2.22527 | 3.625305 | 53.72199 | 7.51E-06 | 0.001486 |
| LOC108594029 | -3.01888 | 3.256255 | 53.17023 | 7.92E-06 | 0.001522 |
| CXCL11 | -7.55535 | 0.9006 | 65.61083 | 8.22E-06 | 0.00153 |
| SCNN1G | -7.51463 | 0.866385 | 65.81933 | 8.10E-06 | 0.00153 |
| LOC100414243 | -2.4316 | 5.932949 | 53.50064 | 8.19E-06 | 0.00153 |
| SAMD3 | -4.29465 | 1.550956 | 52.59235 | 8.37E-06 | 0.001544 |
| CD101 | -3.0362 | 2.93296 | 52.41688 | 8.52E-06 | 0.001545 |
| LOC100897014 | -2.7454 | 3.719812 | 52.13471 | 8.75E-06 | 0.001546 |
| PIGR | -2.37903 | 9.79785 | 51.99356 | 8.88E-06 | 0.001547 |
| SLC7A5 | -2.10015 | 3.931463 | 51.50487 | 9.31E-06 | 0.001583 |
| LOC103793556 | -3.7738 | 2.134295 | 50.81428 | 9.97E-06 | 0.001652 |
| SLAMF6 | -3.01166 | 3.539172 | 51.02124 | 1.03E-05 | 0.001684 |
| RASSF5 | -2.23264 | 4.966034 | 49.67714 | 1.12E-05 | 0.001741 |
| ARHGAP25 | -1.9292 | 5.419824 | 49.15337 | 1.18E-05 | 0.001802 |
| IL18R1 | -3.43323 | 5.014803 | 63.69688 | 1.20E-05 | 0.001812 |
| WIPF1 | -1.6928 | 7.466666 | 48.66034 | 1.24E-05 | 0.001852 |
| ITK | -2.96767 | 4.135691 | 51.62648 | 1.33E-05 | 0.001869 |
| CCR5 | -2.82135 | 2.431277 | 48.12802 | 1.31E-05 | 0.001869 |
| FA2H | -2.60009 | 3.447879 | 48.39421 | 1.28E-05 | 0.001869 |

| | | | | | |
|---|---|---|---|---|---|
| LOC108589742 | -2.31053 | 3.878435 | 48.0025 | 1.33E-05 | 0.001869 |
| UHRF1 | -2.06528 | 4.065442 | 48.25441 | 1.29E-05 | 0.001869 |
| NFATC2 | -1.9582 | 5.824987 | 48.36211 | 1.28E-05 | 0.001869 |
| CYFIP2 | -1.80824 | 5.700274 | 48.1304 | 1.31E-05 | 0.001869 |
| ANKRD44 | -1.75899 | 6.166356 | 48.26159 | 1.29E-05 | 0.001869 |
| SH2D1A | -5.71826 | 2.224777 | 57.28261 | 1.37E-05 | 0.001881 |
| RINL | -2.68988 | 3.489334 | 47.53752 | 1.40E-05 | 0.001881 |
| SERPINB9 | -2.14645 | 3.930694 | 47.58547 | 1.39E-05 | 0.001881 |
| MCM6 | -2.07035 | 4.180517 | 47.42529 | 1.41E-05 | 0.001881 |
| PPIF | -1.79698 | 5.057699 | 47.79246 | 1.36E-05 | 0.001881 |
| CEACAM7 | -2.10634 | 4.215521 | 47.11568 | 1.46E-05 | 0.00191 |
| PIK3CG | -1.99923 | 4.749868 | 46.84007 | 1.50E-05 | 0.001945 |
| LCK | -3.19779 | 4.77935 | 57.03796 | 1.53E-05 | 0.001947 |
| PYCR1 | -2.64223 | 2.915805 | 46.65342 | 1.53E-05 | 0.001947 |
| LOC103794892 | -2.98342 | 2.45813 | 46.25118 | 1.60E-05 | 0.002 |
| CD160 | -1.71795 | 5.289395 | 46.21573 | 1.61E-05 | 0.002 |
| ERAP2 | -2.20919 | 4.367891 | 46.0673 | 1.63E-05 | 0.00202 |
| EPN3 | -2.15927 | 3.771747 | 45.70332 | 1.70E-05 | 0.002088 |
| GCG | -3.47834 | 2.899338 | 47.2521 | 1.72E-05 | 0.002096 |
| RASGEF1A | -3.11621 | 4.040019 | 50.70256 | 1.91E-05 | 0.00219 |
| LOC100403320 | -2.49585 | 3.060367 | 44.8424 | 1.87E-05 | 0.00219 |
| SASH3 | -2.21878 | 5.076017 | 44.95855 | 1.92E-05 | 0.00219 |
| DOCK10 | -1.96867 | 6.568866 | 44.62889 | 1.91E-05 | 0.00219 |
| IL2RB | -3.52644 | 5.754264 | 63.00357 | 1.97E-05 | 0.002235 |
| LOC108590814 | -3.37478 | 3.628161 | 50.42641 | 1.98E-05 | 0.002235 |
| UNC13D | -2.58964 | 4.676631 | 46.90237 | 1.99E-05 | 0.002235 |
| RHOH | -2.81979 | 2.633127 | 44.00828 | 2.05E-05 | 0.002271 |
| NPAS2 | -1.78834 | 4.409956 | 43.51167 | 2.17E-05 | 0.002365 |
| FAM196B | -2.15707 | 3.679289 | 43.2094 | 2.24E-05 | 0.002407 |
| KIAA1462 | -2.06757 | 5.511153 | 43.24858 | 2.23E-05 | 0.002407 |
| HCST | -3.0013 | 2.871142 | 42.7986 | 2.35E-05 | 0.002509 |
| TBX21 | -3.56378 | 2.942825 | 46.01425 | 2.41E-05 | 0.002524 |
| LAPTM5 | -2.4081 | 5.576732 | 45.36701 | 2.42E-05 | 0.002524 |
| SIDT1 | -1.9584 | 4.56028 | 42.5325 | 2.42E-05 | 0.002524 |
| PLXNC1 | -1.70436 | 4.838582 | 42.49593 | 2.43E-05 | 0.002524 |
| CLDND2 | -3.61144 | 2.347142 | 42.77792 | 2.46E-05 | 0.002534 |
| SRGN | -2.06975 | 4.672377 | 42.2775 | 2.49E-05 | 0.002539 |
| LOC103795753 | -7.5475 | 0.888895 | 51.42626 | 2.59E-05 | 0.002587 |
| LOC103788970 | -2.41943 | 3.535154 | 42.00066 | 2.57E-05 | 0.002587 |
| PIK3AP1 | -2.03222 | 5.683863 | 41.95237 | 2.59E-05 | 0.002587 |
| LOC100390041 | -7.20669 | 0.620348 | 50.28719 | 2.70E-05 | 0.00265 |
| LOC108588840 | -7.41562 | 0.781514 | 49.66849 | 2.85E-05 | 0.002769 |
| LOC100415212 | -2.67898 | 4.522366 | 45.16153 | 2.84E-05 | 0.002769 |
| ARAP2 | -1.55318 | 6.574878 | 41.08936 | 2.87E-05 | 0.002773 |
| WFDC2 | -2.77456 | 2.824151 | 40.80786 | 2.96E-05 | 0.002808 |
| IL16 | -1.62846 | 5.01006 | 40.77995 | 2.97E-05 | 0.002808 |
| DENND2D | -1.60035 | 5.820804 | 40.82168 | 2.96E-05 | 0.002808 |
| ARHGDIB | -2.00292 | 5.903045 | 40.30134 | 3.21E-05 | 0.003007 |

| | | | | | |
|---|---|---|---|---|---|
| SP140 | -1.73617 | 4.743155 | 39.99925 | 3.26E-05 | 0.003029 |
| LOC103794640 | -3.83533 | 3.979089 | 54.54004 | 3.40E-05 | 0.003103 |
| P2RX5 | -1.91705 | 5.061523 | 39.62961 | 3.41E-05 | 0.003105 |
| DPEP2 | -2.48265 | 2.670957 | 39.52751 | 3.46E-05 | 0.003105 |
| LOC108588537 | -2.46191 | 2.82944 | 39.53321 | 3.45E-05 | 0.003105 |
| CHST11 | -1.86902 | 5.186879 | 39.31654 | 3.55E-05 | 0.003167 |
| LYZ | -2.00552 | 5.841814 | 39.39863 | 3.64E-05 | 0.003179 |
| CELF2 | -1.57067 | 7.568741 | 39.10067 | 3.64E-05 | 0.003179 |
| TRERF1 | -1.60707 | 5.301953 | 39.03817 | 3.67E-05 | 0.00319 |
| PLA2G3 | -3.06484 | 2.819797 | 39.28745 | 3.71E-05 | 0.003198 |
| LOC108590938 | -2.78749 | 2.673239 | 38.9096 | 3.73E-05 | 0.003198 |
| MKI67 | -2.00064 | 6.588497 | 39.2318 | 3.71E-05 | 0.003198 |
| LOC103790854 | -3.81873 | 1.404015 | 38.83937 | 3.76E-05 | 0.003212 |
| PVRIG | -3.54138 | 1.837874 | 38.72409 | 3.82E-05 | 0.00323 |
| APOBEC3H | -2.69139 | 3.023114 | 38.68918 | 3.83E-05 | 0.00323 |
| LOC100413268 | -1.79731 | 6.469972 | 38.16113 | 4.09E-05 | 0.00342 |
| CHI3L2 | -3.73034 | 2.118481 | 38.86931 | 4.13E-05 | 0.003437 |
| LOC108589056 | -4.74937 | 0.874092 | 37.07326 | 4.70E-05 | 0.003488 |
| LOC108589050 | -3.92154 | 1.022626 | 37.14108 | 4.66E-05 | 0.003488 |
| LOC103793168 | -3.29274 | 1.491242 | 37.15146 | 4.65E-05 | 0.003488 |
| RAC2 | -3.09507 | 5.277363 | 50.06532 | 4.56E-05 | 0.003488 |
| CARD16 | -2.97925 | 2.685697 | 37.93176 | 4.31E-05 | 0.003488 |
| LOC108590090 | -2.84991 | 3.680785 | 40.71955 | 4.36E-05 | 0.003488 |
| RHBDL2 | -2.68489 | 2.574689 | 37.25643 | 4.59E-05 | 0.003488 |
| PPP1R16B | -2.4249 | 5.888575 | 42.26826 | 4.38E-05 | 0.003488 |
| TOX2 | -2.41427 | 4.138392 | 38.46689 | 4.50E-05 | 0.003488 |
| CAMK4 | -1.99919 | 3.174338 | 37.6305 | 4.38E-05 | 0.003488 |
| LIMD2 | -1.91715 | 5.151581 | 37.1108 | 4.68E-05 | 0.003488 |
| ARNTL | -1.79769 | 4.325162 | 37.42072 | 4.49E-05 | 0.003488 |
| CD3E | -3.35224 | 5.389666 | 52.43404 | 4.78E-05 | 0.003524 |
| SPN | -1.89813 | 4.318903 | 36.91389 | 4.80E-05 | 0.003525 |
| SH2D3C | -2.04961 | 4.605284 | 36.86102 | 4.83E-05 | 0.003536 |
| ETS1 | -1.71181 | 7.785176 | 36.7583 | 4.89E-05 | 0.00357 |
| KLRF1 | -3.22424 | 2.512614 | 37.00735 | 5.04E-05 | 0.00361 |
| LOC100409427 | -2.21321 | 4.996993 | 37.94005 | 5.06E-05 | 0.00361 |
| TPST2 | -2.15928 | 4.254814 | 36.48019 | 5.20E-05 | 0.003696 |
| SLAMF1 | -3.22606 | 4.549251 | 47.89252 | 5.22E-05 | 0.003699 |
| LOC100399118 | -2.55402 | 4.314116 | 39.43051 | 5.26E-05 | 0.003707 |
| CCR2 | -2.06846 | 3.111231 | 36.02215 | 5.38E-05 | 0.003773 |
| LOC100895955 | -3.0056 | 3.558101 | 40.00541 | 5.63E-05 | 0.003915 |
| TBC1D10C | -2.84928 | 4.84781 | 43.98 | 5.91E-05 | 0.004052 |
| LOC103794991 | -4.71154 | 1.335008 | 35.48668 | 6.17E-05 | 0.004129 |
| PRKCH | -1.61576 | 6.830715 | 35.08302 | 6.10E-05 | 0.004129 |
| ABLIM2 | -2.33644 | 4.887719 | 37.7941 | 6.33E-05 | 0.004194 |
| LOC100390508 | -3.04819 | 2.072513 | 34.63615 | 6.47E-05 | 0.004248 |
| ARL11 | -2.38399 | 2.420065 | 34.55555 | 6.55E-05 | 0.004266 |
| RGS14 | -2.02938 | 3.882292 | 34.43421 | 6.65E-05 | 0.004322 |
| CYTIP | -3.14488 | 4.186643 | 43.50011 | 6.91E-05 | 0.004373 |

| | | | | | |
|---|---|---|---|---|---|
| SH2D1B | -2.50003 | 2.115368 | 34.18912 | 6.88E-05 | 0.004373 |
| LOC108590082 | -2.43468 | 2.231307 | 34.18948 | 6.88E-05 | 0.004373 |
| TTC39C | -2.02741 | 3.403013 | 34.23953 | 6.83E-05 | 0.004373 |
| LOC100402290 | -1.75585 | 7.04277 | 34.1687 | 6.90E-05 | 0.004373 |
| LOC108589888 | -1.44308 | 6.307214 | 34.27376 | 6.80E-05 | 0.004373 |
| LOC103796360 | -2.15269 | 2.817377 | 34.03717 | 7.02E-05 | 0.00441 |
| KCNAB2 | -1.62502 | 4.899738 | 34.03511 | 7.03E-05 | 0.00441 |
| TRAF3IP3 | -1.96111 | 5.096285 | 34.15849 | 7.17E-05 | 0.004465 |
| LOC100389406 | -4.5627 | 1.200228 | 33.82635 | 7.46E-05 | 0.004563 |
| LOC108593212 | -3.25077 | 2.143551 | 33.81649 | 7.47E-05 | 0.004563 |
| KCNA3 | -2.00814 | 4.166163 | 33.5629 | 7.50E-05 | 0.004567 |
| FLI1 | -1.53714 | 5.070198 | 33.24469 | 7.84E-05 | 0.004665 |
| BIRC5 | -2.41796 | 2.329285 | 32.84833 | 8.28E-05 | 0.004792 |
| GCH1 | -2.0564 | 5.746771 | 34.57953 | 8.21E-05 | 0.004792 |
| SLFN5 | -2.02946 | 4.575697 | 33.12235 | 8.27E-05 | 0.004792 |
| GIMAP4 | -1.83064 | 3.74037 | 32.93077 | 8.19E-05 | 0.004792 |
| LOC103788771 | -2.90376 | 1.801772 | 32.75357 | 8.39E-05 | 0.004813 |
| LOC100392075 | -1.39219 | 6.489949 | 32.76188 | 8.38E-05 | 0.004813 |
| RRM2 | -1.95328 | 3.402605 | 32.55895 | 8.63E-05 | 0.004933 |
| GFOD1 | -1.83738 | 3.636112 | 32.30916 | 8.94E-05 | 0.005081 |
| TIGIT | -4.58965 | 3.16324 | 45.47213 | 9.53E-05 | 0.005287 |
| GFI1 | -2.31524 | 4.17358 | 33.525 | 9.55E-05 | 0.005287 |
| LOC108593906 | -6.76511 | 0.284474 | 37.31493 | 9.65E-05 | 0.005316 |
| LOC100390015 | -2.31437 | 4.18194 | 33.31609 | 9.86E-05 | 0.005383 |
| APOBR | -2.00949 | 5.723247 | 33.02967 | 0.000101 | 0.005472 |
| TNFSF8 | -2.30823 | 2.563769 | 31.31089 | 0.000103 | 0.005504 |
| LOC100895801 | -1.60698 | 4.011792 | 31.33927 | 0.000103 | 0.005504 |
| LOC100411388 | -2.54336 | 3.748517 | 33.48334 | 0.000108 | 0.005646 |
| UBASH3A | -3.36653 | 3.693703 | 40.22887 | 0.000109 | 0.005649 |
| PDE3B | -1.46868 | 6.388859 | 30.99159 | 0.000108 | 0.005649 |
| LOC103789345 | -2.72648 | 2.066012 | 30.7663 | 0.000112 | 0.005752 |
| CD40LG | -4.02652 | 0.770702 | 30.73548 | 0.000112 | 0.005752 |
| DEF6 | -1.68541 | 4.920649 | 30.66717 | 0.000113 | 0.005765 |
| LOC100395199 | -1.57606 | 3.895201 | 30.66218 | 0.000114 | 0.005765 |
| LOC100385441 | -4.39515 | 2.988017 | 41.95953 | 0.000115 | 0.005804 |
| PLEKHF1 | -2.37654 | 4.58464 | 34.08316 | 0.000116 | 0.005804 |
| PSTPIP1 | -2.25268 | 5.462531 | 34.4906 | 0.000116 | 0.005805 |
| LOC100404789 | -2.10152 | 6.196917 | 33.20629 | 0.000118 | 0.005888 |
| ITM2A | -2.30826 | 2.593728 | 30.35314 | 0.000119 | 0.005898 |
| JCHAIN | -3.51595 | 8.120903 | 46.42597 | 0.000119 | 0.005911 |
| ZBP1 | -2.75685 | 2.66913 | 30.45212 | 0.000121 | 0.005953 |
| PIK3CD | -1.84767 | 5.647527 | 30.6563 | 0.000122 | 0.005975 |
| LOC100387227 | -2.38505 | 2.233093 | 30.07492 | 0.000124 | 0.006027 |
| RAB27A | -1.95379 | 5.278798 | 30.9027 | 0.000128 | 0.006144 |
| CEACAM8 | -2.18946 | 2.193359 | 29.82469 | 0.000129 | 0.006166 |
| NCKAP1L | -1.58967 | 5.885705 | 29.69194 | 0.000131 | 0.006252 |
| LY9 | -2.14542 | 4.827529 | 31.86855 | 0.000133 | 0.00631 |
| LOC100409241 | -1.92567 | 11.29177 | 29.58267 | 0.000133 | 0.006319 |

| | | | | | |
|---|---|---|---|---|---|
| BATF | -2.29918 | 3.565098 | 30.03023 | 0.000135 | 0.006365 |
| DNMT1 | -1.52596 | 6.055217 | 29.36343 | 0.000138 | 0.006409 |
| ITGB7 | -2.88602 | 5.883999 | 40.02242 | 0.000139 | 0.006425 |
| DNAH17 | -1.73747 | 4.70195 | 29.26147 | 0.00014 | 0.006448 |
| CD3G | -3.64621 | 3.061931 | 37.00864 | 0.000142 | 0.006449 |
| CD69 | -3.16489 | 1.679313 | 29.17322 | 0.000142 | 0.006449 |
| ASNS | -2.10825 | 4.337124 | 30.15309 | 0.000142 | 0.006449 |
| LOC100400385 | -1.70976 | 4.037526 | 29.21986 | 0.000141 | 0.006449 |
| STAT4 | -2.41596 | 2.536832 | 29.11912 | 0.000143 | 0.006469 |
| LOC103792190 | -2.35598 | 4.096488 | 31.52208 | 0.000145 | 0.006506 |
| BIN2 | -2.14304 | 4.728784 | 31.21538 | 0.000145 | 0.006506 |
| ATP10A | -1.51807 | 5.203592 | 29.04457 | 0.000145 | 0.006506 |
| LOC100396946 | -1.79028 | 4.502933 | 28.98945 | 0.000146 | 0.006509 |
| TXNDC5 | -1.60291 | 5.99409 | 28.94515 | 0.000147 | 0.006539 |
| LOC100404432 | -2.74759 | 6.30078 | 38.49617 | 0.000153 | 0.006686 |
| MCM2 | -1.52846 | 4.380573 | 28.67761 | 0.000153 | 0.006686 |
| IL2RG | -1.46597 | 5.701244 | 28.62461 | 0.000155 | 0.006702 |
| NCR1 | -4.94529 | 2.294029 | 38.20334 | 0.000156 | 0.006722 |
| NR4A2 | -1.88356 | 3.350817 | 28.55658 | 0.000156 | 0.006728 |
| BCL11B | -1.55443 | 5.447909 | 28.53139 | 0.000157 | 0.006728 |
| FAM118A | -1.42458 | 4.564177 | 28.53507 | 0.000157 | 0.006728 |
| CCDC88B | -1.95287 | 5.665876 | 30.1693 | 0.000158 | 0.006753 |
| LOC103796114 | -2.32647 | 2.304209 | 28.42847 | 0.000159 | 0.006763 |
| PRIMPOL | -1.7701 | 3.541243 | 28.46119 | 0.000159 | 0.006763 |
| AKNA | -2.31861 | 7.18861 | 33.78292 | 0.000162 | 0.006847 |
| CARMIL2 | -2.4667 | 5.748379 | 35.199 | 0.000166 | 0.006889 |
| 1-Sep | -2.44377 | 5.046634 | 34.11695 | 0.000165 | 0.006889 |
| FMNL1 | -2.3014 | 5.800247 | 33.48744 | 0.000166 | 0.006889 |
| MVB12B | -1.44579 | 5.315016 | 28.17382 | 0.000166 | 0.006889 |
| LOC108589194 | -5.04242 | 0.459769 | 28.049 | 0.000169 | 0.006925 |
| LOC108593366 | -1.88323 | 3.115434 | 28.04647 | 0.000169 | 0.006925 |
| LOC100395418 | -1.61505 | 5.399182 | 28.06882 | 0.000169 | 0.006925 |
| SAMD9L | -2.01066 | 5.03042 | 29.76218 | 0.00017 | 0.00694 |
| SLC38A5 | -1.6323 | 3.878895 | 27.97543 | 0.000171 | 0.006948 |
| FGD3 | -2.42562 | 5.005666 | 33.62408 | 0.000174 | 0.007026 |
| LOC103788402 | -2.15688 | 2.803743 | 27.77175 | 0.000177 | 0.0071 |
| LOC103794592 | -1.45003 | 5.116876 | 27.69464 | 0.000179 | 0.007144 |
| LOC108590842 | -5.10154 | 0.556946 | 27.60662 | 0.000182 | 0.007157 |
| LMNB1 | -1.48704 | 5.055816 | 27.63778 | 0.000181 | 0.007157 |
| LOC100407409 | -2.8036 | 1.721699 | 27.40365 | 0.000188 | 0.007306 |
| LOC100415219 | -5.19743 | 1.256137 | 30.33514 | 0.000188 | 0.00731 |
| LOC100394455 | -1.75018 | 4.802496 | 27.46621 | 0.000189 | 0.007319 |
| IRF4 | -2.4006 | 3.756598 | 29.42459 | 0.000194 | 0.00745 |
| MB21D1 | -1.76674 | 3.116021 | 27.14906 | 0.000195 | 0.007477 |
| ITGAX | -1.51291 | 5.288418 | 27.10669 | 0.000197 | 0.007513 |
| PTPN7 | -2.65802 | 4.302052 | 33.21233 | 0.000198 | 0.00754 |
| TNFRSF9 | -4.29773 | 1.016634 | 27.19824 | 0.000199 | 0.007565 |
| LOC108588161 | -3.9675 | 4.282406 | 43.46029 | 0.000203 | 0.007603 |

| | | | | | |
|---|---|---|---|---|---|
| SUSD3 | -2.43084 | 2.652676 | 26.88549 | 0.000204 | 0.007603 |
| PARVG | -2.39841 | 3.220572 | 27.58984 | 0.000203 | 0.007603 |
| WAS | -2.08132 | 3.810587 | 27.27434 | 0.000203 | 0.007603 |
| VSIG4 | -1.92301 | 2.622075 | 26.86151 | 0.000205 | 0.007603 |
| GPR68 | -3.0338 | 4.594019 | 36.40419 | 0.000208 | 0.007631 |
| MAPK11 | -1.46762 | 4.034396 | 26.77516 | 0.000208 | 0.007631 |
| PRF1 | -5.04296 | 6.193088 | 56.32974 | 0.00021 | 0.007666 |
| AFMID | -1.83027 | 2.798578 | 26.71246 | 0.00021 | 0.007666 |
| TMC1 | -4.57738 | 2.165536 | 34.0386 | 0.000216 | 0.007825 |
| GPR174 | -3.22335 | 2.729997 | 30.23366 | 0.000217 | 0.007825 |
| LOC100415407 | -1.69903 | 3.292765 | 26.45053 | 0.000219 | 0.007871 |
| LOC108591782 | -2.53752 | 1.711409 | 26.43711 | 0.000219 | 0.007874 |
| AGAP2 | -1.98792 | 4.986875 | 28.25393 | 0.000222 | 0.00793 |
| WNT10B | -2.55492 | 2.712338 | 26.61385 | 0.000223 | 0.007936 |
| CYBB | -1.6952 | 5.063988 | 26.47102 | 0.000224 | 0.007954 |
| ARHGAP9 | -2.26889 | 6.104325 | 31.66394 | 0.000225 | 0.007989 |
| ARRDC2 | -2.03348 | 5.03771 | 28.66727 | 0.000226 | 0.007989 |
| ARHGAP15 | -1.77294 | 3.901362 | 26.22179 | 0.000227 | 0.008012 |
| LOC100388085 | -2.03944 | 2.284944 | 26.2033 | 0.000228 | 0.008022 |
| C2H5orf58 | -2.66813 | 2.986227 | 27.80184 | 0.00023 | 0.008063 |
| CTSS | -1.82001 | 5.883839 | 27.26993 | 0.000233 | 0.008127 |
| KIF18B | -1.94402 | 2.488177 | 25.98805 | 0.000236 | 0.008153 |
| BEND4 | -3.038 | 1.614856 | 25.9106 | 0.00024 | 0.008201 |
| MUC4 | -3.36944 | 1.710954 | 26.32643 | 0.000242 | 0.00826 |
| P4HA3 | -2.84502 | 2.233284 | 26.10118 | 0.000244 | 0.008331 |
| LOC100390394 | -3.19198 | 3.815792 | 34.068 | 0.000249 | 0.008346 |
| GRAP2 | -2.83916 | 5.028774 | 34.87278 | 0.000247 | 0.008346 |
| CARD9 | -1.96269 | 3.083682 | 25.77371 | 0.000245 | 0.008346 |
| KLHL4 | -1.75431 | 3.095847 | 25.69523 | 0.000248 | 0.008346 |
| LOC103795186 | -1.52172 | 3.957316 | 25.75163 | 0.000246 | 0.008346 |
| KCNQ5 | -1.49272 | 4.122485 | 25.66405 | 0.00025 | 0.008346 |
| TNFAIP8 | -1.29488 | 5.196466 | 25.71822 | 0.000247 | 0.008346 |
| M6PR | -1.26721 | 5.597366 | 25.70768 | 0.000248 | 0.008346 |
| STK4 | -1.22299 | 6.08578 | 25.71462 | 0.000247 | 0.008346 |
| ADGRE5 | -1.32451 | 5.767821 | 25.5368 | 0.000255 | 0.008494 |
| ADORA2A | -1.89724 | 2.59204 | 25.49264 | 0.000257 | 0.008514 |
| HN1 | -1.82891 | 4.547685 | 25.69234 | 0.000263 | 0.008598 |
| HSP90B1 | -1.58559 | 8.786654 | 25.36878 | 0.000262 | 0.008598 |
| SORD | -1.51961 | 3.964734 | 25.32589 | 0.000264 | 0.008641 |
| LPCAT2 | -2.22345 | 2.480563 | 25.24231 | 0.000268 | 0.008673 |
| CD48 | -1.92131 | 3.631395 | 25.24244 | 0.000268 | 0.008673 |
| LOC103788928 | -1.82672 | 3.107931 | 25.25083 | 0.000268 | 0.008673 |
| CLCA4 | -4.09636 | 6.711916 | 46.49493 | 0.000273 | 0.008744 |
| KIF11 | -2.16651 | 2.793189 | 25.12409 | 0.000274 | 0.008744 |
| LOC100390922 | -4.00223 | 6.34165 | 45.52654 | 0.00028 | 0.008837 |
| TOP2A | -1.96215 | 4.140089 | 25.64486 | 0.000278 | 0.008837 |
| TMC8 | -1.61492 | 5.429986 | 25.16784 | 0.000278 | 0.008837 |
| C21H21orf62 | -2.4917 | 1.416056 | 24.96214 | 0.000281 | 0.008868 |

| | | | | | |
|---|---|---|---|---|---|
| SERPINB1 | -1.66689 | 4.899919 | 24.96003 | 0.000288 | 0.009053 |
| HMGCS1 | -1.95853 | 7.605188 | 27.13835 | 0.00029 | 0.009105 |
| GSAP | -1.37616 | 4.890106 | 24.743 | 0.000292 | 0.009135 |
| KRT18 | -1.4694 | 6.943123 | 24.69181 | 0.000295 | 0.009172 |
| PF4 | -2.43982 | 1.621424 | 24.66978 | 0.000296 | 0.009192 |
| CARD11 | -2.16511 | 4.632516 | 27.69903 | 0.000296 | 0.009192 |
| GIMAP6 | -1.94307 | 3.789855 | 24.74593 | 0.0003 | 0.009298 |
| KIF14 | -2.20315 | 2.377975 | 24.51476 | 0.000304 | 0.009383 |
| LOC100414172 | -2.39711 | 1.5856 | 24.43473 | 0.000308 | 0.00945 |
| ASPHD2 | -1.86428 | 2.418451 | 24.41796 | 0.000309 | 0.00945 |
| ZDHHC13 | -1.70189 | 4.031259 | 24.42418 | 0.000309 | 0.00945 |
| CCR9 | -3.35389 | 4.291581 | 35.65537 | 0.000312 | 0.009478 |
| MYO1F | -2.42182 | 5.710988 | 31.14754 | 0.000316 | 0.009531 |
| IFNG | -5.00545 | 1.098584 | 26.50245 | 0.000317 | 0.009546 |
| PSME1 | -1.54828 | 3.744093 | 24.14664 | 0.000324 | 0.009684 |
| C1S | -1.44456 | 6.719375 | 24.15084 | 0.000324 | 0.009684 |
| KLHL6 | -1.4498 | 3.848843 | 24.11629 | 0.000326 | 0.009692 |
| LOC108589199 | -1.83404 | 4.619964 | 24.74542 | 0.000329 | 0.00975 |
| CALR | -1.40381 | 9.148537 | 24.06505 | 0.000329 | 0.00975 |
| WARS | -1.87662 | 6.687047 | 26.12278 | 0.000336 | 0.0099 |
| LOC100394277 | -2.60644 | 2.349636 | 23.9661 | 0.000344 | 0.010028 |
| LOC108591734 | -1.49494 | 4.00965 | 23.70861 | 0.00035 | 0.010153 |
| FAM105A | -1.43699 | 5.271296 | 23.67522 | 0.000352 | 0.010188 |
| SLC4A7 | -1.21632 | 6.757496 | 23.6727 | 0.000352 | 0.010188 |
| ASPM | -1.60864 | 3.874132 | 23.55204 | 0.00036 | 0.010333 |
| CDC25B | -1.40481 | 4.606058 | 23.51398 | 0.000362 | 0.010388 |
| LOC108590481 | -2.81433 | 2.538882 | 24.92429 | 0.000371 | 0.010536 |
| RCSD1 | -1.58029 | 5.829761 | 23.48318 | 0.000374 | 0.010604 |
| MCM4 | -1.77981 | 2.954362 | 23.27877 | 0.000378 | 0.010633 |
| FAM46C | -1.30533 | 5.700056 | 23.29387 | 0.000377 | 0.010633 |
| SH3KBP1 | -1.30528 | 4.465109 | 23.29605 | 0.000377 | 0.010633 |
| LSP1 | -1.71533 | 6.304622 | 24.27874 | 0.000379 | 0.010639 |
| LOC103793370 | -4.0263 | 0.782173 | 23.24624 | 0.00038 | 0.010665 |
| LOC100398523 | -1.54517 | 5.173816 | 23.16714 | 0.000388 | 0.010733 |
| ALS2CR12 | -2.15256 | 2.691222 | 23.06548 | 0.000393 | 0.010796 |
| GPR18 | -1.60079 | 3.867508 | 22.97976 | 0.000399 | 0.010894 |
| CYTH4 | -2.14598 | 4.132765 | 25.12536 | 0.000404 | 0.010994 |
| PLA2G2A | -1.85798 | 6.480795 | 25.10918 | 0.000404 | 0.010994 |
| EHF | -2.71838 | 5.218896 | 31.27566 | 0.000409 | 0.011063 |
| EVI2B | -1.60858 | 3.468068 | 22.8065 | 0.000412 | 0.011119 |
| LOC108592856 | -2.17933 | 1.883129 | 22.74884 | 0.000416 | 0.011196 |
| LOC100392020 | -1.81927 | 2.821475 | 22.73175 | 0.000417 | 0.01121 |
| EGR2 | -2.7973 | 1.256404 | 22.69321 | 0.00042 | 0.011216 |
| BARD1 | -1.80858 | 2.65854 | 22.70353 | 0.00042 | 0.011216 |
| LOC108588928 | -1.57877 | 3.366757 | 22.64137 | 0.000424 | 0.011243 |
| LOC100405506 | -2.07078 | 4.737084 | 25.4339 | 0.000433 | 0.011382 |
| ADCY7 | -1.40703 | 6.908597 | 22.40773 | 0.000443 | 0.011549 |
| FAM111B | -2.12276 | 3.982068 | 24.3347 | 0.000444 | 0.011572 |

| | | | | | |
|---|---|---|---|---|---|
| LOC108590106 | -1.90847 | 2.638288 | 22.31143 | 0.000451 | 0.011709 |
| ADRB1 | -1.91286 | 2.612309 | 22.29923 | 0.000452 | 0.011717 |
| LOC100402677 | -1.57465 | 3.10109 | 22.13301 | 0.000466 | 0.012024 |
| LOC103794320 | -2.73873 | 1.697405 | 22.10497 | 0.000469 | 0.012067 |
| PLCB2 | -2.19084 | 6.1748 | 27.28176 | 0.000472 | 0.012121 |
| C12H10orf90 | -2.85586 | 1.451384 | 21.99275 | 0.000479 | 0.012246 |
| RUNX3 | -2.11878 | 4.911107 | 25.7177 | 0.000481 | 0.01229 |
| MYADML2 | -1.85854 | 2.313744 | 21.89904 | 0.000487 | 0.012398 |
| CLEC12A | -2.51961 | 1.377421 | 21.82443 | 0.000494 | 0.01254 |
| FAM65B | -1.29977 | 4.614292 | 21.73202 | 0.000503 | 0.012727 |
| BATF2 | -2.77609 | 3.864013 | 27.68184 | 0.000505 | 0.012753 |
| LOC100411271 | -2.69951 | 3.072004 | 24.71794 | 0.000506 | 0.012753 |
| GPR183 | -1.59394 | 3.22098 | 21.68988 | 0.000507 | 0.012753 |
| LOC103796419 | -1.17272 | 5.911683 | 21.67819 | 0.000508 | 0.012758 |
| MR1 | -1.7927 | 2.578512 | 21.65571 | 0.00051 | 0.012784 |
| LOC108588630 | -1.36192 | 4.849728 | 21.62622 | 0.000513 | 0.012835 |
| LCP2 | -2.28099 | 5.845311 | 27.40051 | 0.000514 | 0.012857 |
| LOC100393894 | -1.9503 | 2.015954 | 21.57409 | 0.000518 | 0.012875 |
| ITPRIPL1 | -1.7523 | 2.595619 | 21.57052 | 0.000518 | 0.012875 |
| NRROS | -1.82041 | 2.938766 | 21.55515 | 0.00052 | 0.012893 |
| USP44 | -2.05283 | 1.741717 | 21.34759 | 0.000541 | 0.013261 |
| LOC100390149 | -2.24584 | 2.701649 | 21.22794 | 0.000567 | 0.01366 |
| SH2D2A | -2.16372 | 4.527803 | 24.65147 | 0.000566 | 0.01366 |
| ITGAL | -3.06836 | 6.737322 | 32.81478 | 0.000574 | 0.013765 |
| LOC108593454 | -1.4577 | 3.270368 | 21.02999 | 0.000574 | 0.013765 |
| LOC103792287 | -3.61398 | 0.967358 | 21.07296 | 0.000577 | 0.013769 |
| LAX1 | -3.30764 | 2.185103 | 23.92105 | 0.000584 | 0.013838 |
| SIPA1L1 | -1.18289 | 6.163044 | 20.92272 | 0.000587 | 0.013866 |
| TSPOAP1 | -1.60709 | 3.205243 | 20.83097 | 0.000597 | 0.013946 |
| VCAN | -1.42106 | 5.451494 | 20.84093 | 0.000596 | 0.013946 |
| VAMP1 | -1.41469 | 3.821659 | 20.83252 | 0.000597 | 0.013946 |
| LOC103791285 | -1.32024 | 3.745 | 20.72258 | 0.00061 | 0.014092 |
| CD52 | -2.32036 | 4.12263 | 24.53863 | 0.00062 | 0.014196 |
| PLBD1 | -1.3096 | 4.977142 | 20.66828 | 0.000616 | 0.014196 |
| SLC17A9 | -1.26673 | 4.146414 | 20.66053 | 0.000617 | 0.014196 |
| FAM49A | -1.15835 | 5.017598 | 20.65377 | 0.000618 | 0.014196 |
| LOC100394108 | -1.92367 | 4.863552 | 22.94697 | 0.000623 | 0.014239 |
| LOC100894856 | -2.48789 | 2.124745 | 20.68657 | 0.000626 | 0.014286 |
| HES7 | -1.63287 | 4.126533 | 20.62053 | 0.00063 | 0.014361 |
| NUP210 | -1.2587 | 6.600381 | 20.51788 | 0.000635 | 0.014421 |
| LOC100388240 | -2.0906 | 1.646669 | 20.48092 | 0.000639 | 0.014482 |
| LOC100405375 | -1.12119 | 6.08948 | 20.42691 | 0.000646 | 0.014569 |
| LOC100896470 | -3.03035 | 1.667729 | 20.85564 | 0.000648 | 0.014569 |
| LOC100404972 | -1.70168 | 3.099777 | 20.41802 | 0.000647 | 0.014569 |
| FNDC9 | -1.70286 | 2.819618 | 20.4013 | 0.000649 | 0.014584 |
| LOC103796527 | -4.60853 | 0.167521 | 20.37059 | 0.000653 | 0.014645 |
| MUC20 | -2.53521 | 1.139919 | 20.35321 | 0.000655 | 0.014645 |
| LOC103796425 | -1.66933 | 3.522088 | 20.35983 | 0.000655 | 0.014645 |

| | | | | | |
|---|---|---|---|---|---|
| PTPRCAP | -2.57292 | 6.080392 | 28.17119 | 0.000661 | 0.014711 |
| LOC108592270 | -1.85159 | 2.707011 | 20.29003 | 0.000664 | 0.014762 |
| PIM2 | -1.18369 | 5.000432 | 20.22186 | 0.000673 | 0.014895 |
| PPP1R18 | -1.31103 | 5.691997 | 20.18641 | 0.000677 | 0.014966 |
| KCNG3 | -1.89462 | 2.970637 | 20.14077 | 0.000684 | 0.015034 |
| SNX10 | -1.64191 | 3.631106 | 20.07327 | 0.000693 | 0.015202 |
| LOC108588781 | -1.66544 | 3.420404 | 20.0617 | 0.000694 | 0.01522 |
| CD44 | -1.15628 | 7.090843 | 20.03247 | 0.000698 | 0.015275 |
| LOC103794276 | -3.17803 | 0.451756 | 20.01314 | 0.000701 | 0.015299 |
| SLA | -1.62879 | 4.29494 | 20.12214 | 0.000702 | 0.015301 |
| LOC100412268 | -1.48311 | 3.689454 | 19.99453 | 0.000704 | 0.015322 |
| LOC100395429 | -1.44144 | 4.583447 | 19.97901 | 0.000706 | 0.015352 |
| LOC108588674 | -1.39237 | 3.630907 | 19.93478 | 0.000712 | 0.01546 |
| KNTC1 | -2.15743 | 2.106509 | 19.91822 | 0.000715 | 0.015474 |
| AGPAT5 | -1.39173 | 3.538434 | 19.89144 | 0.000718 | 0.015474 |
| MFNG | -1.59987 | 4.678478 | 20.14231 | 0.000722 | 0.01552 |
| LOC103789954 | -1.33236 | 9.597203 | 19.81936 | 0.000729 | 0.015607 |
| LOC103790968 | -1.25937 | 4.208313 | 19.82358 | 0.000728 | 0.015607 |
| LOC100411293 | -3.8527 | 0.681601 | 19.79446 | 0.000732 | 0.015651 |
| JAK3 | -2.30794 | 5.182468 | 25.07901 | 0.000739 | 0.015709 |
| CD300C | -2.09005 | 2.777129 | 19.83153 | 0.000741 | 0.015709 |
| ITGB2 | -2.19704 | 6.699464 | 24.87475 | 0.000744 | 0.015737 |
| PRR5L | -2.39746 | 2.12961 | 19.78411 | 0.000748 | 0.015773 |
| TPX2 | -1.63608 | 3.433657 | 19.67506 | 0.00075 | 0.015773 |
| PARP15 | -1.53558 | 3.984943 | 19.67346 | 0.00075 | 0.015773 |
| PDIA4 | -1.1955 | 7.285635 | 19.68051 | 0.000749 | 0.015773 |
| TDRD10 | -3.17323 | 0.459928 | 19.59862 | 0.000762 | 0.015865 |
| MAP3K1 | -1.07872 | 6.518029 | 19.62002 | 0.000759 | 0.015865 |
| LOC100397482 | -1.91085 | 2.939726 | 19.5152 | 0.000775 | 0.016099 |
| LOC100412682 | -1.28784 | 4.176772 | 19.5049 | 0.000776 | 0.016116 |
| TMSB4X | -1.25713 | 6.8301 | 19.43024 | 0.000788 | 0.01631 |
| MSMO1 | -1.40206 | 6.672039 | 19.48071 | 0.000795 | 0.016409 |
| LCN2 | -5.03314 | 5.244569 | 41.8457 | 0.000796 | 0.016425 |
| SEPTIN6 | -1.16252 | 5.336585 | 19.37412 | 0.000797 | 0.016427 |
| ICOSLG | -1.70936 | 3.56912 | 19.34955 | 0.000801 | 0.016482 |
| GPR55 | -2.83769 | 1.742417 | 19.68374 | 0.000805 | 0.016499 |
| PIP4K2A | -1.07251 | 5.592887 | 19.30475 | 0.000809 | 0.016521 |
| LOC100411501 | -2.51573 | 9.166619 | 24.87171 | 0.000813 | 0.016558 |
| ST6GALNAC1 | -1.90001 | 4.173833 | 20.76763 | 0.000814 | 0.016558 |
| LOC103790605 | -4.56891 | 0.173946 | 19.20004 | 0.000826 | 0.016635 |
| SLA2 | -2.82089 | 2.910923 | 22.70024 | 0.000822 | 0.016635 |
| GMFG | -1.6827 | 4.206217 | 19.55121 | 0.000826 | 0.016635 |
| ADCY3 | -1.53402 | 4.631967 | 19.30905 | 0.000826 | 0.016635 |
| GPSM3 | -1.38524 | 4.461877 | 19.21626 | 0.000823 | 0.016635 |
| TK1 | -1.15585 | 4.681683 | 19.21309 | 0.000824 | 0.016635 |
| IQGAP3 | -1.51739 | 2.974991 | 19.1784 | 0.00083 | 0.016662 |
| DUSP5 | -1.14557 | 4.781827 | 19.07236 | 0.000848 | 0.016834 |
| TRAFD1 | -1.19422 | 6.515773 | 19.06418 | 0.000849 | 0.016845 |

| | | | | | |
|---|---|---|---|---|---|
| NDC80 | -2.4246 | 1.43014 | 19.03016 | 0.000855 | 0.016911 |
| PPP1R1B | -1.35538 | 4.982191 | 19.0254 | 0.000856 | 0.016911 |
| LOC100406244 | -2.32733 | 7.908224 | 24.34543 | 0.00086 | 0.016963 |
| LOC100406154 | -3.5814 | 7.201562 | 33.17815 | 0.000862 | 0.016982 |
| SIT1 | -2.22301 | 1.620634 | 18.94389 | 0.000871 | 0.017128 |
| ARHGEF6 | -1.07698 | 5.965272 | 18.92776 | 0.000874 | 0.017151 |
| FAM26F | -2.02389 | 2.766586 | 18.98057 | 0.00088 | 0.017255 |
| RGS19 | -1.58399 | 3.946309 | 18.84859 | 0.000888 | 0.017347 |
| LOC108592819 | -2.47215 | 1.407726 | 18.83552 | 0.000891 | 0.017376 |
| CXHXorf21 | -2.48264 | 2.393214 | 19.50004 | 0.000898 | 0.017457 |
| MAP4K1 | -2.18537 | 5.645087 | 23.65511 | 0.000897 | 0.017457 |
| SEMA4A | -1.43014 | 4.615913 | 18.80395 | 0.000896 | 0.017457 |
| CDC45 | -2.42478 | 1.252386 | 18.75454 | 0.000906 | 0.017574 |
| RFLNB | -1.14409 | 4.594516 | 18.74567 | 0.000907 | 0.017574 |
| MYOF | -1.11099 | 7.126407 | 18.75104 | 0.000906 | 0.017574 |
| CASP8 | -1.4259 | 4.791302 | 18.73369 | 0.00091 | 0.017589 |
| TRAF1 | -1.54542 | 3.761826 | 18.69073 | 0.000918 | 0.017712 |
| CCNB1 | -2.5482 | 1.044028 | 18.52065 | 0.000951 | 0.018226 |
| LOC100396399 | -2.80718 | 2.295028 | 20.20156 | 0.000955 | 0.018253 |
| LOC100396380 | -1.82396 | 2.161572 | 18.45476 | 0.000964 | 0.018323 |
| LOC100411069 | -1.48445 | 3.235371 | 18.44475 | 0.000966 | 0.018323 |
| HCLS1 | -1.43517 | 5.313815 | 18.56341 | 0.000964 | 0.018323 |
| LOC100415684 | -1.35273 | 3.524156 | 18.4461 | 0.000966 | 0.018323 |
| CLEC6A | -2.97364 | 2.337127 | 21.00787 | 0.000968 | 0.018331 |
| LOC103791604 | -2.00322 | 2.796808 | 18.50284 | 0.000971 | 0.018357 |
| NBEAL2 | -1.36413 | 6.833188 | 18.49551 | 0.000973 | 0.018357 |
| RNASE6 | -1.65434 | 4.154538 | 18.69978 | 0.000978 | 0.018363 |
| KRT7 | -1.59961 | 3.609224 | 18.38319 | 0.000979 | 0.018363 |
| ZNF217 | -1.12833 | 6.352567 | 18.39579 | 0.000976 | 0.018363 |
| CCDC88C | -1.06424 | 6.851163 | 18.37429 | 0.000981 | 0.018363 |
| LOC100895417 | -3.82024 | 0.200318 | 18.34979 | 0.000986 | 0.018404 |
| LOC100405222 | -3.735 | 0.567804 | 18.26954 | 0.001002 | 0.018587 |
| MSN | -1.04261 | 7.264583 | 18.25372 | 0.001006 | 0.018622 |
| LOC100387316 | -1.45343 | 4.517227 | 18.22229 | 0.001014 | 0.018708 |
| LOC100398859 | -1.95226 | 7.023627 | 21.87172 | 0.001017 | 0.018746 |
| CDCA2 | -1.78708 | 2.043229 | 18.16901 | 0.001024 | 0.018798 |
| NLRC3 | -2.77199 | 5.762986 | 27.12185 | 0.001025 | 0.018806 |
| CAPN2 | -1.02542 | 6.648347 | 18.1535 | 0.001027 | 0.018825 |
| RBL1 | -1.42928 | 3.988533 | 18.13573 | 0.001031 | 0.018878 |
| XCL1 | -4.37438 | 1.689303 | 22.53831 | 0.001038 | 0.018986 |
| CNGB1 | -4.85134 | 1.363921 | 22.29897 | 0.00104 | 0.019011 |
| LOC108593911 | -1.91789 | 2.618825 | 18.0079 | 0.00106 | 0.019308 |
| CCK | -1.81844 | 3.165559 | 18.08264 | 0.001061 | 0.019312 |
| GPR171 | -3.44086 | 2.644663 | 22.78549 | 0.001087 | 0.01967 |
| BRIP1 | -2.22953 | 1.22533 | 17.89198 | 0.001086 | 0.01967 |
| APBB1IP | -1.26517 | 4.377125 | 17.87771 | 0.001089 | 0.019687 |
| SEMA4D | -1.01646 | 6.156465 | 17.86958 | 0.001091 | 0.019703 |
| FAM83E | -1.93769 | 4.373945 | 20.07567 | 0.001097 | 0.019759 |

| | | | | | |
|---|---|---|---|---|---|
| PLAGL2 | -1.04066 | 6.491958 | 17.80082 | 0.001108 | 0.019831 |
| SFN | -2.08204 | 6.330301 | 22.17292 | 0.001111 | 0.019856 |
| GGTA1P | -1.139 | 5.135611 | 17.78645 | 0.001111 | 0.019856 |
| DAPP1 | -2.40333 | 3.536669 | 21.06016 | 0.001118 | 0.019958 |
| LOC103793724 | -4.41944 | 0.041653 | 17.72589 | 0.001126 | 0.020061 |
| SPAG5 | -1.608 | 2.858669 | 17.71592 | 0.001128 | 0.020086 |
| LOC100393780 | -1.95174 | 1.894186 | 17.70246 | 0.001131 | 0.020092 |
| MSR1 | -1.76451 | 2.358562 | 17.71028 | 0.001129 | 0.020092 |
| LOC100402149 | -1.73743 | 2.302394 | 17.66569 | 0.00114 | 0.020136 |
| LOC100405488 | -3.18677 | 7.037548 | 28.95776 | 0.001149 | 0.020233 |
| CENPU | -1.67781 | 2.588643 | 17.61312 | 0.001153 | 0.020233 |
| CCNF | -1.49674 | 3.004038 | 17.6109 | 0.001154 | 0.020233 |
| STARD4 | -1.33593 | 5.674691 | 17.70084 | 0.001151 | 0.020233 |
| PTTG1 | -2.29443 | 2.154428 | 17.67669 | 0.00116 | 0.02031 |
| SIRPB1 | -1.65651 | 3.059769 | 17.51443 | 0.001178 | 0.020568 |
| GAB3 | -1.53664 | 3.575584 | 17.49351 | 0.001184 | 0.020624 |
| ZBTB1 | -1.10714 | 5.809261 | 17.48858 | 0.001185 | 0.020628 |
| ADAM22 | -1.11755 | 4.404991 | 17.44921 | 0.001195 | 0.02075 |
| CDCA3 | -2.47257 | 1.241704 | 17.41279 | 0.001204 | 0.020841 |
| LOC108589890 | -1.18195 | 3.873491 | 17.33881 | 0.001224 | 0.021086 |
| TDRD6 | -1.72294 | 2.599508 | 17.30822 | 0.001232 | 0.021193 |
| LRP8 | -2.03689 | 3.110569 | 17.94434 | 0.00124 | 0.02128 |
| PARP8 | -1.24445 | 5.264825 | 17.24247 | 0.00125 | 0.021384 |
| TRIM59 | -2.08832 | 1.55024 | 17.23462 | 0.001252 | 0.021403 |
| LOC100895074 | -1.82419 | 2.931337 | 17.21314 | 0.001258 | 0.021449 |
| ZWINT | -1.76395 | 2.194344 | 17.2042 | 0.001261 | 0.021471 |
| UCP2 | -1.44974 | 5.272794 | 17.56002 | 0.001262 | 0.02148 |
| XCR1 | -2.91303 | 0.538786 | 17.18061 | 0.001267 | 0.021545 |
| HTR6 | -2.16418 | 1.662106 | 17.15039 | 0.001276 | 0.021613 |
| LOC100409955 | -1.5013 | 2.978718 | 17.14443 | 0.001277 | 0.021613 |
| LOC103792275 | -3.0422 | 0.972478 | 17.12517 | 0.001283 | 0.021658 |
| LOC108594075 | -1.72934 | 3.452681 | 17.20577 | 0.001285 | 0.021659 |
| LOC108590123 | -2.67974 | 2.266949 | 18.51474 | 0.001291 | 0.021719 |
| GSG2 | -2.12655 | 1.912438 | 17.0517 | 0.001304 | 0.021765 |
| LOC100390753 | -2.03878 | 4.988864 | 20.71115 | 0.001309 | 0.021765 |
| TGM2 | -1.10711 | 7.358478 | 17.03914 | 0.001307 | 0.021765 |
| HMGB2 | -1.05551 | 5.196621 | 17.04091 | 0.001307 | 0.021765 |
| LOC100393273 | -2.41722 | 1.226052 | 16.99773 | 0.001319 | 0.02186 |
| MYH3 | -2.94533 | 1.271461 | 17.02546 | 0.001336 | 0.02205 |
| GINS4 | -2.39577 | 1.712972 | 16.88098 | 0.001354 | 0.022189 |
| LOC103789725 | -2 | 1.949046 | 16.86569 | 0.001358 | 0.022189 |
| ARHGAP8 | -1.92964 | 1.89898 | 16.86641 | 0.001358 | 0.022189 |
| IL27RA | -1.65573 | 3.444952 | 16.9198 | 0.001357 | 0.022189 |
| LOC100403284 | -1.53709 | 2.853973 | 16.86919 | 0.001357 | 0.022189 |
| LOC100403034 | -1.5072 | 5.122865 | 17.48657 | 0.001356 | 0.022189 |
| LOC100402594 | -1.65503 | 4.713171 | 17.97175 | 0.001376 | 0.022393 |
| CASP3 | -1.12754 | 4.888036 | 16.7966 | 0.001379 | 0.02242 |
| P2RX7 | -1.49586 | 3.466587 | 16.74222 | 0.001396 | 0.022618 |

| | | | | | |
|---|---|---|---|---|---|
| C11H11orf21 | -4.49767 | 1.519618 | 20.66817 | 0.001401 | 0.022652 |
| LOC108593028 | -1.64768 | 2.180476 | 16.71854 | 0.001404 | 0.022663 |
| SLC7A9 | -1.24356 | 7.780298 | 16.69299 | 0.001412 | 0.022717 |
| PSD4 | -1.2413 | 6.973407 | 16.7001 | 0.001409 | 0.022717 |
| FGR | -4.14261 | 4.234205 | 29.77132 | 0.001417 | 0.022731 |
| LOC103793673 | -2.50172 | 2.084378 | 17.28014 | 0.001414 | 0.022731 |
| ATP2C2 | -1.07367 | 5.651606 | 16.6759 | 0.001417 | 0.022731 |
| DMBT1 | -3.5622 | 8.210569 | 28.69738 | 0.001424 | 0.022809 |
| CD14 | -1.52589 | 5.362345 | 17.53131 | 0.001432 | 0.022888 |
| RGS3 | -1.12608 | 6.821812 | 16.61912 | 0.001435 | 0.022888 |
| LOC100409919 | -1.08093 | 5.377686 | 16.60394 | 0.00144 | 0.022929 |
| SMAP2 | -1.17415 | 6.5196 | 16.50141 | 0.001474 | 0.023293 |
| NPNT | -1.30176 | 4.785832 | 16.46263 | 0.001487 | 0.023402 |
| RNF207 | -1.086 | 4.444351 | 16.4324 | 0.001497 | 0.023468 |
| ATP1B3 | -1.29165 | 4.55089 | 16.42505 | 0.001499 | 0.023472 |
| LY6E | -1.26317 | 4.31632 | 16.37641 | 0.001516 | 0.023653 |
| LOC100409143 | -2.73507 | 1.904433 | 17.30107 | 0.001536 | 0.023784 |
| LOC103796571 | -1.48358 | 3.589733 | 16.3207 | 0.001535 | 0.023784 |
| TERF2IP | -0.99549 | 5.354849 | 16.31724 | 0.001536 | 0.023784 |
| ACAT2 | -1.30566 | 4.09711 | 16.30706 | 0.00154 | 0.023818 |
| LOC100897022 | -1.12594 | 8.53691 | 16.29914 | 0.001543 | 0.023842 |
| CORO1A | -2.36629 | 6.920521 | 22.37442 | 0.001552 | 0.02393 |
| TMEM206 | -1.4017 | 3.076099 | 16.26126 | 0.001556 | 0.023937 |
| LOC100409132 | -1.91258 | 2.02817 | 16.2522 | 0.001559 | 0.023948 |
| ESRP1 | -1.22478 | 5.089889 | 16.25178 | 0.001559 | 0.023948 |
| ABRACL | -1.31561 | 3.465956 | 16.24296 | 0.001562 | 0.023977 |
| CAJA-DPB1 | -2.39754 | 7.717014 | 22.19661 | 0.001564 | 0.02399 |
| KIF2C | -1.79313 | 1.987378 | 16.22835 | 0.001568 | 0.024 |
| LOC100413233 | -1.53379 | 3.144899 | 16.21468 | 0.001572 | 0.024001 |
| LIG1 | -1.04991 | 4.996555 | 16.21453 | 0.001572 | 0.024001 |
| B2M | -1.98797 | 10.27793 | 18.44117 | 0.001586 | 0.024091 |
| FOXM1 | -1.85872 | 2.275326 | 16.1689 | 0.001589 | 0.024101 |
| ITGAM | -1.3209 | 6.0395 | 16.26341 | 0.001592 | 0.024105 |
| ARRB2 | -1.24035 | 5.270364 | 16.158 | 0.001593 | 0.024105 |
| LOC108590528 | -3.30709 | 2.84352 | 21.2408 | 0.001617 | 0.024421 |
| ACOT7 | -1.91049 | 4.435314 | 18.46905 | 0.001623 | 0.024483 |
| NFIL3 | -1.27875 | 4.489788 | 16.05064 | 0.001632 | 0.024589 |
| E2F8 | -2.52441 | 1.314747 | 16.02568 | 0.001642 | 0.024677 |
| C3 | -1.1471 | 8.950604 | 15.99917 | 0.001652 | 0.024766 |
| CEP55 | -2.81694 | 1.194486 | 15.99264 | 0.001676 | 0.025049 |
| OSTC | -1.52684 | 3.271304 | 15.92972 | 0.001678 | 0.025057 |
| NTNG2 | -1.30691 | 4.07119 | 15.91216 | 0.001685 | 0.025074 |
| PHF19 | -1.14074 | 5.039248 | 15.91351 | 0.001685 | 0.025074 |
| LOC100387926 | -2.15613 | 2.54773 | 16.32531 | 0.001694 | 0.025114 |
| LOC103790084 | -1.59621 | 2.878158 | 15.83601 | 0.001715 | 0.025325 |
| LOC103794136 | -1.81281 | 2.771654 | 15.80664 | 0.001727 | 0.025459 |
| DUOXA2 | -2.48012 | 3.215706 | 18.81334 | 0.001731 | 0.025511 |
| LOC100387406 | -1.30002 | 3.331649 | 15.78 | 0.001737 | 0.025558 |

| | | | | | |
|---|---|---|---|---|---|
| TIFA | -1.33456 | 3.413778 | 15.71687 | 0.001763 | 0.025865 |
| MAPK12 | -1.34671 | 3.440321 | 15.66968 | 0.001782 | 0.026082 |
| LOC108594078 | -3.16793 | 2.305694 | 18.93868 | 0.001801 | 0.026263 |
| LOC103788772 | -4.37088 | 3.01246 | 24.64155 | 0.001827 | 0.02651 |
| LOC100402818 | -2.99269 | 3.830028 | 22.17016 | 0.001827 | 0.02651 |
| VOPP1 | -1.29448 | 4.220573 | 15.55818 | 0.001829 | 0.02651 |
| ORAI1 | -1.17698 | 3.725216 | 15.54504 | 0.001835 | 0.026537 |
| LOC100896919 | -2.27839 | 1.664824 | 15.45919 | 0.001872 | 0.026911 |
| LOC100405150 | -3.75776 | 7.288767 | 28.53389 | 0.001906 | 0.027215 |
| CDC42SE2 | -1.25388 | 6.521563 | 15.44851 | 0.001904 | 0.027215 |
| LOC100386904 | -1.22245 | 3.427404 | 15.38499 | 0.001905 | 0.027215 |
| TPM4 | -1.20066 | 6.895359 | 15.38861 | 0.001903 | 0.027215 |
| SLC16A6 | -1.94774 | 2.261061 | 15.36751 | 0.001913 | 0.02723 |
| GLIPR1 | -1.31516 | 3.585717 | 15.36222 | 0.001915 | 0.02723 |
| CRACR2A | -1.14697 | 5.060541 | 15.37086 | 0.001911 | 0.02723 |
| ORC1 | -1.44939 | 2.550966 | 15.34415 | 0.001923 | 0.027306 |
| IL12RB2 | -3.94411 | 4.88578 | 28.32569 | 0.001941 | 0.027454 |
| IQGAP1 | -1.1003 | 8.456749 | 15.308 | 0.00194 | 0.027454 |
| LOC103794953 | -2.48083 | 5.568344 | 21.81426 | 0.001944 | 0.027477 |
| IL10RA | -1.35794 | 5.792997 | 15.63007 | 0.001947 | 0.0275 |
| LOC103790978 | -1.75793 | 1.721989 | 15.27414 | 0.001955 | 0.0276 |
| LOC100406082 | -1.87647 | 2.689938 | 15.29433 | 0.001974 | 0.027747 |
| CD68 | -1.0285 | 6.411073 | 15.22505 | 0.001978 | 0.027781 |
| LOC100403855 | -1.09869 | 4.941517 | 15.22052 | 0.00198 | 0.027791 |
| SIPA1 | -1.31526 | 6.117659 | 15.42934 | 0.001988 | 0.027879 |
| ADAMTS14 | -1.6848 | 5.302868 | 17.21243 | 0.001996 | 0.027976 |
| LOC108590093 | -1.39206 | 3.058023 | 15.16681 | 0.002005 | 0.028045 |
| HMGA1 | -1.12092 | 6.432597 | 15.11982 | 0.002028 | 0.028238 |
| CD86 | -2.44268 | 4.268252 | 19.79207 | 0.002054 | 0.028513 |
| RFC4 | -1.25708 | 3.485811 | 15.04065 | 0.002066 | 0.028643 |
| GLS | -0.99495 | 7.876759 | 15.01278 | 0.00208 | 0.028802 |
| RRP1B | -1.09578 | 5.71875 | 15.00873 | 0.002082 | 0.028809 |
| ANKRD33B | -1.18267 | 4.369894 | 15.00421 | 0.002084 | 0.028812 |
| LOC103792274 | -2.49782 | 0.6174 | 14.98715 | 0.002093 | 0.028876 |
| GALNT3 | -1.46902 | 4.957239 | 15.63393 | 0.002115 | 0.029125 |
| KIF20B | -1.50507 | 3.383874 | 14.93054 | 0.002121 | 0.029131 |
| PTAFR | -1.474 | 2.456257 | 14.91561 | 0.002129 | 0.029148 |
| DUSP4 | -1.67926 | 4.323169 | 16.02416 | 0.002158 | 0.029525 |
| ARHGAP4 | -1.15552 | 6.243958 | 14.85267 | 0.002161 | 0.029549 |
| ANP32E | -1.18739 | 4.890792 | 14.77339 | 0.002203 | 0.029991 |
| TNFRSF17 | -2.49158 | 1.385388 | 14.74534 | 0.002218 | 0.030131 |
| BOLA3 | -1.70532 | 2.01947 | 14.73581 | 0.002223 | 0.030138 |
| PDIA3 | -1.08097 | 7.838694 | 14.72712 | 0.002227 | 0.030159 |
| LOC100399497 | -1.10844 | 6.891685 | 14.71766 | 0.002232 | 0.030165 |
| CXCL6 | -2.11543 | 3.051456 | 16.01937 | 0.002244 | 0.030259 |
| C1R | -1.22915 | 7.045007 | 14.74815 | 0.002247 | 0.030273 |
| LOC108593572 | -1.98442 | 2.215187 | 14.69511 | 0.002262 | 0.030444 |
| BLM | -1.79955 | 2.249302 | 14.65545 | 0.002266 | 0.030474 |

| | | | | | |
|---|---|---|---|---|---|
| PCSK9 | -1.55983 | 4.344033 | 15.31655 | 0.002289 | 0.030739 |
| FHL3 | -1.33762 | 4.255495 | 14.60079 | 0.002296 | 0.030797 |
| RIPK3 | -1.61696 | 2.600623 | 14.57681 | 0.00231 | 0.030953 |
| RPA2 | -1.48285 | 2.369366 | 14.56952 | 0.002314 | 0.030987 |
| RPIA | -1.11275 | 3.862315 | 14.56113 | 0.002319 | 0.031007 |
| GARS | -1.00262 | 5.41899 | 14.56247 | 0.002318 | 0.031007 |
| CRELD2 | -0.92931 | 5.749309 | 14.5569 | 0.002321 | 0.031018 |
| LOC108590345 | -1.80061 | 1.761414 | 14.53174 | 0.002335 | 0.031165 |
| TRIM5 | -1.34832 | 5.264297 | 14.82035 | 0.002351 | 0.031278 |
| LOC100415230 | -1.5005 | 2.251657 | 14.4913 | 0.002358 | 0.031324 |
| DUOX1 | -1.05 | 4.960025 | 14.4914 | 0.002358 | 0.031324 |
| SESN3 | -1.01501 | 5.712378 | 14.46789 | 0.002372 | 0.031481 |
| CENPM | -1.70096 | 1.83235 | 14.45432 | 0.00238 | 0.031521 |
| THOP1 | -1.05501 | 4.56774 | 14.45609 | 0.002379 | 0.031521 |
| PHGDH | -1.61145 | 2.640141 | 14.4422 | 0.002387 | 0.03155 |
| CELSR3 | -1.32215 | 4.706672 | 14.50937 | 0.002386 | 0.03155 |
| HIVEP3 | -1.18952 | 3.718262 | 14.44385 | 0.002386 | 0.03155 |
| SFMBT2 | -1.15592 | 4.283948 | 14.41162 | 0.002404 | 0.031765 |
| LOC100385039 | -1.81547 | 2.26175 | 14.39443 | 0.002415 | 0.031813 |
| RARRES1 | -1.68355 | 5.87329 | 16.66006 | 0.002421 | 0.031872 |
| HOPX | -1.37522 | 2.66766 | 14.35308 | 0.002439 | 0.032049 |
| TLR3 | -1.25463 | 4.31534 | 14.35447 | 0.002438 | 0.032049 |
| NPRL2 | -1.24764 | 4.378151 | 14.33649 | 0.002449 | 0.032115 |
| MARCH1 | -1.4 | 3.267785 | 14.31762 | 0.00246 | 0.032177 |
| FCHO1 | -1.30473 | 4.574204 | 14.35456 | 0.002471 | 0.03225 |
| SMC4 | -1.02699 | 5.481696 | 14.29263 | 0.002475 | 0.032267 |
| RPL36A | -2.08329 | 1.435065 | 14.26849 | 0.00249 | 0.032382 |
| LOC100896775 | -1.34339 | 3.106518 | 14.26705 | 0.002491 | 0.032382 |
| GALM | -1.28035 | 4.551338 | 14.26976 | 0.002493 | 0.032382 |
| FANCD2 | -1.59104 | 2.466557 | 14.25234 | 0.0025 | 0.032441 |
| LOC103789502 | -1.20719 | 8.64529 | 14.25176 | 0.0025 | 0.032441 |
| TMEM176B | -1.12412 | 6.79962 | 14.2311 | 0.002513 | 0.032559 |
| NPC2 | -1.06337 | 5.374759 | 14.21947 | 0.00252 | 0.032613 |
| CPS1 | -1.11882 | 8.907544 | 14.20775 | 0.002528 | 0.032678 |
| TOX | -0.98294 | 5.084351 | 14.20607 | 0.002529 | 0.032678 |
| LOC100410098 | -2.00188 | 1.734329 | 14.1919 | 0.002538 | 0.032706 |
| LOC100411032 | -1.59593 | 2.481634 | 14.18107 | 0.002544 | 0.032749 |
| ALOX12 | -0.97888 | 5.040969 | 14.17359 | 0.002549 | 0.032788 |
| FADS2 | -1.08277 | 5.25334 | 14.14455 | 0.002567 | 0.032937 |
| MYC | -0.98963 | 4.935731 | 14.14103 | 0.00257 | 0.032944 |
| LOC103787275 | -1.58375 | 2.83541 | 14.10439 | 0.002593 | 0.033135 |
| LOC100395214 | -5.88547 | 2.290449 | 23.28418 | 0.002604 | 0.033165 |
| LOC100395313 | -1.58118 | 3.063144 | 14.09365 | 0.0026 | 0.033165 |
| ZDHHC23 | -0.99721 | 5.135633 | 14.09152 | 0.002601 | 0.033165 |
| LOC108588472 | -1.7547 | 1.674272 | 14.07514 | 0.002612 | 0.033224 |
| LOC100398815 | -1.69083 | 2.261522 | 14.06257 | 0.00262 | 0.033284 |
| SLAMF7 | -3.36323 | 4.622245 | 23.18976 | 0.002641 | 0.033378 |
| LOC103791358 | -2.63465 | 0.862565 | 14.03776 | 0.002636 | 0.033378 |

| | | | | | |
|---|---|---|---|---|---|
| GPR132 | -2.11295 | 3.130278 | 15.60736 | 0.002643 | 0.033378 |
| TACSTD2 | -1.67725 | 2.205528 | 14.03666 | 0.002637 | 0.033378 |
| DMRTA2 | -3.32335 | 0.573891 | 14.03857 | 0.002655 | 0.033457 |
| SSBP4 | -1.22773 | 5.169978 | 14.05754 | 0.002657 | 0.033457 |
| MMS22L | -1.13748 | 4.540863 | 14.00246 | 0.002659 | 0.033457 |
| HYOU1 | -0.94544 | 7.63188 | 14.00475 | 0.002658 | 0.033457 |
| TCIRG1 | -0.9968 | 6.013884 | 13.98422 | 0.002671 | 0.033566 |
| LOC103792212 | -1.19083 | 3.775691 | 13.94489 | 0.002698 | 0.033809 |
| LOC108589454 | -2.11628 | 1.463703 | 13.93857 | 0.002702 | 0.03384 |
| DUSP2 | -2.9206 | 3.014997 | 18.02919 | 0.002704 | 0.033843 |
| NKG7 | -4.46129 | 3.571246 | 24.70367 | 0.002709 | 0.033865 |
| CPVL | -1.69946 | 3.48498 | 14.43057 | 0.002709 | 0.033865 |
| CD163 | -1.43842 | 4.956496 | 14.63203 | 0.002712 | 0.033876 |
| MAD1L1 | -1.1226 | 5.308175 | 13.91787 | 0.002716 | 0.033905 |
| HMCN1 | -1.0103 | 6.68089 | 13.91547 | 0.002717 | 0.033905 |
| DMXL2 | -1.29467 | 7.327064 | 14.15215 | 0.002751 | 0.034223 |
| HSPD1 | -1.04976 | 5.122493 | 13.86533 | 0.002752 | 0.034223 |
| CCR8 | -3.0551 | 0.937239 | 14.02655 | 0.002753 | 0.034223 |
| GATA3 | -2.11343 | 3.665919 | 16.29163 | 0.002765 | 0.034326 |
| TGFB1 | -1.12756 | 5.411716 | 13.83796 | 0.00277 | 0.034326 |
| ASF1B | -2.03373 | 1.030378 | 13.79487 | 0.0028 | 0.034611 |
| TRPC5 | -1.22489 | 4.399334 | 13.76915 | 0.002819 | 0.034768 |
| CHSY1 | -1.00689 | 4.693608 | 13.7597 | 0.002825 | 0.034807 |
| ETV3L | -3.87975 | 0.646721 | 14.39598 | 0.002831 | 0.034851 |
| LOC100398205 | -1.28176 | 4.086569 | 13.7489 | 0.002833 | 0.034857 |
| LOC100390292 | -0.96107 | 4.726659 | 13.73836 | 0.00284 | 0.034906 |
| RDH11 | -1.08689 | 4.614298 | 13.71344 | 0.002858 | 0.035103 |
| NOP56 | -0.91128 | 6.224573 | 13.71095 | 0.00286 | 0.035103 |
| FEN1 | -1.51735 | 2.183612 | 13.66727 | 0.002892 | 0.035219 |
| EML4 | -1.08058 | 6.028523 | 13.68035 | 0.002882 | 0.035219 |
| PRKCB | -0.98332 | 5.270081 | 13.66552 | 0.002893 | 0.035219 |
| SNX20 | -2.03039 | 1.896503 | 13.65654 | 0.002899 | 0.035232 |
| NUAK1 | -1.24369 | 3.976328 | 13.66002 | 0.002897 | 0.035232 |
| LOC100401712 | -1.51479 | 2.81776 | 13.6458 | 0.002907 | 0.035284 |
| WNT1 | -2.29085 | 0.8962 | 13.6236 | 0.002924 | 0.035416 |
| TNFAIP8L2 | -1.56222 | 2.593757 | 13.56148 | 0.00297 | 0.035805 |
| TANK | -1.34689 | 3.485278 | 13.51392 | 0.003006 | 0.036121 |
| CTSW | -3.86622 | 5.281506 | 25.46787 | 0.00301 | 0.036152 |
| LOC103789118 | -1.22689 | 3.133566 | 13.50562 | 0.003012 | 0.036152 |
| CD47 | -1.14521 | 6.195394 | 13.5035 | 0.003022 | 0.036228 |
| LOC100395561 | -0.99709 | 4.809687 | 13.49283 | 0.003022 | 0.036228 |
| LOC100896309 | -2.42659 | 0.75142 | 13.47129 | 0.003038 | 0.036346 |
| MLEC | -0.94464 | 8.153786 | 13.46729 | 0.003041 | 0.036346 |
| LOC100411120 | -1.08063 | 6.325862 | 13.44715 | 0.003057 | 0.03638 |
| LOC100391263 | -1.30757 | 2.816229 | 13.43486 | 0.003067 | 0.036427 |
| LOC103793827 | -3.80231 | 0.203224 | 13.39337 | 0.003099 | 0.036769 |
| MOB3A | -1.09895 | 5.389128 | 13.38852 | 0.003103 | 0.036783 |
| HSP90AA1 | -0.9798 | 7.009219 | 13.38715 | 0.003104 | 0.036783 |

| | | | | | |
|---|---|---|---|---|---|
| KIF20A | -1.51881 | 2.119978 | 13.38213 | 0.003108 | 0.036807 |
| CDKN2A | -1.81408 | 2.217132 | 13.37544 | 0.003113 | 0.036848 |
| C14H2orf16 | -2.12586 | 1.372695 | 13.3606 | 0.003125 | 0.036934 |
| ARHGAP45 | -1.78324 | 6.728299 | 16.0513 | 0.003123 | 0.036934 |
| LOC100402421 | -1.5677 | 2.468966 | 13.35918 | 0.003126 | 0.036934 |
| PDX1 | -2.38874 | 3.339027 | 16.13674 | 0.003134 | 0.036946 |
| IFI30 | -1.01308 | 5.956904 | 13.34841 | 0.003135 | 0.036946 |
| LOC100402737 | -1.51766 | 2.653647 | 13.32924 | 0.00315 | 0.037016 |
| LOC100405988 | -4.95636 | 1.50065 | 17.97765 | 0.003161 | 0.037096 |
| POLR3GL | -1.03244 | 3.864289 | 13.30032 | 0.003174 | 0.037223 |
| SPC24 | -2.28623 | 1.666915 | 13.3301 | 0.003199 | 0.037456 |
| BUB1B | -1.27861 | 3.403895 | 13.26321 | 0.003204 | 0.037488 |
| PDIA6 | -1.05434 | 6.263227 | 13.24963 | 0.003215 | 0.037574 |
| RTKN2 | -3.79162 | 0.196664 | 13.24709 | 0.003217 | 0.037576 |
| LOC100390453 | -1.65762 | 1.784766 | 13.23862 | 0.003224 | 0.037635 |
| TRIM21 | -1.13521 | 4.74679 | 13.22265 | 0.003237 | 0.037767 |
| ADAM17 | -0.93564 | 5.494384 | 13.21497 | 0.003244 | 0.037819 |
| TNFRSF4 | -1.57804 | 2.685237 | 13.19417 | 0.003261 | 0.037895 |
| LOC100392572 | -1.03052 | 8.384286 | 13.19323 | 0.003262 | 0.037895 |
| HMGN1 | -0.99879 | 5.002239 | 13.19839 | 0.003258 | 0.037895 |
| LOC100394285 | -1.78875 | 1.587863 | 13.16424 | 0.003286 | 0.038133 |
| ACAP1 | -1.23417 | 6.193921 | 13.34341 | 0.003323 | 0.038471 |
| AMZ1 | -1.50506 | 2.343782 | 13.10528 | 0.003337 | 0.038533 |
| SQLE | -1.69323 | 6.038347 | 15.46192 | 0.003344 | 0.038564 |
| TDP1 | -1.15098 | 3.287098 | 13.08842 | 0.003351 | 0.038564 |
| LOC108593553 | -4.04183 | 2.105954 | 17.86735 | 0.003401 | 0.038956 |
| KIFC1 | -1.14726 | 3.275078 | 12.97324 | 0.003452 | 0.039406 |
| HPS3 | -1.0117 | 4.323935 | 12.934 | 0.003488 | 0.039707 |
| NUS1 | -0.96205 | 4.883143 | 12.93794 | 0.003484 | 0.039707 |
| LOC100394437 | -1.17775 | 5.364179 | 12.9653 | 0.003502 | 0.039789 |
| LOC108590918 | -1.3325 | 4.083223 | 12.94648 | 0.00352 | 0.039915 |
| H2AFX | -1.08635 | 4.63337 | 12.8748 | 0.003542 | 0.040053 |
| FLRT3 | -1.04593 | 5.936751 | 12.87743 | 0.003539 | 0.040053 |
| RFTN1 | -0.88955 | 5.335594 | 12.87703 | 0.00354 | 0.040053 |
| PCNA | -1.15575 | 3.452584 | 12.8452 | 0.003569 | 0.040232 |
| LYN | -1.14892 | 6.149796 | 12.88997 | 0.00357 | 0.040232 |
| APOC1 | -2.8591 | 2.21031 | 15.16526 | 0.003581 | 0.040244 |
| FLT3 | -2.76293 | 1.003502 | 12.8883 | 0.003579 | 0.040244 |
| LOC100400495 | -2.0925 | 1.986649 | 12.97561 | 0.003581 | 0.040244 |
| LOC100406240 | -1.68766 | 7.081163 | 15.17031 | 0.003577 | 0.040244 |
| PLEKHO2 | -1.0241 | 5.998519 | 12.83789 | 0.003576 | 0.040244 |
| BAIAP2L1 | -1.10437 | 6.756376 | 12.82625 | 0.003587 | 0.040283 |
| LOC100397968 | -1.98942 | 4.658518 | 15.72384 | 0.003635 | 0.040658 |
| FIGNL1 | -1.17538 | 3.161107 | 12.77054 | 0.00364 | 0.040687 |
| LOC103793005 | -3.83391 | 1.42214 | 15.17965 | 0.003688 | 0.041092 |
| TNFAIP3 | -1.20311 | 6.306942 | 12.89408 | 0.003688 | 0.041092 |
| DUSP22 | -0.95878 | 4.503469 | 12.70707 | 0.003701 | 0.041204 |
| LOC103794333 | -1.96245 | 1.755179 | 12.70488 | 0.003703 | 0.041204 |

| | | | | | |
|---|---|---|---|---|---|
| LOC100415238 | -2.30305 | 1.076696 | 12.69335 | 0.003714 | 0.041238 |
| LRFN2 | -1.49456 | 2.318123 | 12.69484 | 0.003712 | 0.041238 |
| LRP4 | -0.97938 | 5.659441 | 12.69305 | 0.003714 | 0.041238 |
| KIAA0101 | -2.40885 | 0.681693 | 12.67817 | 0.003729 | 0.041376 |
| LOC108591194 | -1.1761 | 3.068775 | 12.6666 | 0.00374 | 0.041478 |
| LOC100399132 | -1.94411 | 1.476057 | 12.65311 | 0.003753 | 0.041508 |
| PDE4B | -1.23306 | 5.119919 | 12.80814 | 0.003753 | 0.041508 |
| LOC108594037 | -1.14103 | 3.265715 | 12.6544 | 0.003752 | 0.041508 |
| PTPN22 | -1.94618 | 4.684153 | 15.38197 | 0.003784 | 0.041774 |
| RUNX2 | -1.24983 | 4.373298 | 12.66228 | 0.003788 | 0.041792 |
| RASSF2 | -1.03404 | 4.693526 | 12.6065 | 0.0038 | 0.041855 |
| LOC100409719 | -2.06458 | 1.682422 | 12.5975 | 0.003809 | 0.041923 |
| LOC100415472 | -0.9885 | 4.967646 | 12.59227 | 0.003814 | 0.041941 |
| PREX1 | -1.92853 | 6.365709 | 15.99636 | 0.003816 | 0.041941 |
| DENND1C | -1.13189 | 6.233165 | 12.62477 | 0.003825 | 0.042015 |
| LOC108594024 | -2.65823 | 0.545191 | 12.57611 | 0.00383 | 0.042026 |
| C3H4orf51 | -2.45223 | 0.562618 | 12.57794 | 0.003828 | 0.042026 |
| IL5RA | -2.00007 | 1.284838 | 12.55896 | 0.003848 | 0.042193 |
| LOC108590966 | -1.16136 | 3.968789 | 12.53863 | 0.003868 | 0.042349 |
| RASSF1 | -1.05452 | 6.033302 | 12.49797 | 0.00391 | 0.042637 |
| C7H1orf216 | -1.22137 | 3.302231 | 12.4842 | 0.003924 | 0.0427 |
| TBXAS1 | -1.08656 | 3.988812 | 12.45323 | 0.003957 | 0.042932 |
| CEP128 | -1.01864 | 3.893021 | 12.44896 | 0.003961 | 0.042933 |
| MTHFD2 | -1.21964 | 5.234734 | 12.60473 | 0.003973 | 0.042994 |
| LOC100895081 | -1.13882 | 4.428184 | 12.43743 | 0.003973 | 0.042994 |
| HAAO | -1.81612 | 3.256203 | 13.28864 | 0.003977 | 0.042997 |
| HELB | -0.88498 | 5.443562 | 12.43299 | 0.003978 | 0.042997 |
| SLC9B2 | -1.33616 | 2.756541 | 12.42354 | 0.003988 | 0.043051 |
| CXCR6 | -0.9457 | 5.552731 | 12.42197 | 0.00399 | 0.043051 |
| TLE1 | -0.99727 | 5.053035 | 12.40272 | 0.00401 | 0.043224 |
| LOC108589988 | -2.61428 | 3.174464 | 15.7018 | 0.004018 | 0.043258 |
| NUCB2 | -1.25694 | 4.038062 | 12.37047 | 0.004045 | 0.043453 |
| HSF2BP | -1.53421 | 2.312871 | 12.34988 | 0.004067 | 0.043603 |
| MAN2B2 | -1.30888 | 5.336108 | 12.88429 | 0.004076 | 0.043672 |
| GNLY | -4.25862 | 5.624375 | 24.87496 | 0.00409 | 0.04377 |
| PRC1 | -0.87381 | 5.188091 | 12.31761 | 0.004102 | 0.043853 |
| VARS | -0.8815 | 5.902921 | 12.3106 | 0.00411 | 0.043874 |
| POLE | -0.98702 | 4.270431 | 12.2771 | 0.004147 | 0.044113 |
| PLCG2 | -0.87502 | 6.486689 | 12.26518 | 0.00416 | 0.044206 |
| STIP1 | -0.89127 | 6.288057 | 12.22471 | 0.004205 | 0.044542 |
| NCAPH | -1.39588 | 2.466151 | 12.19185 | 0.004243 | 0.044887 |
| ALDH1L2 | -1.07128 | 4.611606 | 12.17878 | 0.004258 | 0.044997 |
| RASAL3 | -2.31889 | 4.51802 | 16.40851 | 0.004275 | 0.045107 |
| NDUFAF1 | -1.01648 | 4.112513 | 12.16447 | 0.004274 | 0.045107 |
| NPL | -1.55073 | 3.648251 | 12.60985 | 0.004279 | 0.045128 |
| LOC100895618 | -2.21936 | 0.711372 | 12.13967 | 0.004303 | 0.045325 |
| DSC3 | -1.42623 | 3.01427 | 12.13755 | 0.004305 | 0.045327 |
| LOC100392865 | -1.1988 | 3.206498 | 12.1239 | 0.004321 | 0.04542 |

| | | | | | |
|---|---|---|---|---|---|
| RFX5 | -1.13275 | 6.067901 | 12.17477 | 0.004319 | 0.04542 |
| BTK | -1.20958 | 3.217345 | 12.12168 | 0.004323 | 0.045423 |
| SLFN11 | -1.81104 | 2.199988 | 12.12793 | 0.004348 | 0.045556 |
| CD24 | -1.46493 | 3.323169 | 12.14391 | 0.004346 | 0.045556 |
| TOR4A | -1.44043 | 3.082665 | 12.09885 | 0.00435 | 0.045556 |
| SSR1 | -0.99354 | 6.095733 | 12.08771 | 0.004363 | 0.045627 |
| LOC103789338 | -1.34123 | 3.477439 | 12.05849 | 0.004398 | 0.045883 |
| FXYD5 | -1.42094 | 3.450944 | 12.08278 | 0.004417 | 0.045964 |
| STC1 | -1.63444 | 2.47096 | 12.03856 | 0.004421 | 0.045983 |
| MANF | -1.01462 | 4.605598 | 12.01006 | 0.004456 | 0.046265 |
| CALHM2 | -1.11847 | 3.890107 | 12.00734 | 0.004459 | 0.046275 |
| TNFRSF18 | -2.36748 | 3.108305 | 14.31631 | 0.004488 | 0.046498 |
| CCL5 | -4.09487 | 5.985174 | 23.78262 | 0.004491 | 0.046513 |
| AIF1 | -1.18403 | 3.758569 | 11.96743 | 0.004507 | 0.046605 |
| WDR87 | -2.15128 | 1.77536 | 12.09988 | 0.004517 | 0.046656 |
| ALOX5AP | -2.83757 | 3.656102 | 16.7196 | 0.004552 | 0.046894 |
| RUNX1 | -1.3803 | 5.27401 | 12.79264 | 0.004558 | 0.046926 |
| TFCP2L1 | -1.79701 | 1.219678 | 11.89609 | 0.004596 | 0.047119 |
| LOC100399174 | -0.91164 | 4.780468 | 11.87432 | 0.004623 | 0.047348 |
| FBL | -0.95563 | 4.938852 | 11.85859 | 0.004643 | 0.047501 |
| LOC100392893 | -1.49926 | 3.01901 | 11.88072 | 0.004648 | 0.047531 |
| P2RY14 | -1.30951 | 4.087762 | 11.93108 | 0.004664 | 0.04764 |
| LOC100409880 | -0.87452 | 5.65622 | 11.82874 | 0.004681 | 0.047765 |
| LOC103793895 | -1.65564 | 2.222282 | 11.80063 | 0.004717 | 0.047983 |
| LOC100397605 | -2.42612 | 3.172165 | 14.42239 | 0.004727 | 0.048014 |
| CD274 | -2.09006 | 1.534757 | 11.79524 | 0.004724 | 0.048014 |
| PSMB2 | -1.08299 | 4.955026 | 11.79394 | 0.004725 | 0.048014 |
| ZNF267 | -0.93233 | 4.463038 | 11.77438 | 0.004751 | 0.048156 |
| EPHB2 | -1.71033 | 4.766366 | 13.76349 | 0.004761 | 0.048203 |
| LOC100896238 | -1.21968 | 2.937766 | 11.75966 | 0.00477 | 0.048247 |
| PRKX | -1.21319 | 3.1161 | 11.73944 | 0.004796 | 0.048416 |
| IL12RB1 | -2.79276 | 3.204597 | 15.61444 | 0.00481 | 0.048496 |
| LOC103794171 | -1.41795 | 2.930785 | 11.72281 | 0.004818 | 0.048536 |
| AGTRAP | -1.04203 | 3.680224 | 11.71953 | 0.004823 | 0.048554 |
| LOC100413381 | -1.45031 | 2.086047 | 11.71666 | 0.004826 | 0.048568 |
| PXYLP1 | -0.8783 | 5.887614 | 11.67679 | 0.00488 | 0.048926 |
| LOC108588401 | -4.72055 | 5.540643 | 24.67428 | 0.004896 | 0.049016 |
| CD180 | -1.54327 | 3.663793 | 12.15082 | 0.00494 | 0.049276 |
| NCAPG | -1.46902 | 2.588246 | 11.62119 | 0.004955 | 0.049351 |
| TESPA1 | -2.43343 | 2.264067 | 13.05199 | 0.00497 | 0.049475 |
| EVI2A | -1.58181 | 3.175957 | 11.81369 | 0.004978 | 0.049537 |
| ANK1 | -1.06497 | 3.491742 | 11.58755 | 0.005001 | 0.04971 |
| LOC103792196 | -1.22325 | 3.814308 | 11.5854 | 0.005004 | 0.049714 |
| CD2 | -2.01798 | 3.975631 | 13.92281 | 0.005031 | 0.049882 |
| LYSMD2 | -1.9006 | 1.433849 | 11.56284 | 0.005035 | 0.049882 |
| RASA3 | -1.1276 | 5.850856 | 11.64291 | 0.005039 | 0.049882 |
| LOC100409620 | -2.84024 | 3.56466 | 16.10667 | 0.005044 | 0.049883 |
| LOC108592870 | -1.26809 | 2.895925 | 11.55446 | 0.005047 | 0.049887 |

| | | | | | |
|---|---|---|---|---|---|
| NCK1 | -0.97397 | 4.408408 | 11.5434 | 0.005062 | 0.049943 |
| LOC100411778 | -0.92047 | 6.5179 | 11.54333 | 0.005062 | 0.049943 |
| DCP2 | -0.88019 | 5.519903 | 11.54308 | 0.005063 | 0.049943 |
| LOC100388343 | -1.6387 | 3.485042 | 12.20189 | 0.005066 | 0.049949 |

**Supp. Table 7b. Genes upregulated in non-IBD**

| Gene name | logFC | logCPM | F | PValue | FDR |
|---|---|---|---|---|---|
| TMEM252 | 6.19491 | 5.866223 | 206.2875 | 4.25E-09 | 2.77E-05 |
| LOC100397175 | 4.554002 | 4.723648 | 190.2493 | 6.78E-09 | 3.31E-05 |
| PLIN4 | 3.550317 | 6.267744 | 149.6418 | 2.73E-08 | 7.26E-05 |
| AADAC | 4.171532 | 5.48819 | 121.5845 | 8.96E-08 | 0.000146 |
| LOC100389947 | 4.407675 | 5.117669 | 123.1901 | 8.32E-08 | 0.000146 |
| TEF | 3.172237 | 5.671816 | 108.6723 | 1.69E-07 | 0.00022 |
| GGT7 | 3.50294 | 6.094515 | 84.29965 | 7.33E-07 | 0.000456 |
| RGS6 | 3.946081 | 5.703896 | 84.78888 | 1.11E-06 | 0.000534 |
| NELL2 | 4.148902 | 4.55659 | 80.21784 | 1.32E-06 | 0.000561 |
| G0S2 | 2.636739 | 5.748327 | 73.43371 | 1.46E-06 | 0.000593 |
| LOC103789763 | 4.134557 | 2.975444 | 69.03428 | 2.03E-06 | 0.000734 |
| ATP1A2 | 2.525538 | 6.541073 | 68.23001 | 2.16E-06 | 0.000767 |
| LEP | 3.953639 | 2.918747 | 67.64807 | 2.26E-06 | 0.000774 |
| FREM1 | 2.679725 | 3.905973 | 62.31154 | 3.48E-06 | 0.000973 |
| AQP7 | 4.024321 | 4.547448 | 70.1562 | 3.63E-06 | 0.000993 |
| TPSG1 | 3.11692 | 3.37826 | 59.99754 | 4.24E-06 | 0.001064 |
| LOC108592439 | 3.703493 | 3.079778 | 59.64112 | 4.38E-06 | 0.001083 |
| LOC108589026 | 4.49758 | 2.215521 | 59.04757 | 4.61E-06 | 0.0011 |
| MYOM1 | 2.426333 | 4.762947 | 58.40367 | 4.88E-06 | 0.001136 |
| SLC2A4 | 2.296187 | 5.434333 | 57.92743 | 5.09E-06 | 0.001145 |
| ANO1 | 2.457331 | 5.362416 | 57.97466 | 5.07E-06 | 0.001145 |
| DCLK1 | 2.166117 | 5.395598 | 56.85939 | 5.61E-06 | 0.001232 |
| RGMB | 2.028204 | 5.755381 | 56.60467 | 5.74E-06 | 0.001247 |
| HSPB6 | 2.003857 | 7.295012 | 54.64445 | 6.88E-06 | 0.001417 |
| RYR3 | 2.232323 | 5.985692 | 54.66016 | 6.87E-06 | 0.001417 |
| EPHA7 | 2.610796 | 5.773255 | 54.01862 | 7.53E-06 | 0.001486 |
| PER3 | 2.47095 | 6.034521 | 53.39364 | 7.75E-06 | 0.001515 |
| CKM | 2.127659 | 5.328615 | 53.1406 | 7.94E-06 | 0.001522 |
| CIDEA | 4.196708 | 3.143251 | 54.28099 | 8.53E-06 | 0.001545 |
| SPEG | 2.259867 | 6.616302 | 52.13261 | 8.76E-06 | 0.001546 |
| SYT1 | 2.304481 | 4.170779 | 52.10539 | 8.78E-06 | 0.001546 |
| FRZB | 2.221502 | 4.780678 | 51.92153 | 8.94E-06 | 0.001547 |
| COL4A5 | 1.961957 | 5.340855 | 51.57762 | 9.25E-06 | 0.001583 |
| GLIS1 | 2.678286 | 3.937933 | 50.96867 | 9.82E-06 | 0.001643 |
| BMP3 | 2.876879 | 4.254483 | 50.99656 | 9.83E-06 | 0.001643 |
| CACNA1C | 2.06918 | 6.567074 | 50.41918 | 1.04E-05 | 0.001684 |
| SYN2 | 2.241084 | 5.264714 | 50.37145 | 1.04E-05 | 0.001684 |
| SLC34A3 | 2.510446 | 8.502646 | 50.2952 | 1.05E-05 | 0.001684 |
| RBM24 | 2.328334 | 3.809601 | 50.19834 | 1.06E-05 | 0.001686 |
| BMP8A | 2.11047 | 6.028596 | 50.07303 | 1.07E-05 | 0.001694 |
| CHRNA3 | 3.774031 | 2.33637 | 49.64256 | 1.12E-05 | 0.001741 |

| | | | | | |
|---|---|---|---|---|---|
| RPS2 | 2.26099 | 5.714517 | 49.45716 | 1.14E-05 | 0.001761 |
| CBLN2 | 2.88199 | 3.135957 | 49.03907 | 1.19E-05 | 0.001809 |
| DBP | 2.171179 | 6.295702 | 47.93651 | 1.34E-05 | 0.001869 |
| NQO1 | 1.889691 | 6.267965 | 47.41106 | 1.41E-05 | 0.001881 |
| CDH2 | 2.382982 | 3.534421 | 47.46846 | 1.41E-05 | 0.001881 |
| GDPD2 | 2.270071 | 5.699413 | 47.18727 | 1.45E-05 | 0.00191 |
| PRPH | 2.57676 | 3.657857 | 47.07654 | 1.47E-05 | 0.00191 |
| RASD2 | 2.517761 | 4.744531 | 46.64819 | 1.53E-05 | 0.001947 |
| OSR2 | 2.881746 | 3.619542 | 46.31942 | 1.59E-05 | 0.002 |
| RNF224 | 2.402571 | 6.466389 | 45.76737 | 1.75E-05 | 0.002125 |
| LOC108590858 | 2.5923 | 4.28677 | 45.37284 | 1.76E-05 | 0.002125 |
| ADRA1A | 2.028717 | 4.396439 | 45.2781 | 1.78E-05 | 0.002133 |
| LDB3 | 1.988028 | 5.769764 | 45.17314 | 1.80E-05 | 0.002145 |
| MAPK4 | 1.893318 | 4.855432 | 44.69096 | 1.90E-05 | 0.00219 |
| PCLO | 2.028681 | 5.500951 | 44.92649 | 1.85E-05 | 0.00219 |
| GABBR2 | 2.055095 | 4.617509 | 44.71565 | 1.89E-05 | 0.00219 |
| AQP3 | 2.093948 | 6.421409 | 44.11662 | 2.02E-05 | 0.00226 |
| RNF112 | 2.510231 | 5.649482 | 44.95333 | 2.06E-05 | 0.002271 |
| KCNB1 | 2.059455 | 5.829437 | 43.62379 | 2.14E-05 | 0.002348 |
| PRDM8 | 2.180882 | 4.542456 | 43.35512 | 2.20E-05 | 0.002394 |
| TSPAN18 | 1.921805 | 5.301937 | 42.74205 | 2.36E-05 | 0.002511 |
| HLF | 2.346957 | 4.875833 | 42.46642 | 2.44E-05 | 0.002524 |
| SALL2 | 1.913229 | 4.934987 | 42.23397 | 2.51E-05 | 0.002539 |
| SLC1A1 | 2.654432 | 4.598068 | 42.96369 | 2.50E-05 | 0.002539 |
| TMEM82 | 2.077704 | 6.078813 | 41.84781 | 2.62E-05 | 0.002601 |
| ZNF704 | 1.886121 | 5.708206 | 41.6414 | 2.69E-05 | 0.00265 |
| DRP2 | 1.902843 | 5.520802 | 40.87706 | 2.94E-05 | 0.002808 |
| VAT1L | 2.491698 | 3.409287 | 40.881 | 2.94E-05 | 0.002808 |
| HAND1 | 1.91717 | 4.909874 | 40.69421 | 3.00E-05 | 0.002823 |
| ABCB1 | 1.781936 | 8.143904 | 39.95462 | 3.28E-05 | 0.003029 |
| RYR2 | 1.861574 | 6.458454 | 39.94803 | 3.28E-05 | 0.003029 |
| GALR2 | 2.832534 | 2.591109 | 39.69278 | 3.39E-05 | 0.003103 |
| CECR2 | 2.232522 | 4.605591 | 39.51583 | 3.46E-05 | 0.003105 |
| NR1D2 | 1.934459 | 5.5497 | 39.20364 | 3.60E-05 | 0.003168 |
| COL28A1 | 2.040345 | 4.700697 | 39.22607 | 3.59E-05 | 0.003168 |
| LOC100402764 | 6.947089 | 0.854317 | 42.65384 | 3.57E-05 | 0.003168 |
| SOSTDC1 | 2.061916 | 5.524363 | 38.70474 | 3.82E-05 | 0.00323 |
| ST6GALNAC5 | 2.715663 | 3.254308 | 38.43075 | 3.96E-05 | 0.003321 |
| CTNNA3 | 2.544346 | 4.281916 | 38.29513 | 4.20E-05 | 0.003476 |
| LOC103792464 | 1.660165 | 6.280032 | 37.57342 | 4.41E-05 | 0.003488 |
| CASQ2 | 1.67621 | 6.853454 | 37.06834 | 4.70E-05 | 0.003488 |
| CAP2 | 1.756273 | 5.471085 | 37.57057 | 4.41E-05 | 0.003488 |
| POSTN | 1.758602 | 6.952522 | 37.40331 | 4.50E-05 | 0.003488 |
| STUM | 1.797249 | 4.509242 | 37.37607 | 4.52E-05 | 0.003488 |
| PYGM | 1.799335 | 5.158674 | 37.14595 | 4.65E-05 | 0.003488 |
| KIT | 1.949633 | 4.986385 | 37.87627 | 4.24E-05 | 0.003488 |
| MTUS2 | 2.254063 | 4.28872 | 37.76432 | 4.30E-05 | 0.003488 |
| LOC100410434 | 2.26288 | 3.698602 | 37.2935 | 4.57E-05 | 0.003488 |

| | | | | | |
|---|---|---|---|---|---|
| OXCT2 | 2.358479 | 3.422957 | 37.51283 | 4.44E-05 | 0.003488 |
| SNAP25 | 2.437355 | 3.250946 | 37.17411 | 4.64E-05 | 0.003488 |
| STMN2 | 2.640493 | 3.828217 | 38.05953 | 4.29E-05 | 0.003488 |
| DGKG | 2.696469 | 3.783303 | 37.37737 | 4.68E-05 | 0.003488 |
| HTR2A | 2.721696 | 2.401185 | 37.8513 | 4.26E-05 | 0.003488 |
| TMEM151A | 2.743264 | 2.687211 | 37.05414 | 4.71E-05 | 0.003488 |
| KCNIP1 | 3.124636 | 2.045251 | 37.57618 | 4.41E-05 | 0.003488 |
| LUZP2 | 2.499126 | 3.862719 | 36.68698 | 4.94E-05 | 0.003589 |
| CACNA1H | 1.933904 | 6.692667 | 36.53376 | 5.04E-05 | 0.00361 |
| RAB3C | 2.214399 | 4.366046 | 36.52073 | 5.05E-05 | 0.00361 |
| GAL | 2.50452 | 3.151734 | 36.59508 | 5.00E-05 | 0.00361 |
| ITGA7 | 1.905235 | 5.424039 | 36.18596 | 5.27E-05 | 0.003707 |
| PDE4C | 2.784562 | 2.386682 | 35.8327 | 5.52E-05 | 0.003855 |
| DOPEY2 | 1.71681 | 7.383933 | 35.58963 | 5.70E-05 | 0.003952 |
| FRAS1 | 2.124487 | 6.164345 | 35.72933 | 5.82E-05 | 0.004019 |
| LOC103790125 | 1.777665 | 4.349305 | 35.41024 | 5.84E-05 | 0.004019 |
| GSTA3 | 1.789905 | 4.318044 | 35.27116 | 5.95E-05 | 0.004052 |
| PCSK2 | 3.380306 | 2.244319 | 35.26834 | 5.95E-05 | 0.004052 |
| OLFML2B | 1.889384 | 4.631951 | 35.00308 | 6.16E-05 | 0.004129 |
| COL4A6 | 2.025145 | 4.270833 | 35.01585 | 6.15E-05 | 0.004129 |
| TTC36 | 3.780515 | 1.580087 | 35.00634 | 6.16E-05 | 0.004129 |
| LOC108587995 | 2.031894 | 3.690575 | 34.96849 | 6.19E-05 | 0.004132 |
| LOC103796384 | 1.864526 | 9.600279 | 34.86722 | 6.28E-05 | 0.004174 |
| DGKB | 2.095924 | 4.473625 | 34.77507 | 6.35E-05 | 0.004197 |
| TMEM178B | 2.36925 | 3.051017 | 34.71947 | 6.40E-05 | 0.004215 |
| RPRM | 7.199031 | 1.06749 | 37.62736 | 6.53E-05 | 0.004266 |
| RBPMS2 | 1.659394 | 4.815453 | 34.25562 | 6.82E-05 | 0.004373 |
| RBFOX3 | 2.112051 | 3.329091 | 34.2608 | 6.81E-05 | 0.004373 |
| KCNK10 | 3.066094 | 2.812081 | 34.02382 | 7.04E-05 | 0.00441 |
| ARHGEF25 | 1.662859 | 5.587508 | 33.95679 | 7.10E-05 | 0.004436 |
| KCNK3 | 2.349902 | 3.976219 | 33.76812 | 7.29E-05 | 0.004524 |
| PHOX2B | 2.56816 | 2.701769 | 33.7391 | 7.32E-05 | 0.004526 |
| NLGN4X | 2.571327 | 2.538858 | 33.69732 | 7.36E-05 | 0.004526 |
| LOC100391398 | 2.919097 | 2.061735 | 33.69603 | 7.36E-05 | 0.004526 |
| PLD5 | 3.095784 | 1.718446 | 33.47447 | 7.59E-05 | 0.004609 |
| PDE1C | 1.706578 | 5.434919 | 33.37862 | 7.69E-05 | 0.004656 |
| ITIH5 | 1.538476 | 6.425516 | 33.30559 | 7.77E-05 | 0.004665 |
| WNK2 | 1.588299 | 7.850655 | 33.28134 | 7.80E-05 | 0.004665 |
| ADIRF | 1.752414 | 4.998633 | 33.3244 | 7.75E-05 | 0.004665 |
| CHRDL1 | 1.839865 | 4.660631 | 33.23303 | 7.85E-05 | 0.004665 |
| GHR | 2.050393 | 5.207044 | 33.44129 | 7.83E-05 | 0.004665 |
| ARHGEF37 | 1.587678 | 4.996367 | 33.18603 | 7.90E-05 | 0.004681 |
| SYNM | 1.787278 | 9.382107 | 33.09987 | 8.00E-05 | 0.004723 |
| PENK | 2.348864 | 4.068247 | 33.28976 | 8.03E-05 | 0.004729 |
| COL5A3 | 1.902551 | 7.240407 | 32.97859 | 8.13E-05 | 0.004775 |
| PCP4 | 1.922303 | 4.188715 | 32.85348 | 8.28E-05 | 0.004792 |
| CEND1 | 1.901872 | 4.164015 | 32.80642 | 8.33E-05 | 0.004806 |
| CTH | 1.851992 | 4.074545 | 32.50669 | 8.69E-05 | 0.004955 |

| | | | | | |
|---|---|---|---|---|---|
| GRB14 | 3.056891 | 2.11684 | 32.25141 | 9.01E-05 | 0.005108 |
| LOC108588023 | 4.99107 | 1.075377 | 32.15472 | 9.14E-05 | 0.005164 |
| ARHGEF26 | 1.682327 | 5.808882 | 32.06455 | 9.26E-05 | 0.005216 |
| HECW1 | 3.187623 | 1.930607 | 32.04315 | 9.29E-05 | 0.005217 |
| METTL7B | 2.165152 | 5.193406 | 32.74949 | 9.33E-05 | 0.005225 |
| KCNQ3 | 2.684358 | 2.941366 | 31.99127 | 9.35E-05 | 0.005226 |
| LOC103793682 | 2.460309 | 3.341514 | 31.83107 | 9.57E-05 | 0.005287 |
| GRIA1 | 2.784226 | 2.652812 | 31.85549 | 9.54E-05 | 0.005287 |
| TACR2 | 1.672162 | 5.616208 | 31.75261 | 9.68E-05 | 0.005317 |
| CNTFR | 2.207282 | 3.329454 | 31.65995 | 9.81E-05 | 0.005374 |
| SVIL | 1.706418 | 9.070144 | 31.54605 | 9.97E-05 | 0.005433 |
| PTCHD1 | 2.454292 | 2.972083 | 31.45081 | 0.000101 | 0.005478 |
| KCNJ13 | 1.454662 | 6.083811 | 31.4113 | 0.000102 | 0.005494 |
| LMOD1 | 1.583476 | 7.336146 | 31.3097 | 0.000103 | 0.005504 |
| COL9A1 | 1.781857 | 4.339755 | 31.28509 | 0.000104 | 0.005504 |
| PCDH18 | 1.795094 | 4.991902 | 31.30181 | 0.000103 | 0.005504 |
| ADAMTS12 | 1.923224 | 3.644819 | 31.36639 | 0.000102 | 0.005504 |
| LOC100896813 | 2.993013 | 1.789925 | 31.22615 | 0.000104 | 0.005537 |
| PCDH9 | 2.786471 | 2.299116 | 31.0793 | 0.000107 | 0.005642 |
| SRPK3 | 2.172464 | 3.358458 | 31.0551 | 0.000107 | 0.005646 |
| NPTX2 | 4.928248 | 3.700397 | 45.03878 | 0.000108 | 0.005646 |
| LOC103791965 | 2.026059 | 4.185541 | 30.96893 | 0.000109 | 0.005649 |
| KANK1 | 1.428348 | 6.988563 | 30.8926 | 0.00011 | 0.005691 |
| PCDH17 | 1.978962 | 3.796824 | 30.80178 | 0.000111 | 0.005746 |
| SUGCT | 2.240527 | 3.529086 | 30.79077 | 0.000111 | 0.005746 |
| UNC5D | 2.60313 | 2.732283 | 30.73068 | 0.000112 | 0.005752 |
| TNS1 | 1.656806 | 9.011369 | 30.67931 | 0.000113 | 0.005765 |
| LOC108589387 | 2.435966 | 2.984491 | 30.64004 | 0.000114 | 0.005769 |
| LOC100389020 | 1.807083 | 5.313391 | 30.61799 | 0.000114 | 0.005773 |
| PCP4L1 | 1.788637 | 5.052352 | 30.56823 | 0.000115 | 0.005801 |
| PRKG1 | 1.664765 | 6.017526 | 30.45119 | 0.000117 | 0.005842 |
| LOC100399997 | 1.932279 | 4.440908 | 30.26291 | 0.00012 | 0.005942 |
| PLIN1 | 3.37899 | 5.299308 | 41.20389 | 0.000121 | 0.005942 |
| NPTX1 | 2.973769 | 2.347213 | 30.15873 | 0.000122 | 0.005981 |
| LRRC75B | 1.663325 | 4.680359 | 30.1096 | 0.000123 | 0.00601 |
| ZNF503 | 1.643095 | 4.85496 | 30.04756 | 0.000124 | 0.006036 |
| ABCG8 | 1.55784 | 8.22293 | 29.93584 | 0.000127 | 0.006123 |
| TMEM59L | 3.051389 | 1.707329 | 29.91827 | 0.000127 | 0.006124 |
| LOC108589969 | 6.435424 | 0.442842 | 32.06072 | 0.000127 | 0.006129 |
| ADHFE1 | 3.45514 | 3.03354 | 32.89972 | 0.00013 | 0.006195 |
| HRH2 | 2.270008 | 4.258396 | 30.17887 | 0.000131 | 0.006252 |
| CPED1 | 1.511317 | 5.95751 | 29.50092 | 0.000135 | 0.006365 |
| CA10 | 3.698675 | 1.226309 | 29.48679 | 0.000135 | 0.006365 |
| SPTB | 1.628626 | 4.699961 | 29.41751 | 0.000137 | 0.006397 |
| FRY | 1.670308 | 6.591877 | 29.40705 | 0.000137 | 0.006397 |
| FER1L5 | 2.118514 | 5.392811 | 30.50536 | 0.000136 | 0.006397 |
| TAGLN3 | 3.842476 | 1.346069 | 29.36993 | 0.000138 | 0.006409 |
| TRPC4 | 2.335163 | 2.952229 | 29.30473 | 0.000139 | 0.006425 |

| | | | | | |
|---|---|---|---|---|---|
| LOC103791174 | 3.781169 | 3.691566 | 37.7592 | 0.000139 | 0.006425 |
| TUB | 1.534016 | 5.205466 | 29.23794 | 0.000141 | 0.006449 |
| LOC100413669 | 1.537023 | 6.600527 | 29.16635 | 0.000142 | 0.006449 |
| TMOD1 | 2.063598 | 3.433426 | 29.17304 | 0.000142 | 0.006449 |
| PARM1 | 1.449845 | 5.285179 | 29.07215 | 0.000144 | 0.006501 |
| PRNP | 1.592807 | 6.529152 | 28.99483 | 0.000146 | 0.006509 |
| IGSF22 | 2.314142 | 2.296062 | 28.86751 | 0.000149 | 0.006603 |
| SLITRK5 | 1.937029 | 3.623577 | 28.84997 | 0.000149 | 0.006606 |
| PGM5 | 1.686795 | 7.87905 | 28.75739 | 0.000151 | 0.006671 |
| LOC108590272 | 2.248149 | 4.803868 | 30.21071 | 0.000151 | 0.006671 |
| ACACB | 1.599895 | 7.78655 | 28.73059 | 0.000152 | 0.006684 |
| SLC13A2 | 1.862765 | 7.177327 | 28.8902 | 0.000152 | 0.006684 |
| PDLIM3 | 1.605557 | 6.64814 | 28.67066 | 0.000154 | 0.006686 |
| BNC2 | 1.570972 | 5.636102 | 28.64329 | 0.000154 | 0.0067 |
| ATP2B3 | 3.520845 | 1.396782 | 28.61263 | 0.000155 | 0.006702 |
| CNN1 | 1.626473 | 8.519895 | 28.43075 | 0.000159 | 0.006763 |
| ACTG2 | 1.803008 | 10.6776 | 28.45395 | 0.000159 | 0.006763 |
| AMOTL1 | 1.573738 | 7.043951 | 28.36589 | 0.000161 | 0.006814 |
| SMAD9 | 1.563345 | 5.350337 | 28.20726 | 0.000165 | 0.006889 |
| REEP1 | 1.578143 | 4.784817 | 28.2122 | 0.000165 | 0.006889 |
| LOC103790969 | 1.587954 | 5.764329 | 28.23447 | 0.000164 | 0.006889 |
| LOC103791695 | 1.820972 | 4.101002 | 28.21268 | 0.000165 | 0.006889 |
| PCDH10 | 2.317422 | 3.587796 | 28.30644 | 0.000165 | 0.006889 |
| SLC26A10 | 1.668005 | 6.142511 | 28.12021 | 0.000167 | 0.006894 |
| LOC100403233 | 1.933747 | 5.157503 | 28.3253 | 0.000167 | 0.006894 |
| LOC103794263 | 2.208284 | 2.718588 | 28.11542 | 0.000167 | 0.006894 |
| EML6 | 1.717628 | 6.593907 | 27.98284 | 0.000171 | 0.006948 |
| SNAP91 | 3.150641 | 1.612983 | 27.97324 | 0.000171 | 0.006948 |
| MYL9 | 1.659951 | 9.545298 | 27.84985 | 0.000175 | 0.007045 |
| LRRTM3 | 2.329254 | 2.451334 | 27.84607 | 0.000175 | 0.007045 |
| SLC25A45 | 1.905272 | 4.492227 | 27.77284 | 0.000177 | 0.0071 |
| IYD | 2.004284 | 3.978118 | 27.73753 | 0.000178 | 0.007122 |
| MELTF | 4.646032 | 4.035911 | 42.24591 | 0.000178 | 0.007122 |
| SLC5A4 | 1.522744 | 6.557559 | 27.68052 | 0.000179 | 0.007145 |
| HAND2 | 1.67192 | 4.657594 | 27.61723 | 0.000181 | 0.007157 |
| JPH2 | 1.756079 | 6.318007 | 27.63065 | 0.000181 | 0.007157 |
| MEGF10 | 2.473514 | 2.388313 | 27.61384 | 0.000181 | 0.007157 |
| TIMP4 | 2.557213 | 3.583557 | 28.44187 | 0.000183 | 0.007188 |
| C2CD4D | 2.9149 | 2.339098 | 27.56344 | 0.000183 | 0.007188 |
| GRIN2A | 3.391859 | 2.083221 | 27.63925 | 0.000185 | 0.007263 |
| SLC24A3 | 1.677133 | 4.367193 | 27.41687 | 0.000187 | 0.007306 |
| PDZRN4 | 1.840473 | 5.320362 | 27.56439 | 0.000188 | 0.007306 |
| PPP1R3F | 1.929594 | 5.009745 | 27.54808 | 0.000189 | 0.007319 |
| CLIP3 | 1.36213 | 6.776139 | 27.24439 | 0.000192 | 0.007421 |
| SARDH | 2.950452 | 5.885277 | 35.744 | 0.000192 | 0.007421 |
| TMEM184A | 1.654571 | 6.621825 | 27.19951 | 0.000194 | 0.007445 |
| WSCD2 | 2.309825 | 4.401388 | 28.63243 | 0.000194 | 0.007445 |
| MACROD1 | 1.605791 | 5.621175 | 27.01524 | 0.0002 | 0.007581 |

| | | | | | |
|---|---|---|---|---|---|
| MPP7 | 1.523677 | 5.43507 | 26.83264 | 0.000206 | 0.007603 |
| RAPGEFL1 | 1.558034 | 4.872444 | 26.9348 | 0.000202 | 0.007603 |
| MAB21L2 | 1.564447 | 6.804885 | 26.88459 | 0.000204 | 0.007603 |
| PNMA1 | 1.565015 | 6.711437 | 26.87463 | 0.000204 | 0.007603 |
| MOXD1 | 1.609613 | 4.958881 | 26.91353 | 0.000203 | 0.007603 |
| MOGAT1 | 2.046081 | 3.395344 | 26.89182 | 0.000204 | 0.007603 |
| PLPPR2 | 2.197193 | 2.537102 | 26.84849 | 0.000205 | 0.007603 |
| LOC103795901 | 2.91685 | 3.088179 | 28.19309 | 0.000203 | 0.007603 |
| LOC108588695 | 3.218005 | 1.431451 | 26.83801 | 0.000206 | 0.007603 |
| TREH | 2.374469 | 8.799557 | 29.42344 | 0.000207 | 0.007631 |
| CALML3 | 2.460779 | 3.637407 | 27.4876 | 0.000209 | 0.007666 |
| DMPK | 1.554876 | 6.887867 | 26.66837 | 0.000211 | 0.007708 |
| RIC3 | 2.312756 | 2.747587 | 26.63716 | 0.000212 | 0.007733 |
| SMYD1 | 2.020049 | 3.895072 | 26.61689 | 0.000213 | 0.007744 |
| NGFR | 1.565366 | 4.545613 | 26.52296 | 0.000216 | 0.007825 |
| GPR20 | 1.814116 | 3.937003 | 26.4839 | 0.000218 | 0.007856 |
| ANK2 | 1.350402 | 6.659489 | 26.46243 | 0.000219 | 0.00787 |
| LOC108593562 | 3.107954 | 1.886552 | 26.39749 | 0.000221 | 0.007911 |
| MCAM | 1.391158 | 7.354371 | 26.34465 | 0.000223 | 0.007936 |
| SORBS1 | 1.526075 | 9.160055 | 26.25342 | 0.000226 | 0.007999 |
| LOC100399432 | 2.751722 | 1.857152 | 26.24156 | 0.000227 | 0.008 |
| NFATC4 | 1.608358 | 4.993714 | 26.16803 | 0.000229 | 0.008054 |
| PPP2R2C | 2.627441 | 2.051631 | 26.0925 | 0.000232 | 0.008127 |
| PRICKLE2 | 1.432064 | 5.723011 | 26.07088 | 0.000233 | 0.008127 |
| ARSD | 1.459344 | 6.009561 | 26.04725 | 0.000234 | 0.00813 |
| ANXA13 | 1.935721 | 3.916809 | 26.05052 | 0.000234 | 0.00813 |
| LCT | 1.490796 | 7.950912 | 26.00294 | 0.000236 | 0.008153 |
| IL1RAP | 1.657931 | 4.350514 | 25.99074 | 0.000236 | 0.008153 |
| F11 | 4.67229 | 1.987678 | 29.37174 | 0.000235 | 0.008153 |
| ALDH1L1 | 3.673638 | 5.656301 | 39.27895 | 0.000238 | 0.008194 |
| MAP1B | 1.406828 | 7.997955 | 25.91682 | 0.000239 | 0.008201 |
| TENM4 | 1.971073 | 4.078328 | 25.91629 | 0.000239 | 0.008201 |
| KIF1A | 1.984019 | 4.372586 | 25.80789 | 0.00025 | 0.008346 |
| LOC103795648 | 2.062731 | 2.808752 | 25.69125 | 0.000248 | 0.008346 |
| EFNA5 | 2.087034 | 3.663411 | 25.73682 | 0.000247 | 0.008346 |
| GAP43 | 2.236004 | 2.398331 | 25.58259 | 0.000253 | 0.008443 |
| TMTC2 | 1.411188 | 5.271265 | 25.51839 | 0.000256 | 0.008502 |
| ELAVL4 | 2.348187 | 2.618688 | 25.5111 | 0.000256 | 0.008502 |
| NT5C1A | 3.888733 | 0.908724 | 25.47094 | 0.000258 | 0.008531 |
| ADCY5 | 1.447722 | 5.925257 | 25.44199 | 0.000259 | 0.008532 |
| MN1 | 1.5562 | 4.789756 | 25.4464 | 0.000259 | 0.008532 |
| UNC80 | 2.311163 | 2.664656 | 25.43055 | 0.00026 | 0.008532 |
| MAPK10 | 2.660771 | 2.158321 | 25.43782 | 0.000259 | 0.008532 |
| NEFL | 2.479737 | 2.392521 | 25.29566 | 0.000266 | 0.008671 |
| KCNMB1 | 1.435855 | 4.875688 | 25.25441 | 0.000268 | 0.008673 |
| AKR1B10 | 1.701072 | 7.921724 | 25.23584 | 0.000268 | 0.008673 |
| PER2 | 1.765688 | 6.569793 | 25.41224 | 0.000268 | 0.008673 |
| CCDC80 | 1.547115 | 5.267057 | 25.16171 | 0.000272 | 0.008734 |

| | | | | | |
|---|---|---|---|---|---|
| SYNPO2 | 1.590486 | 9.433546 | 25.16417 | 0.000272 | 0.008734 |
| RGMA | 1.616883 | 4.732686 | 25.15607 | 0.000272 | 0.008734 |
| TTLL7 | 1.865741 | 4.600432 | 25.305 | 0.000271 | 0.008734 |
| PTGS2 | 1.893768 | 4.469914 | 25.24121 | 0.000274 | 0.008744 |
| LOC103794993 | 1.489323 | 6.232528 | 25.1017 | 0.000275 | 0.008758 |
| COL27A1 | 1.77787 | 5.853928 | 25.1711 | 0.00028 | 0.008837 |
| LOC100395613 | 2.13522 | 2.894209 | 25.0096 | 0.000279 | 0.008837 |
| LGALS12 | 5.073817 | 1.874788 | 29.00191 | 0.000279 | 0.008837 |
| FAM13C | 1.877043 | 4.632806 | 25.07674 | 0.000282 | 0.00889 |
| YAP1 | 1.427353 | 7.061924 | 24.74656 | 0.000292 | 0.009135 |
| LOC103793212 | 1.691827 | 3.959709 | 24.73055 | 0.000293 | 0.00914 |
| C10H15orf52 | 1.566909 | 4.683985 | 24.70487 | 0.000294 | 0.009166 |
| HSD17B2 | 1.340272 | 5.832424 | 24.56105 | 0.000301 | 0.009323 |
| RUNDC3B | 1.672022 | 3.701904 | 24.41031 | 0.000309 | 0.00945 |
| GRIK3 | 1.785831 | 3.060082 | 24.4644 | 0.000306 | 0.00945 |
| SLN | 2.279856 | 2.253899 | 24.43065 | 0.000308 | 0.00945 |
| ACTN2 | 3.252596 | 2.085436 | 24.58551 | 0.000309 | 0.00945 |
| NKX3-2 | 2.404604 | 2.700846 | 24.38702 | 0.000311 | 0.009474 |
| CPEB1 | 2.315528 | 3.204277 | 24.45737 | 0.000312 | 0.009478 |
| DACT3 | 1.553229 | 5.136629 | 24.34683 | 0.000313 | 0.009496 |
| SLC2A7 | 1.429129 | 4.923781 | 24.29045 | 0.000316 | 0.009531 |
| LOC100405319 | 1.639249 | 8.162265 | 24.30623 | 0.000315 | 0.009531 |
| LOC103787446 | 1.980155 | 3.157177 | 24.31318 | 0.000315 | 0.009531 |
| PTPRS | 1.498007 | 6.678518 | 24.25075 | 0.000318 | 0.009568 |
| C16H8orf88 | 2.60843 | 2.273194 | 24.21471 | 0.00032 | 0.009614 |
| NDRG4 | 1.527236 | 5.097613 | 24.17259 | 0.000322 | 0.00967 |
| STON1 | 1.563009 | 6.768676 | 24.11925 | 0.000325 | 0.009692 |
| DUSP26 | 1.869724 | 3.473993 | 24.13197 | 0.000325 | 0.009692 |
| LOC108588020 | 1.778661 | 3.886337 | 24.05637 | 0.000329 | 0.00975 |
| CADM2 | 2.28936 | 2.831894 | 24.01916 | 0.000331 | 0.009799 |
| GCKR | 3.011479 | 3.102712 | 26.32769 | 0.000335 | 0.009883 |
| PPP1R1A | 2.838692 | 2.027453 | 23.89052 | 0.000339 | 0.009979 |
| SMTN | 1.465041 | 8.228397 | 23.86709 | 0.00034 | 0.009981 |
| IGFBP4 | 1.539038 | 7.653672 | 23.86355 | 0.000341 | 0.009981 |
| FAM83D | 1.611661 | 7.022137 | 23.87381 | 0.00034 | 0.009981 |
| CRYAB | 1.509314 | 4.075105 | 23.84885 | 0.000341 | 0.009992 |
| RBPMS | 1.452884 | 7.215165 | 23.81242 | 0.000344 | 0.010028 |
| DMD | 1.513144 | 8.374305 | 23.75568 | 0.000347 | 0.010113 |
| NPR3 | 1.587908 | 6.488432 | 23.73253 | 0.000349 | 0.01014 |
| NKD2 | 2.091483 | 3.468219 | 23.71587 | 0.00035 | 0.010153 |
| FSTL5 | 2.895987 | 1.826959 | 23.66135 | 0.000353 | 0.010193 |
| FGF1 | 1.679869 | 4.575362 | 23.63225 | 0.000355 | 0.010217 |
| LOC108588075 | 1.747315 | 3.382208 | 23.63155 | 0.000355 | 0.010217 |
| RET | 1.925166 | 3.627441 | 23.56048 | 0.000359 | 0.010332 |
| PDK2 | 1.608719 | 3.985896 | 23.46506 | 0.000366 | 0.01046 |
| SLITRK2 | 2.655079 | 1.650244 | 23.45868 | 0.000366 | 0.01046 |
| DMGDH | 4.353721 | 1.243552 | 23.5517 | 0.000368 | 0.010507 |
| ADAMTSL3 | 1.605537 | 5.349962 | 23.40992 | 0.000369 | 0.010516 |

| | | | | | |
|---|---|---|---|---|---|
| NOX4 | 2.131886 | 2.461581 | 23.40484 | 0.000369 | 0.010516 |
| SLIT3 | 1.45514 | 4.788942 | 23.33461 | 0.000374 | 0.010604 |
| LOC100405958 | 1.634669 | 7.376774 | 23.33476 | 0.000376 | 0.010633 |
| GLRB | 3.64679 | 0.718116 | 23.27891 | 0.000378 | 0.010633 |
| LOC100397143 | 1.375963 | 8.485604 | 23.22772 | 0.000381 | 0.010667 |
| LOC100896731 | 1.485354 | 4.461954 | 23.22862 | 0.000381 | 0.010667 |
| FAT4 | 1.5137 | 8.039851 | 23.20015 | 0.000383 | 0.010667 |
| PVR | 1.543095 | 6.586595 | 23.21118 | 0.000383 | 0.010667 |
| GRIN3A | 2.353131 | 2.354902 | 23.21571 | 0.000382 | 0.010667 |
| NALCN | 2.856638 | 1.519157 | 23.19757 | 0.000384 | 0.010667 |
| LOC103793586 | 2.30673 | 2.539865 | 23.1619 | 0.000386 | 0.01072 |
| PPM1L | 1.454781 | 4.844209 | 23.1532 | 0.000387 | 0.010722 |
| PLXNA4 | 1.757671 | 3.601057 | 23.12646 | 0.000388 | 0.010743 |
| ABCC2 | 1.306988 | 7.568971 | 23.11329 | 0.000389 | 0.010754 |
| ACHE | 1.500326 | 4.670707 | 23.08422 | 0.000391 | 0.010791 |
| ZNF385D | 2.778526 | 1.612191 | 23.07869 | 0.000392 | 0.010791 |
| LOC103792852 | 1.864434 | 2.946799 | 23.06076 | 0.000393 | 0.010796 |
| MRVI1 | 1.345583 | 7.104678 | 23.02106 | 0.000396 | 0.010858 |
| GAS1 | 1.466971 | 5.618587 | 22.9979 | 0.000398 | 0.010889 |
| MTSS1L | 1.275068 | 6.585794 | 22.98126 | 0.000399 | 0.010894 |
| INSR | 1.261318 | 7.104633 | 22.90631 | 0.000404 | 0.010994 |
| LOC108588872 | 2.448239 | 2.08634 | 22.89914 | 0.000405 | 0.010994 |
| MAP1LC3A | 1.404301 | 4.561376 | 22.87152 | 0.000407 | 0.011034 |
| NMNAT2 | 1.755264 | 4.408218 | 22.83623 | 0.00041 | 0.011074 |
| SDSL | 1.752719 | 4.742583 | 22.91141 | 0.000413 | 0.011132 |
| LOC108593703 | 4.646106 | 0.79105 | 22.74625 | 0.000416 | 0.011196 |
| GPAM | 1.341462 | 8.512532 | 22.68784 | 0.000421 | 0.011216 |
| EML1 | 1.347796 | 5.295337 | 22.69689 | 0.00042 | 0.011216 |
| STC2 | 1.516038 | 4.356946 | 22.68244 | 0.000421 | 0.011216 |
| FOXP2 | 1.691552 | 4.395062 | 22.69791 | 0.00042 | 0.011216 |
| PSD | 1.788718 | 5.213692 | 23.11952 | 0.000422 | 0.011216 |
| EPHA4 | 1.683925 | 3.938338 | 22.66101 | 0.000423 | 0.011233 |
| OVOL2 | 1.602941 | 4.959671 | 22.65057 | 0.000424 | 0.011239 |
| LOC100406585 | 1.500184 | 6.68822 | 22.63355 | 0.000425 | 0.011243 |
| LAMA2 | 1.353839 | 4.436489 | 22.59268 | 0.000428 | 0.011313 |
| NRXN2 | 1.368565 | 5.245619 | 22.5666 | 0.00043 | 0.011352 |
| GPR17 | 1.328593 | 5.54186 | 22.54498 | 0.000432 | 0.011382 |
| LOC100406157 | 1.395368 | 7.501267 | 22.49041 | 0.000436 | 0.01145 |
| LOC103789397 | 2.439547 | 1.643004 | 22.49149 | 0.000436 | 0.01145 |
| LOC100402705 | 2.276378 | 1.929642 | 22.47884 | 0.000437 | 0.011459 |
| KCNN3 | 1.428926 | 4.304184 | 22.47149 | 0.000438 | 0.01146 |
| ADGRB3 | 1.973329 | 3.518655 | 22.44755 | 0.00044 | 0.011495 |
| PLXNB3 | 1.750235 | 4.457788 | 22.42978 | 0.000441 | 0.011517 |
| GLI3 | 1.378931 | 5.416456 | 22.35361 | 0.000447 | 0.011634 |
| ADTRP | 1.357085 | 4.340813 | 22.29352 | 0.000452 | 0.011717 |
| LOC103791888 | 1.617909 | 3.444211 | 22.15849 | 0.000464 | 0.011984 |
| KIF26A | 1.815009 | 4.399101 | 22.27154 | 0.000464 | 0.011984 |
| KCNQ4 | 1.490684 | 3.724971 | 22.09984 | 0.000469 | 0.012067 |

| | | | | | |
|---|---|---|---|---|---|
| OTOG | 2.235233 | 2.963261 | 22.06172 | 0.000472 | 0.012121 |
| KIAA2012 | 1.919699 | 2.740104 | 22.04583 | 0.000474 | 0.012141 |
| WFIKKN2 | 2.959515 | 2.982496 | 24.12591 | 0.00048 | 0.012266 |
| ZCCHC12 | 1.870531 | 2.669051 | 21.92825 | 0.000484 | 0.012346 |
| ABCA8 | 1.37679 | 6.130754 | 21.88891 | 0.000488 | 0.012406 |
| GRP | 3.103886 | 1.056922 | 21.75396 | 0.0005 | 0.012691 |
| APBB1 | 1.702847 | 3.378289 | 21.70799 | 0.000505 | 0.012753 |
| LRAT | 2.081711 | 3.987754 | 22.34707 | 0.000507 | 0.012753 |
| HEY2 | 2.289603 | 1.924063 | 21.65383 | 0.00051 | 0.012784 |
| SCN9A | 1.392904 | 4.298722 | 21.58699 | 0.000516 | 0.012875 |
| NPB | 1.963738 | 2.744737 | 21.56925 | 0.000518 | 0.012875 |
| CPM | 2.097976 | 4.307413 | 22.92456 | 0.000516 | 0.012875 |
| CYS1 | 1.952436 | 4.289459 | 22.09884 | 0.00052 | 0.012895 |
| FOLR2 | 1.684212 | 3.494981 | 21.5332 | 0.000522 | 0.012914 |
| SLC24A2 | 2.051532 | 2.36306 | 21.50591 | 0.000525 | 0.012965 |
| AQP1 | 1.373097 | 5.108082 | 21.48243 | 0.000527 | 0.013006 |
| SERTAD4 | 1.88547 | 2.977928 | 21.44599 | 0.000531 | 0.013074 |
| TAC1 | 2.099081 | 3.089442 | 21.44183 | 0.000531 | 0.013074 |
| LOC103790447 | 1.28446 | 4.475476 | 21.41904 | 0.000533 | 0.013114 |
| APBA2 | 1.332574 | 4.532868 | 21.39676 | 0.000536 | 0.013153 |
| BRSK1 | 1.63879 | 5.208347 | 21.43172 | 0.000543 | 0.013304 |
| NPC1L1 | 1.45595 | 10.76394 | 21.28194 | 0.000547 | 0.013394 |
| OPCML | 2.879831 | 1.124502 | 21.23917 | 0.000552 | 0.013488 |
| THBS2 | 1.51391 | 4.993129 | 21.21455 | 0.000554 | 0.013506 |
| NTM | 1.675655 | 3.319086 | 21.21141 | 0.000555 | 0.013506 |
| LOC108590397 | 1.760281 | 3.07219 | 21.21964 | 0.000554 | 0.013506 |
| LOC100401401 | 2.151931 | 2.814675 | 21.2059 | 0.000555 | 0.013506 |
| CACNA1E | 1.635584 | 4.613996 | 21.19026 | 0.000557 | 0.01353 |
| SIK1 | 1.357898 | 4.801754 | 21.1317 | 0.000563 | 0.01366 |
| TUBAL3 | 1.736069 | 7.325451 | 21.69859 | 0.000565 | 0.01366 |
| CDH19 | 1.873935 | 4.010842 | 21.21172 | 0.000566 | 0.01366 |
| LOC108589277 | 2.184154 | 3.276652 | 21.24087 | 0.000565 | 0.01366 |
| BMF | 1.29793 | 5.284012 | 21.08513 | 0.000568 | 0.013687 |
| ADAM23 | 2.085453 | 3.574715 | 21.29086 | 0.000569 | 0.013687 |
| LRTM1 | 3.691999 | 0.753067 | 21.03088 | 0.000574 | 0.013765 |
| RTN2 | 1.279187 | 4.483274 | 21.00354 | 0.000577 | 0.013769 |
| ANKDD1B | 1.386618 | 5.585457 | 21.01488 | 0.000576 | 0.013769 |
| TGFB1I1 | 1.421797 | 6.344994 | 21.00532 | 0.000577 | 0.013769 |
| CSRP1 | 1.357707 | 8.987437 | 20.99386 | 0.000579 | 0.013777 |
| MMP28 | 1.345793 | 5.294414 | 20.97473 | 0.000581 | 0.013811 |
| LOC103796764 | 1.337511 | 5.072061 | 20.9667 | 0.000582 | 0.013816 |
| TTYH2 | 1.8325 | 3.259365 | 20.93957 | 0.000585 | 0.013838 |
| PDILT | 1.842389 | 4.124205 | 21.06676 | 0.000584 | 0.013838 |
| MEIS1 | 1.265369 | 5.808259 | 20.8939 | 0.00059 | 0.013927 |
| SCN7A | 1.584678 | 6.406994 | 20.99017 | 0.000593 | 0.01393 |
| LOC108590733 | 1.981226 | 3.088821 | 20.87802 | 0.000592 | 0.01393 |
| SMPX | 2.734694 | 1.420207 | 20.87123 | 0.000592 | 0.01393 |
| KLHL31 | 3.345539 | 1.214816 | 20.88315 | 0.000591 | 0.01393 |

| | | | | | |
|---|---|---|---|---|---|
| FMO5 | 1.365033 | 5.80434 | 20.85156 | 0.000595 | 0.013945 |
| FBXO32 | 1.945709 | 7.489874 | 22.70792 | 0.000595 | 0.013945 |
| DLC1 | 1.452362 | 7.101734 | 20.80389 | 0.0006 | 0.013995 |
| FMO2 | 2.143207 | 2.321835 | 20.80085 | 0.000601 | 0.013995 |
| MYBL2 | 1.333713 | 5.05488 | 20.77471 | 0.000604 | 0.014049 |
| COL4A3 | 1.434273 | 4.392497 | 20.76237 | 0.000605 | 0.014066 |
| DCST1 | 1.354286 | 6.661349 | 20.75474 | 0.000606 | 0.01407 |
| SLC22A17 | 1.427535 | 5.219352 | 20.74812 | 0.000607 | 0.014072 |
| CRYBG3 | 1.277164 | 5.924301 | 20.73313 | 0.000608 | 0.014092 |
| GARNL3 | 1.639504 | 3.710162 | 20.72852 | 0.000609 | 0.014092 |
| CALCB | 3.337047 | 1.799211 | 21.02396 | 0.000612 | 0.014136 |
| BVES | 1.361416 | 4.570109 | 20.63829 | 0.00062 | 0.014196 |
| AR | 1.797367 | 2.658202 | 20.6635 | 0.000617 | 0.014196 |
| ADH4 | 1.854511 | 3.569643 | 20.63653 | 0.00062 | 0.014196 |
| GYG2 | 1.50994 | 3.977103 | 20.61532 | 0.000623 | 0.014238 |
| MAGI1 | 1.177593 | 7.103935 | 20.51412 | 0.000635 | 0.014421 |
| RBP4 | 1.327305 | 4.445163 | 20.51837 | 0.000635 | 0.014421 |
| LOC108587886 | 1.890508 | 2.922236 | 20.50192 | 0.000637 | 0.014439 |
| PTPRD | 1.165438 | 7.6969 | 20.45582 | 0.000642 | 0.01452 |
| PPP1R14A | 1.770774 | 4.807268 | 20.91302 | 0.000642 | 0.01452 |
| RASL11B | 1.810135 | 3.800802 | 20.44502 | 0.000644 | 0.014534 |
| KLF9 | 1.280025 | 5.90638 | 20.3981 | 0.00065 | 0.014584 |
| NBL1 | 1.463139 | 7.645396 | 20.3554 | 0.000655 | 0.014645 |
| GOLT1A | 1.558148 | 5.174755 | 20.41109 | 0.000656 | 0.014645 |
| ZNF560 | 3.389808 | 1.846102 | 20.90524 | 0.000657 | 0.014645 |
| CHN1 | 1.6888 | 3.510193 | 20.27091 | 0.000666 | 0.014801 |
| PLA2G4F | 1.725309 | 5.576409 | 20.98454 | 0.00067 | 0.014873 |
| CPNE6 | 1.924164 | 2.377914 | 20.22721 | 0.000672 | 0.014895 |
| LOC108588029 | 2.725817 | 3.610631 | 23.54095 | 0.000675 | 0.014929 |
| AFF3 | 1.294608 | 5.196575 | 20.15302 | 0.000682 | 0.015031 |
| RIMS1 | 1.731643 | 4.771735 | 20.46505 | 0.000682 | 0.015031 |
| LOC103789474 | 2.898406 | 1.17107 | 20.14296 | 0.000683 | 0.015034 |
| PNMA2 | 2.513684 | 2.285764 | 20.09434 | 0.00069 | 0.015156 |
| ASB5 | 2.649229 | 2.685327 | 20.65685 | 0.000697 | 0.015256 |
| CDH20 | 2.38109 | 2.015749 | 20.02179 | 0.0007 | 0.01529 |
| TRPC3 | 1.761924 | 3.339409 | 19.93277 | 0.000712 | 0.01546 |
| AKAP6 | 1.276668 | 5.543383 | 19.88968 | 0.000719 | 0.015474 |
| TSHZ2 | 1.28748 | 6.020595 | 19.90807 | 0.000716 | 0.015474 |
| LOC100386548 | 1.331694 | 4.461469 | 19.90495 | 0.000716 | 0.015474 |
| PLCL1 | 1.445777 | 5.084865 | 19.90014 | 0.000717 | 0.015474 |
| SGCA | 2.554089 | 1.777003 | 19.91155 | 0.000715 | 0.015474 |
| COPRS | 1.838026 | 2.948841 | 19.87538 | 0.000721 | 0.015501 |
| CAMKV | 1.852711 | 3.730424 | 19.90719 | 0.000725 | 0.015564 |
| KLF15 | 2.352794 | 3.090394 | 20.33641 | 0.00073 | 0.015612 |
| REEP6 | 2.094077 | 6.779062 | 23.01422 | 0.000734 | 0.015668 |
| IRS2 | 1.342915 | 5.416063 | 19.77231 | 0.000736 | 0.015686 |
| LIMS2 | 1.295253 | 6.325964 | 19.7473 | 0.000739 | 0.015709 |
| LOC100390425 | 1.514242 | 7.876992 | 19.73987 | 0.000741 | 0.015709 |

| | | | | | |
|---|---|---|---|---|---|
| LOC108593351 | 2.019317 | 2.19736 | 19.74998 | 0.000739 | 0.015709 |
| LOC103792971 | 1.568456 | 4.747187 | 19.72971 | 0.000742 | 0.015718 |
| PLPP2 | 1.450356 | 6.649081 | 19.71314 | 0.000744 | 0.015737 |
| DPYSL5 | 2.822378 | 1.11142 | 19.70042 | 0.000746 | 0.01576 |
| FERMT2 | 1.398292 | 6.012752 | 19.66949 | 0.000751 | 0.015773 |
| PDLIM7 | 1.29827 | 6.517715 | 19.64743 | 0.000754 | 0.015809 |
| ERICH3 | 2.243354 | 1.830903 | 19.6512 | 0.000754 | 0.015809 |
| SLC52A3 | 1.532029 | 5.752067 | 19.69193 | 0.000762 | 0.015865 |
| NR1D1 | 2.061196 | 4.313323 | 21.1561 | 0.00076 | 0.015865 |
| ATP1B2 | 2.087967 | 2.277698 | 19.60103 | 0.000761 | 0.015865 |
| GLDN | 3.238719 | 0.856196 | 19.61343 | 0.00076 | 0.015865 |
| RUNX1T1 | 1.229228 | 5.738848 | 19.54291 | 0.00077 | 0.016026 |
| CTNNAL1 | 1.682977 | 3.914551 | 19.46501 | 0.000783 | 0.016229 |
| TRPM5 | 1.711332 | 3.352433 | 19.44156 | 0.000786 | 0.016289 |
| LOC100400153 | 2.652721 | 2.114895 | 19.45418 | 0.000794 | 0.016409 |
| COMT | 1.283634 | 7.183056 | 19.34211 | 0.000803 | 0.016482 |
| DPP6 | 2.746874 | 1.253008 | 19.34666 | 0.000802 | 0.016482 |
| MAPK8IP2 | 1.97804 | 2.326425 | 19.32165 | 0.000806 | 0.016499 |
| DBH | 4.749518 | 0.871745 | 19.46836 | 0.000806 | 0.016499 |
| HSPB7 | 1.623963 | 4.180049 | 19.31026 | 0.000808 | 0.01652 |
| OSBPL5 | 1.339749 | 4.428716 | 19.2946 | 0.00081 | 0.016538 |
| FAM124A | 1.439612 | 4.245357 | 19.26732 | 0.000815 | 0.016558 |
| CFL2 | 1.601644 | 5.737198 | 19.6165 | 0.000816 | 0.016558 |
| SLC7A14 | 1.772221 | 2.770094 | 19.28101 | 0.000813 | 0.016558 |
| MFAP3L | 1.74342 | 3.35611 | 19.25564 | 0.000817 | 0.016567 |
| AGTR1 | 1.696682 | 3.021026 | 19.22117 | 0.000823 | 0.016635 |
| CHAT | 1.690123 | 2.775998 | 19.18784 | 0.000828 | 0.01666 |
| ENPP2 | 1.376848 | 5.722384 | 19.17558 | 0.00083 | 0.016662 |
| EMILIN1 | 1.381918 | 6.062391 | 19.17206 | 0.000831 | 0.016662 |
| CYSRT1 | 1.463532 | 5.03674 | 19.15314 | 0.000834 | 0.016692 |
| LOC103792639 | 2.175239 | 2.054409 | 19.15705 | 0.000833 | 0.016692 |
| INPP5A | 1.146908 | 6.476087 | 19.13349 | 0.000837 | 0.016718 |
| WWTR1 | 1.165255 | 5.298437 | 19.13071 | 0.000838 | 0.016718 |
| MYH11 | 1.602384 | 12.38329 | 19.13753 | 0.000837 | 0.016718 |
| MCF2L2 | 1.340796 | 4.667561 | 19.10058 | 0.000843 | 0.016774 |
| SLC10A2 | 1.395272 | 5.797089 | 19.10632 | 0.000842 | 0.016774 |
| LOC103787870 | 2.24018 | 1.916395 | 19.09962 | 0.000843 | 0.016774 |
| LOC100410935 | 2.097831 | 2.078027 | 19.07708 | 0.000847 | 0.016834 |
| VANGL2 | 1.469754 | 3.582645 | 19.04781 | 0.000852 | 0.016867 |
| LOC103794805 | 4.388537 | 0.641565 | 19.05063 | 0.000852 | 0.016867 |
| BAG2 | 1.479223 | 4.573485 | 18.98094 | 0.000864 | 0.017015 |
| PRKG2 | 1.170094 | 4.917143 | 18.93751 | 0.000872 | 0.017134 |
| SPRY1 | 1.234214 | 4.854506 | 18.88709 | 0.000881 | 0.017261 |
| GPT2 | 1.411307 | 4.419395 | 18.87913 | 0.000883 | 0.017272 |
| JAM3 | 1.245855 | 4.645472 | 18.85755 | 0.000886 | 0.017332 |
| PRRT4 | 4.254613 | 0.482417 | 18.7939 | 0.000898 | 0.017457 |
| EBF1 | 1.303735 | 3.993769 | 18.74276 | 0.000908 | 0.017574 |
| PPM1E | 1.965116 | 2.687407 | 18.69421 | 0.000917 | 0.017712 |

| | | | | | |
|---|---|---|---|---|---|
| LOC108590767 | 1.477819 | 3.636886 | 18.68152 | 0.000919 | 0.017729 |
| LOC108590707 | 1.920009 | 6.530311 | 20.97229 | 0.000927 | 0.017853 |
| LOC108594082 | 2.282263 | 2.154868 | 18.59716 | 0.000936 | 0.018008 |
| LOC108590109 | 2.826279 | 1.466765 | 18.58851 | 0.000937 | 0.018023 |
| RORB | 2.117707 | 2.917623 | 18.61409 | 0.000944 | 0.018122 |
| CLDN5 | 1.822551 | 2.572298 | 18.53327 | 0.000948 | 0.018196 |
| ULK1 | 1.224616 | 5.712476 | 18.50941 | 0.000953 | 0.018251 |
| SERPINF1 | 1.151413 | 4.87158 | 18.50122 | 0.000955 | 0.018253 |
| CDO1 | 2.42381 | 1.610282 | 18.49508 | 0.000956 | 0.018253 |
| THRB | 1.178418 | 5.687641 | 18.44431 | 0.000966 | 0.018323 |
| AIFM3 | 1.364107 | 4.271334 | 18.46632 | 0.000962 | 0.018323 |
| MYLK | 1.401899 | 10.76939 | 18.47184 | 0.000961 | 0.018323 |
| LOC100389414 | 1.730105 | 2.680047 | 18.41259 | 0.000973 | 0.018357 |
| HES3 | 2.152124 | 1.845931 | 18.42307 | 0.00097 | 0.018357 |
| OBSL1 | 1.342758 | 5.019738 | 18.37749 | 0.00098 | 0.018363 |
| MSRB3 | 1.344456 | 4.211921 | 18.38295 | 0.000979 | 0.018363 |
| DPEP1 | 1.598844 | 8.780204 | 18.48815 | 0.000977 | 0.018363 |
| FLRT1 | 1.778702 | 4.622754 | 19.06976 | 0.000977 | 0.018363 |
| PNLIPRP2 | 1.203836 | 6.646638 | 18.35494 | 0.000985 | 0.018402 |
| LOC103789692 | 2.328569 | 1.787331 | 18.35515 | 0.000984 | 0.018402 |
| GRIK4 | 2.20263 | 1.837391 | 18.34555 | 0.000986 | 0.018404 |
| PCDH20 | 1.508858 | 4.048885 | 18.31995 | 0.000992 | 0.018468 |
| HOXB5 | 1.555261 | 3.676396 | 18.32183 | 0.000991 | 0.018468 |
| LOC108591225 | 1.349968 | 4.139372 | 18.31403 | 0.000993 | 0.018474 |
| PIK3IP1 | 1.350345 | 4.218252 | 18.30057 | 0.000996 | 0.018508 |
| CHL1 | 1.199103 | 4.750821 | 18.28946 | 0.000998 | 0.018534 |
| DES | 1.434751 | 10.71938 | 18.26713 | 0.001003 | 0.018587 |
| SEC16B | 1.361348 | 8.046979 | 18.24787 | 0.001007 | 0.018627 |
| FAM151A | 1.382029 | 6.076129 | 18.22481 | 0.001012 | 0.0187 |
| LOC108588886 | 2.779621 | 1.308089 | 18.21398 | 0.001014 | 0.018708 |
| LOC103787495 | 1.241013 | 4.782015 | 18.19327 | 0.001019 | 0.018755 |
| KCNA6 | 1.633833 | 2.879443 | 18.17397 | 0.001023 | 0.018796 |
| LOC103789439 | 1.685164 | 3.597901 | 18.17762 | 0.001022 | 0.018796 |
| LOC100412233 | 1.392655 | 3.771546 | 18.06294 | 0.001047 | 0.019119 |
| NOS1 | 1.89661 | 3.32573 | 18.10233 | 0.001049 | 0.01913 |
| LGMN | 1.087987 | 6.8425 | 17.99761 | 0.001062 | 0.019314 |
| LOC108592094 | 3.395768 | 0.527198 | 17.9897 | 0.001064 | 0.019329 |
| ABCG5 | 1.326479 | 7.296558 | 17.96447 | 0.001069 | 0.019415 |
| NPR1 | 1.355261 | 4.894796 | 17.92067 | 0.00108 | 0.019579 |
| NHS | 1.238991 | 4.285885 | 17.88004 | 0.001089 | 0.019687 |
| EEPD1 | 1.389065 | 5.349128 | 17.86046 | 0.001094 | 0.019724 |
| OSBPL1A | 1.096815 | 7.04426 | 17.8214 | 0.001103 | 0.019798 |
| FILIP1 | 1.280615 | 5.876059 | 17.82204 | 0.001103 | 0.019798 |
| FLNC | 1.406389 | 8.493528 | 17.8317 | 0.0011 | 0.019798 |
| RNF208 | 1.546734 | 5.024565 | 17.9699 | 0.001103 | 0.019798 |
| FAM46B | 1.300964 | 3.876386 | 17.8086 | 0.001106 | 0.019831 |
| ASTN1 | 1.948546 | 2.000321 | 17.80147 | 0.001107 | 0.019831 |
| LOC108591218 | 2.557494 | 1.105607 | 17.72762 | 0.001125 | 0.020061 |

| | | | | | |
|---|---|---|---|---|---|
| EDN1 | 1.939801 | 2.686339 | 17.70187 | 0.001131 | 0.020092 |
| PTN | 1.883294 | 3.698074 | 17.96415 | 0.001134 | 0.020126 |
| AGAP1 | 1.351269 | 7.157396 | 17.66412 | 0.001141 | 0.020136 |
| TAGLN | 1.391415 | 9.161262 | 17.67054 | 0.001139 | 0.020136 |
| ABLIM3 | 1.471433 | 4.183606 | 17.67549 | 0.001138 | 0.020136 |
| NCAM2 | 1.901122 | 2.661843 | 17.67303 | 0.001138 | 0.020136 |
| PCK1 | 2.60912 | 8.715832 | 22.51255 | 0.001141 | 0.020136 |
| ACSL6 | 1.088165 | 5.786993 | 17.62391 | 0.001151 | 0.020233 |
| BICC1 | 1.324068 | 3.962186 | 17.61578 | 0.001153 | 0.020233 |
| CCDC136 | 1.334273 | 4.838816 | 17.6142 | 0.001153 | 0.020233 |
| SGIP1 | 1.283466 | 4.229524 | 17.59536 | 0.001158 | 0.020283 |
| CDON | 1.138856 | 6.410407 | 17.56788 | 0.001165 | 0.020367 |
| LOC108589003 | 4.367694 | 0.568574 | 17.54701 | 0.00117 | 0.020441 |
| HOXB3 | 1.178447 | 4.483507 | 17.50422 | 0.001181 | 0.020595 |
| NR2F2 | 1.122876 | 5.587714 | 17.45955 | 0.001192 | 0.02074 |
| DLG2 | 1.526478 | 4.003466 | 17.44999 | 0.001195 | 0.02075 |
| EFR3B | 1.928421 | 2.286965 | 17.43559 | 0.001199 | 0.020793 |
| LOC103788365 | 1.437473 | 3.574447 | 17.41643 | 0.001204 | 0.020841 |
| TSPAN2 | 1.524902 | 3.684948 | 17.41954 | 0.001203 | 0.020841 |
| FEZ1 | 1.323995 | 4.053959 | 17.40517 | 0.001206 | 0.020857 |
| DCC | 1.469715 | 3.595645 | 17.39445 | 0.001209 | 0.020887 |
| TSPAN12 | 1.414243 | 4.005968 | 17.3879 | 0.001211 | 0.020899 |
| CHADL | 2.79749 | 3.987732 | 21.81758 | 0.001219 | 0.021026 |
| ADD2 | 1.929328 | 2.701744 | 17.30766 | 0.001232 | 0.021193 |
| LOC100385629 | 1.165896 | 6.270452 | 17.30139 | 0.001234 | 0.021203 |
| PIWIL2 | 1.484073 | 3.212179 | 17.27531 | 0.001241 | 0.021287 |
| LOC108593153 | 1.278316 | 4.635729 | 17.26166 | 0.001245 | 0.021332 |
| LOC103793368 | 1.445376 | 5.693138 | 17.33457 | 0.001249 | 0.021384 |
| SLC38A8 | 2.657331 | 1.740943 | 17.22487 | 0.001255 | 0.02143 |
| MFAP4 | 2.266277 | 6.092836 | 21.36023 | 0.001258 | 0.021449 |
| ARMC4 | 2.670507 | 0.962135 | 17.17277 | 0.001269 | 0.021563 |
| LGI3 | 1.96997 | 2.145403 | 17.16572 | 0.001271 | 0.021578 |
| TM4SF4 | 1.877666 | 2.829175 | 17.15936 | 0.001273 | 0.021589 |
| SLC16A1 | 1.246438 | 5.602597 | 17.14115 | 0.001278 | 0.021613 |
| TRHDE | 2.958824 | 0.942634 | 17.13849 | 0.001279 | 0.021613 |
| LIX1 | 3.797117 | 0.787978 | 17.12084 | 0.001284 | 0.021659 |
| CNTN1 | 1.209032 | 4.549709 | 17.10808 | 0.001288 | 0.021683 |
| PCTP | 1.570718 | 2.991168 | 17.08527 | 0.001294 | 0.021755 |
| DKK3 | 1.154108 | 6.188568 | 17.04907 | 0.001304 | 0.021765 |
| PRELP | 1.242941 | 5.542963 | 17.05531 | 0.001303 | 0.021765 |
| PTRF | 1.30089 | 7.221683 | 17.04842 | 0.001305 | 0.021765 |
| FHOD3 | 1.328014 | 4.243101 | 17.07216 | 0.001298 | 0.021765 |
| KCNE4 | 1.551137 | 2.981545 | 17.03376 | 0.001309 | 0.021765 |
| LRP1B | 1.989259 | 2.378094 | 17.03446 | 0.001309 | 0.021765 |
| CLDN1 | 2.270151 | 2.587483 | 17.14165 | 0.001303 | 0.021765 |
| PTPRR | 2.273033 | 1.91063 | 17.03351 | 0.001309 | 0.021765 |
| LOC108590882 | 2.667931 | 1.340263 | 17.04411 | 0.001306 | 0.021765 |
| TACC2 | 1.090367 | 7.567591 | 17.00486 | 0.001317 | 0.02186 |

| | | | | | |
|---|---|---|---|---|---|
| PC | 1.112399 | 6.196991 | 17.00741 | 0.001316 | 0.02186 |
| KIAA0408 | 1.648297 | 5.027842 | 17.66993 | 0.001318 | 0.02186 |
| ZNF135 | 1.449839 | 3.814775 | 16.97942 | 0.001325 | 0.02193 |
| TUBA8 | 2.431749 | 1.180437 | 16.95872 | 0.001331 | 0.022012 |
| LOC103791753 | 1.442496 | 4.054253 | 16.94726 | 0.001334 | 0.02203 |
| ATP8A2 | 1.542481 | 3.28679 | 16.94999 | 0.001333 | 0.02203 |
| CHP2 | 1.30528 | 6.984536 | 16.92156 | 0.001342 | 0.022106 |
| SYT12 | 2.037653 | 4.999074 | 19.60972 | 0.001342 | 0.022106 |
| MXRA8 | 1.185723 | 5.799297 | 16.91026 | 0.001345 | 0.022137 |
| MEIS2 | 1.273284 | 5.893446 | 16.89071 | 0.001351 | 0.022189 |
| ZNF488 | 1.341055 | 4.096359 | 16.86778 | 0.001358 | 0.022189 |
| MMP16 | 2.807667 | 0.826971 | 16.87194 | 0.001357 | 0.022189 |
| KLHL26 | 1.542464 | 2.905394 | 16.85339 | 0.001362 | 0.022231 |
| EPHX1 | 1.043478 | 6.667529 | 16.83387 | 0.001368 | 0.022298 |
| TVP23B | 1.239252 | 5.591896 | 16.83228 | 0.001369 | 0.022298 |
| GNAO1 | 1.106409 | 4.874803 | 16.79879 | 0.001379 | 0.02242 |
| SMARCA1 | 1.111441 | 5.309522 | 16.78277 | 0.001384 | 0.022444 |
| DOCK9 | 1.20203 | 7.469649 | 16.7805 | 0.001384 | 0.022444 |
| SLITRK6 | 1.492101 | 4.187614 | 16.78762 | 0.001382 | 0.022444 |
| IMMP2L | 1.301655 | 3.570826 | 16.73753 | 0.001398 | 0.022623 |
| LOC108589279 | 1.778462 | 2.425137 | 16.72431 | 0.001402 | 0.022652 |
| MGAM | 1.303573 | 9.066416 | 16.69412 | 0.001411 | 0.022717 |
| TMEM63C | 2.206205 | 1.549861 | 16.69939 | 0.00141 | 0.022717 |
| LOC103793230 | 3.762427 | 0.760648 | 16.681 | 0.001415 | 0.022731 |
| ADCYAP1 | 3.230298 | 0.848911 | 16.65408 | 0.001424 | 0.022809 |
| PFKFB1 | 2.100871 | 2.010001 | 16.63935 | 0.001429 | 0.02286 |
| ITGA9 | 1.224219 | 6.742287 | 16.62208 | 0.001434 | 0.022888 |
| HIF3A | 1.612137 | 3.640221 | 16.6206 | 0.001435 | 0.022888 |
| LOC103794633 | 1.579292 | 4.079676 | 16.65438 | 0.001439 | 0.022929 |
| PPFIA3 | 1.181761 | 6.656358 | 16.57798 | 0.001449 | 0.023003 |
| RASGRF2 | 1.452072 | 3.306453 | 16.58287 | 0.001447 | 0.023003 |
| TNNI2 | 1.617419 | 7.22376 | 17.48529 | 0.001448 | 0.023003 |
| LOC100392932 | 2.606727 | 1.143938 | 16.57515 | 0.001449 | 0.023003 |
| GNG8 | 2.047279 | 1.659279 | 16.51322 | 0.00147 | 0.023287 |
| PTPRT | 2.440019 | 1.454532 | 16.51441 | 0.001469 | 0.023287 |
| LOC103796068 | 1.197656 | 4.299038 | 16.50653 | 0.001472 | 0.023293 |
| PPP1R12B | 1.253611 | 9.1158 | 16.50159 | 0.001474 | 0.023293 |
| DIXDC1 | 1.279175 | 6.004847 | 16.48818 | 0.001478 | 0.023343 |
| SLC35F5 | 1.055961 | 6.868488 | 16.47391 | 0.001483 | 0.023399 |
| STXBP1 | 1.151974 | 4.716624 | 16.46838 | 0.001485 | 0.023402 |
| LOC100896436 | 1.336814 | 4.385634 | 16.46305 | 0.001486 | 0.023402 |
| GPC3 | 1.585706 | 3.805072 | 16.45905 | 0.001488 | 0.023402 |
| LOC100894745 | 1.135654 | 4.516425 | 16.45161 | 0.00149 | 0.023421 |
| ME1 | 1.253377 | 5.021231 | 16.44485 | 0.001493 | 0.023421 |
| NPTXR | 1.419544 | 3.308117 | 16.44579 | 0.001492 | 0.023421 |
| HGD | 2.418762 | 1.667881 | 16.42451 | 0.001499 | 0.023472 |
| DTNA | 1.333537 | 5.063875 | 16.395 | 0.001509 | 0.02361 |
| MPP2 | 1.858156 | 2.346113 | 16.37763 | 0.001515 | 0.023653 |

| | | | | | |
|---|---|---|---|---|---|
| PER1 | 2.035422 | 6.963937 | 19.59401 | 0.001514 | 0.023653 |
| CD109 | 1.398049 | 4.983882 | 16.37357 | 0.001519 | 0.023684 |
| LAMB2 | 1.136796 | 7.018489 | 16.34442 | 0.001527 | 0.023768 |
| PTPRZ1 | 1.428926 | 4.017084 | 16.3468 | 0.001526 | 0.023768 |
| DFNA5 | 1.141773 | 7.185531 | 16.31679 | 0.001536 | 0.023784 |
| LOC108592791 | 1.175711 | 4.588004 | 16.32463 | 0.001534 | 0.023784 |
| EPHA5 | 1.500327 | 2.927882 | 16.32215 | 0.001535 | 0.023784 |
| CBLN3 | 1.980159 | 6.089563 | 19.26792 | 0.001529 | 0.023784 |
| STXBP6 | 1.278924 | 3.921095 | 16.28263 | 0.001548 | 0.023893 |
| NAALADL1 | 2.407509 | 10.3356 | 19.472 | 0.001548 | 0.023893 |
| FAXC | 1.365845 | 3.613279 | 16.26836 | 0.001553 | 0.023933 |
| LOC103788847 | 2.003067 | 2.134969 | 16.26065 | 0.001556 | 0.023937 |
| FIGN | 1.452674 | 2.908459 | 16.23279 | 0.001566 | 0.023995 |
| LOC100393583 | 1.477883 | 3.190362 | 16.21519 | 0.001572 | 0.024001 |
| ZNF214 | 1.88672 | 2.198117 | 16.21688 | 0.001572 | 0.024001 |
| ZNF462 | 1.137047 | 6.276094 | 16.19259 | 0.00158 | 0.024046 |
| SSC5D | 1.407421 | 4.502339 | 16.20105 | 0.001577 | 0.024046 |
| SYT14 | 2.020403 | 3.224453 | 16.54657 | 0.001579 | 0.024046 |
| PKP1 | 4.295196 | 0.513616 | 16.19256 | 0.00158 | 0.024046 |
| LOC108590110 | 3.787698 | 0.77976 | 16.17978 | 0.001585 | 0.024091 |
| NKX2-3 | 1.086758 | 5.831548 | 16.17092 | 0.001588 | 0.024101 |
| AADAT | 1.27489 | 4.650726 | 16.15966 | 0.001592 | 0.024105 |
| SOAT2 | 1.148888 | 7.541421 | 16.14445 | 0.001598 | 0.024161 |
| B4GAT1 | 1.295695 | 3.497397 | 16.11167 | 0.00161 | 0.024323 |
| LOC103789105 | 2.610885 | 1.175403 | 16.07281 | 0.001624 | 0.024483 |
| NAP1L3 | 1.819155 | 2.334238 | 16.04729 | 0.001634 | 0.024589 |
| PLCB1 | 1.304606 | 5.284576 | 16.02485 | 0.001642 | 0.024677 |
| RGL3 | 2.353821 | 1.975391 | 16.02115 | 0.001643 | 0.024679 |
| ARMC9 | 1.252098 | 5.089064 | 16.01536 | 0.001646 | 0.024693 |
| LOC103788361 | 1.750045 | 2.309904 | 15.99501 | 0.001653 | 0.024771 |
| LEXM | 1.259465 | 6.020851 | 15.98434 | 0.001657 | 0.024812 |
| MAGI2 | 1.676247 | 3.187616 | 15.96 | 0.001667 | 0.024932 |
| NID2 | 1.231033 | 4.024101 | 15.92505 | 0.00168 | 0.025057 |
| MFSD7 | 1.399514 | 4.729537 | 15.92691 | 0.001679 | 0.025057 |
| PTPRN2 | 1.688408 | 3.505286 | 15.91243 | 0.001685 | 0.025074 |
| MYOCD | 1.224238 | 6.01488 | 15.90802 | 0.001687 | 0.025078 |
| MYH15 | 1.32916 | 4.855625 | 15.90025 | 0.00169 | 0.025104 |
| LOC103795270 | 1.039493 | 5.649999 | 15.8946 | 0.001692 | 0.025114 |
| LOC103792776 | 2.686326 | 1.178484 | 15.88919 | 0.001694 | 0.025114 |
| WFS1 | 1.022185 | 6.1135 | 15.87311 | 0.0017 | 0.025185 |
| SH3TC2 | 2.364715 | 1.752762 | 15.85339 | 0.001708 | 0.025281 |
| LOC108593386 | 1.346198 | 4.847024 | 15.83896 | 0.001714 | 0.025325 |
| BMPR1B | 2.419409 | 2.438113 | 16.19519 | 0.001715 | 0.025325 |
| SLC27A2 | 1.089866 | 7.04739 | 15.80936 | 0.001725 | 0.025459 |
| IL11RA | 1.130551 | 4.35704 | 15.78496 | 0.001735 | 0.025548 |
| HECTD2 | 1.49503 | 4.018156 | 15.76406 | 0.001744 | 0.025633 |
| BGN | 1.596501 | 3.40911 | 15.74118 | 0.001753 | 0.02575 |
| SLC6A19 | 1.289822 | 10.25001 | 15.71543 | 0.001763 | 0.025865 |

| | | | | | |
|---|---|---|---|---|---|
| LOC108592281 | 1.333236 | 3.657242 | 15.70611 | 0.001767 | 0.025902 |
| LRP3 | 1.443354 | 3.681068 | 15.66989 | 0.001782 | 0.026082 |
| SLC5A9 | 1.2296 | 7.22498 | 15.64969 | 0.001791 | 0.026184 |
| SCN3A | 1.823746 | 2.297914 | 15.64072 | 0.001794 | 0.026219 |
| ACSF2 | 1.209067 | 8.078822 | 15.63604 | 0.001796 | 0.026226 |
| POPDC2 | 1.521413 | 3.914829 | 15.63319 | 0.001797 | 0.026226 |
| DCHS2 | 1.689036 | 2.867926 | 15.61544 | 0.001805 | 0.026295 |
| 12-Sep | 1.835114 | 3.117745 | 15.61839 | 0.001821 | 0.026496 |
| LOC100393377 | 3.527847 | 2.170665 | 18.20044 | 0.00182 | 0.026496 |
| ENTPD3 | 1.954021 | 1.964694 | 15.56293 | 0.001827 | 0.02651 |
| LOC100402656 | 3.782656 | 0.871408 | 15.62356 | 0.001828 | 0.02651 |
| SULT1B1 | 1.337767 | 6.159726 | 15.60679 | 0.001835 | 0.026537 |
| LOC108590768 | 2.501702 | 1.412586 | 15.55049 | 0.001832 | 0.026537 |
| LOC100896973 | 6.005384 | 2.034399 | 21.85114 | 0.00184 | 0.026584 |
| FAM188B | 1.341963 | 4.858681 | 15.5156 | 0.001847 | 0.026656 |
| DDIT4L | 3.007223 | 2.807464 | 18.37179 | 0.001846 | 0.026656 |
| MICAL3 | 1.094647 | 7.503257 | 15.50908 | 0.00185 | 0.026677 |
| CDKN1C | 1.577134 | 3.862959 | 15.51935 | 0.001864 | 0.026814 |
| CNR1 | 1.641172 | 3.016923 | 15.48235 | 0.001862 | 0.026814 |
| LOC103788721 | 3.580682 | 1.045801 | 15.55412 | 0.001863 | 0.026814 |
| CHRM2 | 1.558753 | 4.851163 | 15.91052 | 0.001878 | 0.026985 |
| NIPAL4 | 1.382035 | 3.397507 | 15.43909 | 0.001881 | 0.026998 |
| CRISPLD1 | 1.618288 | 4.745681 | 16.06531 | 0.001883 | 0.027015 |
| PLPPR4 | 2.829928 | 2.817 | 17.74933 | 0.001892 | 0.027127 |
| LOC103793578 | 2.070325 | 1.796713 | 15.38764 | 0.001904 | 0.027215 |
| CALD1 | 1.20401 | 9.817335 | 15.3767 | 0.001908 | 0.02723 |
| LOC103795086 | 2.805037 | 1.033663 | 15.36857 | 0.001912 | 0.02723 |
| FCN2 | 4.547091 | 5.228694 | 28.45553 | 0.001914 | 0.02723 |
| NRXN1 | 1.36474 | 3.227842 | 15.35117 | 0.00192 | 0.027281 |
| RAB6B | 1.306368 | 3.611576 | 15.34066 | 0.001925 | 0.027309 |
| PPP1R3C | 1.294565 | 3.964512 | 15.33094 | 0.001929 | 0.027351 |
| LOC100400012 | 4.0246 | 2.027227 | 18.58596 | 0.001933 | 0.027393 |
| LOC100415560 | 1.61197 | 2.721492 | 15.26876 | 0.001958 | 0.027604 |
| CSMD2 | 1.891102 | 3.572871 | 15.78434 | 0.001958 | 0.027604 |
| HOXA3 | 1.073297 | 5.141845 | 15.25676 | 0.001963 | 0.027653 |
| GPAT3 | 1.238753 | 7.707853 | 15.25337 | 0.001965 | 0.027655 |
| FHL1 | 1.212847 | 7.462929 | 15.23957 | 0.001971 | 0.027726 |
| LOC108594175 | 1.582271 | 4.431871 | 15.4895 | 0.002001 | 0.02802 |
| LOC100403206 | 2.116165 | 1.368636 | 15.17026 | 0.002004 | 0.028042 |
| LOC100412437 | 1.110165 | 6.530962 | 15.16315 | 0.002007 | 0.028046 |
| ZNF185 | 1.591429 | 4.047278 | 15.27286 | 0.002008 | 0.028046 |
| CNKSR2 | 2.00877 | 1.784437 | 15.1528 | 0.002012 | 0.028078 |
| LRRN3 | 1.299836 | 3.57448 | 15.142 | 0.002017 | 0.02813 |
| TMEFF2 | 2.146513 | 1.5028 | 15.12209 | 0.002027 | 0.028238 |
| LOC100406601 | 1.087355 | 6.78207 | 15.1128 | 0.002031 | 0.028265 |
| LRIG1 | 1.105071 | 6.149465 | 15.08925 | 0.002043 | 0.028403 |
| GFRA1 | 1.147762 | 5.492383 | 15.07093 | 0.002051 | 0.028507 |
| PPP1R3B | 1.414406 | 3.531908 | 15.06405 | 0.002055 | 0.028513 |

| | | | | | |
|---|---|---|---|---|---|
| ADRA2B | 2.420201 | 2.024978 | 15.10096 | 0.002067 | 0.028643 |
| DYRK1B | 1.221779 | 6.362779 | 14.99946 | 0.002087 | 0.028812 |
| LOC103795612 | 3.476064 | 5.783006 | 24.21062 | 0.002086 | 0.028812 |
| NRCAM | 1.538191 | 2.668841 | 14.9704 | 0.002101 | 0.028971 |
| LRRTM4 | 2.383027 | 1.317769 | 14.96436 | 0.002104 | 0.028993 |
| PYGB | 1.083497 | 6.889024 | 14.92454 | 0.002124 | 0.029131 |
| PHLDB2 | 1.087863 | 6.247267 | 14.93129 | 0.002121 | 0.029131 |
| SLC28A1 | 1.108552 | 6.238531 | 14.93411 | 0.002119 | 0.029131 |
| LOC103793609 | 1.446491 | 3.797552 | 14.92345 | 0.002125 | 0.029131 |
| MYOM2 | 1.607839 | 2.958937 | 14.92097 | 0.002126 | 0.029131 |
| FHL5 | 1.849471 | 1.767906 | 14.9254 | 0.002124 | 0.029131 |
| ATP1A3 | 1.75641 | 3.022421 | 14.84667 | 0.002164 | 0.029571 |
| SGCD | 1.491824 | 3.884919 | 14.8383 | 0.002169 | 0.029609 |
| KY | 2.420946 | 1.001571 | 14.80954 | 0.002184 | 0.029794 |
| PXDN | 1.111654 | 6.300831 | 14.79066 | 0.002194 | 0.029908 |
| TENM3 | 1.033608 | 4.839922 | 14.77805 | 0.0022 | 0.029978 |
| OGN | 2.299299 | 5.01853 | 18.54339 | 0.002205 | 0.030003 |
| DIRAS2 | 1.98536 | 1.611632 | 14.75324 | 0.002213 | 0.030095 |
| LOC100398578 | 1.314065 | 3.287284 | 14.74225 | 0.002219 | 0.030133 |
| HOXB4 | 1.101323 | 5.027544 | 14.73675 | 0.002222 | 0.030138 |
| SLC23A1 | 1.013595 | 6.501616 | 14.73263 | 0.002224 | 0.03014 |
| SLITRK1 | 3.223267 | 0.842744 | 14.7236 | 0.002229 | 0.030164 |
| CD36 | 1.30385 | 5.873838 | 14.77674 | 0.002231 | 0.030165 |
| VIP | 1.764996 | 3.084841 | 14.71424 | 0.002234 | 0.030169 |
| TM6SF2 | 1.080388 | 7.171162 | 14.70316 | 0.00224 | 0.030229 |
| MAOB | 0.960991 | 6.87381 | 14.68216 | 0.002252 | 0.03032 |
| HMP19 | 2.250937 | 1.513933 | 14.62907 | 0.002281 | 0.030648 |
| FAM19A5 | 2.310323 | 1.671028 | 14.60056 | 0.002297 | 0.030797 |
| PCDH15 | 2.702096 | 1.21726 | 14.54274 | 0.002329 | 0.031103 |
| BCAS1 | 1.137389 | 8.215971 | 14.5233 | 0.00234 | 0.031187 |
| LIN7B | 1.460405 | 4.005557 | 14.52465 | 0.002339 | 0.031187 |
| HSPG2 | 1.232261 | 9.266484 | 14.5187 | 0.002343 | 0.0312 |
| KIAA1217 | 1.153859 | 6.822787 | 14.50281 | 0.002352 | 0.031278 |
| ZMYND15 | 1.027741 | 4.727303 | 14.45558 | 0.002379 | 0.031521 |
| KDM4B | 1.082677 | 6.834459 | 14.39935 | 0.002412 | 0.031796 |
| CHD5 | 1.507683 | 2.787229 | 14.40063 | 0.002411 | 0.031796 |
| C10H14orf180 | 3.530181 | 1.330363 | 15.11615 | 0.002411 | 0.031796 |
| ACVR2B | 1.17679 | 3.849835 | 14.37424 | 0.002427 | 0.031927 |
| ARHGEF28 | 1.335699 | 4.38786 | 14.34947 | 0.002441 | 0.032056 |
| ADGRL3 | 1.089693 | 4.791753 | 14.34242 | 0.002445 | 0.03209 |
| PYGO1 | 1.597381 | 3.780318 | 14.39528 | 0.002453 | 0.032146 |
| NHSL1 | 1.40504 | 7.982821 | 14.534 | 0.002458 | 0.032174 |
| ELOVL4 | 2.617206 | 0.952245 | 14.32163 | 0.002458 | 0.032174 |
| SCN11A | 2.55629 | 1.916931 | 14.51452 | 0.00247 | 0.03225 |
| ZDHHC19 | 2.879388 | 3.10052 | 17.26997 | 0.002471 | 0.03225 |
| GNAI1 | 1.202347 | 4.501577 | 14.29593 | 0.002473 | 0.032263 |
| STARD9 | 1.021247 | 6.835384 | 14.28234 | 0.002482 | 0.032327 |
| CHAD | 1.437262 | 6.981451 | 14.77142 | 0.002486 | 0.032368 |

| | | | | | |
|---|---|---|---|---|---|
| SYDE2 | 1.155982 | 4.709869 | 14.23455 | 0.002511 | 0.032557 |
| LRP12 | 1.515428 | 3.840328 | 14.26902 | 0.002514 | 0.032559 |
| LOC100403069 | 1.266895 | 4.105536 | 14.19495 | 0.002536 | 0.032706 |
| CACNA2D1 | 1.293329 | 5.876465 | 14.25556 | 0.002538 | 0.032706 |
| LOC108590153 | 2.925774 | 0.618635 | 14.1984 | 0.002533 | 0.032706 |
| LOC103795378 | 1.778442 | 1.786143 | 14.18193 | 0.002544 | 0.032749 |
| XPNPEP2 | 1.280773 | 9.589463 | 14.15782 | 0.002559 | 0.032881 |
| MEOX2 | 1.929579 | 1.800513 | 14.15683 | 0.00256 | 0.032881 |
| DNAH6 | 1.551768 | 2.561354 | 14.14991 | 0.002564 | 0.032915 |
| JPH1 | 1.152056 | 4.272484 | 14.13842 | 0.002571 | 0.032944 |
| TBCEL | 1.332266 | 3.723981 | 14.12224 | 0.002582 | 0.033054 |
| NID1 | 0.968058 | 7.791328 | 14.11921 | 0.002583 | 0.033057 |
| CRY2 | 1.295655 | 5.902357 | 14.17576 | 0.002588 | 0.033095 |
| SYT8 | 1.645334 | 8.60335 | 15.05956 | 0.002597 | 0.033165 |
| LOC108588061 | 3.863405 | 0.935656 | 14.37374 | 0.002604 | 0.033165 |
| HSPB8 | 1.029202 | 5.200528 | 14.07722 | 0.00261 | 0.033224 |
| HS6ST3 | 1.391825 | 3.456944 | 14.07033 | 0.002615 | 0.033242 |
| FMO1 | 3.557694 | 1.029442 | 14.28472 | 0.002622 | 0.033289 |
| SLC29A1 | 1.057865 | 5.025602 | 14.04399 | 0.002632 | 0.033378 |
| PALLD | 1.090504 | 7.903901 | 14.03063 | 0.002641 | 0.033378 |
| CACNA1B | 1.756764 | 2.682291 | 14.04303 | 0.002633 | 0.033378 |
| LOC108588063 | 1.859261 | 2.928334 | 14.08344 | 0.002642 | 0.033378 |
| LOC103793410 | 2.659828 | 0.983909 | 14.02173 | 0.002647 | 0.033405 |
| CTNNA2 | 1.733809 | 2.175885 | 14.01184 | 0.002653 | 0.033457 |
| TRPM4 | 1.65882 | 9.172672 | 14.86039 | 0.002665 | 0.033515 |
| LOC103789795 | 1.039343 | 5.79726 | 13.96231 | 0.002686 | 0.033728 |
| PAM | 1.049526 | 7.402705 | 13.94864 | 0.002695 | 0.033809 |
| LRFN4 | 1.315062 | 3.472049 | 13.94732 | 0.002696 | 0.033809 |
| FAM149A | 1.063149 | 6.010252 | 13.9114 | 0.00272 | 0.033918 |
| LOC103792661 | 3.00274 | 0.97735 | 13.89301 | 0.002733 | 0.034052 |
| RUSC2 | 1.114963 | 6.708552 | 13.8878 | 0.002736 | 0.034075 |
| CNTN6 | 2.362693 | 1.414644 | 13.85862 | 0.002756 | 0.034237 |
| SLC16A2 | 1.457974 | 3.69928 | 13.84292 | 0.002767 | 0.034326 |
| RASGEF1C | 2.935428 | 0.625582 | 13.83921 | 0.00277 | 0.034326 |
| GPBAR1 | 1.903002 | 1.93699 | 13.83515 | 0.002772 | 0.034329 |
| ZFPM2 | 1.190418 | 4.260695 | 13.80205 | 0.002795 | 0.034592 |
| LOC103793036 | 2.207055 | 1.18382 | 13.79837 | 0.002798 | 0.034602 |
| DAAM2 | 0.938354 | 5.672788 | 13.79214 | 0.002802 | 0.034612 |
| CACNB2 | 0.962036 | 5.695479 | 13.7876 | 0.002806 | 0.03463 |
| PRUNE2 | 1.144907 | 8.152321 | 13.76593 | 0.002821 | 0.034775 |
| CTSF | 1.256313 | 4.173333 | 13.73884 | 0.00284 | 0.034906 |
| TTLL11 | 1.359291 | 3.357272 | 13.70067 | 0.002867 | 0.035171 |
| PTPN14 | 1.258913 | 5.470787 | 13.74557 | 0.002871 | 0.035197 |
| LOC100396974 | 2.429359 | 1.922481 | 13.79734 | 0.002873 | 0.0352 |
| EPS8L3 | 1.125664 | 8.987972 | 13.67512 | 0.002886 | 0.035219 |
| TPM2 | 1.185959 | 9.423247 | 13.67438 | 0.002886 | 0.035219 |
| OLFM2 | 1.273122 | 3.476581 | 13.66754 | 0.002891 | 0.035219 |
| DNAL1 | 1.320467 | 5.085955 | 13.74649 | 0.002878 | 0.035219 |

| | | | | | |
|---|---|---|---|---|---|
| COL21A1 | 1.684996 | 2.423574 | 13.68275 | 0.00288 | 0.035219 |
| KCNA2 | 2.169125 | 1.520738 | 13.67426 | 0.002886 | 0.035219 |
| LOC103793414 | 4.624846 | 2.4471 | 18.9483 | 0.002893 | 0.035219 |
| LOC108590900 | 1.221274 | 3.914421 | 13.65715 | 0.002899 | 0.035232 |
| IFITM10 | 2.361973 | 2.818976 | 14.79266 | 0.002906 | 0.035284 |
| ARHGEF17 | 1.094329 | 6.488591 | 13.63696 | 0.002914 | 0.035341 |
| TMEM253 | 0.98053 | 5.938558 | 13.6251 | 0.002922 | 0.035416 |
| LOC100407224 | 1.051059 | 5.854575 | 13.61623 | 0.002929 | 0.03546 |
| PEBP4 | 2.44159 | 0.989313 | 13.60799 | 0.002935 | 0.035512 |
| LOC100415308 | 1.015722 | 5.621419 | 13.6014 | 0.00294 | 0.035549 |
| LOC108588517 | 1.708672 | 2.377918 | 13.59429 | 0.002945 | 0.035591 |
| ACSS3 | 1.359447 | 3.950998 | 13.5802 | 0.002956 | 0.035696 |
| ABO | 0.954943 | 7.14437 | 13.56018 | 0.002971 | 0.035805 |
| CDR1 | 1.295677 | 8.194894 | 13.60776 | 0.002972 | 0.035805 |
| PDGFD | 1.832598 | 2.659593 | 13.56182 | 0.00297 | 0.035805 |
| LOC103794065 | 2.73799 | 1.527962 | 13.65181 | 0.002977 | 0.035844 |
| LOC100394452 | 4.954878 | 1.869977 | 17.60662 | 0.002992 | 0.036001 |
| NEDD4L | 0.987941 | 7.638226 | 13.52313 | 0.002999 | 0.036059 |
| CNNM1 | 1.20721 | 3.676568 | 13.47079 | 0.003039 | 0.036346 |
| PRKD1 | 1.227046 | 3.498356 | 13.46521 | 0.003043 | 0.036346 |
| TSPO2 | 1.342058 | 4.262447 | 13.46662 | 0.003042 | 0.036346 |
| GALNT16 | 1.376087 | 3.672598 | 13.47211 | 0.003038 | 0.036346 |
| LOC100415297 | 0.970683 | 5.7814 | 13.45714 | 0.003049 | 0.036354 |
| LOC103795688 | 1.275006 | 4.054755 | 13.45758 | 0.003049 | 0.036354 |
| LOC103789410 | 2.918491 | 0.946755 | 13.45804 | 0.003049 | 0.036354 |
| L1CAM | 1.199001 | 4.678643 | 13.45223 | 0.003053 | 0.036377 |
| KIF12 | 2.541279 | 2.384561 | 14.37007 | 0.003056 | 0.03638 |
| CDK5R1 | 1.413455 | 2.942093 | 13.44102 | 0.003062 | 0.036414 |
| FAM171B | 1.39775 | 3.138117 | 13.43715 | 0.003065 | 0.036427 |
| KCNA1 | 1.857286 | 3.107076 | 13.64966 | 0.003091 | 0.036692 |
| ADGRA2 | 1.163189 | 6.605089 | 13.35037 | 0.003133 | 0.036946 |
| GPM6A | 1.82081 | 1.95315 | 13.34966 | 0.003134 | 0.036946 |
| DMTN | 1.71612 | 1.820721 | 13.34554 | 0.003137 | 0.036951 |
| IDUA | 1.349237 | 6.635052 | 13.67196 | 0.00314 | 0.036966 |
| LOC103794023 | 1.181828 | 3.782618 | 13.33725 | 0.003144 | 0.036966 |
| NREP | 1.214229 | 4.147017 | 13.33687 | 0.003144 | 0.036966 |
| LOC108593182 | 1.64403 | 2.176865 | 13.31929 | 0.003158 | 0.037088 |
| HS6ST2 | 2.276278 | 1.236237 | 13.29087 | 0.003181 | 0.037291 |
| B4GALNT1 | 1.296511 | 7.940886 | 13.34425 | 0.003185 | 0.037308 |
| VSTM2A | 4.373297 | 2.573437 | 18.39434 | 0.00321 | 0.03754 |
| OSR1 | 1.530607 | 4.324695 | 13.58687 | 0.003252 | 0.037889 |
| LOC108590003 | 2.303545 | 1.793177 | 13.2185 | 0.003254 | 0.037889 |
| NRXN3 | 1.844278 | 2.477283 | 13.19397 | 0.003261 | 0.037895 |
| LARP6 | 1.382215 | 2.877571 | 13.18994 | 0.003265 | 0.037904 |
| ATOH7 | 3.403772 | 5.474296 | 21.22313 | 0.003289 | 0.038143 |
| LOC108591005 | 2.56018 | 1.40885 | 13.14294 | 0.003304 | 0.038297 |
| CLCN5 | 0.980826 | 6.113786 | 13.12729 | 0.003318 | 0.038429 |
| CCDC120 | 1.182875 | 6.994809 | 13.1175 | 0.003326 | 0.03848 |

| | | | | | |
|---|---|---|---|---|---|
| LOC103791201 | 1.04758 | 6.688004 | 13.10864 | 0.003334 | 0.038533 |
| LOC108590320 | 1.820375 | 3.716692 | 13.93176 | 0.003335 | 0.038533 |
| PLXDC2 | 1.015022 | 4.409433 | 13.09832 | 0.003343 | 0.038564 |
| PFN2 | 1.138825 | 4.548565 | 13.09364 | 0.003347 | 0.038564 |
| CCDC7 | 2.100593 | 2.478807 | 13.27932 | 0.003351 | 0.038564 |
| GRIK1 | 2.130081 | 1.279426 | 13.09273 | 0.003347 | 0.038564 |
| LOC103794490 | 1.275535 | 3.238775 | 13.07979 | 0.003359 | 0.038628 |
| RHOQ | 1.002459 | 5.010326 | 13.06889 | 0.003368 | 0.038714 |
| ARHGAP42 | 0.903559 | 6.349078 | 13.05789 | 0.003378 | 0.038774 |
| CES2 | 1.291115 | 7.76865 | 13.15004 | 0.003379 | 0.038774 |
| OSGIN1 | 1.294062 | 6.870553 | 13.24674 | 0.003378 | 0.038774 |
| CDC14A | 0.951822 | 6.612 | 13.04308 | 0.003391 | 0.038881 |
| NRP2 | 1.015737 | 7.301474 | 13.04087 | 0.003393 | 0.038881 |
| HOXC6 | 1.04509 | 4.369373 | 13.02803 | 0.003404 | 0.038964 |
| ABI3BP | 1.280002 | 6.697031 | 13.18761 | 0.003406 | 0.038972 |
| ST3GAL6 | 1.579327 | 2.86173 | 13.0096 | 0.00342 | 0.039105 |
| CILP | 1.061873 | 4.81277 | 13.00119 | 0.003428 | 0.039167 |
| PDGFC | 1.044096 | 4.295115 | 12.97738 | 0.003449 | 0.039386 |
| CADPS | 1.740325 | 2.784136 | 12.96018 | 0.003464 | 0.039516 |
| ADGRF5 | 1.080889 | 5.418031 | 12.93263 | 0.003489 | 0.039707 |
| LOC100896839 | 2.800656 | 1.324927 | 12.98416 | 0.003485 | 0.039707 |
| CCL24 | 2.410932 | 0.767962 | 12.92342 | 0.003497 | 0.039779 |
| SNX33 | 0.981101 | 6.922555 | 12.91952 | 0.003501 | 0.039789 |
| ABCG4 | 1.48369 | 2.680108 | 12.89693 | 0.003522 | 0.039915 |
| SEMA3E | 1.636945 | 4.917338 | 14.12928 | 0.003516 | 0.039915 |
| CERS1 | 2.491581 | 1.356242 | 12.89829 | 0.00352 | 0.039915 |
| ACSS1 | 0.994776 | 8.081407 | 12.87476 | 0.003542 | 0.040053 |
| EXOC6B | 0.982799 | 7.247197 | 12.84631 | 0.003568 | 0.040232 |
| ARHGAP6 | 1.104933 | 4.390756 | 12.84703 | 0.003568 | 0.040232 |
| LOC103796146 | 2.124191 | 1.859525 | 12.85045 | 0.003564 | 0.040232 |
| ADAMTS13 | 2.77072 | 2.959734 | 15.29551 | 0.003569 | 0.040232 |
| LOC108590291 | 2.43182 | 2.473409 | 13.72293 | 0.003591 | 0.040308 |
| RHOBTB2 | 1.034123 | 6.321757 | 12.80457 | 0.003607 | 0.040465 |
| MRO | 1.094858 | 5.073561 | 12.79955 | 0.003612 | 0.040481 |
| TYRP1 | 3.848909 | 1.602117 | 14.85612 | 0.003613 | 0.040481 |
| ZBTB47 | 1.22384 | 4.156591 | 12.79314 | 0.003618 | 0.040516 |
| GPSM1 | 1.842808 | 4.944135 | 14.84666 | 0.003634 | 0.040658 |
| GUCY1A3 | 1.081993 | 6.415709 | 12.76505 | 0.003645 | 0.040722 |
| MAP1A | 0.928078 | 6.633264 | 12.75649 | 0.003653 | 0.04079 |
| CNTN5 | 2.761929 | 1.095938 | 12.74024 | 0.003669 | 0.04094 |
| RADIL | 2.001922 | 2.615956 | 12.90308 | 0.003679 | 0.041038 |
| NLGN1 | 3.421516 | 3.533054 | 17.83566 | 0.00371 | 0.041238 |
| BCKDHA | 1.011261 | 4.722953 | 12.65721 | 0.003749 | 0.041508 |
| ITGBL1 | 2.547037 | 0.83714 | 12.659 | 0.003748 | 0.041508 |
| GPRC5B | 1.255125 | 4.079038 | 12.63273 | 0.003774 | 0.041708 |
| STRIP2 | 2.138318 | 1.446093 | 12.62246 | 0.003784 | 0.041774 |
| SLC20A2 | 1.007313 | 4.994762 | 12.6159 | 0.00379 | 0.041798 |
| DCLK2 | 1.274543 | 3.938816 | 12.61137 | 0.003795 | 0.041825 |

| | | | | | |
|---|---|---|---|---|---|
| LOC108591283 | 1.739613 | 2.349459 | 12.59601 | 0.00381 | 0.041923 |
| SLC26A3 | 1.482615 | 5.666219 | 13.46102 | 0.003853 | 0.042226 |
| SLC36A4 | 1.474697 | 2.919751 | 12.54777 | 0.003859 | 0.04227 |
| LOC100411741 | 0.931622 | 7.016987 | 12.53218 | 0.003875 | 0.042397 |
| ROR1 | 1.659189 | 2.579403 | 12.52884 | 0.003878 | 0.042411 |
| SLC51A | 0.990947 | 6.501227 | 12.51406 | 0.003894 | 0.042529 |
| FAM181A | 2.05615 | 1.971964 | 12.51486 | 0.003893 | 0.042529 |
| CUX2 | 1.684419 | 2.239006 | 12.50873 | 0.003899 | 0.042566 |
| PREX2 | 0.944794 | 5.839252 | 12.50325 | 0.003905 | 0.042603 |
| SLC25A4 | 1.020499 | 5.417459 | 12.4944 | 0.003914 | 0.042637 |
| CACNA2D3 | 1.710436 | 2.177627 | 12.49394 | 0.003914 | 0.042637 |
| GLP1R | 1.915709 | 2.026518 | 12.48619 | 0.003922 | 0.0427 |
| TPM1 | 1.111271 | 10.02299 | 12.47781 | 0.003931 | 0.042748 |
| RSAD1 | 1.595488 | 3.468083 | 12.55614 | 0.003943 | 0.042825 |
| LOC103787521 | 2.87217 | 4.158164 | 17.16156 | 0.003941 | 0.042825 |
| RNASE4 | 0.937143 | 5.890116 | 12.45426 | 0.003956 | 0.042932 |
| TRIP6 | 1.024041 | 5.472883 | 12.45076 | 0.003959 | 0.042933 |
| CBLN1 | 2.487723 | 0.9956 | 12.44341 | 0.003967 | 0.042973 |
| LOC100394345 | 1.634641 | 2.220096 | 12.43062 | 0.003981 | 0.043 |
| THSD4 | 1.20985 | 6.468237 | 12.46428 | 0.003999 | 0.043129 |
| CSAD | 1.80293 | 4.218227 | 13.79566 | 0.004015 | 0.043256 |
| ANK3 | 0.882666 | 6.358054 | 12.38194 | 0.004032 | 0.043392 |
| PNPLA2 | 1.105657 | 7.560494 | 12.37363 | 0.004041 | 0.043441 |
| LOC108588094 | 2.298312 | 1.053736 | 12.37353 | 0.004042 | 0.043441 |
| LOC103789058 | 1.682047 | 3.099173 | 12.41912 | 0.004047 | 0.043455 |
| IDO2 | 1.012419 | 6.095714 | 12.35179 | 0.004065 | 0.043603 |
| SDK1 | 1.032983 | 4.251674 | 12.34932 | 0.004068 | 0.043603 |
| LRRC10B | 2.509588 | 0.870392 | 12.33165 | 0.004087 | 0.043761 |
| KNDC1 | 1.943885 | 1.562079 | 12.32089 | 0.004099 | 0.043838 |
| FBN2 | 1.058955 | 3.923271 | 12.31359 | 0.004107 | 0.043874 |
| MAF | 1.137778 | 8.104986 | 12.30969 | 0.004111 | 0.043874 |
| RAB3B | 2.165343 | 1.365053 | 12.30665 | 0.004114 | 0.043885 |
| FLNA | 1.199157 | 11.16938 | 12.3025 | 0.004119 | 0.04391 |
| CRYM | 1.12604 | 5.060194 | 12.29503 | 0.004127 | 0.043974 |
| LOC108592492 | 1.047711 | 4.147569 | 12.28299 | 0.00414 | 0.044091 |
| LGI2 | 1.987196 | 1.519906 | 12.27837 | 0.004145 | 0.044113 |
| CADPS2 | 0.924569 | 6.172122 | 12.27415 | 0.00415 | 0.044124 |
| FAM129A | 1.072582 | 7.16593 | 12.25629 | 0.00417 | 0.044287 |
| LOC103792958 | 1.228336 | 3.575763 | 12.25118 | 0.004176 | 0.044323 |
| ARL4A | 0.915127 | 6.179295 | 12.2404 | 0.004188 | 0.044411 |
| ASPA | 1.367836 | 2.667353 | 12.23976 | 0.004188 | 0.044411 |
| EPB41L1 | 0.87198 | 6.063761 | 12.23261 | 0.004197 | 0.044472 |
| TNS2 | 1.064433 | 6.68102 | 12.1967 | 0.004237 | 0.044853 |
| AGT | 1.217766 | 3.037445 | 12.18939 | 0.004245 | 0.044893 |
| SYP | 1.6793 | 1.882593 | 12.17605 | 0.004261 | 0.045005 |
| MYO3B | 1.675681 | 1.879475 | 12.14871 | 0.004292 | 0.045239 |
| EFNA1 | 1.020854 | 6.507144 | 12.12634 | 0.004318 | 0.04542 |
| RNF150 | 0.934564 | 4.991216 | 12.11032 | 0.004337 | 0.045537 |

| | | | | | |
|---|---|---|---|---|---|
| KIAA1161 | 0.930944 | 6.527564 | 12.10666 | 0.004341 | 0.045556 |
| SPARC | 0.945414 | 7.879963 | 12.10214 | 0.004346 | 0.045556 |
| FGF12 | 1.168724 | 3.774085 | 12.08715 | 0.004364 | 0.045627 |
| LOC100411074 | 1.774325 | 3.6213 | 12.81536 | 0.00436 | 0.045627 |
| ZCCHC24 | 0.932036 | 6.360906 | 12.07858 | 0.004374 | 0.045659 |
| LOC103793502 | 1.484354 | 3.063956 | 12.07974 | 0.004373 | 0.045659 |
| CAMK2A | 1.686777 | 2.434156 | 12.08102 | 0.004371 | 0.045659 |
| GABRB2 | 2.510246 | 0.840832 | 12.0547 | 0.004402 | 0.045905 |
| GEM | 1.0945 | 4.348769 | 12.04766 | 0.004411 | 0.045944 |
| LOC100406675 | 1.426687 | 2.670666 | 12.04894 | 0.004409 | 0.045944 |
| NR3C2 | 1.055066 | 7.461125 | 12.04204 | 0.004417 | 0.045964 |
| LOC103793997 | 1.447582 | 3.12624 | 12.03467 | 0.004426 | 0.046007 |
| LOC100415600 | 2.861249 | 7.457633 | 17.81731 | 0.004429 | 0.046018 |
| FLVCR2 | 1.022933 | 7.971612 | 11.99994 | 0.004468 | 0.046343 |
| RBP3 | 1.635118 | 2.130009 | 11.99769 | 0.004471 | 0.046347 |
| SMLR1 | 1.056444 | 5.410193 | 11.97156 | 0.004502 | 0.046594 |
| UNC13C | 2.314811 | 1.117752 | 11.97025 | 0.004504 | 0.046594 |
| GRM8 | 1.610319 | 2.26579 | 11.96288 | 0.004513 | 0.046637 |
| LOC103796472 | 1.246599 | 3.359233 | 11.95386 | 0.004524 | 0.046702 |
| ZNF599 | 0.969257 | 4.831742 | 11.9435 | 0.004537 | 0.046792 |
| LIN7A | 1.781757 | 2.264317 | 11.94292 | 0.004537 | 0.046792 |
| CSMD3 | 1.921521 | 1.469489 | 11.93441 | 0.004548 | 0.046875 |
| SPARCL1 | 1.140562 | 7.802341 | 11.91821 | 0.004568 | 0.047008 |
| GUCY1B3 | 1.124019 | 4.72784 | 11.90829 | 0.00458 | 0.047085 |
| PRKAR1B | 1.130104 | 4.026856 | 11.90978 | 0.004579 | 0.047085 |
| MPRIP | 0.960278 | 8.67155 | 11.90619 | 0.004583 | 0.047087 |
| KCNF1 | 1.311963 | 3.049833 | 11.90055 | 0.00459 | 0.047119 |
| TMEM86B | 1.319124 | 5.255582 | 12.21805 | 0.004595 | 0.047119 |
| CHRNB4 | 1.764406 | 2.168028 | 11.89601 | 0.004596 | 0.047119 |
| LOC103791830 | 1.761563 | 1.671829 | 11.87723 | 0.004619 | 0.047335 |
| LOC108594109 | 1.53198 | 1.989489 | 11.86191 | 0.004639 | 0.047483 |
| LOC108590715 | 2.716477 | 1.28784 | 11.89965 | 0.004651 | 0.047532 |
| LOC100410989 | 2.558604 | 0.939262 | 11.8308 | 0.004678 | 0.047763 |
| LOC108590450 | 3.655884 | 0.676188 | 11.94494 | 0.004686 | 0.047796 |
| GPT | 1.167507 | 6.09324 | 11.8598 | 0.004693 | 0.047839 |
| LOC103794586 | 1.951514 | 1.635384 | 11.81314 | 0.004701 | 0.047894 |
| TTC7B | 0.928866 | 4.614225 | 11.81066 | 0.004704 | 0.047901 |
| DSTN | 1.27104 | 8.528471 | 11.88633 | 0.004711 | 0.04795 |
| MYRFL | 0.963753 | 7.029074 | 11.78912 | 0.004732 | 0.048034 |
| CCS | 0.95987 | 6.437036 | 11.77503 | 0.00475 | 0.048156 |
| LOC103794801 | 1.75104 | 2.554832 | 11.7741 | 0.004751 | 0.048156 |
| TNN | 3.21463 | 0.758011 | 11.7973 | 0.004755 | 0.048168 |
| MEIS3 | 1.390122 | 2.745719 | 11.76319 | 0.004765 | 0.048225 |
| ACSL1 | 0.947426 | 6.132884 | 11.75778 | 0.004772 | 0.048247 |
| PPP1R16A | 1.066974 | 7.412869 | 11.75097 | 0.004781 | 0.048312 |
| OCA2 | 1.262725 | 2.801866 | 11.73743 | 0.004799 | 0.048416 |
| KCNH2 | 1.323736 | 4.72238 | 11.92198 | 0.004798 | 0.048416 |
| RGS2 | 1.074174 | 4.558361 | 11.72768 | 0.004812 | 0.048496 |

| | | | | | |
|---|---|---|---|---|---|
| LOC100399678 | 2.126994 | 1.29971 | 11.71241 | 0.004832 | 0.048599 |
| SCN2B | 1.532209 | 2.179692 | 11.70346 | 0.004844 | 0.048694 |
| ST8SIA3 | 3.330877 | 0.433048 | 11.69558 | 0.004854 | 0.048774 |
| DDR2 | 0.882321 | 6.463604 | 11.69005 | 0.004862 | 0.048798 |
| SNCAIP | 1.576874 | 2.598647 | 11.69094 | 0.004861 | 0.048798 |
| GJA1 | 1.157068 | 6.141705 | 11.71842 | 0.004876 | 0.048912 |
| SV2B | 1.391536 | 2.582289 | 11.66949 | 0.004889 | 0.048974 |
| GRID2 | 2.644607 | 0.670785 | 11.66986 | 0.004889 | 0.048974 |
| ERBB2 | 0.981774 | 8.420384 | 11.6593 | 0.004903 | 0.04906 |
| LOC108591811 | 2.127085 | 0.951695 | 11.65748 | 0.004906 | 0.04906 |
| DHRS7 | 0.961849 | 4.488921 | 11.65259 | 0.004912 | 0.049101 |
| PDE8B | 1.153011 | 4.874525 | 11.64635 | 0.004921 | 0.049161 |
| TAS1R2 | 1.918542 | 1.948477 | 11.63968 | 0.00493 | 0.049226 |
| CDHR5 | 1.213071 | 9.569993 | 11.63612 | 0.004934 | 0.049249 |
| LOC100405885 | 1.154598 | 3.803504 | 11.62643 | 0.004948 | 0.04933 |
| CLSTN2 | 1.711473 | 2.12754 | 11.623 | 0.004952 | 0.049351 |
| SEMA3A | 1.45739 | 5.039259 | 12.34885 | 0.004993 | 0.049658 |
| HDAC10 | 1.152712 | 8.308556 | 11.58049 | 0.005011 | 0.049756 |
| EHD2 | 0.964466 | 6.104574 | 11.5733 | 0.005021 | 0.04983 |
| PAMR1 | 1.372684 | 2.584463 | 11.55992 | 0.005039 | 0.049882 |
| LOC100403412 | 1.943545 | 1.367401 | 11.56258 | 0.005036 | 0.049882 |
| VWC2 | 2.469896 | 1.21137 | 11.55845 | 0.005041 | 0.049882 |
| LOC108591948 | 1.252387 | 3.981708 | 11.54558 | 0.005059 | 0.049943 |
| LOC103793949 | 1.64711 | 2.219298 | 11.53756 | 0.005071 | 0.049969 |

**Supplemental Table 8. GO sets differentially expressed genes in the jejunum between IBD and non-IBD**
**Supp. Table 8a. GO sets upregulated in IBD**

| GO ID | Term | Ont | N | Up | Down | P-value |
|---|---|---|---|---|---|---|
| GO:0002376 | immune system process | BP | 2197 | 90 | 286 | 7.30E-58 |
| GO:0006955 | immune response | BP | 1473 | 39 | 227 | 7.75E-57 |
| GO:0045321 | leukocyte activation | BP | 949 | 24 | 165 | 7.62E-47 |
| GO:0046649 | lymphocyte activation | BP | 516 | 11 | 119 | 4.13E-46 |
| GO:0042110 | T cell activation | BP | 361 | 7 | 97 | 1.32E-43 |
| GO:0002682 | regulation of immune system process | BP | 1154 | 40 | 177 | 1.42E-42 |
| GO:0001775 | cell activation | BP | 1071 | 37 | 169 | 3.75E-42 |
| GO:0002250 | adaptive immune response | BP | 282 | 3 | 81 | 9.97E-39 |
| GO:0002684 | positive regulation of immune system process | BP | 816 | 20 | 136 | 5.02E-36 |
| GO:0050776 | regulation of immune response | BP | 758 | 17 | 130 | 1.07E-35 |
| GO:0006952 | defense response | BP | 1139 | 48 | 161 | 6.45E-34 |
| GO:0002252 | immune effector process | BP | 890 | 19 | 135 | 2.64E-31 |
| GO:0050778 | positive regulation of immune response | BP | 601 | 10 | 104 | 1.02E-28 |
| GO:0002521 | leukocyte differentiation | BP | 402 | 9 | 83 | 2.24E-28 |
| GO:0007159 | leukocyte cell-cell adhesion | BP | 254 | 6 | 64 | 4.13E-27 |
| GO:0002694 | regulation of leukocyte activation | BP | 380 | 12 | 78 | 1.63E-26 |
| GO:0051249 | regulation of lymphocyte activation | BP | 319 | 8 | 71 | 2.00E-26 |
| GO:0001816 | cytokine production | BP | 590 | 20 | 99 | 3.07E-26 |
| GO:0050863 | regulation of T cell activation | BP | 239 | 4 | 61 | 3.33E-26 |
| GO:0030098 | lymphocyte differentiation | BP | 280 | 6 | 66 | 3.96E-26 |
| GO:0050896 | response to stimulus | BP | 6405 | 497 | 466 | 4.17E-26 |
| GO:0030217 | T cell differentiation | BP | 198 | 5 | 55 | 1.09E-25 |
| GO:0002443 | leukocyte mediated immunity | BP | 591 | 14 | 98 | 1.45E-25 |
| GO:0045087 | innate immune response | BP | 647 | 17 | 103 | 2.06E-25 |
| GO:0050865 | regulation of cell activation | BP | 412 | 15 | 79 | 8.26E-25 |
| GO:1903037 | regulation of leukocyte cell-cell adhesion | BP | 228 | 5 | 57 | 4.73E-24 |
| GO:0048583 | regulation of response to stimulus | BP | 3246 | 232 | 285 | 7.93E-24 |
| GO:0030097 | hemopoiesis | BP | 701 | 25 | 105 | 9.74E-24 |
| GO:0002520 | immune system development | BP | 774 | 30 | 110 | 5.67E-23 |
| GO:0048534 | hematopoietic or lymphoid organ development | BP | 734 | 29 | 106 | 1.10E-22 |
| GO:0048584 | positive regulation of response to stimulus | BP | 1816 | 107 | 188 | 2.79E-22 |
| GO:0002703 | regulation of leukocyte mediated immunity | BP | 140 | 4 | 43 | 3.29E-22 |
| GO:0002366 | leukocyte activation involved in immune response | BP | 552 | 11 | 88 | 9.04E-22 |
| GO:0006950 | response to stress | BP | 3015 | 159 | 265 | 9.26E-22 |
| GO:0002263 | cell activation involved in immune response | BP | 555 | 11 | 88 | 1.32E-21 |
| GO:1903039 | positive regulation of leukocyte cell-cell adhesion | BP | 168 | 2 | 46 | 2.28E-21 |
| GO:0001817 | regulation of cytokine production | BP | 533 | 17 | 85 | 4.73E-21 |
| GO:0032609 | interferon-gamma production | BP | 80 | 2 | 32 | 8.10E-21 |
| GO:0007165 | signal transduction | BP | 4276 | 338 | 335 | 3.17E-20 |
| GO:0002253 | activation of immune response | BP | 486 | 6 | 79 | 4.03E-20 |
| GO:0022407 | regulation of cell-cell adhesion | BP | 314 | 11 | 61 | 1.28E-19 |
| GO:0002757 | immune response-activating signal transduction | BP | 434 | 5 | 73 | 1.65E-19 |
| GO:0051716 | cellular response to stimulus | BP | 5380 | 422 | 393 | 2.31E-19 |
| GO:0002449 | lymphocyte mediated immunity | BP | 171 | 3 | 44 | 2.47E-19 |
| GO:0002696 | positive regulation of leukocyte activation | BP | 237 | 5 | 52 | 2.72E-19 |
| GO:0002764 | immune response-regulating signaling pathway | BP | 459 | 7 | 75 | 2.87E-19 |
| GO:0030155 | regulation of cell adhesion | BP | 562 | 36 | 84 | 5.62E-19 |
| GO:0050867 | positive regulation of cell activation | BP | 246 | 6 | 52 | 1.54E-18 |
| GO:0022409 | positive regulation of cell-cell adhesion | BP | 197 | 4 | 46 | 2.49E-18 |
| GO:0046631 | alpha-beta T cell activation | BP | 114 | 2 | 35 | 2.70E-18 |
| GO:0002697 | regulation of immune effector process | BP | 288 | 7 | 56 | 4.15E-18 |
| GO:0051251 | positive regulation of lymphocyte activation | BP | 208 | 4 | 47 | 4.34E-18 |
| GO:0050870 | positive regulation of T cell activation | BP | 155 | 2 | 40 | 9.84E-18 |
| GO:0051707 | response to other organism | BP | 646 | 30 | 89 | 9.87E-18 |
| GO:0043207 | response to external biotic stimulus | BP | 648 | 31 | 89 | 1.21E-17 |
| GO:0009607 | response to biotic stimulus | BP | 676 | 33 | 91 | 1.85E-17 |
| GO:0023052 | signaling | BP | 4643 | 396 | 346 | 2.69E-17 |
| GO:0002274 | myeloid leukocyte activation | BP | 519 | 13 | 77 | 2.93E-17 |
| GO:0071944 | cell periphery | CC | 3744 | 430 | 294 | 5.23E-17 |
| GO:0070661 | leukocyte proliferation | BP | 213 | 9 | 46 | 6.55E-17 |

| | | | | | | |
|---|---|---|---|---|---|---|
| GO:0007154 | cell communication | BP | 4656 | 397 | 345 | 8.48E-17 |
| GO:0009605 | response to external stimulus | BP | 1751 | 148 | 169 | 9.79E-17 |
| GO:0002768 | immune response-regulating cell surface receptor signaling pathwa | BP | 320 | 3 | 57 | 1.38E-16 |
| GO:0071345 | cellular response to cytokine stimulus | BP | 852 | 41 | 103 | 2.28E-16 |
| GO:0046651 | lymphocyte proliferation | BP | 194 | 7 | 43 | 2.54E-16 |
| GO:0002460 | adaptive immune response based on somatic recombination of imm | BP | 186 | 2 | 42 | 2.81E-16 |
| GO:0002429 | immune response-activating cell surface receptor signaling pathwa | BP | 296 | 1 | 54 | 3.11E-16 |
| GO:0005886 | plasma membrane | CC | 3656 | 422 | 286 | 3.68E-16 |
| GO:0032943 | mononuclear cell proliferation | BP | 196 | 7 | 43 | 3.78E-16 |
| GO:0002706 | regulation of lymphocyte mediated immunity | BP | 96 | 3 | 30 | 4.21E-16 |
| GO:0032649 | regulation of interferon-gamma production | BP | 71 | 2 | 26 | 4.88E-16 |
| GO:0007166 | cell surface receptor signaling pathway | BP | 2269 | 182 | 201 | 5.09E-16 |
| GO:0034097 | response to cytokine | BP | 927 | 45 | 108 | 5.30E-16 |
| GO:0031347 | regulation of defense response | BP | 611 | 22 | 82 | 1.04E-15 |
| GO:0070663 | regulation of leukocyte proliferation | BP | 159 | 6 | 38 | 1.04E-15 |
| GO:0050900 | leukocyte migration | BP | 326 | 23 | 56 | 1.36E-15 |
| GO:0098609 | cell-cell adhesion | BP | 638 | 72 | 84 | 1.39E-15 |
| GO:0019221 | cytokine-mediated signaling pathway | BP | 591 | 21 | 80 | 1.47E-15 |
| GO:0001819 | positive regulation of cytokine production | BP | 347 | 10 | 58 | 1.47E-15 |
| GO:0002699 | positive regulation of immune effector process | BP | 153 | 5 | 37 | 1.66E-15 |
| GO:0001906 | cell killing | BP | 87 | 2 | 28 | 1.73E-15 |
| GO:0050851 | antigen receptor-mediated signaling pathway | BP | 199 | 1 | 42 | 3.60E-15 |
| GO:0042098 | T cell proliferation | BP | 133 | 4 | 34 | 3.97E-15 |
| GO:1902105 | regulation of leukocyte differentiation | BP | 209 | 3 | 43 | 4.40E-15 |
| GO:0050670 | regulation of lymphocyte proliferation | BP | 150 | 6 | 36 | 5.21E-15 |
| GO:0032944 | regulation of mononuclear cell proliferation | BP | 151 | 6 | 36 | 6.51E-15 |
| GO:0032729 | positive regulation of interferon-gamma production | BP | 49 | 0 | 21 | 7.81E-15 |
| GO:0002705 | positive regulation of leukocyte mediated immunity | BP | 87 | 2 | 27 | 1.54E-14 |
| GO:0048518 | positive regulation of biological process | BP | 4721 | 353 | 340 | 1.99E-14 |
| GO:0002285 | lymphocyte activation involved in immune response | BP | 133 | 1 | 33 | 2.58E-14 |
| GO:0001909 | leukocyte mediated cytotoxicity | BP | 64 | 2 | 23 | 3.92E-14 |
| GO:0045785 | positive regulation of cell adhesion | BP | 331 | 18 | 54 | 4.16E-14 |
| GO:0006909 | phagocytosis | BP | 223 | 7 | 43 | 4.91E-14 |
| GO:0009897 | external side of plasma membrane | CC | 220 | 17 | 42 | 1.39E-13 |
| GO:0035556 | intracellular signal transduction | BP | 2352 | 171 | 198 | 1.83E-13 |
| GO:0009617 | response to bacterium | BP | 414 | 28 | 60 | 3.41E-13 |
| GO:0006954 | inflammatory response | BP | 542 | 33 | 71 | 3.48E-13 |
| GO:0045088 | regulation of innate immune response | BP | 360 | 7 | 55 | 3.93E-13 |
| GO:0035710 | CD4-positive, alpha-beta T cell activation | BP | 77 | 1 | 24 | 4.06E-13 |
| GO:0042129 | regulation of T cell proliferation | BP | 107 | 3 | 28 | 5.81E-13 |
| GO:0002708 | positive regulation of lymphocyte mediated immunity | BP | 65 | 1 | 22 | 5.84E-13 |
| GO:0046632 | alpha-beta T cell differentiation | BP | 87 | 2 | 25 | 1.03E-12 |
| GO:0044459 | plasma membrane part | CC | 2045 | 269 | 176 | 1.06E-12 |
| GO:0031341 | regulation of cell killing | BP | 55 | 2 | 20 | 1.42E-12 |
| GO:0007155 | cell adhesion | BP | 1113 | 137 | 113 | 1.53E-12 |
| GO:0002444 | myeloid leukocyte mediated immunity | BP | 441 | 10 | 61 | 1.72E-12 |
| GO:0001910 | regulation of leukocyte mediated cytotoxicity | BP | 44 | 2 | 18 | 1.76E-12 |
| GO:0022610 | biological adhesion | BP | 1116 | 137 | 113 | 1.83E-12 |
| GO:0038023 | signaling receptor activity | MF | 703 | 90 | 82 | 2.60E-12 |
| GO:0002683 | negative regulation of immune system process | BP | 344 | 19 | 52 | 2.70E-12 |
| GO:0070665 | positive regulation of leukocyte proliferation | BP | 98 | 3 | 26 | 2.78E-12 |
| GO:0045619 | regulation of lymphocyte differentiation | BP | 131 | 3 | 30 | 3.68E-12 |
| GO:0080134 | regulation of response to stress | BP | 1228 | 51 | 120 | 3.73E-12 |
| GO:0031349 | positive regulation of defense response | BP | 392 | 14 | 56 | 3.96E-12 |
| GO:0043368 | positive T cell selection | BP | 26 | 0 | 14 | 4.95E-12 |
| GO:0098552 | side of membrane | CC | 383 | 30 | 55 | 5.05E-12 |
| GO:0065009 | regulation of molecular function | BP | 2322 | 174 | 191 | 5.39E-12 |
| GO:0042113 | B cell activation | BP | 187 | 4 | 36 | 6.19E-12 |
| GO:0045058 | T cell selection | BP | 37 | 1 | 16 | 1.10E-11 |
| GO:0002695 | negative regulation of leukocyte activation | BP | 128 | 8 | 29 | 1.12E-11 |
| GO:0043299 | leukocyte degranulation | BP | 426 | 10 | 58 | 1.21E-11 |
| GO:1902531 | regulation of intracellular signal transduction | BP | 1532 | 103 | 139 | 1.31E-11 |
| GO:0009986 | cell surface | CC | 598 | 68 | 72 | 1.45E-11 |
| GO:0044093 | positive regulation of molecular function | BP | 1439 | 97 | 132 | 2.32E-11 |

| | | | | | | |
|---|---|---|---|---|---|---|
| GO:0002275 | myeloid cell activation involved in immune response | BP | 433 | 10 | 58 | 2.36E-11 |
| GO:0060089 | molecular transducer activity | MF | 747 | 92 | 83 | 2.43E-11 |
| GO:0045580 | regulation of T cell differentiation | BP | 107 | 2 | 26 | 2.45E-11 |
| GO:0042102 | positive regulation of T cell proliferation | BP | 63 | 1 | 20 | 2.55E-11 |
| GO:0031226 | intrinsic component of plasma membrane | CC | 1165 | 179 | 113 | 2.91E-11 |
| GO:0006968 | cellular defense response | BP | 34 | 1 | 15 | 3.44E-11 |
| GO:0043367 | CD4-positive, alpha-beta T cell differentiation | BP | 64 | 1 | 20 | 3.54E-11 |
| GO:0050671 | positive regulation of lymphocyte proliferation | BP | 93 | 3 | 24 | 3.67E-11 |
| GO:0002228 | natural killer cell mediated immunity | BP | 40 | 2 | 16 | 4.63E-11 |
| GO:0032946 | positive regulation of mononuclear cell proliferation | BP | 94 | 3 | 24 | 4.69E-11 |
| GO:0031343 | positive regulation of cell killing | BP | 35 | 1 | 15 | 5.73E-11 |
| GO:1903706 | regulation of hemopoiesis | BP | 362 | 8 | 51 | 6.35E-11 |
| GO:0004888 | transmembrane signaling receptor activity | MF | 542 | 70 | 66 | 6.70E-11 |
| GO:0048522 | positive regulation of cellular process | BP | 4206 | 319 | 297 | 7.23E-11 |
| GO:0030101 | natural killer cell activation | BP | 47 | 1 | 17 | 7.63E-11 |
| GO:0050790 | regulation of catalytic activity | BP | 1833 | 122 | 156 | 7.96E-11 |
| GO:0050866 | negative regulation of cell activation | BP | 147 | 10 | 30 | 8.07E-11 |
| GO:0009966 | regulation of signal transduction | BP | 2476 | 180 | 196 | 8.60E-11 |
| GO:0046634 | regulation of alpha-beta T cell activation | BP | 74 | 1 | 21 | 8.84E-11 |
| GO:0051250 | negative regulation of lymphocyte activation | BP | 106 | 5 | 25 | 1.19E-10 |
| GO:0002456 | T cell mediated immunity | BP | 62 | 2 | 19 | 1.61E-10 |
| GO:0051240 | positive regulation of multicellular organismal process | BP | 1345 | 140 | 123 | 1.61E-10 |
| GO:0045089 | positive regulation of innate immune response | BP | 304 | 5 | 45 | 1.71E-10 |
| GO:0001912 | positive regulation of leukocyte mediated cytotoxicity | BP | 27 | 1 | 13 | 1.83E-10 |
| GO:0002819 | regulation of adaptive immune response | BP | 117 | 2 | 26 | 2.08E-10 |
| GO:0050852 | T cell receptor signaling pathway | BP | 162 | 1 | 31 | 2.17E-10 |
| GO:0001776 | leukocyte homeostasis | BP | 71 | 2 | 20 | 2.88E-10 |
| GO:0050853 | B cell receptor signaling pathway | BP | 51 | 0 | 17 | 3.38E-10 |
| GO:0036230 | granulocyte activation | BP | 405 | 8 | 53 | 4.24E-10 |
| GO:0009967 | positive regulation of signal transduction | BP | 1310 | 86 | 119 | 5.21E-10 |
| GO:0048870 | cell motility | BP | 1236 | 109 | 114 | 5.26E-10 |
| GO:0051674 | localization of cell | BP | 1236 | 109 | 114 | 5.26E-10 |
| GO:0050789 | regulation of biological process | BP | 8258 | 598 | 503 | 5.73E-10 |
| GO:0005887 | integral component of plasma membrane | CC | 1107 | 166 | 105 | 6.05E-10 |
| GO:0098542 | defense response to other organism | BP | 296 | 4 | 43 | 8.01E-10 |
| GO:0042119 | neutrophil activation | BP | 401 | 8 | 52 | 8.66E-10 |
| GO:0042267 | natural killer cell mediated cytotoxicity | BP | 36 | 2 | 14 | 1.20E-09 |
| GO:0002709 | regulation of T cell mediated immunity | BP | 42 | 2 | 15 | 1.24E-09 |
| GO:0065007 | biological regulation | BP | 8733 | 631 | 524 | 1.38E-09 |
| GO:2000514 | regulation of CD4-positive, alpha-beta T cell activation | BP | 50 | 0 | 16 | 2.23E-09 |
| GO:0040011 | locomotion | BP | 1418 | 129 | 124 | 2.28E-09 |
| GO:0006897 | endocytosis | BP | 590 | 39 | 66 | 2.44E-09 |
| GO:0051239 | regulation of multicellular organismal process | BP | 2424 | 267 | 187 | 2.49E-09 |
| GO:0060326 | cell chemotaxis | BP | 209 | 15 | 34 | 2.60E-09 |
| GO:1902107 | positive regulation of leukocyte differentiation | BP | 113 | 1 | 24 | 2.73E-09 |
| GO:0002237 | response to molecule of bacterial origin | BP | 264 | 19 | 39 | 3.07E-09 |
| GO:1903555 | regulation of tumor necrosis factor superfamily cytokine productio | BP | 115 | 4 | 24 | 3.96E-09 |
| GO:0070887 | cellular response to chemical stimulus | BP | 2488 | 204 | 190 | 3.99E-09 |
| GO:0036037 | CD8-positive, alpha-beta T cell activation | BP | 18 | 0 | 10 | 4.02E-09 |
| GO:0002286 | T cell activation involved in immune response | BP | 74 | 0 | 19 | 4.46E-09 |
| GO:0050868 | negative regulation of T cell activation | BP | 82 | 3 | 20 | 4.62E-09 |
| GO:0043085 | positive regulation of catalytic activity | BP | 1163 | 69 | 106 | 4.63E-09 |
| GO:0016477 | cell migration | BP | 1136 | 106 | 104 | 5.23E-09 |
| GO:0071310 | cellular response to organic substance | BP | 2086 | 163 | 165 | 5.37E-09 |
| GO:0008283 | cell proliferation | BP | 1486 | 117 | 127 | 5.81E-09 |
| GO:0051336 | regulation of hydrolase activity | BP | 1007 | 70 | 95 | 6.00E-09 |
| GO:0002446 | neutrophil mediated immunity | BP | 401 | 8 | 50 | 7.03E-09 |
| GO:0071706 | tumor necrosis factor superfamily cytokine production | BP | 119 | 4 | 24 | 8.12E-09 |
| GO:0005515 | protein binding | MF | 9061 | 582 | 536 | 8.55E-09 |
| GO:0045055 | regulated exocytosis | BP | 651 | 40 | 69 | 9.85E-09 |
| GO:0002822 | regulation of adaptive immune response based on somatic recombi | BP | 103 | 2 | 22 | 1.12E-08 |
| GO:0050663 | cytokine secretion | BP | 169 | 7 | 29 | 1.13E-08 |
| GO:0072676 | lymphocyte migration | BP | 78 | 3 | 19 | 1.15E-08 |
| GO:0010646 | regulation of cell communication | BP | 2778 | 241 | 205 | 1.31E-08 |

| | | | | | | |
|---|---|---|---|---|---|---|
| GO:0007162 | negative regulation of cell adhesion | BP | 223 | 16 | 34 | 1.43E-08 |
| GO:0032663 | regulation of interleukin-2 production | BP | 50 | 1 | 15 | 1.91E-08 |
| GO:0072678 | T cell migration | BP | 50 | 0 | 15 | 1.91E-08 |
| GO:0043374 | CD8-positive, alpha-beta T cell differentiation | BP | 12 | 0 | 8 | 2.10E-08 |
| GO:0043312 | neutrophil degranulation | BP | 390 | 8 | 48 | 2.16E-08 |
| GO:0032623 | interleukin-2 production | BP | 58 | 2 | 16 | 2.45E-08 |
| GO:0002283 | neutrophil activation involved in immune response | BP | 392 | 8 | 48 | 2.54E-08 |
| GO:0046635 | positive regulation of alpha-beta T cell activation | BP | 51 | 1 | 15 | 2.58E-08 |
| GO:0002715 | regulation of natural killer cell mediated immunity | BP | 26 | 2 | 11 | 2.60E-08 |
| GO:0023051 | regulation of signaling | BP | 2819 | 246 | 206 | 2.68E-08 |
| GO:0042742 | defense response to bacterium | BP | 117 | 3 | 23 | 2.76E-08 |
| GO:0002821 | positive regulation of adaptive immune response | BP | 74 | 1 | 18 | 2.86E-08 |
| GO:0045621 | positive regulation of lymphocyte differentiation | BP | 74 | 1 | 18 | 2.86E-08 |
| GO:0030099 | myeloid cell differentiation | BP | 321 | 12 | 42 | 2.94E-08 |
| GO:1903708 | positive regulation of hemopoiesis | BP | 146 | 1 | 26 | 2.97E-08 |
| GO:0050707 | regulation of cytokine secretion | BP | 146 | 4 | 26 | 2.97E-08 |
| GO:0002440 | production of molecular mediator of immune response | BP | 156 | 5 | 27 | 3.01E-08 |
| GO:1903038 | negative regulation of leukocyte cell-cell adhesion | BP | 92 | 4 | 20 | 3.80E-08 |
| GO:0051345 | positive regulation of hydrolase activity | BP | 633 | 36 | 66 | 4.00E-08 |
| GO:0033077 | T cell differentiation in thymus | BP | 60 | 2 | 16 | 4.15E-08 |
| GO:1902533 | positive regulation of intracellular signal transduction | BP | 815 | 54 | 79 | 4.22E-08 |
| GO:0010647 | positive regulation of cell communication | BP | 1428 | 110 | 120 | 4.43E-08 |
| GO:0050794 | regulation of cellular process | BP | 7763 | 550 | 470 | 4.47E-08 |
| GO:0002287 | alpha-beta T cell activation involved in immune response | BP | 53 | 0 | 15 | 4.60E-08 |
| GO:0002293 | alpha-beta T cell differentiation involved in immune response | BP | 53 | 0 | 15 | 4.60E-08 |
| GO:0023056 | positive regulation of signaling | BP | 1434 | 110 | 120 | 5.61E-08 |
| GO:0032680 | regulation of tumor necrosis factor production | BP | 112 | 4 | 22 | 5.61E-08 |
| GO:0002824 | positive regulation of adaptive immune response based on somatic r | BP | 69 | 1 | 17 | 5.66E-08 |
| GO:0043370 | regulation of CD4-positive, alpha-beta T cell differentiation | BP | 40 | 0 | 13 | 5.87E-08 |
| GO:0030595 | leukocyte chemotaxis | BP | 153 | 10 | 26 | 7.97E-08 |
| GO:0048869 | cellular developmental process | BP | 3171 | 313 | 224 | 8.71E-08 |
| GO:0045582 | positive regulation of T cell differentiation | BP | 63 | 1 | 16 | 8.79E-08 |
| GO:0051704 | multi-organism process | BP | 1827 | 85 | 144 | 8.84E-08 |
| GO:0032640 | tumor necrosis factor production | BP | 115 | 4 | 22 | 9.22E-08 |
| GO:0005576 | extracellular region | CC | 2796 | 243 | 202 | 9.83E-08 |
| GO:0042127 | regulation of cell proliferation | BP | 1233 | 99 | 106 | 1.02E-07 |
| GO:0002292 | T cell differentiation involved in immune response | BP | 56 | 0 | 15 | 1.03E-07 |
| GO:0032496 | response to lipopolysaccharide | BP | 253 | 18 | 35 | 1.07E-07 |
| GO:0010033 | response to organic substance | BP | 2529 | 205 | 186 | 1.13E-07 |
| GO:0005615 | extracellular space | CC | 2228 | 185 | 168 | 1.14E-07 |
| GO:0006887 | exocytosis | BP | 749 | 49 | 73 | 1.17E-07 |
| GO:0042221 | response to chemical | BP | 3288 | 285 | 230 | 1.21E-07 |
| GO:0032940 | secretion by cell | BP | 1193 | 100 | 103 | 1.29E-07 |
| GO:0008284 | positive regulation of cell proliferation | BP | 667 | 47 | 67 | 1.31E-07 |
| GO:0006959 | humoral immune response | BP | 127 | 1 | 23 | 1.35E-07 |
| GO:0042269 | regulation of natural killer cell mediated cytotoxicity | BP | 24 | 2 | 10 | 1.36E-07 |
| GO:0022408 | negative regulation of cell-cell adhesion | BP | 138 | 5 | 24 | 1.60E-07 |
| GO:0002573 | myeloid leukocyte differentiation | BP | 159 | 6 | 26 | 1.76E-07 |
| GO:0046903 | secretion | BP | 1294 | 110 | 109 | 1.86E-07 |
| GO:0006928 | movement of cell or subcellular component | BP | 1611 | 154 | 129 | 2.07E-07 |
| GO:0071222 | cellular response to lipopolysaccharide | BP | 150 | 4 | 25 | 2.08E-07 |
| GO:0006935 | chemotaxis | BP | 487 | 53 | 53 | 2.42E-07 |
| GO:0002294 | CD4-positive, alpha-beta T cell differentiation involved in immune | BP | 52 | 0 | 14 | 2.57E-07 |
| GO:0002260 | lymphocyte homeostasis | BP | 52 | 0 | 14 | 2.57E-07 |
| GO:0042330 | taxis | BP | 489 | 53 | 53 | 2.76E-07 |
| GO:0032655 | regulation of interleukin-12 production | BP | 45 | 2 | 13 | 2.80E-07 |
| GO:0002886 | regulation of myeloid leukocyte mediated immunity | BP | 45 | 1 | 13 | 2.80E-07 |
| GO:0098657 | import into cell | BP | 681 | 55 | 67 | 2.85E-07 |
| GO:0031982 | vesicle | CC | 2976 | 206 | 210 | 3.11E-07 |
| GO:0030154 | cell differentiation | BP | 3014 | 301 | 212 | 3.39E-07 |
| GO:0071346 | cellular response to interferon-gamma | BP | 124 | 6 | 22 | 3.67E-07 |
| GO:2001187 | positive regulation of CD8-positive, alpha-beta T cell activation | BP | 5 | 0 | 5 | 3.73E-07 |
| GO:0032620 | interleukin-17 production | BP | 21 | 1 | 9 | 4.38E-07 |
| GO:0071219 | cellular response to molecule of bacterial origin | BP | 156 | 4 | 25 | 4.51E-07 |

| | | | | | | |
|---|---|---|---|---|---|---|
| GO:0038093 | Fc receptor signaling pathway | BP | 167 | 2 | 26 | 4.75E-07 |
| GO:0034341 | response to interferon-gamma | BP | 136 | 6 | 23 | 4.87E-07 |
| GO:0032615 | interleukin-12 production | BP | 47 | 3 | 13 | 4.90E-07 |
| GO:0072540 | T-helper 17 cell lineage commitment | BP | 8 | 0 | 6 | 4.93E-07 |
| GO:0010941 | regulation of cell death | BP | 1321 | 83 | 109 | 5.23E-07 |
| GO:0050854 | regulation of antigen receptor-mediated signaling pathway | BP | 55 | 1 | 14 | 5.47E-07 |
| GO:2001185 | regulation of CD8-positive, alpha-beta T cell activation | BP | 12 | 0 | 7 | 6.24E-07 |
| GO:0002295 | T-helper cell lineage commitment | BP | 12 | 0 | 7 | 6.24E-07 |
| GO:0042981 | regulation of apoptotic process | BP | 1217 | 75 | 102 | 6.26E-07 |
| GO:0004896 | cytokine receptor activity | MF | 72 | 6 | 16 | 6.37E-07 |
| GO:0098772 | molecular function regulator | MF | 1264 | 95 | 105 | 6.39E-07 |
| GO:0043067 | regulation of programmed cell death | BP | 1233 | 77 | 103 | 6.39E-07 |
| GO:0032501 | multicellular organismal process | BP | 5121 | 490 | 327 | 7.31E-07 |
| GO:0002758 | innate immune response-activating signal transduction | BP | 250 | 4 | 33 | 7.41E-07 |
| GO:0002711 | positive regulation of T cell mediated immunity | BP | 28 | 1 | 10 | 7.51E-07 |
| GO:0002367 | cytokine production involved in immune response | BP | 82 | 4 | 17 | 8.16E-07 |
| GO:0002717 | positive regulation of natural killer cell mediated immunity | BP | 17 | 1 | 8 | 8.16E-07 |
| GO:0048513 | animal organ development | BP | 2568 | 248 | 184 | 8.73E-07 |
| GO:0038096 | Fc-gamma receptor signaling pathway involved in phagocytosis | BP | 74 | 0 | 16 | 9.45E-07 |
| GO:0002433 | immune response-regulating cell surface receptor signaling pathwa | BP | 74 | 0 | 16 | 9.45E-07 |
| GO:0048872 | homeostasis of number of cells | BP | 207 | 7 | 29 | 1.02E-06 |
| GO:0042093 | T-helper cell differentiation | BP | 50 | 0 | 13 | 1.07E-06 |
| GO:0002218 | activation of innate immune response | BP | 266 | 4 | 34 | 1.07E-06 |
| GO:0038094 | Fc-gamma receptor signaling pathway | BP | 75 | 0 | 16 | 1.14E-06 |
| GO:0050830 | defense response to Gram-positive bacterium | BP | 43 | 3 | 12 | 1.22E-06 |
| GO:0044421 | extracellular region part | CC | 2377 | 207 | 172 | 1.24E-06 |
| GO:0016020 | membrane | CC | 6576 | 532 | 402 | 1.28E-06 |
| GO:0042100 | B cell proliferation | BP | 67 | 1 | 15 | 1.29E-06 |
| GO:0043373 | CD4-positive, alpha-beta T cell lineage commitment | BP | 13 | 0 | 7 | 1.29E-06 |
| GO:0050864 | regulation of B cell activation | BP | 94 | 2 | 18 | 1.33E-06 |
| GO:0006915 | apoptotic process | BP | 1545 | 93 | 121 | 1.71E-06 |
| GO:0031295 | T cell costimulation | BP | 52 | 0 | 13 | 1.74E-06 |
| GO:0071216 | cellular response to biotic stimulus | BP | 179 | 5 | 26 | 1.85E-06 |
| GO:0030141 | secretory granule | CC | 632 | 32 | 61 | 1.89E-06 |
| GO:0006468 | protein phosphorylation | BP | 1565 | 107 | 122 | 1.95E-06 |
| GO:0012501 | programmed cell death | BP | 1599 | 95 | 124 | 2.06E-06 |
| GO:0002312 | B cell activation involved in immune response | BP | 61 | 1 | 14 | 2.12E-06 |
| GO:0043376 | regulation of CD8-positive, alpha-beta T cell differentiation | BP | 6 | 0 | 5 | 2.14E-06 |
| GO:0042608 | T cell receptor binding | MF | 6 | 0 | 5 | 2.14E-06 |
| GO:0045123 | cellular extravasation | BP | 53 | 4 | 13 | 2.19E-06 |
| GO:0031294 | lymphocyte costimulation | BP | 53 | 0 | 13 | 2.19E-06 |
| GO:0046637 | regulation of alpha-beta T cell differentiation | BP | 53 | 1 | 13 | 2.19E-06 |
| GO:0070821 | tertiary granule membrane | CC | 53 | 3 | 13 | 2.19E-06 |
| GO:0043029 | T cell homeostasis | BP | 31 | 0 | 10 | 2.20E-06 |
| GO:0002700 | regulation of production of molecular mediator of immune response | BP | 107 | 4 | 19 | 2.25E-06 |
| GO:0001773 | myeloid dendritic cell activation | BP | 19 | 0 | 8 | 2.31E-06 |
| GO:1903557 | positive regulation of tumor necrosis factor superfamily cytokine pı | BP | 70 | 3 | 15 | 2.32E-06 |
| GO:0072643 | interferon-gamma secretion | BP | 14 | 0 | 7 | 2.47E-06 |
| GO:0001818 | negative regulation of cytokine production | BP | 205 | 5 | 28 | 2.60E-06 |
| GO:0002431 | Fc receptor mediated stimulatory signaling pathway | BP | 80 | 0 | 16 | 2.83E-06 |
| GO:0002448 | mast cell mediated immunity | BP | 39 | 2 | 11 | 3.03E-06 |
| GO:0005102 | signaling receptor binding | MF | 1074 | 98 | 90 | 3.18E-06 |
| GO:0030667 | secretory granule membrane | CC | 232 | 15 | 30 | 3.61E-06 |
| GO:0001913 | T cell mediated cytotoxicity | BP | 20 | 1 | 8 | 3.68E-06 |
| GO:0045576 | mast cell activation | BP | 48 | 3 | 12 | 4.35E-06 |
| GO:0002363 | alpha-beta T cell lineage commitment | BP | 15 | 0 | 7 | 4.42E-06 |
| GO:0043369 | CD4-positive or CD8-positive, alpha-beta T cell lineage commitme | BP | 15 | 0 | 7 | 4.42E-06 |
| GO:0008219 | cell death | BP | 1695 | 103 | 128 | 5.37E-06 |
| GO:0051094 | positive regulation of developmental process | BP | 1057 | 124 | 88 | 5.38E-06 |
| GO:0030695 | GTPase regulator activity | MF | 261 | 14 | 32 | 5.42E-06 |
| GO:0010818 | T cell chemotaxis | BP | 21 | 0 | 8 | 5.67E-06 |
| GO:0043087 | regulation of GTPase activity | BP | 404 | 28 | 43 | 6.02E-06 |
| GO:0035747 | natural killer cell chemotaxis | BP | 7 | 0 | 5 | 7.18E-06 |
| GO:1902715 | positive regulation of interferon-gamma secretion | BP | 7 | 0 | 5 | 7.18E-06 |

| | | | | | | |
|---|---|---|---|---|---|---|
| GO:0043378 | positive regulation of CD8-positive, alpha-beta T cell differentiatio | BP | 4 | 0 | 4 | 7.23E-06 |
| GO:0042581 | specific granule | CC | 126 | 4 | 20 | 7.33E-06 |
| GO:0002224 | toll-like receptor signaling pathway | BP | 126 | 4 | 20 | 7.33E-06 |
| GO:0032743 | positive regulation of interleukin-2 production | BP | 28 | 0 | 9 | 7.41E-06 |
| GO:0030234 | enzyme regulator activity | MF | 792 | 43 | 70 | 7.50E-06 |
| GO:0045954 | positive regulation of natural killer cell mediated cytotoxicity | BP | 16 | 1 | 7 | 7.51E-06 |
| GO:0045061 | thymic T cell selection | BP | 16 | 1 | 7 | 7.51E-06 |
| GO:0042088 | T-helper 1 type immune response | BP | 35 | 0 | 10 | 7.54E-06 |
| GO:0042592 | homeostatic process | BP | 1496 | 131 | 115 | 7.93E-06 |
| GO:0050715 | positive regulation of cytokine secretion | BP | 96 | 2 | 17 | 7.93E-06 |
| GO:0032760 | positive regulation of tumor necrosis factor production | BP | 68 | 3 | 14 | 8.27E-06 |
| GO:0002718 | regulation of cytokine production involved in immune response | BP | 68 | 3 | 14 | 8.27E-06 |
| GO:2000106 | regulation of leukocyte apoptotic process | BP | 68 | 2 | 14 | 8.27E-06 |
| GO:0019220 | regulation of phosphate metabolic process | BP | 1401 | 97 | 109 | 8.39E-06 |
| GO:0009615 | response to virus | BP | 242 | 3 | 30 | 8.51E-06 |
| GO:0030888 | regulation of B cell proliferation | BP | 51 | 1 | 12 | 8.59E-06 |
| GO:0050793 | regulation of developmental process | BP | 2011 | 209 | 146 | 8.79E-06 |
| GO:0002761 | regulation of myeloid leukocyte differentiation | BP | 87 | 1 | 16 | 8.85E-06 |
| GO:0051174 | regulation of phosphorus metabolic process | BP | 1403 | 97 | 109 | 8.95E-06 |
| GO:0042325 | regulation of phosphorylation | BP | 1244 | 82 | 99 | 9.14E-06 |
| GO:0005096 | GTPase activator activity | MF | 231 | 12 | 29 | 9.39E-06 |
| GO:0046636 | negative regulation of alpha-beta T cell activation | BP | 29 | 1 | 9 | 1.02E-05 |
| GO:0045937 | positive regulation of phosphate metabolic process | BP | 860 | 64 | 74 | 1.05E-05 |
| GO:0010562 | positive regulation of phosphorus metabolic process | BP | 860 | 64 | 74 | 1.05E-05 |
| GO:0060589 | nucleoside-triphosphatase regulator activity | MF | 295 | 16 | 34 | 1.08E-05 |
| GO:0002221 | pattern recognition receptor signaling pathway | BP | 163 | 4 | 23 | 1.17E-05 |
| GO:0042629 | mast cell granule | CC | 17 | 1 | 7 | 1.22E-05 |
| GO:0032660 | regulation of interleukin-17 production | BP | 17 | 1 | 7 | 1.22E-05 |
| GO:0045589 | regulation of regulatory T cell differentiation | BP | 23 | 0 | 8 | 1.25E-05 |
| GO:0070820 | tertiary granule | CC | 120 | 5 | 19 | 1.28E-05 |
| GO:0043303 | mast cell degranulation | BP | 37 | 2 | 10 | 1.30E-05 |
| GO:0019730 | antimicrobial humoral response | BP | 45 | 0 | 11 | 1.38E-05 |
| GO:0030183 | B cell differentiation | BP | 100 | 3 | 17 | 1.39E-05 |
| GO:0033627 | cell adhesion mediated by integrin | BP | 62 | 3 | 13 | 1.41E-05 |
| GO:0030593 | neutrophil chemotaxis | BP | 62 | 4 | 13 | 1.41E-05 |
| GO:0044425 | membrane part | CC | 4548 | 435 | 288 | 1.51E-05 |
| GO:0001932 | regulation of protein phosphorylation | BP | 1118 | 76 | 90 | 1.58E-05 |
| GO:0002279 | mast cell activation involved in immune response | BP | 38 | 2 | 10 | 1.68E-05 |
| GO:0031410 | cytoplasmic vesicle | CC | 1853 | 112 | 135 | 1.78E-05 |
| GO:0045066 | regulatory T cell differentiation | BP | 24 | 0 | 8 | 1.78E-05 |
| GO:0016192 | vesicle-mediated transport | BP | 1671 | 99 | 124 | 1.81E-05 |
| GO:0097708 | intracellular vesicle | CC | 1855 | 112 | 135 | 1.88E-05 |
| GO:0006271 | DNA strand elongation involved in DNA replication | BP | 18 | 0 | 7 | 1.91E-05 |
| GO:0002360 | T cell lineage commitment | BP | 18 | 0 | 7 | 1.91E-05 |
| GO:0070098 | chemokine-mediated signaling pathway | BP | 55 | 4 | 12 | 1.96E-05 |
| GO:0009893 | positive regulation of metabolic process | BP | 2834 | 181 | 192 | 1.99E-05 |
| GO:0002377 | immunoglobulin production | BP | 83 | 1 | 15 | 2.09E-05 |
| GO:0048247 | lymphocyte chemotaxis | BP | 39 | 1 | 10 | 2.16E-05 |
| GO:0032653 | regulation of interleukin-10 production | BP | 39 | 0 | 10 | 2.16E-05 |
| GO:0050727 | regulation of inflammatory response | BP | 280 | 15 | 32 | 2.31E-05 |
| GO:0001934 | positive regulation of protein phosphorylation | BP | 759 | 57 | 66 | 2.33E-05 |
| GO:0006261 | DNA-dependent DNA replication | BP | 136 | 0 | 20 | 2.34E-05 |
| GO:0042108 | positive regulation of cytokine biosynthetic process | BP | 56 | 0 | 12 | 2.38E-05 |
| GO:0042832 | defense response to protozoan | BP | 13 | 0 | 6 | 2.41E-05 |
| GO:0001562 | response to protozoan | BP | 13 | 0 | 6 | 2.41E-05 |
| GO:0042101 | T cell receptor complex | CC | 13 | 0 | 6 | 2.41E-05 |
| GO:0045637 | regulation of myeloid cell differentiation | BP | 182 | 4 | 24 | 2.42E-05 |
| GO:0061756 | leukocyte adhesion to vascular endothelial cell | BP | 32 | 2 | 9 | 2.49E-05 |
| GO:0003823 | antigen binding | MF | 25 | 2 | 8 | 2.50E-05 |
| GO:0031663 | lipopolysaccharide-mediated signaling pathway | BP | 48 | 1 | 11 | 2.66E-05 |
| GO:0051241 | negative regulation of multicellular organismal process | BP | 930 | 88 | 77 | 2.67E-05 |
| GO:0043547 | positive regulation of GTPase activity | BP | 335 | 18 | 36 | 2.80E-05 |
| GO:0043549 | regulation of kinase activity | BP | 720 | 46 | 63 | 2.98E-05 |
| GO:0032635 | interleukin-6 production | BP | 106 | 4 | 17 | 3.06E-05 |

| | | | | | | |
|---|---|---|---|---|---|---|
| GO:0007264 | small GTPase mediated signal transduction | BP | 517 | 38 | 49 | 3.23E-05 |
| GO:0019724 | B cell mediated immunity | BP | 86 | 0 | 15 | 3.24E-05 |
| GO:2000401 | regulation of lymphocyte migration | BP | 49 | 1 | 11 | 3.27E-05 |
| GO:0002704 | negative regulation of leukocyte mediated immunity | BP | 33 | 0 | 9 | 3.27E-05 |
| GO:0031224 | intrinsic component of membrane | CC | 3479 | 366 | 227 | 3.35E-05 |
| GO:0032733 | positive regulation of interleukin-10 production | BP | 26 | 0 | 8 | 3.45E-05 |
| GO:0042105 | alpha-beta T cell receptor complex | CC | 5 | 0 | 4 | 3.46E-05 |
| GO:0048015 | phosphatidylinositol-mediated signaling | BP | 151 | 15 | 21 | 3.48E-05 |
| GO:0048585 | negative regulation of response to stimulus | BP | 1319 | 96 | 101 | 3.64E-05 |
| GO:0032675 | regulation of interleukin-6 production | BP | 97 | 3 | 16 | 3.65E-05 |
| GO:2000026 | regulation of multicellular organismal development | BP | 1597 | 178 | 118 | 3.66E-05 |
| GO:0071887 | leukocyte apoptotic process | BP | 87 | 3 | 15 | 3.73E-05 |
| GO:0032700 | negative regulation of interleukin-17 production | BP | 9 | 1 | 5 | 3.95E-05 |
| GO:0001916 | positive regulation of T cell mediated cytotoxicity | BP | 9 | 1 | 5 | 3.95E-05 |
| GO:0045059 | positive thymic T cell selection | BP | 9 | 0 | 5 | 3.95E-05 |
| GO:1902713 | regulation of interferon-gamma secretion | BP | 9 | 0 | 5 | 3.95E-05 |
| GO:0001914 | regulation of T cell mediated cytotoxicity | BP | 14 | 1 | 6 | 4.03E-05 |
| GO:0051050 | positive regulation of transport | BP | 773 | 83 | 66 | 4.11E-05 |
| GO:0050871 | positive regulation of B cell activation | BP | 59 | 1 | 12 | 4.13E-05 |
| GO:0042327 | positive regulation of phosphorylation | BP | 804 | 58 | 68 | 4.21E-05 |
| GO:0032613 | interleukin-10 production | BP | 42 | 0 | 10 | 4.34E-05 |
| GO:0045859 | regulation of protein kinase activity | BP | 655 | 41 | 58 | 4.46E-05 |
| GO:0099503 | secretory vesicle | CC | 760 | 49 | 65 | 4.48E-05 |
| GO:0055074 | calcium ion homeostasis | BP | 356 | 48 | 37 | 4.49E-05 |
| GO:0048017 | inositol lipid-mediated signaling | BP | 154 | 15 | 21 | 4.68E-05 |
| GO:0002685 | regulation of leukocyte migration | BP | 154 | 11 | 21 | 4.68E-05 |
| GO:0019955 | cytokine binding | MF | 99 | 11 | 16 | 4.72E-05 |
| GO:0014066 | regulation of phosphatidylinositol 3-kinase signaling | BP | 99 | 8 | 16 | 4.72E-05 |
| GO:0010942 | positive regulation of cell death | BP | 584 | 34 | 53 | 5.02E-05 |
| GO:0038083 | peptidyl-tyrosine autophosphorylation | BP | 35 | 1 | 9 | 5.46E-05 |
| GO:2000404 | regulation of T cell migration | BP | 35 | 0 | 9 | 5.46E-05 |
| GO:0050777 | negative regulation of immune response | BP | 111 | 3 | 17 | 5.60E-05 |
| GO:0070062 | extracellular exosome | CC | 1646 | 110 | 120 | 5.64E-05 |
| GO:0045121 | membrane raft | CC | 267 | 31 | 30 | 5.69E-05 |
| GO:0055065 | metal ion homeostasis | BP | 486 | 59 | 46 | 5.83E-05 |
| GO:0043230 | extracellular organelle | CC | 1664 | 115 | 121 | 5.85E-05 |
| GO:0050766 | positive regulation of phagocytosis | BP | 52 | 2 | 11 | 5.88E-05 |
| GO:0006874 | cellular calcium ion homeostasis | BP | 347 | 46 | 36 | 5.89E-05 |
| GO:0051338 | regulation of transferase activity | BP | 813 | 51 | 68 | 5.95E-05 |
| GO:0000082 | G1/S transition of mitotic cell cycle | BP | 217 | 4 | 26 | 6.02E-05 |
| GO:0098857 | membrane microdomain | CC | 268 | 31 | 30 | 6.10E-05 |
| GO:0004869 | cysteine-type endopeptidase inhibitor activity | MF | 28 | 1 | 8 | 6.26E-05 |
| GO:0008305 | integrin complex | CC | 28 | 4 | 8 | 6.26E-05 |
| GO:0014065 | phosphatidylinositol 3-kinase signaling | BP | 123 | 12 | 18 | 6.33E-05 |
| GO:0016493 | C-C chemokine receptor activity | MF | 15 | 0 | 6 | 6.43E-05 |
| GO:1990869 | cellular response to chemokine | BP | 62 | 4 | 12 | 6.92E-05 |
| GO:1990868 | response to chemokine | BP | 62 | 4 | 12 | 6.92E-05 |
| GO:0043300 | regulation of leukocyte degranulation | BP | 36 | 1 | 9 | 6.94E-05 |
| GO:0000793 | condensed chromosome | CC | 170 | 0 | 22 | 6.97E-05 |
| GO:0031325 | positive regulation of cellular metabolic process | BP | 2612 | 159 | 176 | 7.06E-05 |
| GO:0072507 | divalent inorganic cation homeostasis | BP | 378 | 50 | 38 | 7.30E-05 |
| GO:0044433 | cytoplasmic vesicle part | CC | 1213 | 66 | 93 | 7.37E-05 |
| GO:0002702 | positive regulation of production of molecular mediator of immune | BP | 72 | 3 | 13 | 7.43E-05 |
| GO:0048731 | system development | BP | 3556 | 373 | 229 | 7.45E-05 |
| GO:0042035 | regulation of cytokine biosynthetic process | BP | 82 | 1 | 14 | 7.52E-05 |
| GO:0000727 | double-strand break repair via break-induced replication | BP | 10 | 0 | 5 | 7.56E-05 |
| GO:0043235 | receptor complex | CC | 311 | 50 | 33 | 7.69E-05 |
| GO:0032735 | positive regulation of interleukin-12 production | BP | 29 | 2 | 8 | 8.26E-05 |
| GO:0034142 | toll-like receptor 4 signaling pathway | BP | 29 | 1 | 8 | 8.26E-05 |
| GO:0007249 | I-kappaB kinase/NF-kappaB signaling | BP | 234 | 9 | 27 | 8.41E-05 |
| GO:0033260 | nuclear DNA replication | BP | 54 | 0 | 11 | 8.48E-05 |
| GO:0050901 | leukocyte tethering or rolling | BP | 22 | 2 | 7 | 8.51E-05 |
| GO:2000515 | negative regulation of CD4-positive, alpha-beta T cell activation | BP | 22 | 0 | 7 | 8.51E-05 |
| GO:0050855 | regulation of B cell receptor signaling pathway | BP | 22 | 0 | 7 | 8.51E-05 |

| | | | | | | |
|---|---|---|---|---|---|---|
| GO:0072538 | T-helper 17 type immune response | BP | 22 | 0 | 7 | 8.51E-05 |
| GO:1903561 | extracellular vesicle | CC | 1662 | 115 | 120 | 8.59E-05 |
| GO:1990266 | neutrophil migration | BP | 73 | 4 | 13 | 8.62E-05 |
| GO:0032101 | regulation of response to external stimulus | BP | 611 | 43 | 54 | 8.63E-05 |
| GO:0019900 | kinase binding | MF | 641 | 44 | 56 | 8.72E-05 |
| GO:0044786 | cell cycle DNA replication | BP | 64 | 0 | 12 | 9.56E-05 |
| GO:0060333 | interferon-gamma-mediated signaling pathway | BP | 64 | 1 | 12 | 9.56E-05 |
| GO:0060205 | cytoplasmic vesicle lumen | CC | 275 | 6 | 30 | 9.79E-05 |
| GO:0042802 | identical protein binding | MF | 1369 | 98 | 102 | 9.80E-05 |
| GO:0045076 | regulation of interleukin-2 biosynthetic process | BP | 16 | 0 | 6 | 9.84E-05 |
| GO:0023024 | MHC class I protein complex binding | MF | 6 | 0 | 4 | 9.96E-05 |
| GO:0050764 | regulation of phagocytosis | BP | 74 | 3 | 13 | 9.97E-05 |
| GO:0043086 | negative regulation of catalytic activity | BP | 585 | 49 | 52 | 1.01E-04 |
| GO:0008047 | enzyme activator activity | MF | 426 | 19 | 41 | 1.02E-04 |
| GO:0098589 | membrane region | CC | 276 | 31 | 30 | 1.05E-04 |
| GO:0031983 | vesicle lumen | CC | 276 | 6 | 30 | 1.05E-04 |
| GO:0072593 | reactive oxygen species metabolic process | BP | 212 | 15 | 25 | 1.08E-04 |
| GO:0002369 | T cell cytokine production | BP | 30 | 1 | 8 | 1.08E-04 |
| GO:0051247 | positive regulation of protein metabolic process | BP | 1340 | 82 | 100 | 1.09E-04 |
| GO:0016310 | phosphorylation | BP | 1891 | 117 | 133 | 1.11E-04 |
| GO:0016064 | immunoglobulin mediated immune response | BP | 85 | 0 | 14 | 1.13E-04 |
| GO:1902106 | negative regulation of leukocyte differentiation | BP | 75 | 3 | 13 | 1.15E-04 |
| GO:0035579 | specific granule membrane | CC | 75 | 4 | 13 | 1.15E-04 |
| GO:0043065 | positive regulation of apoptotic process | BP | 544 | 33 | 49 | 1.16E-04 |
| GO:0022616 | DNA strand elongation | BP | 23 | 0 | 7 | 1.17E-04 |
| GO:0070664 | negative regulation of leukocyte proliferation | BP | 56 | 3 | 11 | 1.20E-04 |
| GO:0044427 | chromosomal part | CC | 740 | 15 | 62 | 1.21E-04 |
| GO:0051301 | cell division | BP | 487 | 20 | 45 | 1.23E-04 |
| GO:0034612 | response to tumor necrosis factor | BP | 240 | 13 | 27 | 1.29E-04 |
| GO:0071621 | granulocyte chemotaxis | BP | 76 | 4 | 13 | 1.32E-04 |
| GO:0006268 | DNA unwinding involved in DNA replication | BP | 11 | 0 | 5 | 1.33E-04 |
| GO:0071682 | endocytic vesicle lumen | CC | 11 | 1 | 5 | 1.33E-04 |
| GO:0002418 | immune response to tumor cell | BP | 11 | 1 | 5 | 1.33E-04 |
| GO:0006310 | DNA recombination | BP | 215 | 2 | 25 | 1.35E-04 |
| GO:0051056 | regulation of small GTPase mediated signal transduction | BP | 307 | 25 | 32 | 1.37E-04 |
| GO:0032715 | negative regulation of interleukin-6 production | BP | 31 | 0 | 8 | 1.38E-04 |
| GO:2000516 | positive regulation of CD4-positive, alpha-beta T cell activation | BP | 31 | 0 | 8 | 1.38E-04 |
| GO:0098636 | protein complex involved in cell adhesion | CC | 31 | 4 | 8 | 1.38E-04 |
| GO:0007204 | positive regulation of cytosolic calcium ion concentration | BP | 241 | 35 | 27 | 1.39E-04 |
| GO:0043068 | positive regulation of programmed cell death | BP | 548 | 33 | 49 | 1.39E-04 |
| GO:0016513 | core-binding factor complex | CC | 3 | 0 | 3 | 1.40E-04 |
| GO:1903238 | positive regulation of leukocyte tethering or rolling | BP | 3 | 0 | 3 | 1.40E-04 |
| GO:1902565 | positive regulation of neutrophil activation | BP | 3 | 0 | 3 | 1.40E-04 |
| GO:0043315 | positive regulation of neutrophil degranulation | BP | 3 | 0 | 3 | 1.40E-04 |
| GO:0031401 | positive regulation of protein modification process | BP | 946 | 65 | 75 | 1.40E-04 |
| GO:0000280 | nuclear division | BP | 321 | 9 | 33 | 1.41E-04 |
| GO:0019957 | C-C chemokine binding | MF | 17 | 0 | 6 | 1.45E-04 |
| GO:0034113 | heterotypic cell-cell adhesion | BP | 48 | 4 | 10 | 1.46E-04 |
| GO:0005070 | SH3/SH2 adaptor activity | MF | 48 | 4 | 10 | 1.46E-04 |
| GO:0072503 | cellular divalent inorganic cation homeostasis | BP | 363 | 46 | 36 | 1.48E-04 |
| GO:0005694 | chromosome | CC | 838 | 15 | 68 | 1.48E-04 |
| GO:0006796 | phosphate-containing compound metabolic process | BP | 2631 | 172 | 175 | 1.52E-04 |
| GO:0045597 | positive regulation of cell differentiation | BP | 762 | 85 | 63 | 1.56E-04 |
| GO:0046633 | alpha-beta T cell proliferation | BP | 24 | 0 | 7 | 1.58E-04 |
| GO:0019956 | chemokine binding | MF | 24 | 0 | 7 | 1.58E-04 |
| GO:0002888 | positive regulation of myeloid leukocyte mediated immunity | BP | 24 | 1 | 7 | 1.58E-04 |
| GO:0002724 | regulation of T cell cytokine production | BP | 24 | 1 | 7 | 1.58E-04 |
| GO:0044843 | cell cycle G1/S phase transition | BP | 230 | 5 | 26 | 1.58E-04 |
| GO:0051092 | positive regulation of NF-kappaB transcription factor activity | BP | 132 | 8 | 18 | 1.60E-04 |
| GO:0006793 | phosphorus metabolic process | BP | 2653 | 173 | 176 | 1.66E-04 |
| GO:0046638 | positive regulation of alpha-beta T cell differentiation | BP | 40 | 1 | 9 | 1.67E-04 |
| GO:0051924 | regulation of calcium ion transport | BP | 193 | 26 | 23 | 1.71E-04 |
| GO:0051480 | regulation of cytosolic calcium ion concentration | BP | 271 | 41 | 29 | 1.78E-04 |
| GO:0031399 | regulation of protein modification process | BP | 1458 | 94 | 106 | 1.83E-04 |

| | | | | | | |
|---|---|---|---|---|---|---|
| GO:0016021 | integral component of membrane | CC | 3371 | 343 | 216 | 1.86E-04 |
| GO:0000777 | condensed chromosome kinetochore | CC | 89 | 0 | 14 | 1.87E-04 |
| GO:0045595 | regulation of cell differentiation | BP | 1409 | 152 | 103 | 1.87E-04 |
| GO:1903556 | negative regulation of tumor necrosis factor superfamily cytokine p | BP | 41 | 1 | 9 | 2.05E-04 |
| GO:0002763 | positive regulation of myeloid leukocyte differentiation | BP | 41 | 0 | 9 | 2.05E-04 |
| GO:0004950 | chemokine receptor activity | MF | 18 | 1 | 6 | 2.09E-04 |
| GO:0001637 | G protein-coupled chemoattractant receptor activity | MF | 18 | 1 | 6 | 2.09E-04 |
| GO:0048523 | negative regulation of cellular process | BP | 3703 | 259 | 234 | 2.09E-04 |
| GO:0001891 | phagocytic cup | CC | 25 | 1 | 7 | 2.09E-04 |
| GO:0009620 | response to fungus | BP | 25 | 0 | 7 | 2.09E-04 |
| GO:0042089 | cytokine biosynthetic process | BP | 90 | 2 | 14 | 2.11E-04 |
| GO:0000779 | condensed chromosome, centromeric region | CC | 101 | 0 | 15 | 2.15E-04 |
| GO:0002687 | positive regulation of leukocyte migration | BP | 101 | 5 | 15 | 2.15E-04 |
| GO:0023014 | signal transduction by protein phosphorylation | BP | 740 | 56 | 61 | 2.15E-04 |
| GO:0098802 | plasma membrane receptor complex | CC | 147 | 23 | 19 | 2.17E-04 |
| GO:1902969 | mitotic DNA replication | BP | 12 | 0 | 5 | 2.18E-04 |
| GO:0045086 | positive regulation of interleukin-2 biosynthetic process | BP | 12 | 0 | 5 | 2.18E-04 |
| GO:0055080 | cation homeostasis | BP | 544 | 62 | 48 | 2.21E-04 |
| GO:0042107 | cytokine metabolic process | BP | 91 | 2 | 14 | 2.37E-04 |
| GO:1903409 | reactive oxygen species biosynthetic process | BP | 91 | 9 | 14 | 2.37E-04 |
| GO:0018212 | peptidyl-tyrosine modification | BP | 303 | 26 | 31 | 2.43E-04 |
| GO:0007186 | G protein-coupled receptor signaling pathway | BP | 591 | 69 | 51 | 2.43E-04 |
| GO:0097529 | myeloid leukocyte migration | BP | 137 | 11 | 18 | 2.57E-04 |
| GO:0006875 | cellular metal ion homeostasis | BP | 431 | 50 | 40 | 2.65E-04 |
| GO:0045639 | positive regulation of myeloid cell differentiation | BP | 71 | 0 | 12 | 2.67E-04 |
| GO:0000165 | MAPK cascade | BP | 731 | 56 | 60 | 2.71E-04 |
| GO:0044364 | disruption of cells of other organism | BP | 26 | 0 | 7 | 2.74E-04 |
| GO:0031640 | killing of cells of other organism | BP | 26 | 0 | 7 | 2.74E-04 |
| GO:2000406 | positive regulation of T cell migration | BP | 26 | 0 | 7 | 2.74E-04 |
| GO:0030890 | positive regulation of B cell proliferation | BP | 34 | 1 | 8 | 2.77E-04 |
| GO:0051090 | regulation of DNA-binding transcription factor activity | BP | 347 | 21 | 34 | 2.81E-04 |
| GO:1903047 | mitotic cell cycle process | BP | 748 | 21 | 61 | 2.87E-04 |
| GO:0002407 | dendritic cell chemotaxis | BP | 19 | 1 | 6 | 2.91E-04 |
| GO:0042094 | interleukin-2 biosynthetic process | BP | 19 | 1 | 6 | 2.91E-04 |
| GO:0035586 | purinergic receptor activity | MF | 19 | 0 | 6 | 2.91E-04 |
| GO:0050672 | negative regulation of lymphocyte proliferation | BP | 52 | 3 | 10 | 2.92E-04 |
| GO:0032945 | negative regulation of mononuclear cell proliferation | BP | 52 | 3 | 10 | 2.92E-04 |
| GO:0050801 | ion homeostasis | BP | 611 | 67 | 52 | 2.96E-04 |
| GO:0098771 | inorganic ion homeostasis | BP | 551 | 63 | 48 | 2.97E-04 |
| GO:0004715 | non-membrane spanning protein tyrosine kinase activity | MF | 43 | 0 | 9 | 3.00E-04 |
| GO:0070228 | regulation of lymphocyte apoptotic process | BP | 43 | 2 | 9 | 3.00E-04 |
| GO:0000819 | sister chromatid segregation | BP | 163 | 0 | 20 | 3.03E-04 |
| GO:0000070 | mitotic sister chromatid segregation | BP | 139 | 0 | 18 | 3.08E-04 |
| GO:0033673 | negative regulation of kinase activity | BP | 214 | 13 | 24 | 3.15E-04 |
| GO:0065008 | regulation of biological quality | BP | 3132 | 307 | 201 | 3.23E-04 |
| GO:0043011 | myeloid dendritic cell differentiation | BP | 13 | 0 | 5 | 3.39E-04 |
| GO:0046641 | positive regulation of alpha-beta T cell proliferation | BP | 13 | 0 | 5 | 3.39E-04 |
| GO:1904996 | positive regulation of leukocyte adhesion to vascular endothelial ce | BP | 13 | 0 | 5 | 3.39E-04 |
| GO:0032633 | interleukin-4 production | BP | 27 | 1 | 7 | 3.53E-04 |
| GO:0045730 | respiratory burst | BP | 27 | 1 | 7 | 3.53E-04 |
| GO:0042092 | type 2 immune response | BP | 27 | 0 | 7 | 3.53E-04 |
| GO:0051348 | negative regulation of transferase activity | BP | 242 | 15 | 26 | 3.56E-04 |
| GO:0005657 | replication fork | CC | 63 | 0 | 11 | 3.57E-04 |
| GO:0002381 | immunoglobulin production involved in immunoglobulin mediated | BP | 44 | 0 | 9 | 3.60E-04 |
| GO:1902532 | negative regulation of intracellular signal transduction | BP | 438 | 23 | 40 | 3.68E-04 |
| GO:0042326 | negative regulation of phosphorylation | BP | 381 | 26 | 36 | 3.78E-04 |
| GO:0019901 | protein kinase binding | MF | 572 | 34 | 49 | 3.80E-04 |
| GO:0043122 | regulation of I-kappaB kinase/NF-kappaB signaling | BP | 204 | 9 | 23 | 3.86E-04 |
| GO:0099513 | polymeric cytoskeletal fiber | CC | 498 | 44 | 44 | 3.91E-04 |
| GO:0035591 | signaling adaptor activity | MF | 74 | 6 | 12 | 3.97E-04 |
| GO:0046639 | negative regulation of alpha-beta T cell differentiation | BP | 20 | 1 | 6 | 3.98E-04 |
| GO:0032753 | positive regulation of interleukin-4 production | BP | 20 | 0 | 6 | 3.98E-04 |
| GO:0072539 | T-helper 17 cell differentiation | BP | 20 | 0 | 6 | 3.98E-04 |
| GO:1903707 | negative regulation of hemopoiesis | BP | 107 | 5 | 15 | 4.08E-04 |

| | | | | | | |
|---|---|---|---|---|---|---|
| GO:0007275 | multicellular organism development | BP | 3945 | 389 | 245 | 4.20E-04 |
| GO:0042492 | gamma-delta T cell differentiation | BP | 8 | 0 | 4 | 4.27E-04 |
| GO:0002839 | positive regulation of immune response to tumor cell | BP | 8 | 1 | 4 | 4.27E-04 |
| GO:0002836 | positive regulation of response to tumor cell | BP | 8 | 1 | 4 | 4.27E-04 |
| GO:0006590 | thyroid hormone generation | BP | 8 | 1 | 4 | 4.27E-04 |
| GO:0007059 | chromosome segregation | BP | 259 | 2 | 27 | 4.46E-04 |
| GO:1901701 | cellular response to oxygen-containing compound | BP | 903 | 78 | 70 | 4.48E-04 |
| GO:0018108 | peptidyl-tyrosine phosphorylation | BP | 300 | 26 | 30 | 4.48E-04 |
| GO:0048285 | organelle fission | BP | 356 | 10 | 34 | 4.49E-04 |
| GO:0050798 | activated T cell proliferation | BP | 28 | 1 | 7 | 4.50E-04 |
| GO:0071353 | cellular response to interleukin-4 | BP | 28 | 1 | 7 | 4.50E-04 |
| GO:0097028 | dendritic cell differentiation | BP | 28 | 0 | 7 | 4.50E-04 |
| GO:0042554 | superoxide anion generation | BP | 28 | 3 | 7 | 4.50E-04 |
| GO:0003697 | single-stranded DNA binding | MF | 97 | 1 | 14 | 4.66E-04 |
| GO:0051246 | regulation of protein metabolic process | BP | 2197 | 125 | 147 | 4.69E-04 |
| GO:0050708 | regulation of protein secretion | BP | 357 | 30 | 34 | 4.73E-04 |
| GO:0052547 | regulation of peptidase activity | BP | 301 | 20 | 30 | 4.74E-04 |
| GO:0032502 | developmental process | BP | 4618 | 444 | 281 | 4.77E-04 |
| GO:0012506 | vesicle membrane | CC | 639 | 48 | 53 | 4.82E-04 |
| GO:0010959 | regulation of metal ion transport | BP | 302 | 51 | 30 | 5.01E-04 |
| GO:0035743 | CD4-positive, alpha-beta T cell cytokine production | BP | 14 | 0 | 5 | 5.04E-04 |
| GO:0050665 | hydrogen peroxide biosynthetic process | BP | 14 | 1 | 5 | 5.04E-04 |
| GO:0038187 | pattern recognition receptor activity | MF | 14 | 1 | 5 | 5.04E-04 |
| GO:0008329 | signaling pattern recognition receptor activity | MF | 14 | 1 | 5 | 5.04E-04 |
| GO:0032879 | regulation of localization | BP | 2184 | 239 | 146 | 5.12E-04 |
| GO:0050829 | defense response to Gram-negative bacterium | BP | 37 | 2 | 8 | 5.14E-04 |
| GO:0050731 | positive regulation of peptidyl-tyrosine phosphorylation | BP | 145 | 15 | 18 | 5.18E-04 |
| GO:0002688 | regulation of leukocyte chemotaxis | BP | 87 | 6 | 13 | 5.24E-04 |
| GO:0005488 | binding | MF | 10954 | 706 | 600 | 5.25E-04 |
| GO:0036336 | dendritic cell migration | BP | 21 | 1 | 6 | 5.33E-04 |
| GO:0046640 | regulation of alpha-beta T cell proliferation | BP | 21 | 0 | 6 | 5.33E-04 |
| GO:0046696 | lipopolysaccharide receptor complex | CC | 4 | 0 | 3 | 5.37E-04 |
| GO:0031580 | membrane raft distribution | BP | 4 | 0 | 3 | 5.37E-04 |
| GO:0032393 | MHC class I receptor activity | MF | 4 | 0 | 3 | 5.37E-04 |
| GO:2000403 | positive regulation of lymphocyte migration | BP | 29 | 0 | 7 | 5.67E-04 |
| GO:0070670 | response to interleukin-4 | BP | 29 | 1 | 7 | 5.67E-04 |
| GO:0097190 | apoptotic signaling pathway | BP | 507 | 18 | 44 | 5.71E-04 |
| GO:0071356 | cellular response to tumor necrosis factor | BP | 223 | 11 | 24 | 5.77E-04 |
| GO:0098813 | nuclear chromosome segregation | BP | 210 | 2 | 23 | 5.83E-04 |
| GO:1901700 | response to oxygen-containing compound | BP | 1287 | 123 | 93 | 5.84E-04 |
| GO:0051091 | positive regulation of DNA-binding transcription factor activity | BP | 211 | 14 | 23 | 6.24E-04 |
| GO:0043270 | positive regulation of ion transport | BP | 211 | 33 | 23 | 6.24E-04 |
| GO:0140014 | mitotic nuclear division | BP | 238 | 7 | 25 | 6.41E-04 |
| GO:0052548 | regulation of endopeptidase activity | BP | 279 | 19 | 28 | 6.49E-04 |
| GO:0006469 | negative regulation of protein kinase activity | BP | 199 | 10 | 22 | 6.71E-04 |
| GO:0001933 | negative regulation of protein phosphorylation | BP | 350 | 20 | 33 | 6.79E-04 |
| GO:0032703 | negative regulation of interleukin-2 production | BP | 22 | 1 | 6 | 7.01E-04 |
| GO:0006297 | nucleotide-excision repair, DNA gap filling | BP | 22 | 0 | 6 | 7.01E-04 |
| GO:0002825 | regulation of T-helper 1 type immune response | BP | 22 | 0 | 6 | 7.01E-04 |
| GO:0044092 | negative regulation of molecular function | BP | 838 | 69 | 65 | 7.07E-04 |
| GO:0048519 | negative regulation of biological process | BP | 4162 | 287 | 255 | 7.10E-04 |
| GO:0030659 | cytoplasmic vesicle membrane | CC | 619 | 47 | 51 | 7.14E-04 |
| GO:0060249 | anatomical structure homeostasis | BP | 337 | 15 | 32 | 7.23E-04 |
| GO:0001848 | complement binding | MF | 15 | 1 | 5 | 7.24E-04 |
| GO:0001779 | natural killer cell differentiation | BP | 15 | 0 | 5 | 7.24E-04 |
| GO:0016502 | nucleotide receptor activity | MF | 15 | 0 | 5 | 7.24E-04 |
| GO:0001614 | purinergic nucleotide receptor activity | MF | 15 | 0 | 5 | 7.24E-04 |
| GO:0002347 | response to tumor cell | BP | 15 | 1 | 5 | 7.24E-04 |
| GO:0002507 | tolerance induction | BP | 15 | 0 | 5 | 7.24E-04 |
| GO:0006801 | superoxide metabolic process | BP | 58 | 7 | 10 | 7.27E-04 |
| GO:0097530 | granulocyte migration | BP | 90 | 4 | 13 | 7.30E-04 |
| GO:0070486 | leukocyte aggregation | BP | 9 | 0 | 4 | 7.38E-04 |
| GO:0002323 | natural killer cell activation involved in immune response | BP | 9 | 0 | 4 | 7.38E-04 |
| GO:0043383 | negative T cell selection | BP | 9 | 1 | 4 | 7.38E-04 |

| | | | | | | |
|---|---|---|---|---|---|---|
| GO:0010912 | positive regulation of isomerase activity | BP | 9 | 0 | 4 | 7.38E-04 |
| GO:0002837 | regulation of immune response to tumor cell | BP | 9 | 1 | 4 | 7.38E-04 |
| GO:0010911 | regulation of isomerase activity | BP | 9 | 0 | 4 | 7.38E-04 |
| GO:0002834 | regulation of response to tumor cell | BP | 9 | 1 | 4 | 7.38E-04 |
| GO:0045807 | positive regulation of endocytosis | BP | 125 | 8 | 16 | 7.43E-04 |
| GO:0048856 | anatomical structure development | BP | 4314 | 418 | 263 | 7.56E-04 |
| GO:0005884 | actin filament | CC | 102 | 17 | 14 | 7.80E-04 |
| GO:0030316 | osteoclast differentiation | BP | 69 | 2 | 11 | 7.96E-04 |
| GO:0032270 | positive regulation of cellular protein metabolic process | BP | 1250 | 75 | 90 | 8.08E-04 |
| GO:0006260 | DNA replication | BP | 242 | 1 | 25 | 8.18E-04 |
| GO:0010604 | positive regulation of macromolecule metabolic process | BP | 2615 | 163 | 169 | 8.51E-04 |
| GO:0030003 | cellular cation homeostasis | BP | 487 | 51 | 42 | 8.57E-04 |
| GO:0032720 | negative regulation of tumor necrosis factor production | BP | 40 | 1 | 8 | 8.92E-04 |
| GO:0014068 | positive regulation of phosphatidylinositol 3-kinase signaling | BP | 70 | 7 | 11 | 9.01E-04 |
| GO:0061135 | endopeptidase regulator activity | MF | 92 | 9 | 13 | 9.01E-04 |
| GO:0008009 | chemokine activity | MF | 23 | 1 | 6 | 9.08E-04 |
| GO:0002707 | negative regulation of lymphocyte mediated immunity | BP | 23 | 0 | 6 | 9.08E-04 |
| GO:2000108 | positive regulation of leukocyte apoptotic process | BP | 23 | 0 | 6 | 9.08E-04 |
| GO:0032673 | regulation of interleukin-4 production | BP | 23 | 0 | 6 | 9.08E-04 |
| GO:0042534 | regulation of tumor necrosis factor biosynthetic process | BP | 23 | 0 | 6 | 9.08E-04 |
| GO:0042533 | tumor necrosis factor biosynthetic process | BP | 23 | 0 | 6 | 9.08E-04 |
| GO:0019899 | enzyme binding | MF | 1918 | 109 | 129 | 9.23E-04 |
| GO:0045936 | negative regulation of phosphate metabolic process | BP | 474 | 32 | 41 | 9.29E-04 |
| GO:0050920 | regulation of chemotaxis | BP | 165 | 13 | 19 | 9.34E-04 |
| GO:0033209 | tumor necrosis factor-mediated signaling pathway | BP | 140 | 1 | 17 | 9.42E-04 |
| GO:0016444 | somatic cell DNA recombination | BP | 50 | 0 | 9 | 9.64E-04 |
| GO:0002562 | somatic diversification of immune receptors via germline recombin | BP | 50 | 0 | 9 | 9.64E-04 |
| GO:0010563 | negative regulation of phosphorus metabolic process | BP | 475 | 32 | 41 | 9.68E-04 |
| GO:0033631 | cell-cell adhesion mediated by integrin | BP | 16 | 0 | 5 | 1.01E-03 |
| GO:0023023 | MHC protein complex binding | MF | 16 | 0 | 5 | 1.01E-03 |
| GO:0002827 | positive regulation of T-helper 1 type immune response | BP | 16 | 0 | 5 | 1.01E-03 |
| GO:0035590 | purinergic nucleotide receptor signaling pathway | BP | 16 | 0 | 5 | 1.01E-03 |
| GO:0070374 | positive regulation of ERK1 and ERK2 cascade | BP | 141 | 11 | 17 | 1.02E-03 |
| GO:0002791 | regulation of peptide secretion | BP | 373 | 32 | 34 | 1.03E-03 |
| GO:0000724 | double-strand break repair via homologous recombination | BP | 105 | 1 | 14 | 1.04E-03 |
| GO:0071349 | cellular response to interleukin-12 | BP | 41 | 1 | 8 | 1.06E-03 |
| GO:0090342 | regulation of cell aging | BP | 41 | 0 | 8 | 1.06E-03 |
| GO:0002820 | negative regulation of adaptive immune response | BP | 32 | 0 | 7 | 1.07E-03 |
| GO:0051173 | positive regulation of nitrogen compound metabolic process | BP | 2505 | 154 | 162 | 1.10E-03 |
| GO:0051928 | positive regulation of calcium ion transport | BP | 94 | 11 | 13 | 1.11E-03 |
| GO:0006873 | cellular ion homeostasis | BP | 494 | 53 | 42 | 1.13E-03 |
| GO:0009306 | protein secretion | BP | 449 | 45 | 39 | 1.14E-03 |
| GO:0000725 | recombinational repair | BP | 106 | 1 | 14 | 1.15E-03 |
| GO:0098761 | cellular response to interleukin-7 | BP | 24 | 1 | 6 | 1.16E-03 |
| GO:0003688 | DNA replication origin binding | MF | 24 | 0 | 6 | 1.16E-03 |
| GO:0002828 | regulation of type 2 immune response | BP | 24 | 0 | 6 | 1.16E-03 |
| GO:0098760 | response to interleukin-7 | BP | 24 | 1 | 6 | 1.16E-03 |
| GO:0030246 | carbohydrate binding | MF | 181 | 15 | 20 | 1.16E-03 |
| GO:0140097 | catalytic activity, acting on DNA | MF | 181 | 2 | 20 | 1.16E-03 |
| GO:0000228 | nuclear chromosome | CC | 466 | 11 | 40 | 1.23E-03 |
| GO:0034774 | secretory granule lumen | CC | 263 | 6 | 26 | 1.24E-03 |
| GO:0005126 | cytokine receptor binding | MF | 169 | 11 | 19 | 1.25E-03 |
| GO:0042130 | negative regulation of T cell proliferation | BP | 42 | 2 | 8 | 1.25E-03 |
| GO:0006911 | phagocytosis, engulfment | BP | 42 | 3 | 8 | 1.25E-03 |
| GO:0033628 | regulation of cell adhesion mediated by integrin | BP | 42 | 2 | 8 | 1.25E-03 |
| GO:0070671 | response to interleukin-12 | BP | 42 | 1 | 8 | 1.25E-03 |
| GO:0051179 | localization | BP | 5115 | 395 | 304 | 1.26E-03 |
| GO:0001851 | complement component C3b binding | MF | 5 | 0 | 3 | 1.29E-03 |
| GO:0051665 | membrane raft localization | BP | 5 | 0 | 3 | 1.29E-03 |
| GO:2000348 | regulation of CD40 signaling pathway | BP | 5 | 0 | 3 | 1.29E-03 |
| GO:2000501 | regulation of natural killer cell chemotaxis | BP | 5 | 0 | 3 | 1.29E-03 |
| GO:0070432 | regulation of nucleotide-binding oligomerization domain containing | BP | 5 | 0 | 3 | 1.29E-03 |
| GO:2000407 | regulation of T cell extravasation | BP | 5 | 0 | 3 | 1.29E-03 |
| GO:0042379 | chemokine receptor binding | MF | 33 | 1 | 7 | 1.29E-03 |

| | | | | | | |
|---|---|---|---|---|---|---|
| GO:0045005 | DNA-dependent DNA replication maintenance of fidelity | BP | 33 | 0 | 7 | 1.29E-03 |
| GO:0001772 | immunological synapse | CC | 33 | 0 | 7 | 1.29E-03 |
| GO:0034122 | negative regulation of toll-like receptor signaling pathway | BP | 33 | 2 | 7 | 1.29E-03 |
| GO:0050714 | positive regulation of protein secretion | BP | 196 | 10 | 21 | 1.32E-03 |
| GO:0070371 | ERK1 and ERK2 cascade | BP | 223 | 21 | 23 | 1.33E-03 |
| GO:0033554 | cellular response to stress | BP | 1730 | 74 | 117 | 1.36E-03 |
| GO:0043138 | 3'-5' DNA helicase activity | MF | 17 | 0 | 5 | 1.37E-03 |
| GO:0001530 | lipopolysaccharide binding | MF | 17 | 0 | 5 | 1.37E-03 |
| GO:0043371 | negative regulation of CD4-positive, alpha-beta T cell differentiatio | BP | 17 | 0 | 5 | 1.37E-03 |
| GO:0036151 | phosphatidylcholine acyl-chain remodeling | BP | 17 | 2 | 5 | 1.37E-03 |
| GO:0002726 | positive regulation of T cell cytokine production | BP | 17 | 0 | 5 | 1.37E-03 |
| GO:1904994 | regulation of leukocyte adhesion to vascular endothelial cell | BP | 17 | 0 | 5 | 1.37E-03 |
| GO:0060548 | negative regulation of cell death | BP | 764 | 44 | 59 | 1.38E-03 |
| GO:0071396 | cellular response to lipid | BP | 484 | 36 | 41 | 1.39E-03 |
| GO:0032418 | lysosome localization | BP | 63 | 2 | 10 | 1.41E-03 |
| GO:0043069 | negative regulation of programmed cell death | BP | 702 | 40 | 55 | 1.44E-03 |
| GO:0050858 | negative regulation of antigen receptor-mediated signaling pathway | BP | 25 | 1 | 6 | 1.46E-03 |
| GO:0043372 | positive regulation of CD4-positive, alpha-beta T cell differentiatio | BP | 25 | 0 | 6 | 1.46E-03 |
| GO:0035587 | purinergic receptor signaling pathway | BP | 25 | 1 | 6 | 1.46E-03 |
| GO:0046006 | regulation of activated T cell proliferation | BP | 25 | 1 | 6 | 1.46E-03 |
| GO:1903428 | positive regulation of reactive oxygen species biosynthetic process | BP | 43 | 5 | 8 | 1.47E-03 |
| GO:0051208 | sequestering of calcium ion | BP | 109 | 13 | 14 | 1.51E-03 |
| GO:0002437 | inflammatory response to antigenic stimulus | BP | 34 | 4 | 7 | 1.56E-03 |
| GO:0004866 | endopeptidase inhibitor activity | MF | 86 | 9 | 12 | 1.57E-03 |
| GO:0043066 | negative regulation of apoptotic process | BP | 689 | 38 | 54 | 1.58E-03 |
| GO:0000018 | regulation of DNA recombination | BP | 64 | 1 | 10 | 1.59E-03 |
| GO:0000278 | mitotic cell cycle | BP | 866 | 31 | 65 | 1.63E-03 |
| GO:0005125 | cytokine activity | MF | 98 | 5 | 13 | 1.63E-03 |
| GO:0045786 | negative regulation of cell cycle | BP | 535 | 19 | 44 | 1.69E-03 |
| GO:0043410 | positive regulation of MAPK cascade | BP | 399 | 31 | 35 | 1.69E-03 |
| GO:0061900 | glial cell activation | BP | 44 | 1 | 8 | 1.71E-03 |
| GO:0002720 | positive regulation of cytokine production involved in immune resp | BP | 44 | 2 | 8 | 1.71E-03 |
| GO:1901222 | regulation of NIK/NF-kappaB signaling | BP | 87 | 3 | 12 | 1.74E-03 |
| GO:0046629 | gamma-delta T cell activation | BP | 11 | 0 | 4 | 1.78E-03 |
| GO:0048291 | isotype switching to IgG isotypes | BP | 11 | 0 | 4 | 1.78E-03 |
| GO:0002710 | negative regulation of T cell mediated immunity | BP | 11 | 0 | 4 | 1.78E-03 |
| GO:0001846 | opsonin binding | MF | 11 | 1 | 4 | 1.78E-03 |
| GO:0002730 | regulation of dendritic cell cytokine production | BP | 11 | 0 | 4 | 1.78E-03 |
| GO:0002714 | positive regulation of B cell mediated immunity | BP | 26 | 0 | 6 | 1.81E-03 |
| GO:0002891 | positive regulation of immunoglobulin mediated immune response | BP | 26 | 0 | 6 | 1.81E-03 |
| GO:0032201 | telomere maintenance via semi-conservative replication | BP | 26 | 0 | 6 | 1.81E-03 |
| GO:0017000 | antibiotic biosynthetic process | BP | 18 | 2 | 5 | 1.81E-03 |
| GO:0019835 | cytolysis | BP | 18 | 1 | 5 | 1.81E-03 |
| GO:0002313 | mature B cell differentiation involved in immune response | BP | 18 | 1 | 5 | 1.81E-03 |
| GO:0050860 | negative regulation of T cell receptor signaling pathway | BP | 18 | 1 | 5 | 1.81E-03 |
| GO:0042535 | positive regulation of tumor necrosis factor biosynthetic process | BP | 18 | 0 | 5 | 1.81E-03 |
| GO:0043269 | regulation of ion transport | BP | 507 | 92 | 42 | 1.86E-03 |
| GO:2000377 | regulation of reactive oxygen species metabolic process | BP | 149 | 9 | 17 | 1.88E-03 |
| GO:0032103 | positive regulation of response to external stimulus | BP | 229 | 12 | 23 | 1.89E-03 |
| GO:0030545 | receptor regulator activity | MF | 257 | 29 | 25 | 1.91E-03 |
| GO:0050730 | regulation of peptidyl-tyrosine phosphorylation | BP | 202 | 19 | 21 | 1.92E-03 |
| GO:0070201 | regulation of establishment of protein localization | BP | 602 | 45 | 48 | 2.01E-03 |
| GO:0098687 | chromosomal region | CC | 287 | 1 | 27 | 2.09E-03 |
| GO:2000116 | regulation of cysteine-type endopeptidase activity | BP | 190 | 10 | 20 | 2.10E-03 |
| GO:0051272 | positive regulation of cellular component movement | BP | 450 | 33 | 38 | 2.16E-03 |
| GO:0032651 | regulation of interleukin-1 beta production | BP | 56 | 0 | 9 | 2.21E-03 |
| GO:0006284 | base-excision repair | BP | 36 | 0 | 7 | 2.21E-03 |
| GO:0002269 | leukocyte activation involved in inflammatory response | BP | 36 | 1 | 7 | 2.21E-03 |
| GO:0001774 | microglial cell activation | BP | 36 | 1 | 7 | 2.21E-03 |
| GO:0002548 | monocyte chemotaxis | BP | 36 | 3 | 7 | 2.21E-03 |
| GO:0050856 | regulation of T cell receptor signaling pathway | BP | 36 | 1 | 7 | 2.21E-03 |
| GO:0007265 | Ras protein signal transduction | BP | 406 | 29 | 35 | 2.27E-03 |
| GO:0032611 | interleukin-1 beta production | BP | 67 | 1 | 10 | 2.27E-03 |
| GO:0002690 | positive regulation of leukocyte chemotaxis | BP | 67 | 3 | 10 | 2.27E-03 |

| | | | | | | |
|---|---|---|---|---|---|---|
| GO:0051223 | regulation of protein transport | BP | 559 | 45 | 45 | 2.28E-03 |
| GO:0002637 | regulation of immunoglobulin production | BP | 46 | 1 | 8 | 2.30E-03 |
| GO:0002698 | negative regulation of immune effector process | BP | 90 | 0 | 12 | 2.33E-03 |
| GO:0030414 | peptidase inhibitor activity | MF | 90 | 10 | 12 | 2.33E-03 |
| GO:0031579 | membrane raft organization | BP | 19 | 0 | 5 | 2.36E-03 |
| GO:0050857 | positive regulation of antigen receptor-mediated signaling pathway | BP | 19 | 0 | 5 | 2.36E-03 |
| GO:0071677 | positive regulation of mononuclear cell migration | BP | 19 | 2 | 5 | 2.36E-03 |
| GO:0090025 | regulation of monocyte chemotaxis | BP | 19 | 2 | 5 | 2.36E-03 |
| GO:0034138 | toll-like receptor 3 signaling pathway | BP | 19 | 0 | 5 | 2.36E-03 |
| GO:0006816 | calcium ion transport | BP | 333 | 44 | 30 | 2.37E-03 |
| GO:0051302 | regulation of cell division | BP | 127 | 7 | 15 | 2.42E-03 |
| GO:0002793 | positive regulation of peptide secretion | BP | 206 | 12 | 21 | 2.43E-03 |
| GO:0019725 | cellular homeostasis | BP | 751 | 71 | 57 | 2.44E-03 |
| GO:2000147 | positive regulation of cell motility | BP | 438 | 31 | 37 | 2.44E-03 |
| GO:0048871 | multicellular organismal homeostasis | BP | 378 | 40 | 33 | 2.45E-03 |
| GO:0006287 | base-excision repair, gap-filling | BP | 6 | 0 | 3 | 2.48E-03 |
| GO:0035701 | hematopoietic stem cell migration | BP | 6 | 1 | 3 | 2.48E-03 |
| GO:0042824 | MHC class I peptide loading complex | CC | 6 | 0 | 3 | 2.48E-03 |
| GO:0043320 | natural killer cell degranulation | BP | 6 | 0 | 3 | 2.48E-03 |
| GO:0034128 | negative regulation of MyD88-independent toll-like receptor signal | BP | 6 | 0 | 3 | 2.48E-03 |
| GO:0033634 | positive regulation of cell-cell adhesion mediated by integrin | BP | 6 | 0 | 3 | 2.48E-03 |
| GO:1904783 | positive regulation of NMDA glutamate receptor activity | BP | 6 | 0 | 3 | 2.48E-03 |
| GO:0042268 | regulation of cytolysis | BP | 6 | 0 | 3 | 2.48E-03 |
| GO:1903236 | regulation of leukocyte tethering or rolling | BP | 6 | 0 | 3 | 2.48E-03 |
| GO:0033018 | sarcoplasmic reticulum lumen | CC | 6 | 1 | 3 | 2.48E-03 |
| GO:0035744 | T-helper 1 cell cytokine production | BP | 6 | 0 | 3 | 2.48E-03 |
| GO:0002790 | peptide secretion | BP | 469 | 47 | 39 | 2.50E-03 |
| GO:0071674 | mononuclear cell migration | BP | 57 | 5 | 9 | 2.51E-03 |
| GO:0034121 | regulation of toll-like receptor signaling pathway | BP | 57 | 3 | 9 | 2.51E-03 |
| GO:0002200 | somatic diversification of immune receptors | BP | 57 | 0 | 9 | 2.51E-03 |
| GO:0044454 | nuclear chromosome part | CC | 439 | 11 | 37 | 2.54E-03 |
| GO:0002371 | dendritic cell cytokine production | BP | 12 | 1 | 4 | 2.56E-03 |
| GO:0042555 | MCM complex | CC | 12 | 0 | 4 | 2.56E-03 |
| GO:0051709 | regulation of killing of cells of other organism | BP | 12 | 0 | 4 | 2.56E-03 |
| GO:0042403 | thyroid hormone metabolic process | BP | 12 | 3 | 4 | 2.56E-03 |
| GO:0002712 | regulation of B cell mediated immunity | BP | 37 | 0 | 7 | 2.60E-03 |
| GO:0002889 | regulation of immunoglobulin mediated immune response | BP | 37 | 0 | 7 | 2.60E-03 |
| GO:0008360 | regulation of cell shape | BP | 128 | 14 | 15 | 2.62E-03 |
| GO:0035700 | astrocyte chemotaxis | BP | 2 | 0 | 2 | 2.70E-03 |
| GO:0002368 | B cell cytokine production | BP | 2 | 0 | 2 | 2.70E-03 |
| GO:0030156 | benzodiazepine receptor binding | MF | 2 | 0 | 2 | 2.70E-03 |
| GO:0002305 | CD8-positive, gamma-delta intraepithelial T cell differentiation | BP | 2 | 0 | 2 | 2.70E-03 |
| GO:0042610 | CD8 receptor binding | MF | 2 | 0 | 2 | 2.70E-03 |
| GO:0003366 | cell-matrix adhesion involved in ameboidal cell migration | BP | 2 | 0 | 2 | 2.70E-03 |
| GO:0071283 | cellular response to iron(III) ion | BP | 2 | 0 | 2 | 2.70E-03 |
| GO:0045004 | DNA replication proofreading | BP | 2 | 0 | 2 | 2.70E-03 |
| GO:0002304 | gamma-delta intraepithelial T cell differentiation | BP | 2 | 0 | 2 | 2.70E-03 |
| GO:0071613 | granzyme B production | BP | 2 | 0 | 2 | 2.70E-03 |
| GO:0030369 | ICAM-3 receptor activity | MF | 2 | 0 | 2 | 2.70E-03 |
| GO:0034669 | integrin alpha4-beta7 complex | CC | 2 | 0 | 2 | 2.70E-03 |
| GO:0034687 | integrin alphaL-beta2 complex | CC | 2 | 0 | 2 | 2.70E-03 |
| GO:0034688 | integrin alphaM-beta2 complex | CC | 2 | 0 | 2 | 2.70E-03 |
| GO:0042008 | interleukin-18 receptor activity | MF | 2 | 0 | 2 | 2.70E-03 |
| GO:0042612 | MHC class I protein complex | CC | 2 | 0 | 2 | 2.70E-03 |
| GO:0002397 | MHC class I protein complex assembly | BP | 2 | 0 | 2 | 2.70E-03 |
| GO:0032394 | MHC class Ib receptor activity | MF | 2 | 0 | 2 | 2.70E-03 |
| GO:0031262 | Ndc80 complex | CC | 2 | 0 | 2 | 2.70E-03 |
| GO:2000349 | negative regulation of CD40 signaling pathway | BP | 2 | 0 | 2 | 2.70E-03 |
| GO:0002502 | peptide antigen assembly with MHC class I protein complex | BP | 2 | 0 | 2 | 2.70E-03 |
| GO:0002501 | peptide antigen assembly with MHC protein complex | BP | 2 | 0 | 2 | 2.70E-03 |
| GO:1905956 | positive regulation of endothelial tube morphogenesis | BP | 2 | 0 | 2 | 2.70E-03 |
| GO:0071663 | positive regulation of granzyme B production | BP | 2 | 0 | 2 | 2.70E-03 |
| GO:2000473 | positive regulation of hematopoietic stem cell migration | BP | 2 | 0 | 2 | 2.70E-03 |
| GO:1905704 | positive regulation of inhibitory synapse assembly | BP | 2 | 0 | 2 | 2.70E-03 |

| | | | | | | |
|---|---|---|---|---|---|---|
| GO:0045404 | positive regulation of interleukin-4 biosynthetic process | BP | 2 | 0 | 2 | 2.70E-03 |
| GO:0002572 | pro-T cell differentiation | BP | 2 | 0 | 2 | 2.70E-03 |
| GO:2000458 | regulation of astrocyte chemotaxis | BP | 2 | 0 | 2 | 2.70E-03 |
| GO:0071661 | regulation of granzyme B production | BP | 2 | 0 | 2 | 2.70E-03 |
| GO:2000471 | regulation of hematopoietic stem cell migration | BP | 2 | 0 | 2 | 2.70E-03 |
| GO:0070428 | regulation of nucleotide-binding oligomerization domain containing | BP | 2 | 0 | 2 | 2.70E-03 |
| GO:2000609 | regulation of thyroid hormone generation | BP | 2 | 0 | 2 | 2.70E-03 |
| GO:0035985 | senescence-associated heterochromatin focus | CC | 2 | 0 | 2 | 2.70E-03 |
| GO:0035986 | senescence-associated heterochromatin focus assembly | BP | 2 | 0 | 2 | 2.70E-03 |
| GO:0035711 | T-helper 1 cell activation | BP | 2 | 0 | 2 | 2.70E-03 |
| GO:0043408 | regulation of MAPK cascade | BP | 564 | 44 | 45 | 2.71E-03 |
| GO:0002823 | negative regulation of adaptive immune response based on somatic | BP | 28 | 0 | 6 | 2.71E-03 |
| GO:0043548 | phosphatidylinositol 3-kinase binding | MF | 28 | 4 | 6 | 2.71E-03 |
| GO:0002639 | positive regulation of immunoglobulin production | BP | 28 | 1 | 6 | 2.71E-03 |
| GO:0070372 | regulation of ERK1 and ERK2 cascade | BP | 208 | 19 | 21 | 2.74E-03 |
| GO:0033993 | response to lipid | BP | 723 | 64 | 55 | 2.76E-03 |
| GO:0048018 | receptor ligand activity | MF | 236 | 26 | 23 | 2.78E-03 |
| GO:0033674 | positive regulation of kinase activity | BP | 472 | 34 | 39 | 2.79E-03 |
| GO:0050921 | positive regulation of chemotaxis | BP | 104 | 5 | 13 | 2.80E-03 |
| GO:0019207 | kinase regulator activity | MF | 168 | 5 | 18 | 2.82E-03 |
| GO:0005623 | cell | CC | 11901 | 796 | 639 | 2.83E-03 |
| GO:0070227 | lymphocyte apoptotic process | BP | 58 | 3 | 9 | 2.83E-03 |
| GO:0010951 | negative regulation of endopeptidase activity | BP | 142 | 13 | 16 | 2.86E-03 |
| GO:1903530 | regulation of secretion by cell | BP | 567 | 70 | 45 | 2.99E-03 |
| GO:1904951 | positive regulation of establishment of protein localization | BP | 368 | 23 | 32 | 3.00E-03 |
| GO:0009636 | response to toxic substance | BP | 398 | 42 | 34 | 3.01E-03 |
| GO:0000729 | DNA double-strand break processing | BP | 20 | 0 | 5 | 3.01E-03 |
| GO:0044815 | DNA packaging complex | CC | 20 | 0 | 5 | 3.01E-03 |
| GO:1901623 | regulation of lymphocyte chemotaxis | BP | 20 | 0 | 5 | 3.01E-03 |
| GO:0034391 | regulation of smooth muscle cell apoptotic process | BP | 20 | 1 | 5 | 3.01E-03 |
| GO:0034390 | smooth muscle cell apoptotic process | BP | 20 | 1 | 5 | 3.01E-03 |
| GO:2000379 | positive regulation of reactive oxygen species metabolic process | BP | 81 | 8 | 11 | 3.01E-03 |
| GO:0045190 | isotype switching | BP | 38 | 0 | 7 | 3.05E-03 |
| GO:0002208 | somatic diversification of immunoglobulins involved in immune res | BP | 38 | 0 | 7 | 3.05E-03 |
| GO:0002204 | somatic recombination of immunoglobulin genes involved in immu | BP | 38 | 0 | 7 | 3.05E-03 |
| GO:0009968 | negative regulation of signal transduction | BP | 1038 | 73 | 74 | 3.08E-03 |
| GO:0044464 | cell part | CC | 11861 | 795 | 637 | 3.10E-03 |
| GO:0040012 | regulation of locomotion | BP | 808 | 66 | 60 | 3.11E-03 |
| GO:0051049 | regulation of transport | BP | 1444 | 180 | 98 | 3.12E-03 |
| GO:0031400 | negative regulation of protein modification process | BP | 506 | 28 | 41 | 3.13E-03 |
| GO:0009987 | cellular process | BP | 11272 | 768 | 610 | 3.18E-03 |
| GO:1903532 | positive regulation of secretion by cell | BP | 297 | 31 | 27 | 3.37E-03 |
| GO:0000776 | kinetochore | CC | 119 | 0 | 14 | 3.45E-03 |
| GO:0032481 | positive regulation of type I interferon production | BP | 71 | 0 | 10 | 3.51E-03 |
| GO:0040017 | positive regulation of locomotion | BP | 463 | 33 | 38 | 3.52E-03 |
| GO:0019731 | antibacterial humoral response | BP | 13 | 0 | 4 | 3.54E-03 |
| GO:0034185 | apolipoprotein binding | MF | 13 | 0 | 4 | 3.54E-03 |
| GO:0031254 | cell trailing edge | CC | 13 | 0 | 4 | 3.54E-03 |
| GO:0032695 | negative regulation of interleukin-12 production | BP | 13 | 0 | 4 | 3.54E-03 |
| GO:0090026 | positive regulation of monocyte chemotaxis | BP | 13 | 1 | 4 | 3.54E-03 |
| GO:0010820 | positive regulation of T cell chemotaxis | BP | 13 | 0 | 4 | 3.54E-03 |
| GO:0001931 | uropod | CC | 13 | 0 | 4 | 3.54E-03 |
| GO:0043596 | nuclear replication fork | CC | 39 | 0 | 7 | 3.56E-03 |
| GO:0032268 | regulation of cellular protein metabolic process | BP | 2042 | 116 | 132 | 3.58E-03 |
| GO:0071103 | DNA conformation change | BP | 213 | 4 | 21 | 3.63E-03 |
| GO:0046983 | protein dimerization activity | MF | 979 | 80 | 70 | 3.70E-03 |
| GO:0043123 | positive regulation of I-kappaB kinase/NF-kappaB signaling | BP | 159 | 7 | 17 | 3.73E-03 |
| GO:0019887 | protein kinase regulator activity | MF | 146 | 5 | 16 | 3.78E-03 |
| GO:2000482 | regulation of interleukin-8 secretion | BP | 21 | 1 | 5 | 3.79E-03 |
| GO:0004857 | enzyme inhibitor activity | MF | 242 | 18 | 23 | 3.80E-03 |
| GO:2000117 | negative regulation of cysteine-type endopeptidase activity | BP | 72 | 5 | 10 | 3.89E-03 |
| GO:1903426 | regulation of reactive oxygen species biosynthetic process | BP | 72 | 5 | 10 | 3.89E-03 |
| GO:0006323 | DNA packaging | BP | 108 | 4 | 13 | 3.90E-03 |
| GO:0035821 | modification of morphology or physiology of other organism | BP | 108 | 3 | 13 | 3.90E-03 |

| | | | | | | |
|---|---|---|---|---|---|---|
| GO:0050706 | regulation of interleukin-1 beta secretion | BP | 30 | 0 | 6 | 3.92E-03 |
| GO:0044772 | mitotic cell cycle phase transition | BP | 497 | 10 | 40 | 3.93E-03 |
| GO:0070838 | divalent metal ion transport | BP | 375 | 45 | 32 | 4.00E-03 |
| GO:0006898 | receptor-mediated endocytosis | BP | 201 | 14 | 20 | 4.03E-03 |
| GO:0045860 | positive regulation of protein kinase activity | BP | 436 | 32 | 36 | 4.03E-03 |
| GO:0035722 | interleukin-12-mediated signaling pathway | BP | 40 | 1 | 7 | 4.12E-03 |
| GO:0045620 | negative regulation of lymphocyte differentiation | BP | 40 | 3 | 7 | 4.12E-03 |
| GO:0090087 | regulation of peptide transport | BP | 577 | 47 | 45 | 4.14E-03 |
| GO:0023035 | CD40 signaling pathway | BP | 7 | 0 | 3 | 4.17E-03 |
| GO:0098757 | cellular response to interleukin-21 | BP | 7 | 0 | 3 | 4.17E-03 |
| GO:0002524 | hypersensitivity | BP | 7 | 0 | 3 | 4.17E-03 |
| GO:0038114 | interleukin-21-mediated signaling pathway | BP | 7 | 0 | 3 | 4.17E-03 |
| GO:0032627 | interleukin-23 production | BP | 7 | 0 | 3 | 4.17E-03 |
| GO:0030854 | positive regulation of granulocyte differentiation | BP | 7 | 0 | 3 | 4.17E-03 |
| GO:0032825 | positive regulation of natural killer cell differentiation | BP | 7 | 0 | 3 | 4.17E-03 |
| GO:0036388 | pre-replicative complex assembly | BP | 7 | 0 | 3 | 4.17E-03 |
| GO:1902299 | pre-replicative complex assembly involved in cell cycle DNA repli | BP | 7 | 0 | 3 | 4.17E-03 |
| GO:0006267 | pre-replicative complex assembly involved in nuclear cell cycle DN | BP | 7 | 0 | 3 | 4.17E-03 |
| GO:0032667 | regulation of interleukin-23 production | BP | 7 | 0 | 3 | 4.17E-03 |
| GO:0034127 | regulation of MyD88-independent toll-like receptor signaling pathv | BP | 7 | 0 | 3 | 4.17E-03 |
| GO:0043313 | regulation of neutrophil degranulation | BP | 7 | 0 | 3 | 4.17E-03 |
| GO:0070424 | regulation of nucleotide-binding oligomerization domain containing | BP | 7 | 0 | 3 | 4.17E-03 |
| GO:2000551 | regulation of T-helper 2 cell cytokine production | BP | 7 | 0 | 3 | 4.17E-03 |
| GO:0098756 | response to interleukin-21 | BP | 7 | 0 | 3 | 4.17E-03 |
| GO:0016175 | superoxide-generating NADPH oxidase activity | MF | 7 | 1 | 3 | 4.17E-03 |
| GO:0071900 | regulation of protein serine/threonine kinase activity | BP | 422 | 24 | 35 | 4.24E-03 |
| GO:0051046 | regulation of secretion | BP | 610 | 73 | 47 | 4.31E-03 |
| GO:0051222 | positive regulation of protein transport | BP | 332 | 23 | 29 | 4.32E-03 |
| GO:0051130 | positive regulation of cellular component organization | BP | 1069 | 95 | 75 | 4.32E-03 |
| GO:0010466 | negative regulation of peptidase activity | BP | 148 | 14 | 16 | 4.32E-03 |
| GO:0061134 | peptidase regulator activity | MF | 122 | 10 | 14 | 4.33E-03 |
| GO:0002474 | antigen processing and presentation of peptide antigen via MHC cla | BP | 85 | 1 | 11 | 4.40E-03 |
| GO:1901216 | positive regulation of neuron death | BP | 85 | 5 | 11 | 4.40E-03 |
| GO:0050710 | negative regulation of cytokine secretion | BP | 51 | 2 | 8 | 4.47E-03 |
| GO:0099024 | plasma membrane invagination | BP | 51 | 4 | 8 | 4.47E-03 |
| GO:0046209 | nitric oxide metabolic process | BP | 62 | 7 | 9 | 4.49E-03 |
| GO:0001894 | tissue homeostasis | BP | 162 | 7 | 17 | 4.51E-03 |
| GO:0060759 | regulation of response to cytokine stimulus | BP | 149 | 5 | 16 | 4.62E-03 |
| GO:0097242 | amyloid-beta clearance | BP | 31 | 3 | 6 | 4.65E-03 |
| GO:0070266 | necroptotic process | BP | 31 | 1 | 6 | 4.65E-03 |
| GO:0005737 | cytoplasm | CC | 8907 | 540 | 494 | 4.66E-03 |
| GO:0072511 | divalent inorganic cation transport | BP | 379 | 46 | 32 | 4.68E-03 |
| GO:0006302 | double-strand break repair | BP | 190 | 2 | 19 | 4.72E-03 |
| GO:0007259 | JAK-STAT cascade | BP | 98 | 5 | 12 | 4.74E-03 |
| GO:0043124 | negative regulation of I-kappaB kinase/NF-kappaB signaling | BP | 41 | 2 | 7 | 4.75E-03 |
| GO:0045670 | regulation of osteoclast differentiation | BP | 41 | 0 | 7 | 4.75E-03 |
| GO:0042116 | macrophage activation | BP | 74 | 2 | 10 | 4.75E-03 |
| GO:0002467 | germinal center formation | BP | 14 | 0 | 4 | 4.76E-03 |
| GO:0090280 | positive regulation of calcium ion import | BP | 14 | 0 | 4 | 4.76E-03 |
| GO:0010744 | positive regulation of macrophage derived foam cell differentiation | BP | 14 | 3 | 4 | 4.76E-03 |
| GO:0045651 | positive regulation of macrophage differentiation | BP | 14 | 0 | 4 | 4.76E-03 |
| GO:0010819 | regulation of T cell chemotaxis | BP | 14 | 0 | 4 | 4.76E-03 |
| GO:0022402 | cell cycle process | BP | 1125 | 39 | 78 | 4.90E-03 |
| GO:0032479 | regulation of type I interferon production | BP | 111 | 1 | 13 | 4.93E-03 |
| GO:2001057 | reactive nitrogen species metabolic process | BP | 63 | 7 | 9 | 5.00E-03 |
| GO:0150076 | neuroinflammatory response | BP | 52 | 2 | 8 | 5.05E-03 |
| GO:0032652 | regulation of interleukin-1 production | BP | 64 | 0 | 9 | 5.56E-03 |
| GO:0032880 | regulation of protein localization | BP | 830 | 65 | 60 | 5.57E-03 |
| GO:0050729 | positive regulation of inflammatory response | BP | 100 | 7 | 12 | 5.58E-03 |
| GO:0007260 | tyrosine phosphorylation of STAT protein | BP | 53 | 3 | 8 | 5.68E-03 |
| GO:0001959 | regulation of cytokine-mediated signaling pathway | BP | 139 | 5 | 15 | 5.72E-03 |
| GO:0032606 | type I interferon production | BP | 113 | 1 | 13 | 5.74E-03 |
| GO:0032411 | positive regulation of transporter activity | BP | 88 | 15 | 11 | 5.74E-03 |
| GO:0001228 | DNA-binding transcription activator activity, RNA polymerase II-s | MF | 309 | 23 | 27 | 5.74E-03 |

| | | | | | | |
|---|---|---|---|---|---|---|
| GO:0061844 | antimicrobial humoral immune response mediated by antimicrobial | BP | 23 | 0 | 5 | 5.75E-03 |
| GO:0033032 | regulation of myeloid cell apoptotic process | BP | 23 | 0 | 5 | 5.75E-03 |
| GO:0032612 | interleukin-1 production | BP | 76 | 1 | 10 | 5.76E-03 |
| GO:0031334 | positive regulation of protein complex assembly | BP | 236 | 11 | 22 | 5.76E-03 |
| GO:2000145 | regulation of cell motility | BP | 750 | 64 | 55 | 5.88E-03 |
| GO:0051347 | positive regulation of transferase activity | BP | 541 | 36 | 42 | 5.95E-03 |
| GO:0097696 | STAT cascade | BP | 101 | 6 | 12 | 6.04E-03 |
| GO:0004930 | G protein-coupled receptor activity | MF | 237 | 29 | 22 | 6.05E-03 |
| GO:0002262 | myeloid cell homeostasis | BP | 127 | 5 | 14 | 6.19E-03 |
| GO:0030479 | actin cortical patch | CC | 15 | 0 | 4 | 6.23E-03 |
| GO:0050832 | defense response to fungus | BP | 15 | 0 | 4 | 6.23E-03 |
| GO:0061645 | endocytic patch | CC | 15 | 0 | 4 | 6.23E-03 |
| GO:0051782 | negative regulation of cell division | BP | 15 | 0 | 4 | 6.23E-03 |
| GO:0140131 | positive regulation of lymphocyte chemotaxis | BP | 15 | 0 | 4 | 6.23E-03 |
| GO:0016447 | somatic recombination of immunoglobulin gene segments | BP | 43 | 0 | 7 | 6.23E-03 |
| GO:0044770 | cell cycle phase transition | BP | 527 | 11 | 41 | 6.31E-03 |
| GO:0033138 | positive regulation of peptidyl-serine phosphorylation | BP | 77 | 5 | 10 | 6.32E-03 |
| GO:1990782 | protein tyrosine kinase binding | MF | 77 | 5 | 10 | 6.32E-03 |
| GO:0046427 | positive regulation of JAK-STAT cascade | BP | 54 | 4 | 8 | 6.37E-03 |
| GO:0002483 | antigen processing and presentation of endogenous peptide antigen | BP | 8 | 0 | 3 | 6.42E-03 |
| GO:0019885 | antigen processing and presentation of endogenous peptide antigen | BP | 8 | 0 | 3 | 6.42E-03 |
| GO:0042609 | CD4 receptor binding | MF | 8 | 0 | 3 | 6.42E-03 |
| GO:0000796 | condensin complex | CC | 8 | 0 | 3 | 6.42E-03 |
| GO:0045060 | negative thymic T cell selection | BP | 8 | 1 | 3 | 6.42E-03 |
| GO:2000774 | positive regulation of cellular senescence | BP | 8 | 0 | 3 | 6.42E-03 |
| GO:0045591 | positive regulation of regulatory T cell differentiation | BP | 8 | 0 | 3 | 6.42E-03 |
| GO:2000109 | regulation of macrophage apoptotic process | BP | 8 | 0 | 3 | 6.42E-03 |
| GO:0032823 | regulation of natural killer cell differentiation | BP | 8 | 0 | 3 | 6.42E-03 |
| GO:0035745 | T-helper 2 cell cytokine production | BP | 8 | 0 | 3 | 6.42E-03 |
| GO:0000775 | chromosome, centromeric region | CC | 168 | 0 | 17 | 6.49E-03 |
| GO:0097553 | calcium ion transmembrane import into cytosol | BP | 115 | 16 | 13 | 6.64E-03 |
| GO:0000188 | inactivation of MAPK activity | BP | 24 | 0 | 5 | 6.95E-03 |
| GO:0072606 | interleukin-8 secretion | BP | 24 | 2 | 5 | 6.95E-03 |
| GO:0002335 | mature B cell differentiation | BP | 24 | 1 | 5 | 6.95E-03 |
| GO:0032647 | regulation of interferon-alpha production | BP | 24 | 1 | 5 | 6.95E-03 |
| GO:0043304 | regulation of mast cell degranulation | BP | 24 | 1 | 5 | 6.95E-03 |
| GO:0000083 | regulation of transcription involved in G1/S transition of mitotic ce | BP | 24 | 0 | 5 | 6.95E-03 |
| GO:0051209 | release of sequestered calcium ion into cytosol | BP | 103 | 13 | 12 | 7.05E-03 |
| GO:0000731 | DNA synthesis involved in DNA repair | BP | 44 | 0 | 7 | 7.08E-03 |
| GO:0035580 | specific granule lumen | CC | 44 | 0 | 7 | 7.08E-03 |
| GO:0060402 | calcium ion transport into cytosol | BP | 129 | 18 | 14 | 7.09E-03 |
| GO:0032677 | regulation of interleukin-8 production | BP | 55 | 1 | 8 | 7.13E-03 |
| GO:0071901 | negative regulation of protein serine/threonine kinase activity | BP | 116 | 6 | 13 | 7.14E-03 |
| GO:0044444 | cytoplasmic part | CC | 7640 | 438 | 428 | 7.38E-03 |
| GO:0038061 | NIK/NF-kappaB signaling | BP | 143 | 3 | 15 | 7.41E-03 |
| GO:0045581 | negative regulation of T cell differentiation | BP | 34 | 2 | 6 | 7.44E-03 |
| GO:0045429 | positive regulation of nitric oxide biosynthetic process | BP | 34 | 5 | 6 | 7.44E-03 |
| GO:0050704 | regulation of interleukin-1 secretion | BP | 34 | 0 | 6 | 7.44E-03 |
| GO:0043154 | negative regulation of cysteine-type endopeptidase activity involvec | BP | 67 | 5 | 9 | 7.52E-03 |
| GO:0030674 | protein binding, bridging | MF | 157 | 11 | 16 | 7.61E-03 |
| GO:0035690 | cellular response to drug | BP | 301 | 33 | 26 | 7.68E-03 |
| GO:0051047 | positive regulation of secretion | BP | 316 | 33 | 27 | 7.68E-03 |
| GO:0097237 | cellular response to toxic substance | BP | 185 | 15 | 18 | 7.74E-03 |
| GO:0035697 | CD8-positive, alpha-beta T cell extravasation | BP | 3 | 0 | 2 | 7.81E-03 |
| GO:0150062 | complement-mediated synapse pruning | BP | 3 | 0 | 2 | 7.81E-03 |
| GO:0042335 | cuticle development | BP | 3 | 0 | 2 | 7.81E-03 |
| GO:0042010 | interleukin-15 receptor activity | MF | 3 | 0 | 2 | 7.81E-03 |
| GO:0045092 | interleukin-18 receptor complex | CC | 3 | 0 | 2 | 7.81E-03 |
| GO:0019976 | interleukin-2 binding | MF | 3 | 0 | 2 | 7.81E-03 |
| GO:0004911 | interleukin-2 receptor activity | MF | 3 | 0 | 2 | 7.81E-03 |
| GO:0042097 | interleukin-4 biosynthetic process | BP | 3 | 0 | 2 | 7.81E-03 |
| GO:0006272 | leading strand elongation | BP | 3 | 0 | 2 | 7.81E-03 |
| GO:0071593 | lymphocyte aggregation | BP | 3 | 0 | 2 | 7.81E-03 |
| GO:0001766 | membrane raft polarization | BP | 3 | 0 | 2 | 7.81E-03 |

| | | | | | | |
|---|---|---|---|---|---|---|
| GO:0042611 | MHC protein complex | CC | 3 | 0 | 2 | 7.81E-03 |
| GO:0002396 | MHC protein complex assembly | BP | 3 | 0 | 2 | 7.81E-03 |
| GO:0002277 | myeloid dendritic cell activation involved in immune response | BP | 3 | 0 | 2 | 7.81E-03 |
| GO:0045918 | negative regulation of cytolysis | BP | 3 | 0 | 2 | 7.81E-03 |
| GO:1905463 | negative regulation of DNA duplex unwinding | BP | 3 | 0 | 2 | 7.81E-03 |
| GO:0070433 | negative regulation of nucleotide-binding oligomerization domain c | BP | 3 | 0 | 2 | 7.81E-03 |
| GO:0070425 | negative regulation of nucleotide-binding oligomerization domain c | BP | 3 | 0 | 2 | 7.81E-03 |
| GO:0045588 | positive regulation of gamma-delta T cell differentiation | BP | 3 | 0 | 2 | 7.81E-03 |
| GO:1901666 | positive regulation of NAD+ ADP-ribosyltransferase activity | BP | 3 | 0 | 2 | 7.81E-03 |
| GO:2000503 | positive regulation of natural killer cell chemotaxis | BP | 3 | 0 | 2 | 7.81E-03 |
| GO:2000363 | positive regulation of prostaglandin-E synthase activity | BP | 3 | 0 | 2 | 7.81E-03 |
| GO:2000412 | positive regulation of thymocyte migration | BP | 3 | 0 | 2 | 7.81E-03 |
| GO:2000449 | regulation of CD8-positive, alpha-beta T cell extravasation | BP | 3 | 0 | 2 | 7.81E-03 |
| GO:1905702 | regulation of inhibitory synapse assembly | BP | 3 | 0 | 2 | 7.81E-03 |
| GO:0045402 | regulation of interleukin-4 biosynthetic process | BP | 3 | 0 | 2 | 7.81E-03 |
| GO:1901664 | regulation of NAD+ ADP-ribosyltransferase activity | BP | 3 | 0 | 2 | 7.81E-03 |
| GO:2000361 | regulation of prostaglandin-E synthase activity | BP | 3 | 0 | 2 | 7.81E-03 |
| GO:2000328 | regulation of T-helper 17 cell lineage commitment | BP | 3 | 0 | 2 | 7.81E-03 |
| GO:2000410 | regulation of thymocyte migration | BP | 3 | 0 | 2 | 7.81E-03 |
| GO:0150064 | vertebrate eye-specific patterning | BP | 3 | 0 | 2 | 7.81E-03 |
| GO:0051128 | regulation of cellular component organization | BP | 2090 | 174 | 132 | 7.94E-03 |
| GO:1904894 | positive regulation of STAT cascade | BP | 56 | 4 | 8 | 7.95E-03 |
| GO:0008092 | cytoskeletal protein binding | MF | 828 | 101 | 59 | 8.01E-03 |
| GO:0098805 | whole membrane | CC | 1371 | 85 | 91 | 8.02E-03 |
| GO:0030335 | positive regulation of cell migration | BP | 425 | 30 | 34 | 8.19E-03 |
| GO:0051283 | negative regulation of sequestering of calcium ion | BP | 105 | 13 | 12 | 8.20E-03 |
| GO:0030856 | regulation of epithelial cell differentiation | BP | 105 | 11 | 12 | 8.20E-03 |
| GO:0007049 | cell cycle | BP | 1511 | 55 | 99 | 8.22E-03 |
| GO:0032414 | positive regulation of ion transmembrane transporter activity | BP | 80 | 15 | 10 | 8.26E-03 |
| GO:0018958 | phenol-containing compound metabolic process | BP | 68 | 14 | 9 | 8.28E-03 |
| GO:0045577 | regulation of B cell differentiation | BP | 25 | 1 | 5 | 8.33E-03 |
| GO:0002691 | regulation of cellular extravasation | BP | 25 | 2 | 5 | 8.33E-03 |
| GO:0033006 | regulation of mast cell activation involved in immune response | BP | 25 | 1 | 5 | 8.33E-03 |
| GO:1904407 | positive regulation of nitric oxide metabolic process | BP | 35 | 5 | 6 | 8.59E-03 |
| GO:2000772 | regulation of cellular senescence | BP | 35 | 0 | 6 | 8.59E-03 |
| GO:0019882 | antigen processing and presentation | BP | 187 | 7 | 18 | 8.62E-03 |
| GO:0042493 | response to drug | BP | 782 | 92 | 56 | 8.77E-03 |
| GO:0099512 | supramolecular fiber | CC | 667 | 85 | 49 | 8.78E-03 |
| GO:2001251 | negative regulation of chromosome organization | BP | 119 | 0 | 13 | 8.80E-03 |
| GO:0051282 | regulation of sequestering of calcium ion | BP | 106 | 13 | 12 | 8.82E-03 |
| GO:0019210 | kinase inhibitor activity | MF | 57 | 2 | 8 | 8.84E-03 |
| GO:0045132 | meiotic chromosome segregation | BP | 57 | 0 | 8 | 8.84E-03 |
| GO:1904062 | regulation of cation transmembrane transport | BP | 260 | 42 | 23 | 8.94E-03 |
| GO:0048010 | vascular endothelial growth factor receptor signaling pathway | BP | 81 | 3 | 10 | 8.99E-03 |
| GO:0098797 | plasma membrane protein complex | CC | 413 | 74 | 33 | 9.26E-03 |
| GO:0032059 | bleb | CC | 9 | 1 | 3 | 9.27E-03 |
| GO:0071281 | cellular response to iron ion | BP | 9 | 0 | 3 | 9.27E-03 |
| GO:0006957 | complement activation, alternative pathway | BP | 9 | 0 | 3 | 9.27E-03 |
| GO:0000940 | condensed chromosome outer kinetochore | CC | 9 | 0 | 3 | 9.27E-03 |
| GO:0034663 | endoplasmic reticulum chaperone complex | CC | 9 | 0 | 3 | 9.27E-03 |
| GO:0070757 | interleukin-35-mediated signaling pathway | BP | 9 | 0 | 3 | 9.27E-03 |
| GO:0050713 | negative regulation of interleukin-1 beta secretion | BP | 9 | 0 | 3 | 9.27E-03 |
| GO:0001911 | negative regulation of leukocyte mediated cytotoxicity | BP | 9 | 0 | 3 | 9.27E-03 |
| GO:0002829 | negative regulation of type 2 immune response | BP | 9 | 0 | 3 | 9.27E-03 |
| GO:0014049 | positive regulation of glutamate secretion | BP | 9 | 1 | 3 | 9.27E-03 |
| GO:0032740 | positive regulation of interleukin-17 production | BP | 9 | 0 | 3 | 9.27E-03 |
| GO:0051901 | positive regulation of mitochondrial depolarization | BP | 9 | 0 | 3 | 9.27E-03 |
| GO:0071417 | cellular response to organonitrogen compound | BP | 476 | 45 | 37 | 9.31E-03 |
| GO:0051983 | regulation of chromosome segregation | BP | 94 | 0 | 11 | 9.37E-03 |
| GO:0032269 | negative regulation of cellular protein metabolic process | BP | 818 | 47 | 58 | 9.40E-03 |
| GO:0034767 | positive regulation of ion transmembrane transport | BP | 120 | 20 | 13 | 9.41E-03 |
| GO:0033135 | regulation of peptidyl-serine phosphorylation | BP | 107 | 8 | 12 | 9.48E-03 |
| GO:0006270 | DNA replication initiation | BP | 36 | 0 | 6 | 9.87E-03 |
| GO:0097300 | programmed necrotic cell death | BP | 36 | 1 | 6 | 9.87E-03 |

| | | | | | | |
|---|---|---|---|---|---|---|
| GO:0032607 | interferon-alpha production | BP | 26 | 1 | 5 | 9.88E-03 |
| GO:0051019 | mitogen-activated protein kinase binding | MF | 26 | 0 | 5 | 9.88E-03 |
| GO:0033028 | myeloid cell apoptotic process | BP | 26 | 0 | 5 | 9.88E-03 |
| GO:0050869 | negative regulation of B cell activation | BP | 26 | 1 | 5 | 9.88E-03 |
| GO:0004629 | phospholipase C activity | MF | 26 | 2 | 5 | 9.88E-03 |
| GO:0034765 | regulation of ion transmembrane transport | BP | 353 | 64 | 29 | 9.91E-03 |
| GO:0048878 | chemical homeostasis | BP | 904 | 98 | 63 | 9.95E-03 |
| GO:0006925 | inflammatory cell apoptotic process | BP | 17 | 0 | 4 | 9.99E-03 |
| GO:0032691 | negative regulation of interleukin-1 beta production | BP | 17 | 0 | 4 | 9.99E-03 |
| GO:0016646 | oxidoreductase activity, acting on the CH-NH group of donors, NA | MF | 17 | 2 | 4 | 9.99E-03 |
| GO:0017116 | single-stranded DNA-dependent ATP-dependent DNA helicase act | MF | 17 | 0 | 4 | 9.99E-03 |
| GO:0043142 | single-stranded DNA-dependent ATPase activity | MF | 17 | 0 | 4 | 9.99E-03 |
| GO:0045063 | T-helper 1 cell differentiation | BP | 17 | 0 | 4 | 9.99E-03 |
| GO:0046653 | tetrahydrofolate metabolic process | BP | 17 | 1 | 4 | 9.99E-03 |
| GO:0051248 | negative regulation of protein metabolic process | BP | 871 | 51 | 61 | 1.01E-02 |
| GO:0032392 | DNA geometric change | BP | 108 | 2 | 12 | 1.02E-02 |
| GO:0140013 | meiotic nuclear division | BP | 108 | 2 | 12 | 1.02E-02 |
| GO:0099081 | supramolecular polymer | CC | 674 | 88 | 49 | 1.06E-02 |
| GO:0046425 | regulation of JAK-STAT cascade | BP | 83 | 6 | 10 | 1.06E-02 |
| GO:0051235 | maintenance of location | BP | 279 | 24 | 24 | 1.07E-02 |
| GO:0010324 | membrane invagination | BP | 59 | 4 | 8 | 1.08E-02 |
| GO:0006809 | nitric oxide biosynthetic process | BP | 59 | 7 | 8 | 1.08E-02 |
| GO:0099080 | supramolecular complex | CC | 675 | 88 | 49 | 1.08E-02 |
| GO:0030334 | regulation of cell migration | BP | 708 | 62 | 51 | 1.09E-02 |
| GO:0043200 | response to amino acid | BP | 96 | 8 | 11 | 1.09E-02 |
| GO:0003678 | DNA helicase activity | MF | 71 | 1 | 9 | 1.09E-02 |
| GO:0009898 | cytoplasmic side of plasma membrane | CC | 136 | 14 | 14 | 1.11E-02 |
| GO:2000045 | regulation of G1/S transition of mitotic cell cycle | BP | 136 | 3 | 14 | 1.11E-02 |
| GO:0043405 | regulation of MAP kinase activity | BP | 280 | 18 | 24 | 1.11E-02 |
| GO:0050702 | interleukin-1 beta secretion | BP | 37 | 1 | 6 | 1.13E-02 |
| GO:1905819 | negative regulation of chromosome separation | BP | 37 | 0 | 6 | 1.13E-02 |
| GO:2000816 | negative regulation of mitotic sister chromatid separation | BP | 37 | 0 | 6 | 1.13E-02 |
| GO:0019985 | translesion synthesis | BP | 37 | 0 | 6 | 1.13E-02 |
| GO:0007093 | mitotic cell cycle checkpoint | BP | 150 | 2 | 15 | 1.13E-02 |
| GO:0005871 | kinesin complex | CC | 48 | 3 | 7 | 1.14E-02 |
| GO:0045428 | regulation of nitric oxide biosynthetic process | BP | 48 | 5 | 7 | 1.14E-02 |
| GO:0051346 | negative regulation of hydrolase activity | BP | 311 | 27 | 26 | 1.15E-02 |
| GO:0001782 | B cell homeostasis | BP | 27 | 0 | 5 | 1.16E-02 |
| GO:0030261 | chromosome condensation | BP | 27 | 0 | 5 | 1.16E-02 |
| GO:0046875 | ephrin receptor binding | MF | 27 | 6 | 5 | 1.16E-02 |
| GO:0070232 | regulation of T cell apoptotic process | BP | 27 | 1 | 5 | 1.16E-02 |
| GO:0035666 | TRIF-dependent toll-like receptor signaling pathway | BP | 27 | 0 | 5 | 1.16E-02 |
| GO:0032637 | interleukin-8 production | BP | 60 | 2 | 8 | 1.20E-02 |
| GO:0004003 | ATP-dependent DNA helicase activity | MF | 18 | 0 | 4 | 1.23E-02 |
| GO:0008026 | ATP-dependent helicase activity | MF | 18 | 0 | 4 | 1.23E-02 |
| GO:0008574 | ATP-dependent microtubule motor activity, plus-end-directed | MF | 18 | 1 | 4 | 1.23E-02 |
| GO:0046058 | cAMP metabolic process | BP | 18 | 3 | 4 | 1.23E-02 |
| GO:0043027 | cysteine-type endopeptidase inhibitor activity involved in apoptotic | MF | 18 | 1 | 4 | 1.23E-02 |
| GO:0002719 | negative regulation of cytokine production involved in immune resp | BP | 18 | 0 | 4 | 1.23E-02 |
| GO:0043302 | positive regulation of leukocyte degranulation | BP | 18 | 1 | 4 | 1.23E-02 |
| GO:0070035 | purine NTP-dependent helicase activity | MF | 18 | 0 | 4 | 1.23E-02 |
| GO:0031664 | regulation of lipopolysaccharide-mediated signaling pathway | BP | 18 | 1 | 4 | 1.23E-02 |
| GO:0005575 | cellular_component | CC | 12612 | 866 | 667 | 1.24E-02 |
| GO:1904019 | epithelial cell apoptotic process | BP | 85 | 3 | 10 | 1.25E-02 |
| GO:1901214 | regulation of neuron death | BP | 253 | 19 | 22 | 1.25E-02 |
| GO:0002223 | stimulatory C-type lectin receptor signaling pathway | BP | 98 | 0 | 11 | 1.26E-02 |
| GO:0045165 | cell fate commitment | BP | 166 | 12 | 16 | 1.27E-02 |
| GO:0050891 | multicellular organismal water homeostasis | BP | 49 | 6 | 7 | 1.27E-02 |
| GO:2001259 | positive regulation of cation channel activity | BP | 49 | 10 | 7 | 1.27E-02 |
| GO:0016445 | somatic diversification of immunoglobulins | BP | 49 | 0 | 7 | 1.27E-02 |
| GO:0019883 | antigen processing and presentation of endogenous antigen | BP | 10 | 0 | 3 | 1.27E-02 |
| GO:0001608 | G protein-coupled nucleotide receptor activity | MF | 10 | 0 | 3 | 1.27E-02 |
| GO:0045028 | G protein-coupled purinergic nucleotide receptor activity | MF | 10 | 0 | 3 | 1.27E-02 |
| GO:0035589 | G protein-coupled purinergic nucleotide receptor signaling pathway | BP | 10 | 0 | 3 | 1.27E-02 |

| | | | | | | |
|---|---|---|---|---|---|---|
| GO:0071888 | macrophage apoptotic process | BP | 10 | 0 | 3 | 1.27E-02 |
| GO:0043020 | NADPH oxidase complex | CC | 10 | 1 | 3 | 1.27E-02 |
| GO:0031342 | negative regulation of cell killing | BP | 10 | 0 | 3 | 1.27E-02 |
| GO:0070431 | nucleotide-binding oligomerization domain containing 2 signaling p | BP | 10 | 0 | 3 | 1.27E-02 |
| GO:0090343 | positive regulation of cell aging | BP | 10 | 0 | 3 | 1.27E-02 |
| GO:0048302 | regulation of isotype switching to IgG isotypes | BP | 10 | 0 | 3 | 1.27E-02 |
| GO:1902563 | regulation of neutrophil activation | BP | 10 | 0 | 3 | 1.27E-02 |
| GO:0042743 | hydrogen peroxide metabolic process | BP | 38 | 2 | 6 | 1.28E-02 |
| GO:0071675 | regulation of mononuclear cell migration | BP | 38 | 4 | 6 | 1.28E-02 |
| GO:0051668 | localization within membrane | BP | 125 | 11 | 13 | 1.30E-02 |
| GO:1901224 | positive regulation of NIK/NF-kappaB signaling | BP | 61 | 3 | 8 | 1.31E-02 |
| GO:0048365 | Rac GTPase binding | MF | 61 | 3 | 8 | 1.31E-02 |
| GO:0001664 | G protein-coupled receptor binding | MF | 181 | 18 | 17 | 1.32E-02 |
| GO:0060090 | molecular adaptor activity | MF | 210 | 15 | 19 | 1.33E-02 |
| GO:0005764 | lysosome | CC | 552 | 24 | 41 | 1.33E-02 |
| GO:0060401 | cytosolic calcium ion transport | BP | 139 | 18 | 14 | 1.33E-02 |
| GO:0003690 | double-stranded DNA binding | MF | 733 | 45 | 52 | 1.34E-02 |
| GO:1904892 | regulation of STAT cascade | BP | 86 | 7 | 10 | 1.35E-02 |
| GO:0071402 | cellular response to lipoprotein particle stimulus | BP | 28 | 2 | 5 | 1.36E-02 |
| GO:1990752 | microtubule end | CC | 28 | 1 | 5 | 1.36E-02 |
| GO:0006259 | DNA metabolic process | BP | 817 | 14 | 57 | 1.36E-02 |
| GO:0097191 | extrinsic apoptotic signaling pathway | BP | 196 | 8 | 18 | 1.36E-02 |
| GO:0000323 | lytic vacuole | CC | 553 | 24 | 41 | 1.37E-02 |
| GO:0023057 | negative regulation of signaling | BP | 1125 | 89 | 75 | 1.42E-02 |
| GO:0007169 | transmembrane receptor protein tyrosine kinase signaling pathway | BP | 587 | 51 | 43 | 1.42E-02 |
| GO:0044877 | protein-containing complex binding | MF | 938 | 74 | 64 | 1.45E-02 |
| GO:0051270 | regulation of cellular component movement | BP | 820 | 72 | 57 | 1.45E-02 |
| GO:0032508 | DNA duplex unwinding | BP | 100 | 1 | 11 | 1.45E-02 |
| GO:0002220 | innate immune response activating cell surface receptor signaling p | BP | 100 | 0 | 11 | 1.45E-02 |
| GO:0043565 | sequence-specific DNA binding | MF | 821 | 48 | 57 | 1.49E-02 |
| GO:0009913 | epidermal cell differentiation | BP | 141 | 8 | 14 | 1.49E-02 |
| GO:0048020 | CCR chemokine receptor binding | MF | 19 | 1 | 4 | 1.50E-02 |
| GO:0035588 | G protein-coupled purinergic receptor signaling pathway | BP | 19 | 1 | 4 | 1.50E-02 |
| GO:1902993 | positive regulation of amyloid precursor protein catabolic process | BP | 19 | 3 | 4 | 1.50E-02 |
| GO:0033630 | positive regulation of cell adhesion mediated by integrin | BP | 19 | 1 | 4 | 1.50E-02 |
| GO:0032495 | response to muramyl dipeptide | BP | 19 | 0 | 4 | 1.50E-02 |
| GO:0032237 | activation of store-operated calcium channel activity | BP | 4 | 0 | 2 | 1.51E-02 |
| GO:1902292 | cell cycle DNA replication initiation | BP | 4 | 0 | 2 | 1.51E-02 |
| GO:0000942 | condensed nuclear chromosome outer kinetochore | CC | 4 | 0 | 2 | 1.51E-02 |
| GO:0031261 | DNA replication preinitiation complex | CC | 4 | 0 | 2 | 1.51E-02 |
| GO:0034987 | immunoglobulin receptor binding | MF | 4 | 1 | 2 | 1.51E-02 |
| GO:0001875 | lipopolysaccharide receptor activity | MF | 4 | 0 | 2 | 1.51E-02 |
| GO:0004488 | methylenetetrahydrofolate dehydrogenase (NADP+) activity | MF | 4 | 0 | 2 | 1.51E-02 |
| GO:1902975 | mitotic DNA replication initiation | BP | 4 | 0 | 2 | 1.51E-02 |
| GO:0052490 | negative regulation by organism of programmed cell death in other | BP | 4 | 0 | 2 | 1.51E-02 |
| GO:0033668 | negative regulation by symbiont of host apoptotic process | BP | 4 | 0 | 2 | 1.51E-02 |
| GO:0052041 | negative regulation by symbiont of host programmed cell death | BP | 4 | 0 | 2 | 1.51E-02 |
| GO:0045914 | negative regulation of catecholamine metabolic process | BP | 4 | 1 | 2 | 1.51E-02 |
| GO:2001186 | negative regulation of CD8-positive, alpha-beta T cell activation | BP | 4 | 0 | 2 | 1.51E-02 |
| GO:0045963 | negative regulation of dopamine metabolic process | BP | 4 | 1 | 2 | 1.51E-02 |
| GO:2000110 | negative regulation of macrophage apoptotic process | BP | 4 | 0 | 2 | 1.51E-02 |
| GO:0061218 | negative regulation of mesonephros development | BP | 4 | 0 | 2 | 1.51E-02 |
| GO:0034136 | negative regulation of toll-like receptor 2 signaling pathway | BP | 4 | 0 | 2 | 1.51E-02 |
| GO:1901164 | negative regulation of trophoblast cell migration | BP | 4 | 0 | 2 | 1.51E-02 |
| GO:1902315 | nuclear cell cycle DNA replication initiation | BP | 4 | 0 | 2 | 1.51E-02 |
| GO:0070427 | nucleotide-binding oligomerization domain containing 1 signaling p | BP | 4 | 0 | 2 | 1.51E-02 |
| GO:0015671 | oxygen transport | BP | 4 | 0 | 2 | 1.51E-02 |
| GO:0052812 | phosphatidylinositol-3,4-bisphosphate 5-kinase activity | MF | 4 | 0 | 2 | 1.51E-02 |
| GO:0046645 | positive regulation of gamma-delta T cell activation | BP | 4 | 0 | 2 | 1.51E-02 |
| GO:0002885 | positive regulation of hypersensitivity | BP | 4 | 0 | 2 | 1.51E-02 |
| GO:1901341 | positive regulation of store-operated calcium channel activity | BP | 4 | 0 | 2 | 1.51E-02 |
| GO:0008330 | protein tyrosine/threonine phosphatase activity | MF | 4 | 0 | 2 | 1.51E-02 |
| GO:0072679 | thymocyte migration | BP | 4 | 0 | 2 | 1.51E-02 |
| GO:0002513 | tolerance induction to self antigen | BP | 4 | 0 | 2 | 1.51E-02 |

| | | | | | | |
|---|---|---|---|---|---|---|
| GO:0030229 | very-low-density lipoprotein particle receptor activity | MF | 4 | 0 | 2 | 1.51E-02 |
| GO:0030168 | platelet activation | BP | 128 | 11 | 13 | 1.56E-02 |
| GO:0042803 | protein homodimerization activity | MF | 673 | 56 | 48 | 1.56E-02 |
| GO:0043113 | receptor clustering | BP | 51 | 9 | 7 | 1.57E-02 |
| GO:0042509 | regulation of tyrosine phosphorylation of STAT protein | BP | 51 | 3 | 7 | 1.57E-02 |
| GO:0008608 | attachment of spindle microtubules to kinetochore | BP | 29 | 0 | 5 | 1.57E-02 |
| GO:0090279 | regulation of calcium ion import | BP | 29 | 2 | 5 | 1.57E-02 |
| GO:0050690 | regulation of defense response to virus by virus | BP | 29 | 0 | 5 | 1.57E-02 |
| GO:0045622 | regulation of T-helper cell differentiation | BP | 29 | 0 | 5 | 1.57E-02 |
| GO:0030041 | actin filament polymerization | BP | 156 | 13 | 15 | 1.58E-02 |
| GO:0043409 | negative regulation of MAPK cascade | BP | 142 | 9 | 14 | 1.58E-02 |
| GO:0006974 | cellular response to DNA damage stimulus | BP | 724 | 13 | 51 | 1.61E-02 |
| GO:0033048 | negative regulation of mitotic sister chromatid segregation | BP | 40 | 0 | 6 | 1.63E-02 |
| GO:0051234 | establishment of localization | BP | 4066 | 306 | 237 | 1.66E-02 |
| GO:0046578 | regulation of Ras protein signal transduction | BP | 215 | 18 | 19 | 1.67E-02 |
| GO:0008150 | biological_process | BP | 12173 | 834 | 647 | 1.68E-02 |
| GO:0004115 | 3',5'-cyclic-AMP phosphodiesterase activity | MF | 11 | 2 | 3 | 1.68E-02 |
| GO:0051315 | attachment of mitotic spindle microtubules to kinetochore | BP | 11 | 0 | 3 | 1.68E-02 |
| GO:0038110 | interleukin-2-mediated signaling pathway | BP | 11 | 0 | 3 | 1.68E-02 |
| GO:0034362 | low-density lipoprotein particle | CC | 11 | 0 | 3 | 1.68E-02 |
| GO:0017017 | MAP kinase tyrosine/serine/threonine phosphatase activity | MF | 11 | 0 | 3 | 1.68E-02 |
| GO:0050711 | negative regulation of interleukin-1 secretion | BP | 11 | 0 | 3 | 1.68E-02 |
| GO:0045579 | positive regulation of B cell differentiation | BP | 11 | 0 | 3 | 1.68E-02 |
| GO:0034116 | positive regulation of heterotypic cell-cell adhesion | BP | 11 | 0 | 3 | 1.68E-02 |
| GO:1904181 | positive regulation of membrane depolarization | BP | 11 | 2 | 3 | 1.68E-02 |
| GO:0033632 | regulation of cell-cell adhesion mediated by integrin | BP | 11 | 0 | 3 | 1.68E-02 |
| GO:0045346 | regulation of MHC class II biosynthetic process | BP | 11 | 0 | 3 | 1.68E-02 |
| GO:0072683 | T cell extravasation | BP | 11 | 0 | 3 | 1.68E-02 |
| GO:0046717 | acid secretion | BP | 89 | 13 | 10 | 1.69E-02 |
| GO:0016887 | ATPase activity | MF | 306 | 17 | 25 | 1.69E-02 |
| GO:0043281 | regulation of cysteine-type endopeptidase activity involved in apop | BP | 172 | 8 | 16 | 1.73E-02 |
| GO:0043407 | negative regulation of MAP kinase activity | BP | 64 | 2 | 8 | 1.73E-02 |
| GO:0022603 | regulation of anatomical structure morphogenesis | BP | 879 | 93 | 60 | 1.74E-02 |
| GO:0032367 | intracellular cholesterol transport | BP | 20 | 1 | 4 | 1.80E-02 |
| GO:0032366 | intracellular sterol transport | BP | 20 | 1 | 4 | 1.80E-02 |
| GO:0035371 | microtubule plus-end | CC | 20 | 0 | 4 | 1.80E-02 |
| GO:0032692 | negative regulation of interleukin-1 production | BP | 20 | 0 | 4 | 1.80E-02 |
| GO:0032727 | positive regulation of interferon-alpha production | BP | 20 | 0 | 4 | 1.80E-02 |
| GO:0032814 | regulation of natural killer cell activation | BP | 20 | 1 | 4 | 1.80E-02 |
| GO:0010569 | regulation of double-strand break repair via homologous recombina | BP | 30 | 1 | 5 | 1.80E-02 |
| GO:0051607 | defense response to virus | BP | 173 | 1 | 16 | 1.81E-02 |
| GO:1901652 | response to peptide | BP | 418 | 41 | 32 | 1.82E-02 |
| GO:0015629 | actin cytoskeleton | CC | 434 | 55 | 33 | 1.82E-02 |
| GO:0101003 | ficolin-1-rich granule membrane | CC | 41 | 3 | 6 | 1.83E-02 |
| GO:0030225 | macrophage differentiation | BP | 41 | 3 | 6 | 1.83E-02 |
| GO:0070265 | necrotic cell death | BP | 41 | 1 | 6 | 1.83E-02 |
| GO:0042531 | positive regulation of tyrosine phosphorylation of STAT protein | BP | 41 | 3 | 6 | 1.83E-02 |
| GO:0098562 | cytoplasmic side of membrane | CC | 159 | 14 | 15 | 1.85E-02 |
| GO:0010243 | response to organonitrogen compound | BP | 814 | 73 | 56 | 1.85E-02 |
| GO:0010648 | negative regulation of cell communication | BP | 1122 | 89 | 74 | 1.86E-02 |
| GO:1903305 | regulation of regulated secretory pathway | BP | 131 | 20 | 13 | 1.86E-02 |
| GO:0002244 | hematopoietic progenitor cell differentiation | BP | 145 | 9 | 14 | 1.86E-02 |
| GO:0051303 | establishment of chromosome localization | BP | 65 | 2 | 8 | 1.88E-02 |
| GO:0070588 | calcium ion transmembrane transport | BP | 248 | 33 | 21 | 1.90E-02 |
| GO:0034599 | cellular response to oxidative stress | BP | 248 | 14 | 21 | 1.90E-02 |
| GO:0002831 | regulation of response to biotic stimulus | BP | 104 | 2 | 11 | 1.90E-02 |
| GO:0051310 | metaphase plate congression | BP | 53 | 2 | 7 | 1.91E-02 |
| GO:0043550 | regulation of lipid kinase activity | BP | 53 | 3 | 7 | 1.91E-02 |
| GO:0030104 | water homeostasis | BP | 53 | 7 | 7 | 1.91E-02 |
| GO:0006281 | DNA repair | BP | 452 | 5 | 34 | 1.93E-02 |
| GO:0055082 | cellular chemical homeostasis | BP | 616 | 65 | 44 | 1.97E-02 |
| GO:0045861 | negative regulation of proteolysis | BP | 234 | 19 | 20 | 1.98E-02 |
| GO:0051259 | protein complex oligomerization | BP | 437 | 30 | 33 | 1.99E-02 |
| GO:0005874 | microtubule | CC | 342 | 20 | 27 | 2.01E-02 |

| | | | | | | |
|---|---|---|---|---|---|---|
| GO:0050701 | interleukin-1 secretion | BP | 42 | 1 | 6 | 2.05E-02 |
| GO:0007080 | mitotic metaphase plate congression | BP | 42 | 0 | 6 | 2.05E-02 |
| GO:0051985 | negative regulation of chromosome segregation | BP | 42 | 0 | 6 | 2.05E-02 |
| GO:0045824 | negative regulation of innate immune response | BP | 42 | 1 | 6 | 2.05E-02 |
| GO:0033046 | negative regulation of sister chromatid segregation | BP | 42 | 0 | 6 | 2.05E-02 |
| GO:0090183 | regulation of kidney development | BP | 42 | 7 | 6 | 2.05E-02 |
| GO:0009409 | response to cold | BP | 42 | 0 | 6 | 2.05E-02 |
| GO:0050000 | chromosome localization | BP | 66 | 2 | 8 | 2.05E-02 |
| GO:0046470 | phosphatidylcholine metabolic process | BP | 66 | 6 | 8 | 2.05E-02 |
| GO:0010948 | negative regulation of cell cycle process | BP | 296 | 4 | 24 | 2.06E-02 |
| GO:0090077 | foam cell differentiation | BP | 31 | 4 | 5 | 2.06E-02 |
| GO:0010742 | macrophage derived foam cell differentiation | BP | 31 | 4 | 5 | 2.06E-02 |
| GO:0002756 | MyD88-independent toll-like receptor signaling pathway | BP | 31 | 0 | 5 | 2.06E-02 |
| GO:0044853 | plasma membrane raft | CC | 92 | 16 | 10 | 2.08E-02 |
| GO:0004620 | phospholipase activity | MF | 79 | 5 | 9 | 2.10E-02 |
| GO:0030032 | lamellipodium assembly | BP | 54 | 4 | 7 | 2.10E-02 |
| GO:0035306 | positive regulation of dephosphorylation | BP | 54 | 4 | 7 | 2.10E-02 |
| GO:2000779 | regulation of double-strand break repair | BP | 54 | 1 | 7 | 2.10E-02 |
| GO:1901698 | response to nitrogen compound | BP | 871 | 79 | 59 | 2.11E-02 |
| GO:0071377 | cellular response to glucagon stimulus | BP | 21 | 4 | 4 | 2.13E-02 |
| GO:0071542 | dopaminergic neuron differentiation | BP | 21 | 1 | 4 | 2.13E-02 |
| GO:0071813 | lipoprotein particle binding | MF | 21 | 1 | 4 | 2.13E-02 |
| GO:0032689 | negative regulation of interferon-gamma production | BP | 21 | 2 | 4 | 2.13E-02 |
| GO:0071814 | protein-lipid complex binding | MF | 21 | 1 | 4 | 2.13E-02 |
| GO:0031935 | regulation of chromatin silencing | BP | 21 | 0 | 4 | 2.13E-02 |
| GO:0045649 | regulation of macrophage differentiation | BP | 21 | 1 | 4 | 2.13E-02 |
| GO:0033044 | regulation of chromosome organization | BP | 297 | 4 | 24 | 2.13E-02 |
| GO:0007015 | actin filament organization | BP | 344 | 36 | 27 | 2.15E-02 |
| GO:1901991 | negative regulation of mitotic cell cycle phase transition | BP | 206 | 3 | 18 | 2.15E-02 |
| GO:0071352 | cellular response to interleukin-2 | BP | 12 | 0 | 3 | 2.16E-02 |
| GO:0002544 | chronic inflammatory response | BP | 12 | 1 | 3 | 2.16E-02 |
| GO:0021871 | forebrain regionalization | BP | 12 | 1 | 3 | 2.16E-02 |
| GO:0097284 | hepatocyte apoptotic process | BP | 12 | 0 | 3 | 2.16E-02 |
| GO:0033549 | MAP kinase phosphatase activity | MF | 12 | 0 | 3 | 2.16E-02 |
| GO:0045342 | MHC class II biosynthetic process | BP | 12 | 0 | 3 | 2.16E-02 |
| GO:0043031 | negative regulation of macrophage activation | BP | 12 | 2 | 3 | 2.16E-02 |
| GO:0001780 | neutrophil homeostasis | BP | 12 | 1 | 3 | 2.16E-02 |
| GO:0050664 | oxidoreductase activity, acting on NAD(P)H, oxygen as acceptor | MF | 12 | 1 | 3 | 2.16E-02 |
| GO:0034393 | positive regulation of smooth muscle cell apoptotic process | BP | 12 | 0 | 3 | 2.16E-02 |
| GO:0060368 | regulation of Fc receptor mediated stimulatory signaling pathway | BP | 12 | 0 | 3 | 2.16E-02 |
| GO:0014048 | regulation of glutamate secretion | BP | 12 | 1 | 3 | 2.16E-02 |
| GO:0045930 | negative regulation of mitotic cell cycle | BP | 282 | 6 | 23 | 2.17E-02 |
| GO:1902806 | regulation of cell cycle G1/S phase transition | BP | 148 | 4 | 14 | 2.19E-02 |
| GO:0003674 | molecular_function | MF | 12188 | 814 | 647 | 2.20E-02 |
| GO:1901699 | cellular response to nitrogen compound | BP | 522 | 50 | 38 | 2.22E-02 |
| GO:0003777 | microtubule motor activity | MF | 67 | 5 | 8 | 2.22E-02 |
| GO:0005788 | endoplasmic reticulum lumen | CC | 252 | 31 | 21 | 2.23E-02 |
| GO:0098655 | cation transmembrane transport | BP | 605 | 84 | 43 | 2.25E-02 |
| GO:1903307 | positive regulation of regulated secretory pathway | BP | 43 | 8 | 6 | 2.28E-02 |
| GO:0002920 | regulation of humoral immune response | BP | 43 | 0 | 6 | 2.28E-02 |
| GO:0004860 | protein kinase inhibitor activity | MF | 55 | 2 | 7 | 2.30E-02 |
| GO:0038095 | Fc-epsilon receptor signaling pathway | BP | 107 | 1 | 11 | 2.30E-02 |
| GO:0017124 | SH3 domain binding | MF | 121 | 3 | 12 | 2.33E-02 |
| GO:0007520 | myoblast fusion | BP | 32 | 7 | 5 | 2.34E-02 |
| GO:0032350 | regulation of hormone metabolic process | BP | 32 | 3 | 5 | 2.34E-02 |
| GO:0009256 | 10-formyltetrahydrofolate metabolic process | BP | 5 | 1 | 2 | 2.43E-02 |
| GO:0002322 | B cell proliferation involved in immune response | BP | 5 | 0 | 2 | 2.43E-02 |
| GO:0021796 | cerebral cortex regionalization | BP | 5 | 0 | 2 | 2.43E-02 |
| GO:0097278 | complement-dependent cytotoxicity | BP | 5 | 0 | 2 | 2.43E-02 |
| GO:0048248 | CXCR3 chemokine receptor binding | MF | 5 | 0 | 2 | 2.43E-02 |
| GO:0019238 | cyclohydrolase activity | MF | 5 | 0 | 2 | 2.43E-02 |
| GO:0097199 | cysteine-type endopeptidase activity involved in apoptotic signaling | MF | 5 | 0 | 2 | 2.43E-02 |
| GO:0044194 | cytolytic granule | CC | 5 | 0 | 2 | 2.43E-02 |
| GO:0033567 | DNA replication, Okazaki fragment processing | BP | 5 | 0 | 2 | 2.43E-02 |

| | | | | | | |
|---|---|---|---|---|---|---|
| GO:0006269 | DNA replication, synthesis of RNA primer | BP | 5 | 0 | 2 | 2.43E-02 |
| GO:0033227 | dsRNA transport | BP | 5 | 0 | 2 | 2.43E-02 |
| GO:0004931 | extracellularly ATP-gated cation channel activity | MF | 5 | 0 | 2 | 2.43E-02 |
| GO:0001652 | granular component | CC | 5 | 0 | 2 | 2.43E-02 |
| GO:0036042 | long-chain fatty acyl-CoA binding | MF | 5 | 0 | 2 | 2.43E-02 |
| GO:0060374 | mast cell differentiation | BP | 5 | 1 | 2 | 2.43E-02 |
| GO:0014004 | microglia differentiation | BP | 5 | 1 | 2 | 2.43E-02 |
| GO:0052150 | modulation by symbiont of host apoptotic process | BP | 5 | 0 | 2 | 2.43E-02 |
| GO:2000562 | negative regulation of CD4-positive, alpha-beta T cell proliferation | BP | 5 | 0 | 2 | 2.43E-02 |
| GO:0045953 | negative regulation of natural killer cell mediated cytotoxicity | BP | 5 | 0 | 2 | 2.43E-02 |
| GO:0002725 | negative regulation of T cell cytokine production | BP | 5 | 0 | 2 | 2.43E-02 |
| GO:0046934 | phosphatidylinositol-4,5-bisphosphate 3-kinase activity | MF | 5 | 0 | 2 | 2.43E-02 |
| GO:0052813 | phosphatidylinositol bisphosphate kinase activity | MF | 5 | 0 | 2 | 2.43E-02 |
| GO:2000987 | positive regulation of behavioral fear response | BP | 5 | 2 | 2 | 2.43E-02 |
| GO:1903367 | positive regulation of fear response | BP | 5 | 2 | 2 | 2.43E-02 |
| GO:0002857 | positive regulation of natural killer cell mediated immune response | BP | 5 | 1 | 2 | 2.43E-02 |
| GO:1903911 | positive regulation of receptor clustering | BP | 5 | 1 | 2 | 2.43E-02 |
| GO:1904058 | positive regulation of sensory perception of pain | BP | 5 | 0 | 2 | 2.43E-02 |
| GO:2000556 | positive regulation of T-helper 1 cell cytokine production | BP | 5 | 0 | 2 | 2.43E-02 |
| GO:1990405 | protein antigen binding | MF | 5 | 0 | 2 | 2.43E-02 |
| GO:0032596 | protein transport into membrane raft | BP | 5 | 0 | 2 | 2.43E-02 |
| GO:0010533 | regulation of activation of Janus kinase activity | BP | 5 | 1 | 2 | 2.43E-02 |
| GO:0002676 | regulation of chronic inflammatory response | BP | 5 | 0 | 2 | 2.43E-02 |
| GO:1905462 | regulation of DNA duplex unwinding | BP | 5 | 0 | 2 | 2.43E-02 |
| GO:0045586 | regulation of gamma-delta T cell differentiation | BP | 5 | 0 | 2 | 2.43E-02 |
| GO:1902412 | regulation of mitotic cytokinesis | BP | 5 | 0 | 2 | 2.43E-02 |
| GO:2000554 | regulation of T-helper 1 cell cytokine production | BP | 5 | 0 | 2 | 2.43E-02 |
| GO:0010041 | response to iron(III) ion | BP | 5 | 0 | 2 | 2.43E-02 |
| GO:0002457 | T cell antigen processing and presentation | BP | 5 | 0 | 2 | 2.43E-02 |
| GO:0044691 | tooth eruption | BP | 5 | 0 | 2 | 2.43E-02 |
| GO:0031802 | type 5 metabotropic glutamate receptor binding | MF | 5 | 1 | 2 | 2.43E-02 |
| GO:0033093 | Weibel-Palade body | CC | 5 | 1 | 2 | 2.43E-02 |
| GO:0000403 | Y-form DNA binding | MF | 5 | 0 | 2 | 2.43E-02 |
| GO:0044419 | interspecies interaction between organisms | BP | 776 | 13 | 53 | 2.43E-02 |
| GO:1903046 | meiotic cell cycle process | BP | 122 | 2 | 12 | 2.47E-02 |
| GO:0071375 | cellular response to peptide hormone stimulus | BP | 255 | 21 | 21 | 2.50E-02 |
| GO:0006541 | glutamine metabolic process | BP | 22 | 0 | 4 | 2.50E-02 |
| GO:0030159 | receptor signaling complex scaffold activity | MF | 22 | 1 | 4 | 2.50E-02 |
| GO:0090398 | cellular senescence | BP | 56 | 0 | 7 | 2.52E-02 |
| GO:0003684 | damaged DNA binding | MF | 56 | 1 | 7 | 2.52E-02 |
| GO:0008154 | actin polymerization or depolymerization | BP | 180 | 18 | 16 | 2.53E-02 |
| GO:1901615 | organic hydroxy compound metabolic process | BP | 413 | 41 | 31 | 2.53E-02 |
| GO:0051276 | chromosome organization | BP | 966 | 18 | 64 | 2.54E-02 |
| GO:0032559 | adenyl ribonucleotide binding | MF | 1279 | 81 | 82 | 2.54E-02 |
| GO:0000075 | cell cycle checkpoint | BP | 195 | 2 | 17 | 2.55E-02 |
| GO:0008017 | microtubule binding | MF | 210 | 16 | 18 | 2.55E-02 |
| GO:0051897 | positive regulation of protein kinase B signaling | BP | 123 | 11 | 12 | 2.61E-02 |
| GO:0006979 | response to oxidative stress | BP | 366 | 24 | 28 | 2.61E-02 |
| GO:1990939 | ATP-dependent microtubule motor activity | MF | 33 | 2 | 5 | 2.65E-02 |
| GO:0007094 | mitotic spindle assembly checkpoint | BP | 33 | 0 | 5 | 2.65E-02 |
| GO:0071174 | mitotic spindle checkpoint | BP | 33 | 0 | 5 | 2.65E-02 |
| GO:0002833 | positive regulation of response to biotic stimulus | BP | 33 | 1 | 5 | 2.65E-02 |
| GO:0030449 | regulation of complement activation | BP | 33 | 0 | 5 | 2.65E-02 |
| GO:0033003 | regulation of mast cell activation | BP | 33 | 2 | 5 | 2.65E-02 |
| GO:0071173 | spindle assembly checkpoint | BP | 33 | 0 | 5 | 2.65E-02 |
| GO:0031577 | spindle checkpoint | BP | 33 | 0 | 5 | 2.65E-02 |
| GO:0007599 | hemostasis | BP | 257 | 19 | 21 | 2.69E-02 |
| GO:0050700 | CARD domain binding | MF | 13 | 0 | 3 | 2.70E-02 |
| GO:0071350 | cellular response to interleukin-15 | BP | 13 | 1 | 3 | 2.70E-02 |
| GO:0000778 | condensed nuclear chromosome kinetochore | CC | 13 | 0 | 3 | 2.70E-02 |
| GO:0043650 | dicarboxylic acid biosynthetic process | BP | 13 | 0 | 3 | 2.70E-02 |
| GO:0008301 | DNA binding, bending | MF | 13 | 0 | 3 | 2.70E-02 |
| GO:0035723 | interleukin-15-mediated signaling pathway | BP | 13 | 1 | 3 | 2.70E-02 |
| GO:1900451 | positive regulation of glutamate receptor signaling pathway | BP | 13 | 0 | 3 | 2.70E-02 |

| | | | | | | |
|---|---|---|---|---|---|---|
| GO:1901741 | positive regulation of myoblast fusion | BP | 13 | 1 | 3 | 2.70E-02 |
| GO:0030852 | regulation of granulocyte differentiation | BP | 13 | 0 | 3 | 2.70E-02 |
| GO:0070669 | response to interleukin-2 | BP | 13 | 0 | 3 | 2.70E-02 |
| GO:0051775 | response to redox state | BP | 13 | 1 | 3 | 2.70E-02 |
| GO:0045064 | T-helper 2 cell differentiation | BP | 13 | 0 | 3 | 2.70E-02 |
| GO:0030162 | regulation of proteolysis | BP | 563 | 27 | 40 | 2.73E-02 |
| GO:0030139 | endocytic vesicle | CC | 242 | 9 | 20 | 2.73E-02 |
| GO:0030833 | regulation of actin filament polymerization | BP | 138 | 12 | 13 | 2.73E-02 |
| GO:0030100 | regulation of endocytosis | BP | 227 | 14 | 19 | 2.76E-02 |
| GO:1904064 | positive regulation of cation transmembrane transport | BP | 110 | 19 | 11 | 2.76E-02 |
| GO:0043254 | regulation of protein complex assembly | BP | 400 | 24 | 30 | 2.77E-02 |
| GO:1901990 | regulation of mitotic cell cycle phase transition | BP | 368 | 7 | 28 | 2.78E-02 |
| GO:0043536 | positive regulation of blood vessel endothelial cell migration | BP | 45 | 3 | 6 | 2.79E-02 |
| GO:0043551 | regulation of phosphatidylinositol 3-kinase activity | BP | 45 | 3 | 6 | 2.79E-02 |
| GO:0000079 | regulation of cyclin-dependent protein serine/threonine kinase activ | BP | 83 | 4 | 9 | 2.79E-02 |
| GO:0032088 | negative regulation of NF-kappaB transcription factor activity | BP | 70 | 0 | 8 | 2.81E-02 |
| GO:0005773 | vacuole | CC | 631 | 28 | 44 | 2.82E-02 |
| GO:0034762 | regulation of transmembrane transport | BP | 417 | 73 | 31 | 2.84E-02 |
| GO:1990837 | sequence-specific double-stranded DNA binding | MF | 665 | 44 | 46 | 2.84E-02 |
| GO:0006810 | transport | BP | 3967 | 303 | 229 | 2.85E-02 |
| GO:0000902 | cell morphogenesis | BP | 869 | 112 | 58 | 2.85E-02 |
| GO:0046982 | protein heterodimerization activity | MF | 337 | 31 | 26 | 2.86E-02 |
| GO:0072686 | mitotic spindle | CC | 97 | 4 | 10 | 2.89E-02 |
| GO:0010661 | positive regulation of muscle cell apoptotic process | BP | 23 | 2 | 4 | 2.91E-02 |
| GO:0070723 | response to cholesterol | BP | 23 | 0 | 4 | 2.91E-02 |
| GO:0050909 | sensory perception of taste | BP | 23 | 3 | 4 | 2.91E-02 |
| GO:1901653 | cellular response to peptide | BP | 306 | 29 | 24 | 2.91E-02 |
| GO:0031225 | anchored component of membrane | CC | 111 | 24 | 11 | 2.93E-02 |
| GO:0019904 | protein domain specific binding | MF | 616 | 32 | 43 | 2.93E-02 |
| GO:0030278 | regulation of ossification | BP | 154 | 17 | 14 | 2.95E-02 |
| GO:0003779 | actin binding | MF | 370 | 56 | 28 | 2.95E-02 |
| GO:0051726 | regulation of cell cycle | BP | 1009 | 34 | 66 | 2.97E-02 |
| GO:2000257 | regulation of protein activation cascade | BP | 34 | 0 | 5 | 2.98E-02 |
| GO:0071496 | cellular response to external stimulus | BP | 291 | 24 | 23 | 2.99E-02 |
| GO:0010660 | regulation of muscle cell apoptotic process | BP | 58 | 8 | 7 | 2.99E-02 |
| GO:0071897 | DNA biosynthetic process | BP | 169 | 4 | 15 | 3.00E-02 |
| GO:0060993 | kidney morphogenesis | BP | 71 | 9 | 8 | 3.03E-02 |
| GO:0097581 | lamellipodium organization | BP | 71 | 6 | 8 | 3.03E-02 |
| GO:0097006 | regulation of plasma lipoprotein particle levels | BP | 71 | 4 | 8 | 3.03E-02 |
| GO:0005925 | focal adhesion | CC | 387 | 36 | 29 | 3.04E-02 |
| GO:0030554 | adenyl nucleotide binding | MF | 1290 | 82 | 82 | 3.04E-02 |
| GO:0097755 | positive regulation of blood vessel diameter | BP | 46 | 10 | 6 | 3.07E-02 |
| GO:0006301 | postreplication repair | BP | 46 | 0 | 6 | 3.07E-02 |
| GO:0030216 | keratinocyte differentiation | BP | 98 | 6 | 10 | 3.08E-02 |
| GO:0005829 | cytosol | CC | 4237 | 217 | 243 | 3.08E-02 |
| GO:1905477 | positive regulation of protein localization to membrane | BP | 112 | 11 | 11 | 3.10E-02 |
| GO:0031098 | stress-activated protein kinase signaling cascade | BP | 261 | 13 | 21 | 3.11E-02 |
| GO:0030838 | positive regulation of actin filament polymerization | BP | 85 | 6 | 9 | 3.20E-02 |
| GO:0005924 | cell-substrate adherens junction | CC | 389 | 37 | 29 | 3.22E-02 |
| GO:0030971 | receptor tyrosine kinase binding | MF | 59 | 2 | 7 | 3.24E-02 |
| GO:0031348 | negative regulation of defense response | BP | 156 | 6 | 14 | 3.25E-02 |
| GO:0010952 | positive regulation of peptidase activity | BP | 156 | 7 | 14 | 3.25E-02 |
| GO:0034764 | positive regulation of transmembrane transport | BP | 156 | 27 | 14 | 3.25E-02 |
| GO:1901988 | negative regulation of cell cycle phase transition | BP | 216 | 3 | 18 | 3.26E-02 |
| GO:1903725 | regulation of phospholipid metabolic process | BP | 72 | 5 | 8 | 3.26E-02 |
| GO:0030218 | erythrocyte differentiation | BP | 99 | 4 | 10 | 3.27E-02 |
| GO:0045335 | phagocytic vesicle | CC | 113 | 3 | 11 | 3.28E-02 |
| GO:0002438 | acute inflammatory response to antigenic stimulus | BP | 14 | 2 | 3 | 3.31E-02 |
| GO:0035458 | cellular response to interferon-beta | BP | 14 | 0 | 3 | 3.31E-02 |
| GO:0006266 | DNA ligation | BP | 14 | 0 | 3 | 3.31E-02 |
| GO:0070365 | hepatocyte differentiation | BP | 14 | 2 | 3 | 3.31E-02 |
| GO:0001771 | immunological synapse formation | BP | 14 | 0 | 3 | 3.31E-02 |
| GO:0002385 | mucosal immune response | BP | 14 | 0 | 3 | 3.31E-02 |
| GO:0030889 | negative regulation of B cell proliferation | BP | 14 | 0 | 3 | 3.31E-02 |

| | | | | | | |
|---|---|---|---|---|---|---|
| GO:0036152 | phosphatidylethanolamine acyl-chain remodeling | BP | 14 | 1 | 3 | 3.31E-02 |
| GO:0016307 | phosphatidylinositol phosphate kinase activity | MF | 14 | 0 | 3 | 3.31E-02 |
| GO:0042104 | positive regulation of activated T cell proliferation | BP | 14 | 0 | 3 | 3.31E-02 |
| GO:0051957 | positive regulation of amino acid transport | BP | 14 | 2 | 3 | 3.31E-02 |
| GO:2000484 | positive regulation of interleukin-8 secretion | BP | 14 | 0 | 3 | 3.31E-02 |
| GO:0070230 | positive regulation of lymphocyte apoptotic process | BP | 14 | 0 | 3 | 3.31E-02 |
| GO:0032816 | positive regulation of natural killer cell activation | BP | 14 | 0 | 3 | 3.31E-02 |
| GO:0002830 | positive regulation of type 2 immune response | BP | 14 | 0 | 3 | 3.31E-02 |
| GO:0042069 | regulation of catecholamine metabolic process | BP | 14 | 2 | 3 | 3.31E-02 |
| GO:0042053 | regulation of dopamine metabolic process | BP | 14 | 2 | 3 | 3.31E-02 |
| GO:0090266 | regulation of mitotic cell cycle spindle assembly checkpoint | BP | 14 | 0 | 3 | 3.31E-02 |
| GO:1903504 | regulation of mitotic spindle checkpoint | BP | 14 | 0 | 3 | 3.31E-02 |
| GO:0010155 | regulation of proton transport | BP | 14 | 0 | 3 | 3.31E-02 |
| GO:0090231 | regulation of spindle checkpoint | BP | 14 | 0 | 3 | 3.31E-02 |
| GO:0070672 | response to interleukin-15 | BP | 14 | 1 | 3 | 3.31E-02 |
| GO:0035859 | Seh1-associated complex | CC | 14 | 0 | 3 | 3.31E-02 |
| GO:0034134 | toll-like receptor 2 signaling pathway | BP | 14 | 0 | 3 | 3.31E-02 |
| GO:0009712 | catechol-containing compound metabolic process | BP | 35 | 9 | 5 | 3.33E-02 |
| GO:0006584 | catecholamine metabolic process | BP | 35 | 9 | 5 | 3.33E-02 |
| GO:0015026 | coreceptor activity | MF | 35 | 4 | 5 | 3.33E-02 |
| GO:0002755 | MyD88-dependent toll-like receptor signaling pathway | BP | 35 | 1 | 5 | 3.33E-02 |
| GO:1902100 | negative regulation of metaphase/anaphase transition of cell cycle | BP | 35 | 0 | 5 | 3.33E-02 |
| GO:0045841 | negative regulation of mitotic metaphase/anaphase transition | BP | 35 | 0 | 5 | 3.33E-02 |
| GO:0032731 | positive regulation of interleukin-1 beta production | BP | 35 | 0 | 5 | 3.33E-02 |
| GO:2000249 | regulation of actin cytoskeleton reorganization | BP | 35 | 1 | 5 | 3.33E-02 |
| GO:0048538 | thymus development | BP | 35 | 2 | 5 | 3.33E-02 |
| GO:0060627 | regulation of vesicle-mediated transport | BP | 439 | 45 | 32 | 3.33E-02 |
| GO:2001233 | regulation of apoptotic signaling pathway | BP | 342 | 14 | 26 | 3.34E-02 |
| GO:0060706 | cell differentiation involved in embryonic placenta development | BP | 24 | 2 | 4 | 3.35E-02 |
| GO:0006760 | folic acid-containing compound metabolic process | BP | 24 | 2 | 4 | 3.35E-02 |
| GO:0048873 | homeostasis of number of cells within a tissue | BP | 24 | 0 | 4 | 3.35E-02 |
| GO:0016814 | hydrolase activity, acting on carbon-nitrogen (but not peptide) bonc | MF | 24 | 0 | 4 | 3.35E-02 |
| GO:0004435 | phosphatidylinositol phospholipase C activity | MF | 24 | 2 | 4 | 3.35E-02 |
| GO:0031297 | replication fork processing | BP | 24 | 0 | 4 | 3.35E-02 |
| GO:1990776 | response to angiotensin | BP | 24 | 5 | 4 | 3.35E-02 |
| GO:0005164 | tumor necrosis factor receptor binding | MF | 24 | 0 | 4 | 3.35E-02 |
| GO:0042311 | vasodilation | BP | 24 | 4 | 4 | 3.35E-02 |
| GO:0007606 | sensory perception of chemical stimulus | BP | 47 | 4 | 6 | 3.37E-02 |
| GO:0048002 | antigen processing and presentation of peptide antigen | BP | 157 | 5 | 14 | 3.40E-02 |
| GO:0008081 | phosphoric diester hydrolase activity | MF | 86 | 8 | 9 | 3.41E-02 |
| GO:0014074 | response to purine-containing compound | BP | 128 | 17 | 12 | 3.42E-02 |
| GO:0044089 | positive regulation of cellular component biogenesis | BP | 473 | 49 | 34 | 3.43E-02 |
| GO:0042060 | wound healing | BP | 424 | 39 | 31 | 3.45E-02 |
| GO:0046330 | positive regulation of JNK cascade | BP | 114 | 5 | 11 | 3.47E-02 |
| GO:0046849 | bone remodeling | BP | 73 | 5 | 8 | 3.51E-02 |
| GO:0030055 | cell-substrate junction | CC | 392 | 38 | 29 | 3.51E-02 |
| GO:0000922 | spindle pole | CC | 143 | 3 | 13 | 3.51E-02 |
| GO:0030219 | megakaryocyte differentiation | BP | 60 | 3 | 7 | 3.51E-02 |
| GO:0000147 | actin cortical patch assembly | BP | 6 | 0 | 2 | 3.52E-02 |
| GO:0044396 | actin cortical patch organization | BP | 6 | 0 | 2 | 3.52E-02 |
| GO:0043615 | astrocyte cell migration | BP | 6 | 0 | 2 | 3.52E-02 |
| GO:0035381 | ATP-gated ion channel activity | MF | 6 | 0 | 2 | 3.52E-02 |
| GO:0031428 | box C/D snoRNP complex | CC | 6 | 0 | 2 | 3.52E-02 |
| GO:0047498 | calcium-dependent phospholipase A2 activity | MF | 6 | 1 | 2 | 3.52E-02 |
| GO:0031748 | D1 dopamine receptor binding | MF | 6 | 0 | 2 | 3.52E-02 |
| GO:1990130 | GATOR1 complex | CC | 6 | 0 | 2 | 3.52E-02 |
| GO:0008035 | high-density lipoprotein particle binding | MF | 6 | 1 | 2 | 3.52E-02 |
| GO:0002349 | histamine production involved in inflammatory response | BP | 6 | 0 | 2 | 3.52E-02 |
| GO:0002553 | histamine secretion by mast cell | BP | 6 | 0 | 2 | 3.52E-02 |
| GO:0002441 | histamine secretion involved in inflammatory response | BP | 6 | 0 | 2 | 3.52E-02 |
| GO:0038155 | interleukin-23-mediated signaling pathway | BP | 6 | 0 | 2 | 3.52E-02 |
| GO:0005138 | interleukin-6 receptor binding | MF | 6 | 0 | 2 | 3.52E-02 |
| GO:0000800 | lateral element | CC | 6 | 0 | 2 | 3.52E-02 |
| GO:0010216 | maintenance of DNA methylation | BP | 6 | 0 | 2 | 3.52E-02 |

| | | | | | | |
|---|---|---|---|---|---|---|
| GO:0060137 | maternal process involved in parturition | BP | 6 | 1 | 2 | 3.52E-02 |
| GO:0010032 | meiotic chromosome condensation | BP | 6 | 0 | 2 | 3.52E-02 |
| GO:0052433 | modulation by organism of apoptotic process in other organism inv | BP | 6 | 0 | 2 | 3.52E-02 |
| GO:0052040 | modulation by symbiont of host programmed cell death | BP | 6 | 0 | 2 | 3.52E-02 |
| GO:0044532 | modulation of apoptotic process in other organism | BP | 6 | 0 | 2 | 3.52E-02 |
| GO:0044531 | modulation of programmed cell death in other organism | BP | 6 | 0 | 2 | 3.52E-02 |
| GO:0052248 | modulation of programmed cell death in other organism involved in | BP | 6 | 0 | 2 | 3.52E-02 |
| GO:0002423 | natural killer cell mediated immune response to tumor cell | BP | 6 | 1 | 2 | 3.52E-02 |
| GO:0002578 | negative regulation of antigen processing and presentation | BP | 6 | 0 | 2 | 3.52E-02 |
| GO:0033239 | negative regulation of cellular amine metabolic process | BP | 6 | 1 | 2 | 3.52E-02 |
| GO:1903979 | negative regulation of microglial cell activation | BP | 6 | 1 | 2 | 3.52E-02 |
| GO:0002716 | negative regulation of natural killer cell mediated immunity | BP | 6 | 0 | 2 | 3.52E-02 |
| GO:0034392 | negative regulation of smooth muscle cell apoptotic process | BP | 6 | 1 | 2 | 3.52E-02 |
| GO:0002826 | negative regulation of T-helper 1 type immune response | BP | 6 | 0 | 2 | 3.52E-02 |
| GO:0034144 | negative regulation of toll-like receptor 4 signaling pathway | BP | 6 | 1 | 2 | 3.52E-02 |
| GO:0002158 | osteoclast proliferation | BP | 6 | 1 | 2 | 3.52E-02 |
| GO:0046469 | platelet activating factor metabolic process | BP | 6 | 0 | 2 | 3.52E-02 |
| GO:0002866 | positive regulation of acute inflammatory response to antigenic stin | BP | 6 | 1 | 2 | 3.52E-02 |
| GO:0050861 | positive regulation of B cell receptor signaling pathway | BP | 6 | 0 | 2 | 3.52E-02 |
| GO:2000373 | positive regulation of DNA topoisomerase (ATP-hydrolyzing) activ | BP | 6 | 0 | 2 | 3.52E-02 |
| GO:2000553 | positive regulation of T-helper 2 cell cytokine production | BP | 6 | 0 | 2 | 3.52E-02 |
| GO:0002645 | positive regulation of tolerance induction | BP | 6 | 0 | 2 | 3.52E-02 |
| GO:2000822 | regulation of behavioral fear response | BP | 6 | 2 | 2 | 3.52E-02 |
| GO:2000371 | regulation of DNA topoisomerase (ATP-hydrolyzing) activity | BP | 6 | 0 | 2 | 3.52E-02 |
| GO:1901509 | regulation of endothelial tube morphogenesis | BP | 6 | 2 | 2 | 3.52E-02 |
| GO:1903365 | regulation of fear response | BP | 6 | 2 | 2 | 3.52E-02 |
| GO:0046643 | regulation of gamma-delta T cell activation | BP | 6 | 0 | 2 | 3.52E-02 |
| GO:0010728 | regulation of hydrogen peroxide biosynthetic process | BP | 6 | 0 | 2 | 3.52E-02 |
| GO:0002883 | regulation of hypersensitivity | BP | 6 | 0 | 2 | 3.52E-02 |
| GO:0002855 | regulation of natural killer cell mediated immune response to tumor | BP | 6 | 1 | 2 | 3.52E-02 |
| GO:1901339 | regulation of store-operated calcium channel activity | BP | 6 | 0 | 2 | 3.52E-02 |
| GO:0005856 | cytoskeleton | CC | 1709 | 131 | 105 | 3.54E-02 |
| GO:1904029 | regulation of cyclin-dependent protein kinase activity | BP | 87 | 4 | 9 | 3.64E-02 |
| GO:0045839 | negative regulation of mitotic nuclear division | BP | 48 | 0 | 6 | 3.69E-02 |
| GO:0060675 | ureteric bud morphogenesis | BP | 48 | 5 | 6 | 3.69E-02 |
| GO:0002762 | negative regulation of myeloid leukocyte differentiation | BP | 36 | 1 | 5 | 3.71E-02 |
| GO:0005547 | phosphatidylinositol-3,4,5-trisphosphate binding | MF | 36 | 2 | 5 | 3.71E-02 |
| GO:0043331 | response to dsRNA | BP | 36 | 0 | 5 | 3.71E-02 |
| GO:0006812 | cation transport | BP | 831 | 121 | 55 | 3.71E-02 |
| GO:0005524 | ATP binding | MF | 1232 | 76 | 78 | 3.71E-02 |
| GO:0051015 | actin filament binding | MF | 174 | 26 | 15 | 3.74E-02 |
| GO:0031497 | chromatin assembly | BP | 74 | 3 | 8 | 3.76E-02 |
| GO:0010657 | muscle cell apoptotic process | BP | 61 | 9 | 7 | 3.80E-02 |
| GO:0090734 | site of DNA damage | CC | 61 | 0 | 7 | 3.80E-02 |
| GO:1901617 | organic hydroxy compound biosynthetic process | BP | 220 | 20 | 18 | 3.80E-02 |
| GO:1990777 | lipoprotein particle | CC | 25 | 0 | 4 | 3.83E-02 |
| GO:0016645 | oxidoreductase activity, acting on the CH-NH group of donors | MF | 25 | 4 | 4 | 3.83E-02 |
| GO:0005942 | phosphatidylinositol 3-kinase complex | CC | 25 | 0 | 4 | 3.83E-02 |
| GO:0034358 | plasma lipoprotein particle | CC | 25 | 0 | 4 | 3.83E-02 |
| GO:0051482 | positive regulation of cytosolic calcium ion concentration involved | BP | 25 | 5 | 4 | 3.83E-02 |
| GO:0051569 | regulation of histone H3-K4 methylation | BP | 25 | 0 | 4 | 3.83E-02 |
| GO:0045191 | regulation of isotype switching | BP | 25 | 0 | 4 | 3.83E-02 |
| GO:0010591 | regulation of lamellipodium assembly | BP | 25 | 2 | 4 | 3.83E-02 |
| GO:0030894 | replisome | CC | 25 | 0 | 4 | 3.83E-02 |
| GO:0032094 | response to food | BP | 25 | 2 | 4 | 3.83E-02 |
| GO:0030855 | epithelial cell differentiation | BP | 461 | 37 | 33 | 3.85E-02 |
| GO:0046165 | alcohol biosynthetic process | BP | 145 | 12 | 13 | 3.87E-02 |
| GO:0070997 | neuron death | BP | 283 | 21 | 22 | 3.87E-02 |
| GO:0007569 | cell aging | BP | 88 | 2 | 9 | 3.87E-02 |
| GO:0072089 | stem cell proliferation | BP | 88 | 8 | 9 | 3.87E-02 |
| GO:0001503 | ossification | BP | 299 | 31 | 23 | 3.89E-02 |
| GO:0048532 | anatomical structure arrangement | BP | 15 | 4 | 3 | 3.98E-02 |
| GO:0009713 | catechol-containing compound biosynthetic process | BP | 15 | 3 | 3 | 3.98E-02 |
| GO:0042423 | catecholamine biosynthetic process | BP | 15 | 3 | 3 | 3.98E-02 |

| | | | | | | |
|---|---|---|---|---|---|---|
| GO:0038111 | interleukin-7-mediated signaling pathway | BP | 15 | 1 | 3 | 3.98E-02 |
| GO:0048535 | lymph node development | BP | 15 | 1 | 3 | 3.98E-02 |
| GO:0070233 | negative regulation of T cell apoptotic process | BP | 15 | 1 | 3 | 3.98E-02 |
| GO:0015037 | peptide disulfide oxidoreductase activity | MF | 15 | 0 | 3 | 3.98E-02 |
| GO:1902004 | positive regulation of amyloid-beta formation | BP | 15 | 3 | 3 | 3.98E-02 |
| GO:0002693 | positive regulation of cellular extravasation | BP | 15 | 1 | 3 | 3.98E-02 |
| GO:2000696 | regulation of epithelial cell differentiation involved in kidney devel | BP | 15 | 4 | 3 | 3.98E-02 |
| GO:0010715 | regulation of extracellular matrix disassembly | BP | 15 | 2 | 3 | 3.98E-02 |
| GO:0010310 | regulation of hydrogen peroxide metabolic process | BP | 15 | 0 | 3 | 3.98E-02 |
| GO:1901739 | regulation of myoblast fusion | BP | 15 | 1 | 3 | 3.98E-02 |
| GO:2000316 | regulation of T-helper 17 type immune response | BP | 15 | 0 | 3 | 3.98E-02 |
| GO:0034143 | regulation of toll-like receptor 4 signaling pathway | BP | 15 | 1 | 3 | 3.98E-02 |
| GO:0072331 | signal transduction by p53 class mediator | BP | 237 | 5 | 19 | 4.02E-02 |
| GO:0045453 | bone resorption | BP | 49 | 1 | 6 | 4.03E-02 |
| GO:0072171 | mesonephric tubule morphogenesis | BP | 49 | 6 | 6 | 4.03E-02 |
| GO:0010518 | positive regulation of phospholipase activity | BP | 49 | 6 | 6 | 4.03E-02 |
| GO:0035861 | site of double-strand break | CC | 49 | 0 | 6 | 4.03E-02 |
| GO:0009991 | response to extracellular stimulus | BP | 430 | 37 | 31 | 4.05E-02 |
| GO:0060429 | epithelium development | BP | 888 | 70 | 58 | 4.09E-02 |
| GO:0080135 | regulation of cellular response to stress | BP | 614 | 23 | 42 | 4.09E-02 |
| GO:0072562 | blood microparticle | CC | 62 | 3 | 7 | 4.10E-02 |
| GO:0030670 | phagocytic vesicle membrane | CC | 62 | 1 | 7 | 4.10E-02 |
| GO:0030295 | protein kinase activator activity | MF | 62 | 3 | 7 | 4.10E-02 |
| GO:0010564 | regulation of cell cycle process | BP | 648 | 17 | 44 | 4.10E-02 |
| GO:0014047 | glutamate secretion | BP | 37 | 6 | 5 | 4.11E-02 |
| GO:0016684 | oxidoreductase activity, acting on peroxide as acceptor | MF | 37 | 3 | 5 | 4.11E-02 |
| GO:1902743 | regulation of lamellipodium organization | BP | 37 | 4 | 5 | 4.11E-02 |
| GO:0032813 | tumor necrosis factor receptor superfamily binding | MF | 37 | 0 | 5 | 4.11E-02 |
| GO:0051093 | negative regulation of developmental process | BP | 717 | 73 | 48 | 4.18E-02 |
| GO:0030001 | metal ion transport | BP | 666 | 97 | 45 | 4.19E-02 |
| GO:1903169 | regulation of calcium ion transmembrane transport | BP | 118 | 15 | 11 | 4.29E-02 |
| GO:0045596 | negative regulation of cell differentiation | BP | 532 | 53 | 37 | 4.30E-02 |
| GO:0033045 | regulation of sister chromatid segregation | BP | 76 | 0 | 8 | 4.30E-02 |
| GO:0007190 | activation of adenylate cyclase activity | BP | 26 | 4 | 4 | 4.35E-02 |
| GO:0097431 | mitotic spindle pole | CC | 26 | 2 | 4 | 4.35E-02 |
| GO:0045736 | negative regulation of cyclin-dependent protein serine/threonine kir | BP | 26 | 3 | 4 | 4.35E-02 |
| GO:0002701 | negative regulation of production of molecular mediator of immune | BP | 26 | 0 | 4 | 4.35E-02 |
| GO:1905332 | positive regulation of morphogenesis of an epithelium | BP | 26 | 2 | 4 | 4.35E-02 |
| GO:0010743 | regulation of macrophage derived foam cell differentiation | BP | 26 | 3 | 4 | 4.35E-02 |
| GO:0055094 | response to lipoprotein particle | BP | 26 | 2 | 4 | 4.35E-02 |
| GO:0036314 | response to sterol | BP | 26 | 0 | 4 | 4.35E-02 |
| GO:0043434 | response to peptide hormone | BP | 351 | 33 | 26 | 4.36E-02 |
| GO:0021782 | glial cell development | BP | 90 | 11 | 9 | 4.38E-02 |
| GO:0098662 | inorganic cation transmembrane transport | BP | 533 | 74 | 37 | 4.40E-02 |
| GO:0022617 | extracellular matrix disassembly | BP | 63 | 3 | 7 | 4.41E-02 |
| GO:0065003 | protein-containing complex assembly | BP | 1546 | 88 | 95 | 4.42E-02 |
| GO:0097305 | response to alcohol | BP | 178 | 22 | 15 | 4.43E-02 |
| GO:0007346 | regulation of mitotic cell cycle | BP | 551 | 17 | 38 | 4.52E-02 |
| GO:0070192 | chromosome organization involved in meiotic cell cycle | BP | 38 | 0 | 5 | 4.54E-02 |
| GO:2000107 | negative regulation of leukocyte apoptotic process | BP | 38 | 2 | 5 | 4.54E-02 |
| GO:0010863 | positive regulation of phospholipase C activity | BP | 38 | 5 | 5 | 4.54E-02 |
| GO:0043030 | regulation of macrophage activation | BP | 38 | 2 | 5 | 4.54E-02 |
| GO:0012505 | endomembrane system | CC | 3492 | 212 | 201 | 4.60E-02 |
| GO:0032993 | protein-DNA complex | CC | 91 | 0 | 9 | 4.64E-02 |
| GO:0017048 | Rho GTPase binding | MF | 164 | 12 | 14 | 4.64E-02 |
| GO:0008356 | asymmetric cell division | BP | 16 | 0 | 3 | 4.72E-02 |
| GO:0051131 | chaperone-mediated protein complex assembly | BP | 16 | 0 | 3 | 4.72E-02 |
| GO:0070182 | DNA polymerase binding | MF | 16 | 0 | 3 | 4.72E-02 |
| GO:0009084 | glutamine family amino acid biosynthetic process | BP | 16 | 0 | 3 | 4.72E-02 |
| GO:0016864 | intramolecular oxidoreductase activity, transposing S-S bonds | MF | 16 | 1 | 3 | 4.72E-02 |
| GO:0030011 | maintenance of cell polarity | BP | 16 | 2 | 3 | 4.72E-02 |
| GO:0007076 | mitotic chromosome condensation | BP | 16 | 0 | 3 | 4.72E-02 |
| GO:0002251 | organ or tissue specific immune response | BP | 16 | 0 | 3 | 4.72E-02 |
| GO:0004623 | phospholipase A2 activity | MF | 16 | 1 | 3 | 4.72E-02 |

| | | | | | | |
|---|---|---|---|---|---|---|
| GO:0010592 | positive regulation of lamellipodium assembly | BP | 16 | 0 | 3 | 4.72E-02 |
| GO:0032930 | positive regulation of superoxide anion generation | BP | 16 | 1 | 3 | 4.72E-02 |
| GO:0003756 | protein disulfide isomerase activity | MF | 16 | 1 | 3 | 4.72E-02 |
| GO:0034975 | protein folding in endoplasmic reticulum | BP | 16 | 0 | 3 | 4.72E-02 |
| GO:0002577 | regulation of antigen processing and presentation | BP | 16 | 0 | 3 | 4.72E-02 |
| GO:0042481 | regulation of odontogenesis | BP | 16 | 1 | 3 | 4.72E-02 |
| GO:0097503 | sialylation | BP | 16 | 3 | 3 | 4.72E-02 |
| GO:0008373 | sialyltransferase activity | MF | 16 | 3 | 3 | 4.72E-02 |
| GO:0032755 | positive regulation of interleukin-6 production | BP | 64 | 3 | 7 | 4.74E-02 |
| GO:0002673 | regulation of acute inflammatory response | BP | 64 | 4 | 7 | 4.74E-02 |
| GO:0071236 | cellular response to antibiotic | BP | 120 | 10 | 11 | 4.75E-02 |
| GO:0042542 | response to hydrogen peroxide | BP | 120 | 7 | 11 | 4.75E-02 |
| GO:0034627 | 'de novo' NAD biosynthetic process | BP | 7 | 1 | 2 | 4.76E-02 |
| GO:0034354 | 'de novo' NAD biosynthetic process from tryptophan | BP | 7 | 1 | 2 | 4.76E-02 |
| GO:0035005 | 1-phosphatidylinositol-4-phosphate 3-kinase activity | MF | 7 | 0 | 2 | 4.76E-02 |
| GO:0051666 | actin cortical patch localization | BP | 7 | 0 | 2 | 4.76E-02 |
| GO:0070358 | actin polymerization-dependent cell motility | BP | 7 | 1 | 2 | 4.76E-02 |
| GO:0071355 | cellular response to interleukin-9 | BP | 7 | 0 | 2 | 4.76E-02 |
| GO:0051852 | disruption by host of symbiont cells | BP | 7 | 0 | 2 | 4.76E-02 |
| GO:0051103 | DNA ligation involved in DNA repair | BP | 7 | 0 | 2 | 4.76E-02 |
| GO:0006265 | DNA topological change | BP | 7 | 0 | 2 | 4.76E-02 |
| GO:0021873 | forebrain neuroblast division | BP | 7 | 1 | 2 | 4.76E-02 |
| GO:0015669 | gas transport | BP | 7 | 1 | 2 | 4.76E-02 |
| GO:0090713 | immunological memory process | BP | 7 | 0 | 2 | 4.76E-02 |
| GO:0035771 | interleukin-4-mediated signaling pathway | BP | 7 | 0 | 2 | 4.76E-02 |
| GO:0038113 | interleukin-9-mediated signaling pathway | BP | 7 | 0 | 2 | 4.76E-02 |
| GO:0051873 | killing by host of symbiont cells | BP | 7 | 0 | 2 | 4.76E-02 |
| GO:0006273 | lagging strand elongation | BP | 7 | 0 | 2 | 4.76E-02 |
| GO:0035696 | monocyte extravasation | BP | 7 | 2 | 2 | 4.76E-02 |
| GO:0046642 | negative regulation of alpha-beta T cell proliferation | BP | 7 | 0 | 2 | 4.76E-02 |
| GO:2000483 | negative regulation of interleukin-8 secretion | BP | 7 | 1 | 2 | 4.76E-02 |
| GO:0010985 | negative regulation of lipoprotein particle clearance | BP | 7 | 0 | 2 | 4.76E-02 |
| GO:1905064 | negative regulation of vascular smooth muscle cell differentiation | BP | 7 | 1 | 2 | 4.76E-02 |
| GO:1901857 | positive regulation of cellular respiration | BP | 7 | 0 | 2 | 4.76E-02 |
| GO:0010873 | positive regulation of cholesterol esterification | BP | 7 | 2 | 2 | 4.76E-02 |
| GO:0060369 | positive regulation of Fc receptor mediated stimulatory signaling pa | BP | 7 | 0 | 2 | 4.76E-02 |
| GO:0045359 | positive regulation of interferon-beta biosynthetic process | BP | 7 | 0 | 2 | 4.76E-02 |
| GO:0045348 | positive regulation of MHC class II biosynthetic process | BP | 7 | 0 | 2 | 4.76E-02 |
| GO:0033034 | positive regulation of myeloid cell apoptotic process | BP | 7 | 0 | 2 | 4.76E-02 |
| GO:0031077 | post-embryonic camera-type eye development | BP | 7 | 0 | 2 | 4.76E-02 |
| GO:0060558 | regulation of calcidiol 1-monooxygenase activity | BP | 7 | 0 | 2 | 4.76E-02 |
| GO:1905449 | regulation of Fc-gamma receptor signaling pathway involved in pha | BP | 7 | 0 | 2 | 4.76E-02 |
| GO:0032383 | regulation of intracellular cholesterol transport | BP | 7 | 0 | 2 | 4.76E-02 |
| GO:0032377 | regulation of intracellular lipid transport | BP | 7 | 0 | 2 | 4.76E-02 |
| GO:0032380 | regulation of intracellular sterol transport | BP | 7 | 0 | 2 | 4.76E-02 |
| GO:0034756 | regulation of iron ion transport | BP | 7 | 0 | 2 | 4.76E-02 |
| GO:0090235 | regulation of metaphase plate congression | BP | 7 | 0 | 2 | 4.76E-02 |
| GO:0035507 | regulation of myosin-light-chain-phosphatase activity | BP | 7 | 0 | 2 | 4.76E-02 |
| GO:1903995 | regulation of non-membrane spanning protein tyrosine kinase activ | BP | 7 | 1 | 2 | 4.76E-02 |
| GO:0031298 | replication fork protection complex | CC | 7 | 0 | 2 | 4.76E-02 |
| GO:0050913 | sensory perception of bitter taste | BP | 7 | 0 | 2 | 4.76E-02 |
| GO:0007172 | signal complex assembly | BP | 7 | 1 | 2 | 4.76E-02 |
| GO:0021978 | telencephalon regionalization | BP | 7 | 0 | 2 | 4.76E-02 |
| GO:0046654 | tetrahydrofolate biosynthetic process | BP | 7 | 0 | 2 | 4.76E-02 |
| GO:0034447 | very-low-density lipoprotein particle clearance | BP | 7 | 0 | 2 | 4.76E-02 |
| GO:1904888 | cranial skeletal system development | BP | 51 | 3 | 6 | 4.76E-02 |
| GO:0051784 | negative regulation of nuclear division | BP | 51 | 0 | 6 | 4.76E-02 |
| GO:0016853 | isomerase activity | MF | 135 | 3 | 12 | 4.83E-02 |
| GO:0034101 | erythrocyte homeostasis | BP | 106 | 4 | 10 | 4.86E-02 |
| GO:0008528 | G protein-coupled peptide receptor activity | MF | 78 | 8 | 8 | 4.90E-02 |
| GO:0042136 | neurotransmitter biosynthetic process | BP | 78 | 11 | 8 | 4.90E-02 |
| GO:0050650 | chondroitin sulfate proteoglycan biosynthetic process | BP | 27 | 1 | 4 | 4.90E-02 |
| GO:0042417 | dopamine metabolic process | BP | 27 | 8 | 4 | 4.90E-02 |
| GO:1904030 | negative regulation of cyclin-dependent protein kinase activity | BP | 27 | 3 | 4 | 4.90E-02 |

| | | | | | | |
|---|---|---|---|---|---|---|
| GO:0043552 | positive regulation of phosphatidylinositol 3-kinase activity | BP | 27 | 2 | 4 | 4.90E-02 |
| GO:0032947 | protein-containing complex scaffold activity | MF | 27 | 2 | 4 | 4.90E-02 |
| GO:0032994 | protein-lipid complex | CC | 27 | 0 | 4 | 4.90E-02 |
| GO:0016298 | lipase activity | MF | 92 | 7 | 9 | 4.92E-02 |
| GO:0000784 | nuclear chromosome, telomeric region | CC | 92 | 0 | 9 | 4.92E-02 |
| GO:0033043 | regulation of organelle organization | BP | 1074 | 66 | 68 | 4.95E-02 |
| GO:0032732 | positive regulation of interleukin-1 production | BP | 39 | 0 | 5 | 4.99E-02 |

**Supp. Table 8b. GO sets upregulated in non-IBD**

| GO ID | Term | Ont | N | Up | Down | P-value |
|---|---|---|---|---|---|---|
| GO:0071944 | cell periphery | CC | 3744 | 430 | 294 | 3.22E-38 |
| GO:0005886 | plasma membrane | CC | 3656 | 422 | 286 | 9.07E-38 |
| GO:0003008 | system process | BP | 1229 | 204 | 64 | 3.89E-36 |
| GO:0044459 | plasma membrane part | CC | 2045 | 269 | 176 | 2.75E-30 |
| GO:0031226 | intrinsic component of plasma membrane | CC | 1165 | 179 | 113 | 3.89E-27 |
| GO:0045202 | synapse | CC | 977 | 156 | 37 | 2.62E-25 |
| GO:0005887 | integral component of plasma membrane | CC | 1107 | 166 | 105 | 7.04E-24 |
| GO:0099537 | trans-synaptic signaling | BP | 538 | 105 | 20 | 8.64E-24 |
| GO:0032501 | multicellular organismal process | BP | 5121 | 490 | 327 | 1.40E-23 |
| GO:0099536 | synaptic signaling | BP | 543 | 105 | 20 | 1.85E-23 |
| GO:0048731 | system development | BP | 3556 | 373 | 229 | 4.31E-23 |
| GO:0044057 | regulation of system process | BP | 444 | 92 | 17 | 8.25E-23 |
| GO:0031224 | intrinsic component of membrane | CC | 3479 | 366 | 227 | 9.22E-23 |
| GO:0098916 | anterograde trans-synaptic signaling | BP | 530 | 102 | 20 | 1.23E-22 |
| GO:0007268 | chemical synaptic transmission | BP | 530 | 102 | 20 | 1.23E-22 |
| GO:0006936 | muscle contraction | BP | 267 | 66 | 10 | 7.02E-21 |
| GO:0032502 | developmental process | BP | 4618 | 444 | 281 | 1.20E-20 |
| GO:0007399 | nervous system development | BP | 1847 | 226 | 95 | 1.33E-20 |
| GO:0048468 | cell development | BP | 1674 | 210 | 98 | 2.52E-20 |
| GO:0097458 | neuron part | CC | 1383 | 183 | 58 | 4.60E-20 |
| GO:0098590 | plasma membrane region | CC | 938 | 140 | 51 | 7.24E-20 |
| GO:0044456 | synapse part | CC | 776 | 123 | 31 | 1.26E-19 |
| GO:0048856 | anatomical structure development | BP | 4314 | 418 | 263 | 1.43E-19 |
| GO:0044425 | membrane part | CC | 4548 | 435 | 288 | 1.51E-19 |
| GO:0097060 | synaptic membrane | CC | 336 | 73 | 12 | 1.72E-19 |
| GO:0043292 | contractile fiber | CC | 179 | 51 | 6 | 2.92E-19 |
| GO:0003012 | muscle system process | BP | 340 | 73 | 11 | 3.50E-19 |
| GO:0007275 | multicellular organism development | BP | 3945 | 389 | 245 | 4.19E-19 |
| GO:0007610 | behavior | BP | 450 | 86 | 20 | 5.57E-19 |
| GO:0007267 | cell-cell signaling | BP | 1245 | 167 | 62 | 7.67E-19 |
| GO:0043269 | regulation of ion transport | BP | 507 | 92 | 42 | 1.11E-18 |
| GO:0050877 | nervous system process | BP | 664 | 109 | 36 | 1.39E-18 |
| GO:1902495 | transmembrane transporter complex | CC | 221 | 56 | 9 | 2.04E-18 |
| GO:0016021 | integral component of membrane | CC | 3371 | 343 | 216 | 2.08E-18 |
| GO:0051239 | regulation of multicellular organismal process | BP | 2424 | 267 | 187 | 4.04E-18 |
| GO:0030016 | myofibril | CC | 171 | 48 | 5 | 6.60E-18 |
| GO:1990351 | transporter complex | CC | 229 | 56 | 9 | 1.19E-17 |
| GO:0006811 | ion transport | BP | 1224 | 161 | 73 | 2.53E-17 |
| GO:0044449 | contractile fiber part | CC | 170 | 47 | 6 | 2.86E-17 |
| GO:0050804 | modulation of chemical synaptic transmission | BP | 359 | 72 | 14 | 3.25E-17 |
| GO:0099177 | regulation of trans-synaptic signaling | BP | 360 | 72 | 14 | 3.80E-17 |
| GO:0051049 | regulation of transport | BP | 1444 | 180 | 98 | 5.63E-17 |
| GO:0005261 | cation channel activity | MF | 215 | 53 | 14 | 6.23E-17 |
| GO:0022008 | neurogenesis | BP | 1291 | 166 | 74 | 6.62E-17 |
| GO:0042391 | regulation of membrane potential | BP | 317 | 66 | 12 | 1.01E-16 |
| GO:0034703 | cation channel complex | CC | 155 | 44 | 8 | 1.01E-16 |
| GO:0050808 | synapse organization | BP | 334 | 68 | 14 | 1.14E-16 |
| GO:0022839 | ion gated channel activity | MF | 211 | 52 | 13 | 1.25E-16 |
| GO:0034702 | ion channel complex | CC | 204 | 51 | 8 | 1.30E-16 |
| GO:0030017 | sarcomere | CC | 156 | 44 | 5 | 1.32E-16 |
| GO:0009653 | anatomical structure morphogenesis | BP | 2116 | 236 | 121 | 2.02E-16 |
| GO:0006812 | cation transport | BP | 831 | 121 | 55 | 2.41E-16 |
| GO:0022838 | substrate-specific channel activity | MF | 277 | 60 | 16 | 3.82E-16 |

| | | | | | | |
|---|---|---|---|---|---|---|
| GO:0031674 | I band | CC | 110 | 36 | 3 | 4.45E-16 |
| GO:1903522 | regulation of blood circulation | BP | 217 | 52 | 10 | 4.46E-16 |
| GO:0048699 | generation of neurons | BP | 1209 | 156 | 70 | 5.20E-16 |
| GO:0030018 | Z disc | CC | 105 | 35 | 3 | 5.99E-16 |
| GO:0022836 | gated channel activity | MF | 219 | 52 | 14 | 6.74E-16 |
| GO:0007416 | synapse assembly | BP | 138 | 40 | 6 | 1.17E-15 |
| GO:0032879 | regulation of localization | BP | 2184 | 239 | 146 | 1.20E-15 |
| GO:0015267 | channel activity | MF | 292 | 61 | 18 | 1.30E-15 |
| GO:0061061 | muscle structure development | BP | 509 | 86 | 22 | 1.44E-15 |
| GO:0022803 | passive transmembrane transporter activity | MF | 293 | 61 | 18 | 1.54E-15 |
| GO:0001505 | regulation of neurotransmitter levels | BP | 289 | 60 | 16 | 3.03E-15 |
| GO:0003015 | heart process | BP | 212 | 50 | 9 | 3.30E-15 |
| GO:0005216 | ion channel activity | MF | 274 | 58 | 16 | 3.60E-15 |
| GO:0032989 | cellular component morphogenesis | BP | 955 | 130 | 60 | 3.70E-15 |
| GO:0043005 | neuron projection | CC | 1036 | 137 | 49 | 6.31E-15 |
| GO:0048666 | neuron development | BP | 909 | 125 | 50 | 6.34E-15 |
| GO:0098797 | plasma membrane protein complex | CC | 413 | 74 | 33 | 6.75E-15 |
| GO:0006836 | neurotransmitter transport | BP | 217 | 50 | 9 | 8.93E-15 |
| GO:0051960 | regulation of nervous system development | BP | 740 | 108 | 42 | 1.03E-14 |
| GO:0030182 | neuron differentiation | BP | 1086 | 141 | 60 | 1.07E-14 |
| GO:0030154 | cell differentiation | BP | 3014 | 301 | 212 | 1.57E-14 |
| GO:0048869 | cellular developmental process | BP | 3171 | 313 | 224 | 1.67E-14 |
| GO:0006813 | potassium ion transport | BP | 162 | 42 | 6 | 1.70E-14 |
| GO:0098978 | glutamatergic synapse | CC | 292 | 59 | 16 | 1.86E-14 |
| GO:0045211 | postsynaptic membrane | CC | 244 | 53 | 10 | 1.88E-14 |
| GO:0003013 | circulatory system process | BP | 406 | 72 | 21 | 2.76E-14 |
| GO:0098793 | presynapse | CC | 406 | 72 | 16 | 2.76E-14 |
| GO:0046873 | metal ion transmembrane transporter activity | MF | 321 | 62 | 16 | 3.49E-14 |
| GO:0034762 | regulation of transmembrane transport | BP | 417 | 73 | 31 | 3.59E-14 |
| GO:0060047 | heart contraction | BP | 202 | 47 | 8 | 3.91E-14 |
| GO:0098794 | postsynapse | CC | 510 | 83 | 23 | 4.12E-14 |
| GO:0031175 | neuron projection development | BP | 815 | 113 | 44 | 8.65E-14 |
| GO:0065008 | regulation of biological quality | BP | 3132 | 307 | 201 | 9.39E-14 |
| GO:0071804 | cellular potassium ion transport | BP | 143 | 38 | 6 | 1.31E-13 |
| GO:0071805 | potassium ion transmembrane transport | BP | 143 | 38 | 6 | 1.31E-13 |
| GO:0023061 | signal release | BP | 365 | 66 | 15 | 1.35E-13 |
| GO:0015318 | inorganic molecular entity transmembrane transporter activity | MF | 550 | 86 | 27 | 1.44E-13 |
| GO:0008016 | regulation of heart contraction | BP | 180 | 43 | 8 | 1.83E-13 |
| GO:0120025 | plasma membrane bounded cell projection | CC | 1631 | 185 | 84 | 2.28E-13 |
| GO:0008015 | blood circulation | BP | 396 | 69 | 20 | 2.37E-13 |
| GO:0001508 | action potential | BP | 100 | 31 | 1 | 2.62E-13 |
| GO:0034765 | regulation of ion transmembrane transport | BP | 353 | 64 | 29 | 2.83E-13 |
| GO:0042995 | cell projection | CC | 1695 | 190 | 89 | 3.02E-13 |
| GO:0030001 | metal ion transport | BP | 666 | 97 | 45 | 3.10E-13 |
| GO:0022890 | inorganic cation transmembrane transporter activity | MF | 400 | 69 | 20 | 3.90E-13 |
| GO:0022857 | transmembrane transporter activity | MF | 720 | 102 | 31 | 4.19E-13 |
| GO:0034220 | ion transmembrane transport | BP | 848 | 114 | 52 | 5.54E-13 |
| GO:0008324 | cation transmembrane transporter activity | MF | 432 | 72 | 22 | 6.78E-13 |
| GO:0055085 | transmembrane transport | BP | 1167 | 143 | 60 | 7.14E-13 |
| GO:0007269 | neurotransmitter secretion | BP | 145 | 37 | 6 | 1.06E-12 |
| GO:0099643 | signal release from synapse | BP | 146 | 37 | 6 | 1.33E-12 |
| GO:0015672 | monovalent inorganic cation transport | BP | 356 | 63 | 18 | 1.34E-12 |
| GO:0007155 | cell adhesion | BP | 1113 | 137 | 113 | 1.70E-12 |
| GO:0022610 | biological adhesion | BP | 1116 | 137 | 113 | 2.09E-12 |
| GO:0099240 | intrinsic component of synaptic membrane | CC | 114 | 32 | 7 | 2.25E-12 |
| GO:0030054 | cell junction | CC | 1101 | 135 | 64 | 3.40E-12 |
| GO:2000026 | regulation of multicellular organismal development | BP | 1597 | 178 | 118 | 3.55E-12 |
| GO:0120039 | plasma membrane bounded cell projection morphogenesis | BP | 563 | 84 | 35 | 3.78E-12 |
| GO:0051965 | positive regulation of synapse assembly | BP | 57 | 22 | 4 | 5.47E-12 |
| GO:0048858 | cell projection morphogenesis | BP | 567 | 84 | 35 | 5.54E-12 |
| GO:0015075 | ion transmembrane transporter activity | MF | 598 | 87 | 28 | 6.14E-12 |
| GO:0048812 | neuron projection morphogenesis | BP | 549 | 82 | 34 | 6.57E-12 |
| GO:0005215 | transporter activity | MF | 795 | 106 | 35 | 6.72E-12 |
| GO:0051962 | positive regulation of nervous system development | BP | 429 | 69 | 24 | 1.15E-11 |

| | | | | | | |
|---|---|---|---|---|---|---|
| GO:0043266 | regulation of potassium ion transport | BP | 70 | 24 | 3 | 1.17E-11 |
| GO:0035637 | multicellular organismal signaling | BP | 149 | 36 | 2 | 1.22E-11 |
| GO:0000902 | cell morphogenesis | BP | 869 | 112 | 58 | 1.42E-11 |
| GO:0042734 | presynaptic membrane | CC | 129 | 33 | 4 | 1.67E-11 |
| GO:0022832 | voltage-gated channel activity | MF | 129 | 33 | 7 | 1.67E-11 |
| GO:0005244 | voltage-gated ion channel activity | MF | 129 | 33 | 7 | 1.67E-11 |
| GO:0050807 | regulation of synapse organization | BP | 181 | 40 | 11 | 1.80E-11 |
| GO:0000904 | cell morphogenesis involved in differentiation | BP | 632 | 89 | 40 | 2.10E-11 |
| GO:0099699 | integral component of synaptic membrane | CC | 103 | 29 | 6 | 2.23E-11 |
| GO:0050793 | regulation of developmental process | BP | 2011 | 209 | 146 | 2.52E-11 |
| GO:0032990 | cell part morphogenesis | BP | 585 | 84 | 35 | 2.92E-11 |
| GO:0060284 | regulation of cell development | BP | 743 | 99 | 42 | 3.75E-11 |
| GO:0006937 | regulation of muscle contraction | BP | 126 | 32 | 6 | 4.17E-11 |
| GO:0030424 | axon | CC | 510 | 76 | 24 | 4.57E-11 |
| GO:0050803 | regulation of synapse structure or activity | BP | 188 | 40 | 11 | 6.26E-11 |
| GO:0007611 | learning or memory | BP | 204 | 42 | 12 | 6.29E-11 |
| GO:0048667 | cell morphogenesis involved in neuron differentiation | BP | 494 | 74 | 30 | 6.52E-11 |
| GO:0061564 | axon development | BP | 427 | 67 | 29 | 7.09E-11 |
| GO:0051963 | regulation of synapse assembly | BP | 88 | 26 | 5 | 7.43E-11 |
| GO:0050767 | regulation of neurogenesis | BP | 651 | 89 | 37 | 1.04E-10 |
| GO:0036477 | somatodendritic compartment | CC | 672 | 91 | 29 | 1.05E-10 |
| GO:0098655 | cation transmembrane transport | BP | 605 | 84 | 43 | 1.66E-10 |
| GO:0098742 | cell-cell adhesion via plasma-membrane adhesion molecules | BP | 188 | 39 | 11 | 2.41E-10 |
| GO:0007517 | muscle organ development | BP | 299 | 52 | 13 | 2.50E-10 |
| GO:0098660 | inorganic ion transmembrane transport | BP | 580 | 81 | 39 | 2.71E-10 |
| GO:0098984 | neuron to neuron synapse | CC | 309 | 53 | 13 | 2.83E-10 |
| GO:0090257 | regulation of muscle system process | BP | 189 | 39 | 7 | 2.84E-10 |
| GO:0043025 | neuronal cell body | CC | 403 | 63 | 18 | 3.14E-10 |
| GO:0050890 | cognition | BP | 231 | 44 | 13 | 3.16E-10 |
| GO:0048513 | animal organ development | BP | 2568 | 248 | 184 | 3.96E-10 |
| GO:0042692 | muscle cell differentiation | BP | 286 | 50 | 16 | 4.61E-10 |
| GO:0099504 | synaptic vesicle cycle | BP | 168 | 36 | 5 | 4.62E-10 |
| GO:0098936 | intrinsic component of postsynaptic membrane | CC | 82 | 24 | 6 | 4.96E-10 |
| GO:0051094 | positive regulation of developmental process | BP | 1057 | 124 | 88 | 5.44E-10 |
| GO:0008307 | structural constituent of muscle | MF | 37 | 16 | 1 | 5.89E-10 |
| GO:0120036 | plasma membrane bounded cell projection organization | BP | 1232 | 139 | 61 | 6.16E-10 |
| GO:0099055 | integral component of postsynaptic membrane | CC | 77 | 23 | 6 | 7.28E-10 |
| GO:1901379 | regulation of potassium ion transmembrane transport | BP | 59 | 20 | 3 | 7.70E-10 |
| GO:0010959 | regulation of metal ion transport | BP | 302 | 51 | 30 | 1.08E-09 |
| GO:0007423 | sensory organ development | BP | 406 | 62 | 16 | 1.14E-09 |
| GO:0031012 | extracellular matrix | CC | 397 | 61 | 17 | 1.22E-09 |
| GO:0042383 | sarcolemma | CC | 113 | 28 | 2 | 1.27E-09 |
| GO:0044297 | cell body | CC | 457 | 67 | 23 | 1.37E-09 |
| GO:0099081 | supramolecular polymer | CC | 674 | 88 | 49 | 1.42E-09 |
| GO:0099080 | supramolecular complex | CC | 675 | 88 | 49 | 1.53E-09 |
| GO:0007409 | axonogenesis | BP | 390 | 60 | 28 | 1.59E-09 |
| GO:0016020 | membrane | CC | 6576 | 532 | 402 | 1.65E-09 |
| GO:0097479 | synaptic vesicle localization | BP | 145 | 32 | 6 | 1.98E-09 |
| GO:0045595 | regulation of cell differentiation | BP | 1409 | 152 | 103 | 2.11E-09 |
| GO:0098662 | inorganic cation transmembrane transport | BP | 533 | 74 | 37 | 2.19E-09 |
| GO:0038023 | signaling receptor activity | MF | 703 | 90 | 82 | 2.48E-09 |
| GO:0015079 | potassium ion transmembrane transporter activity | MF | 102 | 26 | 5 | 2.57E-09 |
| GO:0045664 | regulation of neuron differentiation | BP | 525 | 73 | 26 | 2.65E-09 |
| GO:0051952 | regulation of amine transport | BP | 63 | 20 | 3 | 2.87E-09 |
| GO:0051146 | striated muscle cell differentiation | BP | 221 | 41 | 12 | 2.89E-09 |
| GO:0030030 | cell projection organization | BP | 1262 | 139 | 63 | 3.10E-09 |
| GO:0008092 | cytoskeletal protein binding | MF | 828 | 101 | 59 | 3.43E-09 |
| GO:0007612 | learning | BP | 118 | 28 | 6 | 3.63E-09 |
| GO:0007154 | cell communication | BP | 4656 | 397 | 345 | 4.44E-09 |
| GO:0023052 | signaling | BP | 4643 | 396 | 346 | 4.58E-09 |
| GO:0099068 | postsynapse assembly | BP | 27 | 13 | 3 | 4.88E-09 |
| GO:0072359 | circulatory system development | BP | 870 | 104 | 37 | 5.90E-09 |
| GO:0043235 | receptor complex | CC | 311 | 50 | 33 | 8.67E-09 |
| GO:0099512 | supramolecular fiber | CC | 667 | 85 | 49 | 8.87E-09 |

| | | | | | | |
|---|---|---|---|---|---|---|
| GO:0015837 | amine transport | BP | 67 | 20 | 3 | 9.49E-09 |
| GO:0015077 | monovalent inorganic cation transmembrane transporter activity | MF | 239 | 42 | 10 | 9.90E-09 |
| GO:0060537 | muscle tissue development | BP | 303 | 49 | 11 | 1.01E-08 |
| GO:0003779 | actin binding | MF | 370 | 56 | 28 | 1.03E-08 |
| GO:0005267 | potassium channel activity | MF | 74 | 21 | 4 | 1.11E-08 |
| GO:0060089 | molecular transducer activity | MF | 747 | 92 | 83 | 1.14E-08 |
| GO:0030029 | actin filament-based process | BP | 651 | 83 | 42 | 1.32E-08 |
| GO:0005198 | structural molecule activity | MF | 536 | 72 | 18 | 1.50E-08 |
| GO:0007600 | sensory perception | BP | 326 | 51 | 21 | 1.57E-08 |
| GO:0062023 | collagen-containing extracellular matrix | CC | 317 | 50 | 12 | 1.65E-08 |
| GO:0032279 | asymmetric synapse | CC | 289 | 47 | 13 | 1.69E-08 |
| GO:0051588 | regulation of neurotransmitter transport | BP | 111 | 26 | 4 | 1.75E-08 |
| GO:0007507 | heart development | BP | 458 | 64 | 12 | 2.18E-08 |
| GO:0086010 | membrane depolarization during action potential | BP | 30 | 13 | 0 | 2.40E-08 |
| GO:0002027 | regulation of heart rate | BP | 77 | 21 | 2 | 2.41E-08 |
| GO:0022843 | voltage-gated cation channel activity | MF | 91 | 23 | 6 | 2.53E-08 |
| GO:0099572 | postsynaptic specialization | CC | 294 | 47 | 13 | 2.92E-08 |
| GO:0098693 | regulation of synaptic vesicle cycle | BP | 99 | 24 | 3 | 3.03E-08 |
| GO:0055001 | muscle cell development | BP | 137 | 29 | 5 | 3.04E-08 |
| GO:0014069 | postsynaptic density | CC | 285 | 46 | 13 | 3.04E-08 |
| GO:0070252 | actin-mediated cell contraction | BP | 92 | 23 | 5 | 3.16E-08 |
| GO:0051899 | membrane depolarization | BP | 72 | 20 | 4 | 3.69E-08 |
| GO:0014706 | striated muscle tissue development | BP | 287 | 46 | 11 | 3.79E-08 |
| GO:0007158 | neuron cell-cell adhesion | BP | 14 | 9 | 0 | 4.31E-08 |
| GO:0007588 | excretion | BP | 42 | 15 | 4 | 4.62E-08 |
| GO:0048589 | developmental growth | BP | 509 | 68 | 25 | 4.77E-08 |
| GO:0150052 | regulation of postsynapse assembly | BP | 11 | 8 | 0 | 5.99E-08 |
| GO:0099150 | regulation of postsynaptic specialization assembly | BP | 11 | 8 | 0 | 5.99E-08 |
| GO:0016079 | synaptic vesicle exocytosis | BP | 103 | 24 | 4 | 6.88E-08 |
| GO:0015850 | organic hydroxy compound transport | BP | 185 | 34 | 8 | 8.26E-08 |
| GO:0099560 | synaptic membrane adhesion | BP | 23 | 11 | 2 | 8.39E-08 |
| GO:0043271 | negative regulation of ion transport | BP | 112 | 25 | 5 | 9.08E-08 |
| GO:0099173 | postsynapse organization | BP | 144 | 29 | 8 | 9.62E-08 |
| GO:0051240 | positive regulation of multicellular organismal process | BP | 1345 | 140 | 123 | 9.83E-08 |
| GO:0042493 | response to drug | BP | 782 | 92 | 56 | 1.03E-07 |
| GO:0098698 | postsynaptic specialization assembly | BP | 19 | 10 | 1 | 1.04E-07 |
| GO:0019932 | second-messenger-mediated signaling | BP | 316 | 48 | 23 | 1.06E-07 |
| GO:0098889 | intrinsic component of presynaptic membrane | CC | 63 | 18 | 4 | 1.10E-07 |
| GO:0030425 | dendrite | CC | 499 | 66 | 22 | 1.11E-07 |
| GO:0097447 | dendritic tree | CC | 499 | 66 | 22 | 1.11E-07 |
| GO:0005201 | extracellular matrix structural constituent | MF | 137 | 28 | 4 | 1.15E-07 |
| GO:0006941 | striated muscle contraction | BP | 129 | 27 | 6 | 1.16E-07 |
| GO:0004888 | transmembrane signaling receptor activity | MF | 542 | 70 | 66 | 1.16E-07 |
| GO:1904062 | regulation of cation transmembrane transport | BP | 260 | 42 | 23 | 1.19E-07 |
| GO:0009410 | response to xenobiotic stimulus | BP | 206 | 36 | 9 | 1.30E-07 |
| GO:0099060 | integral component of postsynaptic specialization membrane | CC | 45 | 15 | 2 | 1.33E-07 |
| GO:0017156 | calcium ion regulated exocytosis | BP | 130 | 27 | 4 | 1.37E-07 |
| GO:0034705 | potassium channel complex | CC | 64 | 18 | 4 | 1.44E-07 |
| GO:0022898 | regulation of transmembrane transporter activity | BP | 198 | 35 | 14 | 1.45E-07 |
| GO:0099634 | postsynaptic specialization membrane | CC | 65 | 18 | 2 | 1.87E-07 |
| GO:0051371 | muscle alpha-actinin binding | MF | 20 | 10 | 0 | 1.95E-07 |
| GO:0099054 | presynapse assembly | BP | 35 | 13 | 2 | 2.15E-07 |
| GO:0032412 | regulation of ion transmembrane transporter activity | BP | 193 | 34 | 14 | 2.38E-07 |
| GO:0055002 | striated muscle cell development | BP | 126 | 26 | 5 | 2.66E-07 |
| GO:0005509 | calcium ion binding | MF | 490 | 64 | 32 | 2.76E-07 |
| GO:0009914 | hormone transport | BP | 240 | 39 | 10 | 2.86E-07 |
| GO:0097480 | establishment of synaptic vesicle localization | BP | 135 | 27 | 5 | 3.07E-07 |
| GO:0048489 | synaptic vesicle transport | BP | 135 | 27 | 5 | 3.07E-07 |
| GO:0031225 | anchored component of membrane | CC | 111 | 24 | 11 | 3.08E-07 |
| GO:0010975 | regulation of neuron projection development | BP | 419 | 57 | 21 | 3.28E-07 |
| GO:0009719 | response to endogenous stimulus | BP | 1287 | 133 | 76 | 3.30E-07 |
| GO:0098948 | intrinsic component of postsynaptic specialization membrane | CC | 48 | 15 | 2 | 3.48E-07 |
| GO:0006940 | regulation of smooth muscle contraction | BP | 48 | 15 | 2 | 3.48E-07 |
| GO:0019229 | regulation of vasoconstriction | BP | 48 | 15 | 2 | 3.48E-07 |

| | | | | | | |
|---|---|---|---|---|---|---|
| GO:0061337 | cardiac conduction | BP | 112 | 24 | 2 | 3.68E-07 |
| GO:0051393 | alpha-actinin binding | MF | 31 | 12 | 1 | 3.75E-07 |
| GO:0099084 | postsynaptic specialization organization | BP | 31 | 12 | 1 | 3.75E-07 |
| GO:0010817 | regulation of hormone levels | BP | 380 | 53 | 17 | 3.88E-07 |
| GO:1901381 | positive regulation of potassium ion transmembrane transport | BP | 26 | 11 | 2 | 3.97E-07 |
| GO:1905874 | regulation of postsynaptic density organization | BP | 13 | 8 | 0 | 4.13E-07 |
| GO:0015081 | sodium ion transmembrane transporter activity | MF | 113 | 24 | 4 | 4.37E-07 |
| GO:0044463 | cell projection part | CC | 1141 | 120 | 53 | 5.49E-07 |
| GO:0120038 | plasma membrane bounded cell projection part | CC | 1141 | 120 | 53 | 5.49E-07 |
| GO:0032409 | regulation of transporter activity | BP | 209 | 35 | 15 | 5.60E-07 |
| GO:0030239 | myofibril assembly | BP | 56 | 16 | 1 | 5.60E-07 |
| GO:0043268 | positive regulation of potassium ion transport | BP | 32 | 12 | 2 | 5.63E-07 |
| GO:0048167 | regulation of synaptic plasticity | BP | 156 | 29 | 5 | 5.73E-07 |
| GO:0006939 | smooth muscle contraction | BP | 84 | 20 | 2 | 5.83E-07 |
| GO:0099151 | regulation of postsynaptic density assembly | BP | 10 | 7 | 0 | 6.48E-07 |
| GO:0050769 | positive regulation of neurogenesis | BP | 376 | 52 | 20 | 6.52E-07 |
| GO:0099172 | presynapse organization | BP | 38 | 13 | 2 | 6.53E-07 |
| GO:1903530 | regulation of secretion by cell | BP | 567 | 70 | 45 | 6.55E-07 |
| GO:0098900 | regulation of action potential | BP | 44 | 14 | 1 | 6.69E-07 |
| GO:0006814 | sodium ion transport | BP | 175 | 31 | 8 | 7.10E-07 |
| GO:0099175 | regulation of postsynapse organization | BP | 85 | 20 | 5 | 7.14E-07 |
| GO:0071495 | cellular response to endogenous stimulus | BP | 1088 | 115 | 63 | 7.60E-07 |
| GO:2001023 | regulation of response to drug | BP | 64 | 17 | 2 | 7.81E-07 |
| GO:0042221 | response to chemical | BP | 3288 | 285 | 230 | 8.13E-07 |
| GO:0007411 | axon guidance | BP | 231 | 37 | 17 | 8.29E-07 |
| GO:0097485 | neuron projection guidance | BP | 231 | 37 | 17 | 8.29E-07 |
| GO:0098982 | GABA-ergic synapse | CC | 51 | 15 | 0 | 8.37E-07 |
| GO:0005604 | basement membrane | CC | 86 | 20 | 2 | 8.72E-07 |
| GO:0042805 | actinin binding | MF | 39 | 13 | 2 | 9.19E-07 |
| GO:0050806 | positive regulation of synaptic transmission | BP | 134 | 26 | 5 | 9.35E-07 |
| GO:0086001 | cardiac muscle cell action potential | BP | 58 | 16 | 1 | 9.45E-07 |
| GO:0097517 | contractile actin filament bundle | CC | 58 | 16 | 3 | 9.45E-07 |
| GO:0001725 | stress fiber | CC | 58 | 16 | 3 | 9.45E-07 |
| GO:0005262 | calcium channel activity | MF | 94 | 21 | 6 | 9.50E-07 |
| GO:0032432 | actin filament bundle | CC | 65 | 17 | 4 | 9.92E-07 |
| GO:0051480 | regulation of cytosolic calcium ion concentration | BP | 271 | 41 | 29 | 1.00E-06 |
| GO:0005516 | calmodulin binding | MF | 169 | 30 | 9 | 1.02E-06 |
| GO:0010720 | positive regulation of cell development | BP | 434 | 57 | 24 | 1.05E-06 |
| GO:0007156 | homophilic cell adhesion via plasma membrane adhesion molecules | BP | 103 | 22 | 4 | 1.19E-06 |
| GO:0007417 | central nervous system development | BP | 757 | 86 | 41 | 1.23E-06 |
| GO:0051046 | regulation of secretion | BP | 610 | 73 | 47 | 1.23E-06 |
| GO:0046928 | regulation of neurotransmitter secretion | BP | 88 | 20 | 3 | 1.29E-06 |
| GO:0045666 | positive regulation of neuron differentiation | BP | 294 | 43 | 11 | 1.38E-06 |
| GO:0060048 | cardiac muscle contraction | BP | 104 | 22 | 4 | 1.41E-06 |
| GO:0035725 | sodium ion transmembrane transport | BP | 112 | 23 | 5 | 1.41E-06 |
| GO:0097106 | postsynaptic density organization | BP | 29 | 11 | 0 | 1.47E-06 |
| GO:0045177 | apical part of cell | CC | 306 | 44 | 20 | 1.67E-06 |
| GO:1904889 | regulation of excitatory synapse assembly | BP | 11 | 7 | 1 | 1.68E-06 |
| GO:0086002 | cardiac muscle cell action potential involved in contraction | BP | 41 | 13 | 1 | 1.75E-06 |
| GO:0048878 | chemical homeostasis | BP | 904 | 98 | 63 | 1.81E-06 |
| GO:0016324 | apical plasma membrane | CC | 248 | 38 | 18 | 1.81E-06 |
| GO:0046883 | regulation of hormone secretion | BP | 201 | 33 | 7 | 1.85E-06 |
| GO:0086003 | cardiac muscle cell contraction | BP | 54 | 15 | 3 | 1.87E-06 |
| GO:0030594 | neurotransmitter receptor activity | MF | 54 | 15 | 2 | 1.87E-06 |
| GO:0030048 | actin filament-based movement | BP | 114 | 23 | 7 | 1.95E-06 |
| GO:0030534 | adult behavior | BP | 114 | 23 | 4 | 1.95E-06 |
| GO:0010927 | cellular component assembly involved in morphogenesis | BP | 91 | 20 | 2 | 2.25E-06 |
| GO:0007215 | glutamate receptor signaling pathway | BP | 76 | 18 | 4 | 2.30E-06 |
| GO:0099056 | integral component of presynaptic membrane | CC | 55 | 15 | 3 | 2.42E-06 |
| GO:0098839 | postsynaptic density membrane | CC | 55 | 15 | 2 | 2.42E-06 |
| GO:0001964 | startle response | BP | 20 | 9 | 1 | 2.49E-06 |
| GO:0090066 | regulation of anatomical structure size | BP | 414 | 54 | 29 | 2.50E-06 |
| GO:0043267 | negative regulation of potassium ion transport | BP | 25 | 10 | 0 | 2.52E-06 |
| GO:0017158 | regulation of calcium ion-dependent exocytosis | BP | 92 | 20 | 3 | 2.70E-06 |

| | | | | | | |
|---|---|---|---|---|---|---|
| GO:0097746 | regulation of blood vessel diameter | BP | 108 | 22 | 7 | 2.74E-06 |
| GO:0050880 | regulation of blood vessel size | BP | 108 | 22 | 7 | 2.74E-06 |
| GO:0035296 | regulation of tube diameter | BP | 108 | 22 | 7 | 2.74E-06 |
| GO:0035150 | regulation of tube size | BP | 108 | 22 | 7 | 2.74E-06 |
| GO:0048738 | cardiac muscle tissue development | BP | 168 | 29 | 5 | 2.77E-06 |
| GO:0060341 | regulation of cellular localization | BP | 784 | 87 | 44 | 2.81E-06 |
| GO:0045597 | positive regulation of cell differentiation | BP | 762 | 85 | 63 | 2.99E-06 |
| GO:0035249 | synaptic transmission, glutamatergic | BP | 70 | 17 | 3 | 3.03E-06 |
| GO:0008076 | voltage-gated potassium channel complex | CC | 56 | 15 | 4 | 3.10E-06 |
| GO:0048880 | sensory system development | BP | 283 | 41 | 11 | 3.10E-06 |
| GO:0060078 | regulation of postsynaptic membrane potential | BP | 101 | 21 | 5 | 3.26E-06 |
| GO:0022604 | regulation of cell morphogenesis | BP | 418 | 54 | 29 | 3.35E-06 |
| GO:0099061 | integral component of postsynaptic density membrane | CC | 37 | 12 | 2 | 3.36E-06 |
| GO:1901016 | regulation of potassium ion transmembrane transporter activity | BP | 37 | 12 | 2 | 3.36E-06 |
| GO:0046879 | hormone secretion | BP | 235 | 36 | 10 | 3.39E-06 |
| GO:0097107 | postsynaptic density assembly | BP | 16 | 8 | 0 | 3.44E-06 |
| GO:0055074 | calcium ion homeostasis | BP | 356 | 48 | 37 | 3.62E-06 |
| GO:0051966 | regulation of synaptic transmission, glutamatergic | BP | 50 | 14 | 2 | 3.72E-06 |
| GO:0015085 | calcium ion transmembrane transporter activity | MF | 110 | 22 | 7 | 3.77E-06 |
| GO:0007613 | memory | BP | 94 | 20 | 5 | 3.84E-06 |
| GO:0120035 | regulation of plasma membrane bounded cell projection organizatic | BP | 562 | 67 | 29 | 3.91E-06 |
| GO:0072507 | divalent inorganic cation homeostasis | BP | 378 | 50 | 38 | 3.95E-06 |
| GO:0006928 | movement of cell or subcellular component | BP | 1611 | 154 | 129 | 3.96E-06 |
| GO:0035235 | ionotropic glutamate receptor signaling pathway | BP | 21 | 9 | 0 | 4.09E-06 |
| GO:0043010 | camera-type eye development | BP | 237 | 36 | 9 | 4.14E-06 |
| GO:0023051 | regulation of signaling | BP | 2819 | 246 | 206 | 4.18E-06 |
| GO:0055078 | sodium ion homeostasis | BP | 32 | 11 | 4 | 4.54E-06 |
| GO:0022835 | transmitter-gated channel activity | MF | 32 | 11 | 0 | 4.54E-06 |
| GO:0022824 | transmitter-gated ion channel activity | MF | 32 | 11 | 0 | 4.54E-06 |
| GO:0015629 | actin cytoskeleton | CC | 434 | 55 | 33 | 4.88E-06 |
| GO:0001540 | amyloid-beta binding | MF | 65 | 16 | 7 | 4.89E-06 |
| GO:0043270 | positive regulation of ion transport | BP | 211 | 33 | 23 | 5.48E-06 |
| GO:0019226 | transmission of nerve impulse | BP | 45 | 13 | 0 | 5.64E-06 |
| GO:2001257 | regulation of cation channel activity | BP | 121 | 23 | 8 | 5.64E-06 |
| GO:1904315 | transmitter-gated ion channel activity involved in regulation of post | MF | 27 | 10 | 0 | 5.74E-06 |
| GO:0034764 | positive regulation of transmembrane transport | BP | 156 | 27 | 14 | 5.76E-06 |
| GO:0007186 | G protein-coupled receptor signaling pathway | BP | 591 | 69 | 51 | 5.79E-06 |
| GO:0031344 | regulation of cell projection organization | BP | 569 | 67 | 30 | 5.95E-06 |
| GO:0040007 | growth | BP | 764 | 84 | 35 | 6.05E-06 |
| GO:0008066 | glutamate receptor activity | MF | 17 | 8 | 0 | 6.10E-06 |
| GO:0045214 | sarcomere organization | BP | 39 | 12 | 1 | 6.26E-06 |
| GO:1904861 | excitatory synapse assembly | BP | 22 | 9 | 1 | 6.51E-06 |
| GO:0019228 | neuronal action potential | BP | 22 | 9 | 0 | 6.51E-06 |
| GO:0005576 | extracellular region | CC | 2796 | 243 | 202 | 6.74E-06 |
| GO:0043204 | perikaryon | CC | 106 | 21 | 3 | 7.29E-06 |
| GO:0009888 | tissue development | BP | 1451 | 140 | 89 | 7.32E-06 |
| GO:0044241 | lipid digestion | BP | 13 | 7 | 0 | 7.73E-06 |
| GO:0042420 | dopamine catabolic process | BP | 6 | 5 | 0 | 8.06E-06 |
| GO:0032222 | regulation of synaptic transmission, cholinergic | BP | 6 | 5 | 1 | 8.06E-06 |
| GO:0010769 | regulation of cell morphogenesis involved in differentiation | BP | 264 | 38 | 16 | 8.25E-06 |
| GO:0005865 | striated muscle thin filament | CC | 28 | 10 | 1 | 8.38E-06 |
| GO:0046658 | anchored component of plasma membrane | CC | 40 | 12 | 5 | 8.39E-06 |
| GO:0099146 | intrinsic component of postsynaptic density membrane | CC | 40 | 12 | 2 | 8.39E-06 |
| GO:0055065 | metal ion homeostasis | BP | 486 | 59 | 46 | 8.56E-06 |
| GO:0010646 | regulation of cell communication | BP | 2778 | 241 | 205 | 8.59E-06 |
| GO:0022834 | ligand-gated channel activity | MF | 83 | 18 | 5 | 8.75E-06 |
| GO:0015276 | ligand-gated ion channel activity | MF | 83 | 18 | 5 | 8.75E-06 |
| GO:0099003 | vesicle-mediated transport in synapse | BP | 178 | 29 | 5 | 8.97E-06 |
| GO:0001654 | eye development | BP | 275 | 39 | 9 | 8.99E-06 |
| GO:0006874 | cellular calcium ion homeostasis | BP | 347 | 46 | 36 | 9.06E-06 |
| GO:0050896 | response to stimulus | BP | 6405 | 497 | 466 | 9.17E-06 |
| GO:0022603 | regulation of anatomical structure morphogenesis | BP | 879 | 93 | 60 | 9.26E-06 |
| GO:0042310 | vasoconstriction | BP | 61 | 15 | 2 | 9.76E-06 |
| GO:1905606 | regulation of presynapse assembly | BP | 23 | 9 | 1 | 1.01E-05 |

| | | | | | | |
|---|---|---|---|---|---|---|
| GO:0034706 | sodium channel complex | CC | 18 | 8 | 1 | 1.03E-05 |
| GO:0042641 | actomyosin | CC | 69 | 16 | 3 | 1.12E-05 |
| GO:0098609 | cell-cell adhesion | BP | 638 | 72 | 84 | 1.19E-05 |
| GO:0015893 | drug transport | BP | 144 | 25 | 5 | 1.20E-05 |
| GO:0036379 | myofilament | CC | 29 | 10 | 1 | 1.20E-05 |
| GO:0086009 | membrane repolarization | BP | 35 | 11 | 2 | 1.22E-05 |
| GO:0150063 | visual system development | BP | 279 | 39 | 11 | 1.27E-05 |
| GO:0031032 | actomyosin structure organization | BP | 172 | 28 | 5 | 1.29E-05 |
| GO:0007187 | G protein-coupled receptor signaling pathway, coupled to cyclic nu | BP | 154 | 26 | 11 | 1.35E-05 |
| GO:0055006 | cardiac cell development | BP | 70 | 16 | 1 | 1.36E-05 |
| GO:1902803 | regulation of synaptic vesicle transport | BP | 70 | 16 | 3 | 1.36E-05 |
| GO:0071466 | cellular response to xenobiotic stimulus | BP | 119 | 22 | 4 | 1.42E-05 |
| GO:1901018 | positive regulation of potassium ion transmembrane transporter act | BP | 14 | 7 | 2 | 1.46E-05 |
| GO:0003014 | renal system process | BP | 94 | 19 | 6 | 1.46E-05 |
| GO:0099174 | regulation of presynapse organization | BP | 24 | 9 | 1 | 1.51E-05 |
| GO:0007204 | positive regulation of cytosolic calcium ion concentration | BP | 241 | 35 | 27 | 1.53E-05 |
| GO:0006816 | calcium ion transport | BP | 333 | 44 | 30 | 1.54E-05 |
| GO:0016529 | sarcoplasmic reticulum | CC | 56 | 14 | 4 | 1.57E-05 |
| GO:0034704 | calcium channel complex | CC | 49 | 13 | 2 | 1.57E-05 |
| GO:0009986 | cell surface | CC | 598 | 68 | 72 | 1.63E-05 |
| GO:0043034 | costamere | CC | 19 | 8 | 0 | 1.68E-05 |
| GO:0051050 | positive regulation of transport | BP | 773 | 83 | 66 | 1.68E-05 |
| GO:0099529 | neurotransmitter receptor activity involved in regulation of postsyn | MF | 30 | 10 | 1 | 1.69E-05 |
| GO:0098960 | postsynaptic neurotransmitter receptor activity | MF | 30 | 10 | 1 | 1.69E-05 |
| GO:0007200 | phospholipase C-activating G protein-coupled receptor signaling pa | BP | 64 | 15 | 4 | 1.82E-05 |
| GO:0008038 | neuron recognition | BP | 43 | 12 | 3 | 1.91E-05 |
| GO:0019233 | sensory perception of pain | BP | 72 | 16 | 6 | 1.98E-05 |
| GO:0086065 | cell communication involved in cardiac conduction | BP | 50 | 13 | 1 | 2.00E-05 |
| GO:0030072 | peptide hormone secretion | BP | 195 | 30 | 6 | 2.01E-05 |
| GO:0010038 | response to metal ion | BP | 285 | 39 | 17 | 2.08E-05 |
| GO:0048646 | anatomical structure formation involved in morphogenesis | BP | 873 | 91 | 51 | 2.09E-05 |
| GO:0097118 | neuroligin clustering involved in postsynaptic membrane assembly | BP | 4 | 4 | 0 | 2.11E-05 |
| GO:0008328 | ionotropic glutamate receptor complex | CC | 37 | 11 | 0 | 2.20E-05 |
| GO:0055013 | cardiac muscle cell development | BP | 65 | 15 | 1 | 2.22E-05 |
| GO:0097756 | negative regulation of blood vessel diameter | BP | 65 | 15 | 2 | 2.22E-05 |
| GO:2000300 | regulation of synaptic vesicle exocytosis | BP | 65 | 15 | 3 | 2.22E-05 |
| GO:0016528 | sarcoplasm | CC | 65 | 15 | 4 | 2.22E-05 |
| GO:1903779 | regulation of cardiac conduction | BP | 58 | 14 | 2 | 2.42E-05 |
| GO:0071248 | cellular response to metal ion | BP | 150 | 25 | 8 | 2.47E-05 |
| GO:0003018 | vascular process in circulatory system | BP | 132 | 23 | 8 | 2.49E-05 |
| GO:0004970 | ionotropic glutamate receptor activity | MF | 15 | 7 | 0 | 2.57E-05 |
| GO:0097091 | synaptic vesicle clustering | BP | 15 | 7 | 1 | 2.57E-05 |
| GO:0005251 | delayed rectifier potassium channel activity | MF | 20 | 8 | 3 | 2.63E-05 |
| GO:0019614 | catechol-containing compound catabolic process | BP | 7 | 5 | 0 | 2.66E-05 |
| GO:0042424 | catecholamine catabolic process | BP | 7 | 5 | 0 | 2.66E-05 |
| GO:0021675 | nerve development | BP | 66 | 15 | 5 | 2.70E-05 |
| GO:0098771 | inorganic ion homeostasis | BP | 551 | 63 | 48 | 2.84E-05 |
| GO:0009887 | animal organ morphogenesis | BP | 820 | 86 | 40 | 2.84E-05 |
| GO:0098878 | neurotransmitter receptor complex | CC | 38 | 11 | 0 | 2.90E-05 |
| GO:0050433 | regulation of catecholamine secretion | BP | 38 | 11 | 1 | 2.90E-05 |
| GO:0072503 | cellular divalent inorganic cation homeostasis | BP | 363 | 46 | 36 | 2.91E-05 |
| GO:0015844 | monoamine transport | BP | 59 | 14 | 1 | 2.97E-05 |
| GO:0030036 | actin cytoskeleton organization | BP | 575 | 65 | 38 | 3.02E-05 |
| GO:1903524 | positive regulation of blood circulation | BP | 52 | 13 | 5 | 3.15E-05 |
| GO:0098688 | parallel fiber to Purkinje cell synapse | CC | 11 | 6 | 0 | 3.30E-05 |
| GO:0097104 | postsynaptic membrane assembly | BP | 11 | 6 | 2 | 3.30E-05 |
| GO:0015696 | ammonium transport | BP | 75 | 16 | 3 | 3.40E-05 |
| GO:0019935 | cyclic-nucleotide-mediated signaling | BP | 144 | 24 | 9 | 3.57E-05 |
| GO:0055080 | cation homeostasis | BP | 544 | 62 | 48 | 3.62E-05 |
| GO:0044421 | extracellular region part | CC | 2377 | 207 | 172 | 3.80E-05 |
| GO:0051716 | cellular response to stimulus | BP | 5380 | 422 | 393 | 3.90E-05 |
| GO:0045956 | positive regulation of calcium ion-dependent exocytosis | BP | 21 | 8 | 0 | 4.00E-05 |
| GO:0006820 | anion transport | BP | 467 | 55 | 21 | 4.08E-05 |
| GO:0048588 | developmental cell growth | BP | 183 | 28 | 12 | 4.15E-05 |

| | | | | | | |
|---|---|---|---|---|---|---|
| GO:0086014 | atrial cardiac muscle cell action potential | BP | 16 | 7 | 1 | 4.30E-05 |
| GO:0086066 | atrial cardiac muscle cell to AV node cell communication | BP | 16 | 7 | 1 | 4.30E-05 |
| GO:0086026 | atrial cardiac muscle cell to AV node cell signaling | BP | 16 | 7 | 1 | 4.30E-05 |
| GO:0046942 | carboxylic acid transport | BP | 253 | 35 | 13 | 4.35E-05 |
| GO:0015849 | organic acid transport | BP | 253 | 35 | 13 | 4.35E-05 |
| GO:0048638 | regulation of developmental growth | BP | 253 | 35 | 11 | 4.35E-05 |
| GO:1901700 | response to oxygen-containing compound | BP | 1287 | 123 | 93 | 4.42E-05 |
| GO:0008306 | associative learning | BP | 61 | 14 | 4 | 4.43E-05 |
| GO:0086019 | cell-cell signaling involved in cardiac conduction | BP | 27 | 9 | 1 | 4.51E-05 |
| GO:0060306 | regulation of membrane repolarization | BP | 27 | 9 | 1 | 4.51E-05 |
| GO:0099094 | ligand-gated cation channel activity | MF | 69 | 15 | 5 | 4.71E-05 |
| GO:0031594 | neuromuscular junction | CC | 69 | 15 | 3 | 4.71E-05 |
| GO:0050432 | catecholamine secretion | BP | 40 | 11 | 1 | 4.92E-05 |
| GO:0005230 | extracellular ligand-gated ion channel activity | MF | 40 | 11 | 2 | 4.92E-05 |
| GO:0007626 | locomotory behavior | BP | 147 | 24 | 5 | 5.03E-05 |
| GO:0030073 | insulin secretion | BP | 167 | 26 | 6 | 5.73E-05 |
| GO:0060076 | excitatory synapse | CC | 34 | 10 | 1 | 5.76E-05 |
| GO:1903959 | regulation of anion transmembrane transport | BP | 22 | 8 | 1 | 5.91E-05 |
| GO:0050801 | ion homeostasis | BP | 611 | 67 | 52 | 5.92E-05 |
| GO:0030299 | intestinal cholesterol absorption | BP | 12 | 6 | 0 | 6.22E-05 |
| GO:0005184 | neuropeptide hormone activity | MF | 12 | 6 | 1 | 6.22E-05 |
| GO:0010644 | cell communication by electrical coupling | BP | 28 | 9 | 0 | 6.24E-05 |
| GO:0051937 | catecholamine transport | BP | 48 | 12 | 1 | 6.32E-05 |
| GO:0044420 | extracellular matrix component | CC | 48 | 12 | 0 | 6.32E-05 |
| GO:0021545 | cranial nerve development | BP | 41 | 11 | 4 | 6.32E-05 |
| GO:0030315 | T-tubule | CC | 41 | 11 | 0 | 6.32E-05 |
| GO:0007167 | enzyme linked receptor protein signaling pathway | BP | 827 | 85 | 50 | 6.54E-05 |
| GO:0014070 | response to organic cyclic compound | BP | 731 | 77 | 43 | 6.55E-05 |
| GO:0035051 | cardiocyte differentiation | BP | 122 | 21 | 5 | 6.66E-05 |
| GO:0014701 | junctional sarcoplasmic reticulum membrane | CC | 8 | 5 | 0 | 6.70E-05 |
| GO:0019336 | phenol-containing compound catabolic process | BP | 8 | 5 | 0 | 6.70E-05 |
| GO:0055007 | cardiac muscle cell differentiation | BP | 96 | 18 | 4 | 6.94E-05 |
| GO:0010976 | positive regulation of neuron projection development | BP | 239 | 33 | 11 | 7.40E-05 |
| GO:0072511 | divalent inorganic cation transport | BP | 379 | 46 | 32 | 8.42E-05 |
| GO:0007271 | synaptic transmission, cholinergic | BP | 23 | 8 | 1 | 8.53E-05 |
| GO:0017157 | regulation of exocytosis | BP | 181 | 27 | 14 | 8.84E-05 |
| GO:0060560 | developmental growth involved in morphogenesis | BP | 191 | 28 | 13 | 8.99E-05 |
| GO:0031589 | cell-substrate adhesion | BP | 315 | 40 | 22 | 9.08E-05 |
| GO:0060538 | skeletal muscle organ development | BP | 125 | 21 | 7 | 9.58E-05 |
| GO:0005911 | cell-cell junction | CC | 383 | 46 | 25 | 1.08E-04 |
| GO:0007519 | skeletal muscle tissue development | BP | 117 | 20 | 7 | 1.09E-04 |
| GO:0033267 | axon part | CC | 329 | 41 | 18 | 1.13E-04 |
| GO:0033017 | sarcoplasmic reticulum membrane | CC | 30 | 9 | 0 | 1.14E-04 |
| GO:0051015 | actin filament binding | MF | 174 | 26 | 15 | 1.15E-04 |
| GO:0005249 | voltage-gated potassium channel activity | MF | 51 | 12 | 4 | 1.19E-04 |
| GO:0031644 | regulation of neurological system process | BP | 100 | 18 | 4 | 1.20E-04 |
| GO:0019842 | vitamin binding | MF | 100 | 18 | 1 | 1.20E-04 |
| GO:0090092 | regulation of transmembrane receptor protein serine/threonine kina: | BP | 175 | 26 | 8 | 1.27E-04 |
| GO:0070838 | divalent metal ion transport | BP | 375 | 45 | 32 | 1.31E-04 |
| GO:0006875 | cellular metal ion homeostasis | BP | 431 | 50 | 40 | 1.32E-04 |
| GO:1904064 | positive regulation of cation transmembrane transport | BP | 110 | 19 | 11 | 1.39E-04 |
| GO:0098802 | plasma membrane receptor complex | CC | 147 | 23 | 19 | 1.39E-04 |
| GO:1903789 | regulation of amino acid transmembrane transport | BP | 9 | 5 | 0 | 1.42E-04 |
| GO:2000807 | regulation of synaptic vesicle clustering | BP | 9 | 5 | 0 | 1.42E-04 |
| GO:0089717 | spanning component of membrane | CC | 9 | 5 | 0 | 1.42E-04 |
| GO:0035265 | organ growth | BP | 138 | 22 | 5 | 1.46E-04 |
| GO:0071363 | cellular response to growth factor stimulus | BP | 536 | 59 | 27 | 1.47E-04 |
| GO:0032970 | regulation of actin filament-based process | BP | 333 | 41 | 23 | 1.48E-04 |
| GO:0010001 | glial cell differentiation | BP | 167 | 25 | 10 | 1.50E-04 |
| GO:0070588 | calcium ion transmembrane transport | BP | 248 | 33 | 21 | 1.52E-04 |
| GO:0050805 | negative regulation of synaptic transmission | BP | 60 | 13 | 0 | 1.54E-04 |
| GO:0018958 | phenol-containing compound metabolic process | BP | 68 | 14 | 9 | 1.55E-04 |
| GO:0034767 | positive regulation of ion transmembrane transport | BP | 120 | 20 | 13 | 1.56E-04 |
| GO:0007178 | transmembrane receptor protein serine/threonine kinase signaling p | BP | 259 | 34 | 11 | 1.58E-04 |

| | | | | | | |
|---|---|---|---|---|---|---|
| GO:0007188 | adenylate cyclase-modulating G protein-coupled receptor signaling | BP | 139 | 22 | 9 | 1.63E-04 |
| GO:0098856 | intestinal lipid absorption | BP | 14 | 6 | 0 | 1.80E-04 |
| GO:0006112 | energy reserve metabolic process | BP | 69 | 14 | 1 | 1.83E-04 |
| GO:0072358 | cardiovascular system development | BP | 588 | 63 | 27 | 1.90E-04 |
| GO:0016358 | dendrite development | BP | 210 | 29 | 11 | 1.98E-04 |
| GO:0005272 | sodium channel activity | MF | 32 | 9 | 1 | 1.98E-04 |
| GO:1902904 | negative regulation of supramolecular fiber organization | BP | 113 | 19 | 7 | 2.01E-04 |
| GO:0035239 | tube morphogenesis | BP | 685 | 71 | 37 | 2.06E-04 |
| GO:0060419 | heart growth | BP | 70 | 14 | 2 | 2.15E-04 |
| GO:0006073 | cellular glucan metabolic process | BP | 62 | 13 | 0 | 2.19E-04 |
| GO:0044042 | glucan metabolic process | BP | 62 | 13 | 0 | 2.19E-04 |
| GO:0019722 | calcium-mediated signaling | BP | 171 | 25 | 14 | 2.20E-04 |
| GO:0071241 | cellular response to inorganic substance | BP | 171 | 25 | 9 | 2.20E-04 |
| GO:0098796 | membrane protein complex | CC | 882 | 87 | 50 | 2.24E-04 |
| GO:0050954 | sensory perception of mechanical stimulus | BP | 114 | 19 | 6 | 2.26E-04 |
| GO:0086005 | ventricular cardiac muscle cell action potential | BP | 26 | 8 | 1 | 2.26E-04 |
| GO:0055082 | cellular chemical homeostasis | BP | 616 | 65 | 44 | 2.32E-04 |
| GO:0001764 | neuron migration | BP | 124 | 20 | 7 | 2.46E-04 |
| GO:0150034 | distal axon | CC | 244 | 32 | 15 | 2.49E-04 |
| GO:0040011 | locomotion | BP | 1418 | 129 | 124 | 2.55E-04 |
| GO:0045933 | positive regulation of muscle contraction | BP | 33 | 9 | 3 | 2.56E-04 |
| GO:0008217 | regulation of blood pressure | BP | 134 | 21 | 8 | 2.62E-04 |
| GO:0007165 | signal transduction | BP | 4276 | 338 | 335 | 2.67E-04 |
| GO:0030241 | skeletal muscle myosin thick filament assembly | BP | 10 | 5 | 0 | 2.69E-04 |
| GO:0071688 | striated muscle myosin thick filament assembly | BP | 10 | 5 | 0 | 2.69E-04 |
| GO:0070848 | response to growth factor | BP | 560 | 60 | 29 | 2.69E-04 |
| GO:0032414 | positive regulation of ion transmembrane transporter activity | BP | 80 | 15 | 10 | 2.74E-04 |
| GO:0035813 | regulation of renal sodium excretion | BP | 15 | 6 | 2 | 2.82E-04 |
| GO:0098910 | regulation of atrial cardiac muscle cell action potential | BP | 6 | 4 | 1 | 2.83E-04 |
| GO:0051093 | negative regulation of developmental process | BP | 717 | 73 | 48 | 2.88E-04 |
| GO:0051648 | vesicle localization | BP | 278 | 35 | 9 | 2.89E-04 |
| GO:0007605 | sensory perception of sound | BP | 98 | 17 | 5 | 2.92E-04 |
| GO:0042562 | hormone binding | MF | 72 | 14 | 1 | 2.93E-04 |
| GO:0006805 | xenobiotic metabolic process | BP | 72 | 14 | 1 | 2.93E-04 |
| GO:0042417 | dopamine metabolic process | BP | 27 | 8 | 4 | 3.03E-04 |
| GO:0055017 | cardiac muscle tissue growth | BP | 64 | 13 | 2 | 3.05E-04 |
| GO:1904063 | negative regulation of cation transmembrane transport | BP | 64 | 13 | 4 | 3.05E-04 |
| GO:0015711 | organic anion transport | BP | 367 | 43 | 18 | 3.10E-04 |
| GO:0097435 | supramolecular fiber organization | BP | 563 | 60 | 37 | 3.10E-04 |
| GO:0033002 | muscle cell proliferation | BP | 155 | 23 | 10 | 3.10E-04 |
| GO:0002074 | extraocular skeletal muscle development | BP | 3 | 3 | 0 | 3.11E-04 |
| GO:0021615 | glossopharyngeal nerve morphogenesis | BP | 3 | 3 | 0 | 3.11E-04 |
| GO:0016715 | oxidoreductase activity, acting on paired donors, with incorporatior | MF | 3 | 3 | 0 | 3.11E-04 |
| GO:0048662 | negative regulation of smooth muscle cell proliferation | BP | 41 | 10 | 3 | 3.19E-04 |
| GO:0086012 | membrane depolarization during cardiac muscle cell action potenti | BP | 21 | 7 | 0 | 3.24E-04 |
| GO:0010460 | positive regulation of heart rate | BP | 21 | 7 | 2 | 3.24E-04 |
| GO:0005245 | voltage-gated calcium channel activity | MF | 34 | 9 | 2 | 3.28E-04 |
| GO:0006942 | regulation of striated muscle contraction | BP | 73 | 14 | 4 | 3.40E-04 |
| GO:0001944 | vasculature development | BP | 579 | 61 | 27 | 3.75E-04 |
| GO:0016477 | cell migration | BP | 1136 | 106 | 104 | 3.85E-04 |
| GO:0035295 | tube development | BP | 835 | 82 | 43 | 3.91E-04 |
| GO:0034766 | negative regulation of ion transmembrane transport | BP | 74 | 14 | 4 | 3.94E-04 |
| GO:0005891 | voltage-gated calcium channel complex | CC | 28 | 8 | 1 | 3.99E-04 |
| GO:0001659 | temperature homeostasis | BP | 148 | 22 | 11 | 4.05E-04 |
| GO:0051179 | localization | BP | 5115 | 395 | 304 | 4.09E-04 |
| GO:0009712 | catechol-containing compound metabolic process | BP | 35 | 9 | 5 | 4.15E-04 |
| GO:0006584 | catecholamine metabolic process | BP | 35 | 9 | 5 | 4.15E-04 |
| GO:0006935 | chemotaxis | BP | 487 | 53 | 53 | 4.17E-04 |
| GO:0044853 | plasma membrane raft | CC | 92 | 16 | 10 | 4.23E-04 |
| GO:0045471 | response to ethanol | BP | 92 | 16 | 8 | 4.23E-04 |
| GO:0048739 | cardiac muscle fiber development | BP | 16 | 6 | 0 | 4.26E-04 |
| GO:0009725 | response to hormone | BP | 776 | 77 | 48 | 4.33E-04 |
| GO:0042330 | taxis | BP | 489 | 53 | 53 | 4.60E-04 |
| GO:0044306 | neuron projection terminus | CC | 111 | 18 | 8 | 4.61E-04 |

| | | | | | | |
|---|---|---|---|---|---|---|
| GO:0031034 | myosin filament assembly | BP | 11 | 5 | 0 | 4.65E-04 |
| GO:0005884 | actin filament | CC | 102 | 17 | 14 | 4.74E-04 |
| GO:0031103 | axon regeneration | BP | 43 | 10 | 0 | 4.82E-04 |
| GO:0007010 | cytoskeleton organization | BP | 1105 | 103 | 68 | 4.85E-04 |
| GO:0005901 | caveola | CC | 67 | 13 | 3 | 4.88E-04 |
| GO:0051051 | negative regulation of transport | BP | 387 | 44 | 22 | 5.08E-04 |
| GO:0031102 | neuron projection regeneration | BP | 51 | 11 | 0 | 5.10E-04 |
| GO:0048813 | dendrite morphogenesis | BP | 131 | 20 | 8 | 5.15E-04 |
| GO:1903305 | regulation of regulated secretory pathway | BP | 131 | 20 | 13 | 5.15E-04 |
| GO:0031672 | A band | CC | 29 | 8 | 1 | 5.19E-04 |
| GO:0045823 | positive regulation of heart contraction | BP | 29 | 8 | 3 | 5.19E-04 |
| GO:0044091 | membrane biogenesis | BP | 36 | 9 | 3 | 5.20E-04 |
| GO:0070161 | anchoring junction | CC | 516 | 55 | 35 | 5.44E-04 |
| GO:1901019 | regulation of calcium ion transmembrane transporter activity | BP | 68 | 13 | 3 | 5.66E-04 |
| GO:0006873 | cellular ion homeostasis | BP | 494 | 53 | 42 | 5.86E-04 |
| GO:0008021 | synaptic vesicle | CC | 162 | 23 | 5 | 5.90E-04 |
| GO:0032386 | regulation of intracellular transport | BP | 367 | 42 | 24 | 5.91E-04 |
| GO:0005912 | adherens junction | CC | 506 | 54 | 34 | 5.92E-04 |
| GO:0045987 | positive regulation of smooth muscle contraction | BP | 23 | 7 | 2 | 6.06E-04 |
| GO:0045907 | positive regulation of vasoconstriction | BP | 23 | 7 | 2 | 6.06E-04 |
| GO:0051592 | response to calcium ion | BP | 123 | 19 | 3 | 6.07E-04 |
| GO:0005179 | hormone activity | MF | 52 | 11 | 3 | 6.08E-04 |
| GO:0051705 | multi-organism behavior | BP | 52 | 11 | 1 | 6.08E-04 |
| GO:0032413 | negative regulation of ion transmembrane transporter activity | BP | 52 | 11 | 2 | 6.08E-04 |
| GO:0051924 | regulation of calcium ion transport | BP | 193 | 26 | 23 | 6.09E-04 |
| GO:0015749 | monosaccharide transmembrane transport | BP | 86 | 15 | 2 | 6.14E-04 |
| GO:0010771 | negative regulation of cell morphogenesis involved in differentiatio | BP | 86 | 15 | 7 | 6.14E-04 |
| GO:0005227 | calcium activated cation channel activity | MF | 17 | 6 | 0 | 6.21E-04 |
| GO:0035812 | renal sodium excretion | BP | 17 | 6 | 2 | 6.21E-04 |
| GO:0045760 | positive regulation of action potential | BP | 7 | 4 | 1 | 6.24E-04 |
| GO:0010958 | regulation of amino acid import across plasma membrane | BP | 7 | 4 | 0 | 6.24E-04 |
| GO:0048514 | blood vessel morphogenesis | BP | 484 | 52 | 25 | 6.37E-04 |
| GO:0060322 | head development | BP | 603 | 62 | 28 | 6.37E-04 |
| GO:0043583 | ear development | BP | 163 | 23 | 10 | 6.44E-04 |
| GO:0090276 | regulation of peptide hormone secretion | BP | 163 | 23 | 6 | 6.44E-04 |
| GO:0033275 | actin-myosin filament sliding | BP | 30 | 8 | 2 | 6.66E-04 |
| GO:0030049 | muscle filament sliding | BP | 30 | 8 | 2 | 6.66E-04 |
| GO:0008037 | cell recognition | BP | 105 | 17 | 9 | 6.69E-04 |
| GO:0044325 | ion channel binding | MF | 105 | 17 | 5 | 6.69E-04 |
| GO:0051384 | response to glucocorticoid | BP | 105 | 17 | 7 | 6.69E-04 |
| GO:0019933 | cAMP-mediated signaling | BP | 124 | 19 | 9 | 6.72E-04 |
| GO:1990845 | adaptive thermogenesis | BP | 134 | 20 | 11 | 6.93E-04 |
| GO:0042737 | drug catabolic process | BP | 87 | 15 | 3 | 6.96E-04 |
| GO:0005977 | glycogen metabolic process | BP | 61 | 12 | 0 | 6.99E-04 |
| GO:0050796 | regulation of insulin secretion | BP | 144 | 21 | 6 | 7.03E-04 |
| GO:0031346 | positive regulation of cell projection organization | BP | 325 | 38 | 16 | 7.06E-04 |
| GO:0050770 | regulation of axonogenesis | BP | 154 | 22 | 11 | 7.06E-04 |
| GO:0014704 | intercalated disc | CC | 45 | 10 | 0 | 7.09E-04 |
| GO:0006865 | amino acid transport | BP | 115 | 18 | 8 | 7.13E-04 |
| GO:1901342 | regulation of vasculature development | BP | 259 | 32 | 19 | 7.17E-04 |
| GO:0106106 | cold-induced thermogenesis | BP | 125 | 19 | 11 | 7.44E-04 |
| GO:0120161 | regulation of cold-induced thermogenesis | BP | 125 | 19 | 11 | 7.44E-04 |
| GO:0006883 | cellular sodium ion homeostasis | BP | 12 | 5 | 1 | 7.54E-04 |
| GO:0031033 | myosin filament organization | BP | 12 | 5 | 0 | 7.54E-04 |
| GO:0098685 | Schaffer collateral - CA1 synapse | CC | 70 | 13 | 4 | 7.55E-04 |
| GO:0032956 | regulation of actin cytoskeleton organization | BP | 293 | 35 | 19 | 7.69E-04 |
| GO:0098739 | import across plasma membrane | BP | 79 | 14 | 3 | 7.83E-04 |
| GO:0034219 | carbohydrate transmembrane transport | BP | 88 | 15 | 2 | 7.87E-04 |
| GO:0032411 | positive regulation of transporter activity | BP | 88 | 15 | 11 | 7.87E-04 |
| GO:0008643 | carbohydrate transport | BP | 116 | 18 | 2 | 7.92E-04 |
| GO:0042133 | neurotransmitter metabolic process | BP | 116 | 18 | 10 | 7.92E-04 |
| GO:0051241 | negative regulation of multicellular organismal process | BP | 930 | 88 | 77 | 7.96E-04 |
| GO:0009250 | glucan biosynthetic process | BP | 38 | 9 | 0 | 7.97E-04 |
| GO:0005978 | glycogen biosynthetic process | BP | 38 | 9 | 0 | 7.97E-04 |

| | | | | | | |
|---|---|---|---|---|---|---|
| GO:0021602 | cranial nerve morphogenesis | BP | 24 | 7 | 3 | 8.05E-04 |
| GO:0031430 | M band | CC | 24 | 7 | 1 | 8.05E-04 |
| GO:0045777 | positive regulation of blood pressure | BP | 24 | 7 | 2 | 8.05E-04 |
| GO:0010035 | response to inorganic substance | BP | 442 | 48 | 29 | 8.06E-04 |
| GO:0048278 | vesicle docking | BP | 62 | 12 | 0 | 8.14E-04 |
| GO:0033500 | carbohydrate homeostasis | BP | 197 | 26 | 3 | 8.31E-04 |
| GO:0042593 | glucose homeostasis | BP | 197 | 26 | 3 | 8.31E-04 |
| GO:0071709 | membrane assembly | BP | 31 | 8 | 3 | 8.44E-04 |
| GO:0097755 | positive regulation of blood vessel diameter | BP | 46 | 10 | 6 | 8.52E-04 |
| GO:0007420 | brain development | BP | 562 | 58 | 25 | 8.63E-04 |
| GO:0089718 | amino acid import across plasma membrane | BP | 18 | 6 | 0 | 8.78E-04 |
| GO:1901380 | negative regulation of potassium ion transmembrane transport | BP | 18 | 6 | 0 | 8.78E-04 |
| GO:0042135 | neurotransmitter catabolic process | BP | 18 | 6 | 0 | 8.78E-04 |
| GO:0044062 | regulation of excretion | BP | 18 | 6 | 3 | 8.78E-04 |
| GO:0070382 | exocytic vesicle | CC | 177 | 24 | 7 | 8.85E-04 |
| GO:0005788 | endoplasmic reticulum lumen | CC | 252 | 31 | 21 | 9.27E-04 |
| GO:0016202 | regulation of striated muscle tissue development | BP | 99 | 16 | 6 | 9.72E-04 |
| GO:0007157 | heterophilic cell-cell adhesion via plasma membrane cell adhesion [illegible] | BP | 39 | 9 | 3 | 9.75E-04 |
| GO:0001568 | blood vessel development | BP | 553 | 57 | 25 | 9.83E-04 |
| GO:0033555 | multicellular organismal response to stress | BP | 55 | 11 | 6 | 1.00E-03 |
| GO:0007160 | cell-matrix adhesion | BP | 200 | 26 | 13 | 1.04E-03 |
| GO:0072001 | renal system development | BP | 243 | 30 | 16 | 1.05E-03 |
| GO:0010880 | regulation of release of sequestered calcium ion into cytosol by sar[illegible] | BP | 25 | 7 | 0 | 1.05E-03 |
| GO:0070296 | sarcoplasmic reticulum calcium ion transport | BP | 32 | 8 | 1 | 1.06E-03 |
| GO:0042592 | homeostatic process | BP | 1496 | 131 | 115 | 1.08E-03 |
| GO:0044291 | cell-cell contact zone | CC | 64 | 12 | 0 | 1.09E-03 |
| GO:0034763 | negative regulation of transmembrane transport | BP | 91 | 15 | 5 | 1.13E-03 |
| GO:0010721 | negative regulation of cell development | BP | 255 | 31 | 14 | 1.13E-03 |
| GO:0140029 | exocytic process | BP | 73 | 13 | 3 | 1.14E-03 |
| GO:0044070 | regulation of anion transport | BP | 73 | 13 | 5 | 1.14E-03 |
| GO:1900025 | negative regulation of substrate adhesion-dependent cell spreading | BP | 13 | 5 | 1 | 1.16E-03 |
| GO:0060004 | reflex | BP | 13 | 5 | 2 | 1.16E-03 |
| GO:0048670 | regulation of collateral sprouting | BP | 13 | 5 | 1 | 1.16E-03 |
| GO:0043083 | synaptic cleft | CC | 13 | 5 | 0 | 1.16E-03 |
| GO:0021563 | glossopharyngeal nerve development | BP | 4 | 3 | 0 | 1.18E-03 |
| GO:0015168 | glycerol transmembrane transporter activity | MF | 4 | 3 | 0 | 1.18E-03 |
| GO:0015793 | glycerol transport | BP | 4 | 3 | 0 | 1.18E-03 |
| GO:0097109 | neuroligin family protein binding | MF | 4 | 3 | 0 | 1.18E-03 |
| GO:1900454 | positive regulation of long-term synaptic depression | BP | 4 | 3 | 0 | 1.18E-03 |
| GO:0032224 | positive regulation of synaptic transmission, cholinergic | BP | 4 | 3 | 1 | 1.18E-03 |
| GO:0015250 | water channel activity | MF | 4 | 3 | 0 | 1.18E-03 |
| GO:0005372 | water transmembrane transporter activity | MF | 4 | 3 | 0 | 1.18E-03 |
| GO:0048639 | positive regulation of developmental growth | BP | 130 | 19 | 6 | 1.21E-03 |
| GO:0016405 | CoA-ligase activity | MF | 19 | 6 | 1 | 1.21E-03 |
| GO:1905809 | negative regulation of synapse organization | BP | 19 | 6 | 1 | 1.21E-03 |
| GO:0010881 | regulation of cardiac muscle contraction by regulation of the releas[illegible] | BP | 19 | 6 | 0 | 1.21E-03 |
| GO:0005913 | cell-cell adherens junction | CC | 101 | 16 | 5 | 1.21E-03 |
| GO:0048562 | embryonic organ morphogenesis | BP | 213 | 27 | 9 | 1.24E-03 |
| GO:0042063 | gliogenesis | BP | 224 | 28 | 13 | 1.26E-03 |
| GO:0030003 | cellular cation homeostasis | BP | 487 | 51 | 42 | 1.27E-03 |
| GO:0040008 | regulation of growth | BP | 523 | 54 | 22 | 1.27E-03 |
| GO:0002028 | regulation of sodium ion transport | BP | 74 | 13 | 5 | 1.30E-03 |
| GO:0005615 | extracellular space | CC | 2228 | 185 | 168 | 1.31E-03 |
| GO:0015872 | dopamine transport | BP | 33 | 8 | 0 | 1.32E-03 |
| GO:0086091 | regulation of heart rate by cardiac conduction | BP | 33 | 8 | 0 | 1.32E-03 |
| GO:0070887 | cellular response to chemical stimulus | BP | 2488 | 204 | 190 | 1.32E-03 |
| GO:0005102 | signaling receptor binding | MF | 1074 | 98 | 90 | 1.35E-03 |
| GO:0048634 | regulation of muscle organ development | BP | 102 | 16 | 6 | 1.35E-03 |
| GO:1901861 | regulation of muscle tissue development | BP | 102 | 16 | 6 | 1.35E-03 |
| GO:0014046 | dopamine secretion | BP | 26 | 7 | 0 | 1.36E-03 |
| GO:0051955 | regulation of amino acid transport | BP | 26 | 7 | 3 | 1.36E-03 |
| GO:0098901 | regulation of cardiac muscle cell action potential | BP | 26 | 7 | 1 | 1.36E-03 |
| GO:0014059 | regulation of dopamine secretion | BP | 26 | 7 | 0 | 1.36E-03 |
| GO:0006904 | vesicle docking involved in exocytosis | BP | 41 | 9 | 0 | 1.43E-03 |

| | | | | | | |
|---|---|---|---|---|---|---|
| GO:2001259 | positive regulation of cation channel activity | BP | 49 | 10 | 7 | 1.43E-03 |
| GO:0004930 | G protein-coupled receptor activity | MF | 237 | 29 | 22 | 1.45E-03 |
| GO:0008645 | hexose transmembrane transport | BP | 84 | 14 | 2 | 1.46E-03 |
| GO:0060415 | muscle tissue morphogenesis | BP | 58 | 11 | 1 | 1.58E-03 |
| GO:0032410 | negative regulation of transporter activity | BP | 58 | 11 | 3 | 1.58E-03 |
| GO:0001956 | positive regulation of neurotransmitter secretion | BP | 20 | 6 | 1 | 1.63E-03 |
| GO:0099623 | regulation of cardiac muscle cell membrane repolarization | BP | 20 | 6 | 1 | 1.63E-03 |
| GO:0016525 | negative regulation of angiogenesis | BP | 85 | 14 | 6 | 1.64E-03 |
| GO:0030198 | extracellular matrix organization | BP | 306 | 35 | 17 | 1.65E-03 |
| GO:0055117 | regulation of cardiac muscle contraction | BP | 67 | 12 | 4 | 1.65E-03 |
| GO:0051128 | regulation of cellular component organization | BP | 2090 | 174 | 132 | 1.67E-03 |
| GO:0048709 | oligodendrocyte differentiation | BP | 76 | 13 | 3 | 1.67E-03 |
| GO:0099601 | regulation of neurotransmitter receptor activity | BP | 50 | 10 | 3 | 1.68E-03 |
| GO:0015370 | solute:sodium symporter activity | MF | 50 | 10 | 1 | 1.68E-03 |
| GO:0044448 | cell cortex part | CC | 154 | 21 | 9 | 1.68E-03 |
| GO:1901017 | negative regulation of potassium ion transmembrane transporter act | BP | 14 | 5 | 0 | 1.70E-03 |
| GO:0051580 | regulation of neurotransmitter uptake | BP | 14 | 5 | 0 | 1.70E-03 |
| GO:0014866 | skeletal myofibril assembly | BP | 14 | 5 | 0 | 1.70E-03 |
| GO:0032890 | regulation of organic acid transport | BP | 42 | 9 | 3 | 1.71E-03 |
| GO:0044060 | regulation of endocrine process | BP | 27 | 7 | 0 | 1.73E-03 |
| GO:0051147 | regulation of muscle cell differentiation | BP | 124 | 18 | 8 | 1.73E-03 |
| GO:0043679 | axon terminus | CC | 95 | 15 | 7 | 1.76E-03 |
| GO:1902903 | regulation of supramolecular fiber organization | BP | 296 | 34 | 17 | 1.78E-03 |
| GO:2000181 | negative regulation of blood vessel morphogenesis | BP | 86 | 14 | 6 | 1.84E-03 |
| GO:0001822 | kidney development | BP | 230 | 28 | 14 | 1.87E-03 |
| GO:0072341 | modified amino acid binding | MF | 68 | 12 | 2 | 1.89E-03 |
| GO:1901343 | negative regulation of vasculature development | BP | 96 | 15 | 8 | 1.96E-03 |
| GO:0044089 | positive regulation of cellular component biogenesis | BP | 473 | 49 | 34 | 1.96E-03 |
| GO:0060420 | regulation of heart growth | BP | 51 | 10 | 2 | 1.96E-03 |
| GO:0001941 | postsynaptic membrane organization | BP | 35 | 8 | 2 | 1.98E-03 |
| GO:0035930 | corticosteroid hormone secretion | BP | 9 | 4 | 0 | 2.01E-03 |
| GO:0021604 | cranial nerve structural organization | BP | 9 | 4 | 1 | 2.01E-03 |
| GO:1990504 | dense core granule exocytosis | BP | 9 | 4 | 0 | 2.01E-03 |
| GO:0048251 | elastic fiber assembly | BP | 9 | 4 | 0 | 2.01E-03 |
| GO:0005916 | fascia adherens | CC | 9 | 4 | 0 | 2.01E-03 |
| GO:0072178 | nephric duct morphogenesis | BP | 9 | 4 | 1 | 2.01E-03 |
| GO:1990535 | neuron projection maintenance | BP | 9 | 4 | 0 | 2.01E-03 |
| GO:2000302 | positive regulation of synaptic vesicle exocytosis | BP | 9 | 4 | 0 | 2.01E-03 |
| GO:0097119 | postsynaptic density protein 95 clustering | BP | 9 | 4 | 0 | 2.01E-03 |
| GO:2000846 | regulation of corticosteroid hormone secretion | BP | 9 | 4 | 0 | 2.01E-03 |
| GO:2001135 | regulation of endocytic recycling | BP | 9 | 4 | 0 | 2.01E-03 |
| GO:2000831 | regulation of steroid hormone secretion | BP | 9 | 4 | 0 | 2.01E-03 |
| GO:0098917 | retrograde trans-synaptic signaling | BP | 9 | 4 | 0 | 2.01E-03 |
| GO:0031960 | response to corticosteroid | BP | 116 | 17 | 8 | 2.07E-03 |
| GO:0050953 | sensory perception of light stimulus | BP | 126 | 18 | 4 | 2.08E-03 |
| GO:0006835 | dicarboxylic acid transport | BP | 78 | 13 | 5 | 2.13E-03 |
| GO:0071277 | cellular response to calcium ion | BP | 69 | 12 | 2 | 2.15E-03 |
| GO:0051149 | positive regulation of muscle cell differentiation | BP | 69 | 12 | 7 | 2.15E-03 |
| GO:0043090 | amino acid import | BP | 21 | 6 | 0 | 2.15E-03 |
| GO:0007413 | axonal fasciculation | BP | 21 | 6 | 3 | 2.15E-03 |
| GO:0106030 | neuron projection fasciculation | BP | 21 | 6 | 3 | 2.15E-03 |
| GO:0009605 | response to external stimulus | BP | 1751 | 148 | 169 | 2.17E-03 |
| GO:0060986 | endocrine hormone secretion | BP | 28 | 7 | 1 | 2.17E-03 |
| GO:0098801 | regulation of renal system process | BP | 28 | 7 | 3 | 2.17E-03 |
| GO:0014808 | release of sequestered calcium ion into cytosol by sarcoplasmic reti | BP | 28 | 7 | 1 | 2.17E-03 |
| GO:0043197 | dendritic spine | CC | 147 | 20 | 7 | 2.20E-03 |
| GO:0010033 | response to organic substance | BP | 2529 | 205 | 186 | 2.26E-03 |
| GO:0048870 | cell motility | BP | 1236 | 109 | 114 | 2.31E-03 |
| GO:0051674 | localization of cell | BP | 1236 | 109 | 114 | 2.31E-03 |
| GO:0045121 | membrane raft | CC | 267 | 31 | 30 | 2.35E-03 |
| GO:0060079 | excitatory postsynaptic potential | BP | 79 | 13 | 4 | 2.39E-03 |
| GO:0048483 | autonomic nervous system development | BP | 36 | 8 | 3 | 2.40E-03 |
| GO:0060038 | cardiac muscle cell proliferation | BP | 36 | 8 | 2 | 2.40E-03 |
| GO:0001504 | neurotransmitter uptake | BP | 36 | 8 | 0 | 2.40E-03 |

| | | | | | | |
|---|---|---|---|---|---|---|
| GO:0015459 | potassium channel regulator activity | MF | 36 | 8 | 1 | 2.40E-03 |
| GO:0050795 | regulation of behavior | BP | 44 | 9 | 3 | 2.40E-03 |
| GO:0043194 | axon initial segment | CC | 15 | 5 | 2 | 2.41E-03 |
| GO:0031045 | dense core granule | CC | 15 | 5 | 0 | 2.41E-03 |
| GO:0072243 | metanephric nephron epithelium development | BP | 15 | 5 | 0 | 2.41E-03 |
| GO:0071855 | neuropeptide receptor binding | MF | 15 | 5 | 0 | 2.41E-03 |
| GO:0098656 | anion transmembrane transport | BP | 212 | 26 | 6 | 2.41E-03 |
| GO:0043178 | alcohol binding | MF | 70 | 12 | 3 | 2.43E-03 |
| GO:1901879 | regulation of protein depolymerization | BP | 70 | 12 | 1 | 2.43E-03 |
| GO:0098857 | membrane microdomain | CC | 268 | 31 | 30 | 2.49E-03 |
| GO:0019725 | cellular homeostasis | BP | 751 | 71 | 57 | 2.58E-03 |
| GO:0044309 | neuron spine | CC | 149 | 20 | 7 | 2.59E-03 |
| GO:0065007 | biological regulation | BP | 8733 | 631 | 524 | 2.59E-03 |
| GO:0032835 | glomerulus development | BP | 53 | 10 | 2 | 2.65E-03 |
| GO:0044304 | main axon | CC | 53 | 10 | 4 | 2.65E-03 |
| GO:0048636 | positive regulation of muscle organ development | BP | 53 | 10 | 4 | 2.65E-03 |
| GO:0045844 | positive regulation of striated muscle tissue development | BP | 53 | 10 | 4 | 2.65E-03 |
| GO:0050905 | neuromuscular process | BP | 80 | 13 | 3 | 2.68E-03 |
| GO:0051590 | positive regulation of neurotransmitter transport | BP | 29 | 7 | 2 | 2.70E-03 |
| GO:1903514 | release of sequestered calcium ion into cytosol by endoplasmic reti | BP | 29 | 7 | 1 | 2.70E-03 |
| GO:0051932 | synaptic transmission, GABAergic | BP | 29 | 7 | 2 | 2.70E-03 |
| GO:0060402 | calcium ion transport into cytosol | BP | 129 | 18 | 14 | 2.71E-03 |
| GO:0060041 | retina development in camera-type eye | BP | 109 | 16 | 2 | 2.72E-03 |
| GO:0043062 | extracellular structure organization | BP | 350 | 38 | 21 | 2.72E-03 |
| GO:0045778 | positive regulation of ossification | BP | 71 | 12 | 6 | 2.75E-03 |
| GO:0019199 | transmembrane receptor protein kinase activity | MF | 71 | 12 | 1 | 2.75E-03 |
| GO:0048644 | muscle organ morphogenesis | BP | 62 | 11 | 1 | 2.75E-03 |
| GO:0055024 | regulation of cardiac muscle tissue development | BP | 62 | 11 | 4 | 2.75E-03 |
| GO:0001655 | urogenital system development | BP | 270 | 31 | 17 | 2.79E-03 |
| GO:0007340 | acrosome reaction | BP | 22 | 6 | 1 | 2.79E-03 |
| GO:0060314 | regulation of ryanodine-sensitive calcium-release channel activity | BP | 22 | 6 | 0 | 2.79E-03 |
| GO:0005004 | GPI-linked ephrin receptor activity | MF | 5 | 3 | 0 | 2.81E-03 |
| GO:0015277 | kainate selective glutamate receptor activity | MF | 5 | 3 | 0 | 2.81E-03 |
| GO:0032983 | kainate selective glutamate receptor complex | CC | 5 | 3 | 0 | 2.81E-03 |
| GO:0086045 | membrane depolarization during AV node cell action potential | BP | 5 | 3 | 0 | 2.81E-03 |
| GO:0072257 | metanephric nephron tubule epithelial cell differentiation | BP | 5 | 3 | 0 | 2.81E-03 |
| GO:0097025 | MPP7-DLG1-LIN7 complex | CC | 5 | 3 | 0 | 2.81E-03 |
| GO:0004499 | N,N-dimethylaniline monooxygenase activity | MF | 5 | 3 | 0 | 2.81E-03 |
| GO:0015205 | nucleobase transmembrane transporter activity | MF | 5 | 3 | 0 | 2.81E-03 |
| GO:2000809 | positive regulation of synaptic vesicle clustering | BP | 5 | 3 | 0 | 2.81E-03 |
| GO:0030300 | regulation of intestinal cholesterol absorption | BP | 5 | 3 | 0 | 2.81E-03 |
| GO:1904729 | regulation of intestinal lipid absorption | BP | 5 | 3 | 0 | 2.81E-03 |
| GO:0072307 | regulation of metanephric nephron tubule epithelial cell differentiat | BP | 5 | 3 | 0 | 2.81E-03 |
| GO:0044214 | spanning component of plasma membrane | CC | 5 | 3 | 0 | 2.81E-03 |
| GO:0070327 | thyroid hormone transport | BP | 5 | 3 | 0 | 2.81E-03 |
| GO:0086007 | voltage-gated calcium channel activity involved in cardiac muscle c | MF | 5 | 3 | 0 | 2.81E-03 |
| GO:0055021 | regulation of cardiac muscle tissue growth | BP | 45 | 9 | 2 | 2.83E-03 |
| GO:0034330 | cell junction organization | BP | 259 | 30 | 6 | 2.84E-03 |
| GO:0009636 | response to toxic substance | BP | 398 | 42 | 34 | 2.85E-03 |
| GO:0030170 | pyridoxal phosphate binding | MF | 37 | 8 | 0 | 2.88E-03 |
| GO:0070279 | vitamin B6 binding | MF | 37 | 8 | 0 | 2.88E-03 |
| GO:0009063 | cellular amino acid catabolic process | BP | 100 | 15 | 3 | 2.95E-03 |
| GO:1904659 | glucose transmembrane transport | BP | 81 | 13 | 2 | 3.00E-03 |
| GO:0050839 | cell adhesion molecule binding | MF | 435 | 45 | 20 | 3.02E-03 |
| GO:0045596 | negative regulation of cell differentiation | BP | 532 | 53 | 37 | 3.04E-03 |
| GO:0002088 | lens development in camera-type eye | BP | 54 | 10 | 3 | 3.06E-03 |
| GO:1901863 | positive regulation of muscle tissue development | BP | 54 | 10 | 4 | 3.06E-03 |
| GO:0005200 | structural constituent of cytoskeleton | MF | 72 | 12 | 3 | 3.10E-03 |
| GO:0001656 | metanephros development | BP | 63 | 11 | 3 | 3.14E-03 |
| GO:0048786 | presynaptic active zone | CC | 63 | 11 | 2 | 3.14E-03 |
| GO:0003996 | acyl-CoA ligase activity | MF | 10 | 4 | 1 | 3.18E-03 |
| GO:0086016 | AV node cell action potential | BP | 10 | 4 | 0 | 3.18E-03 |
| GO:0086027 | AV node cell to bundle of His cell signaling | BP | 10 | 4 | 0 | 3.18E-03 |
| GO:0061577 | calcium ion transmembrane transport via high voltage-gated calciu | BP | 10 | 4 | 0 | 3.18E-03 |

| | | | | | | |
|---|---|---|---|---|---|---|
| GO:0038063 | collagen-activated tyrosine kinase receptor signaling pathway | BP | 10 | 4 | 0 | 3.18E-03 |
| GO:0035815 | positive regulation of renal sodium excretion | BP | 10 | 4 | 1 | 3.18E-03 |
| GO:0072182 | regulation of nephron tubule epithelial cell differentiation | BP | 10 | 4 | 2 | 3.18E-03 |
| GO:0001518 | voltage-gated sodium channel complex | CC | 10 | 4 | 0 | 3.18E-03 |
| GO:0016323 | basolateral plasma membrane | CC | 173 | 22 | 9 | 3.22E-03 |
| GO:0021533 | cell differentiation in hindbrain | BP | 16 | 5 | 0 | 3.31E-03 |
| GO:0030021 | extracellular matrix structural constituent conferring compression r | MF | 16 | 5 | 1 | 3.31E-03 |
| GO:0032809 | neuronal cell body membrane | CC | 16 | 5 | 0 | 3.31E-03 |
| GO:0098831 | presynaptic active zone cytoplasmic component | CC | 16 | 5 | 0 | 3.31E-03 |
| GO:0071625 | vocalization behavior | BP | 16 | 5 | 0 | 3.31E-03 |
| GO:0014855 | striated muscle cell proliferation | BP | 46 | 9 | 2 | 3.31E-03 |
| GO:0015144 | carbohydrate transmembrane transporter activity | MF | 30 | 7 | 0 | 3.32E-03 |
| GO:1901618 | organic hydroxy compound transmembrane transporter activity | MF | 30 | 7 | 1 | 3.32E-03 |
| GO:2000310 | regulation of NMDA receptor activity | BP | 30 | 7 | 3 | 3.32E-03 |
| GO:0033762 | response to glucagon | BP | 30 | 7 | 4 | 3.32E-03 |
| GO:0048871 | multicellular organismal homeostasis | BP | 378 | 40 | 33 | 3.38E-03 |
| GO:0048839 | inner ear development | BP | 142 | 19 | 10 | 3.39E-03 |
| GO:1902305 | regulation of sodium ion transmembrane transport | BP | 55 | 10 | 3 | 3.52E-03 |
| GO:0051130 | positive regulation of cellular component organization | BP | 1069 | 95 | 75 | 3.52E-03 |
| GO:0016878 | acid-thiol ligase activity | MF | 23 | 6 | 1 | 3.56E-03 |
| GO:0086064 | cell communication by electrical coupling involved in cardiac cond | BP | 23 | 6 | 0 | 3.56E-03 |
| GO:0009649 | entrainment of circadian clock | BP | 23 | 6 | 0 | 3.56E-03 |
| GO:0060384 | innervation | BP | 23 | 6 | 1 | 3.56E-03 |
| GO:0048265 | response to pain | BP | 23 | 6 | 2 | 3.56E-03 |
| GO:0051650 | establishment of vesicle localization | BP | 263 | 30 | 7 | 3.57E-03 |
| GO:0060627 | regulation of vesicle-mediated transport | BP | 439 | 45 | 32 | 3.59E-03 |
| GO:1901606 | alpha-amino acid catabolic process | BP | 83 | 13 | 3 | 3.73E-03 |
| GO:0044264 | cellular polysaccharide metabolic process | BP | 83 | 13 | 0 | 3.73E-03 |
| GO:0050789 | regulation of biological process | BP | 8258 | 598 | 503 | 3.78E-03 |
| GO:0046323 | glucose import | BP | 47 | 9 | 0 | 3.85E-03 |
| GO:0090101 | negative regulation of transmembrane receptor protein serine/threo | BP | 93 | 14 | 5 | 3.87E-03 |
| GO:0007601 | visual perception | BP | 123 | 17 | 4 | 3.87E-03 |
| GO:0098589 | membrane region | CC | 276 | 31 | 30 | 3.89E-03 |
| GO:0072006 | nephron development | BP | 113 | 16 | 8 | 3.93E-03 |
| GO:0007389 | pattern specification process | BP | 311 | 34 | 14 | 3.98E-03 |
| GO:0098686 | hippocampal mossy fiber to CA3 synapse | CC | 31 | 7 | 0 | 4.04E-03 |
| GO:0051703 | intraspecies interaction between organisms | BP | 39 | 8 | 0 | 4.07E-03 |
| GO:0035176 | social behavior | BP | 39 | 8 | 0 | 4.07E-03 |
| GO:0016049 | cell growth | BP | 394 | 41 | 19 | 4.07E-03 |
| GO:0099565 | chemical synaptic transmission, postsynaptic | BP | 84 | 13 | 4 | 4.15E-03 |
| GO:0045765 | regulation of angiogenesis | BP | 232 | 27 | 17 | 4.21E-03 |
| GO:0021987 | cerebral cortex development | BP | 94 | 14 | 6 | 4.27E-03 |
| GO:0034446 | substrate adhesion-dependent cell spreading | BP | 94 | 14 | 8 | 4.27E-03 |
| GO:1905952 | regulation of lipid localization | BP | 104 | 15 | 6 | 4.31E-03 |
| GO:0035690 | cellular response to drug | BP | 301 | 33 | 26 | 4.31E-03 |
| GO:0098533 | ATPase dependent transmembrane transport complex | CC | 17 | 5 | 1 | 4.43E-03 |
| GO:0044298 | cell body membrane | CC | 17 | 5 | 0 | 4.43E-03 |
| GO:0072207 | metanephric epithelium development | BP | 17 | 5 | 0 | 4.43E-03 |
| GO:0032982 | myosin filament | CC | 17 | 5 | 1 | 4.43E-03 |
| GO:0051953 | negative regulation of amine transport | BP | 17 | 5 | 0 | 4.43E-03 |
| GO:0010766 | negative regulation of sodium ion transport | BP | 17 | 5 | 1 | 4.43E-03 |
| GO:0048486 | parasympathetic nervous system development | BP | 17 | 5 | 2 | 4.43E-03 |
| GO:0048714 | positive regulation of oligodendrocyte differentiation | BP | 17 | 5 | 0 | 4.43E-03 |
| GO:0060307 | regulation of ventricular cardiac muscle cell membrane repolarizati | BP | 17 | 5 | 1 | 4.43E-03 |
| GO:0048485 | sympathetic nervous system development | BP | 17 | 5 | 1 | 4.43E-03 |
| GO:0019198 | transmembrane receptor protein phosphatase activity | MF | 17 | 5 | 1 | 4.43E-03 |
| GO:0005001 | transmembrane receptor protein tyrosine phosphatase activity | MF | 17 | 5 | 1 | 4.43E-03 |
| GO:0021543 | pallium development | BP | 135 | 18 | 8 | 4.44E-03 |
| GO:0030042 | actin filament depolymerization | BP | 48 | 9 | 1 | 4.46E-03 |
| GO:0048747 | muscle fiber development | BP | 48 | 9 | 2 | 4.46E-03 |
| GO:0001786 | phosphatidylserine binding | MF | 48 | 9 | 1 | 4.46E-03 |
| GO:0010882 | regulation of cardiac muscle contraction by calcium ion signaling | BP | 24 | 6 | 0 | 4.48E-03 |
| GO:0006821 | chloride transport | BP | 66 | 11 | 2 | 4.55E-03 |
| GO:1901698 | response to nitrogen compound | BP | 871 | 79 | 59 | 4.55E-03 |

| | | | | | | |
|---|---|---|---|---|---|---|
| GO:0002790 | peptide secretion | BP | 469 | 47 | 39 | 4.55E-03 |
| GO:0097305 | response to alcohol | BP | 178 | 22 | 15 | 4.56E-03 |
| GO:0048640 | negative regulation of developmental growth | BP | 85 | 13 | 6 | 4.60E-03 |
| GO:0048814 | regulation of dendrite morphogenesis | BP | 85 | 13 | 4 | 4.60E-03 |
| GO:0032526 | response to retinoic acid | BP | 85 | 13 | 2 | 4.60E-03 |
| GO:0047635 | alanine-oxo-acid transaminase activity | MF | 2 | 2 | 0 | 4.60E-03 |
| GO:0006524 | alanine catabolic process | BP | 2 | 2 | 0 | 4.60E-03 |
| GO:0006522 | alanine metabolic process | BP | 2 | 2 | 0 | 4.60E-03 |
| GO:0046305 | alkanesulfonate biosynthetic process | BP | 2 | 2 | 0 | 4.60E-03 |
| GO:0004062 | aryl sulfotransferase activity | MF | 2 | 2 | 0 | 4.60E-03 |
| GO:0090537 | CERF complex | CC | 2 | 2 | 0 | 4.60E-03 |
| GO:0004692 | cGMP-dependent protein kinase activity | MF | 2 | 2 | 0 | 4.60E-03 |
| GO:0004500 | dopamine beta-monooxygenase activity | MF | 2 | 2 | 0 | 4.60E-03 |
| GO:0003199 | endocardial cushion to mesenchymal transition involved in heart va | BP | 2 | 2 | 0 | 4.60E-03 |
| GO:0060752 | intestinal phytosterol absorption | BP | 2 | 2 | 0 | 4.60E-03 |
| GO:0042853 | L-alanine catabolic process | BP | 2 | 2 | 0 | 4.60E-03 |
| GO:0042851 | L-alanine metabolic process | BP | 2 | 2 | 0 | 4.60E-03 |
| GO:0004021 | L-alanine:2-oxoglutarate aminotransferase activity | MF | 2 | 2 | 0 | 4.60E-03 |
| GO:0097089 | methyl-branched fatty acid metabolic process | BP | 2 | 2 | 0 | 4.60E-03 |
| GO:1904027 | negative regulation of collagen fibril organization | BP | 2 | 2 | 0 | 4.60E-03 |
| GO:0045720 | negative regulation of integrin biosynthetic process | BP | 2 | 2 | 0 | 4.60E-03 |
| GO:0045796 | negative regulation of intestinal cholesterol absorption | BP | 2 | 2 | 0 | 4.60E-03 |
| GO:1904730 | negative regulation of intestinal lipid absorption | BP | 2 | 2 | 0 | 4.60E-03 |
| GO:0010949 | negative regulation of intestinal phytosterol absorption | BP | 2 | 2 | 0 | 4.60E-03 |
| GO:0030185 | nitric oxide transport | BP | 2 | 2 | 0 | 4.60E-03 |
| GO:0006589 | octopamine biosynthetic process | BP | 2 | 2 | 0 | 4.60E-03 |
| GO:0046333 | octopamine metabolic process | BP | 2 | 2 | 0 | 4.60E-03 |
| GO:0035106 | operant conditioning | BP | 2 | 2 | 0 | 4.60E-03 |
| GO:0046997 | oxidoreductase activity, acting on the CH-NH group of donors, flav | MF | 2 | 2 | 0 | 4.60E-03 |
| GO:0086097 | phospholipase C-activating angiotensin-activated signaling pathwa | BP | 2 | 2 | 0 | 4.60E-03 |
| GO:1902608 | positive regulation of large conductance calcium-activated potassiu | BP | 2 | 2 | 0 | 4.60E-03 |
| GO:0046878 | positive regulation of saliva secretion | BP | 2 | 2 | 0 | 4.60E-03 |
| GO:1904071 | presynaptic active zone assembly | BP | 2 | 2 | 0 | 4.60E-03 |
| GO:0009080 | pyruvate family amino acid catabolic process | BP | 2 | 2 | 0 | 4.60E-03 |
| GO:0009078 | pyruvate family amino acid metabolic process | BP | 2 | 2 | 0 | 4.60E-03 |
| GO:0051941 | regulation of amino acid uptake involved in synaptic transmission | BP | 2 | 2 | 0 | 4.60E-03 |
| GO:0051946 | regulation of glutamate uptake involved in transmission of nerve in | BP | 2 | 2 | 0 | 4.60E-03 |
| GO:1902606 | regulation of large conductance calcium-activated potassium chann | BP | 2 | 2 | 0 | 4.60E-03 |
| GO:0098907 | regulation of SA node cell action potential | BP | 2 | 2 | 0 | 4.60E-03 |
| GO:0050758 | regulation of thymidylate synthase biosynthetic process | BP | 2 | 2 | 0 | 4.60E-03 |
| GO:0002018 | renin-angiotensin regulation of aldosterone production | BP | 2 | 2 | 0 | 4.60E-03 |
| GO:0099642 | retrograde axonal protein transport | BP | 2 | 2 | 0 | 4.60E-03 |
| GO:0042412 | taurine biosynthetic process | BP | 2 | 2 | 0 | 4.60E-03 |
| GO:0050757 | thymidylate synthase biosynthetic process | BP | 2 | 2 | 0 | 4.60E-03 |
| GO:0097553 | calcium ion transmembrane import into cytosol | BP | 115 | 16 | 13 | 4.68E-03 |
| GO:0019838 | growth factor binding | MF | 115 | 16 | 4 | 4.68E-03 |
| GO:0022804 | active transmembrane transporter activity | MF | 268 | 30 | 7 | 4.69E-03 |
| GO:0009855 | determination of bilateral symmetry | BP | 95 | 14 | 1 | 4.71E-03 |
| GO:0031223 | auditory behavior | BP | 11 | 4 | 0 | 4.73E-03 |
| GO:0086067 | AV node cell to bundle of His cell communication | BP | 11 | 4 | 0 | 4.73E-03 |
| GO:0005513 | detection of calcium ion | BP | 11 | 4 | 0 | 4.73E-03 |
| GO:0050667 | homocysteine metabolic process | BP | 11 | 4 | 1 | 4.73E-03 |
| GO:0005614 | interstitial matrix | CC | 11 | 4 | 0 | 4.73E-03 |
| GO:0045986 | negative regulation of smooth muscle contraction | BP | 11 | 4 | 0 | 4.73E-03 |
| GO:2000650 | negative regulation of sodium ion transmembrane transporter activi | BP | 11 | 4 | 1 | 4.73E-03 |
| GO:0051152 | positive regulation of smooth muscle cell differentiation | BP | 11 | 4 | 1 | 4.73E-03 |
| GO:1902805 | positive regulation of synaptic vesicle transport | BP | 11 | 4 | 0 | 4.73E-03 |
| GO:0060134 | prepulse inhibition | BP | 11 | 4 | 1 | 4.73E-03 |
| GO:0014874 | response to stimulus involved in regulation of muscle adaptation | BP | 11 | 4 | 0 | 4.73E-03 |
| GO:0019841 | retinol binding | MF | 11 | 4 | 0 | 4.73E-03 |
| GO:0035929 | steroid hormone secretion | BP | 11 | 4 | 0 | 4.73E-03 |
| GO:0016081 | synaptic vesicle docking | BP | 11 | 4 | 0 | 4.73E-03 |
| GO:0014820 | tonic smooth muscle contraction | BP | 11 | 4 | 0 | 4.73E-03 |
| GO:0140253 | cell-cell fusion | BP | 40 | 8 | 5 | 4.79E-03 |

| | | | | | | |
|---|---|---|---|---|---|---|
| GO:0030020 | extracellular matrix structural constituent conferring tensile strengt | MF | 40 | 8 | 0 | 4.79E-03 |
| GO:1904645 | response to amyloid-beta | BP | 40 | 8 | 3 | 4.79E-03 |
| GO:0000768 | syncytium formation by plasma membrane fusion | BP | 40 | 8 | 5 | 4.79E-03 |
| GO:0030545 | receptor regulator activity | MF | 257 | 29 | 25 | 4.81E-03 |
| GO:0001525 | angiogenesis | BP | 410 | 42 | 24 | 4.85E-03 |
| GO:1904646 | cellular response to amyloid-beta | BP | 32 | 7 | 3 | 4.87E-03 |
| GO:0007520 | myoblast fusion | BP | 32 | 7 | 5 | 4.87E-03 |
| GO:0010765 | positive regulation of sodium ion transport | BP | 32 | 7 | 1 | 4.87E-03 |
| GO:0071548 | response to dexamethasone | BP | 32 | 7 | 4 | 4.87E-03 |
| GO:0001501 | skeletal system development | BP | 387 | 40 | 23 | 5.07E-03 |
| GO:0060291 | long-term synaptic potentiation | BP | 67 | 11 | 3 | 5.12E-03 |
| GO:0045685 | regulation of glial cell differentiation | BP | 49 | 9 | 1 | 5.14E-03 |
| GO:0014823 | response to activity | BP | 49 | 9 | 0 | 5.14E-03 |
| GO:0046887 | positive regulation of hormone secretion | BP | 96 | 14 | 2 | 5.17E-03 |
| GO:0009799 | specification of symmetry | BP | 96 | 14 | 1 | 5.17E-03 |
| GO:0050773 | regulation of dendrite development | BP | 137 | 18 | 6 | 5.19E-03 |
| GO:0071286 | cellular response to magnesium ion | BP | 6 | 3 | 0 | 5.33E-03 |
| GO:0021684 | cerebellar granular layer formation | BP | 6 | 3 | 0 | 5.33E-03 |
| GO:0021707 | cerebellar granule cell differentiation | BP | 6 | 3 | 0 | 5.33E-03 |
| GO:0005587 | collagen type IV trimer | CC | 6 | 3 | 0 | 5.33E-03 |
| GO:0030314 | junctional membrane complex | CC | 6 | 3 | 0 | 5.33E-03 |
| GO:1990454 | L-type voltage-gated calcium channel complex | CC | 6 | 3 | 0 | 5.33E-03 |
| GO:1904753 | negative regulation of vascular associated smooth muscle cell migra | BP | 6 | 3 | 0 | 5.33E-03 |
| GO:1901166 | neural crest cell migration involved in autonomic nervous system de | BP | 6 | 3 | 0 | 5.33E-03 |
| GO:0042421 | norepinephrine biosynthetic process | BP | 6 | 3 | 1 | 5.33E-03 |
| GO:0019755 | one-carbon compound transport | BP | 6 | 3 | 0 | 5.33E-03 |
| GO:1905065 | positive regulation of vascular smooth muscle cell differentiation | BP | 6 | 3 | 0 | 5.33E-03 |
| GO:0086068 | Purkinje myocyte to ventricular cardiac muscle cell communication | BP | 6 | 3 | 0 | 5.33E-03 |
| GO:0086029 | Purkinje myocyte to ventricular cardiac muscle cell signaling | BP | 6 | 3 | 0 | 5.33E-03 |
| GO:0030900 | forebrain development | BP | 282 | 31 | 18 | 5.34E-03 |
| GO:1990778 | protein localization to cell periphery | BP | 282 | 31 | 12 | 5.34E-03 |
| GO:0009306 | protein secretion | BP | 449 | 45 | 39 | 5.42E-03 |
| GO:0060292 | long-term synaptic depression | BP | 25 | 6 | 0 | 5.57E-03 |
| GO:0051954 | positive regulation of amine transport | BP | 25 | 6 | 3 | 5.57E-03 |
| GO:0032228 | regulation of synaptic transmission, GABAergic | BP | 25 | 6 | 2 | 5.57E-03 |
| GO:0005544 | calcium-dependent phospholipid binding | MF | 41 | 8 | 0 | 5.60E-03 |
| GO:1990138 | neuron projection extension | BP | 138 | 18 | 11 | 5.61E-03 |
| GO:0001101 | response to acid chemical | BP | 283 | 31 | 15 | 5.62E-03 |
| GO:0015293 | symporter activity | MF | 107 | 15 | 2 | 5.65E-03 |
| GO:0007586 | digestion | BP | 97 | 14 | 4 | 5.68E-03 |
| GO:0051216 | cartilage development | BP | 149 | 19 | 9 | 5.76E-03 |
| GO:0044305 | calyx of Held | CC | 18 | 5 | 3 | 5.79E-03 |
| GO:0098644 | complex of collagen trimers | CC | 18 | 5 | 0 | 5.79E-03 |
| GO:1901881 | positive regulation of protein depolymerization | BP | 18 | 5 | 1 | 5.79E-03 |
| GO:0048545 | response to steroid hormone | BP | 307 | 33 | 15 | 5.82E-03 |
| GO:0010828 | positive regulation of glucose transmembrane transport | BP | 33 | 7 | 1 | 5.83E-03 |
| GO:0014074 | response to purine-containing compound | BP | 128 | 17 | 12 | 5.83E-03 |
| GO:0003007 | heart morphogenesis | BP | 204 | 24 | 3 | 5.88E-03 |
| GO:0016328 | lateral plasma membrane | CC | 50 | 9 | 3 | 5.90E-03 |
| GO:1901616 | organic hydroxy compound catabolic process | BP | 50 | 9 | 2 | 5.90E-03 |
| GO:0010827 | regulation of glucose transmembrane transport | BP | 59 | 10 | 2 | 5.92E-03 |
| GO:0090288 | negative regulation of cellular response to growth factor stimulus | BP | 118 | 16 | 6 | 6.02E-03 |
| GO:0060401 | cytosolic calcium ion transport | BP | 139 | 18 | 14 | 6.04E-03 |
| GO:0030336 | negative regulation of cell migration | BP | 216 | 25 | 11 | 6.16E-03 |
| GO:0007015 | actin filament organization | BP | 344 | 36 | 27 | 6.26E-03 |
| GO:0010876 | lipid localization | BP | 297 | 32 | 16 | 6.32E-03 |
| GO:0006810 | transport | BP | 3967 | 303 | 229 | 6.34E-03 |
| GO:0034637 | cellular carbohydrate biosynthetic process | BP | 69 | 11 | 0 | 6.42E-03 |
| GO:0090102 | cochlea development | BP | 42 | 8 | 2 | 6.52E-03 |
| GO:0007218 | neuropeptide signaling pathway | BP | 42 | 8 | 0 | 6.52E-03 |
| GO:0044273 | sulfur compound catabolic process | BP | 42 | 8 | 2 | 6.52E-03 |
| GO:0006949 | syncytium formation | BP | 42 | 8 | 5 | 6.52E-03 |
| GO:0007166 | cell surface receptor signaling pathway | BP | 2269 | 182 | 201 | 6.53E-03 |
| GO:0051494 | negative regulation of cytoskeleton organization | BP | 119 | 16 | 8 | 6.53E-03 |

| | | | | | | |
|---|---|---|---|---|---|---|
| GO:0031099 | regeneration | BP | 151 | 19 | 6 | 6.65E-03 |
| GO:0005518 | collagen binding | MF | 60 | 10 | 2 | 6.68E-03 |
| GO:0046943 | carboxylic acid transmembrane transporter activity | MF | 109 | 15 | 5 | 6.71E-03 |
| GO:0072073 | kidney epithelium development | BP | 109 | 15 | 9 | 6.71E-03 |
| GO:0005342 | organic acid transmembrane transporter activity | MF | 109 | 15 | 5 | 6.71E-03 |
| GO:0021681 | cerebellar granular layer development | BP | 12 | 4 | 1 | 6.71E-03 |
| GO:0038065 | collagen-activated signaling pathway | BP | 12 | 4 | 0 | 6.71E-03 |
| GO:0099243 | extrinsic component of synaptic membrane | CC | 12 | 4 | 0 | 6.71E-03 |
| GO:1902306 | negative regulation of sodium ion transmembrane transport | BP | 12 | 4 | 1 | 6.71E-03 |
| GO:0072160 | nephron tubule epithelial cell differentiation | BP | 12 | 4 | 2 | 6.71E-03 |
| GO:0051428 | peptide hormone receptor binding | MF | 12 | 4 | 2 | 6.71E-03 |
| GO:0042749 | regulation of circadian sleep/wake cycle | BP | 12 | 4 | 1 | 6.71E-03 |
| GO:1902285 | semaphorin-plexin signaling pathway involved in neuron projection | BP | 12 | 4 | 1 | 6.71E-03 |
| GO:0030240 | skeletal muscle thin filament assembly | BP | 12 | 4 | 0 | 6.71E-03 |
| GO:0055008 | cardiac muscle tissue morphogenesis | BP | 51 | 9 | 1 | 6.75E-03 |
| GO:0043113 | receptor clustering | BP | 51 | 9 | 7 | 6.75E-03 |
| GO:0007584 | response to nutrient | BP | 184 | 22 | 11 | 6.75E-03 |
| GO:0010906 | regulation of glucose metabolic process | BP | 99 | 14 | 1 | 6.81E-03 |
| GO:0060359 | response to ammonium ion | BP | 99 | 14 | 4 | 6.81E-03 |
| GO:0046717 | acid secretion | BP | 89 | 13 | 10 | 6.82E-03 |
| GO:0046903 | secretion | BP | 1294 | 110 | 109 | 6.95E-03 |
| GO:0071773 | cellular response to BMP stimulus | BP | 120 | 16 | 3 | 7.07E-03 |
| GO:0071772 | response to BMP | BP | 120 | 16 | 3 | 7.07E-03 |
| GO:0044087 | regulation of cellular component biogenesis | BP | 822 | 74 | 50 | 7.10E-03 |
| GO:0045926 | negative regulation of growth | BP | 196 | 23 | 9 | 7.12E-03 |
| GO:0048568 | embryonic organ development | BP | 335 | 35 | 15 | 7.14E-03 |
| GO:0034329 | cell junction assembly | BP | 219 | 25 | 5 | 7.32E-03 |
| GO:0048522 | positive regulation of cellular process | BP | 4206 | 319 | 297 | 7.36E-03 |
| GO:0072202 | cell differentiation involved in metanephros development | BP | 19 | 5 | 0 | 7.43E-03 |
| GO:0043153 | entrainment of circadian clock by photoperiod | BP | 19 | 5 | 0 | 7.43E-03 |
| GO:0045932 | negative regulation of muscle contraction | BP | 19 | 5 | 1 | 7.43E-03 |
| GO:1903055 | positive regulation of extracellular matrix organization | BP | 19 | 5 | 2 | 7.43E-03 |
| GO:0099625 | ventricular cardiac muscle cell membrane repolarization | BP | 19 | 5 | 1 | 7.43E-03 |
| GO:0006629 | lipid metabolic process | BP | 1098 | 95 | 55 | 7.45E-03 |
| GO:0042752 | regulation of circadian rhythm | BP | 90 | 13 | 3 | 7.49E-03 |
| GO:0008344 | adult locomotory behavior | BP | 61 | 10 | 3 | 7.52E-03 |
| GO:0050919 | negative chemotaxis | BP | 43 | 8 | 3 | 7.54E-03 |
| GO:1903307 | positive regulation of regulated secretory pathway | BP | 43 | 8 | 6 | 7.54E-03 |
| GO:0046324 | regulation of glucose import | BP | 43 | 8 | 0 | 7.54E-03 |
| GO:0007266 | Rho protein signal transduction | BP | 186 | 22 | 14 | 7.65E-03 |
| GO:0030902 | hindbrain development | BP | 121 | 16 | 5 | 7.65E-03 |
| GO:0004497 | monooxygenase activity | MF | 52 | 9 | 3 | 7.68E-03 |
| GO:0005581 | collagen trimer | CC | 71 | 11 | 1 | 7.97E-03 |
| GO:0071560 | cellular response to transforming growth factor beta stimulus | BP | 198 | 23 | 10 | 8.03E-03 |
| GO:0032535 | regulation of cellular component size | BP | 314 | 33 | 22 | 8.10E-03 |
| GO:1903053 | regulation of extracellular matrix organization | BP | 35 | 7 | 3 | 8.15E-03 |
| GO:0010243 | response to organonitrogen compound | BP | 814 | 73 | 56 | 8.16E-03 |
| GO:0001558 | regulation of cell growth | BP | 338 | 35 | 13 | 8.16E-03 |
| GO:0014902 | myotube differentiation | BP | 81 | 12 | 6 | 8.18E-03 |
| GO:0099622 | cardiac muscle cell membrane repolarization | BP | 27 | 6 | 1 | 8.29E-03 |
| GO:0046875 | ephrin receptor binding | MF | 27 | 6 | 5 | 8.29E-03 |
| GO:0070050 | neuron cellular homeostasis | BP | 27 | 6 | 2 | 8.29E-03 |
| GO:1901699 | cellular response to nitrogen compound | BP | 522 | 50 | 38 | 8.30E-03 |
| GO:0032272 | negative regulation of protein polymerization | BP | 62 | 10 | 3 | 8.43E-03 |
| GO:0048678 | response to axon injury | BP | 62 | 10 | 3 | 8.43E-03 |
| GO:0061351 | neural precursor cell proliferation | BP | 112 | 15 | 8 | 8.59E-03 |
| GO:0045178 | basal part of cell | CC | 44 | 8 | 2 | 8.69E-03 |
| GO:0031526 | brush border membrane | CC | 44 | 8 | 1 | 8.69E-03 |
| GO:0061005 | cell differentiation involved in kidney development | BP | 44 | 8 | 4 | 8.69E-03 |
| GO:0006576 | cellular biogenic amine metabolic process | BP | 44 | 8 | 2 | 8.69E-03 |
| GO:0007193 | adenylate cyclase-inhibiting G protein-coupled receptor signaling p | BP | 53 | 9 | 2 | 8.71E-03 |
| GO:0019694 | alkanesulfonate metabolic process | BP | 7 | 3 | 1 | 8.86E-03 |
| GO:0003360 | brainstem development | BP | 7 | 3 | 0 | 8.86E-03 |
| GO:0098645 | collagen network | CC | 7 | 3 | 0 | 8.86E-03 |

| | | | | | | |
|---|---|---|---|---|---|---|
| GO:0098596 | imitative learning | BP | 7 | 3 | 0 | 8.86E-03 |
| GO:0060073 | micturition | BP | 7 | 3 | 0 | 8.86E-03 |
| GO:0048671 | negative regulation of collateral sprouting | BP | 7 | 3 | 0 | 8.86E-03 |
| GO:0098642 | network-forming collagen trimer | CC | 7 | 3 | 0 | 8.86E-03 |
| GO:0015851 | nucleobase transport | BP | 7 | 3 | 0 | 8.86E-03 |
| GO:0070444 | oligodendrocyte progenitor proliferation | BP | 7 | 3 | 1 | 8.86E-03 |
| GO:0031628 | opioid receptor binding | MF | 7 | 3 | 0 | 8.86E-03 |
| GO:0015166 | polyol transmembrane transporter activity | MF | 7 | 3 | 0 | 8.86E-03 |
| GO:2001046 | positive regulation of integrin-mediated signaling pathway | BP | 7 | 3 | 0 | 8.86E-03 |
| GO:0099569 | presynaptic cytoskeleton | CC | 7 | 3 | 0 | 8.86E-03 |
| GO:0034310 | primary alcohol catabolic process | BP | 7 | 3 | 0 | 8.86E-03 |
| GO:0060046 | regulation of acrosome reaction | BP | 7 | 3 | 0 | 8.86E-03 |
| GO:0032276 | regulation of gonadotropin secretion | BP | 7 | 3 | 0 | 8.86E-03 |
| GO:1904478 | regulation of intestinal absorption | BP | 7 | 3 | 0 | 8.86E-03 |
| GO:0070445 | regulation of oligodendrocyte progenitor proliferation | BP | 7 | 3 | 1 | 8.86E-03 |
| GO:0014819 | regulation of skeletal muscle contraction | BP | 7 | 3 | 0 | 8.86E-03 |
| GO:0048495 | Roundabout binding | MF | 7 | 3 | 0 | 8.86E-03 |
| GO:0014731 | spectrin-associated cytoskeleton | CC | 7 | 3 | 1 | 8.86E-03 |
| GO:0005212 | structural constituent of eye lens | MF | 7 | 3 | 0 | 8.86E-03 |
| GO:0019530 | taurine metabolic process | BP | 7 | 3 | 1 | 8.86E-03 |
| GO:0099542 | trans-synaptic signaling by endocannabinoid | BP | 7 | 3 | 0 | 8.86E-03 |
| GO:0099541 | trans-synaptic signaling by lipid | BP | 7 | 3 | 0 | 8.86E-03 |
| GO:0070471 | uterine smooth muscle contraction | BP | 7 | 3 | 0 | 8.86E-03 |
| GO:0042297 | vocal learning | BP | 7 | 3 | 0 | 8.86E-03 |
| GO:0099626 | voltage-gated calcium channel activity involved in regulation of pre | MF | 7 | 3 | 1 | 8.86E-03 |
| GO:0051047 | positive regulation of secretion | BP | 316 | 33 | 27 | 8.88E-03 |
| GO:0005976 | polysaccharide metabolic process | BP | 92 | 13 | 1 | 8.99E-03 |
| GO:0051261 | protein depolymerization | BP | 92 | 13 | 3 | 8.99E-03 |
| GO:0120162 | positive regulation of cold-induced thermogenesis | BP | 82 | 12 | 6 | 9.01E-03 |
| GO:1901615 | organic hydroxy compound metabolic process | BP | 413 | 41 | 31 | 9.02E-03 |
| GO:0048598 | embryonic morphogenesis | BP | 450 | 44 | 20 | 9.07E-03 |
| GO:0110053 | regulation of actin filament organization | BP | 223 | 25 | 16 | 9.14E-03 |
| GO:0021535 | cell migration in hindbrain | BP | 13 | 4 | 0 | 9.18E-03 |
| GO:0016010 | dystrophin-associated glycoprotein complex | CC | 13 | 4 | 0 | 9.18E-03 |
| GO:0090665 | glycoprotein complex | CC | 13 | 4 | 0 | 9.18E-03 |
| GO:0007638 | mechanosensory behavior | BP | 13 | 4 | 0 | 9.18E-03 |
| GO:0072234 | metanephric nephron tubule development | BP | 13 | 4 | 0 | 9.18E-03 |
| GO:0072176 | nephric duct development | BP | 13 | 4 | 1 | 9.18E-03 |
| GO:2001044 | regulation of integrin-mediated signaling pathway | BP | 13 | 4 | 0 | 9.18E-03 |
| GO:1905063 | regulation of vascular smooth muscle cell differentiation | BP | 13 | 4 | 2 | 9.18E-03 |
| GO:0043586 | tongue development | BP | 13 | 4 | 1 | 9.18E-03 |
| GO:0055003 | cardiac myofibril assembly | BP | 20 | 5 | 0 | 9.36E-03 |
| GO:0048668 | collateral sprouting | BP | 20 | 5 | 2 | 9.36E-03 |
| GO:1903792 | negative regulation of anion transport | BP | 20 | 5 | 0 | 9.36E-03 |
| GO:1902307 | positive regulation of sodium ion transmembrane transport | BP | 20 | 5 | 1 | 9.36E-03 |
| GO:0010996 | response to auditory stimulus | BP | 20 | 5 | 0 | 9.36E-03 |
| GO:0005501 | retinoid binding | MF | 20 | 5 | 0 | 9.36E-03 |
| GO:0048511 | rhythmic process | BP | 235 | 26 | 11 | 9.39E-03 |
| GO:0035850 | epithelial cell differentiation involved in kidney development | BP | 36 | 7 | 4 | 9.54E-03 |
| GO:0010463 | mesenchymal cell proliferation | BP | 36 | 7 | 1 | 9.54E-03 |
| GO:0086004 | regulation of cardiac muscle cell contraction | BP | 36 | 7 | 3 | 9.54E-03 |
| GO:0033218 | amide binding | MF | 247 | 27 | 17 | 9.59E-03 |
| GO:0009611 | response to wounding | BP | 514 | 49 | 35 | 9.71E-03 |
| GO:0060135 | maternal process involved in female pregnancy | BP | 54 | 9 | 2 | 9.84E-03 |
| GO:0035418 | protein localization to synapse | BP | 54 | 9 | 1 | 9.84E-03 |
| GO:0048018 | receptor ligand activity | MF | 236 | 26 | 23 | 9.89E-03 |
| GO:0071385 | cellular response to glucocorticoid stimulus | BP | 45 | 8 | 2 | 9.95E-03 |
| GO:0048016 | inositol phosphate-mediated signaling | BP | 45 | 8 | 1 | 9.95E-03 |
| GO:2000243 | positive regulation of reproductive process | BP | 45 | 8 | 2 | 9.95E-03 |
| GO:0030834 | regulation of actin filament depolymerization | BP | 45 | 8 | 1 | 9.95E-03 |
| GO:0046326 | positive regulation of glucose import | BP | 28 | 6 | 0 | 9.96E-03 |
| GO:0060043 | regulation of cardiac muscle cell proliferation | BP | 28 | 6 | 2 | 9.96E-03 |
| GO:0046683 | response to organophosphorus | BP | 114 | 15 | 10 | 1.01E-02 |
| GO:0050662 | coenzyme binding | MF | 248 | 27 | 10 | 1.01E-02 |

| | | | | | | |
|---|---|---|---|---|---|---|
| GO:0048518 | positive regulation of biological process | BP | 4721 | 353 | 340 | 1.05E-02 |
| GO:0032870 | cellular response to hormone stimulus | BP | 554 | 52 | 34 | 1.05E-02 |
| GO:0046677 | response to antibiotic | BP | 261 | 28 | 20 | 1.08E-02 |
| GO:0003300 | cardiac muscle hypertrophy | BP | 74 | 11 | 0 | 1.08E-02 |
| GO:0033273 | response to vitamin | BP | 74 | 11 | 4 | 1.08E-02 |
| GO:0010522 | regulation of calcium ion transport into cytosol | BP | 84 | 12 | 8 | 1.09E-02 |
| GO:1901652 | response to peptide | BP | 418 | 41 | 32 | 1.09E-02 |
| GO:1903532 | positive regulation of secretion by cell | BP | 297 | 31 | 27 | 1.10E-02 |
| GO:0010812 | negative regulation of cell-substrate adhesion | BP | 55 | 9 | 4 | 1.11E-02 |
| GO:0030133 | transport vesicle | CC | 321 | 33 | 10 | 1.11E-02 |
| GO:0060042 | retina morphogenesis in camera-type eye | BP | 37 | 7 | 1 | 1.11E-02 |
| GO:0071526 | semaphorin-plexin signaling pathway | BP | 37 | 7 | 3 | 1.11E-02 |
| GO:0071407 | cellular response to organic cyclic compound | BP | 456 | 44 | 26 | 1.13E-02 |
| GO:0031253 | cell projection membrane | CC | 262 | 28 | 14 | 1.13E-02 |
| GO:0030863 | cortical cytoskeleton | CC | 105 | 14 | 9 | 1.13E-02 |
| GO:2000146 | negative regulation of cell motility | BP | 227 | 25 | 13 | 1.13E-02 |
| GO:0071559 | response to transforming growth factor beta | BP | 204 | 23 | 11 | 1.13E-02 |
| GO:2000649 | regulation of sodium ion transmembrane transporter activity | BP | 46 | 8 | 2 | 1.14E-02 |
| GO:0008227 | G protein-coupled amine receptor activity | MF | 21 | 5 | 2 | 1.16E-02 |
| GO:0019840 | isoprenoid binding | MF | 21 | 5 | 0 | 1.16E-02 |
| GO:0098810 | neurotransmitter reuptake | BP | 21 | 5 | 0 | 1.16E-02 |
| GO:0009648 | photoperiodism | BP | 21 | 5 | 0 | 1.16E-02 |
| GO:0060343 | trabecula formation | BP | 21 | 5 | 1 | 1.16E-02 |
| GO:0055123 | digestive system development | BP | 116 | 15 | 3 | 1.17E-02 |
| GO:0048660 | regulation of smooth muscle cell proliferation | BP | 116 | 15 | 8 | 1.17E-02 |
| GO:0008285 | negative regulation of cell proliferation | BP | 545 | 51 | 34 | 1.18E-02 |
| GO:0010613 | positive regulation of cardiac muscle hypertrophy | BP | 29 | 6 | 0 | 1.18E-02 |
| GO:0014742 | positive regulation of muscle hypertrophy | BP | 29 | 6 | 0 | 1.18E-02 |
| GO:0048713 | regulation of oligodendrocyte differentiation | BP | 29 | 6 | 0 | 1.18E-02 |
| GO:0051150 | regulation of smooth muscle cell differentiation | BP | 29 | 6 | 3 | 1.18E-02 |
| GO:0007189 | adenylate cyclase-activating G protein-coupled receptor signaling p | BP | 85 | 12 | 8 | 1.19E-02 |
| GO:0099106 | ion channel regulator activity | MF | 85 | 12 | 4 | 1.19E-02 |
| GO:0014897 | striated muscle hypertrophy | BP | 75 | 11 | 0 | 1.19E-02 |
| GO:0001503 | ossification | BP | 299 | 31 | 23 | 1.21E-02 |
| GO:0048791 | calcium ion-regulated exocytosis of neurotransmitter | BP | 14 | 4 | 0 | 1.22E-02 |
| GO:0006067 | ethanol metabolic process | BP | 14 | 4 | 0 | 1.22E-02 |
| GO:0072170 | metanephric tubule development | BP | 14 | 4 | 0 | 1.22E-02 |
| GO:0032891 | negative regulation of organic acid transport | BP | 14 | 4 | 0 | 1.22E-02 |
| GO:0006910 | phagocytosis, recognition | BP | 14 | 4 | 0 | 1.22E-02 |
| GO:0098911 | regulation of ventricular cardiac muscle cell action potential | BP | 14 | 4 | 1 | 1.22E-02 |
| GO:0046459 | short-chain fatty acid metabolic process | BP | 14 | 4 | 0 | 1.22E-02 |
| GO:0001222 | transcription corepressor binding | MF | 14 | 4 | 0 | 1.22E-02 |
| GO:1905039 | carboxylic acid transmembrane transport | BP | 106 | 14 | 3 | 1.22E-02 |
| GO:0016247 | channel regulator activity | MF | 106 | 14 | 4 | 1.22E-02 |
| GO:1903825 | organic acid transmembrane transport | BP | 106 | 14 | 3 | 1.22E-02 |
| GO:0045444 | fat cell differentiation | BP | 183 | 21 | 11 | 1.25E-02 |
| GO:0046395 | carboxylic acid catabolic process | BP | 229 | 25 | 9 | 1.26E-02 |
| GO:0016054 | organic acid catabolic process | BP | 229 | 25 | 9 | 1.26E-02 |
| GO:0048659 | smooth muscle cell proliferation | BP | 117 | 15 | 8 | 1.26E-02 |
| GO:0008542 | visual learning | BP | 38 | 7 | 3 | 1.28E-02 |
| GO:0071384 | cellular response to corticosteroid stimulus | BP | 47 | 8 | 2 | 1.29E-02 |
| GO:1900449 | regulation of glutamate receptor signaling pathway | BP | 47 | 8 | 4 | 1.29E-02 |
| GO:0060485 | mesenchyme development | BP | 218 | 24 | 8 | 1.29E-02 |
| GO:1905114 | cell surface receptor signaling pathway involved in cell-cell signali | BP | 510 | 48 | 26 | 1.30E-02 |
| GO:0007368 | determination of left/right symmetry | BP | 86 | 12 | 1 | 1.30E-02 |
| GO:0099501 | exocytic vesicle membrane | CC | 86 | 12 | 4 | 1.30E-02 |
| GO:0030672 | synaptic vesicle membrane | CC | 86 | 12 | 4 | 1.30E-02 |
| GO:0006869 | lipid transport | BP | 265 | 28 | 14 | 1.31E-02 |
| GO:0014896 | muscle hypertrophy | BP | 76 | 11 | 0 | 1.31E-02 |
| GO:0045921 | positive regulation of exocytosis | BP | 76 | 11 | 7 | 1.31E-02 |
| GO:0003987 | acetate-CoA ligase activity | MF | 3 | 2 | 0 | 1.32E-02 |
| GO:0014055 | acetylcholine secretion, neurotransmission | BP | 3 | 2 | 1 | 1.32E-02 |
| GO:0007195 | adenylate cyclase-inhibiting dopamine receptor signaling pathway | BP | 3 | 2 | 0 | 1.32E-02 |
| GO:0007196 | adenylate cyclase-inhibiting G protein-coupled glutamate receptor s | BP | 3 | 2 | 0 | 1.32E-02 |

| | | | | | | |
|---|---|---|---|---|---|---|
| GO:0004936 | alpha-adrenergic receptor activity | MF | 3 | 2 | 0 | 1.32E-02 |
| GO:0098985 | asymmetric, glutamatergic, excitatory synapse | CC | 3 | 2 | 0 | 1.32E-02 |
| GO:0072338 | cellular lactam metabolic process | BP | 3 | 2 | 0 | 1.32E-02 |
| GO:0021824 | cerebral cortex tangential migration using cell-axon interactions | BP | 3 | 2 | 0 | 1.32E-02 |
| GO:0097167 | circadian regulation of translation | BP | 3 | 2 | 0 | 1.32E-02 |
| GO:0008410 | CoA-transferase activity | MF | 3 | 2 | 0 | 1.32E-02 |
| GO:0009093 | cysteine catabolic process | BP | 3 | 2 | 0 | 1.32E-02 |
| GO:1901588 | dendritic microtubule | CC | 3 | 2 | 0 | 1.32E-02 |
| GO:0036023 | embryonic skeletal limb joint morphogenesis | BP | 3 | 2 | 0 | 1.32E-02 |
| GO:0090500 | endocardial cushion to mesenchymal transition | BP | 3 | 2 | 0 | 1.32E-02 |
| GO:1903375 | facioacoustic ganglion development | BP | 3 | 2 | 0 | 1.32E-02 |
| GO:0097116 | gephyrin clustering involved in postsynaptic density assembly | BP | 3 | 2 | 0 | 1.32E-02 |
| GO:0015254 | glycerol channel activity | MF | 3 | 2 | 0 | 1.32E-02 |
| GO:0008184 | glycogen phosphorylase activity | MF | 3 | 2 | 0 | 1.32E-02 |
| GO:0021828 | gonadotrophin-releasing hormone neuronal migration to the hypoth | BP | 3 | 2 | 0 | 1.32E-02 |
| GO:0017095 | heparan sulfate 6-O-sulfotransferase activity | MF | 3 | 2 | 0 | 1.32E-02 |
| GO:0097457 | hippocampal mossy fiber | CC | 3 | 2 | 0 | 1.32E-02 |
| GO:0021856 | hypothalamic tangential migration using cell-axon interactions | BP | 3 | 2 | 0 | 1.32E-02 |
| GO:0021855 | hypothalamus cell migration | BP | 3 | 2 | 0 | 1.32E-02 |
| GO:0005237 | inhibitory extracellular ligand-gated ion channel activity | MF | 3 | 2 | 0 | 1.32E-02 |
| GO:0019448 | L-cysteine catabolic process | BP | 3 | 2 | 0 | 1.32E-02 |
| GO:0036022 | limb joint morphogenesis | BP | 3 | 2 | 0 | 1.32E-02 |
| GO:0102250 | linear malto-oligosaccharide phosphorylase activity | MF | 3 | 2 | 0 | 1.32E-02 |
| GO:0098912 | membrane depolarization during atrial cardiac muscle cell action p | BP | 3 | 2 | 0 | 1.32E-02 |
| GO:0086048 | membrane depolarization during bundle of His cell action potential | BP | 3 | 2 | 0 | 1.32E-02 |
| GO:0072180 | mesonephric duct morphogenesis | BP | 3 | 2 | 0 | 1.32E-02 |
| GO:0031852 | mu-type opioid receptor binding | MF | 3 | 2 | 0 | 1.32E-02 |
| GO:0031523 | Myb complex | CC | 3 | 2 | 0 | 1.32E-02 |
| GO:0070995 | NADPH oxidation | BP | 3 | 2 | 1 | 1.32E-02 |
| GO:0016941 | natriuretic peptide receptor activity | MF | 3 | 2 | 0 | 1.32E-02 |
| GO:1904479 | negative regulation of intestinal absorption | BP | 3 | 2 | 0 | 1.32E-02 |
| GO:0051387 | negative regulation of neurotrophin TRK receptor signaling pathwa | BP | 3 | 2 | 0 | 1.32E-02 |
| GO:0044245 | polysaccharide digestion | BP | 3 | 2 | 0 | 1.32E-02 |
| GO:2000851 | positive regulation of glucocorticoid secretion | BP | 3 | 2 | 0 | 1.32E-02 |
| GO:0005345 | purine nucleobase transmembrane transporter activity | MF | 3 | 2 | 0 | 1.32E-02 |
| GO:0014056 | regulation of acetylcholine secretion, neurotransmission | BP | 3 | 2 | 1 | 1.32E-02 |
| GO:2000722 | regulation of cardiac vascular smooth muscle cell differentiation | BP | 3 | 2 | 0 | 1.32E-02 |
| GO:0001560 | regulation of cell growth by extracellular stimulus | BP | 3 | 2 | 0 | 1.32E-02 |
| GO:0045113 | regulation of integrin biosynthetic process | BP | 3 | 2 | 0 | 1.32E-02 |
| GO:0014722 | regulation of skeletal muscle contraction by calcium ion signaling | BP | 3 | 2 | 0 | 1.32E-02 |
| GO:0106020 | regulation of vesicle docking | BP | 3 | 2 | 0 | 1.32E-02 |
| GO:0102499 | SHG alpha-glucan phosphorylase activity | MF | 3 | 2 | 0 | 1.32E-02 |
| GO:0060084 | synaptic transmission involved in micturition | BP | 3 | 2 | 0 | 1.32E-02 |
| GO:0071918 | urea transmembrane transport | BP | 3 | 2 | 0 | 1.32E-02 |
| GO:0015204 | urea transmembrane transporter activity | MF | 3 | 2 | 0 | 1.32E-02 |
| GO:0015840 | urea transport | BP | 3 | 2 | 0 | 1.32E-02 |
| GO:0060157 | urinary bladder development | BP | 3 | 2 | 0 | 1.32E-02 |
| GO:0086056 | voltage-gated calcium channel activity involved in AV node cell ac | MF | 3 | 2 | 0 | 1.32E-02 |
| GO:0043168 | anion binding | MF | 2241 | 177 | 128 | 1.32E-02 |
| GO:0099026 | anchored component of presynaptic membrane | CC | 8 | 3 | 1 | 1.35E-02 |
| GO:0098651 | basement membrane collagen trimer | CC | 8 | 3 | 0 | 1.35E-02 |
| GO:0071313 | cellular response to caffeine | BP | 8 | 3 | 0 | 1.35E-02 |
| GO:0021683 | cerebellar granular layer morphogenesis | BP | 8 | 3 | 1 | 1.35E-02 |
| GO:0098981 | cholinergic synapse | CC | 8 | 3 | 0 | 1.35E-02 |
| GO:0102391 | decanoate-CoA ligase activity | MF | 8 | 3 | 0 | 1.35E-02 |
| GO:0021612 | facial nerve structural organization | BP | 8 | 3 | 1 | 1.35E-02 |
| GO:0042577 | lipid phosphatase activity | MF | 8 | 3 | 0 | 1.35E-02 |
| GO:0044539 | long-chain fatty acid import | BP | 8 | 3 | 0 | 1.35E-02 |
| GO:0031956 | medium-chain fatty acid-CoA ligase activity | MF | 8 | 3 | 0 | 1.35E-02 |
| GO:0098597 | observational learning | BP | 8 | 3 | 0 | 1.35E-02 |
| GO:0060452 | positive regulation of cardiac muscle contraction | BP | 8 | 3 | 1 | 1.35E-02 |
| GO:0008556 | potassium-transporting ATPase activity | MF | 8 | 3 | 1 | 1.35E-02 |
| GO:1990709 | presynaptic active zone organization | BP | 8 | 3 | 0 | 1.35E-02 |
| GO:1902514 | regulation of calcium ion transmembrane transport via high voltage | BP | 8 | 3 | 0 | 1.35E-02 |

| | | | | | | |
|---|---|---|---|---|---|---|
| GO:0098962 | regulation of postsynaptic neurotransmitter receptor activity | BP | 8 | 3 | 0 | 1.35E-02 |
| GO:0014894 | response to denervation involved in regulation of muscle adaptation | BP | 8 | 3 | 0 | 1.35E-02 |
| GO:0009750 | response to fructose | BP | 8 | 3 | 0 | 1.35E-02 |
| GO:0014877 | response to muscle inactivity involved in regulation of muscle adap | BP | 8 | 3 | 0 | 1.35E-02 |
| GO:0005391 | sodium:potassium-exchanging ATPase activity | MF | 8 | 3 | 1 | 1.35E-02 |
| GO:0061549 | sympathetic ganglion development | BP | 8 | 3 | 0 | 1.35E-02 |
| GO:0099545 | trans-synaptic signaling by trans-synaptic complex | BP | 8 | 3 | 1 | 1.35E-02 |
| GO:0099511 | voltage-gated calcium channel activity involved in regulation of cyt | MF | 8 | 3 | 1 | 1.35E-02 |
| GO:1903169 | regulation of calcium ion transmembrane transport | BP | 118 | 15 | 11 | 1.36E-02 |
| GO:0051270 | regulation of cellular component movement | BP | 820 | 72 | 57 | 1.37E-02 |
| GO:0043244 | regulation of protein complex disassembly | BP | 97 | 13 | 1 | 1.38E-02 |
| GO:0034332 | adherens junction organization | BP | 129 | 16 | 2 | 1.38E-02 |
| GO:0051234 | establishment of localization | BP | 4066 | 306 | 237 | 1.38E-02 |
| GO:0033692 | cellular polysaccharide biosynthetic process | BP | 57 | 9 | 0 | 1.39E-02 |
| GO:0004714 | transmembrane receptor protein tyrosine kinase activity | MF | 57 | 9 | 1 | 1.39E-02 |
| GO:0060421 | positive regulation of heart growth | BP | 30 | 6 | 1 | 1.40E-02 |
| GO:0044262 | cellular carbohydrate metabolic process | BP | 243 | 26 | 5 | 1.41E-02 |
| GO:0061037 | negative regulation of cartilage development | BP | 22 | 5 | 1 | 1.42E-02 |
| GO:0017075 | syntaxin-1 binding | MF | 22 | 5 | 0 | 1.42E-02 |
| GO:0003073 | regulation of systemic arterial blood pressure | BP | 67 | 10 | 2 | 1.44E-02 |
| GO:0032940 | secretion by cell | BP | 1193 | 100 | 103 | 1.44E-02 |
| GO:0021537 | telencephalon development | BP | 197 | 22 | 11 | 1.45E-02 |
| GO:0016459 | myosin complex | CC | 48 | 8 | 2 | 1.46E-02 |
| GO:0035914 | skeletal muscle cell differentiation | BP | 48 | 8 | 5 | 1.46E-02 |
| GO:0071417 | cellular response to organonitrogen compound | BP | 476 | 45 | 37 | 1.47E-02 |
| GO:0097720 | calcineurin-mediated signaling | BP | 39 | 7 | 1 | 1.47E-02 |
| GO:1903115 | regulation of actin filament-based movement | BP | 39 | 7 | 3 | 1.47E-02 |
| GO:1901701 | cellular response to oxygen-containing compound | BP | 903 | 78 | 70 | 1.52E-02 |
| GO:0061448 | connective tissue development | BP | 198 | 22 | 9 | 1.53E-02 |
| GO:0006103 | 2-oxoglutarate metabolic process | BP | 15 | 4 | 0 | 1.57E-02 |
| GO:0048532 | anatomical structure arrangement | BP | 15 | 4 | 3 | 1.57E-02 |
| GO:0042474 | middle ear morphogenesis | BP | 15 | 4 | 0 | 1.57E-02 |
| GO:0021783 | preganglionic parasympathetic fiber development | BP | 15 | 4 | 2 | 1.57E-02 |
| GO:2000696 | regulation of epithelial cell differentiation involved in kidney devel | BP | 15 | 4 | 3 | 1.57E-02 |
| GO:0099509 | regulation of presynaptic cytosolic calcium ion concentration | BP | 15 | 4 | 1 | 1.57E-02 |
| GO:0005248 | voltage-gated sodium channel activity | MF | 15 | 4 | 0 | 1.57E-02 |
| GO:0042136 | neurotransmitter biosynthetic process | BP | 78 | 11 | 8 | 1.58E-02 |
| GO:1903035 | negative regulation of response to wounding | BP | 68 | 10 | 1 | 1.58E-02 |
| GO:0051279 | regulation of release of sequestered calcium ion into cytosol | BP | 68 | 10 | 5 | 1.58E-02 |
| GO:0050768 | negative regulation of neurogenesis | BP | 222 | 24 | 12 | 1.59E-02 |
| GO:0009925 | basal plasma membrane | CC | 31 | 6 | 2 | 1.64E-02 |
| GO:0010665 | regulation of cardiac muscle cell apoptotic process | BP | 31 | 6 | 1 | 1.64E-02 |
| GO:0090313 | regulation of protein targeting to membrane | BP | 31 | 6 | 3 | 1.64E-02 |
| GO:0005938 | cell cortex | CC | 258 | 27 | 16 | 1.64E-02 |
| GO:0030837 | negative regulation of actin filament polymerization | BP | 49 | 8 | 2 | 1.64E-02 |
| GO:0045669 | positive regulation of osteoblast differentiation | BP | 49 | 8 | 3 | 1.64E-02 |
| GO:0060395 | SMAD protein signal transduction | BP | 49 | 8 | 3 | 1.64E-02 |
| GO:0030055 | cell-substrate junction | CC | 392 | 38 | 29 | 1.65E-02 |
| GO:0033993 | response to lipid | BP | 723 | 64 | 55 | 1.66E-02 |
| GO:1901654 | response to ketone | BP | 154 | 18 | 9 | 1.67E-02 |
| GO:0048706 | embryonic skeletal system development | BP | 89 | 12 | 4 | 1.68E-02 |
| GO:0008201 | heparin binding | MF | 121 | 15 | 4 | 1.68E-02 |
| GO:0099738 | cell cortex region | CC | 40 | 7 | 0 | 1.69E-02 |
| GO:0098563 | intrinsic component of synaptic vesicle membrane | CC | 40 | 7 | 1 | 1.69E-02 |
| GO:0015291 | secondary active transmembrane transporter activity | MF | 177 | 20 | 3 | 1.70E-02 |
| GO:0071295 | cellular response to vitamin | BP | 23 | 5 | 1 | 1.72E-02 |
| GO:0019934 | cGMP-mediated signaling | BP | 23 | 5 | 0 | 1.72E-02 |
| GO:0140115 | export across plasma membrane | BP | 23 | 5 | 2 | 1.72E-02 |
| GO:0005217 | intracellular ligand-gated ion channel activity | MF | 23 | 5 | 2 | 1.72E-02 |
| GO:0099604 | ligand-gated calcium channel activity | MF | 23 | 5 | 1 | 1.72E-02 |
| GO:0051968 | positive regulation of synaptic transmission, glutamatergic | BP | 23 | 5 | 2 | 1.72E-02 |
| GO:0010962 | regulation of glucan biosynthetic process | BP | 23 | 5 | 0 | 1.72E-02 |
| GO:0005979 | regulation of glycogen biosynthetic process | BP | 23 | 5 | 0 | 1.72E-02 |
| GO:0014829 | vascular smooth muscle contraction | BP | 23 | 5 | 0 | 1.72E-02 |

| | | | | | | |
|---|---|---|---|---|---|---|
| GO:0030500 | regulation of bone mineralization | BP | 59 | 9 | 6 | 1.72E-02 |
| GO:0022600 | digestive system process | BP | 79 | 11 | 3 | 1.72E-02 |
| GO:0071383 | cellular response to steroid hormone stimulus | BP | 212 | 23 | 8 | 1.73E-02 |
| GO:0098661 | inorganic anion transmembrane transport | BP | 69 | 10 | 2 | 1.74E-02 |
| GO:0090596 | sensory organ morphogenesis | BP | 189 | 21 | 7 | 1.75E-02 |
| GO:0042277 | peptide binding | MF | 201 | 22 | 15 | 1.79E-02 |
| GO:0030282 | bone mineralization | BP | 90 | 12 | 8 | 1.82E-02 |
| GO:0061387 | regulation of extent of cell growth | BP | 90 | 12 | 6 | 1.82E-02 |
| GO:1902476 | chloride transmembrane transport | BP | 50 | 8 | 1 | 1.84E-02 |
| GO:0019748 | secondary metabolic process | BP | 50 | 8 | 1 | 1.84E-02 |
| GO:0030308 | negative regulation of cell growth | BP | 156 | 18 | 6 | 1.88E-02 |
| GO:0042476 | odontogenesis | BP | 101 | 13 | 7 | 1.88E-02 |
| GO:1901021 | positive regulation of calcium ion transmembrane transporter activi | BP | 32 | 6 | 2 | 1.90E-02 |
| GO:0043243 | positive regulation of protein complex disassembly | BP | 32 | 6 | 1 | 1.90E-02 |
| GO:0010107 | potassium ion import | BP | 32 | 6 | 1 | 1.90E-02 |
| GO:0014824 | artery smooth muscle contraction | BP | 9 | 3 | 0 | 1.92E-02 |
| GO:0035988 | chondrocyte proliferation | BP | 9 | 3 | 1 | 1.92E-02 |
| GO:0005984 | disaccharide metabolic process | BP | 9 | 3 | 0 | 1.92E-02 |
| GO:0021561 | facial nerve development | BP | 9 | 3 | 1 | 1.92E-02 |
| GO:0021610 | facial nerve morphogenesis | BP | 9 | 3 | 1 | 1.92E-02 |
| GO:0032274 | gonadotropin secretion | BP | 9 | 3 | 0 | 1.92E-02 |
| GO:0008331 | high voltage-gated calcium channel activity | MF | 9 | 3 | 0 | 1.92E-02 |
| GO:0021932 | hindbrain radial glia guided cell migration | BP | 9 | 3 | 0 | 1.92E-02 |
| GO:0098598 | learned vocalization behavior or vocal learning | BP | 9 | 3 | 0 | 1.92E-02 |
| GO:0051956 | negative regulation of amino acid transport | BP | 9 | 3 | 0 | 1.92E-02 |
| GO:0060044 | negative regulation of cardiac muscle cell proliferation | BP | 9 | 3 | 1 | 1.92E-02 |
| GO:0008195 | phosphatidate phosphatase activity | MF | 9 | 3 | 0 | 1.92E-02 |
| GO:0015791 | polyol transport | BP | 9 | 3 | 0 | 1.92E-02 |
| GO:0097105 | presynaptic membrane assembly | BP | 9 | 3 | 1 | 1.92E-02 |
| GO:2001225 | regulation of chloride transport | BP | 9 | 3 | 1 | 1.92E-02 |
| GO:0080154 | regulation of fertilization | BP | 9 | 3 | 0 | 1.92E-02 |
| GO:0014061 | regulation of norepinephrine secretion | BP | 9 | 3 | 1 | 1.92E-02 |
| GO:0014870 | response to muscle inactivity | BP | 9 | 3 | 0 | 1.92E-02 |
| GO:0007614 | short-term memory | BP | 9 | 3 | 0 | 1.92E-02 |
| GO:0005890 | sodium:potassium-exchanging ATPase complex | CC | 9 | 3 | 1 | 1.92E-02 |
| GO:0032280 | symmetric synapse | CC | 9 | 3 | 0 | 1.92E-02 |
| GO:0006833 | water transport | BP | 9 | 3 | 0 | 1.92E-02 |
| GO:0042044 | fluid transport | BP | 16 | 4 | 0 | 1.99E-02 |
| GO:1904706 | negative regulation of vascular smooth muscle cell proliferation | BP | 16 | 4 | 0 | 1.99E-02 |
| GO:0045663 | positive regulation of myoblast differentiation | BP | 16 | 4 | 0 | 1.99E-02 |
| GO:0033612 | receptor serine/threonine kinase binding | MF | 16 | 4 | 0 | 1.99E-02 |
| GO:0071711 | basement membrane organization | BP | 24 | 5 | 1 | 2.05E-02 |
| GO:0061049 | cell growth involved in cardiac muscle cell development | BP | 24 | 5 | 0 | 2.05E-02 |
| GO:0085029 | extracellular matrix assembly | BP | 24 | 5 | 1 | 2.05E-02 |
| GO:0008045 | motor neuron axon guidance | BP | 24 | 5 | 3 | 2.05E-02 |
| GO:0007274 | neuromuscular synaptic transmission | BP | 24 | 5 | 0 | 2.05E-02 |
| GO:0003301 | physiological cardiac muscle hypertrophy | BP | 24 | 5 | 0 | 2.05E-02 |
| GO:0003298 | physiological muscle hypertrophy | BP | 24 | 5 | 0 | 2.05E-02 |
| GO:1990776 | response to angiotensin | BP | 24 | 5 | 4 | 2.05E-02 |
| GO:0050678 | regulation of epithelial cell proliferation | BP | 263 | 27 | 19 | 2.05E-02 |
| GO:0015698 | inorganic anion transport | BP | 113 | 14 | 3 | 2.06E-02 |
| GO:0055067 | monovalent inorganic cation homeostasis | BP | 113 | 14 | 8 | 2.06E-02 |
| GO:0031333 | negative regulation of protein complex assembly | BP | 113 | 14 | 6 | 2.06E-02 |
| GO:0061035 | regulation of cartilage development | BP | 51 | 8 | 3 | 2.06E-02 |
| GO:0070509 | calcium ion import | BP | 61 | 9 | 6 | 2.11E-02 |
| GO:0031670 | cellular response to nutrient | BP | 61 | 9 | 5 | 2.11E-02 |
| GO:0010657 | muscle cell apoptotic process | BP | 61 | 9 | 7 | 2.11E-02 |
| GO:0048729 | tissue morphogenesis | BP | 538 | 49 | 29 | 2.11E-02 |
| GO:0051961 | negative regulation of nervous system development | BP | 240 | 25 | 12 | 2.14E-02 |
| GO:0003002 | regionalization | BP | 240 | 25 | 11 | 2.14E-02 |
| GO:0009743 | response to carbohydrate | BP | 193 | 21 | 6 | 2.16E-02 |
| GO:0007528 | neuromuscular junction development | BP | 42 | 7 | 1 | 2.17E-02 |
| GO:0090183 | regulation of kidney development | BP | 42 | 7 | 6 | 2.17E-02 |
| GO:0005923 | bicellular tight junction | CC | 103 | 13 | 1 | 2.18E-02 |

| | | | | | | |
|---|---|---|---|---|---|---|
| GO:0051209 | release of sequestered calcium ion into cytosol | BP | 103 | 13 | 12 | 2.18E-02 |
| GO:0010770 | positive regulation of cell morphogenesis involved in differentiation | BP | 136 | 16 | 10 | 2.18E-02 |
| GO:0010659 | cardiac muscle cell apoptotic process | BP | 33 | 6 | 1 | 2.20E-02 |
| GO:0016877 | ligase activity, forming carbon-sulfur bonds | MF | 33 | 6 | 1 | 2.20E-02 |
| GO:0017046 | peptide hormone binding | MF | 33 | 6 | 0 | 2.20E-02 |
| GO:0045687 | positive regulation of glial cell differentiation | BP | 33 | 6 | 1 | 2.20E-02 |
| GO:0010662 | regulation of striated muscle cell apoptotic process | BP | 33 | 6 | 1 | 2.20E-02 |
| GO:0030334 | regulation of cell migration | BP | 708 | 62 | 51 | 2.22E-02 |
| GO:0072009 | nephron epithelium development | BP | 82 | 11 | 8 | 2.23E-02 |
| GO:0097164 | ammonium ion metabolic process | BP | 159 | 18 | 13 | 2.23E-02 |
| GO:0042445 | hormone metabolic process | BP | 159 | 18 | 9 | 2.23E-02 |
| GO:0045927 | positive regulation of growth | BP | 194 | 21 | 10 | 2.27E-02 |
| GO:0046620 | regulation of organ growth | BP | 72 | 10 | 4 | 2.29E-02 |
| GO:0060021 | roof of mouth development | BP | 72 | 10 | 3 | 2.29E-02 |
| GO:0007338 | single fertilization | BP | 72 | 10 | 2 | 2.29E-02 |
| GO:0017022 | myosin binding | MF | 62 | 9 | 3 | 2.32E-02 |
| GO:0005924 | cell-substrate adherens junction | CC | 389 | 37 | 29 | 2.32E-02 |
| GO:0021953 | central nervous system neuron differentiation | BP | 126 | 15 | 8 | 2.35E-02 |
| GO:0005539 | glycosaminoglycan binding | MF | 160 | 18 | 7 | 2.36E-02 |
| GO:0051591 | response to cAMP | BP | 83 | 11 | 7 | 2.41E-02 |
| GO:0071549 | cellular response to dexamethasone stimulus | BP | 25 | 5 | 1 | 2.42E-02 |
| GO:0016709 | oxidoreductase activity, acting on paired donors, with incorporation | MF | 25 | 5 | 2 | 2.42E-02 |
| GO:0051482 | positive regulation of cytosolic calcium ion concentration involved | BP | 25 | 5 | 4 | 2.42E-02 |
| GO:0034284 | response to monosaccharide | BP | 172 | 19 | 4 | 2.43E-02 |
| GO:0048645 | animal organ formation | BP | 43 | 7 | 0 | 2.44E-02 |
| GO:0051926 | negative regulation of calcium ion transport | BP | 43 | 7 | 2 | 2.44E-02 |
| GO:0007632 | visual behavior | BP | 43 | 7 | 3 | 2.44E-02 |
| GO:0003215 | cardiac right ventricle morphogenesis | BP | 17 | 4 | 1 | 2.46E-02 |
| GO:0042745 | circadian sleep/wake cycle | BP | 17 | 4 | 1 | 2.46E-02 |
| GO:0031489 | myosin V binding | MF | 17 | 4 | 1 | 2.46E-02 |
| GO:0030728 | ovulation | BP | 17 | 4 | 0 | 2.46E-02 |
| GO:1903861 | positive regulation of dendrite extension | BP | 17 | 4 | 0 | 2.46E-02 |
| GO:0014850 | response to muscle activity | BP | 17 | 4 | 0 | 2.46E-02 |
| GO:0008090 | retrograde axonal transport | BP | 17 | 4 | 0 | 2.46E-02 |
| GO:0071310 | cellular response to organic substance | BP | 2086 | 163 | 165 | 2.49E-02 |
| GO:0043167 | ion binding | MF | 4777 | 352 | 235 | 2.49E-02 |
| GO:0015294 | solute:cation symporter activity | MF | 73 | 10 | 1 | 2.50E-02 |
| GO:0061526 | acetylcholine secretion | BP | 4 | 2 | 1 | 2.52E-02 |
| GO:0015870 | acetylcholine transport | BP | 4 | 2 | 1 | 2.52E-02 |
| GO:0030043 | actin filament fragmentation | BP | 4 | 2 | 0 | 2.52E-02 |
| GO:0035932 | aldosterone secretion | BP | 4 | 2 | 0 | 2.52E-02 |
| GO:0051933 | amino acid neurotransmitter reuptake | BP | 4 | 2 | 0 | 2.52E-02 |
| GO:0048763 | calcium-induced calcium release activity | MF | 4 | 2 | 0 | 2.52E-02 |
| GO:0035585 | calcium-mediated signaling using extracellular calcium source | BP | 4 | 2 | 1 | 2.52E-02 |
| GO:0003219 | cardiac right ventricle formation | BP | 4 | 2 | 0 | 2.52E-02 |
| GO:0042426 | choline catabolic process | BP | 4 | 2 | 0 | 2.52E-02 |
| GO:0044302 | dentate gyrus mossy fiber | CC | 4 | 2 | 0 | 2.52E-02 |
| GO:0048242 | epinephrine secretion | BP | 4 | 2 | 0 | 2.52E-02 |
| GO:0048241 | epinephrine transport | BP | 4 | 2 | 0 | 2.52E-02 |
| GO:0009812 | flavonoid metabolic process | BP | 4 | 2 | 0 | 2.52E-02 |
| GO:0051935 | glutamate reuptake | BP | 4 | 2 | 0 | 2.52E-02 |
| GO:0004366 | glycerol-3-phosphate O-acyltransferase activity | MF | 4 | 2 | 0 | 2.52E-02 |
| GO:0021888 | hypothalamus gonadotrophin-releasing hormone neuron developme | BP | 4 | 2 | 0 | 2.52E-02 |
| GO:0021886 | hypothalamus gonadotrophin-releasing hormone neuron differentiat | BP | 4 | 2 | 0 | 2.52E-02 |
| GO:0099703 | induction of synaptic vesicle exocytosis by positive regulation of pr | BP | 4 | 2 | 0 | 2.52E-02 |
| GO:0045112 | integrin biosynthetic process | BP | 4 | 2 | 0 | 2.52E-02 |
| GO:0097016 | L27 domain binding | MF | 4 | 2 | 0 | 2.52E-02 |
| GO:0035931 | mineralocorticoid secretion | BP | 4 | 2 | 0 | 2.52E-02 |
| GO:0005862 | muscle thin filament tropomyosin | CC | 4 | 2 | 1 | 2.52E-02 |
| GO:0002318 | myeloid progenitor cell differentiation | BP | 4 | 2 | 0 | 2.52E-02 |
| GO:0031999 | negative regulation of fatty acid beta-oxidation | BP | 4 | 2 | 0 | 2.52E-02 |
| GO:0032277 | negative regulation of gonadotropin secretion | BP | 4 | 2 | 0 | 2.52E-02 |
| GO:1902744 | negative regulation of lamellipodium organization | BP | 4 | 2 | 0 | 2.52E-02 |
| GO:1900028 | negative regulation of ruffle assembly | BP | 4 | 2 | 1 | 2.52E-02 |

| | | | | | | |
|---|---|---|---|---|---|---|
| GO:0042731 | PH domain binding | MF | 4 | 2 | 0 | 2.52E-02 |
| GO:2000344 | positive regulation of acrosome reaction | BP | 4 | 2 | 0 | 2.52E-02 |
| GO:0003321 | positive regulation of blood pressure by epinephrine-norepinephrine | BP | 4 | 2 | 1 | 2.52E-02 |
| GO:2001137 | positive regulation of endocytic recycling | BP | 4 | 2 | 0 | 2.52E-02 |
| GO:0046881 | positive regulation of follicle-stimulating hormone secretion | BP | 4 | 2 | 0 | 2.52E-02 |
| GO:0001996 | positive regulation of heart rate by epinephrine-norepinephrine | BP | 4 | 2 | 1 | 2.52E-02 |
| GO:1901843 | positive regulation of high voltage-gated calcium channel activity | BP | 4 | 2 | 0 | 2.52E-02 |
| GO:0099533 | positive regulation of presynaptic cytosolic calcium concentration | BP | 4 | 2 | 0 | 2.52E-02 |
| GO:0098735 | positive regulation of the force of heart contraction | BP | 4 | 2 | 0 | 2.52E-02 |
| GO:0010898 | positive regulation of triglyceride catabolic process | BP | 4 | 2 | 0 | 2.52E-02 |
| GO:0070474 | positive regulation of uterine smooth muscle contraction | BP | 4 | 2 | 0 | 2.52E-02 |
| GO:0019541 | propionate metabolic process | BP | 4 | 2 | 0 | 2.52E-02 |
| GO:2000858 | regulation of aldosterone secretion | BP | 4 | 2 | 0 | 2.52E-02 |
| GO:0002034 | regulation of blood vessel diameter by renin-angiotensin | BP | 4 | 2 | 0 | 2.52E-02 |
| GO:0014060 | regulation of epinephrine secretion | BP | 4 | 2 | 0 | 2.52E-02 |
| GO:0046880 | regulation of follicle-stimulating hormone secretion | BP | 4 | 2 | 0 | 2.52E-02 |
| GO:0002036 | regulation of L-glutamate import across plasma membrane | BP | 4 | 2 | 0 | 2.52E-02 |
| GO:2000855 | regulation of mineralocorticoid secretion | BP | 4 | 2 | 0 | 2.52E-02 |
| GO:2000437 | regulation of monocyte extravasation | BP | 4 | 2 | 1 | 2.52E-02 |
| GO:0099505 | regulation of presynaptic membrane potential | BP | 4 | 2 | 0 | 2.52E-02 |
| GO:0003072 | renal control of peripheral vascular resistance involved in regulatio | BP | 4 | 2 | 0 | 2.52E-02 |
| GO:1903416 | response to glycoside | BP | 4 | 2 | 0 | 2.52E-02 |
| GO:0035634 | response to stilbenoid | BP | 4 | 2 | 0 | 2.52E-02 |
| GO:0005219 | ryanodine-sensitive calcium-release channel activity | MF | 4 | 2 | 0 | 2.52E-02 |
| GO:0016012 | sarcoglycan complex | CC | 4 | 2 | 0 | 2.52E-02 |
| GO:0102420 | sn-1-glycerol-3-phosphate C16:0-DCA-CoA acyl transferase activi | MF | 4 | 2 | 0 | 2.52E-02 |
| GO:0097490 | sympathetic neuron projection extension | BP | 4 | 2 | 0 | 2.52E-02 |
| GO:0097491 | sympathetic neuron projection guidance | BP | 4 | 2 | 0 | 2.52E-02 |
| GO:0061551 | trigeminal ganglion development | BP | 4 | 2 | 0 | 2.52E-02 |
| GO:0036484 | trunk neural crest cell migration | BP | 4 | 2 | 0 | 2.52E-02 |
| GO:0035290 | trunk segmentation | BP | 4 | 2 | 0 | 2.52E-02 |
| GO:0036486 | ventral trunk neural crest cell migration | BP | 4 | 2 | 0 | 2.52E-02 |
| GO:0099635 | voltage-gated calcium channel activity involved in positive regulati | MF | 4 | 2 | 0 | 2.52E-02 |
| GO:0008559 | xenobiotic transmembrane transporting ATPase activity | MF | 4 | 2 | 0 | 2.52E-02 |
| GO:0043198 | dendritic shaft | CC | 34 | 6 | 1 | 2.52E-02 |
| GO:0001953 | negative regulation of cell-matrix adhesion | BP | 34 | 6 | 2 | 2.52E-02 |
| GO:0010656 | negative regulation of muscle cell apoptotic process | BP | 34 | 6 | 3 | 2.52E-02 |
| GO:0030501 | positive regulation of bone mineralization | BP | 34 | 6 | 4 | 2.52E-02 |
| GO:0048641 | regulation of skeletal muscle tissue development | BP | 34 | 6 | 3 | 2.52E-02 |
| GO:0035094 | response to nicotine | BP | 34 | 6 | 3 | 2.52E-02 |
| GO:0051283 | negative regulation of sequestering of calcium ion | BP | 105 | 13 | 12 | 2.52E-02 |
| GO:0003143 | embryonic heart tube morphogenesis | BP | 53 | 8 | 1 | 2.55E-02 |
| GO:0034308 | primary alcohol metabolic process | BP | 63 | 9 | 1 | 2.55E-02 |
| GO:0007171 | activation of transmembrane receptor protein tyrosine kinase activi | BP | 10 | 3 | 0 | 2.61E-02 |
| GO:0038166 | angiotensin-activated signaling pathway | BP | 10 | 3 | 1 | 2.61E-02 |
| GO:0005952 | cAMP-dependent protein kinase complex | CC | 10 | 3 | 1 | 2.61E-02 |
| GO:0030007 | cellular potassium ion homeostasis | BP | 10 | 3 | 2 | 2.61E-02 |
| GO:1904322 | cellular response to forskolin | BP | 10 | 3 | 1 | 2.61E-02 |
| GO:0031404 | chloride ion binding | MF | 10 | 3 | 0 | 2.61E-02 |
| GO:0010248 | establishment or maintenance of transmembrane electrochemical gr | BP | 10 | 3 | 1 | 2.61E-02 |
| GO:0032836 | glomerular basement membrane development | BP | 10 | 3 | 0 | 2.61E-02 |
| GO:0060347 | heart trabecula formation | BP | 10 | 3 | 0 | 2.61E-02 |
| GO:0070679 | inositol 1,4,5 trisphosphate binding | MF | 10 | 3 | 1 | 2.61E-02 |
| GO:0044224 | juxtaparanode region of axon | CC | 10 | 3 | 1 | 2.61E-02 |
| GO:1902224 | ketone body metabolic process | BP | 10 | 3 | 0 | 2.61E-02 |
| GO:0001527 | microfibril | CC | 10 | 3 | 0 | 2.61E-02 |
| GO:0005859 | muscle myosin complex | CC | 10 | 3 | 1 | 2.61E-02 |
| GO:0051481 | negative regulation of cytosolic calcium ion concentration | BP | 10 | 3 | 0 | 2.61E-02 |
| GO:0017146 | NMDA selective glutamate receptor complex | CC | 10 | 3 | 0 | 2.61E-02 |
| GO:0042415 | norepinephrine metabolic process | BP | 10 | 3 | 1 | 2.61E-02 |
| GO:0048243 | norepinephrine secretion | BP | 10 | 3 | 1 | 2.61E-02 |
| GO:0045989 | positive regulation of striated muscle contraction | BP | 10 | 3 | 1 | 2.61E-02 |
| GO:0032230 | positive regulation of synaptic transmission, GABAergic | BP | 10 | 3 | 1 | 2.61E-02 |
| GO:1902259 | regulation of delayed rectifier potassium channel activity | BP | 10 | 3 | 1 | 2.61E-02 |

| | | | | | | |
|---|---|---|---|---|---|---|
| GO:1900452 | regulation of long-term synaptic depression | BP | 10 | 3 | 0 | 2.61E-02 |
| GO:0002087 | regulation of respiratory gaseous exchange by neurological system | BP | 10 | 3 | 1 | 2.61E-02 |
| GO:1904321 | response to forskolin | BP | 10 | 3 | 1 | 2.61E-02 |
| GO:0014854 | response to inactivity | BP | 10 | 3 | 0 | 2.61E-02 |
| GO:0035864 | response to potassium ion | BP | 10 | 3 | 0 | 2.61E-02 |
| GO:0002116 | semaphorin receptor complex | CC | 10 | 3 | 1 | 2.61E-02 |
| GO:0043492 | ATPase activity, coupled to movement of substances | MF | 84 | 11 | 4 | 2.61E-02 |
| GO:0042626 | ATPase activity, coupled to transmembrane movement of substance | MF | 84 | 11 | 4 | 2.61E-02 |
| GO:0014013 | regulation of gliogenesis | BP | 84 | 11 | 5 | 2.61E-02 |
| GO:0043279 | response to alkaloid | BP | 84 | 11 | 5 | 2.61E-02 |
| GO:0090287 | regulation of cellular response to growth factor stimulus | BP | 209 | 22 | 11 | 2.66E-02 |
| GO:0035023 | regulation of Rho protein signal transduction | BP | 128 | 15 | 9 | 2.67E-02 |
| GO:0045667 | regulation of osteoblast differentiation | BP | 95 | 12 | 6 | 2.68E-02 |
| GO:0045598 | regulation of fat cell differentiation | BP | 106 | 13 | 6 | 2.70E-02 |
| GO:0051282 | regulation of sequestering of calcium ion | BP | 106 | 13 | 12 | 2.70E-02 |
| GO:0032355 | response to estradiol | BP | 106 | 13 | 7 | 2.70E-02 |
| GO:0009612 | response to mechanical stimulus | BP | 174 | 19 | 11 | 2.70E-02 |
| GO:0001667 | ameboidal-type cell migration | BP | 331 | 32 | 21 | 2.71E-02 |
| GO:0070167 | regulation of biomineral tissue development | BP | 74 | 10 | 6 | 2.72E-02 |
| GO:0043502 | regulation of muscle adaptation | BP | 74 | 10 | 1 | 2.72E-02 |
| GO:0120163 | negative regulation of cold-induced thermogenesis | BP | 44 | 7 | 5 | 2.74E-02 |
| GO:0051271 | negative regulation of cellular component movement | BP | 258 | 26 | 15 | 2.78E-02 |
| GO:0003333 | amino acid transmembrane transport | BP | 64 | 9 | 3 | 2.80E-02 |
| GO:0006855 | drug transmembrane transport | BP | 64 | 9 | 1 | 2.80E-02 |
| GO:0000271 | polysaccharide biosynthetic process | BP | 64 | 9 | 1 | 2.80E-02 |
| GO:0032231 | regulation of actin filament bundle assembly | BP | 85 | 11 | 4 | 2.83E-02 |
| GO:0003401 | axis elongation | BP | 26 | 5 | 1 | 2.84E-02 |
| GO:0042596 | fear response | BP | 26 | 5 | 3 | 2.84E-02 |
| GO:0048009 | insulin-like growth factor receptor signaling pathway | BP | 26 | 5 | 1 | 2.84E-02 |
| GO:0055023 | positive regulation of cardiac muscle tissue growth | BP | 26 | 5 | 1 | 2.84E-02 |
| GO:0003091 | renal water homeostasis | BP | 26 | 5 | 3 | 2.84E-02 |
| GO:0014072 | response to isoquinoline alkaloid | BP | 26 | 5 | 2 | 2.84E-02 |
| GO:0043278 | response to morphine | BP | 26 | 5 | 2 | 2.84E-02 |
| GO:0050873 | brown fat cell differentiation | BP | 35 | 6 | 1 | 2.87E-02 |
| GO:0050892 | intestinal absorption | BP | 35 | 6 | 0 | 2.87E-02 |
| GO:2001222 | regulation of neuron migration | BP | 35 | 6 | 1 | 2.87E-02 |
| GO:0010658 | striated muscle cell apoptotic process | BP | 35 | 6 | 1 | 2.87E-02 |
| GO:0000096 | sulfur amino acid metabolic process | BP | 35 | 6 | 1 | 2.87E-02 |
| GO:0019432 | triglyceride biosynthetic process | BP | 35 | 6 | 1 | 2.87E-02 |
| GO:0015718 | monocarboxylic acid transport | BP | 118 | 14 | 6 | 2.87E-02 |
| GO:0048565 | digestive tract development | BP | 107 | 13 | 3 | 2.89E-02 |
| GO:0070160 | tight junction | CC | 107 | 13 | 1 | 2.89E-02 |
| GO:0045665 | negative regulation of neuron differentiation | BP | 176 | 19 | 11 | 2.99E-02 |
| GO:0071875 | adrenergic receptor signaling pathway | BP | 18 | 4 | 1 | 3.00E-02 |
| GO:0015278 | calcium-release channel activity | MF | 18 | 4 | 1 | 3.00E-02 |
| GO:0021846 | cell proliferation in forebrain | BP | 18 | 4 | 3 | 3.00E-02 |
| GO:0042402 | cellular biogenic amine catabolic process | BP | 18 | 4 | 2 | 3.00E-02 |
| GO:0035116 | embryonic hindlimb morphogenesis | BP | 18 | 4 | 0 | 3.00E-02 |
| GO:0099528 | G protein-coupled neurotransmitter receptor activity | MF | 18 | 4 | 2 | 3.00E-02 |
| GO:0086011 | membrane repolarization during action potential | BP | 18 | 4 | 1 | 3.00E-02 |
| GO:0015145 | monosaccharide transmembrane transporter activity | MF | 18 | 4 | 0 | 3.00E-02 |
| GO:0055022 | negative regulation of cardiac muscle tissue growth | BP | 18 | 4 | 1 | 3.00E-02 |
| GO:0051895 | negative regulation of focal adhesion assembly | BP | 18 | 4 | 1 | 3.00E-02 |
| GO:0061117 | negative regulation of heart growth | BP | 18 | 4 | 1 | 3.00E-02 |
| GO:0048520 | positive regulation of behavior | BP | 18 | 4 | 3 | 3.00E-02 |
| GO:0010884 | positive regulation of lipid storage | BP | 18 | 4 | 2 | 3.00E-02 |
| GO:2001025 | positive regulation of response to drug | BP | 18 | 4 | 1 | 3.00E-02 |
| GO:1903859 | regulation of dendrite extension | BP | 18 | 4 | 0 | 3.00E-02 |
| GO:0060390 | regulation of SMAD protein signal transduction | BP | 18 | 4 | 2 | 3.00E-02 |
| GO:0031290 | retinal ganglion cell axon guidance | BP | 18 | 4 | 1 | 3.00E-02 |
| GO:0030431 | sleep | BP | 18 | 4 | 1 | 3.00E-02 |
| GO:0016082 | synaptic vesicle priming | BP | 18 | 4 | 1 | 3.00E-02 |
| GO:0010810 | regulation of cell-substrate adhesion | BP | 188 | 20 | 12 | 3.03E-02 |
| GO:1901681 | sulfur compound binding | MF | 188 | 20 | 6 | 3.03E-02 |

| | | | | | | |
|---|---|---|---|---|---|---|
| GO:0031345 | negative regulation of cell projection organization | BP | 153 | 17 | 10 | 3.04E-02 |
| GO:0008514 | organic anion transmembrane transporter activity | MF | 153 | 17 | 6 | 3.04E-02 |
| GO:0071320 | cellular response to cAMP | BP | 45 | 7 | 5 | 3.07E-02 |
| GO:0003407 | neural retina development | BP | 45 | 7 | 1 | 3.07E-02 |
| GO:0008509 | anion transmembrane transporter activity | MF | 224 | 23 | 7 | 3.07E-02 |
| GO:0071300 | cellular response to retinoic acid | BP | 55 | 8 | 2 | 3.11E-02 |
| GO:0050886 | endocrine process | BP | 55 | 8 | 2 | 3.11E-02 |
| GO:0097110 | scaffold protein binding | MF | 55 | 8 | 4 | 3.11E-02 |
| GO:0019898 | extrinsic component of membrane | CC | 249 | 25 | 16 | 3.18E-02 |
| GO:0030426 | growth cone | CC | 154 | 17 | 9 | 3.21E-02 |
| GO:0030278 | regulation of ossification | BP | 154 | 17 | 14 | 3.21E-02 |
| GO:0050848 | regulation of calcium-mediated signaling | BP | 76 | 10 | 3 | 3.21E-02 |
| GO:0042060 | wound healing | BP | 424 | 39 | 31 | 3.22E-02 |
| GO:0044282 | small molecule catabolic process | BP | 361 | 34 | 12 | 3.25E-02 |
| GO:0017001 | antibiotic catabolic process | BP | 36 | 6 | 3 | 3.25E-02 |
| GO:0098815 | modulation of excitatory postsynaptic potential | BP | 36 | 6 | 0 | 3.25E-02 |
| GO:0042165 | neurotransmitter binding | MF | 36 | 6 | 0 | 3.25E-02 |
| GO:0055025 | positive regulation of cardiac muscle tissue development | BP | 36 | 6 | 3 | 3.25E-02 |
| GO:0010675 | regulation of cellular carbohydrate metabolic process | BP | 120 | 14 | 3 | 3.26E-02 |
| GO:2000145 | regulation of cell motility | BP | 750 | 64 | 55 | 3.28E-02 |
| GO:0008083 | growth factor activity | MF | 87 | 11 | 4 | 3.29E-02 |
| GO:0030551 | cyclic nucleotide binding | MF | 27 | 5 | 2 | 3.30E-02 |
| GO:0007618 | mating | BP | 27 | 5 | 1 | 3.30E-02 |
| GO:1903523 | negative regulation of blood circulation | BP | 27 | 5 | 1 | 3.30E-02 |
| GO:2000171 | negative regulation of dendrite development | BP | 27 | 5 | 1 | 3.30E-02 |
| GO:0090314 | positive regulation of protein targeting to membrane | BP | 27 | 5 | 3 | 3.30E-02 |
| GO:1990573 | potassium ion import across plasma membrane | BP | 27 | 5 | 1 | 3.30E-02 |
| GO:0070873 | regulation of glycogen metabolic process | BP | 27 | 5 | 0 | 3.30E-02 |
| GO:0010837 | regulation of keratinocyte proliferation | BP | 27 | 5 | 2 | 3.30E-02 |
| GO:0030509 | BMP signaling pathway | BP | 109 | 13 | 2 | 3.30E-02 |
| GO:0051208 | sequestering of calcium ion | BP | 109 | 13 | 14 | 3.30E-02 |
| GO:0005925 | focal adhesion | CC | 387 | 36 | 29 | 3.33E-02 |
| GO:2000177 | regulation of neural precursor cell proliferation | BP | 66 | 9 | 4 | 3.34E-02 |
| GO:0007512 | adult heart development | BP | 11 | 3 | 0 | 3.41E-02 |
| GO:0099144 | anchored component of synaptic membrane | CC | 11 | 3 | 1 | 3.41E-02 |
| GO:0003214 | cardiac left ventricle morphogenesis | BP | 11 | 3 | 0 | 3.41E-02 |
| GO:0090533 | cation-transporting ATPase complex | CC | 11 | 3 | 1 | 3.41E-02 |
| GO:0071415 | cellular response to purine-containing compound | BP | 11 | 3 | 0 | 3.41E-02 |
| GO:0010172 | embryonic body morphogenesis | BP | 11 | 3 | 0 | 3.41E-02 |
| GO:0048484 | enteric nervous system development | BP | 11 | 3 | 0 | 3.41E-02 |
| GO:0098712 | L-glutamate import across plasma membrane | BP | 11 | 3 | 0 | 3.41E-02 |
| GO:0061000 | negative regulation of dendritic spine development | BP | 11 | 3 | 0 | 3.41E-02 |
| GO:1903054 | negative regulation of extracellular matrix organization | BP | 11 | 3 | 1 | 3.41E-02 |
| GO:1900272 | negative regulation of long-term synaptic potentiation | BP | 11 | 3 | 0 | 3.41E-02 |
| GO:0021554 | optic nerve development | BP | 11 | 3 | 2 | 3.41E-02 |
| GO:0035810 | positive regulation of urine volume | BP | 11 | 3 | 1 | 3.41E-02 |
| GO:0097090 | presynaptic membrane organization | BP | 11 | 3 | 1 | 3.41E-02 |
| GO:1905939 | regulation of gonad development | BP | 11 | 3 | 0 | 3.41E-02 |
| GO:0051969 | regulation of transmission of nerve impulse | BP | 11 | 3 | 0 | 3.41E-02 |
| GO:0061478 | response to platelet aggregation inhibitor | BP | 11 | 3 | 1 | 3.41E-02 |
| GO:0016918 | retinal binding | MF | 11 | 3 | 0 | 3.41E-02 |
| GO:1902287 | semaphorin-plexin signaling pathway involved in axon guidance | BP | 11 | 3 | 1 | 3.41E-02 |
| GO:0017154 | semaphorin receptor activity | MF | 11 | 3 | 1 | 3.41E-02 |
| GO:0009071 | serine family amino acid catabolic process | BP | 11 | 3 | 0 | 3.41E-02 |
| GO:0036376 | sodium ion export across plasma membrane | BP | 11 | 3 | 1 | 3.41E-02 |
| GO:0050994 | regulation of lipid catabolic process | BP | 46 | 7 | 3 | 3.41E-02 |
| GO:0007631 | feeding behavior | BP | 56 | 8 | 5 | 3.42E-02 |
| GO:0014910 | regulation of smooth muscle cell migration | BP | 56 | 8 | 3 | 3.42E-02 |
| GO:0072347 | response to anesthetic | BP | 56 | 8 | 2 | 3.42E-02 |
| GO:0001649 | osteoblast differentiation | BP | 167 | 18 | 12 | 3.44E-02 |
| GO:1902905 | positive regulation of supramolecular fiber organization | BP | 179 | 19 | 11 | 3.48E-02 |
| GO:0032388 | positive regulation of intracellular transport | BP | 191 | 20 | 13 | 3.50E-02 |
| GO:0043434 | response to peptide hormone | BP | 351 | 33 | 26 | 3.55E-02 |
| GO:0009310 | amine catabolic process | BP | 19 | 4 | 2 | 3.60E-02 |

| | | | | | | |
|---|---|---|---|---|---|---|
| GO:0042219 | cellular modified amino acid catabolic process | BP | 19 | 4 | 1 | 3.60E-02 |
| GO:0021697 | cerebellar cortex formation | BP | 19 | 4 | 0 | 3.60E-02 |
| GO:0051497 | negative regulation of stress fiber assembly | BP | 19 | 4 | 2 | 3.60E-02 |
| GO:0016701 | oxidoreductase activity, acting on single donors with incorporation | MF | 19 | 4 | 3 | 3.60E-02 |
| GO:0016702 | oxidoreductase activity, acting on single donors with incorporation | MF | 19 | 4 | 3 | 3.60E-02 |
| GO:0045761 | regulation of adenylate cyclase activity | BP | 19 | 4 | 2 | 3.60E-02 |
| GO:0072215 | regulation of metanephros development | BP | 19 | 4 | 1 | 3.60E-02 |
| GO:1904752 | regulation of vascular associated smooth muscle cell migration | BP | 19 | 4 | 1 | 3.60E-02 |
| GO:1904738 | vascular associated smooth muscle cell migration | BP | 19 | 4 | 1 | 3.60E-02 |
| GO:0006006 | glucose metabolic process | BP | 180 | 19 | 3 | 3.65E-02 |
| GO:0046463 | acylglycerol biosynthetic process | BP | 37 | 6 | 1 | 3.67E-02 |
| GO:0033173 | calcineurin-NFAT signaling cascade | BP | 37 | 6 | 1 | 3.67E-02 |
| GO:0014047 | glutamate secretion | BP | 37 | 6 | 5 | 3.67E-02 |
| GO:0021575 | hindbrain morphogenesis | BP | 37 | 6 | 2 | 3.67E-02 |
| GO:0046460 | neutral lipid biosynthetic process | BP | 37 | 6 | 1 | 3.67E-02 |
| GO:0046622 | positive regulation of organ growth | BP | 37 | 6 | 2 | 3.67E-02 |
| GO:0032330 | regulation of chondrocyte differentiation | BP | 37 | 6 | 2 | 3.67E-02 |
| GO:0010632 | regulation of epithelial cell migration | BP | 192 | 20 | 12 | 3.67E-02 |
| GO:0015800 | acidic amino acid transport | BP | 57 | 8 | 5 | 3.75E-02 |
| GO:0005546 | phosphatidylinositol-4,5-bisphosphate binding | MF | 57 | 8 | 1 | 3.75E-02 |
| GO:1901655 | cellular response to ketone | BP | 78 | 10 | 4 | 3.75E-02 |
| GO:0032368 | regulation of lipid transport | BP | 78 | 10 | 4 | 3.75E-02 |
| GO:0030427 | site of polarized growth | CC | 157 | 17 | 10 | 3.76E-02 |
| GO:0050840 | extracellular matrix binding | MF | 47 | 7 | 1 | 3.78E-02 |
| GO:0030285 | integral component of synaptic vesicle membrane | CC | 28 | 5 | 1 | 3.80E-02 |
| GO:0040019 | positive regulation of embryonic development | BP | 28 | 5 | 2 | 3.80E-02 |
| GO:0031279 | regulation of cyclase activity | BP | 28 | 5 | 2 | 3.80E-02 |
| GO:0033280 | response to vitamin D | BP | 28 | 5 | 2 | 3.80E-02 |
| GO:0010594 | regulation of endothelial cell migration | BP | 134 | 15 | 10 | 3.80E-02 |
| GO:1901605 | alpha-amino acid metabolic process | BP | 181 | 19 | 8 | 3.83E-02 |
| GO:0060562 | epithelial tube morphogenesis | BP | 266 | 26 | 15 | 3.85E-02 |
| GO:0043296 | apical junction complex | CC | 123 | 14 | 1 | 3.90E-02 |
| GO:0034109 | homotypic cell-cell adhesion | BP | 68 | 9 | 4 | 3.95E-02 |
| GO:0005178 | integrin binding | MF | 112 | 13 | 8 | 3.99E-02 |
| GO:1900619 | acetate ester metabolic process | BP | 5 | 2 | 0 | 4.01E-02 |
| GO:1901374 | acetate ester transport | BP | 5 | 2 | 1 | 4.01E-02 |
| GO:0008291 | acetylcholine metabolic process | BP | 5 | 2 | 0 | 4.01E-02 |
| GO:0060385 | axonogenesis involved in innervation | BP | 5 | 2 | 1 | 4.01E-02 |
| GO:0009374 | biotin binding | MF | 5 | 2 | 0 | 4.01E-02 |
| GO:0008273 | calcium, potassium:sodium antiporter activity | MF | 5 | 2 | 0 | 4.01E-02 |
| GO:0097512 | cardiac myofibril | CC | 5 | 2 | 0 | 4.01E-02 |
| GO:0071284 | cellular response to lead ion | BP | 5 | 2 | 0 | 4.01E-02 |
| GO:0021626 | central nervous system maturation | BP | 5 | 2 | 0 | 4.01E-02 |
| GO:0042584 | chromaffin granule membrane | CC | 5 | 2 | 0 | 4.01E-02 |
| GO:0060029 | convergent extension involved in organogenesis | BP | 5 | 2 | 0 | 4.01E-02 |
| GO:0061550 | cranial ganglion development | BP | 5 | 2 | 0 | 4.01E-02 |
| GO:0016011 | dystroglycan complex | CC | 5 | 2 | 0 | 4.01E-02 |
| GO:0046884 | follicle-stimulating hormone secretion | BP | 5 | 2 | 0 | 4.01E-02 |
| GO:0070095 | fructose-6-phosphate binding | MF | 5 | 2 | 0 | 4.01E-02 |
| GO:0004966 | galanin receptor activity | MF | 5 | 2 | 0 | 4.01E-02 |
| GO:0035933 | glucocorticoid secretion | BP | 5 | 2 | 0 | 4.01E-02 |
| GO:0006114 | glycerol biosynthetic process | BP | 5 | 2 | 0 | 4.01E-02 |
| GO:0015015 | heparan sulfate proteoglycan biosynthetic process, enzymatic modi | BP | 5 | 2 | 0 | 4.01E-02 |
| GO:0021578 | hindbrain maturation | BP | 5 | 2 | 0 | 4.01E-02 |
| GO:0021979 | hypothalamus cell differentiation | BP | 5 | 2 | 0 | 4.01E-02 |
| GO:0005899 | insulin receptor complex | CC | 5 | 2 | 0 | 4.01E-02 |
| GO:0045110 | intermediate filament bundle assembly | BP | 5 | 2 | 0 | 4.01E-02 |
| GO:0014827 | intestine smooth muscle contraction | BP | 5 | 2 | 0 | 4.01E-02 |
| GO:0071694 | maintenance of protein location in extracellular region | BP | 5 | 2 | 1 | 4.01E-02 |
| GO:0003150 | muscular septum morphogenesis | BP | 5 | 2 | 0 | 4.01E-02 |
| GO:0023041 | neuronal signal transduction | BP | 5 | 2 | 0 | 4.01E-02 |
| GO:0003357 | noradrenergic neuron differentiation | BP | 5 | 2 | 0 | 4.01E-02 |
| GO:0021631 | optic nerve morphogenesis | BP | 5 | 2 | 1 | 4.01E-02 |
| GO:0019800 | peptide cross-linking via chondroitin 4-sulfate glycosaminoglycan | BP | 5 | 2 | 0 | 4.01E-02 |

| | | | | | | |
|---|---|---|---|---|---|---|
| GO:2000987 | positive regulation of behavioral fear response | BP | 5 | 2 | 2 | 4.01E-02 |
| GO:2000848 | positive regulation of corticosteroid hormone secretion | BP | 5 | 2 | 0 | 4.01E-02 |
| GO:1903367 | positive regulation of fear response | BP | 5 | 2 | 2 | 4.01E-02 |
| GO:2000543 | positive regulation of gastrulation | BP | 5 | 2 | 0 | 4.01E-02 |
| GO:0032278 | positive regulation of gonadotropin secretion | BP | 5 | 2 | 0 | 4.01E-02 |
| GO:0034112 | positive regulation of homotypic cell-cell adhesion | BP | 5 | 2 | 1 | 4.01E-02 |
| GO:2000020 | positive regulation of male gonad development | BP | 5 | 2 | 0 | 4.01E-02 |
| GO:2000833 | positive regulation of steroid hormone secretion | BP | 5 | 2 | 0 | 4.01E-02 |
| GO:1903818 | positive regulation of voltage-gated potassium channel activity | BP | 5 | 2 | 1 | 4.01E-02 |
| GO:0071692 | protein localization to extracellular region | BP | 5 | 2 | 1 | 4.01E-02 |
| GO:0006863 | purine nucleobase transport | BP | 5 | 2 | 0 | 4.01E-02 |
| GO:0002016 | regulation of blood volume by renin-angiotensin | BP | 5 | 2 | 0 | 4.01E-02 |
| GO:1904026 | regulation of collagen fibril organization | BP | 5 | 2 | 0 | 4.01E-02 |
| GO:2000849 | regulation of glucocorticoid secretion | BP | 5 | 2 | 0 | 4.01E-02 |
| GO:0003062 | regulation of heart rate by chemical signal | BP | 5 | 2 | 0 | 4.01E-02 |
| GO:0033684 | regulation of luteinizing hormone secretion | BP | 5 | 2 | 0 | 4.01E-02 |
| GO:0060405 | regulation of penile erection | BP | 5 | 2 | 0 | 4.01E-02 |
| GO:0046877 | regulation of saliva secretion | BP | 5 | 2 | 0 | 4.01E-02 |
| GO:0060087 | relaxation of vascular smooth muscle | BP | 5 | 2 | 0 | 4.01E-02 |
| GO:0003097 | renal water transport | BP | 5 | 2 | 0 | 4.01E-02 |
| GO:0051414 | response to cortisol | BP | 5 | 2 | 0 | 4.01E-02 |
| GO:0033591 | response to L-ascorbic acid | BP | 5 | 2 | 0 | 4.01E-02 |
| GO:0098921 | retrograde trans-synaptic signaling by endocannabinoid | BP | 5 | 2 | 0 | 4.01E-02 |
| GO:0098920 | retrograde trans-synaptic signaling by lipid | BP | 5 | 2 | 0 | 4.01E-02 |
| GO:0005436 | sodium:phosphate symporter activity | MF | 5 | 2 | 0 | 4.01E-02 |
| GO:1990239 | steroid hormone binding | MF | 5 | 2 | 0 | 4.01E-02 |
| GO:0021825 | substrate-dependent cerebral cortex tangential migration | BP | 5 | 2 | 0 | 4.01E-02 |
| GO:0007217 | tachykinin receptor signaling pathway | BP | 5 | 2 | 0 | 4.01E-02 |
| GO:0021636 | trigeminal nerve morphogenesis | BP | 5 | 2 | 0 | 4.01E-02 |
| GO:0021637 | trigeminal nerve structural organization | BP | 5 | 2 | 0 | 4.01E-02 |
| GO:0006570 | tyrosine metabolic process | BP | 5 | 2 | 0 | 4.01E-02 |
| GO:0007169 | transmembrane receptor protein tyrosine kinase signaling pathway | BP | 587 | 51 | 43 | 4.02E-02 |
| GO:0007045 | cell-substrate adherens junction assembly | BP | 79 | 10 | 2 | 4.04E-02 |
| GO:0048041 | focal adhesion assembly | BP | 79 | 10 | 2 | 4.04E-02 |
| GO:0021761 | limbic system development | BP | 79 | 10 | 5 | 4.04E-02 |
| GO:0009566 | fertilization | BP | 90 | 11 | 2 | 4.07E-02 |
| GO:0030175 | filopodium | CC | 90 | 11 | 4 | 4.07E-02 |
| GO:0021782 | glial cell development | BP | 90 | 11 | 9 | 4.07E-02 |
| GO:0015405 | P-P-bond-hydrolysis-driven transmembrane transporter activity | MF | 90 | 11 | 4 | 4.07E-02 |
| GO:0009416 | response to light stimulus | BP | 243 | 24 | 15 | 4.10E-02 |
| GO:0015108 | chloride transmembrane transporter activity | MF | 58 | 8 | 1 | 4.10E-02 |
| GO:0098659 | inorganic cation import across plasma membrane | BP | 58 | 8 | 3 | 4.10E-02 |
| GO:0099587 | inorganic ion import across plasma membrane | BP | 58 | 8 | 3 | 4.10E-02 |
| GO:0007422 | peripheral nervous system development | BP | 58 | 8 | 5 | 4.10E-02 |
| GO:0010660 | regulation of muscle cell apoptotic process | BP | 58 | 8 | 7 | 4.10E-02 |
| GO:0042127 | regulation of cell proliferation | BP | 1233 | 99 | 106 | 4.12E-02 |
| GO:0061098 | positive regulation of protein tyrosine kinase activity | BP | 48 | 7 | 5 | 4.18E-02 |
| GO:1900024 | regulation of substrate adhesion-dependent cell spreading | BP | 48 | 7 | 4 | 4.18E-02 |
| GO:1901653 | cellular response to peptide | BP | 306 | 29 | 24 | 4.25E-02 |
| GO:0009068 | aspartate family amino acid catabolic process | BP | 20 | 4 | 0 | 4.27E-02 |
| GO:0010667 | negative regulation of cardiac muscle cell apoptotic process | BP | 20 | 4 | 0 | 4.27E-02 |
| GO:0035024 | negative regulation of Rho protein signal transduction | BP | 20 | 4 | 0 | 4.27E-02 |
| GO:0005343 | organic acid:sodium symporter activity | MF | 20 | 4 | 0 | 4.27E-02 |
| GO:0055075 | potassium ion homeostasis | BP | 20 | 4 | 2 | 4.27E-02 |
| GO:0002026 | regulation of the force of heart contraction | BP | 20 | 4 | 1 | 4.27E-02 |
| GO:0048103 | somatic stem cell division | BP | 20 | 4 | 2 | 4.27E-02 |
| GO:0051119 | sugar transmembrane transporter activity | MF | 20 | 4 | 0 | 4.27E-02 |
| GO:0050660 | flavin adenine dinucleotide binding | MF | 69 | 9 | 2 | 4.28E-02 |
| GO:0099513 | polymeric cytoskeletal fiber | CC | 498 | 44 | 44 | 4.28E-02 |
| GO:0048675 | axon extension | BP | 102 | 12 | 7 | 4.32E-02 |
| GO:0042166 | acetylcholine binding | MF | 12 | 3 | 0 | 4.32E-02 |
| GO:0007214 | gamma-aminobutyric acid signaling pathway | BP | 12 | 3 | 0 | 4.32E-02 |
| GO:0061548 | ganglion development | BP | 12 | 3 | 0 | 4.32E-02 |
| GO:0051938 | L-glutamate import | BP | 12 | 3 | 0 | 4.32E-02 |

| | | | | | | |
|---|---|---|---|---|---|---|
| GO:0022840 | leak channel activity | MF | 12 | 3 | 0 | 4.32E-02 |
| GO:0004467 | long-chain fatty acid-CoA ligase activity | MF | 12 | 3 | 0 | 4.32E-02 |
| GO:0072224 | metanephric glomerulus development | BP | 12 | 3 | 0 | 4.32E-02 |
| GO:0022842 | narrow pore channel activity | MF | 12 | 3 | 0 | 4.32E-02 |
| GO:0007194 | negative regulation of adenylate cyclase activity | BP | 12 | 3 | 1 | 4.32E-02 |
| GO:1903960 | negative regulation of anion transmembrane transport | BP | 12 | 3 | 0 | 4.32E-02 |
| GO:0048681 | negative regulation of axon regeneration | BP | 12 | 3 | 0 | 4.32E-02 |
| GO:0042754 | negative regulation of circadian rhythm | BP | 12 | 3 | 0 | 4.32E-02 |
| GO:2000009 | negative regulation of protein localization to cell surface | BP | 12 | 3 | 0 | 4.32E-02 |
| GO:0015874 | norepinephrine transport | BP | 12 | 3 | 1 | 4.32E-02 |
| GO:0030836 | positive regulation of actin filament depolymerization | BP | 12 | 3 | 1 | 4.32E-02 |
| GO:0045725 | positive regulation of glycogen biosynthetic process | BP | 12 | 3 | 0 | 4.32E-02 |
| GO:0090335 | regulation of brown fat cell differentiation | BP | 12 | 3 | 0 | 4.32E-02 |
| GO:1901841 | regulation of high voltage-gated calcium channel activity | BP | 12 | 3 | 1 | 4.32E-02 |
| GO:0032026 | response to magnesium ion | BP | 12 | 3 | 1 | 4.32E-02 |
| GO:0000098 | sulfur amino acid catabolic process | BP | 12 | 3 | 0 | 4.32E-02 |
| GO:0016188 | synaptic vesicle maturation | BP | 12 | 3 | 0 | 4.32E-02 |
| GO:0042403 | thyroid hormone metabolic process | BP | 12 | 3 | 4 | 4.32E-02 |
| GO:0004806 | triglyceride lipase activity | MF | 12 | 3 | 0 | 4.32E-02 |
| GO:0040013 | negative regulation of locomotion | BP | 269 | 26 | 18 | 4.32E-02 |
| GO:0042578 | phosphoric ester hydrolase activity | MF | 319 | 30 | 21 | 4.32E-02 |
| GO:0006790 | sulfur compound metabolic process | BP | 294 | 28 | 10 | 4.33E-02 |
| GO:0031941 | filamentous actin | CC | 29 | 5 | 1 | 4.34E-02 |
| GO:2001258 | negative regulation of cation channel activity | BP | 29 | 5 | 1 | 4.34E-02 |
| GO:0044058 | regulation of digestive system process | BP | 29 | 5 | 0 | 4.34E-02 |
| GO:0051339 | regulation of lyase activity | BP | 29 | 5 | 2 | 4.34E-02 |
| GO:0003009 | skeletal muscle contraction | BP | 29 | 5 | 2 | 4.34E-02 |
| GO:0015399 | primary active transmembrane transporter activity | MF | 91 | 11 | 4 | 4.36E-02 |
| GO:0044706 | multi-multicellular organism process | BP | 160 | 17 | 10 | 4.38E-02 |
| GO:0050673 | epithelial cell proliferation | BP | 307 | 29 | 22 | 4.41E-02 |
| GO:0090130 | tissue migration | BP | 245 | 24 | 15 | 4.44E-02 |
| GO:0036465 | synaptic vesicle recycling | BP | 59 | 8 | 0 | 4.48E-02 |
| GO:0046983 | protein dimerization activity | MF | 979 | 80 | 70 | 4.49E-02 |
| GO:0071326 | cellular response to monosaccharide stimulus | BP | 114 | 13 | 2 | 4.50E-02 |
| GO:0008283 | cell proliferation | BP | 1486 | 117 | 127 | 4.52E-02 |
| GO:0009593 | detection of chemical stimulus | BP | 39 | 6 | 2 | 4.60E-02 |
| GO:0050879 | multicellular organismal movement | BP | 39 | 6 | 2 | 4.60E-02 |
| GO:0050881 | musculoskeletal movement | BP | 39 | 6 | 2 | 4.60E-02 |
| GO:0045661 | regulation of myoblast differentiation | BP | 39 | 6 | 1 | 4.60E-02 |
| GO:0006639 | acylglycerol metabolic process | BP | 103 | 12 | 7 | 4.60E-02 |
| GO:0050680 | negative regulation of epithelial cell proliferation | BP | 103 | 12 | 7 | 4.60E-02 |
| GO:0001755 | neural crest cell migration | BP | 49 | 7 | 3 | 4.60E-02 |
| GO:0140238 | presynaptic endocytosis | BP | 49 | 7 | 0 | 4.60E-02 |
| GO:0051145 | smooth muscle cell differentiation | BP | 49 | 7 | 4 | 4.60E-02 |
| GO:0003707 | steroid hormone receptor activity | MF | 49 | 7 | 1 | 4.60E-02 |
| GO:0048488 | synaptic vesicle endocytosis | BP | 49 | 7 | 0 | 4.60E-02 |
| GO:0007589 | body fluid secretion | BP | 70 | 9 | 0 | 4.63E-02 |
| GO:0001678 | cellular glucose homeostasis | BP | 126 | 14 | 2 | 4.64E-02 |
| GO:2000241 | regulation of reproductive process | BP | 92 | 11 | 6 | 4.66E-02 |
| GO:0106027 | neuron projection organization | BP | 81 | 10 | 5 | 4.68E-02 |
| GO:0110020 | regulation of actomyosin structure organization | BP | 81 | 10 | 4 | 4.68E-02 |
| GO:0005575 | cellular_component | CC | 12612 | 866 | 667 | 4.86E-02 |
| GO:0035050 | embryonic heart tube development | BP | 60 | 8 | 1 | 4.87E-02 |
| GO:0003158 | endothelium development | BP | 104 | 12 | 5 | 4.89E-02 |
| GO:0006638 | neutral lipid metabolic process | BP | 104 | 12 | 7 | 4.89E-02 |
| GO:0015804 | neutral amino acid transport | BP | 30 | 5 | 3 | 4.93E-02 |
| GO:0043394 | proteoglycan binding | MF | 30 | 5 | 2 | 4.93E-02 |
| GO:0007044 | cell-substrate junction assembly | BP | 93 | 11 | 2 | 4.98E-02 |
| GO:0007565 | female pregnancy | BP | 139 | 15 | 8 | 4.99E-02 |
| GO:1903034 | regulation of response to wounding | BP | 139 | 15 | 6 | 4.99E-02 |
| GO:0030301 | cholesterol transport | BP | 71 | 9 | 6 | 4.99E-02 |
| GO:0060993 | kidney morphogenesis | BP | 71 | 9 | 8 | 4.99E-02 |
| GO:0095500 | acetylcholine receptor signaling pathway | BP | 21 | 4 | 0 | 5.00E-02 |
| GO:1904385 | cellular response to angiotensin | BP | 21 | 4 | 2 | 5.00E-02 |

| | | | | | | |
|---|---|---|---|---|---|---|
| GO:0071377 | cellular response to glucagon stimulus | BP | 21 | 4 | 4 | 5.00E-02 |
| GO:0046697 | decidualization | BP | 21 | 4 | 1 | 5.00E-02 |
| GO:0032590 | dendrite membrane | CC | 21 | 4 | 0 | 5.00E-02 |
| GO:0035137 | hindlimb morphogenesis | BP | 21 | 4 | 0 | 5.00E-02 |
| GO:0032232 | negative regulation of actin filament bundle assembly | BP | 21 | 4 | 2 | 5.00E-02 |
| GO:1903392 | negative regulation of adherens junction organization | BP | 21 | 4 | 1 | 5.00E-02 |
| GO:0051197 | positive regulation of coenzyme metabolic process | BP | 21 | 4 | 2 | 5.00E-02 |
| GO:0030813 | positive regulation of nucleotide catabolic process | BP | 21 | 4 | 2 | 5.00E-02 |
| GO:2000810 | regulation of bicellular tight junction assembly | BP | 21 | 4 | 3 | 5.00E-02 |
| GO:0051930 | regulation of sensory perception of pain | BP | 21 | 4 | 2 | 5.00E-02 |
| GO:1903831 | signal transduction involved in cellular response to ammonium ion | BP | 21 | 4 | 0 | 5.00E-02 |
| GO:0010092 | specification of animal organ identity | BP | 21 | 4 | 0 | 5.00E-02 |
| GO:0009404 | toxin metabolic process | BP | 21 | 4 | 0 | 5.00E-02 |
